# Community-wide epigenetics provides novel perspectives on the ecology and evolution of marine microbiome

**DOI:** 10.1101/2021.11.30.470565

**Authors:** Hoon Je Seong, Simon Roux, Chung Yeon Hwang, Woo Jun Sul

## Abstract

DNA methylation in prokaryotes is involved in many different cellular processes including cell cycle regulation and defense against viruses. To date, most prokaryotic methylation systems have been studied in culturable microorganisms, resulting in a limited understanding of DNA methylation from a microbial ecology perspective. Here, we analyze the distribution patterns of several microbial epigenetics marks in the ocean microbiome through genome-centric metagenomics across all domains of life. We show that overall, DNA methylation can readily be detected across dominant oceanic bacterial, archaeal, and viral populations, and microbial epigenetic changes correlate with population differentiation. Furthermore, our genome-wide epigenetic analysis of *Pelagibacter* suggests that GANTC, a DNA methyltransferase target motif, is related to the cell cycle and is affected by environmental conditions. Yet, the presence of this motif also partitions the phylogeny of the *Pelagibacter* phages, possibly hinting at a competitive co-evolutionary history and multiple effects of a single methylation mark.

**One-Sentence Summary:** DNA methylation patterns are associated with ecological changes and virus-host dynamics in the marine microbiome.

## Main text

DNA methylation is an important epigenetic modification in which methyl groups are added to nucleotides by methyltransferases (MTases). Although much of the literature has focused on DNA methylation in eukaryotes, the last decade has seen increased research on DNA methylation in prokaryotes due to the emergence of third-generation sequencing technology. In prokaryotes, DNA methylation has been largely associated with restriction-modification (RM) systems, which protect host cells against invasion by viruses or against horizontal gene transfer of extracellular DNA by distinguishing host DNA from sequence-specific DNA methylation. DNA methylation also has roles in gene expression regulation, virulence, DNA mismatch repair, and cell-cycle regulation in prokaryotes. DNA MTases that methylate nucleotides without cognate restriction enzymes (REases) are referred to as orphan MTases. Some well-studied orphan MTases play important physiological roles beyond RM systems, including transcriptional regulation and cell phenotype variations (*1–4*). Thus, there has been increasing interest in the role of prokaryotic methylation systems in bacterial genetics, phenotypic changes, and pathogenesis. However, the implications for the diversity and meaning of prokaryotic methylation systems in environmental conditions are unclear. The primary reason is that previous research has focused extensively on culturable prokaryotes, whereas the majority of marine bacteria are currently not culturable under laboratory conditions.

The advent of long-read single-molecule real-time (SMRT) sequencing has opened a new era in methylation research. While SMRT sequencing cannot currently identify all known base modifications, it can readily detect the two major DNA modifications found in prokaryotes: *N*^6^-methyladenosine (m^6^A) and *N*^6^-methylcytosine (m^4^C), including from metagenomes. This methylation information has been used in the past to improve prokaryotic genome binning (*5, 6*). However, outside these few examples, few attempts have been made to apply long-read metagenomic sequencing to detect DNA methylation directly in the environmental microbiome (*7*), and link methylation patterns to ongoing eco-evolutionary processes.

Here, we applied meta-epigenomic analysis to genome-centric metagenomics of the ocean microbiome in the northwest Pacific Ocean to reveal the role of DNA methylation in environmental microbial communities. We report that the DNA methylome is differentiated by taxonomic lineage and is affected by the complexity of the community, i.e., the co-existence of multiple closely related strains. We further link methylation patterns to cell cycle regulation and phage defense for *Pelagibacter* genomes, highlighting the multiple roles played by DNA methylation in one of the dominant bacteria of the marine environment.

## Results and Discussion

### Novel microbial genomes from the northwest Pacific Ocean metagenome

In the 2015 Shipborne Pole-to-Pole Observations (SHIPPO) project of the Korea Polar Research Institute, we conducted shotgun metagenomic sequencing using ocean surface samples from 10 stations (referred to as St2–St11) by traveling about 4,000 km from the Pacific Northwest to the Bering Sea during July 22–29, 2015 (fig. S1, table S1).□To capture free-living organisms, we extracted genomic DNA after 0.22–1.6-μm size filtering.□Ten seawater samples were sequenced with 154.4 Gb (read average length: 151 bp) using the Illumina HiSeq 4000 platform and 32.2 Gb (read average length: 797.9 bp) using the Pacific Biosciences (PacBio) RSII (3.2 cells per sample) platform, respectively. Extensive computational analysis was performed on all samples to reconstruct the genomes across the kingdom using a combination of individual, co-, and hybrid assembly, binning, and refinement methods (Fig. 1). This strategy allowed the recovery of a total of 15,056 viral, 252 prokaryotic, 56 giant viral, and 6 eukaryotic metagenome-assembled genomes (MAGs, specifically referred to here as SHIPPO vMAGs, proMAGs, gvMAGs, and eukMAGs, respectively). A total of 252 dereplicated proMAGs (99% average nucleotide identity; ANI) with ≥50% completeness and <10% contamination remained (average completeness: 78.80±14.09, and contamination: 2.76±2.37), 105 originated from the individual binning and 147 from the co-assembly binning. Forty-seven had >90% completeness and <5% contamination (near-complete), of which only three were high-quality MAGs that fit the Minimum Information about a Metagenome-assembled Genome (MIMAG) criteria (*8*) including rRNA and tRNA. These proMAGs mainly consist of the bacterial phyla of Proteobacteria (*n* = 120), Bacteroidota (*n* = 88), Actinobacteriota (*n* = 11), and the archaeal phyla of Thermoplasmatota (*n* = 15) (Fig. 2A).

**Fig. 1.**
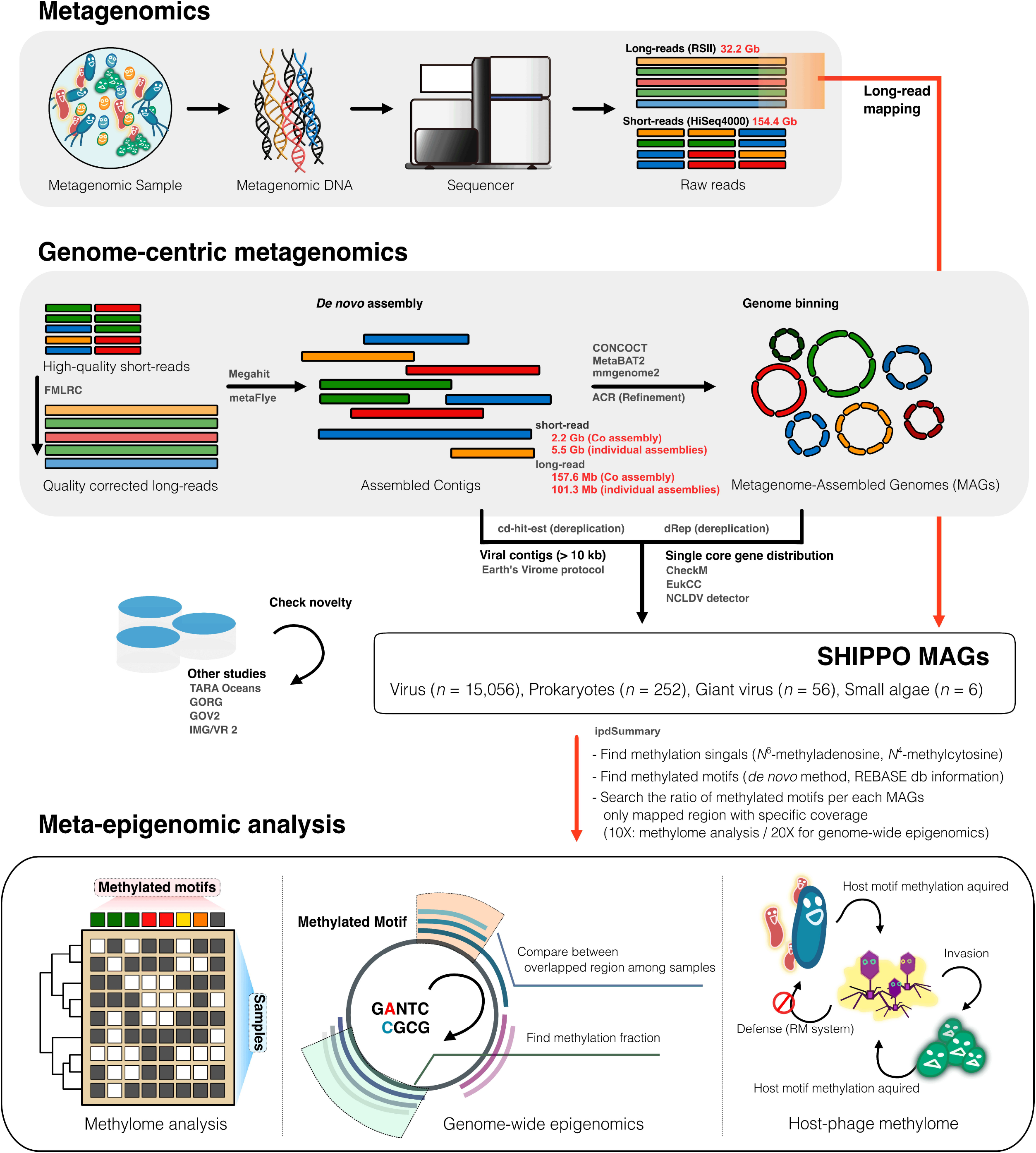
Meta-epigenome analysis scheme of ocean surface samples. A schematic overview of metaepigenomics. Meta-epigenomics using genome-centric metagenomics from the binning approach of short- and long-read assemblies, followed by identifying the epigenetic signals of genomes from long-read mapping.

**Fig. 2.**
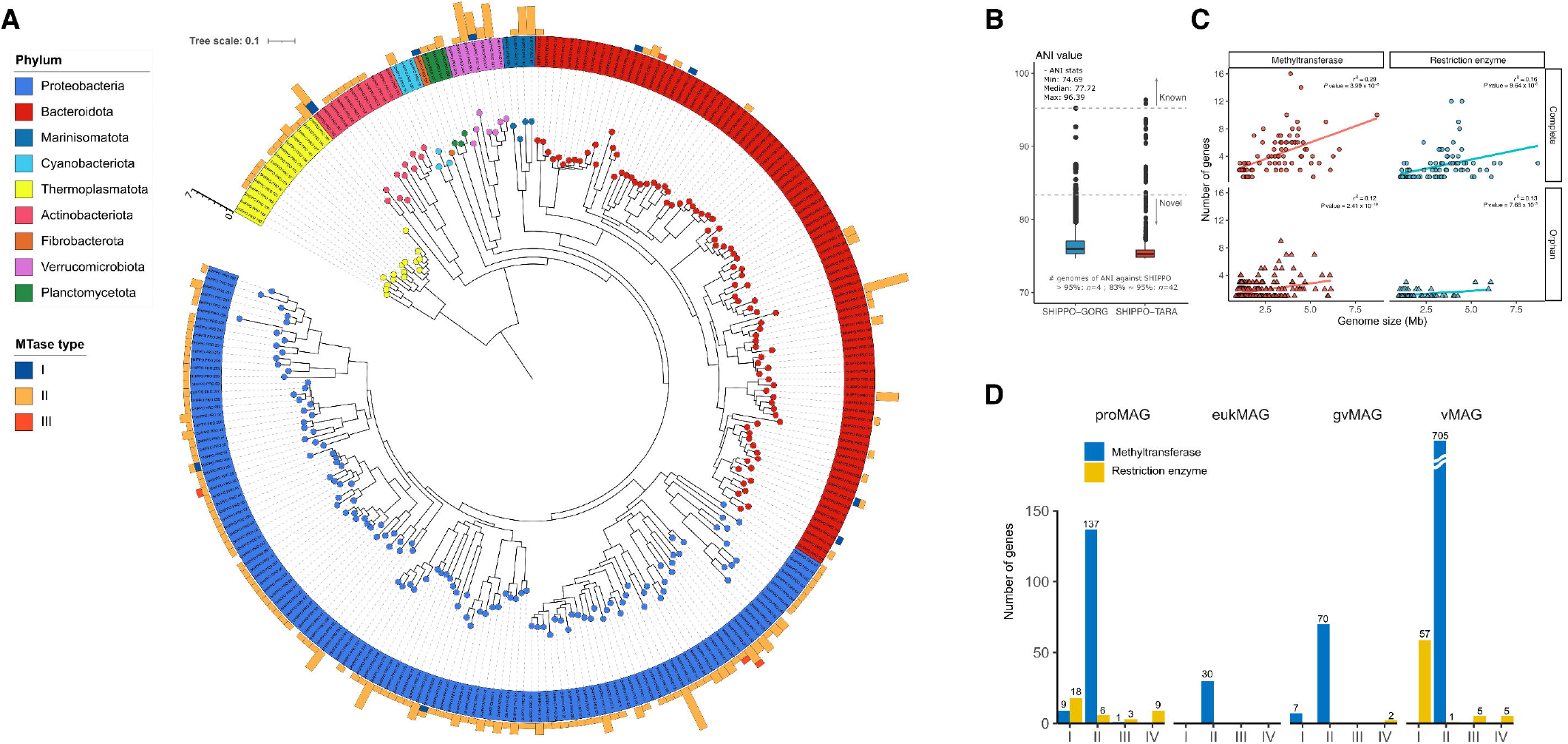
Phylogenetic tree of MAGs obtained from SHIPPO. (**A**) A phylogenetic tree of prokaryotic SHIPPO MAGs using core genes from Phylophlan2. A total of 252 strain-level MAGs were obtained; each bar outside the tree represents the number of methyltransferase (MTase) genes present in each MAG. (**B**) The distribution of restriction enzyme and MTase types from SHIPPO MAGs across kingdoms. (**C**) Prokaryotic SHIPPO MAGs were compared against genomes from *Tara* Oceans (TARA) and Global Ocean Reference Genomes Tropics (GORG-Tropics) datasets using FastANI. (**D**) The number of genes associated with the restriction–modification (RM) system is plotted against the genome size for each ocean microbiome MAG (SHIPPO MAG, TARA, and GORG-Tropics). Points are shaped depending on the type of the complete and orphan RM system. MAG: metagenome-assembled genome; SHIPPO: Shipborne Pole-to-Pole Observations.

This assembly strategy substantially improved the fraction of mapped metagenomic reads. Overall, the average mappability of all samples was 38.03% (std. 2.88) (fig. S2). Most of the reads mapped to vMAGs and proMAGs, with relatively smaller representation of gvMAGs and eukMAGs. We compared our proMAGs with the TARA (9) and Global Ocean Reference Genomes Tropics (GORG-Tropics) (*10*) datasets to evaluate the novelty of our recovered genomes (Fig. 2B). Although these proMAGs and single-cell amplified genomes datasets came from a global-scale study (*9, 10*), only three proMAGs overlapped at the species level (≥95% ANI) with our proMAGs. Furthermore, 95% of the proMAGs obtained here could not be classified at the species level, even though the Genome Taxonomy Database (GTDB) (*11*) includes genomes of uncultured organisms derived from shotgun metagenomics and single-cell genomics. Thus, despite previous extensive ocean metagenomic binning efforts such as those undertaken on data from mega-surveys like the *Tara* Ocean Expedition (TARA) and the Global Ocean Survey (*9, 10*), the northwest Pacific Ocean datasets from this study provide substantial novel genomic information on ocean microbiomes.

### DNA MTases in marine microorganisms are rarely associated with an RM system

To characterize the role of MTases in ocean microbial communities, we first identified the type of MTases and their cognate REases distribution from the genome catalog established here and derived from previous ocean microbiome surveys (GORG, TARA, and SHIPPO). Of the total 5,713 medium-quality proMAGs, we found 67.18% (3,838) and 19.45% (1,111) of proMAGs encoded one or more MTases and REases, respectively (table S2). Among the four MTase types—I, II, III, and IV—type II MTase was found most frequently (94.71%) in proMAGs with MTases, followed by type I (14.02%) and III (2.16%). Of all the proMAGs, only 14.77% had a complete RM system; most consisted of type I and III MTases: 76.39% and 74.70% of type I and III MTases constituted a complete RM system, respectively, whereas only 3.19% of type II MTases constituted an RM system. By contrast, most MTases (86.09%) belonged to orphan MTases, thus lacking counterpart REases, and consisted of type II MTases. To compare the genome size of ocean prokaryotes and the number of genes related to RM systems, 424 near-complete proMAGs were used. Although the number of MTase and REase correlated with genome size, the genome of proMAG with one or more RM system was significantly larger than that of proMAG with orphan MTase (Wilcoxon’s rank-sum *P* value < 2.2 x 10^-16^). In addition, in the case of proMAG with RM system, the correlation between MTase and genome size (*r*^2^: 0.29) was higher than for REase (*r*^2^: 0.16) (Fig. 2C). In the ocean microbial community, loss of the type II RM system may be caused by the selective pressures of genome streamlining in the pelagic environment (12), which contributes to retaining essential cellular mechanisms rather than defense systems. By contrast, type I and III MTases and their cognate REases typically serve as a defense mechanism through the RM system and thus are harbored in genomes of relatively large size.

Beyond MTases detected in proMAGs, a total of 959 MTases were found in the SHIPPO MAGs catalog, including all (*n* = 6) of the eukMAGs, 62.5% (*n* = 35) of the gvMAGs, 36.90% (*n* = 93) of the proMAGs, and 4.28% (*n* = 645) of the vMAGs (Fig. 2D and table S3). Consistent with the abovementioned results, type II MTases were the most frequently detected for all domains (Fig. 2D). All but two MTases were solitary or orphan MTases that have no counterpart REases. Eukaryotes and giant viruses had an average of 5.0 and 2.2 MTases per genome, whereas fewer MTases were found in the prokaryotic and viral genomes (1.58 and 1.09 per genome, respectively).

### DNA methylome of SHIPPO MAGs

We next studied the DNA methylation patterns of the ocean microbiome and compared DNA methylation profiles of each SHIPPO MAG across samples. We first performed a principal coordinate analysis (PCoA) based on the Kulczynski dissimilarity of 5-mer DNA methylation profiles for individual MAG-sample pairs (requiring 10× coverage with 20% genome breadth for proMAGs and gvMAGs, 10% for eukMAGs, and 60% for vMAGs). DNA methylation profiles were grouped clearly by domain, i.e., separating eukaryotes, prokaryotes, and virus MAGs (Fig. 3A). In particular, Alphaproteobacteria harbored distinct methylation profiles compared to all other microbial organisms, and Alphaproteobacteria proMAGs were partitioned from each other down to the family level. However, the methylation profile of 5-mers could not be distinguished at the species level, as in the example of the *Pelagibacteraceae* cluster, which consisted of SHIPP_PRO_33, SHIPP_PRO_245, and SHIPP_PRO_247. Furthermore, proMAGs belonging to Flavobacteriia, Actinobacteria, Gammaproteobacteria, and Bacteroidia were also difficult to distinguish by their methylation profile.

**Fig. 3.**
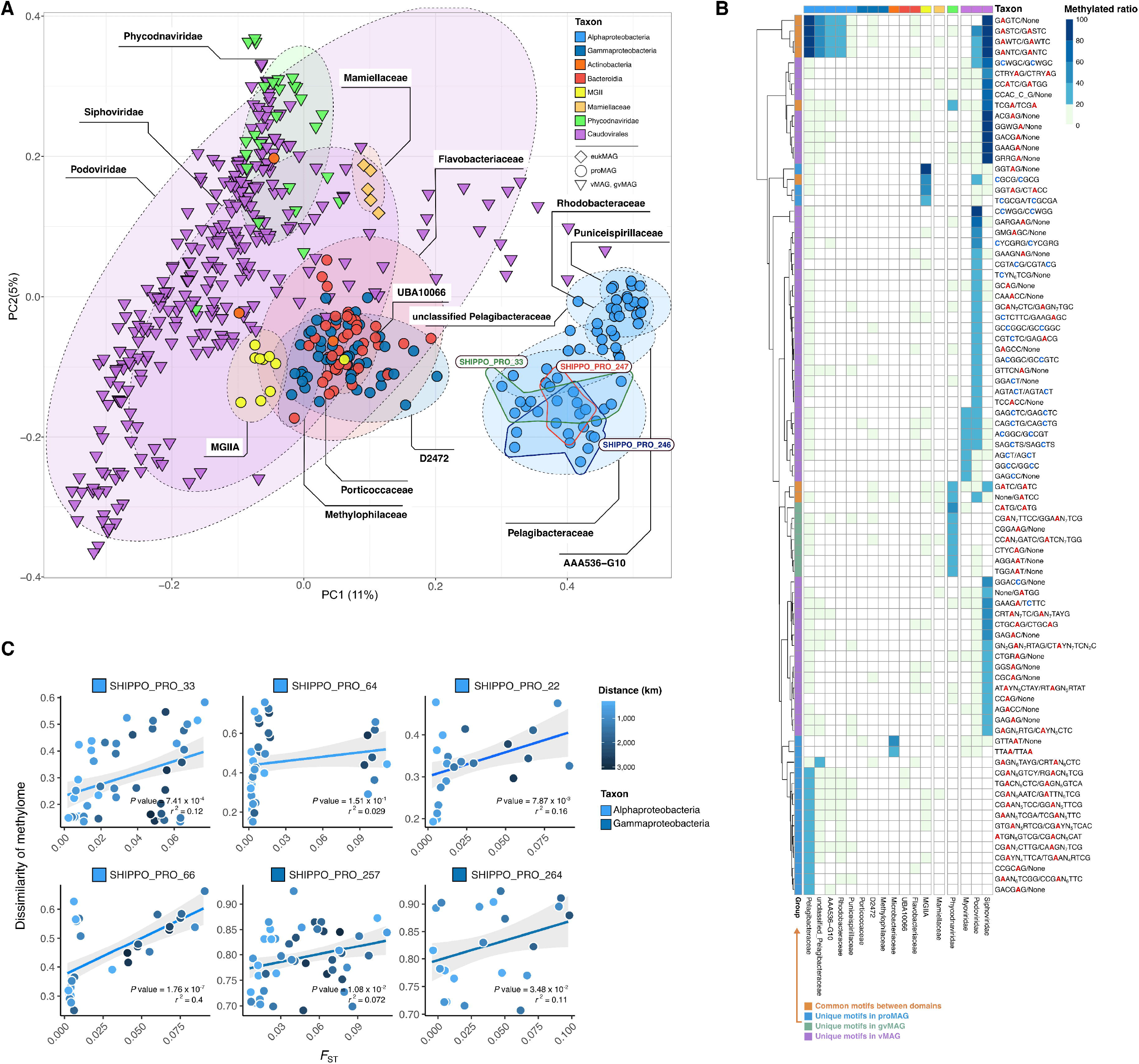
Meta-epigenomic profile of MAGs across all sampling stations. (**A**) Principal coordinate analysis (PCoA) clustering by the 5-mer methylation features of Shipborne Pole-to-Pole Observations (SHIPPO) MAGs based on Kulczynski dissimilarity. Each point represents each species-level MAG in each sample. The black-dashed circles represent family-level clusters of MAGs across samples. The colored-solid lines represent species-level clusters belonging to *Pelagibacteraceae* across samples. (**B**) The maximum methylation ratios of motifs are represented in each MAG at the family level; the highest methylation value among all sample sites is colorized. (**C**) Population differentiation versus methylome across sampling stations. For the most prevalent Shipborne Pole-to-Pole Observations (SHIPPO) MAGs, scatterplots show the relationship of 5-mer methylome dissimilarity based on Bray–Curtis and population differentiation by sampling distance. MAG: metagenome-assembled genome (eukaryotic: eukMAG, prokaryotic: proMAG, viral MAG: vMAG, giant viral MAG: gvMAG); *F*_ST_: fixation index.

To identify the exact DNA methylated motif, the methylated motif information was collected via the Restriction Enzyme database (REBASE) (*13*). Although additional motifs were discovered from the *de novo* motif finding algorithm MultiMotifMaker (*14*) across SHIPPO MAGs, most motifs were discovered in previous studies (G<u>A</u>TC, G<u>A</u>NTC, <u>C</u>GCG, V<u>A</u>TB, underlining indicates methylation position) (*15, 16*). A total of 1,357 motifs were searched along the mapped region of each SHIPPO MAG. When <20% of each motif was methylated in each genome, it was considered noise and excluded from the candidate methylation motifs. Ninety-five candidate methylated motifs were detected, of which 17 and 76 represented m^4^C and m^6^A modifications, respectively. The other two were non-palindromic motifs (CGT<u>C</u>TC/GAG<u>A</u>CG, GAAG<u>A</u>/T<u>C</u>TTC) with both m^4^C and m^6^A methylation. Among the methylated motifs, 13 motifs were shared across domains. G<u>A</u>NTC was found most frequently in several families belonging to Alphaproteobacteria and in *Caudovirales* (Fig. 3B). <u>C</u>GCG and GGT<u>A</u>G were detected in both archaeal proMAGs affiliated to MGIIA and *Caudovirales* vMAGs (Fig. 3B). The G<u>A</u>TC motif was found in gvMAGs (*Phycodnaviridae), Caudovirales*, and unclassified vMAGs (Fig. 3B). In addition, there were methylated motifs unique to vMAGs, such as C<u>C</u>WGG and GG<u>C</u>C, and unique motifs were found in each family, such as TTA<u>A</u> (*Microbacteriaceae*), and T<u>C</u>GCGA (MGIIA) (Fig. 3B). The high diversity of methylated motifs in vMAG suggests the existence of many unknown MTases encoded on prokaryotic and/or viral genomes, and is consistent with a role of viral genome methylation in virus-host arms race.

To match the methylated motifs with MTases of each SHIPPO MAG, candidate MTases were searched against REBASE (*13*) reference MTases with recognition sequence information. Only the recognition sequences of 12 MTases were known, and except for the recognition sequences of one viral MTase, it was confirmed that the methylated motif sequences were identical to the recognition sequence information (table S3). For example, in the seven Alphaproteobacteria proMAGs, we could identify methylation signals from several motifs, including G<u>A</u>NTC, G<u>A</u>STC, G<u>A</u>GTC, and CG<u>A</u>NNNNNNAATC, but these were represented by G<u>A</u>TNC, for which the congruent recognition sequences of the best similarity reference MTase could be found (table S3). There were four methylation motifs in an archaeal proMAG (SHIPPO_PRO_101), and after deduplication, two different motifs (<u>C</u>GCG and GGT<u>A</u>G) were represented (table S3). One congruent MTase could be found (<u>C</u>GCG) among these two methylation motifs, but no MTase matched the other methylation motif (GGT<u>A</u>G). Although the GGT<u>A</u>G motif was novel in that it is previously unreported, this result limited the confirmation of the archaeal novel methylation system, likely due to the fact that this MAGs does not represent a complete genome. Furthermore, except for 12 of the 1,124 MTases, it was either difficult to identify the methylation profile due to the long-read sequencing depth or the lack of MTase recognition sites in the previous database made it difficult to compare between MTases and motif sequences.

### DNA methylation patterns are linked to population genomic structure

Several studies of bacterial DNA methylome suggested that different bacterial strains have different methylome patterns, even within species (*6, 17, 18*). These changes can be caused by the presence of MTases or by phase-variable MTases that respond to changes in the environment (*19*). However, microbial DNA methylation changes in complex environments have not yet been measured directly; therefore, we analyzed intra-species DNA methylation variation at the sampling station. The fixation index (*F*_ST_) was used to compare the similarity in the population differentiation between samples. proMAGs with low base-pair coverage were excluded because the *F*_ST_ had to be calculated by the allele frequencies within the species. *F*_ST_ was calculated for dominant proMAGs using a mapping region that overlaps at least 40% breadth with 10× depth in all samples with short reads. The dissimilarity of methylomes was calculated by the methylation frequency of 5-mer nucleotides in a genome. Therefore, to compare the DNA methylation changes for different sampling stations, only six species-level proMAGs were fulfilled by the sequencing coverage of long- and short reads. In five of the six proMAGs, DNA methylome differences and population differentiation across samples correlated significantly, regardless of the distance between sampling stations (Pearson correlation, *P* value < 0.05; Fig. 3C). A significant correlation was also found in Gammaproteobacteria proMAGs that showed no specific methylated motif (weak methylated motifs ratio <10%). These results may indicate that when multiple strains with different methylation profiles of motifs are present in the environment, they affect the methylation pattern at the species level.

### Genome-wide DNA methylation analysis of Pelagibacter in environmental samples

Next, we analyzed the methylation pattern of dominant proMAGs at the single-nucleotide level to investigate in detail the environmental-dependent changes in methylation. Identifying the single-nucleotide-level DNA methylation from a genome-wide perspective required a relatively deeper and wider read coverage. Four proMAGs (SHIPPO_PRO_33, SHIPPO_PRO_246, SHIPPO_PRO_64, and SHIPPO_PRO_101) were covered over >40% of the breadth of the genome at 20× depth per strand. Of these, only a SHIPPO_PRO_33 proMAG affiliated with *Pelagibacter* overlapped at 65.61% of the genome breadth in all 10 samples. The average breadth of the genome coverage with 20× per strand of SHIPPO_PRO_33 was 90.93% (std. 7.40). The in-depth overlapped coverage of long-reads enabled a comparison of methylation patterns between samples for this specific *Pelagibacter* proMAG.

As mentioned above, the MTases of most marine microorganisms were found without counterpart REases, and conserved MTases were observed without counterpart REases in the four *Pelagibacter* proMAGs. In particular, the *Pelagibacter* proMAGs showed that the methylated motif (GANTC) and the recognition motif of their MTase were consistent. The GANTC motif in the *Pelagibacter* proMAGs was globally uniformly distributed throughout the genome (Shapiro–Wilk test: genic >0.05, intergenic >0.05, and regulatory >0.05). For the bacterial MTases involved in gene regulation, methylated motifs were frequently located upstream of regulated genes (*20*). In this study, we found that GANTC was enriched in the intergenic region of *Pelagibacter*, indicating that GANTC favors an epigenetic role (*P* value = 0.05, fig. S3).

To compensate for the lack of sequencing depth in the genome between samples, we focused on the genome-wide epigenetic analysis of the *Pelagibacter* proMAG (SHIPPO_PRO_33; 12 contigs; 87.30 completeness, 3.79 contamination). Only the nucleotide positions of the overlapped regions that could measure methylation in all samples were compared. A total of 2,494 GANTC motifs were detected on both strands of the SHIPPO_PRO_33 genome, and only 1,719 GANTC (68.93%) were included in the overlapped region (the average depth of reads was 112.85×, std. 60.56) for measuring the methylation at 10 sampling stations. Although most motifs are known to be methylated throughout the prokaryotic genome (*16*), some motif sites have been found that remain unmethylated (*20–22*). These unmethylated sites may be due to competitive binding between MTases and regulatory proteins due to epigenetic regulation (*23, 24*). The methylation rate of GANTC varied in the range 71.31–94.77% depending on the sample, of which 2.44% (42 sites) remain unmethylated in all samples. In most cases, unmethylated sites were frequently observed in intergenic regions (66.67%), including regulatory regions (16.70%) (table S3). The GANTC position of one of these seven unmethylated positions of the regulatory region contains the *sufE* (K04488) regulatory region of the Suf (sulfur-forming) system. This gene is part of the SufSE complex with SufS and is involved in the transport of sulfur to the sulfur mobilization protein. This gene was also significantly more highly expressed in the absence of dimethylsulfoniopropionate (DMSP), sulfur-limited in the laboratory, along with *sufS* (*25*). In addition, we observed unmethylated GANTC in the regulatory region of *groEL* (chaperones gene, K04077), which is also more expressed in sulfur-restricted conditions (25). However, we found that most GANTC motifs (99.86%, 3477/3482) were fully methylated (both strands methylated) under the DMSP-rich laboratory culture condition of *Candidatus* Pelagibacter Giovannoni NP1 (*26*), which is most closely related to SHIPPO_PRO_33 (ANI ≥ 92.36). All but five GANTC sites were methylated throughout the chromosome. Two of the five unmethylated sites were fully methylated in both strands of the tRNA (Phe) region, and the rest were methylated in the intergenic region. However, the SHIPPO_PRO_33 proMAGs from environmental samples lacked genomic integrity and could not be compared with the five unmethylated GANTC positions of *Cand*. P. Giovannoni NP1. The differences in unmethylated GANTCs between environmental and laboratory settings suggest that they are the result of epigenetic regulation in competition for nutrients, such as sulfur.

The DNA methylation signal at the nucleotide resolution is calculated from the pooled interpulse duration (IPD) ratio in separate molecules for each genomic locus. Due to the often-found epigenetic heterogeneity (*27, 28*), these aggregated methylation signals indicate the methylation of cell fractions at the nucleotide level (hereafter referred to as methylation fraction). The methylation fraction of G<u>A</u>NTCs was observed in heterogeneity across samples, particularly in St6, St8, and St9. The differences in the methylation fractions on nucleotide sites were referred to as single nucleotide methylation variation (SNMV). Compared to other samples, the number of unmethylated motifs was higher in St6, St8, and St9 and showed different SNMV patterns at each nucleotide position (Fig. 4A).

**Fig. 4.**
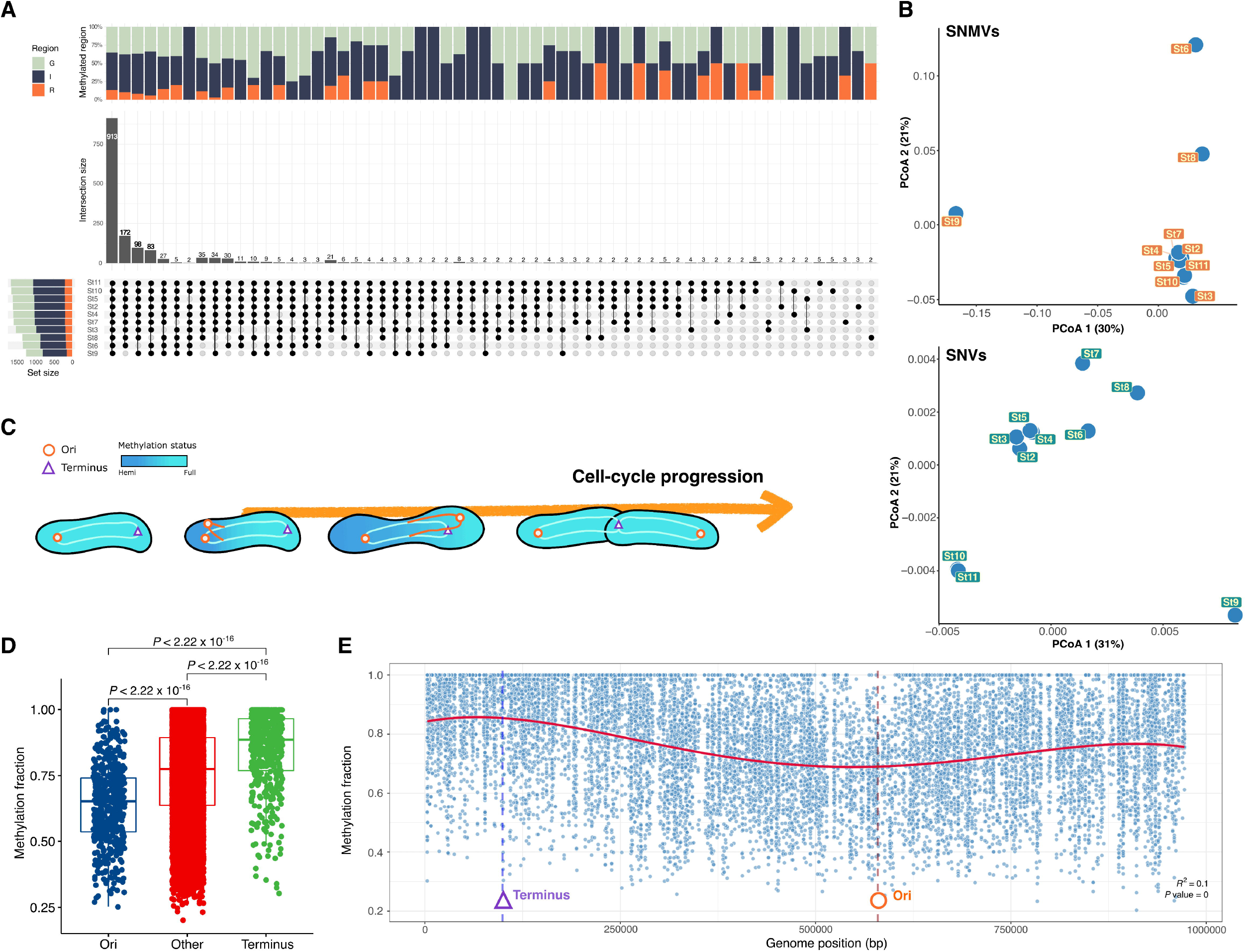
Genome-wide epigenetic analysis of *Pelagibacter* MAG across samples. (**A**) The UpSet plot compares GANTC methylation at each genome position on a *Pelagibacter* MAG (SHIPPO_PRO_33) across samples. More than 0.5 of the methylation fractions were considered methylated at each genome position; the color of each bar depends on the genic (G: green), intergenic (I: navy), and regulatory region (R: orange). The column bar indicates the intersection of the number of methylation positions on the MAG across samples. The left bar represents the total number of methylated positions on the MAG for each sample. (**B**) Principal coordinate analysis (PCoA) clustering by SNMV and SNV based on the Bray–Curtis distance on overlapped regions for all samples. (**C**) A model of the methylation pattern according to the cell-cycle progression in *Alphaproteobacteria*. (**D**) The methylation fraction comparisons of the GANTC motif between genomic regions of *ori* (replication origin), *ter* (replication terminus), and other regions. (**E**) The genome-wide distribution of methylated fractions for the GANTC motif indicates the trend of cell-cycle progress throughout the genome. MAG: metagenome-assembled genome; SHIPPO: Shipborne Pole-to-Pole Observations; SNMV: single nucleotide methylation variation; SNV: single nucleotide variant.

By contrast, all samples, except for St6, St8, and St9, had similar SNMVs to each other and were more closely clustered, regardless of latitude (Fig. 4B). For example, pairs St2 and St3, and St10 and St11, distanced geographically by about 20° latitude and 3,000 km distance, were grouped closely through PCoA (Fig. 4B). In particular, differences in strain-level composition (single nucleotide variants; SNVs) were found between these two groups, but in SNMVs, these groups were clustered together (Fig. 4B). These results indicated that DNA methylation at the single nucleotide level differed under environmental conditions regardless of strain composition, which suggests that dynamic cellular events occur among various *Pelagibacter* in northwest Pacific Ocean surface waters.

### Cell-cycle regulation of Pelagibacter by MTase activity

A gradual decrease was observed in the exponential phase of the genome-wide methylation fraction of GANTC from the replication origin (*ori*) to replication terminus (*ter*) of the *Cand*. P. Giovannoni NP1 chromosome (fig. S4). In addition, the same pattern was observed with the *Pelagibacter* proMAG (SHIPPO_PRO_33) across several samples under real environmental conditions (Fig. 4D). The MTase CcrM, which methylates the GANTC motif, is a representative example of an MTase involved in the cell-cycle regulation in Alphaproteobacteria (*29*). As chromosomal replication proceeds, the methylation status of the parental strand is maintained, whereas GANTC remains unmethylated when a new daughter strand is generated through a replication fork (hemimethylated) (Fig. 4C). *CcrM* is only expressed at the end of chromosome replication, so the chromosome remains hemimethylated until the end of replication. When DNA methylation signals are pooled from separate molecules for each genomic locus, the methylation fraction may result in relatively lower values near the *ori* due to the ratio of hemimethylation to full methylation following different DNA replication instances for different cells in the exponential phase (fig. S4). Based on the above hypothesis, a relatively lower methylation fraction in the *ori* of the *Pelagibacter* chromosome suggests that GANTC methylation may also be involved in the cell cycle of *Pelagibacter*. As in the nutrient-rich laboratory environment, the same pattern of methylation fractions in the *Pelagibacter* chromosome was observed in marine environments (Fig. 4E); this suggests that *Pelagibacter* is also in the exponential phase in real marine environments.

### DNA methylation of the viral genome and the possibility of legacy from the host

The RM system is ubiquitous in about 90% of the bacterial genome (*30*) and acts as a defense mechanism to distinguish between self and non-self DNA. However, due to the lack of cognate REases in most genomes in this study, DNA methylation is thought to be used to regulate the cellular mechanisms rather than defense mechanisms of exogenous invasive DNA, such as the RM system, the BacteRiophage EXclusin (BREX) system (*31*), and the Defense Island System Associated with restriction–modification (DISARM) (*32*) in the ocean microbial communities. We only observed methylation patterns in 83 vMAGs associated with 67 vOTUs (viral operational taxonomic units) in a total of 15,056 putative viral genomes. The 83 vMAGs grouped into five major lineages based on pairwise genome similarity (Fig. 5A). Most of the clades were composed of vMAGs belonging to the *Caudovirales* order, except for clade V which showed genomic similarity to high-quality gvMAGs, and is likely composed of genome fragments related to *Phycodnaviridae*. Within the individual clades, there were typically two or three distinct subclades with outgroups, and some methylation patterns were shared between subclades. For instance in clade-III, specific adenine methylation motifs were found within subclades, which were not shared with other clades. Although these subgroups are phylogenetically related, the two different methylated adenine motifs (GRRGA/None and CCATC/GATGG) did not overlap and represent entirely different motif patterns. Cytosine methylation was dominant throughout clade-II, and methylation was observed in CGCG, GAGCTC, and CYCGRG motifs according to subgroups. On the other hand, both adenine and cytosine methylation were found in clade-IV, and specific methylation was indicated in the GANTC and CCWGG motifs, respectively, depending on the clade. In addition, the methylation in the GATC motif was consistent in clade-I and, together with the motifs found in clades-IV, suggests that it may be associated with bacterial methylation system as a methylation motif frequently found in Proteobacteria. For clade-V, adenine methylation was detected, as previously reported for *Phycodnaviridae* (33); in our study, these were found on GATC and CATG motifs. These two methylated motifs were observed spanning genomes belonging to *Phycodnaviridae*, and importantly the same motif has been previously reported in some of their green algae hosts (15).

**Fig. 5.**
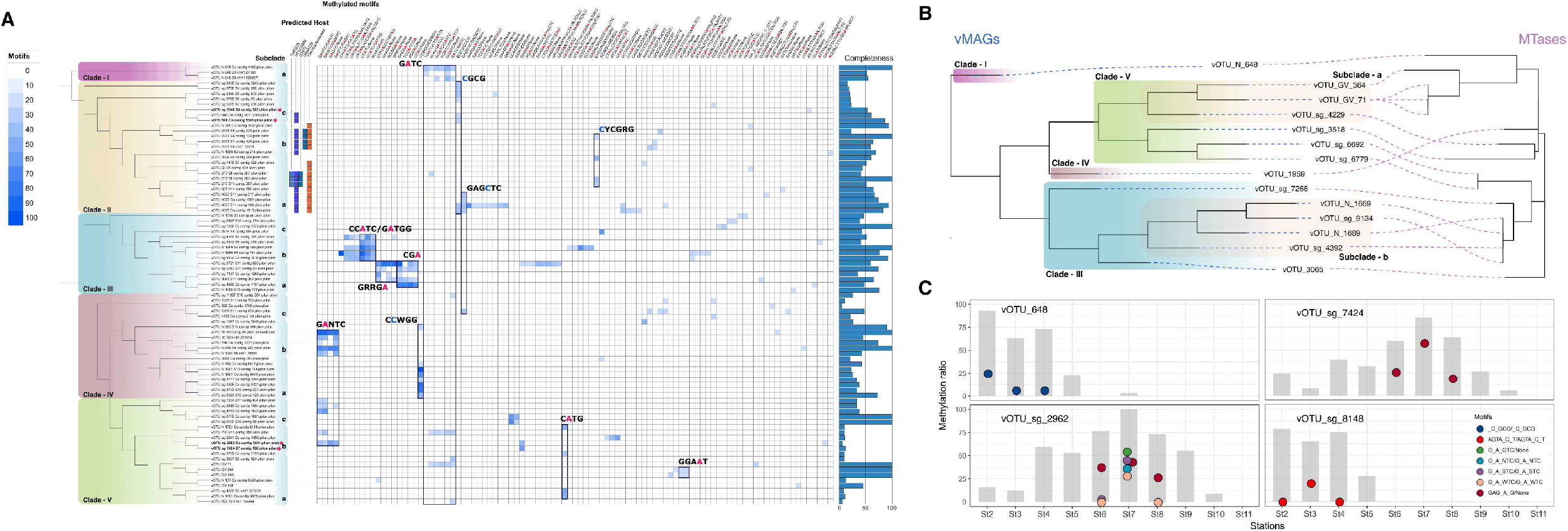
DNA methylation of the viral genome. (**A**) The methylome of 83 prevalent viral Shipborne Pole-to-Pole Observations (SHIPPO) of metagenome-assembled genomes (vMAGs) is indicated by the heatmap with their phylogeny from genome similarities. The star represents vMAGs in (C). (**B**) Phylogenetic comparison of the 14 vMAGs harboring MTase and its MTase genes. (**C**) Changes in vMAG methylation profiles were measured at three sampling stations. Circles represent the methylation ratio of each motif, and bars represent the read mapping breadth of viral genomes with 10× depth.

To investigate how viral methylation patterns are associated with MTase genes, we next evaluated the distribution of MTases according to the phylogenetic distribution of vMAG clades. Of 83 vMAGs, we found only 14 vMAGs encoded MTase genes, all of which belonged to the Type II MTase. Most MTases showed phylogenetic consistency with the vMAGs (Fig. 5B), along with consistencies in the methylated motifs. For instance, 4 vMAGs in the clade-IIIb, all methylated to the CCATC/GATGG motif were found to encode closely related MTases, yet these MTases had less than 50% similarity to compare within the REBASE. This lack of similarity to characterized MTAses was a common observation for vMAG-encoded MTAses, and there was thus only one instance of methylated motifs consistent with the predicted recognition sites based on the closest existing Mtases, for vOTU_N_648 in Clade-I. This suggests that most viral MTase systems are still poorly characterized, and we expect that methylation motifs for these novel MTases could be predicted based on the methylation patterns and phylogenetic distributions of their corresponding vMAGs. Viruses primarily use DNA methylation to counter host defense systems, such as the RM system, although they also exploit methylation to signal the transition in their status from the latent to lytic state (*34*). The methylation of viral DNA is also inherently transient as it is either derived from viral-encoded MTase or the host’s MTase and can be demethylated via passage through an MTase-free host. However, as mentioned above, bacterial DNA methylation in the marine environment does not appear to be a defense mechanism primarily. Viruses use their own MTases as a counter-defense and as a signal to initiate their lysis state and DNA packaging (*35, 36*), however incongruences were observed in our study between the methylome of the viral genome and the recognition motif of the predicted MTase. Therefore, DNA methylation of the viral genome in the marine environment is most likely a marker left behind from a previous host. To support the hypothesis, we further analyzed the DNA methylomes of the viral genomes detected at multiple sampling stations. Only four genomes had >60% of the average breadth of the genome coverage with 10x depth spanning three or more sampling stations. Notably, the DNA methylation profile differed according to the sampling station. For example, in the genome of vOTU sg 2962, most motifs associated with G<u>A</u>NTC (G<u>A</u>GTC, G<u>A</u>WTC, and G<u>A</u>STC) were methylated in St6 and St8, whereas DNA methylation was only detected in GAG<u>A</u>G in St7 (Fig. 5B). Although further studies are needed to understand the biological and ecological significance of DNA methylation in viruses in the real world, an explanation for our results is that different hosts altered the virus methylome with MTases that recognize different motifs.

### Possible implication of methylation in SAR11 phage-host interactions

Most G<u>A</u>NTC positions were methylated throughout the *Cand*. P. Giovannoni NP1 genome (fig. S4), and this methylation mark is thought to be related to cell cycle regulation (see above). However, uneven methylation patterns were observed in specific regions of the genome due to the lack of GANTC motifs in the proviral region (Fig. 6A). This raised questions about the potential use of the same methylation mark as a defense system, and the evasion of this defense system by pelagiphages (phages infecting SAR11 bacteria) at point in the evolutionary history of the phage-host pairs.

**Fig. 6.**
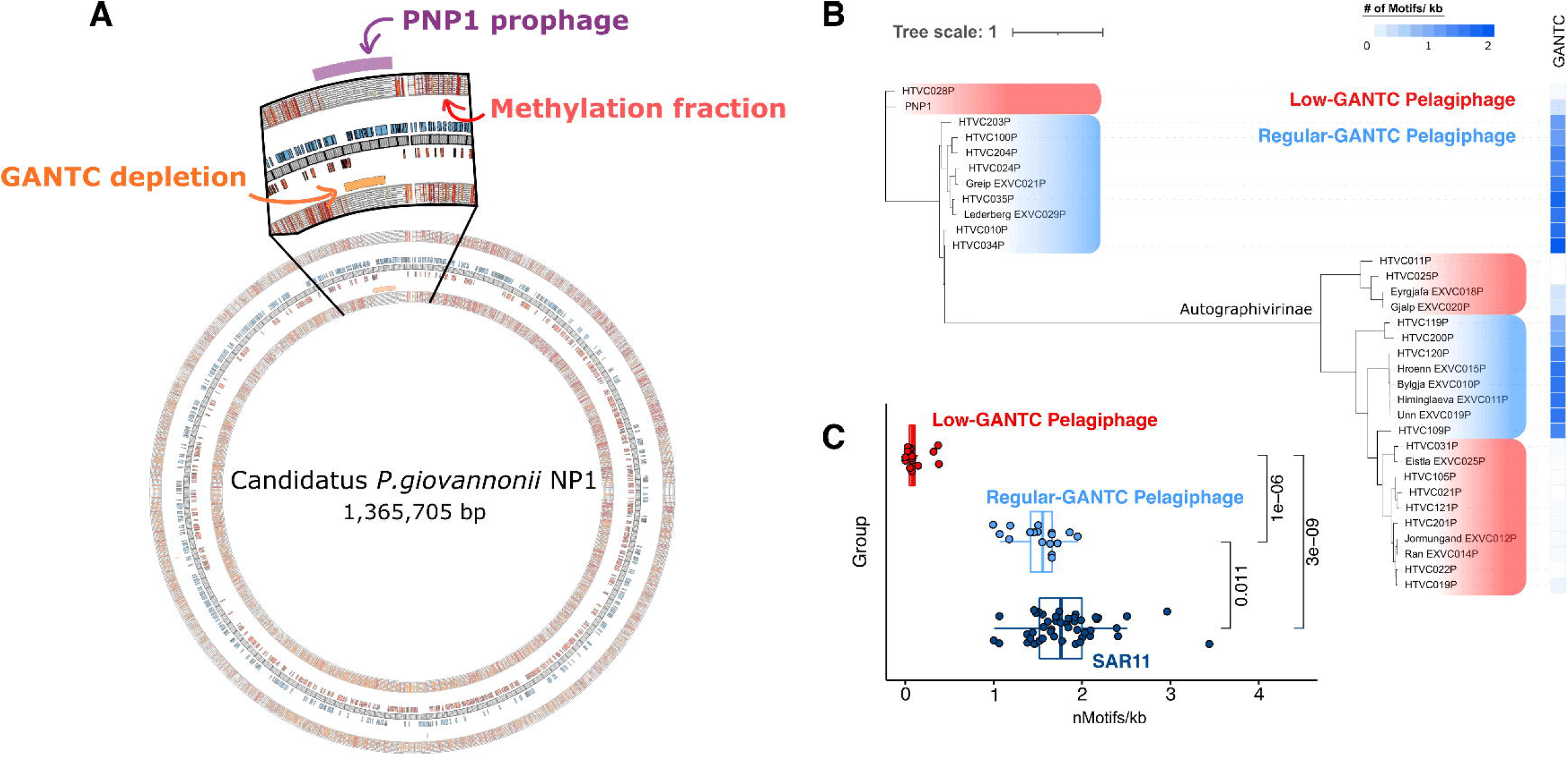
Genomic characterization between *Pelagibacter* and pelagiphage according to the GANTC motif. (**A**) Distribution of GANTC motif methylation in both strands of *Cand*. Pelagibacter Giovannoni NP1 genome. The inner blue and red bars indicate the coding sequence region of the genome. The methylation fraction bar for each strand is red for values ≥0.9, orange for ≥0.6, and bright yellow for ≥0.3. The prophage region of the genome is highlighted by a purple bar from Pleška *et al*. (*39*). The GANTC motif depletion region is marked through a multiscale signal representation (MSR) analysis. (**B**) Phylogenetic tree of pelagiphages of the head–tail connector protein. Red boxes represent the GANTC depletion subgroups of pelagiphage. (**C**) The GANTC motif density comparisons between genomes of *Pelagibacter* and pelagiphage.

In the eight proMAGs of the SHIPPO catalog spanning the order Pelagibacterales, their genomes lacked all genes associated with typical defense mechanisms including CRISPR and RM systems. While SAR11 is exposed to an environment where phage predation is frequent, it is currently thought that by reducing its genome size, it can gain an edge in the environment because of its superior nutrient uptake competitiveness (*37*). We thus interpret the lack of RM systems in SHIPPO *Pelagibacterales* proMAGs as a sign of a strong selection pressure towards genome reduction, suggesting that DNA methylation would not be currently used as an anti-phage defense by these bacterial populations.

We next reconstructed the phylogeny of 33 publicly available pelagiphage genomes based on a head–tail connector protein and compiled the frequency of GANTC motifs in the same genomes (Fig. 6B). Surprisingly, about half of the pelagiphage genomes showed a clear depletion in GANTC motif, and the calculated phylogenetic trees clearly showed that the phages clustered according to the density of GANTC motifs in their genome (Fig. 6B and 6C). This difference was particularly observed in the subfamily *Autographivirinae* of the family *Podoviridae*. This raises the possibility that methylation marks may have been used as anti-phage defense in the past, and some pelagiphages would have retained a genome composition bias since. Alternatively, it is possible that some strains within the SAR11 clade do use RM systems as phage defense, and the difference in GANTC frequency would be related to differences in host range between phage clades.

## Conclusion

Our results describe the biological meaning of DNA methylation in marine microorganisms that have not been previously interpreted from an environmental perspective. Diverse DNA methylation patterns across the kingdoms and changes in marine microbial communities have contributed to a wide range of implications, from the role of DNA methylation in marine microbes to the co-evolution history between phage-hosts. In this meta-epigenomic study, we describe how genome-wide epigenetic analysis and phage-host-related methylation could be implemented as novel interpretations of the microbiome, together with the construction of future metagenome-methylation databases.

## Acknowledgments

The work conducted by the U.S. Department of Energy Joint Genome Institute (SR) is supported by the Office of Science of the U.S. Department of Energy under contract no. DE-AC02-05CH11231.

## Data and materials availability

The raw Illumina and SMRT sequence files used in this study were deposited in the NCBI BioProject under the accession PRJNA784005. Analysis codes of the meta-epigenomics are available at the repository (https://github.com/hoonjeseong/Meta-epigenomics) under a MIT license.

## Supplementary Materials

Materials and Methods

Supplementary Text

fig. S1 to S5

table S1 to S5

