## Supplemental Materials for "Community-wide epigenetics provides novel perspectives on the ecology and evolution of marine microbiome"

**This PDF file includes:**

Materials and Methods
Supplementary Text
figs. S1 to S8
tables S1 to S5

### Materials and Methods

Sample collection and shotgun sequencing

Surface seawater samples (7 m depth) were collected at 10 stations (referred to here as St2–St11) along a transect of 44–63°N latitude (approximately 4,000 km) in the northwest Pacific Ocean during the R/V *ARAON* cruise of the Korea Polar Research Institute (KOPRI) for a Shipborne Pole-to-Pole Observations (SHIPPO) project in July 2015 (fig. S1 and table S1). Onboard, 20 L of seawater at each station was immediately filtered through a 147-mm GF/A glass microfiber filter (Whatman, Maidstone, UK) and subsequently through a 147-mm, 0.22-μm membrane filter (Millipore, Billerica, USA) using a peristaltic pump. These filters were stored at -80°C until DNA extraction. The filters were chopped and dissolved in sucrose–Tris–EDTA buffer (STE buffer; 0.74 M sucrose, 1 M Tris pH 8.3, 0.5 M EDTA, and distilled water). Next, samples were treated with lysozyme (5 mg/mL Tris-HCl), followed by 10% SDS and proteinase K (20 mg/mL). Genomic DNA was extracted using a DNeasy Blood & Tissue Kit (QIAGEN, Hilden, Germany) following the manufacturer-provided protocol. Long-read shotgun sequencing was conducted using the Pacific Biosciences (PacBio) RSII sequencing system (Menlo Park, CA, USA) with P6-C4 chemistry at DNA Link (Seoul, Korea) according to the MagBead OneCellPerWell v1 protocol. Each DNA sample was sequenced in continuous long-reading (CLR) mode with a library size of 5 kb, and three single-molecule real-time (SMRT) cells were used per sample. In addition, for short-read shotgun sequencing, libraries were constructed by TruSeq Nano and sequenced using an Illumina HiSeq 4000 system with 2 × 151 bp paired-end reads at Macrogen (Seoul, Korea).

Read preprocessing and contig assembly

Short-reads were preprocessed with FaQCs (v. 2.09; the default option with -*q* 20) (*38*), and a duplicated read removal process was performed using FastUniq (*39*). Quality-filtered reads of each sample were assembled into contigs using MEGAHIT (v. 1.1.3) (*40*) in "meta-sensitive" mode. In addition, reads of all 10 samples were assembled into contigs through co-assembly with the same options as above.

Long-reads were preprocessed with a hybrid error correction approach using FM-index Long Read Corrector (FMLRC; v. 1.0.0) (*41*) with quality-filtered short-reads. After preprocessing, corrected long-reads of each sample were assembled using Flye (v. 2.4.1) (*42*) with "meta-Flye" and "PacBio corrected reads" options. Co-assembly was also performed on all 10 samples with the same options as above. Furthermore, assembled contigs from long-reads were polished using Pilon (v. 1.23) (*43*) twice.

Genome binning of the northwest Pacific Ocean

Contig coverage was calculated using only short reads, which consisted of a uniform length (151 bp) with the Burrows–Wheeler Aligner tool (BWA-MEM v. 0.7.17) (*44*). Next, contigs from individual assembly and co-assembly were binned separately using CONCOCT (v. 0.5.0) (*45*) and MetaBAT (v. 2.12.1) (*46*), respectively. Additionally, a manual binning process was performed with mmgenome2 (https://github.com/KasperSkytte/mmgenome2). Then, these bins were refined by the integrative *k*-means clustering approach (ACR: https://github.com/hoonjeseong/acr) using contig coverages.

For prokaryotes, the quality of genome bins was estimated using CheckM (*47*). Only bins with better than medium quality (completeness >50% and contamination <10%) (*8*) were referred for metagenome-assembled genomes (MAGs) and used for further analysis. All

MAGs were dereplicated at 99% average nucleotide identity (ANI) value using dRep (*48*) with the options "-comp 50 -con 10" and then dRep was re-run with "-sa 0.95" (95% ANI value clustering) to screen representative species MAGs. To improve the quality of short- and long-read sequencing, a reassembly process was performed on only high-quality MAGs (HQ MAGs, completeness >90% and contamination <5%). Quality-filtered short- and long-reads were mapped against HQ MAGs using bowtie2 (*49*) and PBAlign (https://github.com/PacificBiosciences/pbalign). Then, mapped reads were retrieved using "samtools view" with -f 65, -f 129, and -F 4 options for forward, reverse, and unpaired long reads, respectively (*50*). Each HQ MAG was reassembled with the hybrid assembly approach using Unicycler (*51*). The obtained prokaryotic MAGs (referred to as SHIPPO PRO MAG) were designated as SHPPO_PRO_XX and classified using the Genome Taxonomy Database Toolkit (GTDB-Tk) (*11*). The phylogenic tree was constructed by PhyloPhlAn2 (*52*) and visualized by iTOL v. 6 (*53*).

For eukaryotes, eukaryotic bin classification was performed for those >5 Mb in size by EukRep (*54*). Afterward, a detailed quality evaluation was performed through EukCC (*55*), and eukaryotic MAGs were selected according to the criteria of "Completeness - 5*Contamination > 50." In addition, the MAG with the highest score (Completeness - 5*Contamination + 0.5*log(N50)) in the group within ~99% of the ANI value was selected as the representative eukaryotic MAG. Furthermore, species-level eukaryotic MAGs were chosen by the highest scores within ~95% ANI cluster. The eukaryotic MAGs (referred to as SHIPPO EUK MAG) were designated as SHPPO_EUK_XX, and their taxonomy was classified by phylogenetic topology with MAGs from Delmont *et al.* (*56*). Phylogenetic analysis was processed with two DNA-dependent RNA polymerase (RNAP-a and RNAP-b) subunits. The concatenated protein sequences of RNAP-a and -b were aligned by MAFFT alignment (v. 7.402, FFT-NS-i algorithm with default option) (*57*), and gap sequences were trimmed using trimAl (v. 1.4) (*58*) with the parameter -gt 50. The phylogenetic trees were calculated by IQ-TREE (v. 1.6.12) (*59*) using the following parameters: -m LG+F+R10 -alrt 1000 -bb 1000; LG+F+R10 model, 1,000 replication of the Shimodaira–Hasegawa (SH)-like approximation likelihood ratio (aLRT) and ultrafast bootstrap approximation (UFBoot) (*60*).

The viral contigs were detected using VirSorter (only categories 1, 2, and 3) (*61*) and the Earth's Virome protocol (*62*) from all contigs through individual and co-assemblies. CD-HIT-EST (*63*) clustering was processed with 99% identity to remove duplicate viral contigs, and only >5 kb contigs were subsequently used. Then, viral sequences were clustered as "population" with 95% ANI, which are currently defined as species-level taxonomy in viruses (viral operational taxonomic units: vOTUs)—via all-vs.-all pairwise BLASTn (*64*) using previous viral genome datasets (the Earth's Virome protocol (*65*) and the Global Ocean Viromes 2.0; GOV2.0 (*66*)). vOTUs were clustered with 95% identity across ≥80% aligned fraction, and at least one hit >1 kb in length. Groups with previous study datasets were designated as vOTU_XX, groups of two or more viral contigs only in the SHIPPO set were designated vOTU_N_XX, and singletons were designated vOTU_sg_XX (referred to as SHIPPO vOTU). For giant viruses, dereplicated viral contigs were binned using MetaBAT2 and filtered using NCLDV_detector. Then, giant viral bins were further filtered with the same conditions as in Schulz *et al.* (*67*) through the quality assessment using the copy number of NCOVGs. These giant viral MAGs (referred to as SHIPPO GV) were designated as vOTU_GV_XX. Their ANI values were compared and then selected for representative strain- and species-level genomes with the completion and duplication ratio of NCOVGs. Next, the quality of vOTUs was assessed using CheckV (*68*) and divided into four groups: "complete" (0.02%, $n$ = 4), "high-quality" (2.15%, $n$ = 415), "medium-quality" (5.45%, $n$ = 1,052) and "low-quality" (92.02%, $n$ = 17,758). Taxonomic classification of vOTUs was assigned by

Demovir (https://github.com/feargalr/Demovir) with the comparison of amino acid sequence homologies to a viral subset of the TrEMBL database. Phylogenetic analyses were processed based on the Dice distance using sequence similarity comparisons of each contig using all-vs.-all tBLASTx (≥30% identity, ≥30 amino acids, and an e-value ≤0.01). The Dice distance between contigs A and B ($D_{A,B}$) was calculated as $1 - (2 \times AB / AA + BB)$, where AB is the bit-score sum of all hits between genomes A and B. AA and BB are the bit-score sums of all hits from genomes A and B, respectively, compared to themselves. Then, we obtained a phylogenetic tree from the distance matrix via a neighbor-joining algorithm.

ANI comparison with reference datasets

To ascertain the novelty of SHIPPO MAGs, we collected two genome databases that derived ocean metagenomics (the Tara Oceans survey (*9*) and the Global Ocean Reference Genomes Tropics survey; GORG-Tropics (*10*)) for comparison with our data. The quality of MAGs from Tara Oceans and single-cell assembled genomes (SAGs) from GORG-Tropics were assessed using CheckM. Genomes that were not at least medium-quality were excluded from subsequent analysis. We calculated the ANI values of SHIPPO MAGs for 957 MAGs and 4741 SAGs, respectively. Intra- and inter-species boundaries were differentiated with 95% and 83% ANI values, respectively, as previously studied. Table S1 shows all ANI values.

Sample mapping

Each short-read of a sample was subsampled at 49 million paired reads (minimum sequencing throughput among samples) and mapped to the SHIPPO MAGs using Bowtie2 (v. 2.3.4.3) (*49*) with "--very-sensitive" parameter in end-to-end mode. Additionally, to avoid overestimating the read alignment, we excluded reads whose alignments score was <-20 (tag AS:i). The percentage of mapped reads to SHIPPO MAGs (mappability) was calculated by dividing the aligned reads by subsamples (49 million) paired reads in each sample.

Diversity calculation

To calculate species-level diversity, mapped reads to species-level SHIPPO MAG were further filtered with the read alignment identity (≥95%) using pysam. Macrodiversity was calculated using Shannon diversity by the skbio diversity Python module (http://scikit-bio.org/) for each sample. Before calculating species microdiversity, we observed how many positions within the genome were shared by all samples with sufficient read depth. We only used MAGs with shared mapping positions across all samples wider than 20% of the genome breadth (60% for the viral contig) with ≥10 read depth. Then, single nucleotide variants (SNVs) were called with the following command: "samtools mpileup -l [overlapped region bed file] -0u -C 50 -Af [SHIPPO fasta] [bam files] | bcftools call -vm0 -v -o [output vcf]," where "[overlapped region bed file]" is the bed format of the overlapped region from read-mapping, "[SHIPPO fasta]" is the fasta file of species-level SHIPPO MAG sequence, "[bam files]" are all bam files that result from read mapping to SHIPPO MAG sequence, and "[output vcf]" is the output vcf file of variant information. Each SNV was considered single nucleotide polymorphism (SNP) loci by classical definition as having an alternative allele supported by a frequency >1% and at least four reads. The microdiversity (nucleotide diversity, $\pi$) of each genome was calculated using an equation from Schloissnig *et al.* (*69*). Furthermore, the fixed index ($F_{st}$)—a measure of the population differentiation between samples—was calculated by additional inter-sample nucleotide diversity calculations.

Gene annotation (methyltransferase (MTase) and restriction enzyme (REase))

Gene predictions of prokaryotes, eukaryotes, and viruses of SHIPPO MAG were processed with GeneMarkS-2 (v. 1.09_1.07_lic) (*70*), GeneMark-ES (*71*), and Prodigal (*72*), respectively. We annotated the protein-coding sequences using GhostKOALA (*73*) to generate Kyoto Encyclopedia of Genes and Genomes orthology (KO) classification.

RMS pipeline identified the restriction–modification (RM) systems (https://github.com/oliveira-lab/RMS) using Hidden Markov model (HMM) profiles built from curated reference protein sequences by a type of RM system. In addition, genes related to the RM system were obtained by its type through KO terms, as follows:

- type I RM system (K01153; type I REase, R subunit [EC:3.1.21.3], K01154; type I REase, S subunit [EC:3.1.21.3], K03427; type I REase M protein [EC:2.1.1.72])
- type II RM system (K01155; type II REase [EC:3.1.21.4], K06223; DNA adenine methylase [EC:2.1.1.72], K13581; modification methylase [EC:2.1.1.72], K07317; adenine-specific DNA-MTase [EC:2.1.1.72], K07318; adenine-specific DNA-MTase [EC:2.1.1.72], K07319; adenine-specific DNA-MTase[EC:2.1.1.72], K00571; site-specific DNA-MTase (adenine-specific) [EC:2.1.1.72], K00558; DNA (cytosine-5)-MTase 1 [EC:2.1.1.37], K00590; site-specific DNA-MTase (cytosine-N4-specific) [EC:2.1.1.113])
- type III RM system (K01156; type III REase [EC:3.1.21.5], K07316; adenine-specific DNA-MTase [EC:2.1.1.72])
- type IV RM system (K07451; 5-methylcytosine-specific REase A [EC:3.1.21.-], K07452; 5-methylcytosine-specific REase B [EC:3.1.21.-], K19147; 5-methylcytosine-specific REase subunit McrC, K07448; restriction system protein)

The RM system type was classified by searching genes encoding the MTase and REase that were <10 genes apart.

Host-prediction

We conducted multiple strategies for viral host prediction to identify potential microbial hosts of the vOTUs from SHIPPO MAG. 1) Sequence homology matches: SHIPPO vOTU genomes were aligned against SHIPPO PRO MAG using BLASTn. Among the best hits, hits that met the criterion of ≥75% identity with alignment length ≥2,500 bp were considered. 2) Clustered Regularly Interspaced Short Palindromic Repeats (CRISPR) spacer matches: CRISPR spacers were identified using CRT (CLI v. 1.2) (*74*) and PILER-CR (v. 1.06) (*75*) using the criteria established in the Microbial Genome Annotation Pipeline (MiGAP v. 4) (*76*). Spacer sequences were compared by all-vs.-all BLASTn with "-task BLASTn-short -evalue 1e-10 -perc_identity 95" parameters. 3) tRNA matches: transporter RNAs were obtained using ARAGORN (v. 1.2.38) (*77*) and compared by all-vs.-all BLASTn with "-perc_identity 100" parameter. Only BLAST hits were considered with 100% coverage query length of the alignment. We matched the host of SHIPPO vOTU against SHIPPO PRO MAG through the filtered results from the three abovementioned strategies.

DNA modification and motif detection

DNA methylation was detected with a series of pipelines from BaseMod (https://github.com/ben-lerch/BaseMod-3.0). Briefly, each long-read of samples was mapped to the SHIPPO MAG sequences using PBAlign. Then, the modified positions of genomes were identified through ipdSummary (v. 2.0), which is a *t*-test based comparison of interpulse duration ratios (IPDs) to the "synthetic control" model. These approaches used the same parameters documented in BaseMode. To find novel motifs methylated from each SHIPPO MAG, we found methylated sequence motifs by a *de novo* approach using MultiMotifMaker (*14*). Additionally, we collected the previously curated recognition motifs of MTase from the Restriction Enzyme Database (REBASE; http://rebase.neb.com/cgi-bin/msublist) (*13*) to further capture methylated motifs in SHIPPO MAG.

Epi-metagenomic analysis

Due to the specificity of the metagenome, insufficient reads were sequenced to calculate methylation for all SHIPPO MAGs. Therefore, using a relatively weak criterion of 10× coverage per strand and ≥30 modification quality value, we identified the overall $N^6$-methyladenosine ($m^6A$) and $N^4$-methylcytosine ($m^4C$) DNA modification trends throughout SHIPPO MAGs across the kingdoms of life. To complement this point, the ratio of methylated motifs for each genome in each sample was calculated only within a genomic breadth of coverage in 10×. Additionally, when the genome breadth of coverage was less than a certain percentage of each MAG size (SHIPPO PRO <20%, SHIPPO EUK <10%, SHIPPO vOTU <60%, SHIPPO GV <20%; according to genome size), they were excluded from further analysis.

MTase sequences were downloaded from REBASE (http://rebase.neb.com/cgi-bin/msubprolist), and the most similar MTase was searched for by querying the candidate MTase of SHIPPO MAG with a ≥50% identity parameter using BLASTp in REBASE. Afterward, we compared the motifs obtained from the methylation analysis of SMRT (which were >20% methylated) sequencing with the recognition motif of the most similar reference MTase.

Genome-wide epi-metagenomic analysis

To compare DNA methylation profiles at single-nucleotide resolution across samples, we selected the most dominant MAGs (SHIPPO_PRO_33) that mapped ≥50% of the genomic breadth of 20× per strand in all samples. MAGs consist of fractionated genomes, making it difficult to understand the overall genomic structure, so we rearranged the order of the contigs through genome comparison with the closest reference genome using the Mauve Contig Mover (*78*). Then, the replication origin (*ori*) and terminus (*ter*) were identified using the gc_skew.py script (https://github.com/christophertbrown/iRep) through the concatenate reordered contigs. In addition, methylation fractions (cellular fractions of DNA methylation) in regions overlapped by 20× in all samples were compared at the single nucleotide level.

Motif enrichment

Motif enrichment analysis was processed by calculating the motif densities of three regions: regulatory, genic, and intergenic. Five hundred simulations were performed by shuffling the coding region based on the average gene length and the number of genes present in each genome strand, and the Shapiro test (>0.05) was performed for each of the three regions. Next, *z*-scores were calculated for the motifs following normality and converted into *p*-values. Moreover, we used a multiscale signal representation (MSR) (*79*) method to assess whether the motif for specific regions is enriched or depleted ("section_2" of https://github.com/oliveira-lab/BEAST).

Principal coordinate analysis (PCoA)

PCoA was performed based on the SNVs, methylated motifs, and methylation fractions across samples with the Bray–Curtis distance with R package “vegan” and visualized with “ggplot2”.

**Supplementary Text**

DNA methylome measurement in metagenomic data

The existing methylated motif (methylome) analysis focused on the isolated bacterial genome, which allowed to determine the correct methylated motif from the sufficient coverage of SMRT sequencing. However, because metagenomic data is widely distributed in the sequences of various organisms, reads could not cover the entire genome with sufficient coverage, which is a limitation in analyzing the methylome using existing methods. Therefore, we considered setting the methylome analysis in the metagenomic study to minimize the error of methylome measurement by changing the mapped coverage threshold.

We used the genomes of six bacteria belonging to the Alphaproteobacteria obtained from the Blow *et al*. study (*20*) to benchmark the DNA methylome measurement. Based on the methylated GANTC proportion (the number of a methylated motif/the number of a motif) of the entire genome, we compared how much the methylated GANTC proportion changes depending on the fraction on the genome. The methylation proportion was calculated on the region of the genome fraction selected randomly according to the threshold 1~95% with 100 iterations each.

We calculated the residuals between the predicted value (methylation proportion by genome fraction) and the measured value (methylation proportion of entire genome) (fig. S7). We found that calculating the methylation profile using fractions of 20% or more of the genome is a method with minimal error. These results demonstrate the potential to analyze the profile of methylation motifs in various organisms even with fragmented parts of the metagenomic data.

Benchmarking of viral taxonomy classification

Considering that it is difficult to classify the taxonomy of most marine viruses, we performed a family-level classification benchmarking test to obtain the optimal parameters to minimize false positives. Viral MAGs (vMAGs) constituted 5,282 vOTU clusters with genomes from previous viral genome datasets, of which only 1,176 vOTU clusters were classified at the family level through the GOV2.0 study (*67*) (Not a single Earth virome genome composed of vOTUs with vMAG has been classified). A total of 3,575 vMAGs belonging to 1,176 vOTU clusters were considered true positives (*TP*) when the Demovir results were consistent. Conversely, if the result of vMAG classified through Demovir was an incorrect assignment compared to the family classification of GOV2.0, it was considered a false positive (*FP*).

Precision quantified the quality through the Demovir classification approach in terms of the reliability of its assignment. In addition, the prevalence value was calculated to determine how many vMAGs could be classified appropriately (*TP*) according to the threshold of the number and fraction of viral genes in each vMAG.

$$\text{Precision} = \frac{TP}{TP + FP} \qquad \text{Prevalence} = \frac{TP}{\text{Total number of vMAGs}}$$

We compared the precision and prevalence values to reduce *FP*s and classify as many genomes as possible. When 10% of the genes in each vMAG were viral, the precision value was 92.93, and 89.76% of the vMAGs were correctly classified. Moreover, when 15% of the genes in each vMAG were viral, the precision value was 94.31, and 85.45% of the vMAGs were correctly classified (fig. S8). We have demonstrated that the classification approach using the percentage of viral genes in the genome is efficient, and we believe that it will be helpful to classify unidentified viral genomes.

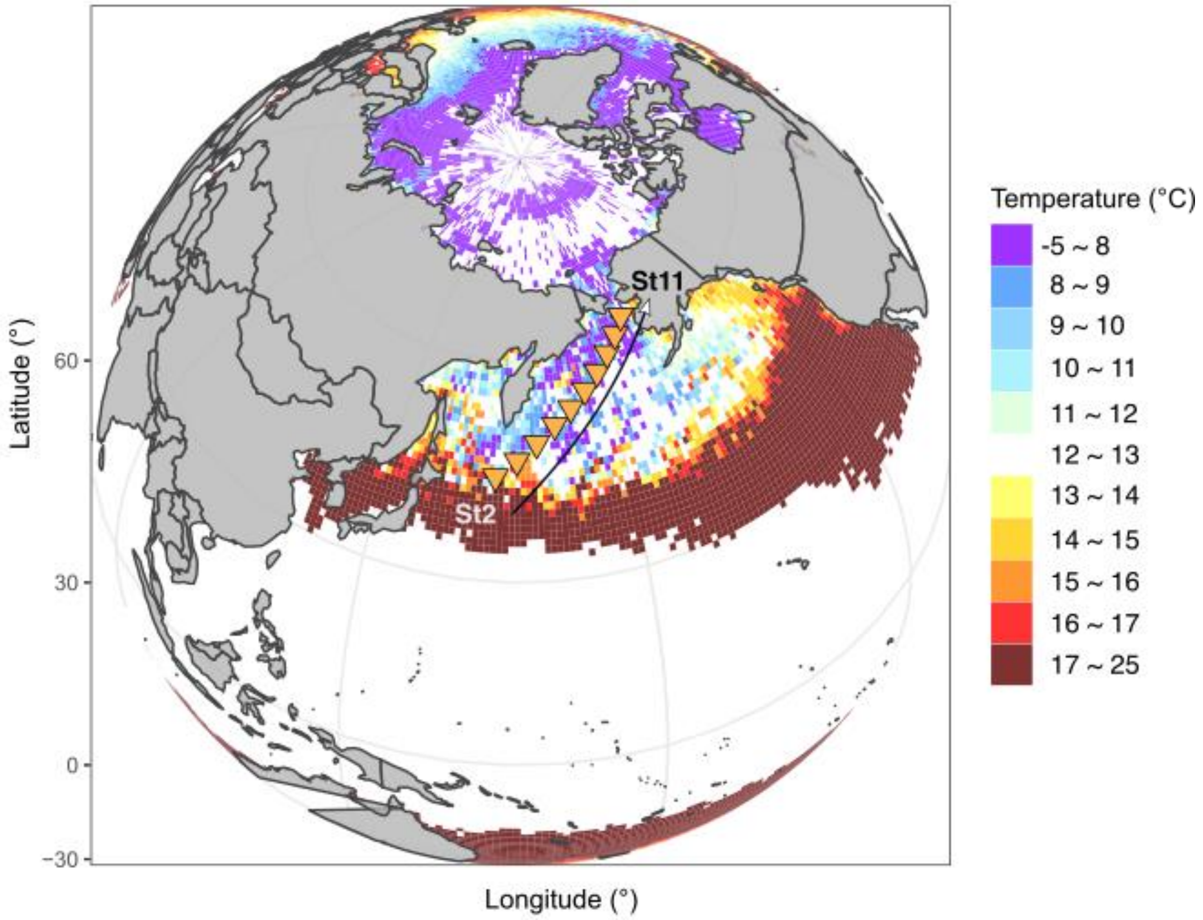

**fig. S1. SHIPPO samples in the northwest Pacific Ocean.**

The yellow triangle represents the 10 sampling stations (Station2–Station11: St2–St11) visited by the icebreaker *ARAON*, which sailed to the North Pole in 2015. SHIPPO: Shipborne Pole-to-Pole Observations.

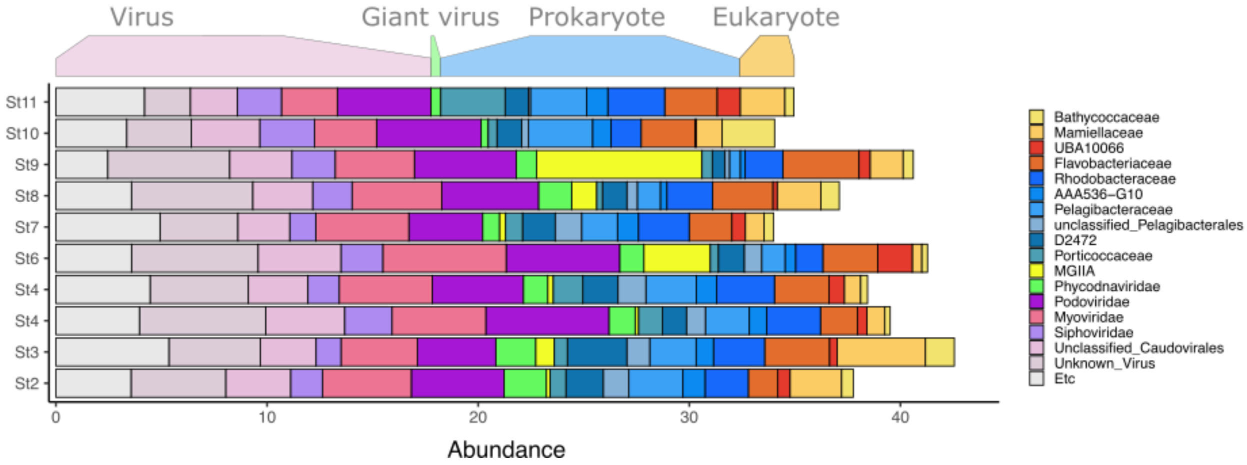

**fig. S2. Compositional profile of microbial communities spanning eukaryotes, prokaryotes, and viruses.**

Each colored bar represents the read-mapped percentage to the phylum level of the assembled genome; the *y*-axis indicates sampling stations St2–St11.

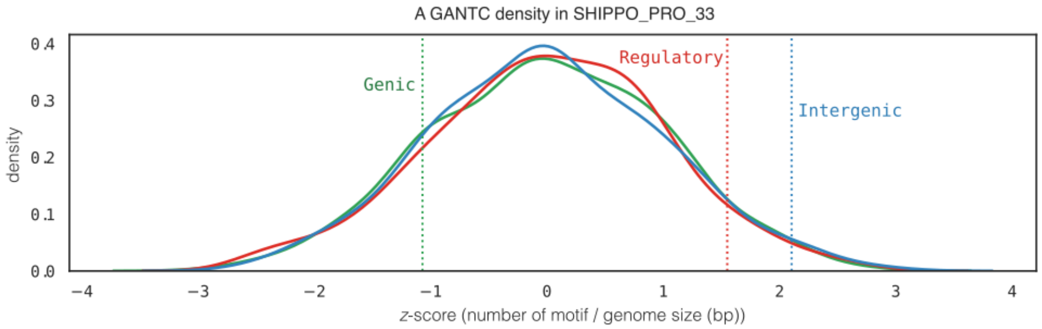

**fig. S3. GANTC motif density in the SHIPPO_PRO_33 genome.**

A randomized density of GANTC motif on the genic, intergenic, and regulatory regions throughout the *Pelagibacter* metagenome-assembled genome (SHIPPO_PRO_33) were calculated as *z*-scores. The dotted lines represent the *z*-scores for the empirical density of GANTC motifs on these regions. SHIPPO: Shipborne Pole-to-Pole Observations.

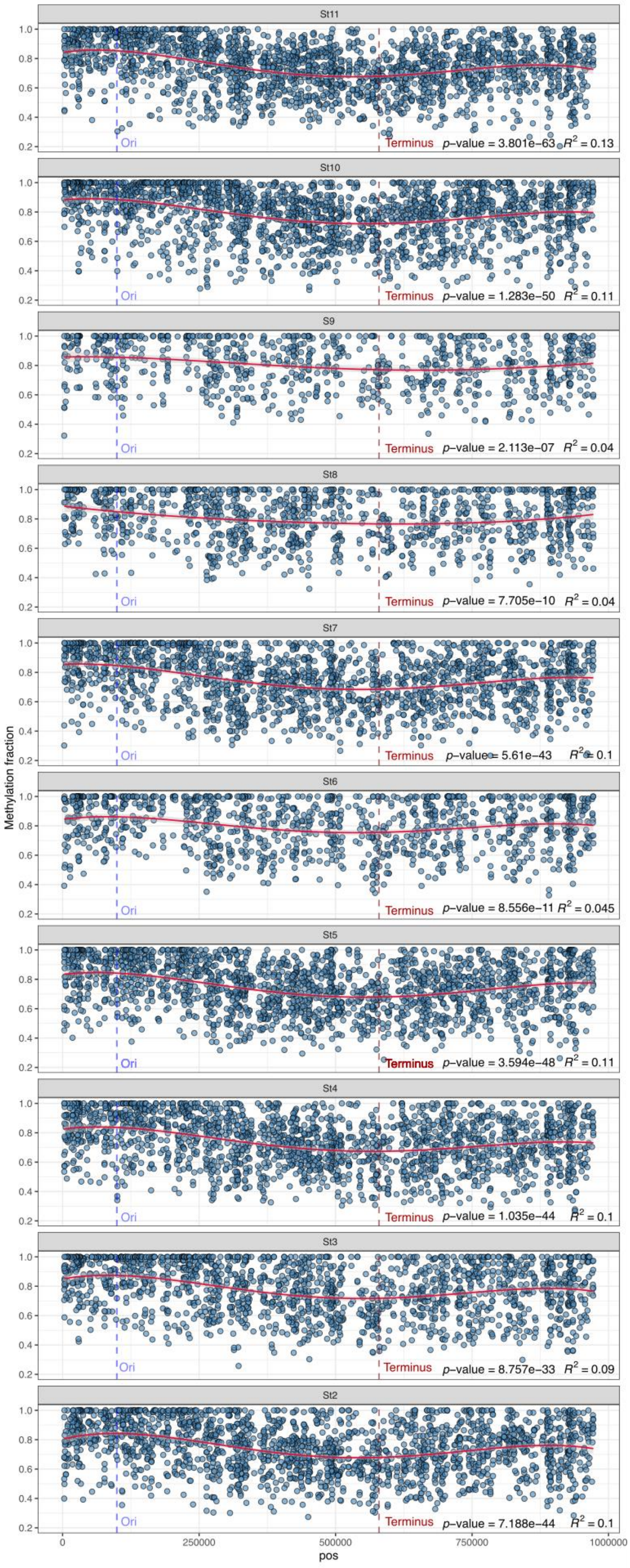

**fig. S4. The *Pelagibacter* (SHIPPO PRO 33) genome-wide distribution of methylated fractions.**

The genome-wide distribution of methylated fractions for GANTC motifs throughout the SHIPPO PRO 33 genome across stations. SHIPPO: Shipborne Pole-to-Pole Observations.

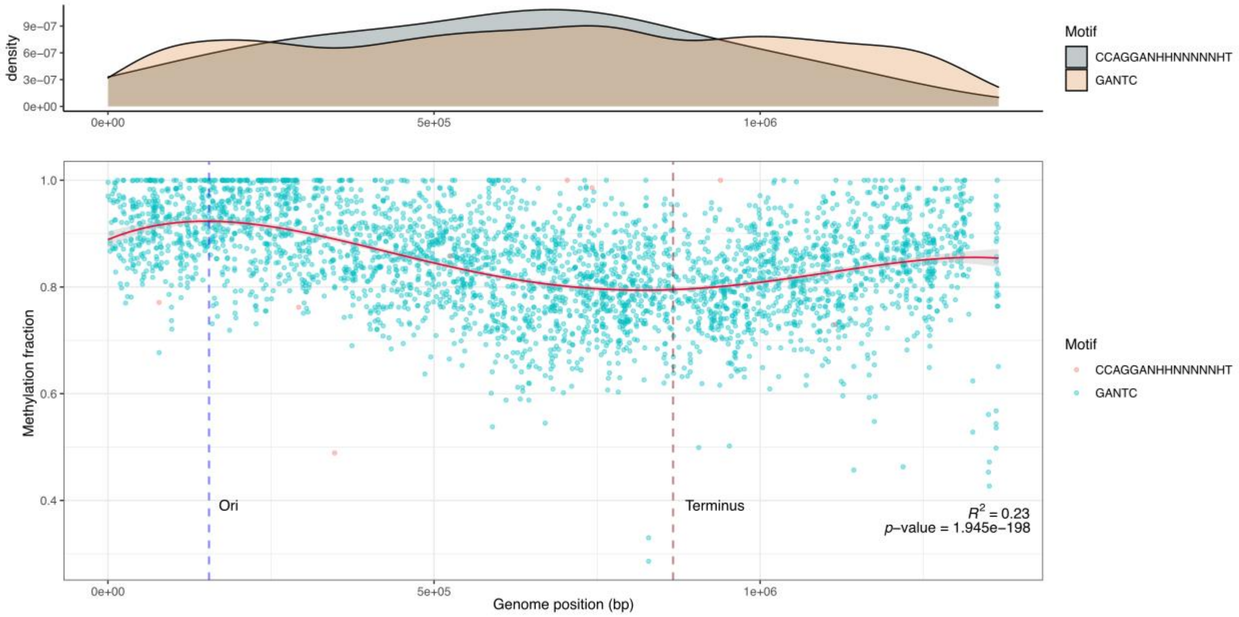

**fig. S5. The *Cand.* P. Giovannoni NP1 genome-wide distribution of methylated fractions.** The genome-wide distribution of methylated fractions for GANTC and CCAGGAHHNNNNNHT motifs throughout the *Candidatus* Pelagibacter Giovannoni NP1 genome.

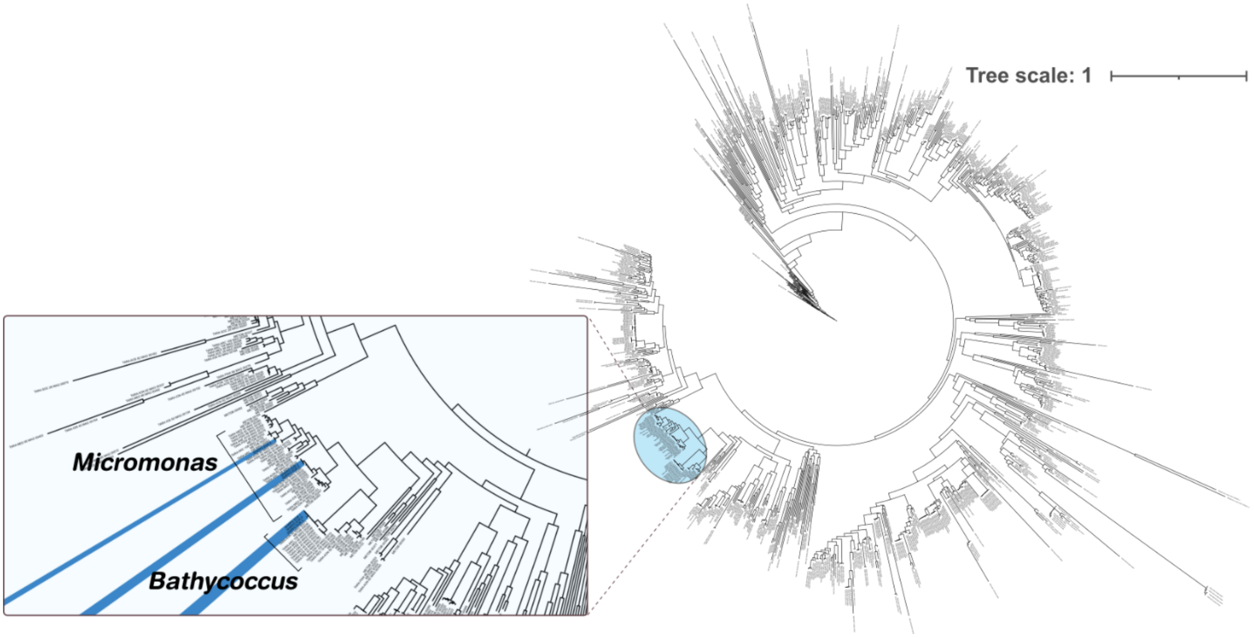

**fig. S6. Phylogenetic tree of eukaryotic plankton MAGs obtained from SHIPPO.**
The phylogenetic trees of MAGs were reconstructed by RNA polymerase protein sequences. Taxonomy of a total of six strain-level eukaryotic MAGs (colored blue) was inferred by comparing RNA polymerase from Delmont *et al.* (15). MAG: metagenome-assembled genome; SHIPPO: Shipborne Pole-to-Pole Observations.

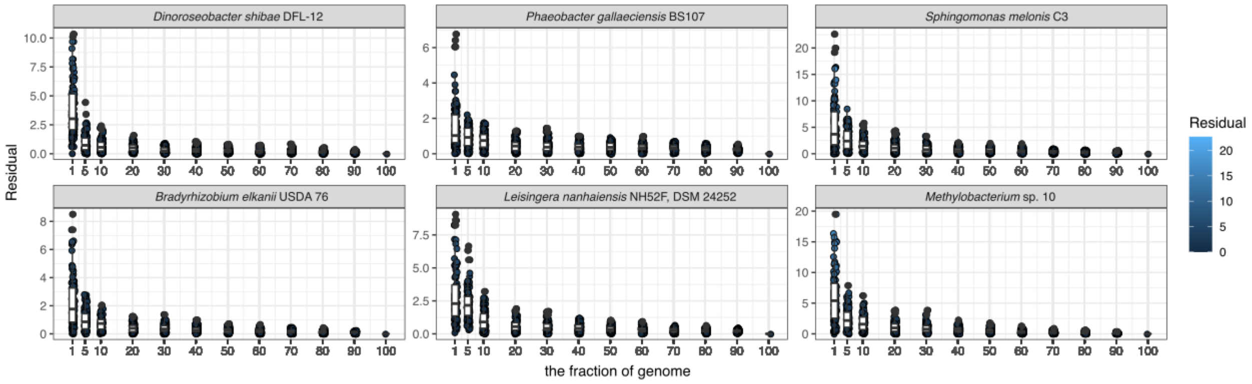

**fig. S7. The GANTC methylation proportion comparison according to the genome fraction in six species belonging to the Alphaproteobacteria.**

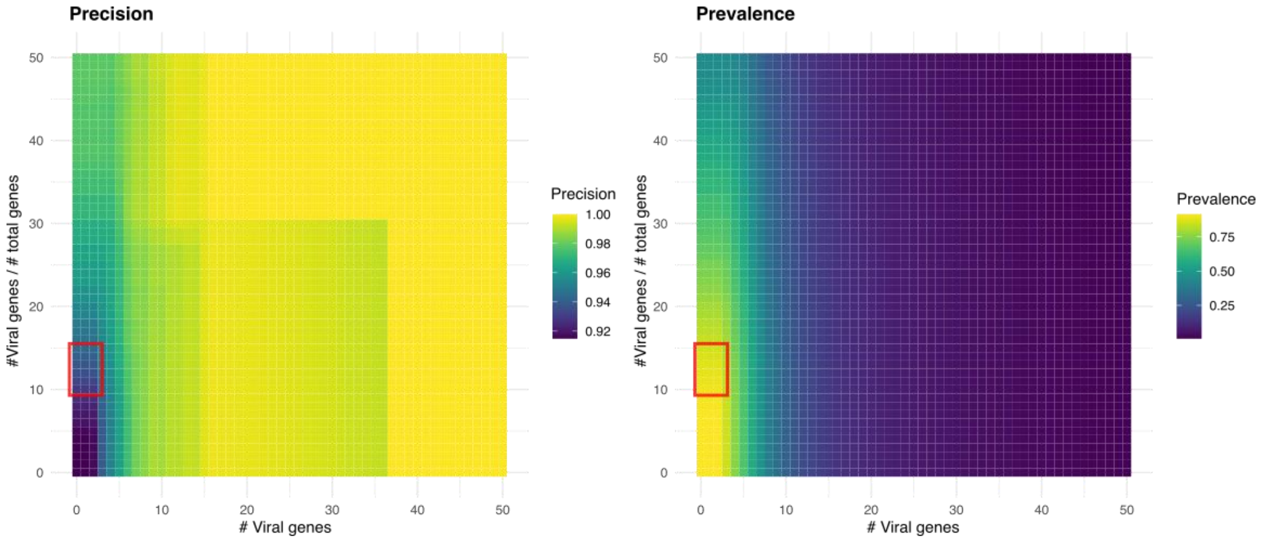

**fig. S8. Benchmarking of viral taxonomy classification according to the number and fraction of viral genes.**

**table S1. SHIPPO samples in the northwest Pacific Ocean.**

| Station | Date | Latitude (°) | Longitude (°) | Depth (m) | Temperature (°C) | Salinity (psu) |
|---|---|---|---|---|---|---|
| St2 | 2015-07-21 | 44.38 (N) | 155.35 (E) | 7 | 12.09 | 32.15 |
| St3 | 2015-07-22 | 46.60 (N) | 159.43 (E) | 7 | 9.46 | 32.23 |
| St4 | 2015-07-23 | 48.86 (N) | 163.52 (E) | 7 | 10.09 | 32.27 |
| St5 | 2015-07-24 | 51.10 (N) | 167.60 (E) | 7 | 10.71 | 32.33 |
| St6 | 2015-07-24 | 53.08 (N) | 171.64 (E) | 7 | 11.47 | 32.17 |
| St7 | 2015-07-25 | 55.07 (N) | 175.68 (E) | 7 | 12.20 | 32.34 |
| St8 | 2015-07-26 | 57.05 (N) | 179.73 (E) | 7 | 12.26 | 32.17 |
| St9 | 2015-07-27 | 59.03 (N) | 176.23 (W) | 7 | 12.55 | 32.16 |
| St10 | 2015-07-28 | 61.02 (N) | 172.19 (W) | 7 | 11.05 | 30.54 |
| St11 | 2015-07-29 | 63.00 (N) | 168.15 (W) | 7 | 10.50 | 31.18 |

SHIPPO: Shipborne Pole-to-Pole Observations.

**table S2. RM system of MAGs from global ocean studies.**

| RM system | Study | Type I | | | Type II | | | Type III | | | Type IV | | | Genome quality | | |
|---|---|---|---|---|---|---|---|---|---|---|---|---|---|---|---|---|
| | | M | R | system | M | R | system | M | R | system | M | R | system | Completeness | Contamination | Genome_size (Mb) |
| TARA_PSE_MAG_00001 | TARA | 0 | 0 | NA | 6 | 0 | orphan | 1 | 0 | orphan | 0 | 0 | NA | 100 | 0 | 3.95 |
| TARA_PSE_MAG_00007 | TARA | 0 | 0 | NA | 7 | 1 | complete | 0 | 0 | NA | 0 | 0 | NA | 100 | 0 | 3.34 |
| TARA_PSE_MAG_00009 | TARA | 1 | 1 | complete | 3 | 0 | orphan | 1 | 1 | complete | 0 | 0 | NA | 100 | 0 | 3.64 |
| TARA_ANW_MAG_00001 | TARA | 1 | 0 | orphan | 9 | 1 | complete | 0 | 0 | NA | 0 | 3 | orphan | 99.57 | 0 | 3.69 |
| TARA_PSE_MAG_00006 | TARA | 0 | 0 | NA | 5 | 1 | complete | 1 | 1 | complete | 0 | 1 | orphan | 99.46 | 0 | 3.4 |
| TARA_RED_MAG_00001 | TARA | 1 | 2 | complete | 3 | 0 | orphan | 0 | 0 | NA | 0 | 0 | NA | 99.14 | 0 | 2.85 |
| TARA_ANE_MAG_00002 | TARA | 0 | 0 | NA | 3 | 0 | orphan | 0 | 0 | NA | 0 | 0 | NA | 99.13 | 0 | 3.13 |
| TARA_PSE_MAG_00018 | TARA | 2 | 2 | complete | 5 | 0 | orphan | 0 | 1 | NA | 0 | 0 | NA | 99.05 | 0 | 4.27 |
| TARA_PSW_MAG_00010 | TARA | 0 | 1 | NA | 2 | 0 | orphan | 0 | 1 | NA | 0 | 0 | NA | 98.92 | 0 | 2.08 |
| TARA_ANW_MAG_00017 | TARA | 2 | 2 | complete | 3 | 0 | orphan | 0 | 0 | NA | 0 | 0 | NA | 98.82 | 0 | 4.8 |
| TARA_PSW_MAG_00003 | TARA | 1 | 1 | complete | 3 | 0 | orphan | 0 | 0 | NA | 0 | 0 | NA | 98.39 | 0 | 2.69 |
| TARA_PSW_MAG_00002 | TARA | 0 | 0 | NA | 1 | 0 | orphan | 0 | 0 | NA | 0 | 0 | NA | 97.89 | 0 | 1.97 |
| TARA_PSW_MAG_00018 | TARA | 1 | 1 | complete | 8 | 0 | orphan | 0 | 1 | NA | 0 | 0 | NA | 97.73 | 0 | 5.86 |
| TARA_ANW_MAG_00015 | TARA | 3 | 3 | complete | 2 | 0 | orphan | 0 | 0 | NA | 0 | 0 | NA | 97.65 | 0 | 4.71 |
| TARA_MED_MAG_00006 | TARA | 0 | 0 | NA | 1 | 0 | orphan | 0 | 0 | NA | 0 | 0 | NA | 97.35 | 0 | 2.95 |
| TARA_PSE_MAG_00004 | TARA | 0 | 0 | NA | 1 | 0 | orphan | 0 | 0 | NA | 0 | 0 | NA | 97.09 | 0 | 2.2 |
| TARA_RED_MAG_00004 | TARA | 0 | 0 | NA | 2 | 0 | orphan | 2 | 0 | orphan | 0 | 0 | NA | 96.81 | 0 | 2.1 |

| | | | | | | | | | | | | | | | | |
|---|---|---|---|---|---|---|---|---|---|---|---|---|---|---|---|---|
| TARA_PSW_MAG_00048 | TARA | 0 | 0 | NA | 1 | 0 | orphan | 0 | 0 | NA | 0 | 0 | NA | 94.33 | 0 | 1.68 |
| TARA_MED_MAG_00022 | TARA | 0 | 0 | NA | 1 | 0 | orphan | 0 | 0 | NA | 0 | 0 | NA | 94.21 | 0 | 1.81 |
| TARA_IOS_MAG_00010 | TARA | 0 | 0 | NA | 1 | 0 | orphan | 0 | 0 | NA | 0 | 0 | NA | 94.2 | 0 | 1.8 |
| TARA_PSW_MAG_00027 | TARA | 0 | 0 | NA | 2 | 0 | orphan | 0 | 0 | NA | 0 | 0 | NA | 94.05 | 0 | 4.65 |
| TARA_PSE_MAG_00023 | TARA | 1 | 1 | complete | 6 | 0 | orphan | 1 | 0 | orphan | 0 | 2 | orphan | 93.18 | 0 | 4 |
| TARA_RED_MAG_00023 | TARA | 0 | 0 | NA | 2 | 0 | orphan | 0 | 0 | NA | 0 | 0 | NA | 92.74 | 0 | 1.53 |
| TARA_MED_MAG_00077 | TARA | 1 | 0 | orphan | 2 | 0 | orphan | 0 | 0 | NA | 0 | 0 | NA | 92.47 | 0 | 2.03 |
| TARA_IOS_MAG_00001 | TARA | 0 | 0 | NA | 1 | 0 | orphan | 0 | 0 | NA | 0 | 0 | NA | 91.22 | 0 | 2.08 |
| TARA_MED_MAG_00021 | TARA | 0 | 0 | NA | 1 | 0 | orphan | 0 | 0 | NA | 0 | 0 | NA | 91.21 | 0 | 2.32 |
| TARA_RED_MAG_00110 | TARA | 0 | 0 | NA | 2 | 0 | orphan | 0 | 0 | NA | 0 | 0 | NA | 89.78 | 0 | 2.15 |
| TARA_ION_MAG_00012 | TARA | 0 | 0 | NA | 1 | 0 | orphan | 0 | 0 | NA | 0 | 0 | NA | 89.54 | 0 | 1.33 |
| TARA_RED_MAG_00018 | TARA | 0 | 0 | NA | 1 | 0 | orphan | 0 | 0 | NA | 0 | 0 | NA | 89.05 | 0 | 1.1 |
| TARA_ANE_MAG_00020 | TARA | 0 | 0 | NA | 2 | 0 | orphan | 0 | 0 | NA | 0 | 0 | NA | 87.29 | 0 | 1.27 |
| TARA_PSE_MAG_00073 | TARA | 0 | 0 | NA | 1 | 0 | orphan | 0 | 0 | NA | 0 | 0 | NA | 86.75 | 0 | 0.77 |
| TARA_MED_MAG_00065 | TARA | 0 | 0 | NA | 3 | 0 | orphan | 0 | 0 | NA | 0 | 0 | NA | 86.11 | 0 | 1.79 |
| TARA_RED_MAG_00041 | TARA | 0 | 0 | NA | 1 | 0 | orphan | 0 | 0 | NA | 0 | 0 | NA | 84.79 | 0 | 2.11 |
| TARA_RED_MAG_00052 | TARA | 0 | 0 | NA | 2 | 0 | orphan | 0 | 0 | NA | 0 | 0 | NA | 84.16 | 0 | 1.2 |
| TARA_MED_MAG_00042 | TARA | 0 | 0 | NA | 1 | 0 | orphan | 0 | 0 | NA | 0 | 0 | NA | 83.52 | 0 | 1.5 |
| TARA_ASW_MAG_00011 | TARA | 0 | 0 | NA | 4 | 0 | orphan | 0 | 0 | NA | 0 | 0 | NA | 83.2 | 0 | 1.96 |
| TARA_RED_MAG_00034 | TARA | 0 | 0 | NA | 1 | 0 | orphan | 0 | 0 | NA | 0 | 1 | orphan | 83.04 | 0 | 2.44 |

| | | | | | | | | | | | | | | | | |
|---|---|---|---|---|---|---|---|---|---|---|---|---|---|---|---|---|
| TARA_ANE_MAG_00024 | TARA | 0 | 0 | NA | 3 | 0 | orphan | 0 | 0 | NA | 0 | 0 | NA | 82.93 | 0 | 1.89 |
| TARA_PSE_MAG_00116 | TARA | 1 | 1 | complete | 2 | 0 | orphan | 0 | 0 | NA | 0 | 1 | orphan | 82.59 | 0 | 4.15 |
| TARA_PSW_MAG_00036 | TARA | 0 | 0 | NA | 1 | 0 | orphan | 0 | 0 | NA | 0 | 0 | NA | 79.29 | 0 | 1.17 |
| TARA_ASW_MAG_00029 | TARA | 4 | 1 | orphan | 3 | 0 | orphan | 1 | 1 | complete | 0 | 4 | orphan | 77.76 | 0 | 3.86 |
| TARA_PSE_MAG_00045 | TARA | 0 | 0 | NA | 1 | 0 | orphan | 0 | 0 | NA | 0 | 0 | NA | 77.37 | 0 | 1.52 |
| TARA_ANW_MAG_00033 | TARA | 0 | 0 | NA | 2 | 0 | orphan | 0 | 0 | NA | 0 | 0 | NA | 76.67 | 0 | 1.32 |
| TARA_PSW_MAG_00057 | TARA | 1 | 0 | orphan | 0 | 0 | NA | 0 | 0 | NA | 0 | 0 | NA | 76.4 | 0 | 1.44 |
| TARA_PSW_MAG_00097 | TARA | 0 | 0 | NA | 2 | 0 | orphan | 0 | 0 | NA | 0 | 0 | NA | 76.06 | 0 | 3.56 |
| TARA_IOS_MAG_00030 | TARA | 0 | 0 | NA | 1 | 0 | orphan | 0 | 0 | NA | 0 | 0 | NA | 76 | 0 | 1.23 |
| TARA_PSE_MAG_00120 | TARA | 0 | 0 | NA | 0 | 0 | NA | 0 | 0 | NA | 0 | 1 | orphan | 75 | 0 | 2.77 |
| TARA_PON_MAG_00043 | TARA | 0 | 0 | NA | 1 | 0 | orphan | 0 | 0 | NA | 0 | 0 | NA | 74.75 | 0 | 1.91 |
| TARA_ANE_MAG_00055 | TARA | 0 | 0 | NA | 1 | 0 | orphan | 0 | 0 | NA | 0 | 0 | NA | 72.69 | 0 | 1.54 |
| TARA_SOC_MAG_00008 | TARA | 0 | 0 | NA | 1 | 0 | orphan | 0 | 0 | NA | 0 | 0 | NA | 72.29 | 0 | 1.15 |
| TARA_IOS_MAG_00063 | TARA | 2 | 2 | complete | 2 | 0 | orphan | 0 | 0 | NA | 0 | 0 | NA | 67.41 | 0 | 2.75 |
| TARA_MED_MAG_00067 | TARA | 0 | 0 | NA | 0 | 0 | NA | 1 | 0 | orphan | 0 | 0 | NA | 66.91 | 0 | 1.3 |
| TARA_ANW_MAG_00045 | TARA | 0 | 0 | NA | 1 | 0 | orphan | 0 | 0 | NA | 0 | 0 | NA | 66.53 | 0 | 1.63 |
| TARA_PON_MAG_00072 | TARA | 0 | 0 | NA | 2 | 0 | orphan | 0 | 0 | NA | 0 | 0 | NA | 66.38 | 0 | 2.33 |
| TARA_PSW_MAG_00123 | TARA | 0 | 0 | NA | 2 | 0 | orphan | 0 | 0 | NA | 0 | 0 | NA | 66.11 | 0 | 2.66 |
| TARA_PSE_MAG_00063 | TARA | 0 | 0 | NA | 1 | 0 | orphan | 0 | 0 | NA | 0 | 0 | NA | 65.73 | 0 | 1.02 |
| TARA_PSE_MAG_00112 | TARA | 0 | 0 | NA | 1 | 0 | orphan | 0 | 0 | NA | 0 | 0 | NA | 64.89 | 0 | 2.66 |

| | | | | | | | | | | | | | | | | |
|---|---|---|---|---|---|---|---|---|---|---|---|---|---|---|---|---|
| TARA_PON_MAG_00041 | TARA | 0 | 0 | NA | 3 | 0 | orphan | 0 | 0 | NA | 0 | 0 | NA | 63.14 | 0 | 2 |
| TARA_PSE_MAG_00084 | TARA | 0 | 0 | NA | 2 | 0 | orphan | 0 | 0 | NA | 0 | 0 | NA | 61.45 | 0 | 0.75 |
| TARA_ANE_MAG_00063 | TARA | 0 | 0 | NA | 1 | 0 | orphan | 0 | 0 | NA | 0 | 0 | NA | 60.82 | 0 | 1.14 |
| TARA_IOS_MAG_00036 | TARA | 0 | 0 | NA | 1 | 0 | orphan | 0 | 0 | NA | 0 | 0 | NA | 58.15 | 0 | 1.11 |
| TARA_PON_MAG_00071 | TARA | 0 | 0 | NA | 1 | 0 | orphan | 0 | 0 | NA | 0 | 0 | NA | 56.71 | 0 | 2.61 |
| TARA_ASW_MAG_00031 | TARA | 3 | 1 | complete | 0 | 0 | NA | 0 | 0 | NA | 0 | 0 | NA | 55.17 | 0 | 2.21 |
| TARA_ANE_MAG_00087 | TARA | 1 | 1 | orphan | 4 | 0 | orphan | 0 | 0 | NA | 0 | 0 | NA | 53.76 | 0 | 3.17 |
| TARA_PON_MAG_00068 | TARA | 1 | 0 | orphan | 7 | 0 | orphan | 1 | 1 | complete | 0 | 2 | orphan | 53.67 | 0 | 3.65 |
| TARA_PSE_MAG_00132 | TARA | 3 | 1 | complete | 3 | 0 | orphan | 0 | 0 | NA | 0 | 0 | NA | 52.9 | 0 | 4.21 |
| TARA_PSE_MAG_00133 | TARA | 1 | 2 | complete | 0 | 0 | NA | 0 | 0 | NA | 0 | 0 | NA | 51.03 | 0 | 2.64 |
| TARA_RED_MAG_00077 | TARA | 0 | 0 | NA | 1 | 0 | orphan | 0 | 0 | NA | 0 | 0 | NA | 50.5 | 0 | 0.91 |
| TARA_PSE_MAG_00131 | TARA | 1 | 1 | complete | 0 | 0 | NA | 0 | 0 | NA | 0 | 0 | NA | 50.17 | 0 | 2.12 |
| TARA_MED_MAG_00015 | TARA | 0 | 0 | NA | 2 | 1 | complete | 0 | 0 | NA | 0 | 0 | NA | 91.98 | 0.03 | 1.78 |
| TARA_ANW_MAG_00024 | TARA | 0 | 0 | NA | 2 | 0 | orphan | 0 | 0 | NA | 0 | 0 | NA | 81.33 | 0.04 | 1.81 |
| TARA_RED_MAG_00055 | TARA | 0 | 0 | NA | 1 | 0 | orphan | 0 | 0 | NA | 0 | 0 | NA | 79.6 | 0.06 | 1.82 |
| TARA_IOS_MAG_00048 | TARA | 0 | 0 | NA | 1 | 0 | orphan | 0 | 0 | NA | 0 | 0 | NA | 54.77 | 0.06 | 1.07 |
| TARA_IOS_MAG_00042 | TARA | 0 | 0 | NA | 1 | 0 | orphan | 0 | 0 | NA | 0 | 0 | NA | 64.11 | 0.07 | 1.09 |
| TARA_IOS_MAG_00006 | TARA | 0 | 0 | NA | 3 | 0 | orphan | 1 | 1 | complete | 0 | 0 | NA | 96.77 | 0.09 | 3.23 |
| TARA_PON_MAG_00008 | TARA | 1 | 1 | complete | 3 | 0 | orphan | 0 | 0 | NA | 0 | 0 | NA | 88.11 | 0.09 | 1.92 |
| TARA_ANE_MAG_00011 | TARA | 0 | 0 | NA | 3 | 0 | orphan | 0 | 0 | NA | 0 | 0 | NA | 94.51 | 0.1 | 2.37 |

| | | | | | | | | | | | | | | | | |
|---|---|---|---|---|---|---|---|---|---|---|---|---|---|---|---|---|
| TARA_MED_MAG_00040 | TARA | 0 | 0 | NA | 1 | 0 | orphan | 0 | 0 | NA | 0 | 0 | NA | 85.16 | 0.1 | 1.72 |
| TARA_ANE_MAG_00053 | TARA | 1 | 0 | orphan | 2 | 0 | orphan | 0 | 0 | NA | 0 | 0 | NA | 63.19 | 0.1 | 0.65 |
| TARA_PSW_MAG_00072 | TARA | 0 | 0 | NA | 2 | 0 | orphan | 0 | 0 | NA | 0 | 0 | NA | 62.12 | 0.1 | 4.1 |
| TARA_SOC_MAG_00004 | TARA | 0 | 0 | NA | 1 | 0 | orphan | 0 | 0 | NA | 0 | 0 | NA | 93.82 | 0.11 | 2.71 |
| TARA_PSW_MAG_00122 | TARA | 0 | 0 | NA | 3 | 0 | orphan | 0 | 0 | NA | 0 | 0 | NA | 65 | 0.11 | 3.24 |
| TARA_PSE_MAG_00041 | TARA | 0 | 0 | NA | 2 | 2 | complete | 0 | 0 | NA | 0 | 0 | NA | 86.98 | 0.13 | 1.11 |
| TARA_MED_MAG_00044 | TARA | 0 | 0 | NA | 1 | 0 | orphan | 0 | 0 | NA | 0 | 0 | NA | 83.95 | 0.14 | 1.98 |
| TARA_ION_MAG_00053 | TARA | 0 | 0 | NA | 1 | 0 | orphan | 0 | 0 | NA | 0 | 0 | NA | 71.08 | 0.14 | 0.64 |
| TARA_PSE_MAG_00005 | TARA | 3 | 2 | complete | 3 | 0 | orphan | 0 | 1 | NA | 0 | 0 | NA | 99.67 | 0.17 | 3.34 |
| TARA_PSE_MAG_00013 | TARA | 1 | 1 | complete | 3 | 0 | orphan | 0 | 0 | NA | 0 | 0 | NA | 98.24 | 0.18 | 3.51 |
| TARA_RED_MAG_00016 | TARA | 0 | 0 | NA | 3 | 0 | orphan | 0 | 0 | NA | 0 | 0 | NA | 88.18 | 0.18 | 1.96 |
| TARA_ANE_MAG_00014 | TARA | 0 | 0 | NA | 1 | 0 | orphan | 0 | 0 | NA | 0 | 0 | NA | 86.13 | 0.18 | 0.95 |
| TARA_PON_MAG_00001 | TARA | 1 | 1 | complete | 2 | 0 | orphan | 0 | 1 | NA | 0 | 0 | NA | 98.71 | 0.2 | 3.58 |
| TARA_ANE_MAG_00006 | TARA | 0 | 0 | NA | 2 | 0 | orphan | 0 | 0 | NA | 0 | 0 | NA | 87.85 | 0.2 | 1.42 |
| TARA_MED_MAG_00002 | TARA | 0 | 0 | NA | 1 | 0 | orphan | 0 | 0 | NA | 0 | 0 | NA | 79.71 | 0.21 | 1.43 |
| TARA_ASW_MAG_00003 | TARA | 0 | 0 | NA | 2 | 0 | orphan | 0 | 0 | NA | 0 | 0 | NA | 73.92 | 0.22 | 1.43 |
| TARA_PON_MAG_00060 | TARA | 1 | 0 | orphan | 9 | 0 | orphan | 3 | 0 | orphan | 0 | 1 | orphan | 50.32 | 0.22 | 4.64 |
| TARA_ION_MAG_00004 | TARA | 1 | 1 | complete | 1 | 0 | orphan | 0 | 0 | NA | 0 | 0 | NA | 99 | 0.24 | 3.38 |
| TARA_ION_MAG_00026 | TARA | 0 | 0 | NA | 1 | 0 | orphan | 0 | 0 | NA | 0 | 0 | NA | 69.85 | 0.24 | 1.67 |
| TARA_RED_MAG_00039 | TARA | 0 | 0 | NA | 2 | 0 | orphan | 0 | 0 | NA | 0 | 0 | NA | 93.4 | 0.25 | 1.88 |

| | | | | | | | | | | | | | | | | |
|---|---|---|---|---|---|---|---|---|---|---|---|---|---|---|---|---|
| TARA_RED_MAG_00028 | TARA | 0 | 0 | NA | 2 | 0 | orphan | 0 | 0 | NA | 0 | 0 | NA | 87.79 | 0.25 | 1.21 |
| TARA_ASW_MAG_00026 | TARA | 1 | 1 | complete | 1 | 0 | orphan | 0 | 0 | NA | 0 | 1 | orphan | 64.02 | 0.25 | 2.01 |
| TARA_PSE_MAG_00002 | TARA | 0 | 0 | NA | 5 | 0 | orphan | 0 | 0 | NA | 0 | 0 | NA | 99.84 | 0.27 | 3.41 |
| TARA_ION_MAG_00023 | TARA | 2 | 1 | complete | 0 | 0 | NA | 0 | 0 | NA | 0 | 0 | NA | 98.05 | 0.29 | 3.08 |
| TARA_RED_MAG_00098 | TARA | 0 | 0 | NA | 2 | 0 | orphan | 0 | 0 | NA | 0 | 0 | NA | 70.89 | 0.34 | 2.23 |
| TARA_ION_MAG_00056 | TARA | 0 | 0 | NA | 2 | 0 | orphan | 0 | 0 | NA | 0 | 0 | NA | 60.84 | 0.34 | 1.88 |
| TARA_ION_MAG_00037 | TARA | 3 | 3 | complete | 2 | 0 | orphan | 0 | 1 | NA | 0 | 3 | orphan | 72.71 | 0.38 | 3.19 |
| TARA_PSW_MAG_00120 | TARA | 3 | 2 | complete | 1 | 0 | orphan | 0 | 0 | NA | 0 | 0 | NA | 54.74 | 0.39 | 2.58 |
| TARA_RED_MAG_00062 | TARA | 0 | 0 | NA | 1 | 0 | orphan | 0 | 0 | NA | 0 | 0 | NA | 74.22 | 0.4 | 1.27 |
| TARA_PSW_MAG_00102 | TARA | 0 | 0 | NA | 1 | 0 | orphan | 0 | 0 | NA | 0 | 0 | NA | 62.81 | 0.4 | 1.51 |
| TARA_ANW_MAG_00003 | TARA | 1 | 1 | complete | 5 | 0 | orphan | 2 | 1 | complete | 0 | 0 | NA | 99.94 | 0.43 | 2.38 |
| TARA_RED_MAG_00007 | TARA | 0 | 0 | NA | 1 | 0 | orphan | 0 | 0 | NA | 0 | 0 | NA | 98.32 | 0.43 | 2.52 |
| TARA_PSE_MAG_00017 | TARA | 0 | 0 | NA | 1 | 0 | orphan | 0 | 0 | NA | 0 | 0 | NA | 92.11 | 0.43 | 1.22 |
| TARA_MED_MAG_00059 | TARA | 0 | 0 | NA | 1 | 0 | orphan | 0 | 0 | NA | 0 | 0 | NA | 83.85 | 0.43 | 1.21 |
| TARA_MED_MAG_00102 | TARA | 1 | 1 | orphan | 1 | 0 | orphan | 0 | 0 | NA | 0 | 0 | NA | 60.31 | 0.43 | 1.45 |
| TARA_PON_MAG_00044 | TARA | 0 | 0 | NA | 2 | 0 | orphan | 0 | 0 | NA | 0 | 0 | NA | 50.04 | 0.43 | 0.58 |
| TARA_MED_MAG_00085 | TARA | 0 | 0 | NA | 1 | 0 | orphan | 0 | 0 | NA | 0 | 0 | NA | 88.58 | 0.45 | 2.6 |
| TARA_ASW_MAG_00002 | TARA | 0 | 0 | NA | 3 | 1 | orphan | 0 | 0 | NA | 0 | 0 | NA | 99.5 | 0.5 | 4.01 |
| TARA_IOS_MAG_00003 | TARA | 0 | 0 | NA | 5 | 0 | orphan | 0 | 0 | NA | 0 | 1 | orphan | 97.9 | 0.5 | 4.06 |
| TARA_MED_MAG_00028 | TARA | 0 | 0 | NA | 1 | 0 | orphan | 0 | 0 | NA | 0 | 0 | NA | 74.25 | 0.5 | 1.54 |

| | | | | | | | | | | | | | | | | |
|---|---|---|---|---|---|---|---|---|---|---|---|---|---|---|---|---|
| TARA_PSW_MAG_00069 | TARA | 0 | 0 | NA | 1 | 0 | orphan | 0 | 0 | NA | 0 | 0 | NA | 82.68 | 0.51 | 1.78 |
| TARA_IOS_MAG_00008 | TARA | 0 | 0 | NA | 9 | 0 | orphan | 0 | 0 | NA | 0 | 0 | NA | 99.84 | 0.52 | 3.36 |
| TARA_PSE_MAG_00016 | TARA | 0 | 0 | NA | 4 | 0 | orphan | 0 | 0 | NA | 0 | 3 | orphan | 99.25 | 0.53 | 6.08 |
| TARA_IOS_MAG_00016 | TARA | 0 | 0 | NA | 1 | 0 | orphan | 0 | 0 | NA | 0 | 0 | NA | 92.11 | 0.53 | 1.77 |
| TARA_MED_MAG_00076 | TARA | 0 | 0 | NA | 1 | 0 | orphan | 0 | 0 | NA | 0 | 0 | NA | 75.6 | 0.53 | 1.48 |
| TARA_PSE_MAG_00050 | TARA | 0 | 0 | NA | 1 | 1 | complete | 0 | 0 | NA | 0 | 0 | NA | 89.25 | 0.54 | 1.49 |
| TARA_PSE_MAG_00033 | TARA | 0 | 0 | NA | 4 | 0 | orphan | 0 | 0 | NA | 0 | 0 | NA | 80.48 | 0.54 | 1.41 |
| TARA_PON_MAG_00046 | TARA | 0 | 0 | NA | 3 | 0 | orphan | 1 | 1 | complete | 0 | 0 | NA | 76.71 | 0.55 | 1.44 |
| TARA_ION_MAG_00020 | TARA | 0 | 0 | NA | 1 | 0 | orphan | 0 | 0 | NA | 0 | 0 | NA | 62.05 | 0.55 | 0.94 |
| TARA_MED_MAG_00100 | TARA | 0 | 0 | NA | 1 | 0 | orphan | 0 | 0 | NA | 0 | 0 | NA | 58.46 | 0.55 | 1.55 |
| TARA_PON_MAG_00005 | TARA | 1 | 1 | complete | 2 | 0 | orphan | 0 | 0 | NA | 0 | 2 | orphan | 95.66 | 0.58 | 2.74 |
| TARA_ANW_MAG_00051 | TARA | 0 | 0 | NA | 1 | 0 | orphan | 0 | 0 | NA | 0 | 0 | NA | 52.79 | 0.6 | 0.71 |
| TARA_SOC_MAG_00011 | TARA | 1 | 0 | orphan | 2 | 1 | complete | 0 | 0 | NA | 0 | 0 | NA | 84.2 | 0.64 | 2.52 |
| TARA_RED_MAG_00109 | TARA | 0 | 0 | NA | 1 | 0 | orphan | 0 | 0 | NA | 0 | 0 | NA | 96.09 | 0.65 | 2.71 |
| TARA_PSW_MAG_00074 | TARA | 0 | 0 | NA | 1 | 0 | orphan | 0 | 0 | NA | 0 | 0 | NA | 82.28 | 0.65 | 1.19 |
| TARA_PON_MAG_00063 | TARA | 0 | 0 | NA | 3 | 0 | orphan | 0 | 0 | NA | 0 | 0 | NA | 51.43 | 0.65 | 2.6 |
| TARA_PSW_MAG_00118 | TARA | 0 | 0 | NA | 0 | 0 | NA | 1 | 0 | orphan | 0 | 0 | NA | 50.69 | 0.65 | 2.09 |
| TARA_PSE_MAG_00026 | TARA | 1 | 2 | complete | 2 | 0 | orphan | 0 | 0 | NA | 0 | 2 | orphan | 100 | 0.66 | 3.97 |
| TARA_PSE_MAG_00038 | TARA | 0 | 0 | NA | 2 | 0 | orphan | 0 | 0 | NA | 0 | 0 | NA | 97.16 | 0.66 | 2.04 |
| TARA_RED_MAG_00038 | TARA | 0 | 0 | NA | 1 | 0 | orphan | 0 | 0 | NA | 0 | 0 | NA | 87.79 | 0.66 | 1.32 |

| | | | | | | | | | | | | | | | | |
|---|---|---|---|---|---|---|---|---|---|---|---|---|---|---|---|---|
| TARA_PSE_MAG_00066 | TARA | 0 | 0 | NA | 2 | 0 | orphan | 0 | 0 | NA | 0 | 0 | NA | 89.58 | 0.67 | 2.4 |
| TARA_MED_MAG_00036 | TARA | 0 | 1 | NA | 1 | 0 | orphan | 0 | 0 | NA | 0 | 0 | NA | 93.42 | 0.68 | 2.21 |
| TARA_IOS_MAG_00035 | TARA | 0 | 0 | NA | 1 | 0 | orphan | 0 | 0 | NA | 0 | 0 | NA | 90.31 | 0.68 | 3.25 |
| TARA_PON_MAG_00056 | TARA | 0 | 0 | NA | 1 | 0 | orphan | 0 | 0 | NA | 0 | 0 | NA | 81.64 | 0.68 | 3.84 |
| TARA_PSW_MAG_00080 | TARA | 0 | 0 | NA | 1 | 0 | orphan | 0 | 0 | NA | 0 | 0 | NA | 69.9 | 0.68 | 2.32 |
| TARA_PSW_MAG_00116 | TARA | 0 | 0 | NA | 1 | 1 | complete | 0 | 0 | NA | 0 | 0 | NA | 89.6 | 0.69 | 2.41 |
| TARA_IOS_MAG_00011 | TARA | 0 | 0 | NA | 1 | 0 | orphan | 0 | 0 | NA | 0 | 0 | NA | 88.03 | 0.71 | 1.77 |
| TARA_PSW_MAG_00068 | TARA | 0 | 0 | NA | 1 | 0 | orphan | 0 | 0 | NA | 0 | 0 | NA | 76.86 | 0.72 | 1.33 |
| TARA_ION_MAG_00036 | TARA | 0 | 0 | NA | 1 | 0 | orphan | 0 | 0 | NA | 0 | 0 | NA | 71.9 | 0.72 | 1.73 |
| TARA_PSE_MAG_00104 | TARA | 0 | 0 | NA | 3 | 0 | orphan | 0 | 0 | NA | 0 | 0 | NA | 94.96 | 0.74 | 4.7 |
| TARA_ION_MAG_00051 | TARA | 0 | 0 | NA | 1 | 0 | orphan | 0 | 0 | NA | 0 | 0 | NA | 67.18 | 0.74 | 2.59 |
| TARA_PSE_MAG_00024 | TARA | 0 | 0 | NA | 2 | 0 | orphan | 0 | 0 | NA | 0 | 1 | orphan | 97.34 | 0.75 | 4.25 |
| TARA_PSE_MAG_00025 | TARA | 0 | 0 | NA | 3 | 0 | orphan | 0 | 1 | NA | 0 | 0 | NA | 92.77 | 0.76 | 2.97 |
| TARA_ASE_MAG_00022 | TARA | 0 | 0 | NA | 1 | 0 | orphan | 0 | 0 | NA | 0 | 0 | NA | 79.96 | 0.77 | 2.27 |
| TARA_PSE_MAG_00107 | TARA | 1 | 0 | orphan | 2 | 0 | orphan | 0 | 0 | NA | 0 | 0 | NA | 57.42 | 0.79 | 2.02 |
| TARA_IOS_MAG_00022 | TARA | 0 | 0 | NA | 1 | 0 | orphan | 0 | 0 | NA | 0 | 0 | NA | 80.67 | 0.8 | 1.34 |
| TARA_MED_MAG_00080 | TARA | 0 | 0 | NA | 1 | 0 | orphan | 0 | 0 | NA | 0 | 0 | NA | 79.73 | 0.8 | 1.77 |
| TARA_PSW_MAG_00055 | TARA | 0 | 0 | NA | 1 | 0 | orphan | 0 | 0 | NA | 0 | 0 | NA | 78.74 | 0.8 | 1.66 |
| TARA_ANE_MAG_00042 | TARA | 0 | 0 | NA | 1 | 0 | orphan | 0 | 0 | NA | 0 | 0 | NA | 76 | 0.8 | 1.9 |
| TARA_ANE_MAG_00073 | TARA | 0 | 0 | NA | 1 | 0 | orphan | 0 | 0 | NA | 0 | 0 | NA | 75.94 | 0.8 | 1.47 |

| | | | | | | | | | | | | | | | | |
|---|---|---|---|---|---|---|---|---|---|---|---|---|---|---|---|---|
| TARA_PSE_MAG_00070 | TARA | 0 | 0 | NA | 1 | 0 | orphan | 0 | 0 | NA | 0 | 0 | NA | 64.94 | 0.8 | 1.14 |
| TARA_PON_MAG_00007 | TARA | 0 | 0 | NA | 1 | 0 | orphan | 0 | 0 | NA | 0 | 0 | NA | 92.74 | 0.81 | 1.5 |
| TARA_RED_MAG_00092 | TARA | 0 | 0 | NA | 1 | 0 | orphan | 0 | 0 | NA | 0 | 1 | orphan | 86.97 | 0.85 | 2.07 |
| TARA_ANW_MAG_00028 | TARA | 0 | 0 | NA | 1 | 0 | orphan | 0 | 0 | NA | 0 | 0 | NA | 62.65 | 0.85 | 0.87 |
| TARA_PON_MAG_00017 | TARA | 0 | 0 | NA | 3 | 0 | orphan | 0 | 0 | NA | 0 | 0 | NA | 61.68 | 0.85 | 0.86 |
| TARA_ANW_MAG_00027 | TARA | 0 | 1 | NA | 2 | 0 | orphan | 0 | 0 | NA | 0 | 1 | orphan | 60.66 | 0.85 | 0.63 |
| TARA_MED_MAG_00109 | TARA | 0 | 0 | NA | 1 | 0 | orphan | 0 | 0 | NA | 0 | 2 | orphan | 70.37 | 0.86 | 0.91 |
| TARA_ION_MAG_00067 | TARA | 1 | 2 | complete | 0 | 0 | NA | 0 | 0 | NA | 0 | 0 | NA | 68.45 | 0.86 | 2.94 |
| TARA_ANW_MAG_00004 | TARA | 0 | 0 | NA | 2 | 0 | orphan | 0 | 0 | NA | 0 | 1 | orphan | 99.5 | 0.87 | 2.83 |
| TARA_SOC_MAG_00001 | TARA | 0 | 0 | NA | 3 | 0 | orphan | 0 | 0 | NA | 0 | 0 | NA | 95.88 | 0.87 | 2.32 |
| TARA_MED_MAG_00083 | TARA | 0 | 0 | NA | 3 | 0 | orphan | 0 | 0 | NA | 0 | 0 | NA | 94.38 | 0.87 | 1.57 |
| TARA_RED_MAG_00022 | TARA | 2 | 3 | complete | 4 | 0 | orphan | 0 | 0 | NA | 0 | 1 | orphan | 97.64 | 0.89 | 3.66 |
| TARA_PSW_MAG_00016 | TARA | 1 | 1 | complete | 1 | 0 | orphan | 2 | 1 | complete | 0 | 0 | NA | 97.32 | 0.89 | 3.02 |
| TARA_ASE_MAG_00002 | TARA | 0 | 0 | NA | 1 | 0 | orphan | 1 | 1 | complete | 0 | 3 | orphan | 93.3 | 0.89 | 3.67 |
| TARA_PON_MAG_00029 | TARA | 1 | 1 | complete | 1 | 0 | orphan | 0 | 0 | NA | 0 | 0 | NA | 84.93 | 0.91 | 3.02 |
| TARA_RED_MAG_00104 | TARA | 3 | 5 | complete | 1 | 0 | orphan | 1 | 1 | complete | 0 | 1 | orphan | 59.9 | 0.92 | 2.93 |
| TARA_PSW_MAG_00004 | TARA | 1 | 1 | complete | 5 | 0 | orphan | 3 | 1 | complete | 0 | 2 | orphan | 99.63 | 0.93 | 4.27 |
| TARA_PON_MAG_00023 | TARA | 4 | 5 | complete | 1 | 0 | orphan | 1 | 1 | complete | 0 | 3 | orphan | 98.58 | 0.93 | 4.29 |
| TARA_ION_MAG_00033 | TARA | 0 | 0 | NA | 1 | 0 | orphan | 0 | 0 | NA | 0 | 0 | NA | 83.27 | 0.93 | 1.51 |
| TARA_MED_MAG_00072 | TARA | 0 | 0 | NA | 1 | 0 | orphan | 0 | 0 | NA | 0 | 0 | NA | 78.65 | 0.93 | 2 |

| | | | | | | | | | | | | | | | | |
|---|---|---|---|---|---|---|---|---|---|---|---|---|---|---|---|---|
| TARA_MED_MAG_00097 | TARA | 0 | 0 | NA | 1 | 0 | orphan | 0 | 0 | NA | 0 | 0 | NA | 63.97 | 0.93 | 0.98 |
| TARA_MED_MAG_00032 | TARA | 0 | 0 | NA | 1 | 0 | orphan | 0 | 0 | NA | 0 | 0 | NA | 83.63 | 0.96 | 1.25 |
| TARA_RED_MAG_00049 | TARA | 0 | 0 | NA | 3 | 0 | orphan | 0 | 0 | NA | 0 | 0 | NA | 74.56 | 0.96 | 1.57 |
| TARA_ION_MAG_00002 | TARA | 0 | 0 | NA | 6 | 1 | orphan | 1 | 1 | complete | 0 | 0 | NA | 99.76 | 0.97 | 3.23 |
| TARA_PSW_MAG_00071 | TARA | 0 | 0 | NA | 2 | 0 | orphan | 0 | 0 | NA | 0 | 0 | NA | 86.48 | 0.97 | 4.68 |
| TARA_RED_MAG_00014 | TARA | 0 | 0 | NA | 3 | 0 | orphan | 0 | 0 | NA | 0 | 0 | NA | 90.26 | 0.99 | 1.61 |
| TARA_ANE_MAG_00028 | TARA | 0 | 0 | NA | 1 | 0 | orphan | 0 | 0 | NA | 0 | 0 | NA | 89.74 | 0.99 | 1.24 |
| TARA_ANE_MAG_00001 | TARA | 0 | 0 | NA | 1 | 0 | orphan | 0 | 0 | NA | 0 | 0 | NA | 88.22 | 1.01 | 1.83 |
| TARA_RED_MAG_00011 | TARA | 0 | 0 | NA | 1 | 0 | orphan | 0 | 0 | NA | 0 | 0 | NA | 84.12 | 1.01 | 3.45 |
| TARA_PSE_MAG_00072 | TARA | 0 | 0 | NA | 4 | 0 | orphan | 0 | 0 | NA | 0 | 0 | NA | 68.08 | 1.02 | 2.63 |
| TARA_MED_MAG_00057 | TARA | 0 | 0 | NA | 1 | 0 | orphan | 0 | 0 | NA | 0 | 1 | orphan | 97.03 | 1.03 | 1.81 |
| TARA_MED_MAG_00011 | TARA | 0 | 0 | NA | 2 | 0 | orphan | 0 | 0 | NA | 0 | 0 | NA | 92.32 | 1.03 | 2.17 |
| TARA_ION_MAG_00001 | TARA | 1 | 1 | complete | 3 | 0 | orphan | 0 | 0 | NA | 0 | 0 | NA | 90.66 | 1.05 | 2.07 |
| TARA_PSW_MAG_00029 | TARA | 1 | 2 | complete | 5 | 0 | orphan | 0 | 0 | NA | 0 | 0 | NA | 98.13 | 1.08 | 3.03 |
| TARA_SOC_MAG_00012 | TARA | 0 | 0 | NA | 6 | 0 | orphan | 0 | 0 | NA | 0 | 0 | NA | 92.17 | 1.08 | 3.25 |
| TARA_ION_MAG_00006 | TARA | 0 | 0 | NA | 1 | 1 | orphan | 0 | 0 | NA | 0 | 0 | NA | 86.56 | 1.08 | 1.4 |
| TARA_IOS_MAG_00056 | TARA | 0 | 0 | NA | 1 | 0 | orphan | 0 | 0 | NA | 0 | 0 | NA | 61.52 | 1.08 | 1.34 |
| TARA_PSW_MAG_00046 | TARA | 0 | 0 | NA | 2 | 0 | orphan | 0 | 0 | NA | 0 | 0 | NA | 98.56 | 1.09 | 4.42 |
| TARA_RED_MAG_00019 | TARA | 0 | 0 | NA | 2 | 0 | orphan | 0 | 0 | NA | 0 | 0 | NA | 84.55 | 1.09 | 1.85 |
| TARA_RED_MAG_00002 | TARA | 0 | 0 | NA | 1 | 0 | orphan | 0 | 0 | NA | 0 | 0 | NA | 96.7 | 1.1 | 2.18 |

| | | | | | | | | | | | | | | | | |
|---|---|---|---|---|---|---|---|---|---|---|---|---|---|---|---|---|
| TARA_MED_MAG_00016 | TARA | 0 | 0 | NA | 1 | 0 | orphan | 0 | 0 | NA | 0 | 0 | NA | 96.15 | 1.1 | 1.64 |
| TARA_MED_MAG_00020 | TARA | 0 | 0 | NA | 1 | 0 | orphan | 0 | 0 | NA | 0 | 0 | NA | 94.51 | 1.1 | 1.58 |
| TARA_PSW_MAG_00030 | TARA | 0 | 0 | NA | 2 | 0 | orphan | 0 | 0 | NA | 0 | 0 | NA | 93.23 | 1.1 | 4.53 |
| TARA_IOS_MAG_00017 | TARA | 0 | 0 | NA | 5 | 0 | orphan | 0 | 0 | NA | 0 | 1 | orphan | 91.57 | 1.1 | 2.26 |
| TARA_RED_MAG_00042 | TARA | 0 | 0 | NA | 2 | 0 | orphan | 0 | 0 | NA | 0 | 0 | NA | 87.36 | 1.1 | 1.69 |
| TARA_MED_MAG_00050 | TARA | 0 | 0 | NA | 1 | 0 | orphan | 0 | 0 | NA | 0 | 0 | NA | 81.02 | 1.1 | 1.4 |
| TARA_IOS_MAG_00052 | TARA | 0 | 0 | NA | 1 | 0 | orphan | 0 | 0 | NA | 0 | 0 | NA | 80.44 | 1.1 | 1.76 |
| TARA_MED_MAG_00116 | TARA | 0 | 0 | NA | 2 | 0 | orphan | 0 | 0 | NA | 0 | 0 | NA | 58.04 | 1.1 | 0.8 |
| TARA_ANW_MAG_00067 | TARA | 1 | 0 | orphan | 2 | 0 | orphan | 0 | 0 | NA | 0 | 0 | NA | 51.25 | 1.1 | 2.24 |
| TARA_RED_MAG_00020 | TARA | 0 | 0 | NA | 2 | 0 | orphan | 0 | 0 | NA | 0 | 0 | NA | 98.33 | 1.11 | 5.28 |
| TARA_RED_MAG_00050 | TARA | 0 | 0 | NA | 2 | 0 | orphan | 0 | 0 | NA | 0 | 0 | NA | 84.1 | 1.11 | 5.4 |
| TARA_MED_MAG_00132 | TARA | 1 | 0 | orphan | 0 | 0 | NA | 0 | 0 | NA | 0 | 0 | NA | 66.48 | 1.12 | 2.06 |
| TARA_ANE_MAG_00050 | TARA | 0 | 0 | NA | 1 | 0 | orphan | 0 | 0 | NA | 0 | 0 | NA | 69.19 | 1.13 | 0.96 |
| TARA_PSW_MAG_00023 | TARA | 0 | 0 | NA | 4 | 0 | orphan | 0 | 0 | NA | 0 | 0 | NA | 97.73 | 1.14 | 3.01 |
| TARA_ANE_MAG_00016 | TARA | 0 | 0 | NA | 3 | 0 | orphan | 0 | 0 | NA | 0 | 0 | NA | 94.89 | 1.14 | 1.99 |
| TARA_ANW_MAG_00041 | TARA | 0 | 0 | NA | 2 | 0 | orphan | 0 | 0 | NA | 0 | 0 | NA | 67.86 | 1.14 | 1.68 |
| TARA_PSW_MAG_00020 | TARA | 1 | 1 | complete | 4 | 1 | complete | 0 | 0 | NA | 0 | 1 | orphan | 98.59 | 1.15 | 6.61 |
| TARA_ION_MAG_00007 | TARA | 0 | 0 | NA | 2 | 0 | orphan | 0 | 0 | NA | 0 | 0 | NA | 97.7 | 1.15 | 3.41 |
| TARA_MED_MAG_00039 | TARA | 0 | 0 | NA | 1 | 0 | orphan | 0 | 0 | NA | 0 | 0 | NA | 90.11 | 1.16 | 1.37 |
| TARA_ASE_MAG_00006 | TARA | 2 | 4 | complete | 0 | 0 | NA | 0 | 0 | NA | 0 | 1 | orphan | 76.46 | 1.16 | 2.98 |

| | | | | | | | | | | | | | | | | |
|---|---|---|---|---|---|---|---|---|---|---|---|---|---|---|---|---|
| TARA_PSW_MAG_00070 | TARA | 0 | 0 | NA | 1 | 0 | orphan | 0 | 0 | NA | 0 | 0 | NA | 69.08 | 1.16 | 2.55 |
| TARA_PSE_MAG_00093 | TARA | 0 | 0 | NA | 1 | 0 | orphan | 0 | 0 | NA | 0 | 0 | NA | 84.43 | 1.17 | 1.62 |
| TARA_PSW_MAG_00082 | TARA | 4 | 5 | complete | 3 | 0 | orphan | 1 | 2 | complete | 0 | 2 | orphan | 70.67 | 1.17 | 3.17 |
| TARA_MED_MAG_00055 | TARA | 0 | 1 | NA | 2 | 0 | orphan | 0 | 0 | NA | 0 | 0 | NA | 84.29 | 1.18 | 1.38 |
| TARA_ANW_MAG_00002 | TARA | 2 | 2 | complete | 1 | 0 | orphan | 0 | 1 | NA | 0 | 0 | NA | 98.37 | 1.19 | 3.03 |
| TARA_PSW_MAG_00107 | TARA | 2 | 2 | complete | 4 | 1 | orphan | 0 | 0 | NA | 0 | 3 | orphan | 68.41 | 1.19 | 4.55 |
| TARA_ASE_MAG_00010 | TARA | 0 | 0 | NA | 1 | 0 | orphan | 0 | 0 | NA | 0 | 0 | NA | 81.16 | 1.2 | 3.18 |
| TARA_PSE_MAG_00089 | TARA | 0 | 0 | NA | 1 | 0 | orphan | 0 | 0 | NA | 0 | 0 | NA | 79.8 | 1.2 | 2.03 |
| TARA_MED_MAG_00107 | TARA | 0 | 0 | NA | 3 | 0 | orphan | 0 | 0 | NA | 0 | 0 | NA | 79.01 | 1.2 | 0.62 |
| TARA_MED_MAG_00081 | TARA | 0 | 0 | NA | 1 | 0 | orphan | 0 | 0 | NA | 0 | 0 | NA | 73.49 | 1.2 | 0.73 |
| TARA_RED_MAG_00105 | TARA | 0 | 0 | NA | 1 | 0 | orphan | 0 | 0 | NA | 0 | 0 | NA | 69.75 | 1.2 | 2.07 |
| TARA_ANW_MAG_00005 | TARA | 3 | 3 | complete | 3 | 0 | orphan | 0 | 0 | NA | 0 | 3 | orphan | 97.64 | 1.21 | 2.34 |
| TARA_IOS_MAG_00044 | TARA | 0 | 0 | NA | 2 | 0 | orphan | 0 | 0 | NA | 0 | 0 | NA | 77.17 | 1.21 | 2.27 |
| TARA_PSE_MAG_00062 | TARA | 0 | 0 | NA | 1 | 0 | orphan | 0 | 0 | NA | 0 | 0 | NA | 67.69 | 1.22 | 1.59 |
| TARA_ION_MAG_00005 | TARA | 3 | 3 | complete | 3 | 0 | orphan | 0 | 0 | NA | 0 | 0 | NA | 97.33 | 1.24 | 4.16 |
| TARA_MED_MAG_00125 | TARA | 0 | 0 | NA | 1 | 0 | orphan | 0 | 0 | NA | 0 | 0 | NA | 62.73 | 1.24 | 1.21 |
| TARA_ANW_MAG_00059 | TARA | 0 | 0 | NA | 1 | 0 | orphan | 0 | 0 | NA | 0 | 0 | NA | 58.5 | 1.24 | 3.33 |
| TARA_ANW_MAG_00007 | TARA | 0 | 0 | NA | 2 | 0 | orphan | 0 | 0 | NA | 0 | 0 | NA | 95.81 | 1.25 | 4.42 |
| TARA_SOC_MAG_00002 | TARA | 1 | 1 | complete | 1 | 0 | orphan | 0 | 0 | NA | 0 | 1 | orphan | 97.19 | 1.26 | 4.43 |
| TARA_ION_MAG_00032 | TARA | 3 | 4 | complete | 3 | 0 | orphan | 0 | 1 | NA | 0 | 0 | NA | 92.98 | 1.26 | 3.68 |

| | | | | | | | | | | | | | | | |
|---|---|---|---|---|---|---|---|---|---|---|---|---|---|---|---|
| TARA_ION_MAG_00015 | TARA | 0 | 0 | NA | 3 | 0 | orphan | 2 | 1 | complete | 0 | 0 | NA | 98.04 | 1.28 | 2.98 |
| TARA_ION_MAG_00058 | TARA | 1 | 1 | complete | 3 | 0 | orphan | 0 | 0 | NA | 0 | 2 | orphan | 93.03 | 1.28 | 5.12 |
| TARA_ANE_MAG_00017 | TARA | 0 | 0 | NA | 1 | 0 | orphan | 0 | 0 | NA | 0 | 0 | NA | 90.36 | 1.28 | 0.85 |
| TARA_PSW_MAG_00093 | TARA | 0 | 0 | NA | 1 | 0 | orphan | 0 | 0 | NA | 0 | 0 | NA | 84.47 | 1.29 | 3.26 |
| TARA_IOS_MAG_00004 | TARA | 0 | 0 | NA | 1 | 0 | orphan | 0 | 0 | NA | 0 | 0 | NA | 97.39 | 1.3 | 2.98 |
| TARA_ANE_MAG_00058 | TARA | 0 | 0 | NA | 1 | 0 | orphan | 0 | 0 | NA | 0 | 0 | NA | 73.84 | 1.3 | 1.51 |
| TARA_PON_MAG_00053 | TARA | 0 | 0 | NA | 1 | 0 | orphan | 0 | 0 | NA | 0 | 0 | NA | 63.16 | 1.3 | 2.64 |
| TARA_RED_MAG_00021 | TARA | 0 | 0 | NA | 1 | 0 | orphan | 0 | 0 | NA | 0 | 1 | orphan | 95.59 | 1.31 | 1.19 |
| TARA_ANW_MAG_00022 | TARA | 0 | 0 | NA | 2 | 0 | orphan | 0 | 0 | NA | 0 | 0 | NA | 85.56 | 1.31 | 5.42 |
| TARA_PON_MAG_00038 | TARA | 0 | 1 | NA | 1 | 0 | orphan | 0 | 0 | NA | 0 | 0 | NA | 78.3 | 1.32 | 1.53 |
| TARA_PSE_MAG_00055 | TARA | 0 | 0 | NA | 1 | 0 | orphan | 2 | 0 | orphan | 0 | 0 | NA | 77.94 | 1.32 | 1.96 |
| TARA_RED_MAG_00078 | TARA | 0 | 0 | NA | 3 | 0 | orphan | 0 | 0 | NA | 0 | 0 | NA | 70.81 | 1.32 | 4.31 |
| TARA_PSE_MAG_00083 | TARA | 0 | 0 | NA | 6 | 0 | orphan | 1 | 1 | complete | 0 | 2 | orphan | 95.76 | 1.33 | 3.85 |
| TARA_PSE_MAG_00032 | TARA | 0 | 0 | NA | 1 | 0 | orphan | 0 | 0 | NA | 0 | 0 | NA | 87.65 | 1.34 | 2.85 |
| TARA_MED_MAG_00127 | TARA | 0 | 0 | NA | 1 | 0 | orphan | 0 | 0 | NA | 0 | 0 | NA | 84.67 | 1.34 | 2.36 |
| TARA_MED_MAG_00014 | TARA | 1 | 0 | orphan | 1 | 0 | orphan | 0 | 0 | NA | 0 | 0 | NA | 97.3 | 1.35 | 4.39 |
| TARA_RED_MAG_00070 | TARA | 0 | 0 | NA | 1 | 0 | orphan | 0 | 0 | NA | 0 | 0 | NA | 71.53 | 1.35 | 1.58 |
| TARA_PON_MAG_00050 | TARA | 0 | 0 | NA | 1 | 0 | orphan | 0 | 0 | NA | 0 | 0 | NA | 71.01 | 1.35 | 2.09 |
| TARA_PON_MAG_00011 | TARA | 0 | 0 | NA | 1 | 0 | orphan | 0 | 0 | NA | 0 | 0 | NA | 97.62 | 1.36 | 3.39 |
| TARA_MED_MAG_00041 | TARA | 0 | 0 | NA | 1 | 0 | orphan | 0 | 0 | NA | 0 | 0 | NA | 91.84 | 1.36 | 3.62 |

| | | | | | | | | | | | | | | | | |
|---|---|---|---|---|---|---|---|---|---|---|---|---|---|---|---|---|
| TARA_PON_MAG_00066 | TARA | 0 | 0 | NA | 2 | 0 | orphan | 0 | 0 | NA | 0 | 0 | NA | 52.64 | 1.36 | 3.2 |
| TARA_ANW_MAG_00011 | TARA | 0 | 0 | NA | 1 | 0 | orphan | 0 | 0 | NA | 0 | 0 | NA | 97.05 | 1.37 | 4.12 |
| TARA_PSE_MAG_00100 | TARA | 0 | 0 | NA | 3 | 0 | orphan | 0 | 0 | NA | 0 | 0 | NA | 50.02 | 1.38 | 2.69 |
| TARA_PSW_MAG_00067 | TARA | 7 | 7 | complete | 2 | 0 | orphan | 0 | 0 | NA | 0 | 0 | NA | 76.29 | 1.42 | 3.37 |
| TARA_PON_MAG_00027 | TARA | 4 | 3 | complete | 8 | 1 | orphan | 2 | 1 | complete | 0 | 0 | NA | 75.08 | 1.42 | 3.16 |
| TARA_PSW_MAG_00086 | TARA | 0 | 0 | NA | 1 | 0 | orphan | 0 | 0 | NA | 0 | 0 | NA | 70 | 1.42 | 1.91 |
| TARA_ANE_MAG_00076 | TARA | 0 | 0 | NA | 1 | 0 | orphan | 0 | 0 | NA | 0 | 0 | NA | 67.74 | 1.44 | 1.18 |
| TARA_ANE_MAG_00008 | TARA | 2 | 3 | complete | 4 | 0 | orphan | 0 | 0 | NA | 0 | 0 | NA | 87.71 | 1.45 | 1.71 |
| TARA_PSE_MAG_00106 | TARA | 0 | 0 | NA | 3 | 0 | orphan | 0 | 0 | NA | 0 | 1 | orphan | 87.13 | 1.46 | 3.41 |
| TARA_PSE_MAG_00103 | TARA | 1 | 1 | orphan | 4 | 0 | orphan | 1 | 1 | complete | 0 | 1 | orphan | 69.04 | 1.46 | 3.24 |
| TARA_ANW_MAG_00055 | TARA | 0 | 0 | NA | 0 | 0 | NA | 0 | 1 | NA | 0 | 0 | NA | 69.55 | 1.49 | 1.12 |
| TARA_PSW_MAG_00013 | TARA | 1 | 0 | orphan | 6 | 0 | orphan | 0 | 0 | NA | 0 | 0 | NA | 98.13 | 1.5 | 3.67 |
| TARA_MED_MAG_00017 | TARA | 0 | 0 | NA | 4 | 0 | orphan | 0 | 0 | NA | 0 | 0 | NA | 88.66 | 1.52 | 2.87 |
| TARA_RED_MAG_00029 | TARA | 0 | 0 | NA | 1 | 0 | orphan | 0 | 0 | NA | 0 | 0 | NA | 80.14 | 1.52 | 1.97 |
| TARA_PSE_MAG_00014 | TARA | 1 | 1 | complete | 1 | 0 | orphan | 0 | 0 | NA | 0 | 0 | NA | 98.51 | 1.55 | 6.12 |
| TARA_ANE_MAG_00009 | TARA | 1 | 0 | orphan | 6 | 0 | orphan | 0 | 0 | NA | 0 | 3 | orphan | 96.26 | 1.56 | 5.23 |
| TARA_PSE_MAG_00111 | TARA | 3 | 3 | complete | 6 | 0 | orphan | 0 | 0 | NA | 0 | 1 | orphan | 89.75 | 1.58 | 2.97 |
| TARA_MED_MAG_00092 | TARA | 0 | 0 | NA | 1 | 0 | orphan | 0 | 0 | NA | 0 | 0 | NA | 60.42 | 1.58 | 1.41 |
| TARA_ANW_MAG_00009 | TARA | 1 | 1 | complete | 2 | 0 | orphan | 0 | 0 | NA | 0 | 0 | NA | 98.97 | 1.59 | 5.05 |
| TARA_IOS_MAG_00023 | TARA | 1 | 1 | complete | 5 | 0 | orphan | 0 | 0 | NA | 0 | 1 | orphan | 98.32 | 1.59 | 4.14 |

| | | | | | | | | | | | | | | | | |
|---|---|---|---|---|---|---|---|---|---|---|---|---|---|---|---|---|
| TARA_ASW_MAG_00012 | TARA | 0 | 0 | NA | 4 | 0 | orphan | 0 | 0 | NA | 0 | 0 | NA | 99.01 | 1.6 | 4.45 |
| TARA_ANE_MAG_00019 | TARA | 0 | 0 | NA | 3 | 0 | orphan | 0 | 0 | NA | 0 | 0 | NA | 84.53 | 1.6 | 1.85 |
| TARA_RED_MAG_00072 | TARA | 0 | 0 | NA | 1 | 0 | orphan | 0 | 0 | NA | 0 | 0 | NA | 71.73 | 1.6 | 1.64 |
| TARA_RED_MAG_00090 | TARA | 0 | 0 | NA | 1 | 0 | orphan | 0 | 0 | NA | 0 | 0 | NA | 66.63 | 1.6 | 1.63 |
| TARA_PSE_MAG_00010 | TARA | 2 | 2 | complete | 4 | 0 | orphan | 0 | 0 | NA | 0 | 1 | orphan | 98.73 | 1.64 | 4.59 |
| TARA_ION_MAG_00014 | TARA | 2 | 2 | complete | 1 | 1 | orphan | 1 | 1 | complete | 0 | 1 | orphan | 99.35 | 1.67 | 4.47 |
| TARA_PON_MAG_00061 | TARA | 8 | 8 | complete | 4 | 0 | orphan | 2 | 1 | complete | 0 | 3 | orphan | 96.27 | 1.68 | 4.48 |
| TARA_PSW_MAG_00024 | TARA | 2 | 1 | complete | 1 | 0 | orphan | 0 | 0 | NA | 0 | 0 | NA | 90.57 | 1.68 | 1.57 |
| TARA_RED_MAG_00009 | TARA | 0 | 1 | NA | 0 | 0 | NA | 0 | 0 | NA | 0 | 0 | NA | 82.77 | 1.68 | 0.93 |
| TARA_IOS_MAG_00040 | TARA | 0 | 0 | NA | 2 | 1 | complete | 0 | 0 | NA | 0 | 0 | NA | 60.15 | 1.68 | 1.15 |
| TARA_RED_MAG_00074 | TARA | 0 | 1 | NA | 1 | 0 | orphan | 0 | 0 | NA | 0 | 0 | NA | 67.33 | 1.69 | 1.99 |
| TARA_PSE_MAG_00058 | TARA | 0 | 0 | NA | 1 | 0 | orphan | 0 | 0 | NA | 0 | 0 | NA | 95.65 | 1.7 | 2.52 |
| TARA_ANW_MAG_00018 | TARA | 0 | 0 | NA | 2 | 0 | orphan | 0 | 0 | NA | 0 | 0 | NA | 92.3 | 1.71 | 2.3 |
| TARA_ANW_MAG_00054 | TARA | 0 | 0 | NA | 1 | 0 | orphan | 0 | 0 | NA | 0 | 0 | NA | 76.84 | 1.71 | 1.42 |
| TARA_IOS_MAG_00027 | TARA | 0 | 0 | NA | 1 | 0 | orphan | 0 | 0 | NA | 0 | 0 | NA | 74.74 | 1.71 | 2.16 |
| TARA_IOS_MAG_00067 | TARA | 1 | 1 | complete | 0 | 0 | NA | 0 | 0 | NA | 0 | 1 | orphan | 83.3 | 1.72 | 2.93 |
| TARA_SOC_MAG_00007 | TARA | 0 | 1 | NA | 0 | 0 | NA | 0 | 0 | NA | 0 | 0 | NA | 78.26 | 1.73 | 2.92 |
| TARA_MED_MAG_00034 | TARA | 1 | 0 | orphan | 1 | 0 | orphan | 0 | 0 | NA | 0 | 0 | NA | 89.36 | 1.74 | 1.96 |
| TARA_MED_MAG_00064 | TARA | 0 | 0 | NA | 1 | 0 | orphan | 0 | 0 | NA | 0 | 0 | NA | 75.55 | 1.74 | 1.11 |
| TARA_ASW_MAG_00010 | TARA | 0 | 0 | NA | 1 | 0 | orphan | 0 | 0 | NA | 0 | 0 | NA | 92.59 | 1.75 | 2.24 |

| | | | | | | | | | | | | | | | | |
|---|---|---|---|---|---|---|---|---|---|---|---|---|---|---|---|---|
| TARA_PSE_MAG_00090 | TARA | 0 | 0 | NA | 1 | 0 | orphan | 0 | 0 | NA | 0 | 0 | NA | 69.53 | 1.75 | 1.22 |
| TARA_RED_MAG_00094 | TARA | 0 | 0 | NA | 1 | 0 | orphan | 0 | 0 | NA | 0 | 0 | NA | 61.92 | 1.75 | 2.52 |
| TARA_IOS_MAG_00066 | TARA | 0 | 0 | NA | 2 | 0 | orphan | 0 | 1 | NA | 0 | 0 | NA | 54.16 | 1.75 | 3.09 |
| TARA_ANE_MAG_00007 | TARA | 1 | 1 | orphan | 3 | 0 | orphan | 0 | 0 | NA | 0 | 0 | NA | 88.3 | 1.77 | 2.45 |
| TARA_ANW_MAG_00020 | TARA | 0 | 0 | NA | 3 | 0 | orphan | 0 | 0 | NA | 0 | 0 | NA | 97.09 | 1.78 | 3.99 |
| TARA_ION_MAG_00029 | TARA | 0 | 0 | NA | 1 | 1 | orphan | 0 | 0 | NA | 0 | 0 | NA | 81.89 | 1.78 | 2.68 |
| TARA_PSW_MAG_00117 | TARA | 0 | 0 | NA | 1 | 0 | orphan | 0 | 0 | NA | 0 | 0 | NA | 50.23 | 1.78 | 2.24 |
| TARA_PSW_MAG_00114 | TARA | 1 | 1 | orphan | 0 | 0 | NA | 0 | 0 | NA | 0 | 0 | NA | 92.78 | 1.79 | 2.59 |
| TARA_ANE_MAG_00031 | TARA | 0 | 1 | NA | 3 | 1 | complete | 0 | 0 | NA | 0 | 1 | orphan | 74.84 | 1.81 | 2.16 |
| TARA_MED_MAG_00111 | TARA | 0 | 0 | NA | 1 | 0 | orphan | 0 | 0 | NA | 0 | 0 | NA | 59.27 | 1.81 | 0.85 |
| TARA_MED_MAG_00060 | TARA | 0 | 0 | NA | 1 | 0 | orphan | 0 | 0 | NA | 0 | 0 | NA | 77.14 | 1.82 | 1.62 |
| TARA_ASW_MAG_00017 | TARA | 0 | 0 | NA | 5 | 0 | orphan | 0 | 0 | NA | 0 | 0 | NA | 71.69 | 1.87 | 1.83 |
| TARA_ANW_MAG_00057 | TARA | 0 | 0 | NA | 2 | 0 | orphan | 0 | 0 | NA | 0 | 0 | NA | 92.98 | 1.88 | 1.98 |
| TARA_PSW_MAG_00017 | TARA | 0 | 1 | NA | 1 | 0 | orphan | 0 | 0 | NA | 0 | 1 | orphan | 98.25 | 1.9 | 5.96 |
| TARA_ANE_MAG_00040 | TARA | 0 | 0 | NA | 1 | 0 | orphan | 0 | 0 | NA | 0 | 0 | NA | 92.83 | 1.9 | 1.64 |
| TARA_ASW_MAG_00004 | TARA | 4 | 4 | complete | 0 | 0 | NA | 0 | 0 | NA | 0 | 1 | orphan | 92.28 | 1.91 | 3.2 |
| TARA_PSE_MAG_00080 | TARA | 1 | 1 | complete | 1 | 0 | orphan | 0 | 0 | NA | 0 | 0 | NA | 68.81 | 1.91 | 1.16 |
| TARA_RED_MAG_00081 | TARA | 0 | 0 | NA | 2 | 0 | orphan | 0 | 0 | NA | 0 | 0 | NA | 80.74 | 1.92 | 4.16 |
| TARA_ANE_MAG_00064 | TARA | 0 | 0 | NA | 1 | 0 | orphan | 0 | 0 | NA | 0 | 0 | NA | 59.62 | 1.92 | 1.23 |
| TARA_MED_MAG_00098 | TARA | 0 | 0 | NA | 0 | 1 | NA | 0 | 0 | NA | 0 | 0 | NA | 74.75 | 1.93 | 1.29 |

| | | | | | | | | | | | | | | | | |
|---|---|---|---|---|---|---|---|---|---|---|---|---|---|---|---|---|
| TARA_PSW_MAG_00026 | TARA | 0 | 0 | NA | 1 | 0 | orphan | 0 | 0 | NA | 0 | 0 | NA | 98.28 | 1.94 | 6 |
| TARA_ANW_MAG_00035 | TARA | 0 | 0 | NA | 1 | 0 | orphan | 0 | 0 | NA | 0 | 0 | NA | 84.83 | 1.95 | 2.66 |
| TARA_MED_MAG_00106 | TARA | 0 | 0 | NA | 1 | 0 | orphan | 0 | 0 | NA | 0 | 0 | NA | 69.09 | 1.95 | 1.59 |
| TARA_ION_MAG_00049 | TARA | 0 | 0 | NA | 1 | 0 | orphan | 0 | 0 | NA | 0 | 0 | NA | 62.39 | 1.95 | 1.26 |
| TARA_RED_MAG_00006 | TARA | 0 | 0 | NA | 1 | 0 | orphan | 0 | 0 | NA | 0 | 0 | NA | 97.22 | 1.96 | 3.75 |
| TARA_ANW_MAG_00019 | TARA | 5 | 4 | complete | 0 | 0 | NA | 1 | 2 | complete | 0 | 0 | NA | 91.3 | 1.96 | 3.99 |
| TARA_PSW_MAG_00064 | TARA | 0 | 0 | NA | 2 | 0 | orphan | 0 | 0 | NA | 0 | 0 | NA | 74.1 | 1.97 | 3.02 |
| TARA_PSE_MAG_00119 | TARA | 0 | 0 | NA | 1 | 0 | orphan | 0 | 0 | NA | 0 | 1 | orphan | 51.47 | 1.98 | 2.06 |
| TARA_PSW_MAG_00040 | TARA | 0 | 0 | NA | 1 | 0 | orphan | 0 | 0 | NA | 0 | 0 | NA | 96.42 | 1.99 | 3.1 |
| TARA_PSW_MAG_00062 | TARA | 4 | 4 | complete | 3 | 0 | orphan | 0 | 3 | NA | 0 | 0 | NA | 79.61 | 1.99 | 3.47 |
| TARA_PSE_MAG_00052 | TARA | 0 | 0 | NA | 1 | 0 | orphan | 0 | 0 | NA | 0 | 0 | NA | 74.79 | 1.99 | 1.75 |
| TARA_RED_MAG_00102 | TARA | 0 | 0 | NA | 2 | 0 | orphan | 0 | 0 | NA | 0 | 0 | NA | 63.16 | 1.99 | 2.17 |
| TARA_PSE_MAG_00042 | TARA | 3 | 3 | complete | 3 | 0 | orphan | 2 | 2 | complete | 0 | 3 | orphan | 91.27 | 2 | 4.35 |
| TARA_RED_MAG_00045 | TARA | 0 | 0 | NA | 1 | 0 | orphan | 0 | 0 | NA | 0 | 0 | NA | 90.49 | 2 | 2.49 |
| TARA_ANW_MAG_00030 | TARA | 1 | 0 | orphan | 6 | 1 | complete | 0 | 0 | NA | 0 | 0 | NA | 90.01 | 2 | 1.87 |
| TARA_PSE_MAG_00109 | TARA | 0 | 1 | NA | 1 | 0 | orphan | 0 | 0 | NA | 0 | 0 | NA | 74.9 | 2.01 | 2.53 |
| TARA_MED_MAG_00058 | TARA | 2 | 2 | complete | 3 | 0 | orphan | 0 | 0 | NA | 0 | 2 | orphan | 83.88 | 2.02 | 3.39 |
| TARA_ASW_MAG_00016 | TARA | 0 | 0 | NA | 1 | 0 | orphan | 0 | 0 | NA | 0 | 0 | NA | 81.1 | 2.02 | 2.62 |
| TARA_PSE_MAG_00105 | TARA | 0 | 0 | NA | 1 | 0 | orphan | 0 | 0 | NA | 0 | 0 | NA | 55.8 | 2.03 | 2.34 |
| TARA_PSE_MAG_00044 | TARA | 0 | 0 | NA | 1 | 0 | orphan | 0 | 0 | NA | 0 | 0 | NA | 86.23 | 2.04 | 3.8 |

| | | | | | | | | | | | | | | | | |
|---|---|---|---|---|---|---|---|---|---|---|---|---|---|---|---|---|
| TARA_MED_MAG_00128 | TARA | 0 | 0 | NA | 0 | 0 | NA | 0 | 0 | NA | 0 | 1 | orphan | 72.1 | 2.04 | 2.43 |
| TARA_PON_MAG_00058 | TARA | 0 | 0 | NA | 1 | 0 | orphan | 0 | 0 | NA | 0 | 1 | orphan | 83.31 | 2.06 | 2.06 |
| TARA_PSW_MAG_00094 | TARA | 0 | 0 | NA | 2 | 0 | orphan | 0 | 0 | NA | 0 | 0 | NA | 76.6 | 2.06 | 1.77 |
| TARA_PON_MAG_00062 | TARA | 0 | 0 | NA | 1 | 0 | orphan | 1 | 1 | complete | 0 | 0 | NA | 65.38 | 2.07 | 3.64 |
| TARA_PSW_MAG_00022 | TARA | 0 | 0 | NA | 2 | 0 | orphan | 0 | 0 | NA | 0 | 0 | NA | 86.53 | 2.08 | 4.26 |
| TARA_PSW_MAG_00044 | TARA | 0 | 0 | NA | 3 | 0 | orphan | 0 | 0 | NA | 0 | 0 | NA | 88.4 | 2.09 | 1.15 |
| TARA_PSW_MAG_00065 | TARA | 1 | 1 | orphan | 4 | 0 | orphan | 0 | 0 | NA | 0 | 0 | NA | 91.86 | 2.1 | 3.07 |
| TARA_IOS_MAG_00055 | TARA | 0 | 0 | NA | 1 | 0 | orphan | 0 | 0 | NA | 0 | 0 | NA | 67.62 | 2.1 | 1.98 |
| TARA_PSW_MAG_00106 | TARA | 0 | 0 | NA | 2 | 0 | orphan | 0 | 0 | NA | 0 | 0 | NA | 71.87 | 2.12 | 2.25 |
| TARA_PON_MAG_00042 | TARA | 1 | 1 | orphan | 2 | 0 | orphan | 0 | 0 | NA | 0 | 0 | NA | 94.27 | 2.13 | 2.28 |
| TARA_PSW_MAG_00049 | TARA | 0 | 0 | NA | 1 | 0 | orphan | 0 | 0 | NA | 0 | 0 | NA | 91.62 | 2.14 | 1.82 |
| TARA_PSW_MAG_00066 | TARA | 0 | 0 | NA | 1 | 0 | orphan | 0 | 0 | NA | 0 | 0 | NA | 84.19 | 2.14 | 2.03 |
| TARA_PSW_MAG_00077 | TARA | 0 | 0 | NA | 2 | 0 | orphan | 0 | 0 | NA | 0 | 0 | NA | 80.04 | 2.16 | 1.57 |
| TARA_PSE_MAG_00019 | TARA | 1 | 0 | orphan | 4 | 0 | orphan | 0 | 0 | NA | 0 | 0 | NA | 98.26 | 2.17 | 2.47 |
| TARA_RED_MAG_00037 | TARA | 0 | 0 | NA | 4 | 0 | orphan | 0 | 0 | NA | 0 | 0 | NA | 89.71 | 2.2 | 2.02 |
| TARA_MED_MAG_00054 | TARA | 0 | 0 | NA | 1 | 0 | orphan | 0 | 0 | NA | 0 | 0 | NA | 84.62 | 2.2 | 1.23 |
| TARA_PSE_MAG_00088 | TARA | 0 | 0 | NA | 1 | 0 | orphan | 0 | 0 | NA | 0 | 0 | NA | 62.2 | 2.2 | 1.02 |
| TARA_RED_MAG_00067 | TARA | 1 | 0 | orphan | 0 | 0 | NA | 0 | 0 | NA | 0 | 1 | orphan | 56.61 | 2.2 | 1.89 |
| TARA_RED_MAG_00043 | TARA | 0 | 0 | NA | 1 | 0 | orphan | 0 | 0 | NA | 0 | 0 | NA | 80.22 | 2.21 | 1.63 |
| TARA_RED_MAG_00036 | TARA | 0 | 0 | NA | 2 | 0 | orphan | 0 | 0 | NA | 0 | 0 | NA | 92.22 | 2.22 | 5.79 |

| | | | | | | | | | | | | | | | | |
|---|---|---|---|---|---|---|---|---|---|---|---|---|---|---|---|---|
| TARA_MED_MAG_00029 | TARA | 0 | 0 | NA | 1 | 0 | orphan | 0 | 0 | NA | 0 | 0 | NA | 97.44 | 2.23 | 2.3 |
| TARA_ANW_MAG_00040 | TARA | 0 | 0 | NA | 5 | 0 | orphan | 2 | 1 | complete | 0 | 0 | NA | 86.02 | 2.23 | 3.17 |
| TARA_PSE_MAG_00071 | TARA | 0 | 0 | NA | 2 | 0 | orphan | 0 | 0 | NA | 0 | 0 | NA | 60.32 | 2.23 | 1.59 |
| TARA_ANE_MAG_00032 | TARA | 0 | 0 | NA | 2 | 0 | orphan | 0 | 0 | NA | 0 | 0 | NA | 85.5 | 2.24 | 1.81 |
| TARA_IOS_MAG_00057 | TARA | 0 | 0 | NA | 2 | 0 | orphan | 0 | 0 | NA | 0 | 0 | NA | 76.72 | 2.26 | 3.01 |
| TARA_PSE_MAG_00028 | TARA | 0 | 0 | NA | 2 | 0 | orphan | 0 | 0 | NA | 0 | 0 | NA | 97.73 | 2.27 | 2.54 |
| TARA_ANW_MAG_00014 | TARA | 0 | 0 | NA | 2 | 0 | orphan | 0 | 0 | NA | 0 | 0 | NA | 92.98 | 2.27 | 2.77 |
| TARA_PSW_MAG_00113 | TARA | 0 | 0 | NA | 2 | 0 | orphan | 0 | 0 | NA | 0 | 0 | NA | 84.21 | 2.27 | 2.44 |
| TARA_PSE_MAG_00061 | TARA | 0 | 0 | NA | 1 | 0 | orphan | 0 | 0 | NA | 0 | 0 | NA | 75.12 | 2.27 | 1.64 |
| TARA_MED_MAG_00136 | TARA | 0 | 0 | NA | 1 | 0 | orphan | 0 | 0 | NA | 0 | 0 | NA | 79.9 | 2.28 | 2.05 |
| TARA_IOS_MAG_00059 | TARA | 1 | 0 | orphan | 2 | 0 | orphan | 0 | 0 | NA | 0 | 0 | NA | 61.54 | 2.3 | 2.32 |
| TARA_RED_MAG_00064 | TARA | 0 | 1 | NA | 0 | 0 | NA | 0 | 0 | NA | 0 | 0 | NA | 87.95 | 2.31 | 1.44 |
| TARA_PSE_MAG_00095 | TARA | 0 | 0 | NA | 2 | 0 | orphan | 0 | 0 | NA | 0 | 0 | NA | 82.01 | 2.34 | 2.78 |
| TARA_IOS_MAG_00034 | TARA | 0 | 0 | NA | 2 | 0 | orphan | 0 | 0 | NA | 0 | 0 | NA | 77.47 | 2.34 | 2.27 |
| TARA_PSE_MAG_00128 | TARA | 0 | 0 | NA | 1 | 0 | orphan | 0 | 0 | NA | 0 | 0 | NA | 61.48 | 2.34 | 2.56 |
| TARA_RED_MAG_00093 | TARA | 0 | 0 | NA | 2 | 0 | orphan | 0 | 0 | NA | 0 | 0 | NA | 91.49 | 2.35 | 6.14 |
| TARA_ANW_MAG_00049 | TARA | 0 | 0 | NA | 2 | 0 | orphan | 0 | 0 | NA | 0 | 0 | NA | 65.99 | 2.35 | 1.4 |
| TARA_RED_MAG_00096 | TARA | 0 | 0 | NA | 4 | 1 | complete | 0 | 0 | NA | 0 | 1 | orphan | 55.88 | 2.36 | 2.06 |
| TARA_PSE_MAG_00101 | TARA | 1 | 1 | complete | 3 | 0 | orphan | 0 | 1 | NA | 0 | 0 | NA | 80.57 | 2.37 | 2.56 |
| TARA_PSE_MAG_00126 | TARA | 0 | 0 | NA | 2 | 0 | orphan | 0 | 1 | NA | 0 | 0 | NA | 59.6 | 2.37 | 2.06 |

| | | | | | | | | | | | | | | | | |
|---|---|---|---|---|---|---|---|---|---|---|---|---|---|---|---|---|
| TARA_PSE_MAG_00077 | TARA | 0 | 0 | NA | 2 | 0 | orphan | 0 | 0 | NA | 0 | 1 | orphan | 98.96 | 2.4 | 2.77 |
| TARA_ANW_MAG_00038 | TARA | 0 | 0 | NA | 2 | 0 | orphan | 0 | 0 | NA | 0 | 0 | NA | 72.89 | 2.41 | 0.75 |
| TARA_ASW_MAG_00008 | TARA | 0 | 0 | NA | 2 | 0 | orphan | 0 | 0 | NA | 0 | 1 | orphan | 95.65 | 2.42 | 3.57 |
| TARA_ANE_MAG_00037 | TARA | 2 | 3 | complete | 1 | 0 | orphan | 0 | 1 | NA | 0 | 1 | orphan | 93.58 | 2.42 | 3.27 |
| TARA_ION_MAG_00057 | TARA | 0 | 0 | NA | 2 | 0 | orphan | 0 | 0 | NA | 0 | 0 | NA | 71.2 | 2.44 | 1.55 |
| TARA_PSE_MAG_00082 | TARA | 0 | 0 | NA | 2 | 0 | orphan | 0 | 0 | NA | 0 | 0 | NA | 67.17 | 2.44 | 2.67 |
| TARA_PSE_MAG_00118 | TARA | 0 | 0 | NA | 2 | 0 | orphan | 0 | 0 | NA | 0 | 0 | NA | 65.58 | 2.48 | 3.65 |
| TARA_PSE_MAG_00092 | TARA | 0 | 0 | NA | 1 | 0 | orphan | 0 | 0 | NA | 0 | 0 | NA | 63.25 | 2.48 | 0.62 |
| TARA_ANE_MAG_00069 | TARA | 0 | 0 | NA | 2 | 0 | orphan | 0 | 0 | NA | 0 | 0 | NA | 84.34 | 2.5 | 2.09 |
| TARA_MED_MAG_00103 | TARA | 0 | 0 | NA | 1 | 0 | orphan | 0 | 0 | NA | 0 | 0 | NA | 79.69 | 2.55 | 1.72 |
| TARA_ANW_MAG_00032 | TARA | 0 | 0 | NA | 2 | 0 | orphan | 0 | 0 | NA | 0 | 0 | NA | 78.07 | 2.55 | 2.01 |
| TARA_PSE_MAG_00097 | TARA | 2 | 3 | complete | 1 | 0 | orphan | 2 | 0 | orphan | 0 | 0 | NA | 64.45 | 2.55 | 3.89 |
| TARA_PON_MAG_00009 | TARA | 0 | 0 | NA | 1 | 0 | orphan | 0 | 0 | NA | 0 | 0 | NA | 91.03 | 2.56 | 1.93 |
| TARA_ANW_MAG_00010 | TARA | 2 | 1 | complete | 3 | 0 | orphan | 0 | 0 | NA | 0 | 3 | orphan | 87.72 | 2.56 | 2.73 |
| TARA_MED_MAG_00131 | TARA | 0 | 0 | NA | 1 | 0 | orphan | 0 | 0 | NA | 0 | 0 | NA | 86.4 | 2.56 | 2.6 |
| TARA_IOS_MAG_00033 | TARA | 0 | 0 | NA | 2 | 0 | orphan | 0 | 0 | NA | 0 | 0 | NA | 84.79 | 2.56 | 2 |
| TARA_RED_MAG_00083 | TARA | 0 | 0 | NA | 1 | 0 | orphan | 0 | 0 | NA | 0 | 0 | NA | 68.35 | 2.56 | 1.2 |
| TARA_RED_MAG_00032 | TARA | 0 | 0 | NA | 4 | 0 | orphan | 0 | 0 | NA | 0 | 1 | orphan | 81.77 | 2.58 | 4.2 |
| TARA_MED_MAG_00137 | TARA | 0 | 0 | NA | 1 | 0 | orphan | 0 | 0 | NA | 0 | 0 | NA | 63.09 | 2.59 | 3.11 |
| TARA_RED_MAG_00026 | TARA | 0 | 0 | NA | 1 | 0 | orphan | 0 | 0 | NA | 0 | 0 | NA | 93.7 | 2.62 | 3.59 |

| | | | | | | | | | | | | | | | |
|---|---|---|---|---|---|---|---|---|---|---|---|---|---|---|---|
| TARA_PSW_MAG_00028 | TARA | 1 | 1 | complete | 5 | 1 | complete | 0 | 1 | NA | 0 | 0 | NA | 99.2 | 2.67 | 3.97 |
| TARA_PSW_MAG_00032 | TARA | 1 | 1 | complete | 3 | 0 | orphan | 0 | 0 | NA | 0 | 1 | orphan | 98.07 | 2.69 | 4.51 |
| TARA_ION_MAG_00025 | TARA | 1 | 1 | complete | 3 | 0 | orphan | 0 | 0 | NA | 0 | 2 | orphan | 93.65 | 2.69 | 6.27 |
| TARA_ANW_MAG_00052 | TARA | 0 | 0 | NA | 1 | 0 | orphan | 0 | 0 | NA | 0 | 0 | NA | 77.82 | 2.71 | 1.36 |
| TARA_IOS_MAG_00005 | TARA | 1 | 0 | orphan | 1 | 0 | orphan | 0 | 0 | NA | 0 | 0 | NA | 97.97 | 2.73 | 3.24 |
| TARA_PSE_MAG_00065 | TARA | 0 | 0 | NA | 5 | 0 | orphan | 0 | 0 | NA | 0 | 0 | NA | 69.79 | 2.74 | 4.51 |
| TARA_ION_MAG_00038 | TARA | 0 | 0 | NA | 1 | 0 | orphan | 0 | 0 | NA | 0 | 0 | NA | 82.42 | 2.75 | 1.22 |
| TARA_PSW_MAG_00085 | TARA | 0 | 0 | NA | 1 | 0 | orphan | 0 | 0 | NA | 0 | 0 | NA | 68.4 | 2.75 | 1.17 |
| TARA_PON_MAG_00065 | TARA | 2 | 0 | orphan | 2 | 0 | orphan | 0 | 0 | NA | 0 | 0 | NA | 51.4 | 2.75 | 3.62 |
| TARA_MED_MAG_00091 | TARA | 0 | 0 | NA | 1 | 0 | orphan | 0 | 0 | NA | 0 | 0 | NA | 62.63 | 2.77 | 1.93 |
| TARA_PSE_MAG_00035 | TARA | 0 | 1 | NA | 0 | 0 | NA | 0 | 0 | NA | 0 | 0 | NA | 90.14 | 2.79 | 3.38 |
| TARA_IOS_MAG_00019 | TARA | 0 | 0 | NA | 1 | 0 | orphan | 0 | 0 | NA | 0 | 0 | NA | 85.38 | 2.79 | 2.18 |
| TARA_ANW_MAG_00012 | TARA | 0 | 0 | NA | 1 | 0 | orphan | 0 | 0 | NA | 0 | 0 | NA | 87.91 | 2.85 | 2.19 |
| TARA_ANE_MAG_00030 | TARA | 0 | 0 | NA | 2 | 0 | orphan | 0 | 0 | NA | 0 | 0 | NA | 80.4 | 2.86 | 1.65 |
| TARA_PSE_MAG_00121 | TARA | 2 | 0 | orphan | 3 | 0 | orphan | 1 | 0 | orphan | 0 | 0 | NA | 55.62 | 2.9 | 2.36 |
| TARA_RED_MAG_00101 | TARA | 0 | 0 | NA | 1 | 0 | orphan | 0 | 0 | NA | 0 | 0 | NA | 56.28 | 2.91 | 2.5 |
| TARA_MED_MAG_00023 | TARA | 0 | 0 | NA | 1 | 0 | orphan | 0 | 0 | NA | 0 | 0 | NA | 95.73 | 2.99 | 1.93 |
| TARA_MED_MAG_00024 | TARA | 0 | 0 | NA | 1 | 0 | orphan | 0 | 0 | NA | 0 | 0 | NA | 91.45 | 2.99 | 1.87 |
| TARA_PSW_MAG_00090 | TARA | 0 | 0 | NA | 1 | 1 | orphan | 0 | 0 | NA | 0 | 0 | NA | 83.76 | 2.99 | 2.08 |
| TARA_PSE_MAG_00027 | TARA | 0 | 0 | NA | 1 | 0 | orphan | 0 | 0 | NA | 0 | 0 | NA | 95.39 | 3.03 | 3.57 |

| | | | | | | | | | | | | | | | | |
|---|---|---|---|---|---|---|---|---|---|---|---|---|---|---|---|---|
| TARA_RED_MAG_00091 | TARA | 0 | 0 | NA | 1 | 0 | orphan | 0 | 0 | NA | 0 | 0 | NA | 78.37 | 3.04 | 2.28 |
| TARA_PSE_MAG_00046 | TARA | 0 | 0 | NA | 1 | 0 | orphan | 0 | 0 | NA | 0 | 0 | NA | 94.58 | 3.07 | 1.92 |
| TARA_ANW_MAG_00068 | TARA | 1 | 0 | orphan | 0 | 0 | NA | 0 | 0 | NA | 0 | 0 | NA | 80.9 | 3.12 | 2.73 |
| TARA_IOS_MAG_00049 | TARA | 0 | 0 | NA | 1 | 0 | orphan | 0 | 0 | NA | 0 | 0 | NA | 70.75 | 3.16 | 1.94 |
| TARA_ANE_MAG_00038 | TARA | 0 | 0 | NA | 1 | 0 | orphan | 0 | 0 | NA | 0 | 0 | NA | 85.36 | 3.2 | 3.56 |
| TARA_IOS_MAG_00031 | TARA | 1 | 0 | orphan | 2 | 0 | orphan | 0 | 0 | NA | 0 | 0 | NA | 77.83 | 3.23 | 1.47 |
| TARA_ANW_MAG_00031 | TARA | 0 | 0 | NA | 1 | 0 | orphan | 0 | 0 | NA | 0 | 0 | NA | 75.12 | 3.23 | 2.16 |
| TARA_RED_MAG_00086 | TARA | 0 | 0 | NA | 2 | 0 | orphan | 0 | 0 | NA | 0 | 0 | NA | 89.06 | 3.36 | 1.27 |
| TARA_PSW_MAG_00053 | TARA | 0 | 0 | NA | 4 | 0 | orphan | 0 | 0 | NA | 0 | 0 | NA | 92.15 | 3.4 | 5.65 |
| TARA_RED_MAG_00106 | TARA | 0 | 0 | NA | 1 | 0 | orphan | 0 | 0 | NA | 0 | 0 | NA | 72.88 | 3.4 | 2.44 |
| TARA_MED_MAG_00074 | TARA | 1 | 1 | complete | 2 | 0 | orphan | 0 | 0 | NA | 0 | 1 | orphan | 88.95 | 3.41 | 2.45 |
| TARA_IOS_MAG_00014 | TARA | 0 | 0 | NA | 1 | 0 | orphan | 0 | 0 | NA | 0 | 0 | NA | 95.95 | 3.44 | 3.61 |
| TARA_ION_MAG_00016 | TARA | 1 | 1 | complete | 1 | 0 | orphan | 0 | 0 | NA | 0 | 0 | NA | 90.53 | 3.48 | 2.57 |
| TARA_PSE_MAG_00022 | TARA | 5 | 5 | complete | 6 | 0 | orphan | 5 | 4 | complete | 0 | 3 | orphan | 94.3 | 3.51 | 3.89 |
| TARA_ION_MAG_00064 | TARA | 0 | 0 | NA | 2 | 0 | orphan | 0 | 0 | NA | 0 | 0 | NA | 63.92 | 3.57 | 2.13 |
| TARA_PSW_MAG_00098 | TARA | 0 | 0 | NA | 1 | 0 | orphan | 0 | 0 | NA | 0 | 0 | NA | 64.51 | 3.61 | 0.72 |
| TARA_PON_MAG_00047 | TARA | 0 | 0 | NA | 1 | 0 | orphan | 0 | 0 | NA | 0 | 0 | NA | 63.56 | 3.66 | 1.22 |
| TARA_RED_MAG_00030 | TARA | 0 | 0 | NA | 2 | 0 | orphan | 0 | 0 | NA | 0 | 0 | NA | 90.6 | 3.69 | 4.02 |
| TARA_ION_MAG_00041 | TARA | 0 | 0 | NA | 1 | 0 | orphan | 0 | 0 | NA | 0 | 0 | NA | 80.96 | 3.72 | 2.02 |
| TARA_PSW_MAG_00083 | TARA | 0 | 0 | NA | 2 | 0 | orphan | 0 | 0 | NA | 0 | 0 | NA | 70.17 | 3.72 | 2.53 |

| | | | | | | | | | | | | | | | | |
|---|---|---|---|---|---|---|---|---|---|---|---|---|---|---|---|---|
| TARA_PSW_MAG_00115 | TARA | 0 | 0 | NA | 2 | 0 | orphan | 0 | 0 | NA | 0 | 0 | NA | 66.48 | 3.77 | 2.69 |
| TARA_ANE_MAG_00044 | TARA | 0 | 0 | NA | 2 | 1 | complete | 0 | 0 | NA | 0 | 0 | NA | 83.74 | 3.85 | 1.11 |
| TARA_PSW_MAG_00019 | TARA | 0 | 0 | NA | 2 | 0 | orphan | 0 | 0 | NA | 0 | 0 | NA | 88.8 | 3.88 | 2.26 |
| TARA_ASE_MAG_00011 | TARA | 0 | 0 | NA | 3 | 0 | orphan | 0 | 0 | NA | 0 | 0 | NA | 61.07 | 4.04 | 1.47 |
| TARA_PSW_MAG_00058 | TARA | 3 | 3 | complete | 4 | 0 | orphan | 0 | 0 | NA | 0 | 1 | orphan | 81.42 | 4.05 | 4.64 |
| TARA_PSE_MAG_00114 | TARA | 5 | 6 | complete | 2 | 0 | orphan | 0 | 0 | NA | 0 | 4 | orphan | 56.66 | 4.07 | 2.26 |
| TARA_ANE_MAG_00035 | TARA | 0 | 0 | NA | 3 | 0 | orphan | 0 | 0 | NA | 0 | 0 | NA | 83.33 | 4.22 | 0.87 |
| TARA_PON_MAG_00015 | TARA | 0 | 0 | NA | 2 | 0 | orphan | 0 | 0 | NA | 0 | 0 | NA | 86.4 | 4.24 | 4.8 |
| TARA_PON_MAG_00064 | TARA | 0 | 0 | NA | 1 | 0 | orphan | 0 | 0 | NA | 0 | 0 | NA | 59.66 | 4.27 | 2.17 |
| TARA_ASE_MAG_00004 | TARA | 0 | 0 | NA | 1 | 0 | orphan | 0 | 0 | NA | 0 | 0 | NA | 92.31 | 4.32 | 2.39 |
| TARA_ASE_MAG_00014 | TARA | 0 | 0 | NA | 1 | 0 | orphan | 0 | 0 | NA | 0 | 0 | NA | 79.88 | 4.32 | 1.58 |
| TARA_RED_MAG_00060 | TARA | 0 | 0 | NA | 1 | 0 | orphan | 0 | 0 | NA | 0 | 1 | orphan | 72.41 | 4.34 | 3.81 |
| TARA_PSW_MAG_00087 | TARA | 11 | 10 | complete | 6 | 0 | orphan | 3 | 3 | complete | 0 | 3 | orphan | 82.24 | 4.39 | 3.85 |
| TARA_RED_MAG_00084 | TARA | 1 | 0 | orphan | 2 | 0 | orphan | 0 | 0 | NA | 0 | 2 | orphan | 65.93 | 4.4 | 5.14 |
| TARA_IOS_MAG_00053 | TARA | 0 | 0 | NA | 1 | 0 | orphan | 0 | 0 | NA | 0 | 0 | NA | 55.3 | 4.4 | 0.7 |
| TARA_ANW_MAG_00056 | TARA | 0 | 0 | NA | 2 | 0 | orphan | 0 | 0 | NA | 0 | 0 | NA | 84.58 | 4.42 | 2.84 |
| TARA_ANW_MAG_00006 | TARA | 1 | 1 | complete | 2 | 1 | complete | 0 | 0 | NA | 0 | 0 | NA | 98.7 | 4.43 | 5.49 |
| TARA_ANW_MAG_00036 | TARA | 2 | 3 | complete | 2 | 0 | orphan | 1 | 1 | complete | 0 | 1 | orphan | 94.48 | 4.49 | 3.39 |
| TARA_PSW_MAG_00100 | TARA | 0 | 0 | NA | 1 | 0 | orphan | 0 | 0 | NA | 0 | 0 | NA | 79.12 | 4.61 | 1.38 |
| TARA_PSE_MAG_00102 | TARA | 0 | 0 | NA | 2 | 0 | orphan | 0 | 0 | NA | 0 | 0 | NA | 55.58 | 4.62 | 2.71 |

| | | | | | | | | | | | | | | | | |
|---|---|---|---|---|---|---|---|---|---|---|---|---|---|---|---|---|
| TARA_PON_MAG_00039 | TARA | 0 | 0 | NA | 1 | 0 | orphan | 0 | 0 | NA | 0 | 0 | NA | 77.84 | 4.67 | 2.28 |
| TARA_PSE_MAG_00039 | TARA | 2 | 2 | complete | 11 | 0 | orphan | 0 | 0 | NA | 0 | 0 | NA | 85.34 | 4.73 | 8.21 |
| TARA_PSE_MAG_00015 | TARA | 1 | 1 | complete | 0 | 0 | NA | 0 | 0 | NA | 0 | 0 | NA | 92.9 | 4.8 | 5.17 |
| TARA_PON_MAG_00006 | TARA | 4 | 3 | complete | 6 | 0 | orphan | 0 | 0 | NA | 0 | 0 | NA | 93.1 | 4.92 | 8.7 |
| TARA_MED_MAG_00135 | TARA | 0 | 0 | NA | 1 | 0 | orphan | 0 | 0 | NA | 0 | 0 | NA | 67.24 | 4.95 | 2.04 |
| TARA_RED_MAG_00103 | TARA | 2 | 1 | complete | 7 | 0 | orphan | 0 | 0 | NA | 0 | 0 | NA | 87.9 | 5.01 | 2.98 |
| TARA_MED_MAG_00129 | TARA | 0 | 0 | NA | 4 | 0 | orphan | 0 | 0 | NA | 0 | 0 | NA | 65.83 | 5.05 | 2.4 |
| TARA_RED_MAG_00057 | TARA | 0 | 0 | NA | 1 | 0 | orphan | 0 | 0 | NA | 0 | 0 | NA | 95.17 | 5.41 | 5.34 |
| TARA_PSW_MAG_00084 | TARA | 0 | 0 | NA | 3 | 0 | orphan | 0 | 0 | NA | 0 | 0 | NA | 86.81 | 5.43 | 2.97 |
| TARA_RED_MAG_00024 | TARA | 0 | 0 | NA | 3 | 0 | orphan | 0 | 0 | NA | 0 | 0 | NA | 92.9 | 5.51 | 4.06 |
| TARA_ANE_MAG_00046 | TARA | 0 | 0 | NA | 0 | 0 | NA | 0 | 0 | NA | 0 | 1 | orphan | 73.09 | 5.54 | 1.9 |
| TARA_ASE_MAG_00016 | TARA | 0 | 0 | NA | 1 | 0 | orphan | 0 | 0 | NA | 0 | 0 | NA | 70.56 | 5.78 | 1.35 |
| TARA_ANW_MAG_00065 | TARA | 1 | 0 | orphan | 2 | 0 | orphan | 0 | 0 | NA | 0 | 0 | NA | 88.61 | 5.97 | 5.06 |
| TARA_ASW_MAG_00024 | TARA | 0 | 0 | NA | 1 | 0 | orphan | 0 | 0 | NA | 0 | 0 | NA | 64.13 | 6 | 1.09 |
| TARA_RED_MAG_00008 | TARA | 0 | 0 | NA | 1 | 0 | orphan | 0 | 0 | NA | 0 | 0 | NA | 94.82 | 6.03 | 3.02 |
| TARA_PSE_MAG_00075 | TARA | 0 | 0 | NA | 3 | 0 | orphan | 0 | 0 | NA | 0 | 0 | NA | 55.51 | 6.27 | 0.9 |
| TARA_ASE_MAG_00003 | TARA | 2 | 0 | orphan | 0 | 0 | NA | 0 | 0 | NA | 0 | 0 | NA | 87.46 | 6.56 | 4.24 |
| TARA_RED_MAG_00040 | TARA | 0 | 0 | NA | 1 | 0 | orphan | 0 | 0 | NA | 0 | 0 | NA | 88 | 6.59 | 2.11 |
| TARA_PSE_MAG_00037 | TARA | 2 | 3 | complete | 8 | 0 | orphan | 0 | 0 | NA | 0 | 0 | NA | 94.09 | 6.63 | 10.72 |
| TARA_RED_MAG_00048 | TARA | 0 | 0 | NA | 2 | 0 | orphan | 0 | 0 | NA | 0 | 0 | NA | 93.59 | 6.84 | 2.8 |

| | | | | | | | | | | | | | | | | |
|---|---|---|---|---|---|---|---|---|---|---|---|---|---|---|---|---|
| TARA_ANW_MAG_00026 | TARA | 0 | 0 | NA | 2 | 0 | orphan | 0 | 0 | NA | 0 | 0 | NA | 93.01 | 7.14 | 1.6 |
| TARA_PON_MAG_00013 | TARA | 1 | 1 | complete | 7 | 0 | orphan | 1 | 0 | orphan | 0 | 0 | NA | 96.97 | 7.17 | 10.15 |
| TARA_ANW_MAG_00042 | TARA | 0 | 0 | NA | 1 | 0 | orphan | 0 | 0 | NA | 0 | 0 | NA | 84.24 | 7.29 | 1.26 |
| TARA_PSW_MAG_00112 | TARA | 0 | 0 | NA | 5 | 0 | orphan | 1 | 0 | orphan | 0 | 2 | orphan | 59.93 | 7.56 | 3.08 |
| TARA_ANW_MAG_00062 | TARA | 0 | 0 | NA | 1 | 0 | orphan | 0 | 0 | NA | 0 | 0 | NA | 66.22 | 7.83 | 2.16 |
| TARA_MED_MAG_00078 | TARA | 0 | 0 | NA | 2 | 0 | orphan | 0 | 0 | NA | 0 | 0 | NA | 83.38 | 8.12 | 1.72 |
| TARA_PON_MAG_00054 | TARA | 0 | 0 | NA | 0 | 0 | NA | 0 | 0 | NA | 0 | 1 | orphan | 62.02 | 8.12 | 1.65 |
| TARA_PSE_MAG_00125 | TARA | 0 | 0 | NA | 1 | 0 | orphan | 0 | 0 | NA | 0 | 0 | NA | 71.82 | 8.14 | 2.53 |
| TARA_RED_MAG_00035 | TARA | 0 | 0 | NA | 1 | 0 | orphan | 0 | 0 | NA | 0 | 0 | NA | 89.01 | 8.79 | 1.18 |
| TARA_MED_MAG_00104 | TARA | 0 | 0 | NA | 0 | 0 | NA | 1 | 0 | orphan | 0 | 0 | NA | 73.49 | 9.64 | 0.74 |
| SHIPPO_MAG_177 | SHIPPO | 0 | 0 | NA | 0 | 0 | NA | 1 | 0 | orphan | 0 | 0 | NA | 98.39 | 0 | 2.52 |
| SHIPPO_MAG_27 | SHIPPO | 0 | 1 | NA | 2 | 0 | orphan | 0 | 0 | NA | 0 | 0 | NA | 93.2 | 0 | 2.57 |
| SHIPPO_MAG_179 | SHIPPO | 0 | 2 | NA | 0 | 0 | NA | 0 | 0 | NA | 0 | 0 | NA | 92.74 | 0 | 1.94 |
| SHIPPO_MAG_64 | SHIPPO | 0 | 0 | NA | 1 | 0 | orphan | 0 | 0 | NA | 0 | 0 | NA | 90.36 | 0 | 1.15 |
| SHIPPO_MAG_128 | SHIPPO | 0 | 0 | NA | 1 | 0 | orphan | 0 | 0 | NA | 0 | 0 | NA | 84.42 | 0 | 2.44 |
| SHIPPO_MAG_132 | SHIPPO | 0 | 0 | NA | 1 | 0 | orphan | 0 | 0 | NA | 0 | 0 | NA | 82.7 | 0 | 1.04 |
| SHIPPO_MAG_102 | SHIPPO | 0 | 0 | NA | 1 | 0 | orphan | 0 | 0 | NA | 0 | 0 | NA | 82.4 | 0 | 1.93 |
| SHIPPO_MAG_105 | SHIPPO | 0 | 0 | NA | 1 | 0 | orphan | 0 | 0 | NA | 0 | 0 | NA | 77.73 | 0 | 1.8 |
| SHIPPO_MAG_282 | SHIPPO | 0 | 0 | NA | 2 | 0 | orphan | 0 | 0 | NA | 0 | 1 | orphan | 74.14 | 0 | 2.47 |
| SHIPPO_MAG_287 | SHIPPO | 0 | 0 | NA | 2 | 0 | orphan | 0 | 0 | NA | 0 | 0 | NA | 68.45 | 0 | 3.06 |

| | | | | | | | | | | | | | | | | |
|---|---|---|---|---|---|---|---|---|---|---|---|---|---|---|---|---|
| SHIPPO_MAG_238 | SHIPPO | 0 | 0 | NA | 1 | 0 | orphan | 0 | 0 | NA | 0 | 0 | NA | 65.52 | 0 | 1.42 |
| SHIPPO_MAG_203 | SHIPPO | 0 | 0 | NA | 1 | 0 | orphan | 0 | 0 | NA | 0 | 0 | NA | 59.77 | 0 | 1.6 |
| SHIPPO_MAG_289 | SHIPPO | 0 | 0 | NA | 1 | 0 | orphan | 0 | 0 | NA | 0 | 0 | NA | 59.48 | 0 | 1.2 |
| SHIPPO_MAG_248 | SHIPPO | 0 | 0 | NA | 1 | 0 | orphan | 0 | 0 | NA | 0 | 0 | NA | 53.57 | 0 | 0.55 |
| SHIPPO_MAG_135 | SHIPPO | 1 | 0 | orphan | 0 | 0 | NA | 0 | 0 | NA | 0 | 0 | NA | 79.98 | 0.02 | 1.73 |
| SHIPPO_MAG_8 | SHIPPO | 0 | 0 | NA | 1 | 0 | orphan | 0 | 0 | NA | 0 | 0 | NA | 92.73 | 0.06 | 3.34 |
| SHIPPO_MAG_180 | SHIPPO | 0 | 0 | NA | 1 | 0 | orphan | 0 | 0 | NA | 0 | 0 | NA | 100 | 0.11 | 2.77 |
| SHIPPO_MAG_101 | SHIPPO | 0 | 0 | NA | 1 | 0 | orphan | 0 | 0 | NA | 0 | 0 | NA | 78.96 | 0.12 | 1.98 |
| SHIPPO_MAG_77 | SHIPPO | 0 | 0 | NA | 1 | 0 | orphan | 0 | 0 | NA | 0 | 0 | NA | 95.27 | 0.2 | 1.24 |
| SHIPPO_MAG_245 | SHIPPO | 0 | 0 | NA | 2 | 0 | orphan | 0 | 0 | NA | 0 | 0 | NA | 58.55 | 0.24 | 0.58 |
| SHIPPO_MAG_104 | SHIPPO | 0 | 0 | NA | 1 | 0 | orphan | 0 | 0 | NA | 0 | 0 | NA | 83.73 | 0.4 | 1.94 |
| SHIPPO_MAG_181 | SHIPPO | 0 | 1 | NA | 0 | 0 | NA | 0 | 0 | NA | 0 | 1 | orphan | 94.51 | 0.55 | 2.78 |
| SHIPPO_MAG_2 | SHIPPO | 0 | 0 | NA | 1 | 0 | orphan | 0 | 0 | NA | 0 | 0 | NA | 90.07 | 0.62 | 2.2 |
| SHIPPO_MAG_17 | SHIPPO | 0 | 0 | NA | 1 | 0 | orphan | 0 | 0 | NA | 0 | 0 | NA | 89.05 | 0.73 | 2.08 |
| SHIPPO_MAG_136 | SHIPPO | 0 | 1 | NA | 0 | 0 | NA | 0 | 0 | NA | 0 | 0 | NA | 95.55 | 0.74 | 1.77 |
| SHIPPO_MAG_161 | SHIPPO | 0 | 0 | NA | 1 | 0 | orphan | 0 | 0 | NA | 0 | 0 | NA | 92.77 | 0.8 | 1.21 |
| SHIPPO_MAG_184 | SHIPPO | 0 | 0 | NA | 2 | 0 | orphan | 0 | 0 | NA | 0 | 0 | NA | 72.49 | 0.8 | 1.73 |
| SHIPPO_MAG_187 | SHIPPO | 0 | 0 | NA | 1 | 0 | orphan | 0 | 0 | NA | 0 | 0 | NA | 68.13 | 0.8 | 1.38 |
| SHIPPO_MAG_185 | SHIPPO | 0 | 0 | NA | 3 | 0 | orphan | 0 | 0 | NA | 0 | 0 | NA | 53.12 | 0.8 | 1.29 |
| SHIPPO_MAG_98 | SHIPPO | 0 | 0 | NA | 1 | 0 | orphan | 0 | 0 | NA | 0 | 0 | NA | 84.53 | 0.84 | 1.95 |

| | | | | | | | | | | | | | | | | |
|---|---|---|---|---|---|---|---|---|---|---|---|---|---|---|---|---|
| SHIPPO_MAG_73 | SHIPPO | 0 | 0 | NA | 1 | 0 | orphan | 0 | 0 | NA | 0 | 0 | NA | 85.86 | 0.93 | 2.57 |
| SHIPPO_MAG_188 | SHIPPO | 0 | 0 | NA | 1 | 0 | orphan | 0 | 0 | NA | 0 | 0 | NA | 57.89 | 1 | 1.22 |
| SHIPPO_MAG_94 | SHIPPO | 0 | 0 | NA | 4 | 0 | orphan | 0 | 0 | NA | 0 | 1 | orphan | 77.49 | 1.1 | 2.62 |
| SHIPPO_MAG_95 | SHIPPO | 0 | 0 | NA | 1 | 0 | orphan | 0 | 0 | NA | 0 | 0 | NA | 95.65 | 1.16 | 2.6 |
| SHIPPO_MAG_152 | SHIPPO | 0 | 0 | NA | 4 | 1 | complete | 0 | 0 | NA | 0 | 0 | NA | 85.97 | 1.33 | 1.84 |
| SHIPPO_MAG_52 | SHIPPO | 0 | 0 | NA | 1 | 0 | orphan | 0 | 0 | NA | 0 | 0 | NA | 91.3 | 1.34 | 3.04 |
| SHIPPO_MAG_82 | SHIPPO | 0 | 0 | NA | 2 | 0 | orphan | 0 | 0 | NA | 0 | 0 | NA | 89.58 | 1.34 | 2.08 |
| SHIPPO_MAG_176 | SHIPPO | 0 | 0 | NA | 2 | 0 | orphan | 0 | 0 | NA | 0 | 0 | NA | 89.99 | 1.4 | 2.77 |
| SHIPPO_MAG_5 | SHIPPO | 0 | 0 | NA | 1 | 0 | orphan | 0 | 0 | NA | 0 | 0 | NA | 89.59 | 1.41 | 1.31 |
| SHIPPO_MAG_22 | SHIPPO | 0 | 0 | NA | 1 | 0 | orphan | 0 | 0 | NA | 0 | 0 | NA | 89.7 | 1.5 | 2.27 |
| SHIPPO_MAG_244 | SHIPPO | 0 | 0 | NA | 1 | 0 | orphan | 0 | 0 | NA | 0 | 0 | NA | 68.46 | 1.52 | 1.54 |
| SHIPPO_MAG_281 | SHIPPO | 0 | 0 | NA | 1 | 0 | orphan | 0 | 0 | NA | 0 | 0 | NA | 73.28 | 1.72 | 2.12 |
| SHIPPO_MAG_280 | SHIPPO | 0 | 0 | NA | 1 | 0 | orphan | 0 | 0 | NA | 0 | 0 | NA | 63.24 | 1.72 | 1.96 |
| SHIPPO_MAG_71 | SHIPPO | 0 | 0 | NA | 1 | 0 | orphan | 0 | 0 | NA | 0 | 0 | NA | 90.73 | 1.77 | 1.98 |
| SHIPPO_MAG_277 | SHIPPO | 0 | 1 | NA | 1 | 0 | orphan | 0 | 0 | NA | 0 | 0 | NA | 56.67 | 1.85 | 1.17 |
| SHIPPO_MAG_6 | SHIPPO | 0 | 0 | NA | 1 | 0 | orphan | 0 | 0 | NA | 0 | 0 | NA | 92.59 | 1.99 | 2.62 |
| SHIPPO_MAG_147 | SHIPPO | 1 | 1 | orphan | 0 | 0 | NA | 0 | 0 | NA | 0 | 0 | NA | 98.66 | 2 | 1.69 |
| SHIPPO_MAG_174 | SHIPPO | 0 | 0 | NA | 1 | 0 | orphan | 0 | 0 | NA | 0 | 0 | NA | 77.04 | 2.14 | 1.82 |
| SHIPPO_MAG_195 | SHIPPO | 0 | 0 | NA | 1 | 0 | orphan | 0 | 0 | NA | 0 | 0 | NA | 65.59 | 2.25 | 0.88 |
| SHIPPO_MAG_186 | SHIPPO | 0 | 0 | NA | 1 | 0 | orphan | 0 | 0 | NA | 0 | 0 | NA | 70.93 | 2.4 | 1.65 |

| | | | | | | | | | | | | | | | | |
|---|---|---|---|---|---|---|---|---|---|---|---|---|---|---|---|---|
| SHIPPO_MAG_13 | SHIPPO | 0 | 0 | NA | 1 | 0 | orphan | 0 | 0 | NA | 0 | 0 | NA | 93.02 | 2.42 | 2.23 |
| SHIPPO_MAG_198 | SHIPPO | 0 | 0 | NA | 1 | 0 | orphan | 0 | 0 | NA | 0 | 0 | NA | 74.23 | 2.54 | 1.59 |
| SHIPPO_MAG_140 | SHIPPO | 0 | 1 | NA | 0 | 0 | NA | 0 | 0 | NA | 0 | 0 | NA | 95.59 | 2.57 | 1.8 |
| SHIPPO_MAG_30 | SHIPPO | 1 | 0 | orphan | 4 | 0 | orphan | 0 | 0 | NA | 0 | 0 | NA | 81.98 | 2.59 | 2.95 |
| SHIPPO_MAG_194 | SHIPPO | 0 | 3 | NA | 1 | 0 | orphan | 0 | 0 | NA | 0 | 0 | NA | 74.52 | 2.59 | 1.99 |
| SHIPPO_MAG_294 | SHIPPO | 0 | 0 | NA | 2 | 0 | orphan | 0 | 0 | NA | 0 | 0 | NA | 55.8 | 2.59 | 1.6 |
| SHIPPO_MAG_242 | SHIPPO | 0 | 0 | NA | 5 | 0 | orphan | 0 | 0 | NA | 0 | 0 | NA | 74.92 | 2.65 | 2.07 |
| SHIPPO_MAG_103 | SHIPPO | 0 | 0 | NA | 1 | 0 | orphan | 0 | 0 | NA | 0 | 0 | NA | 81.33 | 2.92 | 1.88 |
| SHIPPO_MAG_21 | SHIPPO | 0 | 0 | NA | 1 | 0 | orphan | 0 | 0 | NA | 0 | 0 | NA | 89.73 | 2.98 | 2.68 |
| SHIPPO_MAG_62 | SHIPPO | 0 | 0 | NA | 1 | 0 | orphan | 0 | 0 | NA | 0 | 0 | NA | 97.44 | 2.99 | 3.05 |
| SHIPPO_MAG_233 | SHIPPO | 1 | 2 | orphan | 0 | 0 | NA | 0 | 0 | NA | 0 | 0 | NA | 67.78 | 3.15 | 1.37 |
| SHIPPO_MAG_127 | SHIPPO | 0 | 0 | NA | 1 | 0 | orphan | 0 | 0 | NA | 0 | 0 | NA | 80.41 | 3.16 | 2.59 |
| SHIPPO_MAG_249 | SHIPPO | 0 | 0 | NA | 1 | 0 | orphan | 0 | 0 | NA | 0 | 0 | NA | 72.12 | 3.28 | 1.4 |
| SHIPPO_MAG_49 | SHIPPO | 0 | 0 | NA | 1 | 0 | orphan | 0 | 0 | NA | 0 | 0 | NA | 84.45 | 3.3 | 3.09 |
| SHIPPO_MAG_288 | SHIPPO | 0 | 0 | NA | 1 | 0 | orphan | 0 | 0 | NA | 0 | 0 | NA | 74.17 | 3.31 | 1.17 |
| SHIPPO_MAG_227 | SHIPPO | 1 | 0 | orphan | 0 | 0 | NA | 0 | 0 | NA | 0 | 0 | NA | 65.47 | 3.31 | 1.24 |
| SHIPPO_MAG_241 | SHIPPO | 0 | 0 | NA | 1 | 0 | orphan | 0 | 0 | NA | 0 | 0 | NA | 63.59 | 3.39 | 1.3 |
| SHIPPO_MAG_283 | SHIPPO | 0 | 0 | NA | 1 | 0 | orphan | 0 | 0 | NA | 0 | 0 | NA | 62.93 | 3.45 | 2.11 |
| SHIPPO_MAG_204 | SHIPPO | 0 | 0 | NA | 1 | 0 | orphan | 0 | 0 | NA | 0 | 0 | NA | 53.45 | 3.45 | 1.67 |
| SHIPPO_MAG_272 | SHIPPO | 0 | 0 | NA | 2 | 0 | orphan | 0 | 0 | NA | 0 | 0 | NA | 53 | 3.57 | 1.17 |

| | | | | | | | | | | | | | | | | |
|---|---|---|---|---|---|---|---|---|---|---|---|---|---|---|---|---|
| SHIPPO_MAG_279 | SHIPPO | 0 | 0 | NA | 1 | 0 | orphan | 0 | 0 | NA | 0 | 0 | NA | 74.6 | 3.6 | 2.17 |
| SHIPPO_MAG_224 | SHIPPO | 1 | 0 | orphan | 0 | 0 | NA | 0 | 0 | NA | 0 | 0 | NA | 52.15 | 3.6 | 1.3 |
| SHIPPO_MAG_225 | SHIPPO | 0 | 0 | NA | 1 | 0 | orphan | 0 | 0 | NA | 0 | 0 | NA | 57.18 | 3.68 | 0.94 |
| SHIPPO_MAG_33 | SHIPPO | 0 | 0 | NA | 1 | 0 | orphan | 0 | 0 | NA | 0 | 0 | NA | 87.3 | 3.79 | 0.98 |
| SHIPPO_MAG_66 | SHIPPO | 0 | 0 | NA | 1 | 0 | orphan | 0 | 0 | NA | 0 | 0 | NA | 83.38 | 3.91 | 2 |
| SHIPPO_MAG_240 | SHIPPO | 0 | 0 | NA | 2 | 0 | orphan | 0 | 0 | NA | 0 | 0 | NA | 61.25 | 3.91 | 1.37 |
| SHIPPO_MAG_200 | SHIPPO | 0 | 0 | NA | 1 | 0 | orphan | 0 | 0 | NA | 0 | 0 | NA | 66.96 | 4.08 | 1.88 |
| SHIPPO_MAG_173 | SHIPPO | 0 | 0 | NA | 2 | 0 | orphan | 0 | 0 | NA | 0 | 0 | NA | 87.62 | 4.12 | 2.21 |
| SHIPPO_MAG_16 | SHIPPO | 0 | 0 | NA | 1 | 0 | orphan | 0 | 0 | NA | 0 | 0 | NA | 91.35 | 4.22 | 2.03 |
| SHIPPO_MAG_63 | SHIPPO | 0 | 3 | NA | 1 | 0 | orphan | 0 | 0 | NA | 0 | 0 | NA | 85.47 | 4.27 | 3.35 |
| SHIPPO_MAG_285 | SHIPPO | 1 | 0 | orphan | 0 | 0 | NA | 0 | 0 | NA | 0 | 0 | NA | 72.34 | 4.41 | 2.65 |
| SHIPPO_MAG_130 | SHIPPO | 0 | 0 | NA | 2 | 0 | orphan | 0 | 0 | NA | 0 | 0 | NA | 80.33 | 4.46 | 3.22 |
| SHIPPO_MAG_143 | SHIPPO | 0 | 0 | NA | 2 | 0 | orphan | 0 | 0 | NA | 0 | 0 | NA | 91.05 | 4.64 | 2.13 |
| SHIPPO_MAG_164 | SHIPPO | 0 | 0 | NA | 0 | 1 | NA | 0 | 0 | NA | 0 | 0 | NA | 84.32 | 4.71 | 2.11 |
| SHIPPO_MAG_254 | SHIPPO | 0 | 0 | NA | 7 | 0 | orphan | 0 | 0 | NA | 0 | 0 | NA | 60.25 | 4.79 | 1.97 |
| SHIPPO_MAG_68 | SHIPPO | 0 | 0 | NA | 1 | 0 | orphan | 0 | 0 | NA | 0 | 0 | NA | 82.89 | 5.15 | 2.11 |
| SHIPPO_MAG_284 | SHIPPO | 0 | 0 | NA | 1 | 0 | orphan | 0 | 0 | NA | 0 | 0 | NA | 60.34 | 5.17 | 2.18 |
| SHIPPO_MAG_23 | SHIPPO | 0 | 0 | NA | 2 | 0 | orphan | 0 | 0 | NA | 0 | 0 | NA | 78.13 | 5.27 | 3.26 |
| SHIPPO_MAG_84 | SHIPPO | 0 | 0 | NA | 1 | 0 | orphan | 0 | 0 | NA | 0 | 0 | NA | 81.87 | 5.33 | 2.27 |
| SHIPPO_MAG_278 | SHIPPO | 0 | 0 | NA | 2 | 0 | orphan | 0 | 0 | NA | 0 | 0 | NA | 66.33 | 6.28 | 1.86 |

| | | | | | | | | | | | | | | | | |
|---|---|---|---|---|---|---|---|---|---|---|---|---|---|---|---|---|
| SHIPPO_MAG_85 | SHIPPO | 0 | 0 | NA | 0 | 0 | NA | 0 | 1 | NA | 0 | 0 | NA | 76.99 | 6.44 | 2.01 |
| SHIPPO_MAG_262 | SHIPPO | 0 | 0 | NA | 0 | 1 | NA | 0 | 0 | NA | 0 | 0 | NA | 56.66 | 6.5 | 1.38 |
| SHIPPO_MAG_67 | SHIPPO | 0 | 0 | NA | 1 | 0 | orphan | 0 | 0 | NA | 0 | 0 | NA | 91.14 | 6.74 | 1.71 |
| SHIPPO_MAG_263 | SHIPPO | 0 | 0 | NA | 0 | 1 | NA | 0 | 0 | NA | 0 | 0 | NA | 52.34 | 6.91 | 0.96 |
| SHIPPO_MAG_246 | SHIPPO | 0 | 0 | NA | 1 | 0 | orphan | 0 | 0 | NA | 0 | 0 | NA | 53.83 | 6.93 | 1.33 |
| SHIPPO_MAG_202 | SHIPPO | 2 | 1 | complete | 3 | 0 | orphan | 0 | 1 | NA | 0 | 2 | orphan | 73.5 | 7.36 | 1.22 |
| SHIPPO_MAG_141 | SHIPPO | 0 | 0 | NA | 1 | 0 | orphan | 0 | 0 | NA | 0 | 0 | NA | 87.14 | 7.84 | 2.11 |
| SHIPPO_MAG_65 | SHIPPO | 0 | 0 | NA | 1 | 1 | orphan | 0 | 0 | NA | 0 | 0 | NA | 81.68 | 8.34 | 1.57 |
| SHIPPO_MAG_210 | SHIPPO | 0 | 1 | NA | 5 | 0 | orphan | 0 | 1 | NA | 0 | 2 | orphan | 55.22 | 8.63 | 2.5 |
| SHIPPO_MAG_265 | SHIPPO | 0 | 0 | NA | 0 | 1 | NA | 0 | 0 | NA | 0 | 0 | NA | 74.14 | 8.78 | 0.97 |
| SHIPPO_MAG_253 | SHIPPO | 0 | 0 | NA | 6 | 0 | orphan | 0 | 0 | NA | 0 | 0 | NA | 61.79 | 9.95 | 1.69 |
| SHIPPO_MAG_171 | SHIPPO | 0 | 0 | NA | 5 | 0 | orphan | 0 | 0 | NA | 0 | 2 | orphan | 82.04 | 9.99 | 2.7 |
| AG-414-L04 | GORG | 0 | 0 | NA | 1 | 0 | orphan | 0 | 0 | NA | 0 | 0 | NA | 98.8 | 0 | 1.21 |
| AG-422-C21 | GORG | 0 | 0 | NA | 1 | 0 | orphan | 0 | 0 | NA | 0 | 0 | NA | 98.8 | 0 | 1.28 |
| AG-891-K05 | GORG | 0 | 0 | NA | 2 | 0 | orphan | 0 | 1 | NA | 0 | 0 | NA | 98.8 | 0 | 1.65 |
| AG-892-F15 | GORG | 0 | 0 | NA | 1 | 0 | orphan | 0 | 0 | NA | 0 | 0 | NA | 98.8 | 0 | 1.4 |
| AG-893-A06 | GORG | 0 | 0 | NA | 1 | 0 | orphan | 0 | 0 | NA | 0 | 0 | NA | 98.8 | 0 | 1.59 |
| AG-900-I13 | GORG | 2 | 1 | complete | 1 | 0 | orphan | 0 | 0 | NA | 0 | 0 | NA | 98.8 | 0 | 1.31 |
| AG-912-G22 | GORG | 0 | 0 | NA | 1 | 0 | orphan | 0 | 0 | NA | 0 | 0 | NA | 98.8 | 0 | 1.19 |
| AG-920-L11 | GORG | 0 | 0 | NA | 2 | 0 | orphan | 0 | 0 | NA | 0 | 0 | NA | 98.8 | 0 | 1.25 |
| AG-917-K06 | GORG | 0 | 0 | NA | 1 | 0 | orphan | 0 | 0 | NA | 0 | 0 | NA | 98.65 | 0 | 1.19 |
| AG-893-K09 | GORG | 0 | 0 | NA | 1 | 0 | orphan | 0 | 0 | NA | 0 | 2 | orphan | 98.39 | 0 | 1.15 |

| | | | | | | | | | | | | | | | | |
|---|---|---|---|---|---|---|---|---|---|---|---|---|---|---|---|---|
| AG-891-I18 | GORG | 0 | 0 | NA | 1 | 0 | orphan | 0 | 0 | NA | 0 | 0 | NA | 97.63 | 0 | 1.38 |
| AG-891-J07 | GORG | 0 | 0 | NA | 2 | 0 | orphan | 0 | 0 | NA | 0 | 0 | NA | 97.63 | 0 | 1.21 |
| AG-891-C11 | GORG | 0 | 0 | NA | 1 | 0 | orphan | 0 | 0 | NA | 0 | 0 | NA | 97.59 | 0 | 1.57 |
| AG-891-J04 | GORG | 0 | 0 | NA | 3 | 0 | orphan | 0 | 0 | NA | 0 | 0 | NA | 97.59 | 0 | 1.35 |
| AG-904-K03 | GORG | 1 | 1 | complete | 1 | 0 | orphan | 0 | 0 | NA | 0 | 0 | NA | 97.59 | 0 | 1.35 |
| AG-905-D16 | GORG | 0 | 0 | NA | 1 | 0 | orphan | 0 | 0 | NA | 0 | 0 | NA | 97.59 | 0 | 1.26 |
| AG-909-F14 | GORG | 0 | 0 | NA | 1 | 0 | orphan | 0 | 0 | NA | 0 | 0 | NA | 97.59 | 0 | 1.3 |
| AG-917-F18 | GORG | 0 | 0 | NA | 1 | 0 | orphan | 0 | 0 | NA | 0 | 0 | NA | 97.59 | 0 | 1.08 |
| AG-920-L07 | GORG | 0 | 0 | NA | 1 | 0 | orphan | 0 | 0 | NA | 0 | 0 | NA | 97.59 | 0 | 1.09 |
| AG-918-G22 | GORG | 0 | 0 | NA | 1 | 0 | orphan | 0 | 0 | NA | 0 | 0 | NA | 97.48 | 0 | 1.36 |
| AG-918-N23 | GORG | 0 | 0 | NA | 2 | 0 | orphan | 0 | 0 | NA | 0 | 0 | NA | 97.19 | 0 | 1.01 |
| AG-899-E14 | GORG | 0 | 0 | NA | 2 | 1 | orphan | 0 | 0 | NA | 0 | 0 | NA | 96.99 | 0 | 1.05 |
| AG-913-I14 | GORG | 0 | 0 | NA | 1 | 0 | orphan | 0 | 0 | NA | 0 | 0 | NA | 96.99 | 0 | 1.05 |
| AG-325-E20 | GORG | 0 | 0 | NA | 1 | 0 | orphan | 0 | 0 | NA | 0 | 0 | NA | 96.39 | 0 | 1.07 |
| AG-325-F23 | GORG | 0 | 0 | NA | 1 | 0 | orphan | 0 | 0 | NA | 0 | 0 | NA | 96.39 | 0 | 1.62 |
| AG-422-L15 | GORG | 0 | 0 | NA | 2 | 0 | orphan | 0 | 0 | NA | 0 | 0 | NA | 96.39 | 0 | 1.04 |
| AG-461-E04 | GORG | 0 | 0 | NA | 1 | 0 | orphan | 0 | 0 | NA | 0 | 0 | NA | 96.39 | 0 | 1.01 |
| AG-893-J08 | GORG | 0 | 0 | NA | 2 | 0 | orphan | 0 | 0 | NA | 0 | 0 | NA | 96.39 | 0 | 1.34 |
| AG-896-A02 | GORG | 0 | 0 | NA | 2 | 0 | orphan | 0 | 0 | NA | 0 | 0 | NA | 96.39 | 0 | 1.23 |
| AG-897-G09 | GORG | 0 | 0 | NA | 2 | 0 | orphan | 0 | 0 | NA | 0 | 0 | NA | 96.39 | 0 | 1.3 |
| AG-899-O04 | GORG | 0 | 0 | NA | 2 | 0 | orphan | 0 | 0 | NA | 0 | 0 | NA | 96.39 | 0 | 1.23 |
| AG-905-M05 | GORG | 0 | 0 | NA | 2 | 0 | orphan | 0 | 0 | NA | 0 | 0 | NA | 96.39 | 0 | 1.18 |
| AG-917-D14 | GORG | 0 | 0 | NA | 1 | 0 | orphan | 0 | 0 | NA | 0 | 0 | NA | 96.39 | 0 | 1.23 |
| AG-900-I19 | GORG | 0 | 0 | NA | 1 | 0 | orphan | 0 | 0 | NA | 0 | 0 | NA | 96.28 | 0 | 1.19 |
| AG-913-L03 | GORG | 0 | 0 | NA | 1 | 0 | orphan | 0 | 0 | NA | 0 | 0 | NA | 96.21 | 0 | 1.29 |
| AG-911-G13 | GORG | 0 | 0 | NA | 1 | 0 | orphan | 0 | 0 | NA | 0 | 0 | NA | 95.98 | 0 | 1 |
| AG-892-K05 | GORG | 0 | 0 | NA | 1 | 0 | orphan | 0 | 0 | NA | 0 | 0 | NA | 95.97 | 0 | 1.04 |

| | | | | | | | | | | | | | | | | |
|---|---|---|---|---|---|---|---|---|---|---|---|---|---|---|---|---|
| AG-410-L19 | GORG | 0 | 0 | NA | 2 | 1 | complet e | 0 | 0 | NA | 0 | 0 | NA | 95.95 | 0 | 1.07 |
| AG-916-G19 | GORG | 0 | 0 | NA | 1 | 0 | orphan | 0 | 0 | NA | 0 | 0 | NA | 95.73 | 0 | 1.13 |
| AG-900-K21 | GORG | 0 | 0 | NA | 2 | 0 | orphan | 0 | 0 | NA | 0 | 0 | NA | 95.5 | 0 | 1.14 |
| AG-895-J15 | GORG | 0 | 0 | NA | 1 | 0 | orphan | 0 | 0 | NA | 0 | 0 | NA | 95.45 | 0 | 1.37 |
| AG-892-B11 | GORG | 0 | 0 | NA | 2 | 0 | orphan | 0 | 0 | NA | 0 | 0 | NA | 95.29 | 0 | 1.31 |
| AG-912-L11 | GORG | 0 | 0 | NA | 1 | 0 | orphan | 0 | 0 | NA | 0 | 0 | NA | 95.29 | 0 | 1.23 |
| AG-422-E02 | GORG | 0 | 0 | NA | 2 | 0 | orphan | 0 | 1 | NA | 0 | 0 | NA | 95.18 | 0 | 1.63 |
| AG-892-J03 | GORG | 0 | 0 | NA | 1 | 0 | orphan | 0 | 0 | NA | 0 | 0 | NA | 95.18 | 0 | 1.38 |
| AG-894-B06 | GORG | 1 | 1 | complet e | 1 | 0 | orphan | 0 | 0 | NA | 0 | 0 | NA | 95.18 | 0 | 1.23 |
| AG-903-L07 | GORG | 0 | 0 | NA | 1 | 0 | orphan | 0 | 0 | NA | 0 | 0 | NA | 95.18 | 0 | 1.26 |
| AG-919-I13 | GORG | 0 | 0 | NA | 1 | 0 | orphan | 0 | 0 | NA | 0 | 0 | NA | 95.18 | 0 | 1.04 |
| AG-915-K13 | GORG | 0 | 0 | NA | 3 | 0 | orphan | 0 | 0 | NA | 0 | 0 | NA | 95.02 | 0 | 1.25 |
| AG-913-M13 | GORG | 0 | 0 | NA | 2 | 0 | orphan | 0 | 0 | NA | 0 | 0 | NA | 94.94 | 0 | 1.18 |
| AG-390-M23 | GORG | 0 | 0 | NA | 2 | 0 | orphan | 0 | 0 | NA | 0 | 0 | NA | 94.85 | 0 | 1.58 |
| AG-900-G16 | GORG | 0 | 0 | NA | 2 | 0 | orphan | 0 | 0 | NA | 0 | 0 | NA | 94.83 | 0 | 1.31 |
| AG-895-J19 | GORG | 0 | 0 | NA | 2 | 0 | orphan | 0 | 0 | NA | 0 | 0 | NA | 94.79 | 0 | 1.08 |
| AG-891-C15 | GORG | 0 | 0 | NA | 2 | 0 | orphan | 0 | 0 | NA | 0 | 0 | NA | 94.78 | 0 | 1.17 |
| AG-913-I06 | GORG | 0 | 0 | NA | 2 | 0 | orphan | 0 | 0 | NA | 0 | 0 | NA | 94.78 | 0 | 1.03 |
| AG-390-C19 | GORG | 0 | 0 | NA | 2 | 0 | orphan | 0 | 0 | NA | 0 | 0 | NA | 94.58 | 0 | 1.16 |
| AG-893-E23 | GORG | 0 | 0 | NA | 2 | 0 | orphan | 0 | 0 | NA | 0 | 0 | NA | 94.58 | 0 | 1.39 |
| AG-893-F05 | GORG | 0 | 0 | NA | 1 | 0 | orphan | 0 | 0 | NA | 0 | 0 | NA | 94.58 | 0 | 1.45 |
| AG-426-P08 | GORG | 0 | 0 | NA | 2 | 0 | orphan | 0 | 0 | NA | 0 | 0 | NA | 94.55 | 0 | 1.13 |
| AG-894-N05 | GORG | 1 | 0 | orphan | 0 | 0 | NA | 0 | 0 | NA | 0 | 1 | orpha n | 94.43 | 0 | 1.54 |
| AG-422-C20 | GORG | 0 | 0 | NA | 1 | 0 | orphan | 0 | 0 | NA | 0 | 0 | NA | 94.31 | 0 | 1.31 |
| AG-453-G16 | GORG | 0 | 0 | NA | 3 | 0 | orphan | 0 | 0 | NA | 0 | 0 | NA | 94.22 | 0 | 1.8 |
| AG-913-I09 | GORG | 0 | 0 | NA | 1 | 0 | orphan | 0 | 0 | NA | 0 | 0 | NA | 94.16 | 0 | 1.28 |

| | | | | | | | | | | | | | | | | |
|---|---|---|---|---|---|---|---|---|---|---|---|---|---|---|---|---|
| AG-893-K07 | GORG | 0 | 0 | NA | 2 | 0 | orphan | 0 | 0 | NA | 0 | 0 | NA | 94.14 | 0 | 2.11 |
| AG-894-L13 | GORG | 0 | 0 | NA | 1 | 0 | orphan | 0 | 0 | NA | 0 | 0 | NA | 94.08 | 0 | 1.12 |
| AG-447-N08 | GORG | 0 | 0 | NA | 1 | 0 | orphan | 0 | 0 | NA | 0 | 0 | NA | 94.05 | 0 | 1.18 |
| AG-895-L23 | GORG | 0 | 0 | NA | 1 | 0 | orphan | 0 | 0 | NA | 0 | 0 | NA | 94.05 | 0 | 0.97 |
| AG-919-L02 | GORG | 0 | 0 | NA | 2 | 0 | orphan | 0 | 0 | NA | 0 | 0 | NA | 94 | 0 | 1.14 |
| AG-426-I04 | GORG | 0 | 0 | NA | 2 | 0 | orphan | 0 | 0 | NA | 0 | 1 | orphan | 93.98 | 0 | 1.52 |
| AG-365-B19 | GORG | 0 | 0 | NA | 1 | 0 | orphan | 0 | 0 | NA | 0 | 0 | NA | 93.98 | 0 | 1.04 |
| AG-404-B13 | GORG | 0 | 0 | NA | 1 | 0 | orphan | 0 | 0 | NA | 0 | 0 | NA | 93.98 | 0 | 1.43 |
| AG-430-F03 | GORG | 0 | 0 | NA | 1 | 0 | orphan | 0 | 0 | NA | 0 | 0 | NA | 93.98 | 0 | 1.01 |
| AG-439-C18 | GORG | 0 | 0 | NA | 1 | 0 | orphan | 0 | 0 | NA | 0 | 0 | NA | 93.98 | 0 | 1.14 |
| AG-892-M06 | GORG | 1 | 1 | complete | 1 | 0 | orphan | 0 | 0 | NA | 0 | 0 | NA | 93.98 | 0 | 1.24 |
| AG-914-J05 | GORG | 1 | 1 | complete | 1 | 0 | orphan | 0 | 0 | NA | 0 | 0 | NA | 93.98 | 0 | 1.04 |
| AG-917-D03 | GORG | 0 | 0 | NA | 1 | 0 | orphan | 0 | 0 | NA | 0 | 0 | NA | 93.98 | 0 | 1.28 |
| AG-911-K15 | GORG | 0 | 0 | NA | 4 | 0 | orphan | 0 | 0 | NA | 0 | 0 | NA | 93.91 | 0 | 1.14 |
| AG-916-A16 | GORG | 0 | 0 | NA | 2 | 0 | orphan | 0 | 0 | NA | 0 | 0 | NA | 93.84 | 0 | 1.13 |
| AH-321-M13 | GORG | 0 | 0 | NA | 2 | 0 | orphan | 0 | 0 | NA | 0 | 0 | NA | 93.84 | 0 | 1.14 |
| AG-893-J23 | GORG | 1 | 1 | complete | 0 | 0 | NA | 0 | 0 | NA | 0 | 1 | orphan | 93.75 | 0 | 1.52 |
| AG-920-B11 | GORG | 0 | 0 | NA | 1 | 0 | orphan | 0 | 0 | NA | 0 | 0 | NA | 93.75 | 0 | 1.83 |
| AG-919-J11 | GORG | 1 | 1 | complete | 0 | 0 | NA | 0 | 0 | NA | 0 | 0 | NA | 93.66 | 0 | 1.5 |
| AG-894-M09 | GORG | 1 | 1 | complete | 2 | 0 | orphan | 0 | 0 | NA | 0 | 0 | NA | 93.61 | 0 | 2.17 |
| AG-917-B07 | GORG | 0 | 0 | NA | 1 | 0 | orphan | 0 | 0 | NA | 0 | 0 | NA | 93.57 | 0 | 1.09 |
| AG-917-C08 | GORG | 0 | 0 | NA | 1 | 0 | orphan | 1 | 1 | complete | 0 | 0 | NA | 93.41 | 0 | 1.22 |
| AG-916-N15 | GORG | 0 | 0 | NA | 1 | 0 | orphan | 0 | 0 | NA | 0 | 2 | orphan | 93.37 | 0 | 1.26 |
| AG-337-E15 | GORG | 0 | 0 | NA | 1 | 0 | orphan | 0 | 0 | NA | 0 | 0 | NA | 93.37 | 0 | 1.01 |

| | | | | | | | | | | | | | | | | |
|---|---|---|---|---|---|---|---|---|---|---|---|---|---|---|---|---|
| AG-892-O09 | GORG | 0 | 0 | NA | 1 | 0 | orphan | 0 | 0 | NA | 0 | 0 | NA | 93.37 | 0 | 1.25 |
| AG-893-E21 | GORG | 1 | 1 | complete | 1 | 0 | orphan | 0 | 0 | NA | 0 | 0 | NA | 93.37 | 0 | 1.3 |
| AG-916-M14 | GORG | 0 | 0 | NA | 1 | 0 | orphan | 0 | 0 | NA | 0 | 0 | NA | 93.37 | 0 | 1.37 |
| AG-917-M08 | GORG | 0 | 0 | NA | 2 | 0 | orphan | 0 | 0 | NA | 0 | 0 | NA | 93.37 | 0 | 1.39 |
| AH-324-F03 | GORG | 0 | 0 | NA | 3 | 0 | orphan | 0 | 0 | NA | 0 | 0 | NA | 93.37 | 0 | 1.22 |
| AG-895-F17 | GORG | 0 | 0 | NA | 1 | 0 | orphan | 0 | 0 | NA | 0 | 0 | NA | 93.36 | 0 | 1.2 |
| AG-447-P22 | GORG | 0 | 0 | NA | 4 | 0 | orphan | 0 | 0 | NA | 0 | 0 | NA | 93.33 | 0 | 1.12 |
| AG-895-K10 | GORG | 0 | 0 | NA | 1 | 0 | orphan | 0 | 0 | NA | 0 | 0 | NA | 93.33 | 0 | 1.11 |
| AG-894-P13 | GORG | 1 | 1 | complete | 0 | 0 | NA | 0 | 0 | NA | 0 | 1 | orphan | 93.25 | 0 | 1.44 |
| AG-390-G13 | GORG | 0 | 0 | NA | 1 | 0 | orphan | 0 | 0 | NA | 0 | 0 | NA | 93.23 | 0 | 1.09 |
| AG-390-C17 | GORG | 1 | 0 | orphan | 1 | 0 | orphan | 0 | 0 | NA | 0 | 0 | NA | 93.21 | 0 | 1.29 |
| AG-910-B05 | GORG | 0 | 0 | NA | 2 | 0 | orphan | 0 | 0 | NA | 0 | 0 | NA | 93.19 | 0 | 1.08 |
| AG-891-J21 | GORG | 0 | 0 | NA | 2 | 0 | orphan | 0 | 0 | NA | 0 | 2 | orphan | 93.01 | 0 | 1.36 |
| AG-916-I16 | GORG | 1 | 1 | complete | 1 | 0 | orphan | 0 | 0 | NA | 0 | 0 | NA | 92.99 | 0 | 1.21 |
| AH-321-C14 | GORG | 1 | 1 | complete | 2 | 0 | orphan | 0 | 0 | NA | 0 | 0 | NA | 92.93 | 0 | 1.53 |
| AG-359-M21 | GORG | 0 | 0 | NA | 1 | 1 | orphan | 0 | 0 | NA | 0 | 0 | NA | 92.89 | 0 | 1.17 |
| AG-325-G21 | GORG | 0 | 0 | NA | 1 | 0 | orphan | 0 | 0 | NA | 0 | 0 | NA | 92.88 | 0 | 1.09 |
| AG-904-F11 | GORG | 0 | 0 | NA | 2 | 0 | orphan | 0 | 0 | NA | 0 | 0 | NA | 92.81 | 0 | 0.96 |
| AG-422-J16 | GORG | 0 | 0 | NA | 1 | 0 | orphan | 0 | 0 | NA | 0 | 1 | orphan | 92.77 | 0 | 1.47 |
| AG-426-O05 | GORG | 0 | 0 | NA | 1 | 0 | orphan | 0 | 0 | NA | 0 | 1 | orphan | 92.77 | 0 | 1 |
| AG-390-G04 | GORG | 0 | 0 | NA | 1 | 0 | orphan | 0 | 0 | NA | 0 | 0 | NA | 92.77 | 0 | 1.64 |
| AG-426-I21 | GORG | 0 | 0 | NA | 2 | 1 | complete | 0 | 0 | NA | 0 | 0 | NA | 92.77 | 0 | 1.27 |
| AG-439-D15 | GORG | 0 | 0 | NA | 3 | 0 | orphan | 0 | 0 | NA | 0 | 0 | NA | 92.77 | 0 | 1.15 |
| AG-461-L09 | GORG | 0 | 0 | NA | 1 | 0 | orphan | 0 | 0 | NA | 0 | 0 | NA | 92.77 | 0 | 1.24 |

| | | | | | | | | | | | | | | | | |
|---|---|---|---|---|---|---|---|---|---|---|---|---|---|---|---|---|
| AG-891-D10 | GORG | 0 | 0 | NA | 1 | 0 | orphan | 0 | 0 | NA | 0 | 0 | NA | 92.77 | 0 | 1.07 |
| AG-905-E11 | GORG | 1 | 1 | complete | 1 | 0 | orphan | 0 | 0 | NA | 0 | 0 | NA | 92.77 | 0 | 1.32 |
| AG-912-P19 | GORG | 0 | 0 | NA | 1 | 0 | orphan | 0 | 0 | NA | 0 | 0 | NA | 92.77 | 0 | 0.92 |
| AG-914-I16 | GORG | 0 | 0 | NA | 2 | 0 | orphan | 0 | 0 | NA | 0 | 0 | NA | 92.77 | 0 | 0.99 |
| AG-917-F16 | GORG | 0 | 0 | NA | 1 | 0 | orphan | 0 | 0 | NA | 0 | 0 | NA | 92.77 | 0 | 1.16 |
| AG-919-F21 | GORG | 0 | 0 | NA | 2 | 0 | orphan | 0 | 0 | NA | 0 | 0 | NA | 92.77 | 0 | 1.06 |
| AG-919-M05 | GORG | 0 | 0 | NA | 1 | 0 | orphan | 0 | 0 | NA | 0 | 0 | NA | 92.77 | 0 | 1.04 |
| AH-324-F15 | GORG | 0 | 0 | NA | 1 | 1 | orphan | 0 | 0 | NA | 0 | 0 | NA | 92.77 | 0 | 1.48 |
| AG-893-E15 | GORG | 0 | 0 | NA | 1 | 0 | orphan | 0 | 0 | NA | 0 | 0 | NA | 92.72 | 0 | 1.3 |
| AH-321-D17 | GORG | 0 | 0 | NA | 2 | 0 | orphan | 0 | 0 | NA | 0 | 0 | NA | 92.42 | 0 | 1.1 |
| AG-893-A21 | GORG | 0 | 0 | NA | 1 | 0 | orphan | 0 | 0 | NA | 0 | 0 | NA | 92.37 | 0 | 1.54 |
| AG-919-F03 | GORG | 0 | 0 | NA | 2 | 0 | orphan | 0 | 1 | NA | 0 | 0 | NA | 92.37 | 0 | 1.12 |
| AG-920-L19 | GORG | 0 | 0 | NA | 2 | 0 | orphan | 0 | 0 | NA | 0 | 0 | NA | 92.37 | 0 | 1.07 |
| AG-920-L20 | GORG | 0 | 0 | NA | 1 | 0 | orphan | 0 | 0 | NA | 0 | 0 | NA | 92.33 | 0 | 1.28 |
| AG-916-G20 | GORG | 0 | 0 | NA | 1 | 0 | orphan | 0 | 0 | NA | 0 | 0 | NA | 92.28 | 0 | 1.43 |
| AG-337-G11 | GORG | 0 | 0 | NA | 1 | 0 | orphan | 0 | 0 | NA | 0 | 0 | NA | 92.23 | 0 | 1.06 |
| AG-893-L07 | GORG | 1 | 1 | complete | 1 | 0 | orphan | 0 | 0 | NA | 0 | 0 | NA | 92.22 | 0 | 1.32 |
| AH-321-P04 | GORG | 0 | 0 | NA | 2 | 0 | orphan | 0 | 0 | NA | 0 | 0 | NA | 92.18 | 0 | 1.02 |
| AG-892-D04 | GORG | 0 | 0 | NA | 1 | 0 | orphan | 0 | 0 | NA | 0 | 0 | NA | 92.17 | 0 | 1.17 |
| AG-901-E23 | GORG | 0 | 0 | NA | 2 | 0 | orphan | 0 | 0 | NA | 0 | 0 | NA | 92.17 | 0 | 1.29 |
| AG-912-J07 | GORG | 0 | 0 | NA | 2 | 0 | orphan | 0 | 0 | NA | 0 | 0 | NA | 92.17 | 0 | 1.24 |
| AG-918-E02 | GORG | 0 | 1 | NA | 2 | 2 | complete | 0 | 0 | NA | 0 | 0 | NA | 92.17 | 0 | 1.26 |
| AG-918-M15 | GORG | 0 | 0 | NA | 1 | 0 | orphan | 0 | 0 | NA | 0 | 0 | NA | 92.17 | 0 | 0.99 |
| AG-907-E04 | GORG | 0 | 0 | NA | 1 | 0 | orphan | 0 | 0 | NA | 0 | 0 | NA | 92 | 0 | 1.03 |
| AG-900-M14 | GORG | 0 | 0 | NA | 2 | 0 | orphan | 0 | 0 | NA | 0 | 1 | orphan | 91.95 | 0 | 1.26 |
| AG-905-A07 | GORG | 0 | 0 | NA | 2 | 0 | orphan | 0 | 1 | NA | 0 | 0 | NA | 91.95 | 0 | 1.12 |

| | | | | | | | | | | | | | | | | |
|---|---|---|---|---|---|---|---|---|---|---|---|---|---|---|---|---|
| AG-325-C18 | GORG | 1 | 0 | orphan | 2 | 0 | orphan | 0 | 0 | NA | 0 | 0 | NA | 91.94 | 0 | 1.07 |
| AG-913-C06 | GORG | 1 | 1 | complete | 1 | 0 | orphan | 0 | 0 | NA | 0 | 0 | NA | 91.91 | 0 | 1.73 |
| AG-359-D09 | GORG | 0 | 0 | NA | 1 | 0 | orphan | 0 | 0 | NA | 0 | 1 | orphan | 91.86 | 0 | 1.2 |
| AG-430-G11 | GORG | 0 | 0 | NA | 1 | 0 | orphan | 0 | 0 | NA | 0 | 0 | NA | 91.77 | 0 | 1.16 |
| AH-287-G05 | GORG | 0 | 0 | NA | 1 | 0 | orphan | 0 | 0 | NA | 0 | 0 | NA | 91.76 | 0 | 1.13 |
| AG-447-K09 | GORG | 0 | 0 | NA | 3 | 0 | orphan | 0 | 0 | NA | 0 | 0 | NA | 91.71 | 0 | 1.19 |
| AG-892-A22 | GORG | 1 | 1 | complete | 1 | 0 | orphan | 0 | 0 | NA | 0 | 0 | NA | 91.71 | 0 | 1.19 |
| AG-896-K10 | GORG | 0 | 0 | NA | 2 | 0 | orphan | 0 | 0 | NA | 0 | 0 | NA | 91.71 | 0 | 1.11 |
| AG-917-J06 | GORG | 0 | 0 | NA | 3 | 0 | orphan | 0 | 0 | NA | 0 | 0 | NA | 91.63 | 0 | 1.21 |
| AG-414-N23 | GORG | 1 | 0 | orphan | 1 | 0 | orphan | 0 | 0 | NA | 0 | 2 | orphan | 91.57 | 0 | 1.5 |
| AG-422-N15 | GORG | 0 | 0 | NA | 1 | 0 | orphan | 0 | 0 | NA | 0 | 2 | orphan | 91.57 | 0 | 1.04 |
| AG-426-F15 | GORG | 0 | 0 | NA | 1 | 0 | orphan | 0 | 0 | NA | 0 | 0 | NA | 91.57 | 0 | 1.36 |
| AG-426-L15 | GORG | 0 | 0 | NA | 1 | 0 | orphan | 0 | 0 | NA | 0 | 0 | NA | 91.57 | 0 | 0.96 |
| AG-891-O15 | GORG | 0 | 0 | NA | 1 | 0 | orphan | 0 | 0 | NA | 0 | 0 | NA | 91.57 | 0 | 1.04 |
| AG-913-B05 | GORG | 0 | 0 | NA | 1 | 0 | orphan | 0 | 0 | NA | 0 | 0 | NA | 91.57 | 0 | 1.46 |
| AH-287-E19 | GORG | 0 | 0 | NA | 1 | 0 | orphan | 0 | 0 | NA | 0 | 0 | NA | 91.57 | 0 | 0.97 |
| AH-287-O09 | GORG | 0 | 0 | NA | 2 | 0 | orphan | 0 | 0 | NA | 0 | 0 | NA | 91.57 | 0 | 1.16 |
| AH-321-I23 | GORG | 0 | 0 | NA | 1 | 0 | orphan | 0 | 0 | NA | 0 | 0 | NA | 91.57 | 0 | 0.91 |
| AG-895-D05 | GORG | 0 | 0 | NA | 0 | 0 | NA | 0 | 0 | NA | 0 | 1 | orphan | 91.49 | 0 | 1.42 |
| AG-918-K13 | GORG | 0 | 0 | NA | 2 | 0 | orphan | 0 | 0 | NA | 0 | 0 | NA | 91.24 | 0 | 1.27 |
| AG-920-M14 | GORG | 0 | 0 | NA | 0 | 0 | NA | 0 | 0 | NA | 0 | 1 | orphan | 91.17 | 0 | 1.5 |
| AG-917-N15 | GORG | 0 | 0 | NA | 1 | 0 | orphan | 0 | 0 | NA | 0 | 0 | NA | 91.14 | 0 | 1.21 |
| AG-359-P18 | GORG | 0 | 0 | NA | 1 | 0 | orphan | 0 | 0 | NA | 0 | 0 | NA | 91.1 | 0 | 1.07 |
| AG-333-D04 | GORG | 0 | 0 | NA | 1 | 0 | orphan | 0 | 0 | NA | 0 | 0 | NA | 91.09 | 0 | 1.68 |
| AG-461-M18 | GORG | 0 | 0 | NA | 1 | 0 | orphan | 0 | 0 | NA | 0 | 0 | NA | 91.09 | 0 | 1.02 |

| | | | | | | | | | | | | | | | | |
|---|---|---|---|---|---|---|---|---|---|---|---|---|---|---|---|---|
| AG-917-P07 | GORG | 0 | 0 | NA | 1 | 0 | orphan | 0 | 0 | NA | 0 | 0 | NA | 91 | 0 | 1.16 |
| AG-893-P02 | GORG | 0 | 0 | NA | 2 | 0 | orphan | 0 | 0 | NA | 0 | 0 | NA | 90.96 | 0 | 1.08 |
| AG-916-I19 | GORG | 0 | 0 | NA | 1 | 0 | orphan | 0 | 0 | NA | 0 | 0 | NA | 90.96 | 0 | 1.43 |
| AG-422-K10 | GORG | 0 | 0 | NA | 0 | 0 | NA | 0 | 0 | NA | 0 | 3 | orphan | 90.85 | 0 | 1.38 |
| AG-447-F10 | GORG | 0 | 0 | NA | 1 | 0 | orphan | 0 | 0 | NA | 0 | 0 | NA | 90.84 | 0 | 1.06 |
| AG-913-K11 | GORG | 0 | 0 | NA | 1 | 0 | orphan | 1 | 1 | complete | 0 | 0 | NA | 90.81 | 0 | 1.48 |
| AG-430-A11 | GORG | 0 | 0 | NA | 3 | 0 | orphan | 0 | 0 | NA | 0 | 0 | NA | 90.76 | 0 | 1.22 |
| AG-426-I10 | GORG | 0 | 0 | NA | 1 | 0 | orphan | 0 | 0 | NA | 0 | 0 | NA | 90.74 | 0 | 1.14 |
| AG-908-C18 | GORG | 0 | 0 | NA | 1 | 0 | orphan | 0 | 0 | NA | 0 | 0 | NA | 90.73 | 0 | 1.16 |
| AG-919-C02 | GORG | 0 | 0 | NA | 2 | 0 | orphan | 0 | 0 | NA | 0 | 0 | NA | 90.73 | 0 | 1.13 |
| AG-349-G23 | GORG | 1 | 1 | complete | 0 | 0 | NA | 0 | 0 | NA | 0 | 2 | orphan | 90.68 | 0 | 1.41 |
| AG-410-O02 | GORG | 0 | 0 | NA | 1 | 0 | orphan | 0 | 0 | NA | 0 | 0 | NA | 90.64 | 0 | 1.12 |
| AG-893-M17 | GORG | 0 | 0 | NA | 1 | 0 | orphan | 0 | 0 | NA | 0 | 0 | NA | 90.64 | 0 | 1.28 |
| AG-891-J06 | GORG | 1 | 1 | complete | 1 | 0 | orphan | 0 | 0 | NA | 0 | 0 | NA | 90.62 | 0 | 1.32 |
| AG-917-M22 | GORG | 1 | 0 | orphan | 1 | 0 | orphan | 0 | 0 | NA | 0 | 0 | NA | 90.52 | 0 | 1.13 |
| AG-919-L14 | GORG | 0 | 0 | NA | 2 | 0 | orphan | 0 | 0 | NA | 0 | 0 | NA | 90.52 | 0 | 0.98 |
| AG-349-I05 | GORG | 1 | 1 | complete | 1 | 0 | orphan | 0 | 0 | NA | 0 | 1 | orphan | 90.36 | 0 | 1.14 |
| AG-914-N11 | GORG | 0 | 0 | NA | 1 | 0 | orphan | 0 | 0 | NA | 0 | 1 | orphan | 90.36 | 0 | 0.96 |
| AG-891-P20 | GORG | 0 | 0 | NA | 3 | 0 | orphan | 0 | 0 | NA | 0 | 0 | NA | 90.36 | 0 | 1.48 |
| AG-894-F15 | GORG | 1 | 0 | orphan | 1 | 0 | orphan | 0 | 0 | NA | 0 | 0 | NA | 90.36 | 0 | 1.23 |
| AG-897-O16 | GORG | 0 | 0 | NA | 1 | 0 | orphan | 0 | 0 | NA | 0 | 0 | NA | 90.36 | 0 | 1.18 |
| AG-900-K14 | GORG | 0 | 0 | NA | 1 | 0 | orphan | 0 | 0 | NA | 0 | 0 | NA | 90.36 | 0 | 1.26 |
| AG-903-K07 | GORG | 0 | 0 | NA | 1 | 0 | orphan | 0 | 0 | NA | 0 | 0 | NA | 90.36 | 0 | 1.07 |
| AG-905-O08 | GORG | 0 | 0 | NA | 1 | 0 | orphan | 0 | 0 | NA | 0 | 0 | NA | 90.36 | 0 | 1.21 |
| AG-915-G13 | GORG | 0 | 0 | NA | 1 | 0 | orphan | 0 | 0 | NA | 0 | 0 | NA | 90.36 | 0 | 1.24 |

| | | | | | | | | | | | | | | | | |
|---|---|---|---|---|---|---|---|---|---|---|---|---|---|---|---|---|
| AG-916-G09 | GORG | 0 | 0 | NA | 1 | 0 | orphan | 0 | 0 | NA | 0 | 0 | NA | 90.36 | 0 | 1.27 |
| AG-916-O13 | GORG | 0 | 0 | NA | 2 | 0 | orphan | 0 | 0 | NA | 0 | 0 | NA | 90.36 | 0 | 1.08 |
| AG-917-E05 | GORG | 1 | 0 | orphan | 0 | 0 | NA | 0 | 0 | NA | 0 | 0 | NA | 90.36 | 0 | 1.21 |
| AG-917-K20 | GORG | 0 | 0 | NA | 1 | 0 | orphan | 0 | 0 | NA | 0 | 0 | NA | 90.36 | 0 | 1.03 |
| AG-422-C15 | GORG | 0 | 0 | NA | 1 | 0 | orphan | 0 | 0 | NA | 0 | 1 | orphan | 90.35 | 0 | 1.48 |
| AG-912-F16 | GORG | 0 | 0 | NA | 1 | 0 | orphan | 0 | 0 | NA | 0 | 0 | NA | 90.28 | 0 | 1.15 |
| AH-324-K07 | GORG | 0 | 0 | NA | 1 | 0 | orphan | 0 | 0 | NA | 0 | 0 | NA | 90.15 | 0 | 1.06 |
| AG-333-A04 | GORG | 0 | 0 | NA | 2 | 0 | orphan | 0 | 0 | NA | 0 | 0 | NA | 90.05 | 0 | 1.32 |
| AG-390-M14 | GORG | 0 | 0 | NA | 2 | 0 | orphan | 0 | 0 | NA | 0 | 0 | NA | 90.05 | 0 | 1.09 |
| AG-920-L18 | GORG | 0 | 0 | NA | 2 | 0 | orphan | 0 | 0 | NA | 0 | 0 | NA | 90.04 | 0 | 0.96 |
| AG-894-A11 | GORG | 1 | 1 | complete | 1 | 0 | orphan | 0 | 0 | NA | 0 | 0 | NA | 90.01 | 0 | 1.28 |
| AG-916-J11 | GORG | 0 | 0 | NA | 1 | 0 | orphan | 0 | 0 | NA | 0 | 0 | NA | 90.01 | 0 | 1.1 |
| AG-901-I21 | GORG | 0 | 0 | NA | 1 | 0 | orphan | 0 | 0 | NA | 0 | 2 | orphan | 89.96 | 0 | 1.01 |
| AG-911-N14 | GORG | 0 | 0 | NA | 1 | 0 | orphan | 0 | 0 | NA | 0 | 0 | NA | 89.96 | 0 | 1.18 |
| AG-915-D08 | GORG | 1 | 1 | complete | 1 | 0 | orphan | 0 | 0 | NA | 0 | 0 | NA | 89.96 | 0 | 0.94 |
| AG-896-E23 | GORG | 0 | 0 | NA | 1 | 0 | orphan | 0 | 0 | NA | 0 | 1 | orphan | 89.95 | 0 | 1.45 |
| AG-313-D08 | GORG | 0 | 0 | NA | 2 | 0 | orphan | 0 | 0 | NA | 0 | 0 | NA | 89.9 | 0 | 1.65 |
| AG-915-G14 | GORG | 0 | 0 | NA | 1 | 0 | orphan | 0 | 0 | NA | 0 | 0 | NA | 89.89 | 0 | 1.32 |
| AG-893-E16 | GORG | 0 | 0 | NA | 1 | 0 | orphan | 0 | 0 | NA | 0 | 0 | NA | 89.87 | 0 | 1.37 |
| AH-324-M03 | GORG | 0 | 0 | NA | 3 | 0 | orphan | 0 | 0 | NA | 0 | 0 | NA | 89.81 | 0 | 0.93 |
| AG-349-D08 | GORG | 0 | 0 | NA | 1 | 0 | orphan | 0 | 0 | NA | 0 | 0 | NA | 89.76 | 0 | 0.9 |
| AG-426-J10 | GORG | 0 | 0 | NA | 1 | 0 | orphan | 0 | 0 | NA | 0 | 0 | NA | 89.76 | 0 | 1.45 |
| AG-911-E22 | GORG | 0 | 0 | NA | 2 | 0 | orphan | 0 | 0 | NA | 0 | 0 | NA | 89.76 | 0 | 1.3 |
| AG-912-D10 | GORG | 0 | 0 | NA | 1 | 0 | orphan | 0 | 0 | NA | 0 | 0 | NA | 89.76 | 0 | 0.98 |
| AG-919-N08 | GORG | 1 | 0 | orphan | 1 | 0 | orphan | 0 | 0 | NA | 0 | 0 | NA | 89.76 | 0 | 1.33 |

| | | | | | | | | | | | | | | | | |
|---|---|---|---|---|---|---|---|---|---|---|---|---|---|---|---|---|
| AG-896-O18 | GORG | 0 | 0 | NA | 1 | 0 | orphan | 0 | 0 | NA | 0 | 0 | NA | 89.71 | 0 | 1.29 |
| AG-900-K16 | GORG | 0 | 0 | NA | 1 | 0 | orphan | 0 | 0 | NA | 0 | 0 | NA | 89.71 | 0 | 1.35 |
| AH-287-O21 | GORG | 0 | 0 | NA | 2 | 1 | orphan | 0 | 0 | NA | 0 | 0 | NA | 89.68 | 0 | 2.01 |
| AG-894-O09 | GORG | 0 | 0 | NA | 1 | 0 | orphan | 0 | 0 | NA | 0 | 0 | NA | 89.57 | 0 | 1.2 |
| AG-903-F09 | GORG | 0 | 0 | NA | 2 | 0 | orphan | 0 | 0 | NA | 0 | 0 | NA | 89.57 | 0 | 1.08 |
| AH-324-J07 | GORG | 0 | 0 | NA | 2 | 2 | complete | 0 | 0 | NA | 0 | 0 | NA | 89.57 | 0 | 1.31 |
| AG-892-F02 | GORG | 0 | 0 | NA | 1 | 0 | orphan | 0 | 0 | NA | 0 | 0 | NA | 89.56 | 0 | 1.16 |
| AG-899-G06 | GORG | 0 | 0 | NA | 1 | 0 | orphan | 0 | 0 | NA | 0 | 0 | NA | 89.56 | 0 | 0.97 |
| AG-904-O22 | GORG | 1 | 1 | complete | 0 | 0 | NA | 0 | 0 | NA | 0 | 1 | orphan | 89.54 | 0 | 1.39 |
| AG-439-D05 | GORG | 0 | 0 | NA | 1 | 0 | orphan | 0 | 0 | NA | 0 | 0 | NA | 89.53 | 0 | 1.19 |
| AG-911-G06 | GORG | 0 | 0 | NA | 2 | 0 | orphan | 0 | 0 | NA | 0 | 0 | NA | 89.52 | 0 | 0.97 |
| AG-439-I19 | GORG | 0 | 0 | NA | 1 | 0 | orphan | 0 | 0 | NA | 0 | 0 | NA | 89.51 | 0 | 1.09 |
| AG-470-L20 | GORG | 0 | 0 | NA | 2 | 0 | orphan | 0 | 0 | NA | 0 | 0 | NA | 89.41 | 0 | 1.07 |
| AG-914-I23 | GORG | 0 | 0 | NA | 0 | 0 | NA | 0 | 0 | NA | 0 | 1 | orphan | 89.4 | 0 | 1.36 |
| AG-914-A09 | GORG | 0 | 0 | NA | 1 | 0 | orphan | 0 | 0 | NA | 0 | 0 | NA | 89.37 | 0 | 0.97 |
| AG-908-L09 | GORG | 0 | 0 | NA | 1 | 0 | orphan | 0 | 0 | NA | 0 | 0 | NA | 89.3 | 0 | 1.27 |
| AG-893-M22 | GORG | 0 | 0 | NA | 2 | 0 | orphan | 0 | 0 | NA | 0 | 0 | NA | 89.27 | 0 | 1.33 |
| AG-430-N16 | GORG | 0 | 0 | NA | 1 | 0 | orphan | 0 | 0 | NA | 0 | 0 | NA | 89.16 | 0 | 1.06 |
| AG-470-D16 | GORG | 0 | 0 | NA | 1 | 0 | orphan | 0 | 0 | NA | 0 | 0 | NA | 89.16 | 0 | 1 |
| AG-893-I03 | GORG | 0 | 0 | NA | 1 | 0 | orphan | 0 | 0 | NA | 0 | 0 | NA | 89.16 | 0 | 1.48 |
| AG-899-D19 | GORG | 1 | 0 | orphan | 1 | 0 | orphan | 0 | 0 | NA | 0 | 0 | NA | 89.16 | 0 | 1.22 |
| AG-899-M20 | GORG | 0 | 0 | NA | 3 | 0 | orphan | 0 | 0 | NA | 0 | 0 | NA | 89.16 | 0 | 0.91 |
| AG-911-J19 | GORG | 1 | 0 | orphan | 1 | 0 | orphan | 0 | 0 | NA | 0 | 0 | NA | 89.16 | 0 | 1.26 |
| AG-916-O20 | GORG | 1 | 1 | complete | 1 | 0 | orphan | 0 | 0 | NA | 0 | 0 | NA | 89.16 | 0 | 1.22 |
| AG-913-J04 | GORG | 0 | 0 | NA | 2 | 0 | orphan | 0 | 0 | NA | 0 | 0 | NA | 89.08 | 0 | 1.85 |
| AG-390-E04 | GORG | 0 | 0 | NA | 2 | 0 | orphan | 0 | 0 | NA | 0 | 0 | NA | 89.05 | 0 | 1.14 |

| | | | | | | | | | | | | | | | | |
|---|---|---|---|---|---|---|---|---|---|---|---|---|---|---|---|---|
| AG-894-L19 | GORG | 0 | 0 | NA | 1 | 0 | orphan | 0 | 0 | NA | 0 | 0 | NA | 88.98 | 0 | 1.16 |
| AG-912-J06 | GORG | 0 | 0 | NA | 2 | 0 | orphan | 0 | 1 | NA | 0 | 0 | NA | 88.87 | 0 | 0.92 |
| AG-359-G18 | GORG | 0 | 0 | NA | 1 | 0 | orphan | 1 | 1 | complete | 0 | 0 | NA | 88.86 | 0 | 1.03 |
| AG-904-L02 | GORG | 0 | 0 | NA | 2 | 0 | orphan | 0 | 0 | NA | 0 | 0 | NA | 88.81 | 0 | 1.06 |
| AG-896-E11 | GORG | 1 | 1 | complete | 2 | 0 | orphan | 0 | 0 | NA | 0 | 0 | NA | 88.8 | 0 | 1.3 |
| AG-430-G14 | GORG | 0 | 0 | NA | 1 | 0 | orphan | 0 | 0 | NA | 0 | 0 | NA | 88.76 | 0 | 0.98 |
| AG-430-I18 | GORG | 1 | 1 | complete | 1 | 0 | orphan | 0 | 0 | NA | 0 | 0 | NA | 88.76 | 0 | 1.18 |
| AG-447-A02 | GORG | 2 | 1 | complete | 1 | 0 | orphan | 0 | 0 | NA | 0 | 0 | NA | 88.76 | 0 | 1.43 |
| AG-915-K06 | GORG | 0 | 0 | NA | 0 | 0 | NA | 0 | 0 | NA | 0 | 1 | orphan | 88.72 | 0 | 1.43 |
| AG-893-P10 | GORG | 1 | 1 | complete | 1 | 0 | orphan | 0 | 0 | NA | 0 | 0 | NA | 88.65 | 0 | 1.03 |
| AG-901-J22 | GORG | 0 | 0 | NA | 2 | 0 | orphan | 0 | 0 | NA | 0 | 0 | NA | 88.64 | 0 | 1.41 |
| AG-904-A08 | GORG | 0 | 1 | NA | 3 | 0 | orphan | 0 | 0 | NA | 0 | 0 | NA | 88.63 | 0 | 1.08 |
| AG-426-F10 | GORG | 0 | 0 | NA | 1 | 0 | orphan | 0 | 0 | NA | 0 | 0 | NA | 88.61 | 0 | 1.11 |
| AH-321-N11 | GORG | 0 | 0 | NA | 3 | 0 | orphan | 0 | 0 | NA | 0 | 0 | NA | 88.61 | 0 | 1.04 |
| AG-894-G20 | GORG | 0 | 0 | NA | 3 | 0 | orphan | 0 | 0 | NA | 0 | 0 | NA | 88.58 | 0 | 1.84 |
| AG-447-E07 | GORG | 1 | 1 | complete | 1 | 0 | orphan | 0 | 0 | NA | 0 | 0 | NA | 88.55 | 0 | 1.14 |
| AG-894-N13 | GORG | 0 | 0 | NA | 2 | 0 | orphan | 1 | 1 | complete | 0 | 0 | NA | 88.55 | 0 | 1.54 |
| AG-899-C09 | GORG | 0 | 0 | NA | 1 | 0 | orphan | 0 | 0 | NA | 0 | 0 | NA | 88.55 | 0 | 1.21 |
| AG-912-B21 | GORG | 0 | 0 | NA | 1 | 0 | orphan | 0 | 0 | NA | 0 | 0 | NA | 88.55 | 0 | 1.44 |
| AG-900-E18 | GORG | 0 | 0 | NA | 1 | 0 | orphan | 0 | 0 | NA | 0 | 0 | NA | 88.46 | 0 | 1.01 |
| AG-911-C08 | GORG | 0 | 0 | NA | 1 | 0 | orphan | 1 | 1 | complete | 0 | 0 | NA | 88.43 | 0 | 2.29 |
| AG-907-E05 | GORG | 0 | 0 | NA | 1 | 0 | orphan | 0 | 0 | NA | 0 | 0 | NA | 88.38 | 0 | 2.24 |
| AG-457-M14 | GORG | 0 | 0 | NA | 1 | 0 | orphan | 0 | 0 | NA | 0 | 0 | NA | 88.35 | 0 | 1.13 |
| AG-913-A04 | GORG | 0 | 0 | NA | 2 | 1 | complete | 0 | 0 | NA | 0 | 0 | NA | 88.35 | 0 | 1.08 |

| | | | | | | | | | | | | | | | | |
|---|---|---|---|---|---|---|---|---|---|---|---|---|---|---|---|---|
| AH-321-B19 | GORG | 0 | 0 | NA | 1 | 0 | orphan | 0 | 0 | NA | 0 | 1 | orphan | 88.22 | 0 | 1.57 |
| AG-404-C02 | GORG | 0 | 0 | NA | 1 | 0 | orphan | 0 | 0 | NA | 0 | 0 | NA | 88.17 | 0 | 1.1 |
| AG-910-M21 | GORG | 0 | 0 | NA | 1 | 0 | orphan | 0 | 0 | NA | 0 | 0 | NA | 88.1 | 0 | 0.91 |
| AG-447-E19 | GORG | 0 | 0 | NA | 1 | 0 | orphan | 0 | 0 | NA | 0 | 1 | orphan | 88.06 | 0 | 1.19 |
| AG-908-C17 | GORG | 0 | 0 | NA | 1 | 0 | orphan | 0 | 0 | NA | 0 | 0 | NA | 88.06 | 0 | 1.19 |
| AG-349-F22 | GORG | 0 | 0 | NA | 0 | 1 | NA | 0 | 0 | NA | 0 | 0 | NA | 87.95 | 0 | 0.97 |
| AG-359-B05 | GORG | 0 | 0 | NA | 2 | 0 | orphan | 0 | 0 | NA | 0 | 0 | NA | 87.95 | 0 | 1.12 |
| AG-422-K17 | GORG | 0 | 0 | NA | 2 | 0 | orphan | 0 | 0 | NA | 0 | 0 | NA | 87.95 | 0 | 0.96 |
| AG-430-I13 | GORG | 1 | 2 | complete | 1 | 0 | orphan | 0 | 0 | NA | 0 | 0 | NA | 87.95 | 0 | 0.91 |
| AG-891-J18 | GORG | 0 | 0 | NA | 1 | 0 | orphan | 0 | 0 | NA | 0 | 0 | NA | 87.95 | 0 | 1.25 |
| AG-891-P06 | GORG | 0 | 0 | NA | 2 | 0 | orphan | 0 | 0 | NA | 0 | 0 | NA | 87.95 | 0 | 1.31 |
| AG-893-M03 | GORG | 1 | 1 | complete | 0 | 0 | NA | 0 | 0 | NA | 0 | 0 | NA | 87.95 | 0 | 1.3 |
| AG-899-G03 | GORG | 0 | 0 | NA | 1 | 0 | orphan | 0 | 0 | NA | 0 | 0 | NA | 87.95 | 0 | 0.96 |
| AG-903-C11 | GORG | 1 | 1 | complete | 1 | 0 | orphan | 0 | 0 | NA | 0 | 0 | NA | 87.95 | 0 | 1.19 |
| AG-905-B19 | GORG | 0 | 0 | NA | 1 | 0 | orphan | 0 | 0 | NA | 0 | 0 | NA | 87.95 | 0 | 0.96 |
| AG-912-F03 | GORG | 0 | 0 | NA | 1 | 0 | orphan | 0 | 0 | NA | 0 | 0 | NA | 87.95 | 0 | 1.43 |
| AG-916-L09 | GORG | 0 | 0 | NA | 1 | 0 | orphan | 0 | 0 | NA | 0 | 0 | NA | 87.95 | 0 | 0.89 |
| AG-917-D23 | GORG | 0 | 0 | NA | 2 | 0 | orphan | 0 | 0 | NA | 0 | 0 | NA | 87.95 | 0 | 1.23 |
| AH-324-K09 | GORG | 0 | 0 | NA | 1 | 0 | orphan | 0 | 0 | NA | 0 | 0 | NA | 87.95 | 0 | 1.07 |
| AG-337-K10 | GORG | 1 | 1 | complete | 1 | 0 | orphan | 0 | 0 | NA | 0 | 0 | NA | 87.91 | 0 | 0.91 |
| AG-390-J21 | GORG | 0 | 0 | NA | 4 | 1 | complete | 0 | 0 | NA | 0 | 0 | NA | 87.91 | 0 | 1.1 |
| AG-894-P05 | GORG | 0 | 0 | NA | 2 | 0 | orphan | 0 | 0 | NA | 0 | 0 | NA | 87.91 | 0 | 0.91 |
| AH-321-D15 | GORG | 0 | 0 | NA | 2 | 0 | orphan | 0 | 0 | NA | 0 | 0 | NA | 87.91 | 0 | 1.29 |
| AG-916-I17 | GORG | 0 | 0 | NA | 1 | 0 | orphan | 0 | 0 | NA | 0 | 0 | NA | 87.88 | 0 | 1.01 |
| AG-895-C20 | GORG | 0 | 0 | NA | 1 | 0 | orphan | 0 | 0 | NA | 0 | 0 | NA | 87.87 | 0 | 1.44 |

| | | | | | | | | | | | | | | | | |
|---|---|---|---|---|---|---|---|---|---|---|---|---|---|---|---|---|
| AG-894-I05 | GORG | 1 | 1 | complet e | 1 | 0 | orphan | 0 | 0 | NA | 0 | 0 | NA | 87.84 | 0 | 0.95 |
| AG-899-B09 | GORG | 0 | 0 | NA | 2 | 0 | orphan | 0 | 0 | NA | 0 | 0 | NA | 87.84 | 0 | 1.22 |
| AG-422-G03 | GORG | 0 | 1 | NA | 1 | 0 | orphan | 0 | 0 | NA | 0 | 0 | NA | 87.76 | 0 | 1.17 |
| AG-899-I09 | GORG | 1 | 1 | complet e | 1 | 0 | orphan | 0 | 0 | NA | 0 | 0 | NA | 87.76 | 0 | 1.27 |
| AG-359-G17 | GORG | 0 | 0 | NA | 1 | 0 | orphan | 0 | 0 | NA | 0 | 0 | NA | 87.73 | 0 | 1.16 |
| AG-453-G04 | GORG | 0 | 0 | NA | 1 | 0 | orphan | 0 | 0 | NA | 0 | 0 | NA | 87.73 | 0 | 1.02 |
| AH-321-K15 | GORG | 0 | 0 | NA | 1 | 0 | orphan | 0 | 0 | NA | 0 | 0 | NA | 87.6 | 0 | 1.03 |
| AG-461-B13 | GORG | 0 | 0 | NA | 1 | 0 | orphan | 0 | 0 | NA | 0 | 0 | NA | 87.58 | 0 | 1.07 |
| AG-414-B15 | GORG | 0 | 0 | NA | 1 | 0 | orphan | 0 | 0 | NA | 0 | 0 | NA | 87.55 | 0 | 0.97 |
| AG-917-M15 | GORG | 0 | 0 | NA | 1 | 0 | orphan | 0 | 0 | NA | 0 | 0 | NA | 87.55 | 0 | 1.06 |
| AG-903-M21 | GORG | 0 | 0 | NA | 1 | 0 | orphan | 0 | 0 | NA | 0 | 0 | NA | 87.51 | 0 | 1 |
| AG-404-F06 | GORG | 0 | 0 | NA | 1 | 0 | orphan | 0 | 0 | NA | 0 | 0 | NA | 87.5 | 0 | 1.73 |
| AG-390-N04 | GORG | 0 | 0 | NA | 1 | 0 | orphan | 0 | 0 | NA | 0 | 0 | NA | 87.43 | 0 | 1.34 |
| AG-899-C14 | GORG | 0 | 0 | NA | 1 | 0 | orphan | 0 | 0 | NA | 0 | 0 | NA | 87.43 | 0 | 1.25 |
| AG-920-I23 | GORG | 0 | 0 | NA | 1 | 0 | orphan | 0 | 0 | NA | 0 | 0 | NA | 87.4 | 0 | 1.1 |
| AG-404-P20 | GORG | 0 | 0 | NA | 1 | 1 | orphan | 0 | 0 | NA | 0 | 0 | NA | 87.35 | 0 | 1.06 |
| AG-892-I02 | GORG | 0 | 0 | NA | 1 | 0 | orphan | 0 | 0 | NA | 0 | 0 | NA | 87.35 | 0 | 1.47 |
| AG-899-M17 | GORG | 0 | 0 | NA | 2 | 0 | orphan | 0 | 0 | NA | 0 | 0 | NA | 87.35 | 0 | 1.41 |
| AG-905-O21 | GORG | 0 | 0 | NA | 1 | 0 | orphan | 0 | 0 | NA | 0 | 0 | NA | 87.35 | 0 | 1.36 |
| AH-287-C13 | GORG | 0 | 0 | NA | 1 | 0 | orphan | 0 | 0 | NA | 0 | 0 | NA | 87.35 | 0 | 1.03 |
| AG-414-P09 | GORG | 0 | 0 | NA | 2 | 0 | orphan | 0 | 1 | NA | 0 | 0 | NA | 87.3 | 0 | 0.93 |
| AG-899-M19 | GORG | 1 | 1 | complet e | 0 | 0 | NA | 0 | 0 | NA | 0 | 0 | NA | 87.3 | 0 | 1.11 |
| AG-898-P13 | GORG | 0 | 0 | NA | 1 | 0 | orphan | 0 | 0 | NA | 0 | 0 | NA | 87.26 | 0 | 1.04 |
| AG-447-M10 | GORG | 0 | 0 | NA | 1 | 0 | orphan | 0 | 0 | NA | 0 | 0 | NA | 87.25 | 0 | 1.13 |
| AG-410-K05 | GORG | 0 | 0 | NA | 1 | 0 | orphan | 0 | 0 | NA | 0 | 0 | NA | 87.22 | 0 | 1.09 |
| AG-426-C04 | GORG | 0 | 0 | NA | 2 | 0 | orphan | 0 | 0 | NA | 0 | 0 | NA | 87.21 | 0 | 1.04 |

| | | | | | | | | | | | | | | | | |
|---|---|---|---|---|---|---|---|---|---|---|---|---|---|---|---|---|
| AG-325-L02 | GORG | 0 | 0 | NA | 1 | 0 | orphan | 0 | 0 | NA | 0 | 0 | NA | 87.2 | 0 | 1.04 |
| AG-920-C09 | GORG | 0 | 0 | NA | 1 | 0 | orphan | 0 | 0 | NA | 0 | 0 | NA | 87.12 | 0 | 0.93 |
| AG-404-J20 | GORG | 0 | 0 | NA | 1 | 0 | orphan | 0 | 0 | NA | 0 | 0 | NA | 87.1 | 0 | 1.07 |
| AG-898-C07 | GORG | 0 | 0 | NA | 1 | 0 | orphan | 0 | 0 | NA | 0 | 0 | NA | 86.97 | 0 | 1.13 |
| AG-900-M23 | GORG | 0 | 0 | NA | 1 | 0 | orphan | 0 | 0 | NA | 0 | 0 | NA | 86.97 | 0 | 1.07 |
| AG-920-I07 | GORG | 0 | 0 | NA | 2 | 0 | orphan | 0 | 0 | NA | 0 | 0 | NA | 86.97 | 0 | 0.96 |
| AG-920-J16 | GORG | 0 | 0 | NA | 0 | 0 | NA | 0 | 0 | NA | 0 | 1 | orphan | 86.96 | 0 | 1.44 |
| AG-404-P01 | GORG | 0 | 0 | NA | 1 | 0 | orphan | 0 | 0 | NA | 0 | 0 | NA | 86.95 | 0 | 1.09 |
| AG-390-I07 | GORG | 0 | 0 | NA | 2 | 0 | orphan | 0 | 0 | NA | 0 | 0 | NA | 86.91 | 0 | 1.19 |
| AG-904-G21 | GORG | 0 | 0 | NA | 1 | 0 | orphan | 0 | 0 | NA | 0 | 0 | NA | 86.91 | 0 | 1.23 |
| AG-426-B11 | GORG | 1 | 1 | complete | 2 | 0 | orphan | 0 | 0 | NA | 0 | 0 | NA | 86.9 | 0 | 1 |
| AG-439-G13 | GORG | 0 | 0 | NA | 1 | 0 | orphan | 0 | 0 | NA | 0 | 0 | NA | 86.88 | 0 | 1.18 |
| AG-447-K11 | GORG | 0 | 0 | NA | 1 | 0 | orphan | 0 | 0 | NA | 0 | 0 | NA | 86.81 | 0 | 1.21 |
| AG-916-N03 | GORG | 0 | 0 | NA | 2 | 0 | orphan | 0 | 0 | NA | 0 | 0 | NA | 86.81 | 0 | 0.92 |
| AG-891-G23 | GORG | 1 | 1 | complete | 2 | 0 | orphan | 0 | 0 | NA | 0 | 0 | NA | 86.8 | 0 | 1.35 |
| AG-901-B17 | GORG | 1 | 0 | orphan | 0 | 0 | NA | 0 | 0 | NA | 0 | 0 | NA | 86.77 | 0 | 0.99 |
| AG-359-O11 | GORG | 0 | 0 | NA | 1 | 0 | orphan | 0 | 0 | NA | 0 | 1 | orphan | 86.75 | 0 | 1.14 |
| AG-319-D07 | GORG | 0 | 0 | NA | 3 | 1 | complete | 0 | 0 | NA | 0 | 0 | NA | 86.75 | 0 | 1.33 |
| AG-343-E17 | GORG | 0 | 0 | NA | 1 | 0 | orphan | 0 | 0 | NA | 0 | 0 | NA | 86.75 | 0 | 0.99 |
| AG-422-F15 | GORG | 0 | 0 | NA | 1 | 0 | orphan | 0 | 0 | NA | 0 | 0 | NA | 86.75 | 0 | 0.99 |
| AG-893-O06 | GORG | 0 | 0 | NA | 1 | 0 | orphan | 0 | 0 | NA | 0 | 0 | NA | 86.75 | 0 | 1.4 |
| AG-895-E15 | GORG | 0 | 0 | NA | 1 | 0 | orphan | 0 | 0 | NA | 0 | 0 | NA | 86.75 | 0 | 0.96 |
| AG-896-G23 | GORG | 0 | 0 | NA | 1 | 0 | orphan | 0 | 0 | NA | 0 | 0 | NA | 86.75 | 0 | 1.28 |
| AG-898-G21 | GORG | 0 | 0 | NA | 1 | 1 | complete | 0 | 0 | NA | 0 | 0 | NA | 86.75 | 0 | 1.15 |
| AG-907-M07 | GORG | 0 | 0 | NA | 2 | 0 | orphan | 0 | 0 | NA | 0 | 0 | NA | 86.75 | 0 | 1.08 |

| | | | | | | | | | | | | | | | | |
|---|---|---|---|---|---|---|---|---|---|---|---|---|---|---|---|---|
| AG-908-I21 | GORG | 0 | 0 | NA | 2 | 0 | orphan | 0 | 0 | NA | 0 | 0 | NA | 86.75 | 0 | 1.12 |
| AG-909-N22 | GORG | 0 | 0 | NA | 1 | 0 | orphan | 0 | 0 | NA | 0 | 0 | NA | 86.75 | 0 | 0.91 |
| AG-912-M06 | GORG | 0 | 0 | NA | 1 | 0 | orphan | 0 | 0 | NA | 0 | 0 | NA | 86.75 | 0 | 1.09 |
| AG-914-M14 | GORG | 0 | 0 | NA | 2 | 1 | complete | 0 | 0 | NA | 0 | 0 | NA | 86.75 | 0 | 1.01 |
| AG-915-C07 | GORG | 0 | 0 | NA | 2 | 0 | orphan | 0 | 0 | NA | 0 | 0 | NA | 86.75 | 0 | 1.01 |
| AG-916-M11 | GORG | 0 | 0 | NA | 3 | 0 | orphan | 0 | 0 | NA | 0 | 0 | NA | 86.75 | 0 | 0.97 |
| AG-917-E14 | GORG | 0 | 0 | NA | 1 | 0 | orphan | 0 | 0 | NA | 0 | 0 | NA | 86.75 | 0 | 0.9 |
| AH-324-B10 | GORG | 0 | 0 | NA | 1 | 0 | orphan | 0 | 1 | NA | 0 | 0 | NA | 86.75 | 0 | 1 |
| AH-287-J18 | GORG | 0 | 0 | NA | 1 | 0 | orphan | 0 | 0 | NA | 0 | 0 | NA | 86.73 | 0 | 1.03 |
| AG-896-N14 | GORG | 0 | 0 | NA | 0 | 0 | NA | 0 | 1 | NA | 0 | 0 | NA | 86.71 | 0 | 1.89 |
| AG-918-P10 | GORG | 0 | 0 | NA | 1 | 0 | orphan | 0 | 0 | NA | 0 | 0 | NA | 86.71 | 0 | 1.17 |
| AG-337-K17 | GORG | 0 | 0 | NA | 2 | 0 | orphan | 2 | 1 | complete | 0 | 0 | NA | 86.7 | 0 | 1.67 |
| AH-324-K11 | GORG | 0 | 0 | NA | 2 | 0 | orphan | 0 | 0 | NA | 0 | 0 | NA | 86.65 | 0 | 1.07 |
| AG-891-G06 | GORG | 0 | 0 | NA | 3 | 0 | orphan | 0 | 0 | NA | 0 | 0 | NA | 86.58 | 0 | 1.35 |
| AG-470-D02 | GORG | 0 | 0 | NA | 1 | 0 | orphan | 0 | 0 | NA | 0 | 0 | NA | 86.57 | 0 | 1.03 |
| AG-349-C11 | GORG | 0 | 0 | NA | 1 | 0 | orphan | 0 | 0 | NA | 0 | 0 | NA | 86.56 | 0 | 0.95 |
| AG-439-O05 | GORG | 0 | 0 | NA | 1 | 0 | orphan | 0 | 0 | NA | 0 | 0 | NA | 86.53 | 0 | 0.96 |
| AH-324-M21 | GORG | 0 | 0 | NA | 1 | 0 | orphan | 1 | 1 | orphan | 0 | 0 | NA | 86.49 | 0 | 1.58 |
| AG-917-N20 | GORG | 0 | 0 | NA | 2 | 0 | orphan | 0 | 0 | NA | 0 | 0 | NA | 86.47 | 0 | 1.34 |
| AH-326-M23 | GORG | 0 | 0 | NA | 1 | 0 | orphan | 0 | 0 | NA | 0 | 0 | NA | 86.42 | 0 | 0.84 |
| AG-404-O18 | GORG | 0 | 0 | NA | 2 | 0 | orphan | 0 | 0 | NA | 0 | 0 | NA | 86.35 | 0 | 1.02 |
| AG-337-C14 | GORG | 0 | 0 | NA | 1 | 0 | orphan | 0 | 0 | NA | 0 | 0 | NA | 86.31 | 0 | 1.12 |
| AG-895-G15 | GORG | 0 | 0 | NA | 2 | 0 | orphan | 0 | 0 | NA | 0 | 0 | NA | 86.25 | 0 | 1.08 |
| AG-422-C13 | GORG | 0 | 0 | NA | 1 | 0 | orphan | 0 | 0 | NA | 0 | 0 | NA | 86.18 | 0 | 1.01 |
| AG-894-P23 | GORG | 0 | 0 | NA | 2 | 0 | orphan | 0 | 0 | NA | 0 | 0 | NA | 86.18 | 0 | 1.18 |
| AG-894-O02 | GORG | 0 | 0 | NA | 1 | 0 | orphan | 0 | 0 | NA | 0 | 0 | NA | 86.14 | 0 | 1.19 |

| | | | | | | | | | | | | | | | | |
|---|---|---|---|---|---|---|---|---|---|---|---|---|---|---|---|---|
| AG-895-E07 | GORG | 0 | 0 | NA | 1 | 0 | orphan | 0 | 0 | NA | 0 | 0 | NA | 86.14 | 0 | 1.19 |
| AG-899-A06 | GORG | 0 | 0 | NA | 1 | 0 | orphan | 0 | 0 | NA | 0 | 0 | NA | 86.14 | 0 | 1.22 |
| AG-912-D17 | GORG | 0 | 0 | NA | 2 | 0 | orphan | 0 | 0 | NA | 0 | 0 | NA | 86.14 | 0 | 0.95 |
| AG-916-M13 | GORG | 1 | 1 | complete | 1 | 0 | orphan | 0 | 0 | NA | 0 | 0 | NA | 86.14 | 0 | 1.1 |
| AG-918-E06 | GORG | 0 | 0 | NA | 1 | 0 | orphan | 0 | 0 | NA | 0 | 0 | NA | 86.14 | 0 | 0.97 |
| AG-919-C22 | GORG | 0 | 0 | NA | 2 | 0 | orphan | 0 | 0 | NA | 0 | 0 | NA | 86.14 | 0 | 0.86 |
| AG-907-C04 | GORG | 0 | 0 | NA | 1 | 0 | orphan | 0 | 0 | NA | 0 | 0 | NA | 86.1 | 0 | 1.13 |
| AG-426-C16 | GORG | 0 | 0 | NA | 1 | 0 | orphan | 0 | 0 | NA | 0 | 0 | NA | 86.09 | 0 | 1.14 |
| AG-390-M15 | GORG | 0 | 0 | NA | 2 | 0 | orphan | 0 | 0 | NA | 0 | 0 | NA | 86.08 | 0 | 0.96 |
| AG-894-K14 | GORG | 0 | 0 | NA | 1 | 0 | orphan | 0 | 0 | NA | 0 | 0 | NA | 86.06 | 0 | 1.06 |
| AG-894-B21 | GORG | 0 | 0 | NA | 0 | 0 | NA | 0 | 0 | NA | 0 | 1 | orphan | 86.05 | 0 | 1.39 |
| AG-430-J03 | GORG | 0 | 0 | NA | 1 | 0 | orphan | 0 | 0 | NA | 0 | 0 | NA | 86.03 | 0 | 1.27 |
| AG-892-G02 | GORG | 0 | 0 | NA | 1 | 0 | orphan | 0 | 0 | NA | 0 | 0 | NA | 86.03 | 0 | 1 |
| AG-901-D23 | GORG | 0 | 0 | NA | 1 | 0 | orphan | 1 | 1 | complete | 0 | 0 | NA | 86.02 | 0 | 1.44 |
| AG-337-D08 | GORG | 0 | 0 | NA | 1 | 0 | orphan | 0 | 0 | NA | 0 | 0 | NA | 85.97 | 0 | 1.43 |
| AG-908-F20 | GORG | 0 | 0 | NA | 1 | 0 | orphan | 0 | 0 | NA | 0 | 0 | NA | 85.97 | 0 | 1.06 |
| AG-447-O14 | GORG | 0 | 0 | NA | 1 | 0 | orphan | 0 | 0 | NA | 0 | 0 | NA | 85.94 | 0 | 1.12 |
| AG-894-L17 | GORG | 0 | 0 | NA | 1 | 0 | orphan | 0 | 0 | NA | 0 | 0 | NA | 85.94 | 0 | 1.27 |
| AG-908-I10 | GORG | 0 | 0 | NA | 2 | 0 | orphan | 0 | 0 | NA | 0 | 0 | NA | 85.89 | 0 | 1.02 |
| AG-911-D18 | GORG | 0 | 0 | NA | 1 | 0 | orphan | 0 | 0 | NA | 0 | 1 | orphan | 85.78 | 0 | 1.01 |
| AG-404-B04 | GORG | 0 | 0 | NA | 1 | 0 | orphan | 0 | 0 | NA | 0 | 0 | NA | 85.76 | 0 | 1.07 |
| AG-426-F20 | GORG | 0 | 0 | NA | 3 | 0 | orphan | 0 | 0 | NA | 0 | 0 | NA | 85.75 | 0 | 1.34 |
| AH-287-L13 | GORG | 1 | 1 | complete | 1 | 0 | orphan | 0 | 0 | NA | 0 | 0 | NA | 85.75 | 0 | 2.01 |
| AG-893-O16 | GORG | 0 | 0 | NA | 2 | 1 | complete | 0 | 0 | NA | 0 | 0 | NA | 85.68 | 0 | 1.17 |
| AG-325-I09 | GORG | 0 | 0 | NA | 1 | 0 | orphan | 0 | 0 | NA | 0 | 0 | NA | 85.66 | 0 | 1.37 |

| | | | | | | | | | | | | | | | | |
|---|---|---|---|---|---|---|---|---|---|---|---|---|---|---|---|---|
| AG-914-L20 | GORG | 0 | 0 | NA | 2 | 0 | orphan | 0 | 0 | NA | 0 | 0 | NA | 85.66 | 0 | 1.17 |
| AG-313-C04 | GORG | 0 | 0 | NA | 1 | 0 | orphan | 0 | 0 | NA | 0 | 0 | NA | 85.6 | 0 | 1.52 |
| AG-917-I17 | GORG | 1 | 1 | complete | 2 | 0 | orphan | 0 | 0 | NA | 0 | 0 | NA | 85.58 | 0 | 1.18 |
| AG-899-C10 | GORG | 0 | 0 | NA | 0 | 0 | NA | 1 | 1 | complete | 0 | 0 | NA | 85.56 | 0 | 2.11 |
| AG-903-K22 | GORG | 1 | 1 | complete | 0 | 0 | NA | 0 | 0 | NA | 0 | 0 | NA | 85.56 | 0 | 1.07 |
| AG-422-I21 | GORG | 0 | 2 | NA | 0 | 0 | NA | 0 | 0 | NA | 0 | 1 | orphan | 85.55 | 0 | 1.48 |
| AG-891-G02 | GORG | 0 | 0 | NA | 1 | 0 | orphan | 0 | 0 | NA | 0 | 0 | NA | 85.55 | 0 | 1.27 |
| AG-893-M20 | GORG | 1 | 0 | orphan | 1 | 0 | orphan | 0 | 0 | NA | 0 | 1 | orphan | 85.54 | 0 | 1.47 |
| AG-325-P23 | GORG | 0 | 0 | NA | 1 | 0 | orphan | 0 | 0 | NA | 0 | 0 | NA | 85.54 | 0 | 1.05 |
| AG-390-I22 | GORG | 0 | 0 | NA | 1 | 0 | orphan | 0 | 0 | NA | 0 | 0 | NA | 85.54 | 0 | 0.96 |
| AG-422-N03 | GORG | 0 | 0 | NA | 1 | 0 | orphan | 0 | 0 | NA | 0 | 0 | NA | 85.54 | 0 | 1.3 |
| AG-464-G03 | GORG | 0 | 0 | NA | 1 | 0 | orphan | 0 | 0 | NA | 0 | 0 | NA | 85.54 | 0 | 1.51 |
| AG-893-N07 | GORG | 0 | 0 | NA | 1 | 0 | orphan | 0 | 0 | NA | 0 | 0 | NA | 85.54 | 0 | 1.26 |
| AG-896-D15 | GORG | 0 | 0 | NA | 1 | 0 | orphan | 0 | 0 | NA | 0 | 0 | NA | 85.54 | 0 | 1.07 |
| AG-901-G03 | GORG | 1 | 1 | complete | 1 | 0 | orphan | 0 | 0 | NA | 0 | 0 | NA | 85.54 | 0 | 1.19 |
| AG-913-N09 | GORG | 1 | 1 | complete | 1 | 0 | orphan | 0 | 0 | NA | 0 | 0 | NA | 85.54 | 0 | 1.15 |
| AG-914-P22 | GORG | 0 | 0 | NA | 1 | 0 | orphan | 0 | 0 | NA | 0 | 0 | NA | 85.54 | 0 | 1.11 |
| AG-916-B11 | GORG | 0 | 0 | NA | 2 | 0 | orphan | 0 | 0 | NA | 0 | 0 | NA | 85.54 | 0 | 0.93 |
| AG-919-K20 | GORG | 0 | 0 | NA | 2 | 0 | orphan | 0 | 0 | NA | 0 | 0 | NA | 85.54 | 0 | 0.9 |
| AH-321-L23 | GORG | 1 | 0 | orphan | 2 | 0 | orphan | 0 | 0 | NA | 0 | 0 | NA | 85.49 | 0 | 1.03 |
| AG-365-C14 | GORG | 0 | 0 | NA | 1 | 0 | orphan | 0 | 0 | NA | 0 | 0 | NA | 85.48 | 0 | 1.08 |
| AG-349-M19 | GORG | 0 | 0 | NA | 1 | 0 | orphan | 0 | 0 | NA | 0 | 0 | NA | 85.47 | 0 | 1 |
| AG-910-G11 | GORG | 0 | 0 | NA | 2 | 0 | orphan | 0 | 0 | NA | 0 | 0 | NA | 85.47 | 0 | 0.94 |
| AG-359-B04 | GORG | 0 | 0 | NA | 1 | 0 | orphan | 0 | 0 | NA | 0 | 1 | orphan | 85.41 | 0 | 0.92 |

| | | | | | | | | | | | | | | | | |
|---|---|---|---|---|---|---|---|---|---|---|---|---|---|---|---|---|
| AG-901-I06 | GORG | 0 | 0 | NA | 1 | 0 | orphan | 0 | 0 | NA | 0 | 0 | NA | 85.4 | 0 | 1.28 |
| AG-404-B06 | GORG | 0 | 0 | NA | 4 | 0 | orphan | 0 | 0 | NA | 0 | 0 | NA | 85.31 | 0 | 1.09 |
| AG-426-M05 | GORG | 0 | 0 | NA | 1 | 0 | orphan | 0 | 0 | NA | 0 | 0 | NA | 85.31 | 0 | 1.22 |
| AG-903-O08 | GORG | 0 | 0 | NA | 1 | 0 | orphan | 0 | 0 | NA | 0 | 0 | NA | 85.31 | 0 | 1.07 |
| AG-917-M09 | GORG | 0 | 0 | NA | 0 | 0 | NA | 0 | 0 | NA | 0 | 2 | orphan | 85.29 | 0 | 1.72 |
| AG-439-B12 | GORG | 1 | 1 | complete | 2 | 0 | orphan | 0 | 0 | NA | 0 | 0 | NA | 85.25 | 0 | 1.14 |
| AG-918-C08 | GORG | 0 | 0 | NA | 1 | 0 | orphan | 0 | 0 | NA | 0 | 0 | NA | 85.25 | 0 | 0.99 |
| AG-894-O05 | GORG | 0 | 0 | NA | 0 | 0 | NA | 0 | 0 | NA | 0 | 1 | orphan | 85.18 | 0 | 1.45 |
| AG-894-I22 | GORG | 1 | 1 | complete | 1 | 0 | orphan | 0 | 0 | NA | 0 | 0 | NA | 85.16 | 0 | 1.25 |
| AG-896-L20 | GORG | 0 | 0 | NA | 2 | 0 | orphan | 0 | 0 | NA | 0 | 2 | orphan | 85.12 | 0 | 1.08 |
| AG-916-E17 | GORG | 0 | 0 | NA | 1 | 0 | orphan | 0 | 0 | NA | 0 | 0 | NA | 85.1 | 0 | 0.99 |
| AG-893-N22 | GORG | 0 | 0 | NA | 0 | 0 | NA | 2 | 1 | complete | 0 | 0 | NA | 85.07 | 0 | 1.08 |
| AG-913-K21 | GORG | 0 | 0 | NA | 1 | 0 | orphan | 0 | 0 | NA | 0 | 0 | NA | 85.07 | 0 | 1.1 |
| AG-333-K06 | GORG | 0 | 0 | NA | 1 | 0 | orphan | 0 | 0 | NA | 0 | 0 | NA | 84.94 | 0 | 1.01 |
| AG-422-B23 | GORG | 0 | 0 | NA | 1 | 0 | orphan | 0 | 0 | NA | 0 | 0 | NA | 84.94 | 0 | 0.9 |
| AG-891-G05 | GORG | 1 | 1 | complete | 1 | 0 | orphan | 0 | 0 | NA | 0 | 0 | NA | 84.94 | 0 | 1.24 |
| AG-894-C10 | GORG | 0 | 0 | NA | 2 | 0 | orphan | 0 | 0 | NA | 0 | 0 | NA | 84.94 | 0 | 1.21 |
| AG-907-B04 | GORG | 0 | 0 | NA | 1 | 0 | orphan | 0 | 0 | NA | 0 | 0 | NA | 84.94 | 0 | 1.12 |
| AG-914-B16 | GORG | 0 | 0 | NA | 1 | 0 | orphan | 0 | 0 | NA | 0 | 0 | NA | 84.94 | 0 | 1.12 |
| AG-915-K07 | GORG | 0 | 0 | NA | 2 | 0 | orphan | 0 | 0 | NA | 0 | 0 | NA | 84.94 | 0 | 1.18 |
| AG-918-N11 | GORG | 0 | 0 | NA | 1 | 0 | orphan | 0 | 0 | NA | 0 | 0 | NA | 84.94 | 0 | 1.28 |
| AG-920-K17 | GORG | 1 | 0 | orphan | 1 | 0 | orphan | 0 | 0 | NA | 0 | 0 | NA | 84.94 | 0 | 1.23 |
| AG-891-N17 | GORG | 0 | 0 | NA | 1 | 0 | orphan | 0 | 0 | NA | 0 | 0 | NA | 84.93 | 0 | 1.15 |
| AG-911-E05 | GORG | 0 | 0 | NA | 2 | 0 | orphan | 0 | 0 | NA | 0 | 0 | NA | 84.76 | 0 | 0.96 |
| AG-404-C23 | GORG | 0 | 0 | NA | 1 | 0 | orphan | 0 | 0 | NA | 0 | 0 | NA | 84.74 | 0 | 0.97 |

| | | | | | | | | | | | | | | | | |
|---|---|---|---|---|---|---|---|---|---|---|---|---|---|---|---|---|
| AG-917-N11 | GORG | 0 | 0 | NA | 2 | 0 | orphan | 0 | 0 | NA | 0 | 0 | NA | 84.74 | 0 | 1.29 |
| AG-422-J20 | GORG | 0 | 0 | NA | 2 | 0 | orphan | 0 | 0 | NA | 0 | 0 | NA | 84.69 | 0 | 0.99 |
| AG-430-K19 | GORG | 0 | 0 | NA | 1 | 0 | orphan | 0 | 0 | NA | 0 | 0 | NA | 84.68 | 0 | 1.04 |
| AG-893-L06 | GORG | 0 | 0 | NA | 1 | 0 | orphan | 0 | 0 | NA | 0 | 0 | NA | 84.6 | 0 | 1.02 |
| AG-911-B04 | GORG | 1 | 0 | orphan | 2 | 1 | complete | 0 | 0 | NA | 0 | 0 | NA | 84.6 | 0 | 0.92 |
| AG-899-P08 | GORG | 1 | 1 | complete | 0 | 0 | NA | 0 | 0 | NA | 0 | 2 | orphan | 84.59 | 0 | 1.39 |
| AG-430-L17 | GORG | 0 | 0 | NA | 1 | 0 | orphan | 0 | 0 | NA | 0 | 0 | NA | 84.56 | 0 | 1.14 |
| AH-321-D13 | GORG | 1 | 1 | complete | 2 | 0 | orphan | 0 | 0 | NA | 0 | 0 | NA | 84.56 | 0 | 0.99 |
| AG-422-J03 | GORG | 1 | 1 | orphan | 1 | 0 | orphan | 0 | 0 | NA | 0 | 0 | NA | 84.54 | 0 | 1.05 |
| AG-907-B09 | GORG | 0 | 0 | NA | 3 | 0 | orphan | 0 | 0 | NA | 0 | 0 | NA | 84.52 | 0 | 0.99 |
| AG-349-C13 | GORG | 0 | 0 | NA | 0 | 0 | NA | 0 | 0 | NA | 0 | 1 | orphan | 84.51 | 0 | 1.52 |
| AG-911-L11 | GORG | 0 | 0 | NA | 2 | 0 | orphan | 0 | 0 | NA | 0 | 0 | NA | 84.43 | 0 | 1.09 |
| AG-919-D03 | GORG | 1 | 1 | complete | 1 | 0 | orphan | 0 | 0 | NA | 0 | 0 | NA | 84.41 | 0 | 1.65 |
| AG-901-E05 | GORG | 0 | 0 | NA | 1 | 0 | orphan | 0 | 0 | NA | 0 | 0 | NA | 84.38 | 0 | 1.71 |
| AG-914-F07 | GORG | 0 | 0 | NA | 0 | 0 | NA | 0 | 0 | NA | 0 | 1 | orphan | 84.36 | 0 | 1.32 |
| AG-447-O23 | GORG | 0 | 0 | NA | 1 | 0 | orphan | 0 | 0 | NA | 0 | 0 | NA | 84.36 | 0 | 1.07 |
| AG-909-J05 | GORG | 0 | 0 | NA | 1 | 0 | orphan | 0 | 0 | NA | 0 | 0 | NA | 84.36 | 0 | 1.03 |
| AG-337-E19 | GORG | 0 | 0 | NA | 3 | 0 | orphan | 0 | 0 | NA | 0 | 0 | NA | 84.34 | 0 | 1.12 |
| AG-337-F02 | GORG | 0 | 0 | NA | 2 | 0 | orphan | 0 | 0 | NA | 0 | 0 | NA | 84.34 | 0 | 1.38 |
| AG-414-E06 | GORG | 0 | 0 | NA | 1 | 0 | orphan | 0 | 0 | NA | 0 | 0 | NA | 84.34 | 0 | 1.35 |
| AG-430-J04 | GORG | 0 | 0 | NA | 2 | 0 | orphan | 0 | 0 | NA | 0 | 0 | NA | 84.34 | 0 | 1 |
| AG-457-A08 | GORG | 0 | 0 | NA | 1 | 0 | orphan | 0 | 0 | NA | 0 | 0 | NA | 84.34 | 0 | 1.42 |
| AG-904-K16 | GORG | 0 | 0 | NA | 2 | 0 | orphan | 0 | 0 | NA | 0 | 0 | NA | 84.34 | 0 | 1.25 |
| AG-910-M23 | GORG | 0 | 0 | NA | 3 | 0 | orphan | 0 | 0 | NA | 0 | 0 | NA | 84.34 | 0 | 1.02 |
| AG-914-C06 | GORG | 0 | 0 | NA | 1 | 0 | orphan | 0 | 0 | NA | 0 | 0 | NA | 84.34 | 0 | 1.18 |

| | | | | | | | | | | | | | | | | |
|---|---|---|---|---|---|---|---|---|---|---|---|---|---|---|---|---|
| AG-916-B06 | GORG | 0 | 0 | NA | 2 | 0 | orphan | 0 | 0 | NA | 0 | 0 | NA | 84.34 | 0 | 1.18 |
| AG-911-I17 | GORG | 0 | 0 | NA | 2 | 0 | orphan | 0 | 0 | NA | 0 | 0 | NA | 84.17 | 0 | 1.89 |
| AG-891-M18 | GORG | 0 | 0 | NA | 2 | 0 | orphan | 0 | 0 | NA | 0 | 0 | NA | 84.11 | 0 | 1.01 |
| AG-919-C15 | GORG | 1 | 1 | complete | 0 | 0 | NA | 0 | 0 | NA | 0 | 0 | NA | 84.1 | 0 | 1.39 |
| AG-447-I09 | GORG | 0 | 0 | NA | 1 | 0 | orphan | 0 | 0 | NA | 0 | 0 | NA | 84.08 | 0 | 1.12 |
| AG-920-K11 | GORG | 0 | 0 | NA | 1 | 0 | orphan | 0 | 0 | NA | 0 | 0 | NA | 84.04 | 0 | 0.99 |
| AG-916-M06 | GORG | 0 | 0 | NA | 2 | 0 | orphan | 0 | 0 | NA | 0 | 0 | NA | 84.03 | 0 | 1.07 |
| AG-430-K18 | GORG | 0 | 0 | NA | 1 | 0 | orphan | 0 | 0 | NA | 0 | 0 | NA | 83.94 | 0 | 0.96 |
| AG-900-G22 | GORG | 0 | 0 | NA | 3 | 0 | orphan | 0 | 0 | NA | 0 | 0 | NA | 83.91 | 0 | 1.13 |
| AG-910-E10 | GORG | 1 | 1 | complete | 0 | 0 | NA | 0 | 0 | NA | 0 | 0 | NA | 83.87 | 0 | 1.25 |
| AG-422-G06 | GORG | 0 | 0 | NA | 1 | 0 | orphan | 0 | 0 | NA | 0 | 0 | NA | 83.82 | 0 | 1.77 |
| AG-422-E09 | GORG | 0 | 0 | NA | 2 | 0 | orphan | 0 | 0 | NA | 0 | 0 | NA | 83.8 | 0 | 1.15 |
| AG-439-D18 | GORG | 1 | 1 | complete | 3 | 0 | orphan | 0 | 0 | NA | 0 | 0 | NA | 83.76 | 0 | 1.89 |
| AG-897-P03 | GORG | 0 | 0 | NA | 0 | 0 | NA | 0 | 0 | NA | 0 | 1 | orphan | 83.74 | 0 | 1.33 |
| AG-430-A02 | GORG | 0 | 0 | NA | 1 | 0 | orphan | 0 | 0 | NA | 0 | 0 | NA | 83.73 | 0 | 0.89 |
| AG-439-E23 | GORG | 0 | 0 | NA | 1 | 0 | orphan | 0 | 0 | NA | 0 | 0 | NA | 83.73 | 0 | 1.11 |
| AG-891-K19 | GORG | 0 | 0 | NA | 1 | 0 | orphan | 0 | 0 | NA | 0 | 0 | NA | 83.73 | 0 | 1.39 |
| AG-894-K20 | GORG | 0 | 0 | NA | 1 | 0 | orphan | 0 | 0 | NA | 0 | 0 | NA | 83.73 | 0 | 1.27 |
| AG-895-A09 | GORG | 0 | 0 | NA | 1 | 0 | orphan | 0 | 0 | NA | 0 | 0 | NA | 83.73 | 0 | 1.27 |
| AG-913-M07 | GORG | 0 | 0 | NA | 2 | 0 | orphan | 0 | 0 | NA | 0 | 0 | NA | 83.73 | 0 | 1.5 |
| AH-324-F14 | GORG | 0 | 0 | NA | 1 | 0 | orphan | 0 | 0 | NA | 0 | 0 | NA | 83.72 | 0 | 1.36 |
| AG-916-G05 | GORG | 0 | 0 | NA | 3 | 0 | orphan | 0 | 0 | NA | 0 | 0 | NA | 83.68 | 0 | 0.84 |
| AG-447-C19 | GORG | 1 | 1 | complete | 2 | 0 | orphan | 0 | 0 | NA | 0 | 2 | orphan | 83.66 | 0 | 1.1 |
| AG-390-G11 | GORG | 2 | 1 | complete | 1 | 0 | orphan | 0 | 0 | NA | 0 | 0 | NA | 83.65 | 0 | 1.28 |
| AG-426-K19 | GORG | 1 | 1 | complete | 1 | 0 | orphan | 0 | 0 | NA | 0 | 0 | NA | 83.65 | 0 | 1.03 |

| | | | | | | | | | | | | | | | | |
|---|---|---|---|---|---|---|---|---|---|---|---|---|---|---|---|---|
| AG-461-C04 | GORG | 0 | 0 | NA | 1 | 0 | orphan | 0 | 0 | NA | 0 | 0 | NA | 83.63 | 0 | 1.07 |
| AG-892-A23 | GORG | 0 | 0 | NA | 0 | 0 | NA | 0 | 0 | NA | 0 | 1 | orphan | 83.61 | 0 | 1.33 |
| AG-919-G06 | GORG | 0 | 0 | NA | 2 | 0 | orphan | 0 | 0 | NA | 0 | 0 | NA | 83.52 | 0 | 1.18 |
| AG-349-A08 | GORG | 0 | 0 | NA | 1 | 0 | orphan | 0 | 0 | NA | 0 | 0 | NA | 83.45 | 0 | 0.96 |
| AG-915-E06 | GORG | 0 | 0 | NA | 2 | 0 | orphan | 0 | 0 | NA | 0 | 0 | NA | 83.41 | 0 | 1.04 |
| AH-326-D06 | GORG | 0 | 0 | NA | 1 | 0 | orphan | 0 | 0 | NA | 0 | 0 | NA | 83.32 | 0 | 0.93 |
| AG-365-P03 | GORG | 0 | 0 | NA | 1 | 0 | orphan | 0 | 0 | NA | 0 | 2 | orphan | 83.24 | 0 | 1.1 |
| AG-910-E21 | GORG | 1 | 1 | complete | 2 | 0 | orphan | 0 | 0 | NA | 0 | 0 | NA | 83.24 | 0 | 1.04 |
| AG-439-N16 | GORG | 0 | 0 | NA | 1 | 0 | orphan | 0 | 0 | NA | 0 | 0 | NA | 83.13 | 0 | 0.91 |
| AG-447-L18 | GORG | 0 | 0 | NA | 1 | 0 | orphan | 0 | 0 | NA | 0 | 0 | NA | 83.13 | 0 | 1.49 |
| AG-453-K22 | GORG | 0 | 0 | NA | 3 | 0 | orphan | 0 | 0 | NA | 0 | 0 | NA | 83.13 | 0 | 0.98 |
| AG-891-N03 | GORG | 0 | 0 | NA | 1 | 0 | orphan | 0 | 0 | NA | 0 | 0 | NA | 83.13 | 0 | 1.2 |
| AG-894-G18 | GORG | 0 | 0 | NA | 1 | 0 | orphan | 0 | 0 | NA | 0 | 0 | NA | 83.13 | 0 | 1.08 |
| AG-896-G07 | GORG | 1 | 1 | complete | 1 | 0 | orphan | 0 | 0 | NA | 0 | 0 | NA | 83.13 | 0 | 1.24 |
| AG-911-E02 | GORG | 0 | 0 | NA | 2 | 0 | orphan | 0 | 0 | NA | 0 | 0 | NA | 83.13 | 0 | 1.24 |
| AG-913-O18 | GORG | 1 | 1 | complete | 1 | 0 | orphan | 0 | 0 | NA | 0 | 0 | NA | 83.13 | 0 | 1.2 |
| AG-917-D09 | GORG | 0 | 0 | NA | 2 | 0 | orphan | 0 | 0 | NA | 0 | 0 | NA | 83.13 | 0 | 1.22 |
| AG-919-N17 | GORG | 0 | 0 | NA | 1 | 0 | orphan | 0 | 0 | NA | 0 | 0 | NA | 83.13 | 0 | 1.1 |
| AH-287-I02 | GORG | 0 | 0 | NA | 1 | 0 | orphan | 0 | 0 | NA | 0 | 0 | NA | 83.13 | 0 | 0.84 |
| AH-321-O20 | GORG | 1 | 0 | orphan | 0 | 0 | NA | 0 | 0 | NA | 0 | 0 | NA | 83.13 | 0 | 0.98 |
| AG-891-G19 | GORG | 0 | 0 | NA | 1 | 0 | orphan | 1 | 1 | complete | 0 | 0 | NA | 83.09 | 0 | 1.32 |
| AG-447-I15 | GORG | 0 | 0 | NA | 1 | 0 | orphan | 0 | 0 | NA | 0 | 0 | NA | 83.07 | 0 | 1.51 |
| AG-349-D19 | GORG | 0 | 0 | NA | 1 | 0 | orphan | 0 | 0 | NA | 0 | 0 | NA | 83.03 | 0 | 0.92 |
| AG-426-N15 | GORG | 0 | 0 | NA | 1 | 0 | orphan | 0 | 0 | NA | 0 | 0 | NA | 82.96 | 0 | 1.07 |
| AG-426-K09 | GORG | 0 | 0 | NA | 1 | 0 | orphan | 0 | 0 | NA | 0 | 0 | NA | 82.92 | 0 | 1.23 |

| | | | | | | | | | | | | | | | | |
|---|---|---|---|---|---|---|---|---|---|---|---|---|---|---|---|---|
| AG-894-C11 | GORG | 0 | 0 | NA | 1 | 0 | orphan | 0 | 0 | NA | 0 | 2 | orphan | 82.88 | 0 | 1.29 |
| AG-914-P23 | GORG | 0 | 0 | NA | 1 | 0 | orphan | 0 | 0 | NA | 0 | 0 | NA | 82.88 | 0 | 1.08 |
| AG-917-N02 | GORG | 0 | 0 | NA | 1 | 0 | orphan | 0 | 0 | NA | 0 | 0 | NA | 82.86 | 0 | 0.95 |
| AG-439-D21 | GORG | 0 | 0 | NA | 3 | 0 | orphan | 0 | 0 | NA | 0 | 0 | NA | 82.84 | 0 | 3.22 |
| AG-892-E23 | GORG | 0 | 0 | NA | 2 | 0 | orphan | 0 | 0 | NA | 0 | 0 | NA | 82.75 | 0 | 1.18 |
| AG-917-G09 | GORG | 0 | 0 | NA | 2 | 0 | orphan | 0 | 0 | NA | 0 | 0 | NA | 82.73 | 0 | 1.16 |
| AG-892-E21 | GORG | 0 | 0 | NA | 1 | 0 | orphan | 0 | 0 | NA | 0 | 0 | NA | 82.72 | 0 | 1.19 |
| AG-426-L04 | GORG | 0 | 0 | NA | 2 | 0 | orphan | 0 | 0 | NA | 0 | 0 | NA | 82.67 | 0 | 0.95 |
| AG-899-G02 | GORG | 0 | 0 | NA | 1 | 0 | orphan | 0 | 0 | NA | 0 | 0 | NA | 82.64 | 0 | 0.97 |
| AG-918-M19 | GORG | 0 | 0 | NA | 2 | 0 | orphan | 0 | 0 | NA | 0 | 0 | NA | 82.64 | 0 | 1.13 |
| AG-439-C21 | GORG | 0 | 0 | NA | 1 | 0 | orphan | 0 | 0 | NA | 0 | 0 | NA | 82.56 | 0 | 1.15 |
| AG-359-G09 | GORG | 1 | 1 | complete | 1 | 0 | orphan | 0 | 0 | NA | 0 | 0 | NA | 82.53 | 0 | 1.32 |
| AG-439-G19 | GORG | 0 | 0 | NA | 1 | 0 | orphan | 0 | 0 | NA | 0 | 0 | NA | 82.53 | 0 | 1.11 |
| AG-892-O19 | GORG | 0 | 0 | NA | 1 | 0 | orphan | 0 | 0 | NA | 0 | 0 | NA | 82.53 | 0 | 0.94 |
| AG-915-O06 | GORG | 0 | 0 | NA | 2 | 0 | orphan | 0 | 0 | NA | 0 | 0 | NA | 82.53 | 0 | 1.31 |
| AG-916-I06 | GORG | 0 | 0 | NA | 1 | 0 | orphan | 0 | 0 | NA | 0 | 0 | NA | 82.53 | 0 | 1.27 |
| AG-453-B11 | GORG | 1 | 1 | complete | 0 | 0 | NA | 0 | 0 | NA | 0 | 0 | NA | 82.52 | 0 | 1.28 |
| AG-909-K20 | GORG | 0 | 0 | NA | 2 | 0 | orphan | 0 | 0 | NA | 0 | 0 | NA | 82.48 | 0 | 1.19 |
| AH-287-E17 | GORG | 0 | 0 | NA | 2 | 1 | orphan | 0 | 0 | NA | 0 | 0 | NA | 82.46 | 0 | 1.07 |
| AG-917-N08 | GORG | 0 | 0 | NA | 1 | 0 | orphan | 0 | 0 | NA | 0 | 0 | NA | 82.42 | 0 | 0.95 |
| AH-287-F08 | GORG | 0 | 0 | NA | 2 | 0 | orphan | 0 | 0 | NA | 0 | 0 | NA | 82.42 | 0 | 0.92 |
| AH-321-M16 | GORG | 1 | 0 | orphan | 2 | 0 | orphan | 0 | 0 | NA | 0 | 0 | NA | 82.42 | 0 | 2.17 |
| AG-337-I22 | GORG | 0 | 0 | NA | 1 | 0 | orphan | 0 | 0 | NA | 0 | 0 | NA | 82.41 | 0 | 1.22 |
| AH-321-G15 | GORG | 0 | 0 | NA | 3 | 0 | orphan | 0 | 0 | NA | 0 | 0 | NA | 82.4 | 0 | 0.91 |
| AG-359-J06 | GORG | 0 | 0 | NA | 2 | 0 | orphan | 0 | 0 | NA | 0 | 0 | NA | 82.39 | 0 | 1.07 |
| AG-359-G22 | GORG | 0 | 0 | NA | 2 | 0 | orphan | 0 | 0 | NA | 0 | 0 | NA | 82.38 | 0 | 0.98 |

| | | | | | | | | | | | | | | | | |
|---|---|---|---|---|---|---|---|---|---|---|---|---|---|---|---|---|
| AG-325-B10 | GORG | 1 | 1 | complete | 1 | 0 | orphan | 0 | 0 | NA | 0 | 0 | NA | 82.37 | 0 | 0.9 |
| AG-894-F03 | GORG | 0 | 0 | NA | 2 | 0 | orphan | 0 | 0 | NA | 0 | 0 | NA | 82.28 | 0 | 1.07 |
| AH-287-E04 | GORG | 0 | 0 | NA | 1 | 0 | orphan | 0 | 0 | NA | 0 | 0 | NA | 82.28 | 0 | 1.35 |
| AG-447-O08 | GORG | 0 | 0 | NA | 1 | 0 | orphan | 0 | 0 | NA | 0 | 1 | orphan | 82.23 | 0 | 1.11 |
| AG-892-C11 | GORG | 0 | 0 | NA | 1 | 0 | orphan | 0 | 0 | NA | 0 | 0 | NA | 82.23 | 0 | 0.97 |
| AG-916-J09 | GORG | 0 | 0 | NA | 3 | 1 | orphan | 0 | 0 | NA | 0 | 0 | NA | 82.23 | 0 | 0.88 |
| AH-321-A17 | GORG | 0 | 0 | NA | 2 | 0 | orphan | 0 | 0 | NA | 0 | 0 | NA | 82.23 | 0 | 1.18 |
| AH-321-L02 | GORG | 0 | 0 | NA | 3 | 1 | complete | 0 | 0 | NA | 0 | 2 | orphan | 82.22 | 0 | 1.31 |
| AG-894-B13 | GORG | 0 | 0 | NA | 1 | 0 | orphan | 0 | 0 | NA | 0 | 0 | NA | 82.21 | 0 | 1 |
| AG-917-B15 | GORG | 0 | 0 | NA | 1 | 0 | orphan | 0 | 0 | NA | 0 | 0 | NA | 82.2 | 0 | 0.96 |
| AG-390-I11 | GORG | 0 | 0 | NA | 1 | 1 | complete | 0 | 0 | NA | 0 | 0 | NA | 82.15 | 0 | 1.3 |
| AG-892-D10 | GORG | 0 | 0 | NA | 1 | 0 | orphan | 0 | 0 | NA | 0 | 0 | NA | 82.13 | 0 | 1.13 |
| AG-898-F23 | GORG | 0 | 0 | NA | 1 | 0 | orphan | 0 | 0 | NA | 0 | 0 | NA | 82.12 | 0 | 1.1 |
| AG-896-N10 | GORG | 0 | 0 | NA | 1 | 0 | orphan | 0 | 0 | NA | 0 | 0 | NA | 82.11 | 0 | 0.91 |
| AG-365-N20 | GORG | 0 | 0 | NA | 1 | 0 | orphan | 0 | 0 | NA | 0 | 0 | NA | 82.09 | 0 | 1.4 |
| AG-430-O11 | GORG | 0 | 0 | NA | 2 | 0 | orphan | 0 | 0 | NA | 0 | 0 | NA | 82.09 | 0 | 0.98 |
| AG-912-B19 | GORG | 0 | 0 | NA | 1 | 0 | orphan | 0 | 0 | NA | 0 | 0 | NA | 82.09 | 0 | 1.18 |
| AG-911-P17 | GORG | 0 | 0 | NA | 1 | 0 | orphan | 0 | 0 | NA | 0 | 0 | NA | 82.08 | 0 | 0.92 |
| AG-426-M22 | GORG | 0 | 0 | NA | 2 | 0 | orphan | 0 | 0 | NA | 0 | 0 | NA | 82.04 | 0 | 1.04 |
| AG-390-I17 | GORG | 0 | 0 | NA | 3 | 0 | orphan | 0 | 0 | NA | 0 | 0 | NA | 81.99 | 0 | 1.01 |
| AG-447-K20 | GORG | 0 | 0 | NA | 2 | 1 | complete | 0 | 0 | NA | 0 | 0 | NA | 81.99 | 0 | 1.36 |
| AG-337-E07 | GORG | 1 | 1 | complete | 2 | 0 | orphan | 0 | 0 | NA | 0 | 0 | NA | 81.93 | 0 | 1.11 |
| AG-404-B11 | GORG | 0 | 0 | NA | 1 | 0 | orphan | 1 | 1 | complete | 0 | 0 | NA | 81.93 | 0 | 0.82 |
| AG-457-O03 | GORG | 0 | 0 | NA | 1 | 0 | orphan | 0 | 0 | NA | 0 | 0 | NA | 81.93 | 0 | 1.07 |
| AG-896-M05 | GORG | 0 | 0 | NA | 1 | 0 | orphan | 0 | 0 | NA | 0 | 0 | NA | 81.93 | 0 | 1.08 |

| | | | | | | | | | | | | | | | | |
|---|---|---|---|---|---|---|---|---|---|---|---|---|---|---|---|---|
| AG-896-O16 | GORG | 0 | 0 | NA | 2 | 0 | orphan | 0 | 0 | NA | 0 | 0 | NA | 81.93 | 0 | 0.84 |
| AG-909-A17 | GORG | 0 | 0 | NA | 1 | 0 | orphan | 0 | 0 | NA | 0 | 0 | NA | 81.93 | 0 | 1.05 |
| AG-911-K05 | GORG | 0 | 0 | NA | 1 | 0 | orphan | 0 | 0 | NA | 0 | 0 | NA | 81.93 | 0 | 1.12 |
| AG-915-C06 | GORG | 0 | 0 | NA | 2 | 0 | orphan | 1 | 1 | complete | 0 | 0 | NA | 81.93 | 0 | 1.26 |
| AG-916-O06 | GORG | 0 | 0 | NA | 2 | 0 | orphan | 0 | 0 | NA | 0 | 0 | NA | 81.93 | 0 | 0.92 |
| AG-917-M17 | GORG | 0 | 0 | NA | 1 | 0 | orphan | 0 | 0 | NA | 0 | 0 | NA | 81.93 | 0 | 1.09 |
| AG-909-N06 | GORG | 0 | 0 | NA | 2 | 0 | orphan | 1 | 1 | complete | 0 | 0 | NA | 81.9 | 0 | 1.05 |
| AG-359-E10 | GORG | 0 | 0 | NA | 2 | 0 | orphan | 1 | 1 | complete | 0 | 0 | NA | 81.87 | 0 | 1.05 |
| AG-410-O14 | GORG | 0 | 0 | NA | 1 | 0 | orphan | 0 | 0 | NA | 0 | 0 | NA | 81.86 | 0 | 1 |
| AG-893-G21 | GORG | 0 | 0 | NA | 1 | 0 | orphan | 0 | 0 | NA | 0 | 0 | NA | 81.83 | 0 | 2.11 |
| AG-349-N13 | GORG | 0 | 0 | NA | 2 | 0 | orphan | 0 | 0 | NA | 0 | 0 | NA | 81.82 | 0 | 0.91 |
| AG-893-F20 | GORG | 0 | 0 | NA | 0 | 0 | NA | 0 | 0 | NA | 0 | 1 | orphan | 81.7 | 0 | 1.26 |
| AG-895-M18 | GORG | 0 | 0 | NA | 0 | 0 | NA | 0 | 0 | NA | 0 | 1 | orphan | 81.7 | 0 | 1.31 |
| AG-439-I14 | GORG | 0 | 0 | NA | 1 | 0 | orphan | 0 | 0 | NA | 0 | 0 | NA | 81.7 | 0 | 0.95 |
| AG-913-A07 | GORG | 0 | 0 | NA | 2 | 0 | orphan | 0 | 0 | NA | 0 | 0 | NA | 81.7 | 0 | 1.04 |
| AG-909-E04 | GORG | 0 | 0 | NA | 2 | 0 | orphan | 0 | 0 | NA | 0 | 0 | NA | 81.69 | 0 | 0.92 |
| AG-390-L19 | GORG | 0 | 0 | NA | 2 | 0 | orphan | 0 | 0 | NA | 0 | 0 | NA | 81.62 | 0 | 1.52 |
| AG-912-K10 | GORG | 0 | 0 | NA | 1 | 0 | orphan | 0 | 0 | NA | 0 | 0 | NA | 81.59 | 0 | 1.05 |
| AG-359-I06 | GORG | 0 | 0 | NA | 3 | 0 | orphan | 0 | 0 | NA | 0 | 0 | NA | 81.58 | 0 | 1.15 |
| AG-895-B19 | GORG | 0 | 0 | NA | 1 | 0 | orphan | 0 | 0 | NA | 0 | 0 | NA | 81.53 | 0 | 1.14 |
| AG-916-J06 | GORG | 0 | 0 | NA | 1 | 0 | orphan | 0 | 0 | NA | 0 | 0 | NA | 81.52 | 0 | 1.1 |
| AG-913-E11 | GORG | 0 | 0 | NA | 2 | 0 | orphan | 0 | 0 | NA | 0 | 0 | NA | 81.44 | 0 | 1.05 |
| AH-321-F10 | GORG | 0 | 0 | NA | 2 | 0 | orphan | 0 | 0 | NA | 0 | 0 | NA | 81.43 | 0 | 0.91 |
| AG-404-N21 | GORG | 0 | 0 | NA | 1 | 0 | orphan | 0 | 0 | NA | 0 | 0 | NA | 81.38 | 0 | 1.09 |
| AG-903-N02 | GORG | 0 | 0 | NA | 1 | 0 | orphan | 0 | 0 | NA | 0 | 0 | NA | 81.37 | 0 | 1.2 |

| | | | | | | | | | | | | | | | |
|---|---|---|---|---|---|---|---|---|---|---|---|---|---|---|---|
| AG-908-O10 | GORG | 0 | 0 | NA | 1 | 0 | orphan | 0 | 0 | NA | 0 | 0 | NA | 81.35 | 0 | 1.02 |
| AG-915-C08 | GORG | 0 | 0 | NA | 1 | 0 | orphan | 0 | 0 | NA | 0 | 0 | NA | 81.34 | 0 | 1 |
| AG-892-N06 | GORG | 0 | 0 | NA | 1 | 0 | orphan | 0 | 0 | NA | 0 | 0 | NA | 81.33 | 0 | 1.11 |
| AG-911-J07 | GORG | 0 | 0 | NA | 3 | 0 | orphan | 0 | 0 | NA | 0 | 0 | NA | 81.33 | 0 | 0.98 |
| AG-915-I10 | GORG | 0 | 0 | NA | 2 | 0 | orphan | 0 | 0 | NA | 0 | 0 | NA | 81.33 | 0 | 1.18 |
| AG-916-D16 | GORG | 0 | 0 | NA | 1 | 0 | orphan | 0 | 0 | NA | 0 | 0 | NA | 81.33 | 0 | 1.02 |
| AG-920-I06 | GORG | 0 | 0 | NA | 1 | 0 | orphan | 0 | 0 | NA | 0 | 0 | NA | 81.33 | 0 | 0.98 |
| AG-911-F13 | GORG | 0 | 0 | NA | 1 | 0 | orphan | 0 | 0 | NA | 0 | 0 | NA | 81.32 | 0 | 1.21 |
| AG-414-A03 | GORG | 0 | 0 | NA | 1 | 0 | orphan | 0 | 0 | NA | 0 | 1 | orphan | 81.28 | 0 | 0.89 |
| AG-422-K13 | GORG | 0 | 0 | NA | 1 | 0 | orphan | 0 | 0 | NA | 0 | 0 | NA | 81.25 | 0 | 1.41 |
| AG-426-L19 | GORG | 0 | 0 | NA | 1 | 0 | orphan | 0 | 0 | NA | 0 | 0 | NA | 81.25 | 0 | 0.94 |
| AG-893-J15 | GORG | 1 | 1 | complete | 0 | 0 | NA | 0 | 0 | NA | 0 | 0 | NA | 81.2 | 0 | 1.24 |
| AG-901-K05 | GORG | 0 | 0 | NA | 1 | 0 | orphan | 0 | 0 | NA | 0 | 0 | NA | 81.2 | 0 | 0.96 |
| AG-917-O11 | GORG | 0 | 0 | NA | 1 | 0 | orphan | 0 | 0 | NA | 0 | 0 | NA | 81.2 | 0 | 1.13 |
| AG-461-P16 | GORG | 1 | 0 | orphan | 0 | 0 | NA | 0 | 0 | NA | 0 | 0 | NA | 81.16 | 0 | 1.36 |
| AG-894-M18 | GORG | 0 | 0 | NA | 2 | 0 | orphan | 0 | 1 | NA | 0 | 0 | NA | 81.16 | 0 | 1.09 |
| AG-914-E06 | GORG | 0 | 0 | NA | 1 | 0 | orphan | 0 | 0 | NA | 0 | 0 | NA | 81.16 | 0 | 0.89 |
| AG-422-N16 | GORG | 0 | 0 | NA | 2 | 0 | orphan | 0 | 0 | NA | 0 | 0 | NA | 81.14 | 0 | 0.89 |
| AG-390-M07 | GORG | 0 | 0 | NA | 1 | 0 | orphan | 0 | 0 | NA | 0 | 0 | NA | 81.13 | 0 | 1.11 |
| AG-896-F06 | GORG | 0 | 0 | NA | 1 | 0 | orphan | 0 | 0 | NA | 0 | 0 | NA | 81.13 | 0 | 1.35 |
| AG-447-P19 | GORG | 0 | 0 | NA | 1 | 0 | orphan | 0 | 0 | NA | 0 | 0 | NA | 81.12 | 0 | 0.99 |
| AG-913-E07 | GORG | 0 | 0 | NA | 2 | 0 | orphan | 1 | 1 | complete | 0 | 0 | NA | 81.12 | 0 | 1.07 |
| AG-337-N04 | GORG | 0 | 0 | NA | 1 | 0 | orphan | 0 | 0 | NA | 0 | 0 | NA | 81.1 | 0 | 0.85 |
| AG-899-C17 | GORG | 0 | 0 | NA | 2 | 0 | orphan | 0 | 0 | NA | 0 | 0 | NA | 81.08 | 0 | 1.11 |
| AG-914-K18 | GORG | 0 | 0 | NA | 1 | 0 | orphan | 0 | 0 | NA | 0 | 0 | NA | 81.08 | 0 | 0.82 |
| AG-912-P07 | GORG | 0 | 0 | NA | 1 | 0 | orphan | 0 | 0 | NA | 0 | 2 | orphan | 81.07 | 0 | 1.33 |

| | | | | | | | | | | | | | | | | |
|---|---|---|---|---|---|---|---|---|---|---|---|---|---|---|---|---|
| AH-324-I20 | GORG | 0 | 0 | NA | 3 | 0 | orphan | 0 | 0 | NA | 0 | 0 | NA | 81.04 | 0 | 1.06 |
| AG-447-C06 | GORG | 0 | 0 | NA | 1 | 0 | orphan | 0 | 0 | NA | 0 | 0 | NA | 80.95 | 0 | 0.93 |
| AG-900-P18 | GORG | 0 | 0 | NA | 1 | 0 | orphan | 0 | 0 | NA | 0 | 0 | NA | 80.95 | 0 | 0.99 |
| AG-359-N17 | GORG | 0 | 0 | NA | 1 | 0 | orphan | 0 | 0 | NA | 0 | 0 | NA | 80.94 | 0 | 0.96 |
| AG-916-K14 | GORG | 0 | 0 | NA | 1 | 0 | orphan | 0 | 0 | NA | 0 | 0 | NA | 80.94 | 0 | 0.94 |
| AH-321-K14 | GORG | 0 | 0 | NA | 3 | 0 | orphan | 0 | 0 | NA | 0 | 0 | NA | 80.89 | 0 | 1.7 |
| AG-899-P09 | GORG | 0 | 0 | NA | 1 | 0 | orphan | 0 | 0 | NA | 0 | 0 | NA | 80.86 | 0 | 0.92 |
| AG-894-O14 | GORG | 0 | 1 | NA | 0 | 0 | NA | 0 | 0 | NA | 0 | 0 | NA | 80.85 | 0 | 1.14 |
| AG-908-F18 | GORG | 0 | 0 | NA | 1 | 0 | orphan | 0 | 0 | NA | 0 | 0 | NA | 80.85 | 0 | 1.18 |
| AH-321-C19 | GORG | 0 | 0 | NA | 1 | 0 | orphan | 0 | 0 | NA | 0 | 0 | NA | 80.84 | 0 | 1.24 |
| AG-915-K19 | GORG | 0 | 0 | NA | 1 | 0 | orphan | 0 | 0 | NA | 0 | 0 | NA | 80.8 | 0 | 1.23 |
| AG-337-N02 | GORG | 1 | 1 | complete | 1 | 0 | orphan | 0 | 0 | NA | 0 | 0 | NA | 80.76 | 0 | 1.34 |
| AG-911-G16 | GORG | 0 | 0 | NA | 1 | 0 | orphan | 0 | 0 | NA | 0 | 0 | NA | 80.75 | 0 | 1.16 |
| AG-894-D05 | GORG | 0 | 0 | NA | 0 | 1 | NA | 0 | 0 | NA | 0 | 1 | orphan | 80.72 | 0 | 1.27 |
| AG-916-G03 | GORG | 0 | 0 | NA | 2 | 0 | orphan | 0 | 1 | NA | 0 | 1 | orphan | 80.72 | 0 | 0.93 |
| AG-422-B19 | GORG | 0 | 0 | NA | 1 | 0 | orphan | 0 | 0 | NA | 0 | 0 | NA | 80.72 | 0 | 1.03 |
| AG-430-J21 | GORG | 1 | 0 | orphan | 1 | 0 | orphan | 0 | 0 | NA | 0 | 0 | NA | 80.72 | 0 | 1.04 |
| AG-430-K07 | GORG | 0 | 0 | NA | 2 | 0 | orphan | 0 | 0 | NA | 0 | 0 | NA | 80.72 | 0 | 1.13 |
| AG-461-B21 | GORG | 0 | 0 | NA | 1 | 0 | orphan | 0 | 0 | NA | 0 | 0 | NA | 80.72 | 0 | 0.82 |
| AG-891-K21 | GORG | 1 | 1 | complete | 1 | 0 | orphan | 0 | 0 | NA | 0 | 0 | NA | 80.72 | 0 | 1.35 |
| AG-892-B06 | GORG | 0 | 0 | NA | 1 | 0 | orphan | 0 | 0 | NA | 0 | 0 | NA | 80.72 | 0 | 1.29 |
| AG-904-G09 | GORG | 0 | 0 | NA | 1 | 0 | orphan | 0 | 0 | NA | 0 | 0 | NA | 80.72 | 0 | 1.12 |
| AG-910-D13 | GORG | 0 | 0 | NA | 3 | 0 | orphan | 0 | 0 | NA | 0 | 0 | NA | 80.72 | 0 | 1.12 |
| AG-910-J22 | GORG | 1 | 1 | complete | 1 | 0 | orphan | 0 | 0 | NA | 0 | 0 | NA | 80.72 | 0 | 1.1 |
| AG-920-I21 | GORG | 0 | 0 | NA | 2 | 0 | orphan | 0 | 0 | NA | 0 | 0 | NA | 80.72 | 0 | 0.93 |

| | | | | | | | | | | | | | | | |
|---|---|---|---|---|---|---|---|---|---|---|---|---|---|---|---|
| AG-914-P03 | GORG | 0 | 0 | NA | 0 | 0 | NA | 0 | 0 | NA | 0 | 1 | orpha n | 80.71 | 0 | 1.28 |
| AG-895-F20 | GORG | 0 | 0 | NA | 1 | 0 | orphan | 0 | 0 | NA | 0 | 1 | orpha n | 80.65 | 0 | 1.17 |
| AG-916-B16 | GORG | 0 | 0 | NA | 1 | 0 | orphan | 0 | 0 | NA | 0 | 1 | orpha n | 80.65 | 0 | 1.29 |
| AG-907-A11 | GORG | 0 | 0 | NA | 1 | 0 | orphan | 0 | 0 | NA | 0 | 0 | NA | 80.65 | 0 | 0.97 |
| AH-321-F16 | GORG | 0 | 0 | NA | 1 | 0 | orphan | 0 | 0 | NA | 0 | 0 | NA | 80.65 | 0 | 1.02 |
| AG-435-M21 | GORG | 1 | 1 | complet e | 2 | 0 | orphan | 0 | 0 | NA | 0 | 0 | NA | 80.61 | 0 | 0.85 |
| AG-461-E05 | GORG | 0 | 0 | NA | 1 | 0 | orphan | 0 | 0 | NA | 0 | 0 | NA | 80.61 | 0 | 1.03 |
| AG-891-K23 | GORG | 0 | 0 | NA | 2 | 0 | orphan | 0 | 0 | NA | 0 | 0 | NA | 80.58 | 0 | 1.22 |
| AG-426-E13 | GORG | 0 | 0 | NA | 1 | 0 | orphan | 0 | 0 | NA | 0 | 0 | NA | 80.57 | 0 | 1.2 |
| AG-325-C21 | GORG | 0 | 0 | NA | 1 | 0 | orphan | 0 | 0 | NA | 0 | 0 | NA | 80.56 | 0 | 1.04 |
| AG-359-N22 | GORG | 0 | 0 | NA | 1 | 0 | orphan | 0 | 0 | NA | 0 | 0 | NA | 80.52 | 0 | 1.08 |
| AG-447-C16 | GORG | 0 | 0 | NA | 0 | 0 | NA | 0 | 0 | NA | 0 | 1 | orpha n | 80.49 | 0 | 1.13 |
| AG-390-E21 | GORG | 0 | 0 | NA | 1 | 0 | orphan | 0 | 0 | NA | 0 | 0 | NA | 80.48 | 0 | 1.01 |
| AG-892-C21 | GORG | 1 | 0 | orphan | 0 | 0 | NA | 0 | 0 | NA | 0 | 0 | NA | 80.46 | 0 | 1.16 |
| AG-913-G10 | GORG | 0 | 0 | NA | 1 | 0 | orphan | 0 | 0 | NA | 0 | 0 | NA | 80.43 | 0 | 1.1 |
| AG-916-L11 | GORG | 0 | 0 | NA | 1 | 0 | orphan | 0 | 0 | NA | 0 | 0 | NA | 80.42 | 0 | 1.1 |
| AG-915-J15 | GORG | 0 | 0 | NA | 1 | 0 | orphan | 0 | 0 | NA | 0 | 0 | NA | 80.4 | 0 | 1.02 |
| AG-325-K03 | GORG | 0 | 0 | NA | 2 | 0 | orphan | 0 | 0 | NA | 0 | 0 | NA | 80.39 | 0 | 1.07 |
| AG-918-B08 | GORG | 0 | 0 | NA | 1 | 0 | orphan | 0 | 0 | NA | 0 | 0 | NA | 80.37 | 0 | 0.95 |
| AG-895-B06 | GORG | 0 | 0 | NA | 2 | 0 | orphan | 0 | 0 | NA | 0 | 2 | orpha n | 80.36 | 0 | 1.02 |
| AG-390-J13 | GORG | 0 | 0 | NA | 1 | 0 | orphan | 0 | 0 | NA | 0 | 0 | NA | 80.33 | 0 | 1.05 |
| AG-464-P13 | GORG | 0 | 0 | NA | 1 | 0 | orphan | 0 | 0 | NA | 0 | 0 | NA | 80.32 | 0 | 1.18 |
| AG-422-N14 | GORG | 0 | 0 | NA | 1 | 0 | orphan | 0 | 0 | NA | 0 | 0 | NA | 80.3 | 0 | 1.07 |
| AG-430-K14 | GORG | 0 | 0 | NA | 4 | 0 | orphan | 0 | 0 | NA | 0 | 0 | NA | 80.3 | 0 | 0.94 |
| AG-905-A10 | GORG | 0 | 0 | NA | 1 | 0 | orphan | 0 | 0 | NA | 0 | 0 | NA | 80.25 | 0 | 0.99 |

| | | | | | | | | | | | | | | | | |
|---|---|---|---|---|---|---|---|---|---|---|---|---|---|---|---|---|
| AG-919-A02 | GORG | 1 | 1 | complet e | 1 | 0 | orphan | 0 | 0 | NA | 0 | 0 | NA | 80.21 | 0 | 1.18 |
| AG-337-G21 | GORG | 0 | 0 | NA | 1 | 0 | orphan | 0 | 0 | NA | 0 | 0 | NA | 80.19 | 0 | 0.95 |
| AG-913-I22 | GORG | 0 | 0 | NA | 1 | 0 | orphan | 0 | 0 | NA | 0 | 0 | NA | 80.16 | 0 | 0.87 |
| AG-901-C10 | GORG | 2 | 1 | complet e | 0 | 0 | NA | 0 | 0 | NA | 0 | 0 | NA | 80.15 | 0 | 1.21 |
| AG-908-A02 | GORG | 0 | 0 | NA | 1 | 0 | orphan | 0 | 0 | NA | 0 | 0 | NA | 80.15 | 0 | 1.32 |
| AG-470-O10 | GORG | 0 | 0 | NA | 3 | 0 | orphan | 0 | 0 | NA | 0 | 0 | NA | 80.12 | 0 | 1.13 |
| AG-893-L04 | GORG | 0 | 0 | NA | 1 | 0 | orphan | 0 | 0 | NA | 0 | 0 | NA | 80.12 | 0 | 1.12 |
| AG-894-P17 | GORG | 0 | 0 | NA | 1 | 0 | orphan | 0 | 1 | NA | 0 | 0 | NA | 80.12 | 0 | 1.03 |
| AG-912-E13 | GORG | 0 | 0 | NA | 1 | 0 | orphan | 0 | 1 | NA | 0 | 0 | NA | 80.12 | 0 | 0.91 |
| AG-917-A15 | GORG | 0 | 0 | NA | 1 | 0 | orphan | 0 | 0 | NA | 0 | 0 | NA | 80.12 | 0 | 1.02 |
| AH-287-I04 | GORG | 0 | 0 | NA | 1 | 1 | orphan | 0 | 0 | NA | 0 | 0 | NA | 80.12 | 0 | 0.82 |
| AG-894-K15 | GORG | 0 | 0 | NA | 1 | 0 | orphan | 1 | 1 | complet e | 0 | 0 | NA | 80.09 | 0 | 1.01 |
| AG-422-G15 | GORG | 0 | 0 | NA | 1 | 0 | orphan | 0 | 0 | NA | 0 | 0 | NA | 80.02 | 0 | 0.98 |
| AG-905-M18 | GORG | 0 | 0 | NA | 2 | 0 | orphan | 0 | 0 | NA | 0 | 0 | NA | 80.01 | 0 | 0.9 |
| AG-916-E15 | GORG | 0 | 0 | NA | 1 | 0 | orphan | 0 | 0 | NA | 0 | 0 | NA | 79.96 | 0 | 1.28 |
| AG-903-E15 | GORG | 0 | 0 | NA | 2 | 0 | orphan | 0 | 0 | NA | 0 | 0 | NA | 79.94 | 0 | 0.96 |
| AG-910-C16 | GORG | 0 | 0 | NA | 3 | 0 | orphan | 0 | 0 | NA | 0 | 0 | NA | 79.94 | 0 | 0.8 |
| AG-915-P01 | GORG | 0 | 0 | NA | 1 | 0 | orphan | 0 | 0 | NA | 0 | 0 | NA | 79.94 | 0 | 1.02 |
| AG-349-E20 | GORG | 0 | 0 | NA | 1 | 0 | orphan | 0 | 0 | NA | 0 | 0 | NA | 79.92 | 0 | 0.86 |
| AG-893-B07 | GORG | 0 | 0 | NA | 1 | 0 | orphan | 0 | 0 | NA | 0 | 0 | NA | 79.92 | 0 | 1.1 |
| AG-914-D08 | GORG | 0 | 0 | NA | 1 | 0 | orphan | 0 | 0 | NA | 0 | 0 | NA | 79.89 | 0 | 1.01 |
| AG-911-G14 | GORG | 0 | 0 | NA | 2 | 1 | complet e | 0 | 0 | NA | 0 | 0 | NA | 79.84 | 0 | 1.28 |
| AH-321-N17 | GORG | 0 | 0 | NA | 2 | 0 | orphan | 0 | 0 | NA | 0 | 0 | NA | 79.82 | 0 | 1.12 |
| AG-918-M20 | GORG | 0 | 0 | NA | 2 | 0 | orphan | 0 | 0 | NA | 0 | 0 | NA | 79.74 | 0 | 0.93 |
| AG-904-D16 | GORG | 0 | 0 | NA | 1 | 0 | orphan | 0 | 0 | NA | 0 | 0 | NA | 79.7 | 0 | 1 |
| AG-918-L06 | GORG | 1 | 1 | complet | 1 | 0 | orphan | 0 | 0 | NA | 0 | 0 | NA | 79.69 | 0 | 1.29 |

| | | | | | | | | | | | | | | | | |
|---|---|---|---|---|---|---|---|---|---|---|---|---|---|---|---|---|
| | | | | e | | | | | | | | | | | | |
| AG-414-F02 | GORG | 0 | 0 | NA | 1 | 0 | orphan | 0 | 0 | NA | 0 | 0 | NA | 79.67 | 0 | 0.89 |
| AG-426-P19 | GORG | 0 | 0 | NA | 2 | 0 | orphan | 0 | 0 | NA | 0 | 0 | NA | 79.67 | 0 | 1.49 |
| AG-894-I07 | GORG | 0 | 0 | NA | 1 | 0 | orphan | 0 | 0 | NA | 0 | 0 | NA | 79.66 | 0 | 1.11 |
| AG-337-J22 | GORG | 0 | 0 | NA | 1 | 0 | orphan | 0 | 0 | NA | 0 | 0 | NA | 79.65 | 0 | 1.04 |
| AG-909-I04 | GORG | 0 | 0 | NA | 2 | 0 | orphan | 0 | 0 | NA | 0 | 0 | NA | 79.62 | 0 | 0.95 |
| AG-453-G21 | GORG | 1 | 0 | orphan | 1 | 0 | orphan | 0 | 0 | NA | 0 | 0 | NA | 79.58 | 0 | 0.92 |
| AG-414-K15 | GORG | 0 | 0 | NA | 1 | 0 | orphan | 0 | 0 | NA | 0 | 0 | NA | 79.52 | 0 | 1.05 |
| AG-422-A21 | GORG | 0 | 0 | NA | 1 | 0 | orphan | 0 | 0 | NA | 0 | 0 | NA | 79.52 | 0 | 0.97 |
| AG-426-F11 | GORG | 0 | 0 | NA | 1 | 0 | orphan | 0 | 0 | NA | 0 | 0 | NA | 79.52 | 0 | 1.1 |
| AG-461-P14 | GORG | 1 | 1 | complet e | 1 | 0 | orphan | 0 | 0 | NA | 0 | 0 | NA | 79.52 | 0 | 1.18 |
| AG-894-P18 | GORG | 0 | 0 | NA | 1 | 1 | orphan | 0 | 0 | NA | 0 | 0 | NA | 79.52 | 0 | 0.86 |
| AG-896-M10 | GORG | 0 | 0 | NA | 2 | 0 | orphan | 0 | 0 | NA | 0 | 0 | NA | 79.52 | 0 | 1.12 |
| AG-904-N09 | GORG | 0 | 0 | NA | 1 | 0 | orphan | 0 | 0 | NA | 0 | 0 | NA | 79.52 | 0 | 1.11 |
| AG-904-O04 | GORG | 0 | 0 | NA | 1 | 0 | orphan | 0 | 0 | NA | 0 | 0 | NA | 79.52 | 0 | 1.16 |
| AG-913-K15 | GORG | 0 | 0 | NA | 2 | 0 | orphan | 0 | 0 | NA | 0 | 0 | NA | 79.52 | 0 | 1 |
| AG-915-B23 | GORG | 0 | 0 | NA | 1 | 0 | orphan | 0 | 0 | NA | 0 | 0 | NA | 79.52 | 0 | 0.95 |
| AG-919-I11 | GORG | 0 | 0 | NA | 1 | 0 | orphan | 0 | 0 | NA | 0 | 0 | NA | 79.52 | 0 | 1.05 |
| AH-326-G20 | GORG | 0 | 0 | NA | 1 | 0 | orphan | 0 | 0 | NA | 0 | 0 | NA | 79.52 | 0 | 0.89 |
| AG-359-P21 | GORG | 1 | 1 | complet e | 1 | 0 | orphan | 0 | 0 | NA | 0 | 0 | NA | 79.41 | 0 | 1.11 |
| AG-915-C22 | GORG | 0 | 0 | NA | 1 | 0 | orphan | 0 | 0 | NA | 0 | 0 | NA | 79.4 | 0 | 1.04 |
| AG-899-O16 | GORG | 0 | 0 | NA | 1 | 0 | orphan | 0 | 0 | NA | 0 | 2 | orpha n | 79.39 | 0 | 1.17 |
| AG-430-D06 | GORG | 0 | 0 | NA | 1 | 0 | orphan | 0 | 0 | NA | 0 | 0 | NA | 79.38 | 0 | 0.89 |
| AG-430-F02 | GORG | 0 | 0 | NA | 1 | 0 | orphan | 0 | 0 | NA | 0 | 0 | NA | 79.34 | 0 | 0.89 |
| AG-920-G11 | GORG | 0 | 0 | NA | 2 | 0 | orphan | 0 | 0 | NA | 0 | 2 | orpha n | 79.32 | 0 | 1.58 |
| AG-333-M06 | GORG | 0 | 0 | NA | 1 | 0 | orphan | 0 | 0 | NA | 0 | 0 | NA | 79.31 | 0 | 0.89 |

| | | | | | | | | | | | | | | | | |
|---|---|---|---|---|---|---|---|---|---|---|---|---|---|---|---|---|
| AG-912-F21 | GORG | 0 | 0 | NA | 4 | 0 | orphan | 0 | 0 | NA | 0 | 0 | NA | 79.31 | 0 | 1.17 |
| AG-904-G02 | GORG | 0 | 0 | NA | 1 | 0 | orphan | 0 | 0 | NA | 0 | 1 | orphan | 79.26 | 0 | 1.32 |
| AG-919-F09 | GORG | 0 | 0 | NA | 2 | 0 | orphan | 0 | 0 | NA | 0 | 1 | orphan | 79.25 | 0 | 1.06 |
| AG-439-C08 | GORG | 0 | 0 | NA | 1 | 0 | orphan | 0 | 0 | NA | 0 | 0 | NA | 79.25 | 0 | 0.99 |
| AG-893-I20 | GORG | 0 | 0 | NA | 1 | 0 | orphan | 0 | 0 | NA | 0 | 0 | NA | 79.25 | 0 | 1.07 |
| AG-916-B15 | GORG | 0 | 0 | NA | 1 | 0 | orphan | 0 | 0 | NA | 0 | 0 | NA | 79.25 | 0 | 1.24 |
| AG-894-G16 | GORG | 0 | 0 | NA | 0 | 0 | NA | 0 | 0 | NA | 0 | 1 | orphan | 79.21 | 0 | 1.31 |
| AG-349-O03 | GORG | 0 | 0 | NA | 1 | 0 | orphan | 0 | 0 | NA | 0 | 0 | NA | 79.2 | 0 | 1 |
| AG-313-M10 | GORG | 0 | 0 | NA | 2 | 1 | complete | 0 | 0 | NA | 0 | 0 | NA | 79.14 | 0 | 0.97 |
| AG-404-P22 | GORG | 0 | 0 | NA | 2 | 0 | orphan | 0 | 0 | NA | 0 | 0 | NA | 79.12 | 0 | 1.28 |
| AG-891-L15 | GORG | 0 | 0 | NA | 2 | 0 | orphan | 0 | 0 | NA | 0 | 0 | NA | 79.09 | 0 | 0.98 |
| AH-324-B22 | GORG | 0 | 0 | NA | 1 | 0 | orphan | 0 | 0 | NA | 0 | 0 | NA | 79.09 | 0 | 0.89 |
| AG-912-L10 | GORG | 0 | 0 | NA | 3 | 0 | orphan | 0 | 0 | NA | 0 | 0 | NA | 79.05 | 0 | 0.99 |
| AG-912-I09 | GORG | 0 | 0 | NA | 1 | 0 | orphan | 0 | 0 | NA | 0 | 0 | NA | 79.03 | 0 | 0.85 |
| AG-919-P14 | GORG | 1 | 1 | orphan | 1 | 0 | orphan | 0 | 0 | NA | 0 | 0 | NA | 79.03 | 0 | 1.13 |
| AG-447-B21 | GORG | 0 | 0 | NA | 2 | 0 | orphan | 0 | 0 | NA | 0 | 0 | NA | 78.99 | 0 | 1.15 |
| AG-919-O08 | GORG | 0 | 0 | NA | 1 | 0 | orphan | 0 | 0 | NA | 0 | 0 | NA | 78.97 | 0 | 0.97 |
| AG-919-O13 | GORG | 0 | 0 | NA | 1 | 0 | orphan | 0 | 0 | NA | 0 | 0 | NA | 78.97 | 0 | 0.97 |
| AG-337-L15 | GORG | 0 | 0 | NA | 1 | 0 | orphan | 0 | 0 | NA | 0 | 0 | NA | 78.92 | 0 | 1.15 |
| AG-414-G15 | GORG | 0 | 0 | NA | 2 | 0 | orphan | 0 | 0 | NA | 0 | 0 | NA | 78.92 | 0 | 1.01 |
| AG-430-N23 | GORG | 0 | 0 | NA | 1 | 0 | orphan | 0 | 0 | NA | 0 | 0 | NA | 78.92 | 0 | 0.89 |
| AG-894-P02 | GORG | 0 | 0 | NA | 2 | 0 | orphan | 0 | 0 | NA | 0 | 0 | NA | 78.92 | 0 | 1.2 |
| AG-912-N10 | GORG | 0 | 0 | NA | 1 | 0 | orphan | 0 | 0 | NA | 0 | 0 | NA | 78.92 | 0 | 1.37 |
| AG-913-B14 | GORG | 1 | 0 | orphan | 1 | 0 | orphan | 0 | 0 | NA | 0 | 0 | NA | 78.92 | 0 | 1.11 |
| AG-915-C05 | GORG | 0 | 0 | NA | 1 | 0 | orphan | 0 | 1 | NA | 0 | 0 | NA | 78.92 | 0 | 0.94 |
| AG-917-K23 | GORG | 0 | 0 | NA | 0 | 0 | NA | 0 | 1 | NA | 0 | 0 | NA | 78.92 | 0 | 1.18 |

| | | | | | | | | | | | | | | | | |
|---|---|---|---|---|---|---|---|---|---|---|---|---|---|---|---|---|
| AG-390-N19 | GORG | 0 | 0 | NA | 1 | 0 | orphan | 0 | 0 | NA | 0 | 0 | NA | 78.91 | 0 | 1.04 |
| AG-916-P17 | GORG | 0 | 0 | NA | 1 | 0 | orphan | 0 | 0 | NA | 0 | 0 | NA | 78.84 | 0 | 1.04 |
| AH-326-C02 | GORG | 0 | 0 | NA | 1 | 0 | orphan | 0 | 0 | NA | 0 | 0 | NA | 78.81 | 0 | 0.83 |
| AG-325-L14 | GORG | 0 | 0 | NA | 3 | 0 | orphan | 0 | 0 | NA | 0 | 0 | NA | 78.8 | 0 | 0.92 |
| AH-287-M04 | GORG | 0 | 0 | NA | 1 | 0 | orphan | 0 | 0 | NA | 0 | 0 | NA | 78.8 | 0 | 1.42 |
| AG-912-P08 | GORG | 0 | 0 | NA | 1 | 0 | orphan | 0 | 0 | NA | 0 | 0 | NA | 78.79 | 0 | 0.94 |
| AG-359-P07 | GORG | 0 | 0 | NA | 1 | 0 | orphan | 0 | 0 | NA | 0 | 0 | NA | 78.78 | 0 | 1 |
| AG-447-L10 | GORG | 0 | 0 | NA | 1 | 0 | orphan | 0 | 0 | NA | 0 | 0 | NA | 78.75 | 0 | 1.1 |
| AG-439-K10 | GORG | 0 | 0 | NA | 1 | 0 | orphan | 0 | 0 | NA | 0 | 0 | NA | 78.74 | 0 | 0.93 |
| AG-912-F07 | GORG | 0 | 0 | NA | 3 | 0 | orphan | 0 | 0 | NA | 0 | 0 | NA | 78.73 | 0 | 0.92 |
| AG-901-N10 | GORG | 0 | 0 | NA | 1 | 0 | orphan | 0 | 0 | NA | 0 | 0 | NA | 78.72 | 0 | 2.07 |
| AG-907-P16 | GORG | 1 | 1 | complete | 0 | 0 | NA | 0 | 0 | NA | 0 | 0 | NA | 78.72 | 0 | 0.95 |
| AG-404-B14 | GORG | 0 | 0 | NA | 1 | 0 | orphan | 0 | 0 | NA | 0 | 1 | orphan | 78.71 | 0 | 1.08 |
| AG-891-E08 | GORG | 0 | 0 | NA | 1 | 0 | orphan | 0 | 0 | NA | 0 | 0 | NA | 78.71 | 0 | 1.17 |
| AG-914-J20 | GORG | 0 | 0 | NA | 1 | 0 | orphan | 0 | 0 | NA | 0 | 0 | NA | 78.71 | 0 | 0.96 |
| AG-893-M16 | GORG | 0 | 0 | NA | 1 | 0 | orphan | 0 | 0 | NA | 0 | 0 | NA | 78.67 | 0 | 0.96 |
| AG-915-G02 | GORG | 0 | 0 | NA | 1 | 0 | orphan | 0 | 0 | NA | 0 | 0 | NA | 78.67 | 0 | 1.22 |
| AG-414-B09 | GORG | 0 | 0 | NA | 2 | 0 | orphan | 0 | 0 | NA | 0 | 0 | NA | 78.64 | 0 | 0.95 |
| AG-349-L04 | GORG | 0 | 0 | NA | 1 | 0 | orphan | 0 | 0 | NA | 0 | 0 | NA | 78.62 | 0 | 1.04 |
| AG-919-K14 | GORG | 0 | 0 | NA | 1 | 0 | orphan | 0 | 0 | NA | 0 | 0 | NA | 78.6 | 0 | 1.01 |
| AG-893-B10 | GORG | 0 | 0 | NA | 1 | 0 | orphan | 0 | 0 | NA | 0 | 0 | NA | 78.59 | 0 | 0.98 |
| AG-325-K02 | GORG | 0 | 0 | NA | 1 | 0 | orphan | 0 | 0 | NA | 0 | 0 | NA | 78.57 | 0 | 0.93 |
| AG-894-A09 | GORG | 0 | 0 | NA | 1 | 0 | orphan | 0 | 0 | NA | 0 | 0 | NA | 78.57 | 0 | 0.96 |
| AH-324-L09 | GORG | 0 | 0 | NA | 1 | 0 | orphan | 0 | 0 | NA | 0 | 0 | NA | 78.57 | 0 | 0.96 |
| AG-365-O14 | GORG | 0 | 0 | NA | 1 | 0 | orphan | 0 | 0 | NA | 0 | 0 | NA | 78.51 | 0 | 1.07 |
| AG-430-G21 | GORG | 0 | 0 | NA | 1 | 0 | orphan | 0 | 0 | NA | 0 | 0 | NA | 78.48 | 0 | 0.95 |

| | | | | | | | | | | | | | | | | |
|---|---|---|---|---|---|---|---|---|---|---|---|---|---|---|---|---|
| AG-414-K04 | GORG | 0 | 0 | NA | 1 | 0 | orphan | 0 | 0 | NA | 0 | 0 | NA | 78.42 | 0 | 1.04 |
| AG-905-G23 | GORG | 0 | 0 | NA | 2 | 0 | orphan | 0 | 0 | NA | 0 | 0 | NA | 78.42 | 0 | 1.2 |
| AG-895-J06 | GORG | 0 | 0 | NA | 1 | 0 | orphan | 0 | 0 | NA | 0 | 0 | NA | 78.4 | 0 | 1.18 |
| AG-333-P13 | GORG | 0 | 0 | NA | 2 | 0 | orphan | 0 | 0 | NA | 0 | 0 | NA | 78.38 | 0 | 0.89 |
| AG-920-B13 | GORG | 0 | 0 | NA | 1 | 0 | orphan | 0 | 0 | NA | 0 | 0 | NA | 78.38 | 0 | 0.97 |
| AG-337-F15 | GORG | 0 | 0 | NA | 1 | 0 | orphan | 0 | 0 | NA | 0 | 0 | NA | 78.37 | 0 | 1.18 |
| AG-899-O13 | GORG | 0 | 0 | NA | 1 | 0 | orphan | 0 | 0 | NA | 0 | 0 | NA | 78.32 | 0 | 0.93 |
| AG-333-D21 | GORG | 0 | 0 | NA | 3 | 0 | orphan | 0 | 0 | NA | 0 | 0 | NA | 78.31 | 0 | 1.17 |
| AG-343-D15 | GORG | 0 | 0 | NA | 1 | 0 | orphan | 0 | 0 | NA | 0 | 0 | NA | 78.31 | 0 | 0.96 |
| AG-365-C20 | GORG | 0 | 0 | NA | 2 | 0 | orphan | 0 | 0 | NA | 0 | 0 | NA | 78.31 | 0 | 1.03 |
| AG-414-I09 | GORG | 0 | 0 | NA | 1 | 0 | orphan | 0 | 0 | NA | 0 | 0 | NA | 78.31 | 0 | 0.78 |
| AG-457-L23 | GORG | 0 | 0 | NA | 2 | 0 | orphan | 0 | 0 | NA | 0 | 0 | NA | 78.31 | 0 | 0.84 |
| AG-457-M20 | GORG | 0 | 0 | NA | 2 | 0 | orphan | 0 | 0 | NA | 0 | 0 | NA | 78.31 | 0 | 0.87 |
| AG-470-I11 | GORG | 0 | 0 | NA | 1 | 0 | orphan | 0 | 0 | NA | 0 | 0 | NA | 78.31 | 0 | 0.9 |
| AG-893-E11 | GORG | 0 | 0 | NA | 1 | 0 | orphan | 0 | 0 | NA | 0 | 0 | NA | 78.31 | 0 | 1.42 |
| AG-898-E17 | GORG | 0 | 0 | NA | 1 | 0 | orphan | 0 | 0 | NA | 0 | 0 | NA | 78.31 | 0 | 1.03 |
| AG-898-P18 | GORG | 0 | 0 | NA | 1 | 0 | orphan | 0 | 0 | NA | 0 | 0 | NA | 78.31 | 0 | 1.13 |
| AG-904-I05 | GORG | 0 | 0 | NA | 2 | 0 | orphan | 0 | 0 | NA | 0 | 0 | NA | 78.31 | 0 | 0.82 |
| AG-905-B22 | GORG | 0 | 0 | NA | 1 | 0 | orphan | 0 | 0 | NA | 0 | 0 | NA | 78.31 | 0 | 0.81 |
| AG-908-D05 | GORG | 1 | 1 | complete | 1 | 0 | orphan | 0 | 0 | NA | 0 | 0 | NA | 78.31 | 0 | 0.97 |
| AG-909-F08 | GORG | 0 | 0 | NA | 1 | 0 | orphan | 0 | 0 | NA | 0 | 0 | NA | 78.31 | 0 | 1.03 |
| AG-910-K02 | GORG | 0 | 0 | NA | 1 | 0 | orphan | 0 | 0 | NA | 0 | 0 | NA | 78.31 | 0 | 1.15 |
| AG-913-O21 | GORG | 0 | 0 | NA | 1 | 0 | orphan | 0 | 0 | NA | 0 | 0 | NA | 78.31 | 0 | 0.96 |
| AG-919-F04 | GORG | 0 | 0 | NA | 1 | 0 | orphan | 0 | 0 | NA | 0 | 0 | NA | 78.31 | 0 | 0.88 |
| AG-910-O03 | GORG | 1 | 1 | complete | 1 | 0 | orphan | 0 | 0 | NA | 0 | 0 | NA | 78.28 | 0 | 0.93 |
| AG-919-P07 | GORG | 0 | 0 | NA | 1 | 0 | orphan | 0 | 0 | NA | 0 | 0 | NA | 78.28 | 0 | 0.89 |

| | | | | | | | | | | | | | | | | |
|---|---|---|---|---|---|---|---|---|---|---|---|---|---|---|---|---|
| AG-920-K09 | GORG | 0 | 0 | NA | 1 | 0 | orphan | 0 | 0 | NA | 0 | 0 | NA | 78.26 | 0 | 0.88 |
| AG-360-K10 | GORG | 0 | 0 | NA | 1 | 0 | orphan | 0 | 0 | NA | 0 | 0 | NA | 78.22 | 0 | 1.06 |
| AG-410-P11 | GORG | 0 | 0 | NA | 2 | 0 | orphan | 0 | 0 | NA | 0 | 0 | NA | 78.22 | 0 | 0.9 |
| AG-900-O15 | GORG | 0 | 0 | NA | 1 | 0 | orphan | 0 | 0 | NA | 0 | 0 | NA | 78.2 | 0 | 1.09 |
| AH-321-A03 | GORG | 0 | 0 | NA | 1 | 0 | orphan | 0 | 0 | NA | 0 | 0 | NA | 78.2 | 0 | 0.88 |
| AG-894-A18 | GORG | 0 | 0 | NA | 2 | 0 | orphan | 0 | 0 | NA | 0 | 0 | NA | 78.19 | 0 | 1.28 |
| AG-453-A22 | GORG | 0 | 0 | NA | 1 | 0 | orphan | 0 | 0 | NA | 0 | 0 | NA | 78.11 | 0 | 1.01 |
| AG-910-E04 | GORG | 0 | 0 | NA | 2 | 0 | orphan | 0 | 0 | NA | 0 | 0 | NA | 78.1 | 0 | 0.98 |
| AG-461-K07 | GORG | 0 | 0 | NA | 1 | 0 | orphan | 0 | 0 | NA | 0 | 1 | orphan | 78.08 | 0 | 0.83 |
| AG-914-D07 | GORG | 0 | 0 | NA | 1 | 0 | orphan | 0 | 1 | NA | 0 | 0 | NA | 78.07 | 0 | 1.05 |
| AH-287-C09 | GORG | 0 | 0 | NA | 2 | 0 | orphan | 0 | 0 | NA | 0 | 0 | NA | 78.07 | 0 | 1.01 |
| AG-895-O20 | GORG | 0 | 0 | NA | 1 | 0 | orphan | 0 | 0 | NA | 0 | 0 | NA | 78.02 | 0 | 0.92 |
| AG-439-O06 | GORG | 0 | 0 | NA | 3 | 0 | orphan | 0 | 0 | NA | 0 | 0 | NA | 77.99 | 0 | 0.91 |
| AG-900-E06 | GORG | 0 | 0 | NA | 1 | 0 | orphan | 0 | 0 | NA | 0 | 0 | NA | 77.97 | 0 | 1.3 |
| AG-916-F23 | GORG | 0 | 0 | NA | 2 | 0 | orphan | 0 | 0 | NA | 0 | 1 | orphan | 77.96 | 0 | 1 |
| AG-435-G19 | GORG | 0 | 0 | NA | 1 | 0 | orphan | 0 | 0 | NA | 0 | 0 | NA | 77.94 | 0 | 1.02 |
| AG-916-A14 | GORG | 0 | 0 | NA | 2 | 0 | orphan | 0 | 0 | NA | 0 | 0 | NA | 77.94 | 0 | 0.94 |
| AG-916-F16 | GORG | 0 | 0 | NA | 1 | 0 | orphan | 1 | 1 | complete | 0 | 0 | NA | 77.94 | 0 | 1.25 |
| AG-360-M22 | GORG | 0 | 0 | NA | 1 | 0 | orphan | 0 | 0 | NA | 0 | 0 | NA | 77.91 | 0 | 0.98 |
| AH-324-E14 | GORG | 1 | 1 | complete | 1 | 0 | orphan | 0 | 0 | NA | 0 | 0 | NA | 77.91 | 0 | 1.01 |
| AG-333-D23 | GORG | 0 | 0 | NA | 4 | 0 | orphan | 0 | 0 | NA | 0 | 0 | NA | 77.89 | 0 | 1.18 |
| AG-337-G05 | GORG | 0 | 0 | NA | 1 | 0 | orphan | 0 | 0 | NA | 0 | 0 | NA | 77.89 | 0 | 1 |
| AG-447-E20 | GORG | 0 | 0 | NA | 1 | 0 | orphan | 0 | 0 | NA | 0 | 0 | NA | 77.82 | 0 | 1.08 |
| AG-333-N15 | GORG | 0 | 0 | NA | 1 | 0 | orphan | 0 | 0 | NA | 0 | 0 | NA | 77.8 | 0 | 0.99 |
| AG-447-F03 | GORG | 0 | 0 | NA | 1 | 0 | orphan | 0 | 0 | NA | 0 | 0 | NA | 77.8 | 0 | 0.94 |
| AG-911-E04 | GORG | 0 | 0 | NA | 1 | 0 | orphan | 0 | 0 | NA | 0 | 0 | NA | 77.78 | 0 | 1.09 |

| | | | | | | | | | | | | | | | | |
|---|---|---|---|---|---|---|---|---|---|---|---|---|---|---|---|---|
| AG-426-A05 | GORG | 0 | 0 | NA | 1 | 0 | orphan | 0 | 1 | NA | 0 | 0 | NA | 77.76 | 0 | 1.09 |
| AG-435-N04 | GORG | 0 | 0 | NA | 1 | 0 | orphan | 0 | 0 | NA | 0 | 0 | NA | 77.75 | 0 | 0.86 |
| AG-917-P19 | GORG | 0 | 0 | NA | 1 | 0 | orphan | 0 | 0 | NA | 0 | 0 | NA | 77.75 | 0 | 0.97 |
| AG-911-F11 | GORG | 0 | 0 | NA | 1 | 0 | orphan | 0 | 0 | NA | 0 | 0 | NA | 77.73 | 0 | 1.01 |
| AG-891-F06 | GORG | 0 | 0 | NA | 2 | 0 | orphan | 0 | 0 | NA | 0 | 0 | NA | 77.72 | 0 | 2.14 |
| AG-439-O15 | GORG | 0 | 0 | NA | 2 | 0 | orphan | 0 | 0 | NA | 0 | 0 | NA | 77.71 | 0 | 1.07 |
| AG-901-B05 | GORG | 0 | 0 | NA | 3 | 0 | orphan | 0 | 0 | NA | 0 | 0 | NA | 77.71 | 0 | 0.86 |
| AG-920-K10 | GORG | 0 | 0 | NA | 1 | 0 | orphan | 0 | 0 | NA | 0 | 0 | NA | 77.71 | 0 | 1.38 |
| AH-321-F23 | GORG | 0 | 0 | NA | 2 | 0 | orphan | 0 | 0 | NA | 0 | 0 | NA | 77.71 | 0 | 0.92 |
| AG-899-E11 | GORG | 0 | 0 | NA | 2 | 0 | orphan | 0 | 0 | NA | 0 | 0 | NA | 77.69 | 0 | 1.93 |
| AG-430-F10 | GORG | 0 | 0 | NA | 1 | 0 | orphan | 0 | 0 | NA | 0 | 0 | NA | 77.66 | 0 | 0.85 |
| AG-898-I03 | GORG | 0 | 0 | NA | 1 | 0 | orphan | 0 | 0 | NA | 0 | 0 | NA | 77.6 | 0 | 1.35 |
| AG-917-F11 | GORG | 0 | 0 | NA | 2 | 0 | orphan | 0 | 0 | NA | 0 | 0 | NA | 77.6 | 0 | 1.01 |
| AG-349-O22 | GORG | 0 | 0 | NA | 1 | 0 | orphan | 0 | 0 | NA | 0 | 0 | NA | 77.55 | 0 | 1.02 |
| AH-321-P23 | GORG | 1 | 1 | complete | 0 | 0 | NA | 0 | 0 | NA | 0 | 0 | NA | 77.55 | 0 | 1.16 |
| AG-896-M03 | GORG | 0 | 0 | NA | 1 | 0 | orphan | 1 | 1 | complete | 0 | 0 | NA | 77.53 | 0 | 1.97 |
| AG-913-G03 | GORG | 0 | 0 | NA | 1 | 0 | orphan | 0 | 0 | NA | 0 | 0 | NA | 77.52 | 0 | 0.99 |
| AG-390-F15 | GORG | 0 | 0 | NA | 1 | 0 | orphan | 0 | 0 | NA | 0 | 0 | NA | 77.51 | 0 | 1.23 |
| AG-911-A21 | GORG | 0 | 0 | NA | 1 | 0 | orphan | 0 | 0 | NA | 0 | 0 | NA | 77.51 | 0 | 0.76 |
| AG-920-G17 | GORG | 0 | 0 | NA | 2 | 0 | orphan | 0 | 0 | NA | 0 | 0 | NA | 77.51 | 0 | 0.8 |
| AG-337-O05 | GORG | 0 | 0 | NA | 1 | 0 | orphan | 0 | 0 | NA | 0 | 0 | NA | 77.46 | 0 | 1.06 |
| AG-896-K17 | GORG | 0 | 0 | NA | 2 | 0 | orphan | 0 | 0 | NA | 0 | 0 | NA | 77.46 | 0 | 0.9 |
| AG-910-L19 | GORG | 0 | 0 | NA | 1 | 0 | orphan | 0 | 0 | NA | 0 | 0 | NA | 77.38 | 0 | 0.97 |
| AG-914-E21 | GORG | 0 | 0 | NA | 4 | 0 | orphan | 0 | 0 | NA | 0 | 0 | NA | 77.38 | 0 | 0.94 |
| AG-439-P01 | GORG | 0 | 0 | NA | 1 | 0 | orphan | 0 | 0 | NA | 0 | 0 | NA | 77.36 | 0 | 0.93 |
| AG-896-G08 | GORG | 0 | 0 | NA | 1 | 0 | orphan | 0 | 0 | NA | 0 | 0 | NA | 77.36 | 0 | 1.27 |

| | | | | | | | | | | | | | | | | |
|---|---|---|---|---|---|---|---|---|---|---|---|---|---|---|---|---|
| AG-903-M09 | GORG | 1 | 1 | complete | 1 | 0 | orphan | 0 | 0 | NA | 0 | 0 | NA | 77.36 | 0 | 1.15 |
| AG-390-N18 | GORG | 0 | 0 | NA | 3 | 0 | orphan | 0 | 0 | NA | 0 | 0 | NA | 77.34 | 0 | 1.08 |
| AG-430-P01 | GORG | 0 | 0 | NA | 1 | 0 | orphan | 0 | 0 | NA | 0 | 0 | NA | 77.29 | 0 | 0.79 |
| AG-913-C04 | GORG | 0 | 0 | NA | 0 | 0 | NA | 0 | 0 | NA | 0 | 1 | orphan | 77.26 | 0 | 1.26 |
| AG-911-B02 | GORG | 0 | 0 | NA | 2 | 0 | orphan | 0 | 0 | NA | 0 | 0 | NA | 77.25 | 0 | 1.02 |
| AG-904-D14 | GORG | 0 | 0 | NA | 1 | 0 | orphan | 0 | 0 | NA | 0 | 0 | NA | 77.22 | 0 | 1.15 |
| AG-904-P17 | GORG | 0 | 0 | NA | 2 | 0 | orphan | 0 | 0 | NA | 0 | 2 | orphan | 77.21 | 0 | 1.25 |
| AG-912-I18 | GORG | 0 | 0 | NA | 1 | 0 | orphan | 0 | 0 | NA | 0 | 1 | orphan | 77.17 | 0 | 0.86 |
| AG-920-O10 | GORG | 0 | 0 | NA | 1 | 0 | orphan | 0 | 0 | NA | 0 | 0 | NA | 77.15 | 0 | 1.04 |
| AH-287-A18 | GORG | 0 | 0 | NA | 1 | 0 | orphan | 0 | 0 | NA | 0 | 0 | NA | 77.14 | 0 | 0.94 |
| AG-893-M02 | GORG | 0 | 0 | NA | 0 | 0 | NA | 0 | 0 | NA | 0 | 1 | orphan | 77.12 | 0 | 1.19 |
| AG-430-D17 | GORG | 0 | 0 | NA | 1 | 0 | orphan | 0 | 0 | NA | 0 | 0 | NA | 77.12 | 0 | 0.81 |
| AG-912-A14 | GORG | 0 | 0 | NA | 1 | 0 | orphan | 0 | 0 | NA | 0 | 1 | orphan | 77.11 | 0 | 1.09 |
| AG-414-I04 | GORG | 0 | 0 | NA | 2 | 0 | orphan | 0 | 0 | NA | 0 | 0 | NA | 77.11 | 0 | 1.02 |
| AG-414-M09 | GORG | 0 | 0 | NA | 1 | 0 | orphan | 0 | 0 | NA | 0 | 0 | NA | 77.11 | 0 | 0.83 |
| AG-426-P23 | GORG | 0 | 0 | NA | 2 | 0 | orphan | 0 | 0 | NA | 0 | 0 | NA | 77.11 | 0 | 1.01 |
| AG-891-E15 | GORG | 0 | 0 | NA | 1 | 0 | orphan | 0 | 0 | NA | 0 | 0 | NA | 77.11 | 0 | 1.36 |
| AG-892-G22 | GORG | 0 | 0 | NA | 2 | 0 | orphan | 0 | 0 | NA | 0 | 0 | NA | 77.11 | 0 | 1.17 |
| AG-909-M20 | GORG | 0 | 0 | NA | 2 | 0 | orphan | 0 | 0 | NA | 0 | 0 | NA | 77.11 | 0 | 0.88 |
| AG-914-P04 | GORG | 1 | 1 | complete | 1 | 0 | orphan | 0 | 0 | NA | 0 | 0 | NA | 77.11 | 0 | 1.07 |
| AG-917-E07 | GORG | 0 | 0 | NA | 1 | 0 | orphan | 0 | 0 | NA | 0 | 0 | NA | 77.11 | 0 | 0.93 |
| AG-919-O06 | GORG | 0 | 0 | NA | 4 | 0 | orphan | 0 | 0 | NA | 0 | 0 | NA | 77.11 | 0 | 0.77 |
| AG-919-P09 | GORG | 1 | 1 | complete | 0 | 0 | NA | 0 | 0 | NA | 0 | 0 | NA | 77.11 | 0 | 1.15 |
| AG-920-C15 | GORG | 0 | 0 | NA | 2 | 0 | orphan | 0 | 0 | NA | 0 | 0 | NA | 77.11 | 0 | 1.05 |

| | | | | | | | | | | | | | | | | |
|---|---|---|---|---|---|---|---|---|---|---|---|---|---|---|---|---|
| AG-349-M02 | GORG | 0 | 0 | NA | 1 | 0 | orphan | 0 | 0 | NA | 0 | 1 | orpha n | 77.1 | 0 | 1.24 |
| AG-439-B17 | GORG | 1 | 1 | complet e | 0 | 0 | NA | 0 | 0 | NA | 0 | 0 | NA | 77.1 | 0 | 1.27 |
| AG-896-L17 | GORG | 0 | 0 | NA | 2 | 0 | orphan | 0 | 0 | NA | 0 | 0 | NA | 77.09 | 0 | 0.95 |
| AG-904-J10 | GORG | 0 | 0 | NA | 1 | 0 | orphan | 0 | 0 | NA | 0 | 0 | NA | 77.09 | 0 | 1 |
| AG-903-J17 | GORG | 0 | 0 | NA | 2 | 0 | orphan | 0 | 0 | NA | 0 | 0 | NA | 77.01 | 0 | 0.9 |
| AG-899-B06 | GORG | 0 | 0 | NA | 1 | 0 | orphan | 0 | 0 | NA | 0 | 0 | NA | 77 | 0 | 1.4 |
| AH-321-K03 | GORG | 0 | 0 | NA | 1 | 0 | orphan | 0 | 0 | NA | 0 | 1 | orpha n | 76.99 | 0 | 1.24 |
| AG-461-P02 | GORG | 0 | 0 | NA | 0 | 0 | NA | 0 | 0 | NA | 0 | 1 | orpha n | 76.95 | 0 | 1.25 |
| AG-891-I13 | GORG | 0 | 0 | NA | 1 | 0 | orphan | 0 | 0 | NA | 0 | 0 | NA | 76.94 | 0 | 0.9 |
| AG-898-E05 | GORG | 0 | 0 | NA | 1 | 0 | orphan | 0 | 0 | NA | 0 | 0 | NA | 76.93 | 0 | 1.18 |
| AG-404-A03 | GORG | 0 | 0 | NA | 1 | 0 | orphan | 0 | 0 | NA | 0 | 0 | NA | 76.92 | 0 | 1.06 |
| AG-893-L10 | GORG | 0 | 0 | NA | 2 | 0 | orphan | 0 | 0 | NA | 0 | 0 | NA | 76.9 | 0 | 1.96 |
| AG-390-K13 | GORG | 0 | 0 | NA | 1 | 0 | orphan | 0 | 0 | NA | 0 | 0 | NA | 76.87 | 0 | 0.99 |
| AG-461-E21 | GORG | 0 | 0 | NA | 1 | 0 | orphan | 0 | 0 | NA | 0 | 0 | NA | 76.74 | 0 | 1.51 |
| AG-908-E21 | GORG | 0 | 0 | NA | 1 | 0 | orphan | 0 | 0 | NA | 0 | 0 | NA | 76.73 | 0 | 0.94 |
| AG-894-L18 | GORG | 0 | 0 | NA | 1 | 0 | orphan | 0 | 0 | NA | 0 | 0 | NA | 76.72 | 0 | 1.68 |
| AG-916-F06 | GORG | 0 | 0 | NA | 1 | 0 | orphan | 0 | 0 | NA | 0 | 0 | NA | 76.71 | 0 | 0.92 |
| AG-447-N02 | GORG | 2 | 1 | complet e | 1 | 0 | orphan | 0 | 0 | NA | 0 | 0 | NA | 76.7 | 0 | 1.09 |
| AG-470-M04 | GORG | 0 | 0 | NA | 2 | 0 | orphan | 0 | 0 | NA | 0 | 0 | NA | 76.67 | 0 | 0.86 |
| AG-903-E10 | GORG | 1 | 1 | complet e | 0 | 0 | NA | 0 | 0 | NA | 0 | 1 | orpha n | 76.62 | 0 | 1.2 |
| AG-920-P10 | GORG | 0 | 0 | NA | 1 | 0 | orphan | 0 | 0 | NA | 0 | 0 | NA | 76.58 | 0 | 0.87 |
| AG-894-A03 | GORG | 0 | 0 | NA | 2 | 0 | orphan | 0 | 0 | NA | 0 | 0 | NA | 76.56 | 0 | 1.25 |
| AG-909-A05 | GORG | 0 | 0 | NA | 1 | 0 | orphan | 0 | 0 | NA | 0 | 0 | NA | 76.56 | 0 | 0.9 |
| AG-892-L17 | GORG | 0 | 0 | NA | 1 | 0 | orphan | 0 | 0 | NA | 0 | 0 | NA | 76.54 | 0 | 1.08 |
| AG-903-C04 | GORG | 0 | 0 | NA | 1 | 0 | orphan | 0 | 0 | NA | 0 | 0 | NA | 76.54 | 0 | 1.03 |

| | | | | | | | | | | | | | | | | |
|---|---|---|---|---|---|---|---|---|---|---|---|---|---|---|---|---|
| AG-910-C11 | GORG | 0 | 0 | NA | 1 | 0 | orphan | 0 | 0 | NA | 0 | 0 | NA | 76.54 | 0 | 0.62 |
| AG-337-A23 | GORG | 0 | 0 | NA | 1 | 0 | orphan | 0 | 0 | NA | 0 | 0 | NA | 76.53 | 0 | 0.87 |
| AG-919-F22 | GORG | 0 | 0 | NA | 1 | 0 | orphan | 0 | 0 | NA | 0 | 0 | NA | 76.52 | 0 | 1.09 |
| AG-915-O08 | GORG | 0 | 0 | NA | 1 | 0 | orphan | 0 | 0 | NA | 0 | 1 | orpha n | 76.51 | 0 | 1.16 |
| AG-337-C09 | GORG | 1 | 0 | orphan | 1 | 0 | orphan | 0 | 0 | NA | 0 | 0 | NA | 76.51 | 0 | 1.21 |
| AG-337-L16 | GORG | 0 | 0 | NA | 2 | 0 | orphan | 0 | 0 | NA | 0 | 0 | NA | 76.51 | 0 | 1.14 |
| AG-414-D15 | GORG | 0 | 0 | NA | 4 | 0 | orphan | 0 | 2 | NA | 0 | 0 | NA | 76.51 | 0 | 0.99 |
| AG-453-N02 | GORG | 0 | 0 | NA | 1 | 0 | orphan | 0 | 0 | NA | 0 | 0 | NA | 76.51 | 0 | 1.25 |
| AG-895-D04 | GORG | 0 | 0 | NA | 1 | 0 | orphan | 0 | 0 | NA | 0 | 0 | NA | 76.51 | 0 | 1.41 |
| AG-910-A06 | GORG | 0 | 0 | NA | 1 | 0 | orphan | 0 | 0 | NA | 0 | 0 | NA | 76.51 | 0 | 1.01 |
| AH-324-C16 | GORG | 0 | 0 | NA | 2 | 0 | orphan | 0 | 0 | NA | 0 | 0 | NA | 76.51 | 0 | 0.99 |
| AH-326-B08 | GORG | 0 | 0 | NA | 1 | 0 | orphan | 0 | 0 | NA | 0 | 0 | NA | 76.51 | 0 | 1.02 |
| AG-349-O20 | GORG | 0 | 0 | NA | 1 | 0 | orphan | 0 | 0 | NA | 0 | 0 | NA | 76.5 | 0 | 1.19 |
| AG-896-E15 | GORG | 0 | 0 | NA | 2 | 1 | complet e | 0 | 0 | NA | 0 | 0 | NA | 76.47 | 0 | 1.12 |
| AG-913-M18 | GORG | 0 | 0 | NA | 1 | 0 | orphan | 0 | 0 | NA | 0 | 0 | NA | 76.47 | 0 | 1.75 |
| AG-404-O05 | GORG | 0 | 0 | NA | 2 | 0 | orphan | 0 | 0 | NA | 0 | 0 | NA | 76.45 | 0 | 1.08 |
| AG-915-J16 | GORG | 0 | 0 | NA | 1 | 0 | orphan | 0 | 0 | NA | 0 | 0 | NA | 76.45 | 0 | 0.92 |
| AG-905-O07 | GORG | 0 | 0 | NA | 2 | 0 | orphan | 0 | 0 | NA | 0 | 0 | NA | 76.42 | 0 | 1.1 |
| AG-422-C22 | GORG | 0 | 0 | NA | 1 | 0 | orphan | 0 | 0 | NA | 0 | 0 | NA | 76.4 | 0 | 0.92 |
| AG-898-E11 | GORG | 0 | 0 | NA | 1 | 0 | orphan | 0 | 1 | NA | 0 | 0 | NA | 76.31 | 0 | 1.12 |
| AG-903-O18 | GORG | 0 | 0 | NA | 1 | 0 | orphan | 0 | 0 | NA | 0 | 0 | NA | 76.31 | 0 | 0.85 |
| AG-892-A11 | GORG | 0 | 0 | NA | 1 | 0 | orphan | 0 | 0 | NA | 0 | 0 | NA | 76.28 | 0 | 0.97 |
| AG-909-P19 | GORG | 0 | 0 | NA | 1 | 0 | orphan | 0 | 0 | NA | 0 | 0 | NA | 76.28 | 0 | 0.77 |
| AG-914-F18 | GORG | 0 | 0 | NA | 1 | 0 | orphan | 0 | 0 | NA | 0 | 0 | NA | 76.26 | 0 | 1.16 |
| AG-904-A06 | GORG | 2 | 2 | complet e | 0 | 0 | NA | 0 | 0 | NA | 0 | 0 | NA | 76.2 | 0 | 1.13 |
| AG-912-N08 | GORG | 0 | 0 | NA | 1 | 0 | orphan | 0 | 0 | NA | 0 | 0 | NA | 76.19 | 0 | 1.05 |

| | | | | | | | | | | | | | | | | |
|---|---|---|---|---|---|---|---|---|---|---|---|---|---|---|---|---|
| AH-287-G22 | GORG | 0 | 0 | NA | 1 | 0 | orphan | 0 | 0 | NA | 0 | 0 | NA | 76.19 | 0 | 0.96 |
| AG-414-N03 | GORG | 0 | 0 | NA | 3 | 0 | orphan | 0 | 0 | NA | 0 | 0 | NA | 76.15 | 0 | 0.86 |
| AG-464-K04 | GORG | 0 | 0 | NA | 1 | 0 | orphan | 0 | 0 | NA | 0 | 0 | NA | 76.15 | 0 | 0.92 |
| AG-896-M04 | GORG | 0 | 0 | NA | 1 | 0 | orphan | 0 | 0 | NA | 0 | 0 | NA | 76.15 | 0 | 1.02 |
| AH-324-G13 | GORG | 0 | 0 | NA | 1 | 0 | orphan | 0 | 0 | NA | 0 | 0 | NA | 76.15 | 0 | 0.98 |
| AG-918-G14 | GORG | 0 | 0 | NA | 2 | 0 | orphan | 0 | 0 | NA | 0 | 0 | NA | 76.11 | 0 | 0.97 |
| AG-430-F07 | GORG | 0 | 0 | NA | 1 | 0 | orphan | 0 | 0 | NA | 0 | 0 | NA | 76.09 | 0 | 0.92 |
| AG-390-A22 | GORG | 0 | 0 | NA | 1 | 0 | orphan | 0 | 0 | NA | 0 | 0 | NA | 76.07 | 0 | 0.78 |
| AG-457-M22 | GORG | 0 | 0 | NA | 1 | 0 | orphan | 0 | 0 | NA | 0 | 0 | NA | 76.06 | 0 | 0.82 |
| AG-900-D23 | GORG | 0 | 0 | NA | 2 | 0 | orphan | 0 | 1 | NA | 0 | 0 | NA | 76.05 | 0 | 1.18 |
| AG-899-K21 | GORG | 0 | 0 | NA | 1 | 0 | orphan | 0 | 0 | NA | 0 | 0 | NA | 75.96 | 0 | 1.16 |
| AG-333-O05 | GORG | 0 | 0 | NA | 1 | 0 | orphan | 0 | 0 | NA | 0 | 0 | NA | 75.9 | 0 | 0.81 |
| AG-426-J21 | GORG | 0 | 0 | NA | 2 | 0 | orphan | 0 | 0 | NA | 0 | 0 | NA | 75.9 | 0 | 1.09 |
| AG-457-L11 | GORG | 0 | 0 | NA | 1 | 0 | orphan | 0 | 0 | NA | 0 | 0 | NA | 75.9 | 0 | 1.07 |
| AG-891-B11 | GORG | 0 | 0 | NA | 1 | 0 | orphan | 0 | 0 | NA | 0 | 0 | NA | 75.9 | 0 | 1.12 |
| AG-894-N17 | GORG | 0 | 0 | NA | 2 | 0 | orphan | 0 | 0 | NA | 0 | 0 | NA | 75.9 | 0 | 1.16 |
| AG-895-J17 | GORG | 0 | 0 | NA | 1 | 0 | orphan | 0 | 0 | NA | 0 | 0 | NA | 75.9 | 0 | 0.96 |
| AG-898-N21 | GORG | 0 | 0 | NA | 1 | 0 | orphan | 0 | 0 | NA | 0 | 0 | NA | 75.9 | 0 | 1.08 |
| AG-899-O02 | GORG | 0 | 0 | NA | 1 | 0 | orphan | 0 | 0 | NA | 0 | 0 | NA | 75.9 | 0 | 1.14 |
| AG-904-D18 | GORG | 0 | 0 | NA | 1 | 0 | orphan | 0 | 0 | NA | 0 | 0 | NA | 75.9 | 0 | 1.17 |
| AG-917-G13 | GORG | 1 | 1 | complete | 1 | 0 | orphan | 0 | 0 | NA | 0 | 0 | NA | 75.9 | 0 | 1.22 |
| AG-918-A19 | GORG | 0 | 0 | NA | 1 | 0 | orphan | 0 | 0 | NA | 0 | 0 | NA | 75.9 | 0 | 0.94 |
| AG-919-K16 | GORG | 0 | 0 | NA | 1 | 0 | orphan | 0 | 0 | NA | 0 | 0 | NA | 75.9 | 0 | 1.08 |
| AG-900-B23 | GORG | 0 | 0 | NA | 1 | 0 | orphan | 0 | 0 | NA | 0 | 0 | NA | 75.87 | 0 | 0.87 |
| AG-912-P16 | GORG | 0 | 0 | NA | 1 | 0 | orphan | 0 | 0 | NA | 0 | 0 | NA | 75.86 | 0 | 1 |
| AG-349-N10 | GORG | 0 | 0 | NA | 0 | 0 | NA | 0 | 0 | NA | 0 | 1 | orphan | 75.84 | 0 | 1.15 |

| | | | | | | | | | | | | | | | | |
|---|---|---|---|---|---|---|---|---|---|---|---|---|---|---|---|---|
| AG-453-L05 | GORG | 0 | 0 | NA | 1 | 0 | orphan | 0 | 0 | NA | 0 | 0 | NA | 75.82 | 0 | 0.71 |
| AG-904-P13 | GORG | 0 | 0 | NA | 1 | 0 | orphan | 0 | 0 | NA | 0 | 0 | NA | 75.79 | 0 | 1.02 |
| AG-891-E04 | GORG | 1 | 0 | orphan | 1 | 0 | orphan | 0 | 0 | NA | 0 | 0 | NA | 75.78 | 0 | 0.99 |
| AG-325-D16 | GORG | 0 | 0 | NA | 1 | 0 | orphan | 0 | 0 | NA | 0 | 0 | NA | 75.74 | 0 | 1.26 |
| AG-917-F06 | GORG | 0 | 0 | NA | 1 | 0 | orphan | 0 | 0 | NA | 0 | 0 | NA | 75.74 | 0 | 0.95 |
| AG-404-F07 | GORG | 0 | 0 | NA | 1 | 0 | orphan | 0 | 0 | NA | 0 | 0 | NA | 75.73 | 0 | 1.29 |
| AG-359-I17 | GORG | 0 | 0 | NA | 2 | 0 | orphan | 0 | 0 | NA | 0 | 0 | NA | 75.69 | 0 | 0.99 |
| AG-908-P04 | GORG | 0 | 0 | NA | 1 | 0 | orphan | 0 | 0 | NA | 0 | 0 | NA | 75.66 | 0 | 1.24 |
| AG-917-A18 | GORG | 0 | 0 | NA | 2 | 0 | orphan | 0 | 0 | NA | 0 | 0 | NA | 75.65 | 0 | 1.07 |
| AG-426-E03 | GORG | 1 | 0 | orphan | 1 | 0 | orphan | 0 | 0 | NA | 0 | 0 | NA | 75.64 | 0 | 0.99 |
| AG-908-M04 | GORG | 0 | 0 | NA | 1 | 0 | orphan | 0 | 0 | NA | 0 | 0 | NA | 75.64 | 0 | 0.84 |
| AG-908-E18 | GORG | 0 | 0 | NA | 1 | 0 | orphan | 0 | 0 | NA | 0 | 0 | NA | 75.58 | 0 | 0.84 |
| AG-918-A01 | GORG | 0 | 0 | NA | 1 | 0 | orphan | 0 | 0 | NA | 0 | 0 | NA | 75.56 | 0 | 1 |
| AG-915-E10 | GORG | 0 | 0 | NA | 2 | 0 | orphan | 0 | 0 | NA | 0 | 0 | NA | 75.5 | 0 | 0.93 |
| AG-915-K10 | GORG | 0 | 0 | NA | 1 | 0 | orphan | 0 | 0 | NA | 0 | 0 | NA | 75.5 | 0 | 0.94 |
| AG-333-K07 | GORG | 0 | 0 | NA | 1 | 0 | orphan | 0 | 0 | NA | 0 | 0 | NA | 75.47 | 0 | 1.11 |
| AG-390-P07 | GORG | 0 | 0 | NA | 1 | 0 | orphan | 0 | 0 | NA | 0 | 0 | NA | 75.47 | 0 | 1.24 |
| AG-422-K08 | GORG | 0 | 0 | NA | 1 | 0 | orphan | 0 | 0 | NA | 0 | 0 | NA | 75.47 | 0 | 0.88 |
| AG-453-A09 | GORG | 0 | 0 | NA | 1 | 0 | orphan | 0 | 0 | NA | 0 | 0 | NA | 75.47 | 0 | 0.93 |
| AG-893-J21 | GORG | 0 | 0 | NA | 1 | 0 | orphan | 0 | 0 | NA | 0 | 0 | NA | 75.47 | 0 | 0.71 |
| AG-908-B06 | GORG | 0 | 0 | NA | 1 | 0 | orphan | 0 | 0 | NA | 0 | 0 | NA | 75.47 | 0 | 0.88 |
| AG-913-N11 | GORG | 0 | 0 | NA | 1 | 0 | orphan | 0 | 0 | NA | 0 | 0 | NA | 75.47 | 0 | 1.12 |
| AG-895-M05 | GORG | 0 | 0 | NA | 0 | 0 | NA | 0 | 0 | NA | 0 | 1 | orphan | 75.43 | 0 | 1.24 |
| AH-287-C20 | GORG | 2 | 2 | complete | 0 | 0 | NA | 0 | 0 | NA | 0 | 1 | orphan | 75.41 | 0 | 1.23 |
| AG-900-G06 | GORG | 0 | 0 | NA | 1 | 0 | orphan | 0 | 0 | NA | 0 | 0 | NA | 75.41 | 0 | 1.14 |
| AG-918-F23 | GORG | 0 | 0 | NA | 2 | 0 | orphan | 0 | 0 | NA | 0 | 0 | NA | 75.39 | 0 | 0.97 |

| | | | | | | | | | | | | | | | | |
|---|---|---|---|---|---|---|---|---|---|---|---|---|---|---|---|---|
| AG-422-M21 | GORG | 0 | 0 | NA | 1 | 0 | orphan | 0 | 0 | NA | 0 | 0 | NA | 75.38 | 0 | 1.23 |
| AG-426-F21 | GORG | 2 | 0 | orphan | 1 | 0 | orphan | 0 | 0 | NA | 0 | 0 | NA | 75.33 | 0 | 0.91 |
| AG-898-P10 | GORG | 0 | 0 | NA | 1 | 0 | orphan | 0 | 0 | NA | 0 | 0 | NA | 75.3 | 0 | 0.98 |
| AG-905-C11 | GORG | 0 | 0 | NA | 1 | 0 | orphan | 0 | 0 | NA | 0 | 0 | NA | 75.3 | 0 | 0.93 |
| AG-917-C05 | GORG | 0 | 0 | NA | 2 | 0 | orphan | 0 | 0 | NA | 0 | 0 | NA | 75.3 | 0 | 0.88 |
| AH-326-M11 | GORG | 0 | 0 | NA | 1 | 1 | orphan | 0 | 0 | NA | 0 | 0 | NA | 75.3 | 0 | 0.78 |
| AG-894-M20 | GORG | 0 | 0 | NA | 0 | 0 | NA | 0 | 0 | NA | 0 | 1 | orphan | 75.27 | 0 | 1.2 |
| AG-914-F19 | GORG | 0 | 0 | NA | 2 | 0 | orphan | 0 | 0 | NA | 0 | 0 | NA | 75.26 | 0 | 1.13 |
| AG-913-J11 | GORG | 0 | 0 | NA | 1 | 0 | orphan | 0 | 0 | NA | 0 | 0 | NA | 75.23 | 0 | 1.18 |
| AG-426-F02 | GORG | 0 | 0 | NA | 2 | 0 | orphan | 0 | 0 | NA | 0 | 0 | NA | 75.2 | 0 | 1.35 |
| AG-903-O04 | GORG | 1 | 1 | complete | 1 | 0 | orphan | 0 | 0 | NA | 0 | 0 | NA | 75.19 | 0 | 0.85 |
| AG-461-J19 | GORG | 0 | 0 | NA | 1 | 0 | orphan | 0 | 0 | NA | 0 | 0 | NA | 75.17 | 0 | 0.81 |
| AG-447-F16 | GORG | 0 | 0 | NA | 2 | 0 | orphan | 0 | 0 | NA | 0 | 0 | NA | 75.15 | 0 | 1.03 |
| AG-912-P01 | GORG | 0 | 0 | NA | 2 | 0 | orphan | 0 | 0 | NA | 0 | 0 | NA | 75.12 | 0 | 0.9 |
| AG-912-P21 | GORG | 0 | 0 | NA | 1 | 0 | orphan | 0 | 0 | NA | 0 | 0 | NA | 75.1 | 0 | 0.79 |
| AG-918-I15 | GORG | 0 | 0 | NA | 1 | 0 | orphan | 0 | 0 | NA | 0 | 0 | NA | 75.05 | 0 | 0.93 |
| AG-422-E20 | GORG | 0 | 0 | NA | 1 | 0 | orphan | 0 | 0 | NA | 0 | 0 | NA | 75.04 | 0 | 0.84 |
| AG-461-O10 | GORG | 0 | 0 | NA | 1 | 0 | orphan | 0 | 0 | NA | 0 | 0 | NA | 75.04 | 0 | 0.88 |
| AG-911-I23 | GORG | 0 | 0 | NA | 1 | 0 | orphan | 0 | 0 | NA | 0 | 0 | NA | 75.04 | 0 | 1.12 |
| AG-390-C05 | GORG | 0 | 0 | NA | 1 | 0 | orphan | 0 | 0 | NA | 0 | 0 | NA | 75 | 0 | 1.21 |
| AG-912-C06 | GORG | 1 | 1 | complete | 0 | 0 | NA | 0 | 0 | NA | 0 | 0 | NA | 75 | 0 | 1.36 |
| AG-904-K09 | GORG | 0 | 0 | NA | 1 | 0 | orphan | 0 | 0 | NA | 0 | 0 | NA | 74.95 | 0 | 1.08 |
| AG-915-N05 | GORG | 0 | 0 | NA | 1 | 0 | orphan | 0 | 0 | NA | 0 | 0 | NA | 74.9 | 0 | 1.12 |
| AG-912-C16 | GORG | 0 | 0 | NA | 2 | 0 | orphan | 0 | 0 | NA | 0 | 0 | NA | 74.88 | 0 | 0.97 |
| AG-349-D14 | GORG | 0 | 0 | NA | 0 | 0 | NA | 0 | 0 | NA | 0 | 1 | orphan | 74.82 | 0 | 1.2 |
| AG-916-E22 | GORG | 0 | 1 | NA | 0 | 0 | NA | 0 | 0 | NA | 0 | 0 | NA | 74.82 | 0 | 1.22 |

| | | | | | | | | | | | | | | | |
|---|---|---|---|---|---|---|---|---|---|---|---|---|---|---|---|
| AG-365-O07 | GORG | 0 | 0 | NA | 1 | 0 | orphan | 0 | 0 | NA | 0 | 0 | NA | 74.8 | 0 | 1.29 |
| AG-900-M07 | GORG | 0 | 1 | NA | 2 | 1 | complete | 0 | 0 | NA | 0 | 0 | NA | 74.79 | 0 | 1.28 |
| AH-326-C13 | GORG | 1 | 1 | complete | 2 | 0 | orphan | 0 | 0 | NA | 0 | 0 | NA | 74.73 | 0 | 1.04 |
| AG-325-J11 | GORG | 0 | 0 | NA | 1 | 0 | orphan | 0 | 0 | NA | 0 | 0 | NA | 74.71 | 0 | 1.08 |
| AG-333-L20 | GORG | 0 | 0 | NA | 1 | 0 | orphan | 0 | 0 | NA | 0 | 0 | NA | 74.7 | 0 | 1 |
| AG-430-E16 | GORG | 0 | 0 | NA | 1 | 0 | orphan | 0 | 0 | NA | 0 | 0 | NA | 74.7 | 0 | 1.29 |
| AG-904-C07 | GORG | 0 | 0 | NA | 1 | 0 | orphan | 0 | 0 | NA | 0 | 0 | NA | 74.7 | 0 | 1.14 |
| AG-908-K09 | GORG | 0 | 0 | NA | 1 | 0 | orphan | 0 | 0 | NA | 0 | 0 | NA | 74.7 | 0 | 0.87 |
| AG-911-A16 | GORG | 0 | 0 | NA | 1 | 0 | orphan | 0 | 0 | NA | 0 | 0 | NA | 74.7 | 0 | 0.94 |
| AG-914-A17 | GORG | 0 | 0 | NA | 2 | 2 | complete | 0 | 0 | NA | 0 | 0 | NA | 74.7 | 0 | 1.03 |
| AG-920-A01 | GORG | 0 | 0 | NA | 1 | 0 | orphan | 0 | 0 | NA | 0 | 0 | NA | 74.69 | 0 | 0.88 |
| AG-349-C06 | GORG | 0 | 0 | NA | 2 | 0 | orphan | 0 | 0 | NA | 0 | 0 | NA | 74.68 | 0 | 0.88 |
| AG-464-C10 | GORG | 0 | 0 | NA | 3 | 0 | orphan | 0 | 0 | NA | 0 | 0 | NA | 74.68 | 0 | 0.73 |
| AG-895-B08 | GORG | 0 | 0 | NA | 2 | 1 | complete | 0 | 0 | NA | 0 | 0 | NA | 74.64 | 0 | 1.09 |
| AH-326-E22 | GORG | 1 | 1 | complete | 1 | 0 | orphan | 0 | 0 | NA | 0 | 0 | NA | 74.64 | 0 | 1.96 |
| AG-901-F16 | GORG | 0 | 0 | NA | 1 | 0 | orphan | 0 | 0 | NA | 0 | 0 | NA | 74.59 | 0 | 0.86 |
| AG-414-J19 | GORG | 0 | 0 | NA | 1 | 0 | orphan | 0 | 0 | NA | 0 | 0 | NA | 74.53 | 0 | 0.96 |
| AH-321-B02 | GORG | 0 | 0 | NA | 1 | 0 | orphan | 0 | 0 | NA | 0 | 0 | NA | 74.53 | 0 | 1.33 |
| AG-894-G11 | GORG | 0 | 0 | NA | 1 | 0 | orphan | 0 | 0 | NA | 0 | 0 | NA | 74.52 | 0 | 1.12 |
| AG-912-A04 | GORG | 0 | 0 | NA | 2 | 0 | orphan | 0 | 0 | NA | 0 | 0 | NA | 74.51 | 0 | 0.89 |
| AG-447-F22 | GORG | 0 | 0 | NA | 3 | 1 | complete | 0 | 0 | NA | 0 | 0 | NA | 74.5 | 0 | 1.11 |
| AG-907-G18 | GORG | 0 | 0 | NA | 2 | 0 | orphan | 0 | 0 | NA | 0 | 0 | NA | 74.49 | 0 | 0.88 |
| AG-911-P20 | GORG | 0 | 0 | NA | 0 | 0 | NA | 0 | 0 | NA | 0 | 1 | orphan | 74.48 | 0 | 1.16 |
| AG-912-J16 | GORG | 0 | 0 | NA | 1 | 0 | orphan | 0 | 0 | NA | 0 | 0 | NA | 74.47 | 0 | 1.19 |
| AG-461-C05 | GORG | 0 | 0 | NA | 3 | 0 | orphan | 0 | 0 | NA | 0 | 0 | NA | 74.46 | 0 | 0.78 |

| | | | | | | | | | | | | | | | | |
|---|---|---|---|---|---|---|---|---|---|---|---|---|---|---|---|---|
| AG-909-I23 | GORG | 0 | 0 | NA | 1 | 0 | orphan | 0 | 0 | NA | 0 | 0 | NA | 74.42 | 0 | 0.94 |
| AG-319-J06 | GORG | 0 | 0 | NA | 2 | 0 | orphan | 0 | 0 | NA | 0 | 0 | NA | 74.4 | 0 | 1.26 |
| AH-287-A19 | GORG | 0 | 0 | NA | 1 | 0 | orphan | 0 | 0 | NA | 0 | 0 | NA | 74.37 | 0 | 0.68 |
| AG-422-A05 | GORG | 0 | 0 | NA | 3 | 0 | orphan | 0 | 0 | NA | 0 | 0 | NA | 74.31 | 0 | 0.91 |
| AG-365-F20 | GORG | 1 | 1 | complete | 1 | 0 | orphan | 0 | 0 | NA | 0 | 0 | NA | 74.3 | 0 | 1.03 |
| AG-414-A23 | GORG | 0 | 0 | NA | 2 | 0 | orphan | 0 | 0 | NA | 0 | 0 | NA | 74.3 | 0 | 0.92 |
| AH-321-O23 | GORG | 0 | 0 | NA | 2 | 0 | orphan | 0 | 0 | NA | 0 | 0 | NA | 74.3 | 0 | 0.85 |
| AG-453-D05 | GORG | 0 | 0 | NA | 1 | 0 | orphan | 0 | 0 | NA | 0 | 0 | NA | 74.25 | 0 | 0.91 |
| AG-426-N16 | GORG | 0 | 0 | NA | 1 | 0 | orphan | 0 | 0 | NA | 0 | 0 | NA | 74.23 | 0 | 1.36 |
| AG-349-P01 | GORG | 0 | 0 | NA | 1 | 0 | orphan | 0 | 0 | NA | 0 | 0 | NA | 74.17 | 0 | 1.1 |
| AG-917-M03 | GORG | 0 | 0 | NA | 1 | 0 | orphan | 0 | 0 | NA | 0 | 0 | NA | 74.17 | 0 | 0.78 |
| AG-917-I05 | GORG | 1 | 1 | complete | 1 | 0 | orphan | 0 | 0 | NA | 0 | 0 | NA | 74.16 | 0 | 1.15 |
| AG-895-O19 | GORG | 0 | 0 | NA | 1 | 0 | orphan | 0 | 0 | NA | 0 | 0 | NA | 74.14 | 0 | 1.01 |
| AG-914-O05 | GORG | 0 | 0 | NA | 2 | 0 | orphan | 0 | 0 | NA | 0 | 0 | NA | 74.14 | 0 | 1.07 |
| AG-404-O17 | GORG | 0 | 0 | NA | 1 | 0 | orphan | 0 | 0 | NA | 0 | 0 | NA | 74.1 | 0 | 0.92 |
| AG-916-P23 | GORG | 0 | 0 | NA | 1 | 0 | orphan | 0 | 0 | NA | 0 | 0 | NA | 74.1 | 0 | 1.02 |
| AG-918-O11 | GORG | 0 | 0 | NA | 1 | 0 | orphan | 0 | 0 | NA | 0 | 0 | NA | 74.1 | 0 | 1.1 |
| AG-901-F04 | GORG | 1 | 1 | complete | 2 | 0 | orphan | 0 | 0 | NA | 0 | 0 | NA | 74.05 | 0 | 0.89 |
| AG-891-F22 | GORG | 0 | 0 | NA | 1 | 0 | orphan | 0 | 0 | NA | 0 | 0 | NA | 74.03 | 0 | 1.15 |
| AG-337-D19 | GORG | 0 | 0 | NA | 2 | 0 | orphan | 0 | 0 | NA | 0 | 0 | NA | 73.99 | 0 | 0.93 |
| AG-904-M07 | GORG | 0 | 0 | NA | 1 | 0 | orphan | 0 | 0 | NA | 0 | 0 | NA | 73.99 | 0 | 1.22 |
| AG-917-K10 | GORG | 0 | 0 | NA | 1 | 0 | orphan | 0 | 0 | NA | 0 | 0 | NA | 73.99 | 0 | 1.02 |
| AG-464-J09 | GORG | 0 | 1 | NA | 0 | 0 | NA | 0 | 0 | NA | 0 | 0 | NA | 73.97 | 0 | 1.08 |
| AG-349-M18 | GORG | 0 | 0 | NA | 4 | 0 | orphan | 0 | 0 | NA | 0 | 0 | NA | 73.93 | 0 | 1.08 |
| AG-901-G09 | GORG | 1 | 1 | complete | 2 | 1 | complete | 0 | 0 | NA | 0 | 0 | NA | 73.93 | 0 | 0.99 |
| AG-913-M06 | GORG | 0 | 0 | NA | 2 | 0 | orphan | 0 | 0 | NA | 0 | 0 | NA | 73.93 | 0 | 0.92 |

| | | | | | | | | | | | | | | | |
|---|---|---|---|---|---|---|---|---|---|---|---|---|---|---|---|
| AG-422-F18 | GORG | 0 | 0 | NA | 0 | 0 | NA | 0 | 0 | NA | 0 | 1 | orphan | 73.91 | 0 | 1.24 |
| AG-914-F13 | GORG | 0 | 0 | NA | 1 | 0 | orphan | 0 | 0 | NA | 0 | 0 | NA | 73.91 | 0 | 1.12 |
| AG-439-L10 | GORG | 0 | 0 | NA | 1 | 0 | orphan | 0 | 0 | NA | 0 | 0 | NA | 73.9 | 0 | 0.87 |
| AG-349-C16 | GORG | 0 | 0 | NA | 0 | 0 | NA | 0 | 0 | NA | 0 | 3 | orphan | 73.88 | 0 | 1.12 |
| AG-914-P08 | GORG | 0 | 0 | NA | 1 | 0 | orphan | 0 | 0 | NA | 0 | 0 | NA | 73.88 | 0 | 1.74 |
| AG-453-G13 | GORG | 0 | 0 | NA | 3 | 0 | orphan | 0 | 0 | NA | 0 | 0 | NA | 73.86 | 0 | 1.24 |
| AG-908-O05 | GORG | 0 | 0 | NA | 2 | 1 | orphan | 0 | 0 | NA | 0 | 0 | NA | 73.78 | 0 | 0.83 |
| AG-325-K10 | GORG | 0 | 0 | NA | 1 | 0 | orphan | 0 | 0 | NA | 0 | 0 | NA | 73.76 | 0 | 0.82 |
| AG-426-N14 | GORG | 0 | 0 | NA | 1 | 0 | orphan | 0 | 0 | NA | 0 | 0 | NA | 73.69 | 0 | 1.25 |
| AG-349-C10 | GORG | 0 | 0 | NA | 1 | 1 | complete | 0 | 0 | NA | 0 | 0 | NA | 73.68 | 0 | 1.14 |
| AG-337-N08 | GORG | 0 | 0 | NA | 1 | 0 | orphan | 0 | 0 | NA | 0 | 0 | NA | 73.66 | 0 | 0.99 |
| AG-907-P15 | GORG | 0 | 0 | NA | 0 | 0 | NA | 0 | 0 | NA | 0 | 1 | orphan | 73.64 | 0 | 1.16 |
| AG-337-B08 | GORG | 0 | 0 | NA | 1 | 0 | orphan | 0 | 0 | NA | 0 | 1 | orphan | 73.63 | 0 | 1.1 |
| AG-359-L13 | GORG | 0 | 0 | NA | 1 | 0 | orphan | 0 | 0 | NA | 0 | 0 | NA | 73.62 | 0 | 0.94 |
| AG-894-F23 | GORG | 2 | 1 | complete | 0 | 0 | NA | 0 | 0 | NA | 0 | 0 | NA | 73.61 | 0 | 1.17 |
| AG-404-E19 | GORG | 0 | 0 | NA | 0 | 0 | NA | 0 | 0 | NA | 0 | 1 | orphan | 73.6 | 0 | 1.28 |
| AG-912-A22 | GORG | 0 | 0 | NA | 1 | 0 | orphan | 0 | 0 | NA | 0 | 0 | NA | 73.6 | 0 | 0.88 |
| AG-365-G16 | GORG | 0 | 0 | NA | 2 | 0 | orphan | 0 | 0 | NA | 0 | 0 | NA | 73.58 | 0 | 0.95 |
| AG-907-C16 | GORG | 0 | 0 | NA | 2 | 0 | orphan | 0 | 0 | NA | 0 | 0 | NA | 73.58 | 0 | 1.18 |
| AG-912-P09 | GORG | 0 | 1 | NA | 1 | 0 | orphan | 0 | 0 | NA | 0 | 0 | NA | 73.57 | 0 | 0.98 |
| AG-905-D17 | GORG | 0 | 0 | NA | 2 | 1 | complete | 0 | 0 | NA | 0 | 0 | NA | 73.55 | 0 | 0.87 |
| AG-337-A10 | GORG | 1 | 1 | complete | 0 | 0 | NA | 0 | 0 | NA | 0 | 0 | NA | 73.54 | 0 | 0.95 |
| AG-359-I20 | GORG | 0 | 0 | NA | 3 | 0 | orphan | 0 | 0 | NA | 0 | 0 | NA | 73.52 | 0 | 0.97 |
| AG-903-D08 | GORG | 0 | 0 | NA | 2 | 0 | orphan | 0 | 0 | NA | 0 | 0 | NA | 73.51 | 0 | 0.86 |

| | | | | | | | | | | | | | | | | |
|---|---|---|---|---|---|---|---|---|---|---|---|---|---|---|---|---|
| AG-892-M07 | GORG | 0 | 0 | NA | 1 | 0 | orphan | 0 | 0 | NA | 0 | 1 | orpha n | 73.49 | 0 | 1.18 |
| AG-453-E13 | GORG | 0 | 0 | NA | 1 | 0 | orphan | 0 | 0 | NA | 0 | 0 | NA | 73.49 | 0 | 0.88 |
| AG-901-M14 | GORG | 0 | 0 | NA | 2 | 0 | orphan | 0 | 0 | NA | 0 | 0 | NA | 73.49 | 0 | 1.07 |
| AG-904-B17 | GORG | 0 | 0 | NA | 1 | 0 | orphan | 0 | 0 | NA | 0 | 0 | NA | 73.49 | 0 | 1.1 |
| AG-907-A13 | GORG | 0 | 0 | NA | 1 | 0 | orphan | 0 | 0 | NA | 0 | 0 | NA | 73.49 | 0 | 0.89 |
| AG-914-B17 | GORG | 0 | 0 | NA | 2 | 0 | orphan | 0 | 0 | NA | 0 | 0 | NA | 73.49 | 0 | 0.87 |
| AG-915-D02 | GORG | 2 | 1 | complet e | 0 | 0 | NA | 0 | 0 | NA | 0 | 0 | NA | 73.43 | 0 | 0.81 |
| AG-439-E13 | GORG | 0 | 0 | NA | 1 | 0 | orphan | 0 | 0 | NA | 0 | 0 | NA | 73.41 | 0 | 1.26 |
| AG-912-K05 | GORG | 1 | 1 | complet e | 0 | 0 | NA | 0 | 0 | NA | 0 | 1 | orpha n | 73.38 | 0 | 0.91 |
| AG-439-M12 | GORG | 0 | 0 | NA | 3 | 0 | orphan | 0 | 0 | NA | 0 | 0 | NA | 73.38 | 0 | 0.91 |
| AG-893-O11 | GORG | 0 | 0 | NA | 3 | 2 | complet e | 0 | 0 | NA | 0 | 0 | NA | 73.38 | 0 | 1.2 |
| AG-895-L21 | GORG | 0 | 0 | NA | 1 | 0 | orphan | 0 | 0 | NA | 0 | 0 | NA | 73.38 | 0 | 0.9 |
| AG-904-D05 | GORG | 0 | 0 | NA | 1 | 0 | orphan | 0 | 0 | NA | 0 | 0 | NA | 73.37 | 0 | 1.12 |
| AG-426-I09 | GORG | 0 | 0 | NA | 1 | 1 | complet e | 0 | 0 | NA | 0 | 0 | NA | 73.36 | 0 | 1.16 |
| AG-447-O22 | GORG | 0 | 0 | NA | 1 | 0 | orphan | 0 | 0 | NA | 0 | 0 | NA | 73.33 | 0 | 0.67 |
| AG-897-C02 | GORG | 0 | 0 | NA | 2 | 0 | orphan | 0 | 0 | NA | 0 | 0 | NA | 73.29 | 0 | 1.04 |
| AG-895-E05 | GORG | 0 | 0 | NA | 1 | 1 | orphan | 0 | 0 | NA | 0 | 0 | NA | 73.28 | 0 | 1.18 |
| AG-913-I05 | GORG | 0 | 0 | NA | 1 | 0 | orphan | 0 | 0 | NA | 0 | 0 | NA | 73.27 | 0 | 0.91 |
| AG-337-M15 | GORG | 0 | 0 | NA | 2 | 1 | complet e | 0 | 0 | NA | 0 | 0 | NA | 73.22 | 0 | 0.74 |
| AG-349-A02 | GORG | 0 | 0 | NA | 1 | 0 | orphan | 0 | 0 | NA | 0 | 0 | NA | 73.22 | 0 | 0.88 |
| AG-349-K23 | GORG | 0 | 1 | NA | 1 | 0 | orphan | 0 | 0 | NA | 0 | 0 | NA | 73.22 | 0 | 0.91 |
| AH-287-D11 | GORG | 0 | 0 | NA | 1 | 0 | orphan | 0 | 0 | NA | 0 | 0 | NA | 73.17 | 0 | 0.9 |
| AG-915-O22 | GORG | 1 | 1 | complet e | 1 | 0 | orphan | 0 | 0 | NA | 0 | 0 | NA | 73.1 | 0 | 1.34 |
| AG-414-M10 | GORG | 0 | 0 | NA | 1 | 0 | orphan | 0 | 0 | NA | 0 | 0 | NA | 73.09 | 0 | 0.99 |
| AG-457-O17 | GORG | 0 | 0 | NA | 1 | 0 | orphan | 0 | 0 | NA | 0 | 0 | NA | 73.09 | 0 | 0.86 |

| | | | | | | | | | | | | | | | |
|---|---|---|---|---|---|---|---|---|---|---|---|---|---|---|---|
| AG-899-L21 | GORG | 0 | 0 | NA | 1 | 0 | orphan | 0 | 0 | NA | 0 | 0 | NA | 73.07 | 0 | 0.74 |
| AG-349-E23 | GORG | 0 | 0 | NA | 0 | 0 | NA | 0 | 0 | NA | 0 | 1 | orphan | 73.05 | 0 | 1.19 |
| AG-390-D03 | GORG | 0 | 0 | NA | 1 | 0 | orphan | 0 | 0 | NA | 0 | 0 | NA | 73.03 | 0 | 1 |
| AG-313-F06 | GORG | 0 | 0 | NA | 1 | 0 | orphan | 0 | 0 | NA | 0 | 0 | NA | 73.02 | 0 | 2.05 |
| AG-899-E18 | GORG | 0 | 0 | NA | 0 | 0 | NA | 0 | 0 | NA | 0 | 1 | orphan | 73.01 | 0 | 1.2 |
| AH-287-O17 | GORG | 1 | 1 | complete | 0 | 0 | NA | 0 | 0 | NA | 0 | 1 | orphan | 72.93 | 0 | 1.18 |
| AG-464-B02 | GORG | 1 | 1 | complete | 0 | 0 | NA | 0 | 0 | NA | 0 | 1 | orphan | 72.91 | 0 | 1.36 |
| AG-325-A09 | GORG | 1 | 0 | orphan | 0 | 0 | NA | 0 | 0 | NA | 0 | 0 | NA | 72.91 | 0 | 0.97 |
| AG-414-I15 | GORG | 0 | 0 | NA | 1 | 0 | orphan | 0 | 0 | NA | 0 | 0 | NA | 72.89 | 0 | 0.84 |
| AG-907-C03 | GORG | 0 | 0 | NA | 1 | 0 | orphan | 0 | 0 | NA | 0 | 0 | NA | 72.89 | 0 | 1.15 |
| AG-911-A15 | GORG | 0 | 0 | NA | 1 | 0 | orphan | 0 | 0 | NA | 0 | 0 | NA | 72.89 | 0 | 0.8 |
| AG-915-G17 | GORG | 0 | 0 | NA | 2 | 0 | orphan | 0 | 0 | NA | 0 | 0 | NA | 72.89 | 0 | 1.11 |
| AG-901-F21 | GORG | 0 | 0 | NA | 2 | 0 | orphan | 0 | 0 | NA | 0 | 0 | NA | 72.87 | 0 | 0.79 |
| AG-333-J17 | GORG | 0 | 0 | NA | 2 | 0 | orphan | 0 | 1 | NA | 0 | 0 | NA | 72.86 | 0 | 0.9 |
| AG-914-J02 | GORG | 0 | 0 | NA | 1 | 0 | orphan | 0 | 0 | NA | 0 | 0 | NA | 72.86 | 0 | 0.66 |
| AH-287-P07 | GORG | 0 | 0 | NA | 3 | 0 | orphan | 0 | 0 | NA | 0 | 1 | orphan | 72.82 | 0 | 1.66 |
| AG-426-N02 | GORG | 0 | 0 | NA | 0 | 0 | NA | 0 | 0 | NA | 0 | 1 | orphan | 72.8 | 0 | 1.14 |
| AG-918-J23 | GORG | 0 | 0 | NA | 1 | 0 | orphan | 0 | 0 | NA | 0 | 0 | NA | 72.8 | 0 | 0.8 |
| AG-910-M02 | GORG | 0 | 0 | NA | 3 | 0 | orphan | 0 | 0 | NA | 0 | 0 | NA | 72.79 | 0 | 1.13 |
| AG-913-B15 | GORG | 0 | 0 | NA | 1 | 0 | orphan | 0 | 0 | NA | 0 | 0 | NA | 72.79 | 0 | 1.13 |
| AG-892-N18 | GORG | 0 | 0 | NA | 1 | 0 | orphan | 0 | 0 | NA | 0 | 0 | NA | 72.77 | 0 | 1.62 |
| AG-337-M21 | GORG | 0 | 0 | NA | 1 | 0 | orphan | 0 | 0 | NA | 0 | 0 | NA | 72.76 | 0 | 0.83 |
| AG-390-E23 | GORG | 0 | 0 | NA | 2 | 0 | orphan | 0 | 0 | NA | 0 | 0 | NA | 72.71 | 0 | 1.13 |
| AG-319-G18 | GORG | 0 | 0 | NA | 2 | 0 | orphan | 0 | 0 | NA | 0 | 0 | NA | 72.67 | 0 | 1.19 |
| AG-912-I11 | GORG | 0 | 0 | NA | 1 | 0 | orphan | 0 | 0 | NA | 0 | 0 | NA | 72.67 | 0 | 1.05 |

| | | | | | | | | | | | | | | | | |
|---|---|---|---|---|---|---|---|---|---|---|---|---|---|---|---|---|
| AG-337-I18 | GORG | 0 | 0 | NA | 2 | 0 | orphan | 0 | 0 | NA | 0 | 0 | NA | 72.6 | 0 | 1.15 |
| AG-453-C08 | GORG | 0 | 0 | NA | 1 | 0 | orphan | 0 | 0 | NA | 0 | 0 | NA | 72.56 | 0 | 0.8 |
| AG-915-N20 | GORG | 1 | 1 | complete | 0 | 0 | NA | 0 | 0 | NA | 0 | 0 | NA | 72.55 | 0 | 1.27 |
| AG-916-L21 | GORG | 0 | 0 | NA | 1 | 0 | orphan | 0 | 0 | NA | 0 | 0 | NA | 72.55 | 0 | 0.88 |
| AG-453-K19 | GORG | 1 | 1 | complete | 1 | 0 | orphan | 0 | 0 | NA | 0 | 0 | NA | 72.53 | 0 | 0.97 |
| AG-918-E13 | GORG | 0 | 0 | NA | 1 | 1 | complete | 0 | 0 | NA | 0 | 0 | NA | 72.53 | 0 | 1.25 |
| AG-453-F05 | GORG | 0 | 0 | NA | 1 | 0 | orphan | 0 | 0 | NA | 0 | 0 | NA | 72.51 | 0 | 0.87 |
| AG-895-L02 | GORG | 0 | 0 | NA | 1 | 0 | orphan | 0 | 0 | NA | 0 | 0 | NA | 72.46 | 0 | 1.13 |
| AG-914-O08 | GORG | 1 | 1 | complete | 3 | 0 | orphan | 0 | 0 | NA | 0 | 0 | NA | 72.45 | 0 | 1.13 |
| AG-313-M11 | GORG | 0 | 0 | NA | 1 | 0 | orphan | 0 | 0 | NA | 0 | 0 | NA | 72.41 | 0 | 1.11 |
| AG-892-I03 | GORG | 0 | 0 | NA | 1 | 1 | complete | 0 | 0 | NA | 0 | 0 | NA | 72.4 | 0 | 1.07 |
| AG-910-C14 | GORG | 0 | 0 | NA | 2 | 0 | orphan | 0 | 0 | NA | 0 | 0 | NA | 72.4 | 0 | 0.97 |
| AG-435-K07 | GORG | 0 | 0 | NA | 1 | 0 | orphan | 0 | 0 | NA | 0 | 0 | NA | 72.38 | 0 | 1.12 |
| AG-461-O21 | GORG | 0 | 0 | NA | 1 | 0 | orphan | 0 | 0 | NA | 0 | 0 | NA | 72.37 | 0 | 1.24 |
| AG-916-K03 | GORG | 0 | 0 | NA | 2 | 0 | orphan | 0 | 0 | NA | 0 | 0 | NA | 72.34 | 0 | 1.07 |
| AG-365-M23 | GORG | 0 | 0 | NA | 2 | 0 | orphan | 0 | 0 | NA | 0 | 0 | NA | 72.33 | 0 | 0.85 |
| AG-414-P14 | GORG | 0 | 1 | NA | 2 | 0 | orphan | 0 | 0 | NA | 0 | 0 | NA | 72.33 | 0 | 0.88 |
| AG-337-O16 | GORG | 0 | 0 | NA | 1 | 0 | orphan | 0 | 0 | NA | 0 | 0 | NA | 72.29 | 0 | 0.87 |
| AG-359-E20 | GORG | 0 | 0 | NA | 1 | 0 | orphan | 0 | 0 | NA | 0 | 0 | NA | 72.29 | 0 | 1 |
| AG-457-E22 | GORG | 0 | 0 | NA | 1 | 0 | orphan | 0 | 0 | NA | 0 | 0 | NA | 72.29 | 0 | 1.11 |
| AG-892-G03 | GORG | 0 | 0 | NA | 1 | 0 | orphan | 0 | 0 | NA | 0 | 0 | NA | 72.29 | 0 | 1.16 |
| AG-911-J02 | GORG | 0 | 0 | NA | 2 | 0 | orphan | 0 | 0 | NA | 0 | 0 | NA | 72.29 | 0 | 1.38 |
| AG-913-A05 | GORG | 0 | 0 | NA | 2 | 0 | orphan | 0 | 0 | NA | 0 | 0 | NA | 72.29 | 0 | 0.86 |
| AG-915-B19 | GORG | 0 | 0 | NA | 1 | 0 | orphan | 0 | 0 | NA | 0 | 0 | NA | 72.29 | 0 | 0.82 |
| AG-430-P11 | GORG | 0 | 0 | NA | 1 | 0 | orphan | 0 | 0 | NA | 0 | 0 | NA | 72.2 | 0 | 0.94 |

| | | | | | | | | | | | | | | | | |
|---|---|---|---|---|---|---|---|---|---|---|---|---|---|---|---|---|
| AG-910-K08 | GORG | 0 | 0 | NA | 2 | 0 | orphan | 0 | 0 | NA | 0 | 0 | NA | 72.18 | 0 | 1 |
| AG-905-K02 | GORG | 0 | 0 | NA | 1 | 0 | orphan | 0 | 0 | NA | 0 | 0 | NA | 72.11 | 0 | 0.77 |
| AG-907-P23 | GORG | 0 | 0 | NA | 1 | 0 | orphan | 0 | 0 | NA | 0 | 0 | NA | 72.1 | 0 | 0.86 |
| AH-326-B17 | GORG | 0 | 0 | NA | 1 | 0 | orphan | 0 | 0 | NA | 0 | 0 | NA | 72.09 | 0 | 0.83 |
| AG-430-N20 | GORG | 0 | 0 | NA | 2 | 0 | orphan | 0 | 2 | NA | 0 | 0 | NA | 72.06 | 0 | 1.03 |
| AG-892-D16 | GORG | 0 | 0 | NA | 1 | 0 | orphan | 0 | 0 | NA | 0 | 0 | NA | 72.06 | 0 | 1.56 |
| AG-439-H11 | GORG | 0 | 0 | NA | 1 | 0 | orphan | 0 | 0 | NA | 0 | 0 | NA | 72.04 | 0 | 0.92 |
| AG-325-O19 | GORG | 0 | 0 | NA | 2 | 0 | orphan | 0 | 0 | NA | 0 | 0 | NA | 72.03 | 0 | 0.82 |
| AG-920-B06 | GORG | 0 | 0 | NA | 1 | 0 | orphan | 0 | 0 | NA | 0 | 0 | NA | 72.01 | 0 | 0.91 |
| AG-911-N03 | GORG | 1 | 1 | complete | 0 | 0 | NA | 0 | 0 | NA | 0 | 0 | NA | 71.98 | 0 | 1.03 |
| AG-913-K22 | GORG | 0 | 1 | NA | 0 | 0 | NA | 0 | 0 | NA | 0 | 0 | NA | 71.98 | 0 | 1.12 |
| AH-321-A16 | GORG | 0 | 0 | NA | 2 | 0 | orphan | 0 | 0 | NA | 0 | 0 | NA | 71.96 | 0 | 0.8 |
| AG-903-N11 | GORG | 0 | 0 | NA | 0 | 0 | NA | 0 | 0 | NA | 0 | 1 | orphan | 71.95 | 0 | 1.16 |
| AG-891-D16 | GORG | 0 | 0 | NA | 2 | 0 | orphan | 0 | 0 | NA | 0 | 0 | NA | 71.89 | 0 | 1.13 |
| AG-903-M03 | GORG | 0 | 0 | NA | 1 | 0 | orphan | 0 | 1 | NA | 0 | 0 | NA | 71.89 | 0 | 0.82 |
| AG-917-G23 | GORG | 0 | 0 | NA | 2 | 0 | orphan | 0 | 0 | NA | 0 | 0 | NA | 71.89 | 0 | 0.96 |
| AG-907-K11 | GORG | 0 | 0 | NA | 1 | 0 | orphan | 0 | 0 | NA | 0 | 0 | NA | 71.87 | 0 | 1.06 |
| AH-326-O04 | GORG | 0 | 0 | NA | 1 | 0 | orphan | 0 | 0 | NA | 0 | 0 | NA | 71.87 | 0 | 1.05 |
| AG-894-N09 | GORG | 1 | 1 | complete | 1 | 0 | orphan | 0 | 0 | NA | 0 | 0 | NA | 71.85 | 0 | 1.02 |
| AG-916-I03 | GORG | 0 | 0 | NA | 0 | 0 | NA | 0 | 0 | NA | 0 | 1 | orphan | 71.8 | 0 | 1.04 |
| AG-365-N08 | GORG | 0 | 0 | NA | 1 | 1 | orphan | 0 | 0 | NA | 0 | 0 | NA | 71.74 | 0 | 0.79 |
| AG-917-O21 | GORG | 0 | 0 | NA | 1 | 0 | orphan | 0 | 0 | NA | 0 | 0 | NA | 71.71 | 0 | 0.76 |
| AG-893-K16 | GORG | 0 | 0 | NA | 2 | 0 | orphan | 0 | 0 | NA | 0 | 1 | orphan | 71.7 | 0 | 1.19 |
| AG-359-N03 | GORG | 0 | 0 | NA | 1 | 0 | orphan | 0 | 0 | NA | 0 | 0 | NA | 71.7 | 0 | 1.14 |
| AG-365-E11 | GORG | 0 | 0 | NA | 3 | 0 | orphan | 0 | 0 | NA | 0 | 0 | NA | 71.7 | 0 | 0.98 |

| | | | | | | | | | | | | | | | | |
|---|---|---|---|---|---|---|---|---|---|---|---|---|---|---|---|---|
| AG-447-G14 | GORG | 0 | 0 | NA | 1 | 0 | orphan | 0 | 0 | NA | 0 | 0 | NA | 71.7 | 0 | 0.85 |
| AG-895-I14 | GORG | 0 | 0 | NA | 3 | 0 | orphan | 0 | 0 | NA | 0 | 0 | NA | 71.7 | 0 | 0.95 |
| AG-907-N15 | GORG | 0 | 0 | NA | 1 | 0 | orphan | 0 | 0 | NA | 0 | 0 | NA | 71.7 | 0 | 0.98 |
| AG-325-E02 | GORG | 0 | 0 | NA | 3 | 0 | orphan | 0 | 0 | NA | 0 | 0 | NA | 71.69 | 0 | 0.77 |
| AG-897-D19 | GORG | 0 | 0 | NA | 2 | 0 | orphan | 0 | 0 | NA | 0 | 0 | NA | 71.69 | 0 | 1.1 |
| AG-905-N15 | GORG | 0 | 0 | NA | 0 | 0 | NA | 1 | 0 | orphan | 0 | 0 | NA | 71.69 | 0 | 1.07 |
| AG-911-L18 | GORG | 0 | 0 | NA | 1 | 0 | orphan | 0 | 0 | NA | 0 | 0 | NA | 71.69 | 0 | 1.11 |
| AH-326-B11 | GORG | 0 | 0 | NA | 3 | 0 | orphan | 0 | 0 | NA | 0 | 0 | NA | 71.69 | 0 | 0.93 |
| AG-390-B03 | GORG | 0 | 0 | NA | 1 | 0 | orphan | 0 | 0 | NA | 0 | 0 | NA | 71.66 | 0 | 1.04 |
| AG-461-E16 | GORG | 0 | 0 | NA | 0 | 0 | NA | 0 | 0 | NA | 0 | 1 | orphan | 71.65 | 0 | 1 |
| AG-343-I10 | GORG | 0 | 0 | NA | 1 | 0 | orphan | 0 | 0 | NA | 0 | 0 | NA | 71.6 | 0 | 1.17 |
| AH-321-P05 | GORG | 0 | 0 | NA | 1 | 0 | orphan | 0 | 0 | NA | 0 | 0 | NA | 71.6 | 0 | 1.23 |
| AG-908-J14 | GORG | 0 | 0 | NA | 1 | 0 | orphan | 0 | 0 | NA | 0 | 0 | NA | 71.57 | 0 | 1.22 |
| AG-915-F20 | GORG | 0 | 0 | NA | 1 | 0 | orphan | 0 | 0 | NA | 0 | 1 | orphan | 71.56 | 0 | 0.76 |
| AG-410-A05 | GORG | 0 | 0 | NA | 2 | 0 | orphan | 0 | 0 | NA | 0 | 0 | NA | 71.47 | 0 | 1.44 |
| AG-893-B02 | GORG | 0 | 0 | NA | 1 | 0 | orphan | 0 | 0 | NA | 0 | 0 | NA | 71.47 | 0 | 1.1 |
| AG-891-D22 | GORG | 0 | 0 | NA | 1 | 0 | orphan | 0 | 0 | NA | 0 | 0 | NA | 71.43 | 0 | 0.91 |
| AH-321-A07 | GORG | 0 | 0 | NA | 1 | 0 | orphan | 0 | 0 | NA | 0 | 0 | NA | 71.32 | 0 | 1.58 |
| AG-915-K22 | GORG | 0 | 0 | NA | 1 | 0 | orphan | 0 | 0 | NA | 0 | 0 | NA | 71.3 | 0 | 0.92 |
| AG-470-I02 | GORG | 0 | 0 | NA | 0 | 0 | NA | 0 | 0 | NA | 0 | 1 | orphan | 71.26 | 0 | 1.02 |
| AH-326-M17 | GORG | 0 | 0 | NA | 1 | 0 | orphan | 0 | 0 | NA | 0 | 0 | NA | 71.25 | 0 | 0.92 |
| AG-905-F19 | GORG | 0 | 0 | NA | 1 | 0 | orphan | 0 | 0 | NA | 0 | 0 | NA | 71.22 | 0 | 0.92 |
| AG-901-N02 | GORG | 0 | 0 | NA | 1 | 0 | orphan | 0 | 0 | NA | 0 | 0 | NA | 71.18 | 0 | 0.92 |
| AG-390-E16 | GORG | 0 | 0 | NA | 1 | 0 | orphan | 0 | 0 | NA | 0 | 0 | NA | 71.15 | 0 | 0.78 |
| AG-447-I11 | GORG | 0 | 0 | NA | 1 | 0 | orphan | 0 | 0 | NA | 0 | 0 | NA | 71.14 | 0 | 0.97 |
| AG-325-L22 | GORG | 0 | 0 | NA | 1 | 0 | orphan | 0 | 0 | NA | 0 | 0 | NA | 71.08 | 0 | 0.84 |

| | | | | | | | | | | | | | | | | |
|---|---|---|---|---|---|---|---|---|---|---|---|---|---|---|---|---|
| AG-426-K21 | GORG | 1 | 1 | complet e | 1 | 0 | orphan | 0 | 0 | NA | 0 | 0 | NA | 71.08 | 0 | 0.95 |
| AG-457-G04 | GORG | 0 | 0 | NA | 1 | 0 | orphan | 0 | 0 | NA | 0 | 0 | NA | 71.08 | 0 | 0.91 |
| AG-898-A06 | GORG | 0 | 0 | NA | 1 | 0 | orphan | 0 | 0 | NA | 0 | 0 | NA | 71.08 | 0 | 1.26 |
| AG-903-K09 | GORG | 0 | 0 | NA | 1 | 0 | orphan | 0 | 0 | NA | 0 | 0 | NA | 71.08 | 0 | 0.82 |
| AG-911-E03 | GORG | 0 | 0 | NA | 1 | 0 | orphan | 0 | 0 | NA | 0 | 0 | NA | 71.08 | 0 | 1.14 |
| AG-916-G11 | GORG | 0 | 0 | NA | 1 | 0 | orphan | 0 | 0 | NA | 0 | 0 | NA | 71.08 | 0 | 0.87 |
| AG-349-I11 | GORG | 0 | 0 | NA | 0 | 0 | NA | 0 | 0 | NA | 0 | 1 | orpha n | 71.06 | 0 | 1.16 |
| AG-349-K10 | GORG | 0 | 0 | NA | 0 | 0 | NA | 0 | 0 | NA | 0 | 1 | orpha n | 71.06 | 0 | 1.12 |
| AG-343-M19 | GORG | 0 | 0 | NA | 2 | 0 | orphan | 0 | 0 | NA | 0 | 0 | NA | 71.06 | 0 | 0.9 |
| AG-404-D18 | GORG | 0 | 0 | NA | 3 | 0 | orphan | 0 | 0 | NA | 0 | 0 | NA | 71.06 | 0 | 1.29 |
| AG-435-J18 | GORG | 0 | 0 | NA | 1 | 0 | orphan | 0 | 0 | NA | 0 | 0 | NA | 71.04 | 0 | 0.96 |
| AG-435-B08 | GORG | 0 | 0 | NA | 1 | 0 | orphan | 0 | 0 | NA | 0 | 0 | NA | 71.03 | 0 | 0.99 |
| AG-410-A18 | GORG | 0 | 0 | NA | 1 | 0 | orphan | 0 | 0 | NA | 0 | 0 | NA | 71.02 | 0 | 1.03 |
| AG-915-K15 | GORG | 0 | 0 | NA | 1 | 0 | orphan | 0 | 0 | NA | 0 | 0 | NA | 71.01 | 0 | 0.79 |
| AG-899-I06 | GORG | 1 | 1 | complet e | 0 | 0 | NA | 0 | 0 | NA | 0 | 0 | NA | 70.98 | 0 | 1 |
| AG-470-I07 | GORG | 0 | 0 | NA | 1 | 0 | orphan | 0 | 0 | NA | 0 | 0 | NA | 70.96 | 0 | 0.99 |
| AG-919-M14 | GORG | 0 | 1 | NA | 0 | 0 | NA | 0 | 0 | NA | 0 | 0 | NA | 70.95 | 0 | 1.32 |
| AG-911-E15 | GORG | 0 | 0 | NA | 1 | 0 | orphan | 0 | 0 | NA | 0 | 0 | NA | 70.93 | 0 | 1.08 |
| AG-911-A02 | GORG | 0 | 0 | NA | 2 | 0 | orphan | 0 | 1 | NA | 0 | 0 | NA | 70.92 | 0 | 0.96 |
| AG-916-A10 | GORG | 0 | 0 | NA | 1 | 0 | orphan | 0 | 0 | NA | 0 | 0 | NA | 70.91 | 0 | 0.83 |
| AG-916-N21 | GORG | 0 | 0 | NA | 1 | 0 | orphan | 0 | 0 | NA | 0 | 0 | NA | 70.9 | 0 | 0.71 |
| AG-915-A01 | GORG | 1 | 1 | complet e | 0 | 0 | NA | 0 | 0 | NA | 0 | 0 | NA | 70.89 | 0 | 1.74 |
| AG-898-P07 | GORG | 0 | 0 | NA | 2 | 0 | orphan | 0 | 0 | NA | 0 | 0 | NA | 70.88 | 0 | 1.22 |
| AG-907-B07 | GORG | 1 | 0 | orphan | 1 | 0 | orphan | 0 | 0 | NA | 0 | 0 | NA | 70.88 | 0 | 1.03 |
| AG-914-G18 | GORG | 0 | 0 | NA | 1 | 0 | orphan | 0 | 0 | NA | 0 | 0 | NA | 70.87 | 0 | 1.21 |

| | | | | | | | | | | | | | | | | |
|---|---|---|---|---|---|---|---|---|---|---|---|---|---|---|---|---|
| AG-917-I15 | GORG | 0 | 0 | NA | 1 | 0 | orphan | 0 | 0 | NA | 0 | 0 | NA | 70.85 | 0 | 0.87 |
| AG-457-M19 | GORG | 0 | 0 | NA | 1 | 0 | orphan | 0 | 0 | NA | 0 | 0 | NA | 70.83 | 0 | 0.97 |
| AG-390-J22 | GORG | 0 | 0 | NA | 1 | 0 | orphan | 0 | 0 | NA | 0 | 0 | NA | 70.81 | 0 | 1.41 |
| AG-907-F04 | GORG | 1 | 1 | complete | 0 | 0 | NA | 0 | 0 | NA | 0 | 0 | NA | 70.79 | 0 | 1.06 |
| AG-915-M19 | GORG | 0 | 0 | NA | 1 | 0 | orphan | 0 | 0 | NA | 0 | 0 | NA | 70.76 | 0 | 0.92 |
| AG-898-G03 | GORG | 0 | 0 | NA | 1 | 0 | orphan | 0 | 0 | NA | 0 | 1 | orphan | 70.74 | 0 | 1.31 |
| AH-287-L15 | GORG | 0 | 1 | NA | 0 | 0 | NA | 0 | 0 | NA | 0 | 0 | NA | 70.74 | 0 | 1.11 |
| AG-430-E19 | GORG | 0 | 0 | NA | 3 | 0 | orphan | 0 | 0 | NA | 0 | 0 | NA | 70.7 | 0 | 0.81 |
| AG-908-E14 | GORG | 0 | 0 | NA | 1 | 0 | orphan | 0 | 0 | NA | 0 | 0 | NA | 70.69 | 0 | 1.38 |
| AG-439-A02 | GORG | 0 | 0 | NA | 2 | 0 | orphan | 0 | 0 | NA | 0 | 0 | NA | 70.68 | 0 | 0.9 |
| AG-908-F08 | GORG | 0 | 0 | NA | 3 | 0 | orphan | 0 | 0 | NA | 0 | 0 | NA | 70.68 | 0 | 0.86 |
| AG-422-I20 | GORG | 0 | 0 | NA | 1 | 0 | orphan | 0 | 0 | NA | 0 | 0 | NA | 70.67 | 0 | 1.16 |
| AH-326-F10 | GORG | 0 | 0 | NA | 1 | 0 | orphan | 1 | 1 | complete | 0 | 0 | NA | 70.66 | 0 | 0.82 |
| AG-892-K10 | GORG | 1 | 1 | complete | 0 | 0 | NA | 0 | 0 | NA | 0 | 0 | NA | 70.65 | 0 | 1.2 |
| AG-904-F02 | GORG | 0 | 0 | NA | 2 | 0 | orphan | 0 | 0 | NA | 0 | 0 | NA | 70.62 | 0 | 0.95 |
| AG-439-K12 | GORG | 1 | 1 | complete | 2 | 0 | orphan | 0 | 0 | NA | 0 | 0 | NA | 70.59 | 0 | 1.41 |
| AG-899-G21 | GORG | 0 | 0 | NA | 1 | 0 | orphan | 0 | 0 | NA | 0 | 0 | NA | 70.59 | 0 | 0.85 |
| AG-916-A02 | GORG | 0 | 0 | NA | 1 | 0 | orphan | 0 | 0 | NA | 0 | 0 | NA | 70.59 | 0 | 0.81 |
| AG-404-P15 | GORG | 1 | 0 | orphan | 0 | 0 | NA | 0 | 0 | NA | 0 | 0 | NA | 70.58 | 0 | 1.37 |
| AG-919-K05 | GORG | 0 | 0 | NA | 1 | 0 | orphan | 0 | 0 | NA | 0 | 0 | NA | 70.58 | 0 | 0.84 |
| AG-470-C13 | GORG | 1 | 1 | complete | 1 | 0 | orphan | 0 | 0 | NA | 0 | 0 | NA | 70.57 | 0 | 0.94 |
| AG-897-O19 | GORG | 0 | 0 | NA | 1 | 0 | orphan | 0 | 0 | NA | 0 | 1 | orphan | 70.56 | 0 | 1.06 |
| AG-319-A09 | GORG | 0 | 0 | NA | 1 | 1 | complete | 0 | 0 | NA | 0 | 0 | NA | 70.55 | 0 | 1.41 |
| AH-326-P06 | GORG | 0 | 0 | NA | 1 | 0 | orphan | 0 | 0 | NA | 0 | 0 | NA | 70.54 | 0 | 0.89 |

| | | | | | | | | | | | | | | | | |
|---|---|---|---|---|---|---|---|---|---|---|---|---|---|---|---|---|
| AG-343-C23 | GORG | 0 | 0 | NA | 1 | 0 | orphan | 0 | 0 | NA | 0 | 0 | NA | 70.48 | 0 | 0.9 |
| AG-893-P11 | GORG | 0 | 0 | NA | 1 | 0 | orphan | 0 | 0 | NA | 0 | 0 | NA | 70.48 | 0 | 0.89 |
| AG-894-D19 | GORG | 0 | 0 | NA | 1 | 0 | orphan | 0 | 0 | NA | 0 | 0 | NA | 70.48 | 0 | 1.27 |
| AG-903-E09 | GORG | 0 | 0 | NA | 1 | 0 | orphan | 0 | 0 | NA | 0 | 0 | NA | 70.48 | 0 | 0.68 |
| AG-909-E20 | GORG | 1 | 0 | orphan | 0 | 0 | NA | 0 | 0 | NA | 0 | 0 | NA | 70.48 | 0 | 1.09 |
| AG-910-K17 | GORG | 0 | 0 | NA | 1 | 0 | orphan | 0 | 0 | NA | 0 | 0 | NA | 70.48 | 0 | 0.93 |
| AG-914-P16 | GORG | 0 | 0 | NA | 2 | 0 | orphan | 0 | 0 | NA | 0 | 0 | NA | 70.48 | 0 | 0.85 |
| AH-321-J11 | GORG | 0 | 0 | NA | 1 | 0 | orphan | 0 | 0 | NA | 0 | 0 | NA | 70.48 | 0 | 1.03 |
| AG-390-L11 | GORG | 0 | 0 | NA | 2 | 0 | orphan | 0 | 0 | NA | 0 | 0 | NA | 70.47 | 0 | 1.29 |
| AG-892-N21 | GORG | 1 | 1 | complete | 1 | 0 | orphan | 0 | 0 | NA | 0 | 0 | NA | 70.46 | 0 | 1.78 |
| AG-905-O19 | GORG | 0 | 0 | NA | 1 | 0 | orphan | 0 | 0 | NA | 0 | 0 | NA | 70.45 | 0 | 0.92 |
| AG-470-J14 | GORG | 0 | 0 | NA | 1 | 0 | orphan | 0 | 0 | NA | 0 | 0 | NA | 70.44 | 0 | 0.83 |
| AG-911-O15 | GORG | 0 | 0 | NA | 1 | 0 | orphan | 0 | 0 | NA | 0 | 0 | NA | 70.42 | 0 | 0.91 |
| AG-893-P07 | GORG | 0 | 0 | NA | 2 | 0 | orphan | 0 | 0 | NA | 0 | 0 | NA | 70.4 | 0 | 0.93 |
| AG-893-L15 | GORG | 0 | 0 | NA | 1 | 0 | orphan | 0 | 0 | NA | 0 | 0 | NA | 70.39 | 0 | 1.06 |
| AG-909-N10 | GORG | 0 | 0 | NA | 1 | 0 | orphan | 0 | 0 | NA | 0 | 0 | NA | 70.38 | 0 | 0.81 |
| AG-464-N14 | GORG | 0 | 0 | NA | 3 | 0 | orphan | 0 | 0 | NA | 0 | 0 | NA | 70.36 | 0 | 0.8 |
| AG-390-C08 | GORG | 1 | 1 | complete | 0 | 0 | NA | 0 | 0 | NA | 0 | 0 | NA | 70.34 | 0 | 1.28 |
| AG-453-P15 | GORG | 0 | 0 | NA | 2 | 0 | orphan | 0 | 0 | NA | 0 | 0 | NA | 70.34 | 0 | 0.84 |
| AG-426-J23 | GORG | 0 | 0 | NA | 1 | 0 | orphan | 0 | 1 | NA | 0 | 0 | NA | 70.33 | 0 | 1.3 |
| AG-901-C17 | GORG | 0 | 0 | NA | 2 | 0 | orphan | 0 | 0 | NA | 0 | 0 | NA | 70.33 | 0 | 0.76 |
| AG-349-M05 | GORG | 0 | 0 | NA | 1 | 0 | orphan | 0 | 0 | NA | 0 | 0 | NA | 70.3 | 0 | 0.91 |
| AG-900-N04 | GORG | 0 | 0 | NA | 1 | 0 | orphan | 0 | 0 | NA | 0 | 0 | NA | 70.3 | 0 | 1.04 |
| AG-426-B19 | GORG | 0 | 0 | NA | 1 | 0 | orphan | 0 | 0 | NA | 0 | 1 | orphan | 70.29 | 0 | 0.96 |
| AG-914-I15 | GORG | 0 | 0 | NA | 1 | 0 | orphan | 0 | 0 | NA | 0 | 0 | NA | 70.23 | 0 | 1.17 |
| AG-333-E10 | GORG | 0 | 0 | NA | 3 | 0 | orphan | 0 | 0 | NA | 0 | 0 | NA | 70.18 | 0 | 1 |

| | | | | | | | | | | | | | | | | |
|---|---|---|---|---|---|---|---|---|---|---|---|---|---|---|---|---|
| AG-447-K23 | GORG | 0 | 0 | NA | 2 | 1 | complet e | 0 | 0 | NA | 0 | 0 | NA | 70.17 | 0 | 1.27 |
| AG-909-D07 | GORG | 1 | 1 | complet e | 0 | 0 | NA | 0 | 0 | NA | 0 | 0 | NA | 70.15 | 0 | 1.48 |
| AG-895-G06 | GORG | 0 | 0 | NA | 1 | 0 | orphan | 0 | 0 | NA | 0 | 0 | NA | 70.14 | 0 | 0.86 |
| AG-414-B18 | GORG | 0 | 0 | NA | 1 | 0 | orphan | 0 | 0 | NA | 0 | 0 | NA | 70.13 | 0 | 1.24 |
| AG-892-G09 | GORG | 0 | 0 | NA | 1 | 0 | orphan | 0 | 0 | NA | 0 | 1 | orpha n | 70.12 | 0 | 1.17 |
| AG-914-K21 | GORG | 0 | 0 | NA | 2 | 0 | orphan | 0 | 0 | NA | 0 | 0 | NA | 70.09 | 0 | 0.97 |
| AH-324-P18 | GORG | 0 | 0 | NA | 2 | 0 | orphan | 0 | 0 | NA | 0 | 1 | orpha n | 70.06 | 0 | 2.56 |
| AG-898-E04 | GORG | 0 | 0 | NA | 1 | 0 | orphan | 0 | 0 | NA | 0 | 0 | NA | 70.04 | 0 | 0.73 |
| AG-333-I15 | GORG | 0 | 0 | NA | 1 | 0 | orphan | 0 | 0 | NA | 0 | 0 | NA | 70 | 0 | 1.09 |
| AG-910-B16 | GORG | 0 | 0 | NA | 2 | 1 | complet e | 0 | 0 | NA | 0 | 0 | NA | 70 | 0 | 0.87 |
| AG-422-F10 | GORG | 0 | 0 | NA | 2 | 0 | orphan | 0 | 0 | NA | 0 | 0 | NA | 69.99 | 0 | 0.91 |
| AG-911-D03 | GORG | 0 | 0 | NA | 1 | 0 | orphan | 0 | 0 | NA | 0 | 0 | NA | 69.99 | 0 | 0.93 |
| AG-914-A08 | GORG | 0 | 0 | NA | 2 | 1 | complet e | 0 | 0 | NA | 0 | 0 | NA | 69.99 | 0 | 1.13 |
| AG-894-K22 | GORG | 0 | 0 | NA | 1 | 0 | orphan | 0 | 0 | NA | 0 | 1 | orpha n | 69.98 | 0 | 0.94 |
| AG-908-P13 | GORG | 0 | 0 | NA | 1 | 0 | orphan | 0 | 0 | NA | 0 | 0 | NA | 69.98 | 0 | 0.79 |
| AG-911-A04 | GORG | 2 | 1 | orphan | 0 | 0 | NA | 0 | 0 | NA | 0 | 0 | NA | 69.97 | 0 | 1.26 |
| AG-917-O03 | GORG | 0 | 0 | NA | 2 | 0 | orphan | 0 | 0 | NA | 0 | 0 | NA | 69.96 | 0 | 0.87 |
| AG-915-D19 | GORG | 0 | 0 | NA | 2 | 1 | complet e | 0 | 0 | NA | 0 | 0 | NA | 69.95 | 0 | 1.04 |
| AG-422-B14 | GORG | 0 | 0 | NA | 1 | 0 | orphan | 0 | 0 | NA | 0 | 1 | orpha n | 69.93 | 0 | 1.04 |
| AG-899-N03 | GORG | 0 | 0 | NA | 1 | 0 | orphan | 0 | 0 | NA | 0 | 0 | NA | 69.93 | 0 | 0.98 |
| AG-343-F07 | GORG | 0 | 0 | NA | 2 | 0 | orphan | 0 | 0 | NA | 0 | 0 | NA | 69.88 | 0 | 0.72 |
| AG-343-O17 | GORG | 0 | 0 | NA | 1 | 1 | complet e | 0 | 0 | NA | 0 | 0 | NA | 69.88 | 0 | 0.87 |
| AG-349-L11 | GORG | 0 | 0 | NA | 2 | 0 | orphan | 0 | 0 | NA | 0 | 0 | NA | 69.88 | 0 | 0.89 |
| AG-359-K16 | GORG | 0 | 0 | NA | 2 | 0 | orphan | 0 | 2 | NA | 0 | 0 | NA | 69.88 | 0 | 1.03 |

| | | | | | | | | | | | | | | | |
|---|---|---|---|---|---|---|---|---|---|---|---|---|---|---|---|
| AG-422-K23 | GORG | 0 | 0 | NA | 3 | 0 | orphan | 0 | 0 | NA | 0 | 0 | NA | 69.88 | 0 | 0.92 |
| AG-447-E23 | GORG | 0 | 0 | NA | 1 | 0 | orphan | 0 | 0 | NA | 0 | 0 | NA | 69.88 | 0 | 1.04 |
| AG-457-M16 | GORG | 0 | 0 | NA | 1 | 0 | orphan | 0 | 0 | NA | 0 | 0 | NA | 69.88 | 0 | 0.84 |
| AG-892-B03 | GORG | 0 | 0 | NA | 2 | 0 | orphan | 0 | 0 | NA | 0 | 0 | NA | 69.88 | 0 | 1.05 |
| AG-911-G10 | GORG | 0 | 0 | NA | 1 | 0 | orphan | 0 | 0 | NA | 0 | 0 | NA | 69.88 | 0 | 1.11 |
| AG-913-J21 | GORG | 0 | 0 | NA | 1 | 0 | orphan | 0 | 0 | NA | 0 | 0 | NA | 69.88 | 0 | 0.79 |
| AG-914-F14 | GORG | 0 | 0 | NA | 2 | 0 | orphan | 0 | 0 | NA | 0 | 0 | NA | 69.88 | 0 | 1.39 |
| AH-321-J23 | GORG | 0 | 0 | NA | 1 | 0 | orphan | 0 | 0 | NA | 0 | 0 | NA | 69.88 | 0 | 1.01 |
| AH-326-I05 | GORG | 0 | 0 | NA | 2 | 0 | orphan | 0 | 0 | NA | 0 | 0 | NA | 69.88 | 0 | 0.84 |
| AG-325-M19 | GORG | 0 | 0 | NA | 1 | 0 | orphan | 0 | 0 | NA | 0 | 0 | NA | 69.85 | 0 | 0.99 |
| AG-390-O23 | GORG | 0 | 0 | NA | 1 | 0 | orphan | 0 | 0 | NA | 0 | 0 | NA | 69.85 | 0 | 1.08 |
| AG-908-D17 | GORG | 0 | 0 | NA | 2 | 0 | orphan | 0 | 0 | NA | 0 | 0 | NA | 69.83 | 0 | 0.89 |
| AG-909-M13 | GORG | 0 | 0 | NA | 1 | 0 | orphan | 0 | 1 | NA | 0 | 0 | NA | 69.83 | 0 | 0.95 |
| AG-360-K19 | GORG | 0 | 0 | NA | 1 | 0 | orphan | 0 | 0 | NA | 0 | 0 | NA | 69.81 | 0 | 0.82 |
| AG-390-G10 | GORG | 0 | 0 | NA | 2 | 0 | orphan | 0 | 0 | NA | 0 | 0 | NA | 69.81 | 0 | 1.12 |
| AG-414-E07 | GORG | 0 | 0 | NA | 2 | 0 | orphan | 0 | 0 | NA | 0 | 0 | NA | 69.81 | 0 | 0.82 |
| AG-426-J11 | GORG | 0 | 0 | NA | 1 | 0 | orphan | 0 | 0 | NA | 0 | 0 | NA | 69.81 | 0 | 1.21 |
| AG-430-C08 | GORG | 1 | 1 | complet e | 1 | 0 | orphan | 0 | 0 | NA | 0 | 0 | NA | 69.81 | 0 | 0.97 |
| AG-447-L03 | GORG | 0 | 0 | NA | 1 | 0 | orphan | 0 | 0 | NA | 0 | 0 | NA | 69.81 | 0 | 1.25 |
| AG-470-K18 | GORG | 0 | 0 | NA | 1 | 0 | orphan | 0 | 0 | NA | 0 | 0 | NA | 69.81 | 0 | 0.81 |
| AG-891-N16 | GORG | 0 | 0 | NA | 1 | 0 | orphan | 0 | 0 | NA | 0 | 0 | NA | 69.81 | 0 | 0.84 |
| AG-894-C21 | GORG | 0 | 0 | NA | 3 | 0 | orphan | 0 | 0 | NA | 0 | 0 | NA | 69.81 | 0 | 1.01 |
| AG-899-B05 | GORG | 0 | 0 | NA | 1 | 0 | orphan | 0 | 0 | NA | 0 | 0 | NA | 69.81 | 0 | 1.17 |
| AG-907-J15 | GORG | 0 | 0 | NA | 1 | 0 | orphan | 0 | 0 | NA | 0 | 0 | NA | 69.81 | 0 | 0.97 |
| AG-913-O23 | GORG | 0 | 0 | NA | 2 | 0 | orphan | 0 | 0 | NA | 0 | 0 | NA | 69.81 | 0 | 0.93 |
| AG-916-F19 | GORG | 0 | 0 | NA | 1 | 0 | orphan | 0 | 0 | NA | 0 | 0 | NA | 69.81 | 0 | 0.82 |
| AG-917-I08 | GORG | 1 | 1 | orphan | 1 | 0 | orphan | 0 | 0 | NA | 0 | 0 | NA | 69.81 | 0 | 0.91 |

| | | | | | | | | | | | | | | | | |
|---|---|---|---|---|---|---|---|---|---|---|---|---|---|---|---|---|
| AG-365-J21 | GORG | 0 | 0 | NA | 1 | 0 | orphan | 0 | 0 | NA | 0 | 0 | NA | 69.76 | 0 | 0.87 |
| AG-439-E16 | GORG | 0 | 0 | NA | 2 | 0 | orphan | 0 | 0 | NA | 0 | 0 | NA | 69.74 | 0 | 0.83 |
| AG-426-K03 | GORG | 0 | 0 | NA | 1 | 0 | orphan | 0 | 0 | NA | 0 | 0 | NA | 69.73 | 0 | 0.96 |
| AG-390-F17 | GORG | 0 | 0 | NA | 1 | 0 | orphan | 0 | 0 | NA | 0 | 0 | NA | 69.72 | 0 | 1.05 |
| AG-915-N07 | GORG | 0 | 0 | NA | 2 | 0 | orphan | 0 | 0 | NA | 0 | 0 | NA | 69.72 | 0 | 1.11 |
| AG-893-G11 | GORG | 0 | 0 | NA | 1 | 0 | orphan | 0 | 0 | NA | 0 | 0 | NA | 69.7 | 0 | 1.18 |
| AG-907-I07 | GORG | 0 | 0 | NA | 1 | 0 | orphan | 0 | 0 | NA | 0 | 0 | NA | 69.59 | 0 | 1.01 |
| AH-324-G23 | GORG | 1 | 1 | complete | 0 | 0 | NA | 0 | 0 | NA | 0 | 2 | orphan | 69.57 | 0 | 1.04 |
| AH-287-E03 | GORG | 0 | 0 | NA | 1 | 0 | orphan | 0 | 0 | NA | 0 | 0 | NA | 69.55 | 0 | 0.64 |
| AG-899-C16 | GORG | 0 | 0 | NA | 1 | 0 | orphan | 0 | 0 | NA | 0 | 0 | NA | 69.54 | 0 | 0.88 |
| AG-414-M06 | GORG | 0 | 0 | NA | 1 | 0 | orphan | 0 | 0 | NA | 0 | 0 | NA | 69.53 | 0 | 1.14 |
| AG-891-N20 | GORG | 0 | 0 | NA | 2 | 0 | orphan | 0 | 1 | NA | 0 | 0 | NA | 69.51 | 0 | 1.52 |
| AG-414-J16 | GORG | 0 | 0 | NA | 1 | 0 | orphan | 0 | 0 | NA | 0 | 0 | NA | 69.48 | 0 | 1 |
| AG-913-A14 | GORG | 0 | 0 | NA | 1 | 0 | orphan | 0 | 0 | NA | 0 | 0 | NA | 69.47 | 0 | 1.09 |
| AH-287-B08 | GORG | 0 | 0 | NA | 0 | 0 | NA | 0 | 0 | NA | 0 | 1 | orphan | 69.43 | 0 | 1.16 |
| AH-324-F23 | GORG | 0 | 0 | NA | 1 | 0 | orphan | 0 | 0 | NA | 0 | 0 | NA | 69.38 | 0 | 2.37 |
| AG-337-O06 | GORG | 0 | 0 | NA | 1 | 0 | orphan | 0 | 0 | NA | 0 | 0 | NA | 69.33 | 0 | 1.18 |
| AG-893-K04 | GORG | 0 | 0 | NA | 2 | 0 | orphan | 0 | 0 | NA | 0 | 0 | NA | 69.33 | 0 | 0.82 |
| AG-907-B03 | GORG | 0 | 0 | NA | 1 | 0 | orphan | 0 | 0 | NA | 0 | 0 | NA | 69.33 | 0 | 0.85 |
| AG-911-J10 | GORG | 0 | 0 | NA | 1 | 0 | orphan | 0 | 0 | NA | 0 | 0 | NA | 69.33 | 0 | 1.49 |
| AG-343-C05 | GORG | 1 | 1 | complete | 2 | 0 | orphan | 0 | 0 | NA | 0 | 0 | NA | 69.32 | 0 | 1.8 |
| AG-447-K22 | GORG | 1 | 1 | complete | 0 | 0 | NA | 0 | 0 | NA | 0 | 0 | NA | 69.32 | 0 | 1.25 |
| AG-470-F09 | GORG | 1 | 1 | complete | 1 | 0 | orphan | 0 | 0 | NA | 0 | 0 | NA | 69.3 | 0 | 0.93 |
| AG-430-E17 | GORG | 0 | 0 | NA | 1 | 0 | orphan | 0 | 0 | NA | 0 | 0 | NA | 69.28 | 0 | 1.09 |
| AG-447-J18 | GORG | 0 | 0 | NA | 2 | 0 | orphan | 0 | 0 | NA | 0 | 0 | NA | 69.28 | 0 | 1.17 |

| | | | | | | | | | | | | | | | | |
|---|---|---|---|---|---|---|---|---|---|---|---|---|---|---|---|---|
| AG-908-E17 | GORG | 0 | 0 | NA | 1 | 0 | orphan | 0 | 1 | NA | 0 | 0 | NA | 69.26 | 0 | 0.69 |
| AG-919-G23 | GORG | 2 | 1 | complete | 0 | 0 | NA | 0 | 0 | NA | 0 | 0 | NA | 69.26 | 0 | 0.71 |
| AG-907-M08 | GORG | 0 | 0 | NA | 2 | 0 | orphan | 0 | 0 | NA | 0 | 0 | NA | 69.23 | 0 | 0.84 |
| AG-390-L17 | GORG | 0 | 0 | NA | 2 | 0 | orphan | 0 | 0 | NA | 0 | 0 | NA | 69.18 | 0 | 0.91 |
| AG-914-O19 | GORG | 0 | 0 | NA | 1 | 0 | orphan | 0 | 0 | NA | 0 | 0 | NA | 69.18 | 0 | 0.72 |
| AG-912-P10 | GORG | 0 | 0 | NA | 1 | 0 | orphan | 0 | 1 | NA | 0 | 0 | NA | 69.16 | 0 | 0.84 |
| AG-313-L18 | GORG | 1 | 1 | complete | 1 | 0 | orphan | 0 | 0 | NA | 0 | 0 | NA | 69.15 | 0 | 1.3 |
| AG-390-O21 | GORG | 0 | 0 | NA | 1 | 0 | orphan | 0 | 0 | NA | 0 | 0 | NA | 69.09 | 0 | 2.58 |
| AG-337-C13 | GORG | 0 | 0 | NA | 1 | 1 | orphan | 0 | 0 | NA | 0 | 0 | NA | 69.08 | 0 | 0.74 |
| AG-913-J17 | GORG | 0 | 0 | NA | 1 | 0 | orphan | 0 | 0 | NA | 0 | 0 | NA | 69.06 | 0 | 0.96 |
| AG-917-I09 | GORG | 0 | 0 | NA | 3 | 0 | orphan | 0 | 0 | NA | 0 | 0 | NA | 69.06 | 0 | 1.09 |
| AG-404-G08 | GORG | 0 | 0 | NA | 1 | 0 | orphan | 0 | 0 | NA | 0 | 0 | NA | 69.04 | 0 | 1 |
| AG-325-E07 | GORG | 0 | 0 | NA | 3 | 0 | orphan | 0 | 0 | NA | 0 | 0 | NA | 69.02 | 0 | 1.12 |
| AG-349-F19 | GORG | 0 | 0 | NA | 1 | 0 | orphan | 0 | 0 | NA | 0 | 0 | NA | 69.01 | 0 | 0.99 |
| AG-919-O05 | GORG | 0 | 0 | NA | 2 | 1 | orphan | 0 | 0 | NA | 0 | 0 | NA | 69 | 0 | 0.86 |
| AG-895-O17 | GORG | 0 | 0 | NA | 1 | 0 | orphan | 0 | 0 | NA | 0 | 0 | NA | 68.99 | 0 | 0.7 |
| AG-430-M14 | GORG | 0 | 0 | NA | 3 | 0 | orphan | 0 | 0 | NA | 0 | 0 | NA | 68.97 | 0 | 0.88 |
| AG-447-P04 | GORG | 0 | 0 | NA | 1 | 0 | orphan | 0 | 0 | NA | 0 | 0 | NA | 68.93 | 0 | 1.12 |
| AG-907-B23 | GORG | 0 | 0 | NA | 1 | 0 | orphan | 0 | 0 | NA | 0 | 0 | NA | 68.93 | 0 | 1.06 |
| AG-891-F11 | GORG | 0 | 0 | NA | 0 | 0 | NA | 0 | 0 | NA | 0 | 2 | orphan | 68.92 | 0 | 1.09 |
| AH-326-C06 | GORG | 1 | 1 | complete | 0 | 0 | NA | 0 | 0 | NA | 0 | 0 | NA | 68.92 | 0 | 1.26 |
| AG-439-O21 | GORG | 1 | 1 | complete | 0 | 0 | NA | 0 | 0 | NA | 0 | 0 | NA | 68.89 | 0 | 1.08 |
| AG-457-A20 | GORG | 0 | 0 | NA | 1 | 0 | orphan | 0 | 0 | NA | 0 | 0 | NA | 68.89 | 0 | 1.09 |
| AG-891-O22 | GORG | 0 | 0 | NA | 1 | 0 | orphan | 0 | 0 | NA | 0 | 0 | NA | 68.89 | 0 | 1.12 |
| AG-343-P20 | GORG | 0 | 0 | NA | 1 | 0 | orphan | 0 | 0 | NA | 0 | 0 | NA | 68.87 | 0 | 2.09 |

| | | | | | | | | | | | | | | | | |
|---|---|---|---|---|---|---|---|---|---|---|---|---|---|---|---|---|
| AG-426-C20 | GORG | 0 | 0 | NA | 2 | 0 | orphan | 0 | 0 | NA | 0 | 0 | NA | 68.87 | 0 | 1.08 |
| AG-901-J09 | GORG | 0 | 0 | NA | 1 | 0 | orphan | 0 | 0 | NA | 0 | 0 | NA | 68.87 | 0 | 0.78 |
| AH-287-A09 | GORG | 0 | 0 | NA | 1 | 0 | orphan | 0 | 0 | NA | 0 | 0 | NA | 68.87 | 0 | 0.85 |
| AG-414-E08 | GORG | 0 | 0 | NA | 1 | 0 | orphan | 0 | 0 | NA | 0 | 0 | NA | 68.83 | 0 | 0.87 |
| AG-907-L10 | GORG | 2 | 2 | complete | 1 | 0 | orphan | 0 | 0 | NA | 0 | 0 | NA | 68.8 | 0 | 1.2 |
| AG-907-B17 | GORG | 0 | 0 | NA | 2 | 0 | orphan | 0 | 0 | NA | 0 | 0 | NA | 68.78 | 0 | 0.82 |
| AG-912-J15 | GORG | 0 | 0 | NA | 1 | 0 | orphan | 0 | 0 | NA | 0 | 0 | NA | 68.78 | 0 | 0.82 |
| AG-414-M08 | GORG | 0 | 0 | NA | 0 | 0 | NA | 0 | 0 | NA | 0 | 1 | orphan | 68.75 | 0 | 0.83 |
| AG-892-B09 | GORG | 0 | 0 | NA | 0 | 0 | NA | 1 | 0 | orphan | 0 | 0 | NA | 68.75 | 0 | 1.49 |
| AG-900-K04 | GORG | 0 | 0 | NA | 2 | 0 | orphan | 0 | 0 | NA | 0 | 0 | NA | 68.75 | 0 | 1 |
| AG-894-F05 | GORG | 0 | 0 | NA | 2 | 0 | orphan | 0 | 0 | NA | 0 | 0 | NA | 68.73 | 0 | 1.17 |
| AG-439-O04 | GORG | 0 | 0 | NA | 1 | 0 | orphan | 0 | 0 | NA | 0 | 0 | NA | 68.72 | 0 | 1.04 |
| AG-464-D09 | GORG | 0 | 0 | NA | 2 | 0 | orphan | 0 | 0 | NA | 0 | 0 | NA | 68.72 | 0 | 0.89 |
| AG-414-P05 | GORG | 0 | 0 | NA | 1 | 0 | orphan | 0 | 0 | NA | 0 | 0 | NA | 68.67 | 0 | 1.13 |
| AG-891-F15 | GORG | 0 | 0 | NA | 1 | 0 | orphan | 0 | 0 | NA | 0 | 0 | NA | 68.67 | 0 | 1.29 |
| AG-913-D08 | GORG | 0 | 0 | NA | 2 | 0 | orphan | 0 | 0 | NA | 0 | 0 | NA | 68.67 | 0 | 1.16 |
| AG-913-F06 | GORG | 0 | 0 | NA | 1 | 0 | orphan | 0 | 0 | NA | 0 | 0 | NA | 68.67 | 0 | 1.2 |
| AH-326-B09 | GORG | 0 | 0 | NA | 1 | 0 | orphan | 0 | 0 | NA | 0 | 0 | NA | 68.67 | 0 | 0.89 |
| AG-430-N14 | GORG | 1 | 0 | orphan | 1 | 0 | orphan | 0 | 0 | NA | 0 | 0 | NA | 68.66 | 0 | 1.7 |
| AG-430-N08 | GORG | 0 | 0 | NA | 2 | 1 | complete | 0 | 0 | NA | 0 | 0 | NA | 68.64 | 0 | 0.93 |
| AG-447-K19 | GORG | 0 | 0 | NA | 1 | 0 | orphan | 0 | 1 | NA | 0 | 0 | NA | 68.6 | 0 | 1.34 |
| AG-920-K18 | GORG | 0 | 0 | NA | 1 | 0 | orphan | 0 | 0 | NA | 0 | 0 | NA | 68.6 | 0 | 1.11 |
| AG-917-M04 | GORG | 0 | 0 | NA | 1 | 0 | orphan | 0 | 0 | NA | 0 | 0 | NA | 68.58 | 0 | 1.06 |
| AG-915-A06 | GORG | 1 | 1 | complete | 1 | 0 | orphan | 0 | 0 | NA | 0 | 0 | NA | 68.57 | 0 | 1.04 |
| AG-447-A19 | GORG | 0 | 0 | NA | 1 | 0 | orphan | 0 | 0 | NA | 0 | 0 | NA | 68.56 | 0 | 1.15 |
| AG-918-B20 | GORG | 0 | 0 | NA | 2 | 0 | orphan | 0 | 0 | NA | 0 | 0 | NA | 68.54 | 0 | 0.89 |

| | | | | | | | | | | | | | | | | |
|---|---|---|---|---|---|---|---|---|---|---|---|---|---|---|---|---|
| AG-903-M14 | GORG | 0 | 0 | NA | 2 | 0 | orphan | 0 | 0 | NA | 0 | 0 | NA | 68.52 | 0 | 0.72 |
| AG-917-A21 | GORG | 0 | 0 | NA | 1 | 0 | orphan | 0 | 0 | NA | 0 | 0 | NA | 68.52 | 0 | 1.11 |
| AG-912-J21 | GORG | 0 | 0 | NA | 1 | 0 | orphan | 0 | 0 | NA | 0 | 0 | NA | 68.51 | 0 | 0.92 |
| AH-287-E21 | GORG | 0 | 0 | NA | 2 | 0 | orphan | 0 | 0 | NA | 0 | 0 | NA | 68.49 | 0 | 0.85 |
| AG-894-P19 | GORG | 0 | 0 | NA | 1 | 0 | orphan | 0 | 0 | NA | 0 | 0 | NA | 68.46 | 0 | 1.03 |
| AG-426-M10 | GORG | 0 | 0 | NA | 1 | 0 | orphan | 0 | 0 | NA | 0 | 0 | NA | 68.44 | 0 | 1.25 |
| AG-919-F05 | GORG | 0 | 0 | NA | 1 | 0 | orphan | 0 | 0 | NA | 0 | 0 | NA | 68.42 | 0 | 0.92 |
| AG-447-B20 | GORG | 0 | 0 | NA | 1 | 0 | orphan | 0 | 0 | NA | 0 | 0 | NA | 68.4 | 0 | 0.88 |
| AG-893-J16 | GORG | 1 | 1 | complete | 0 | 0 | NA | 0 | 0 | NA | 0 | 1 | orphan | 68.39 | 0 | 1.08 |
| AG-892-L04 | GORG | 1 | 0 | orphan | 0 | 0 | NA | 0 | 0 | NA | 0 | 0 | NA | 68.39 | 0 | 1.07 |
| AG-313-J22 | GORG | 0 | 0 | NA | 1 | 0 | orphan | 0 | 0 | NA | 0 | 0 | NA | 68.38 | 0 | 1.29 |
| AG-365-D18 | GORG | 0 | 0 | NA | 3 | 0 | orphan | 0 | 0 | NA | 0 | 0 | NA | 68.38 | 0 | 0.96 |
| AG-325-E04 | GORG | 0 | 1 | NA | 0 | 0 | NA | 0 | 0 | NA | 0 | 0 | NA | 68.36 | 0 | 1.67 |
| AG-901-D09 | GORG | 0 | 0 | NA | 2 | 0 | orphan | 0 | 0 | NA | 0 | 0 | NA | 68.33 | 0 | 0.88 |
| AG-916-G07 | GORG | 0 | 0 | NA | 1 | 0 | orphan | 0 | 0 | NA | 0 | 0 | NA | 68.33 | 0 | 1.22 |
| AG-920-J03 | GORG | 0 | 0 | NA | 1 | 0 | orphan | 0 | 0 | NA | 0 | 0 | NA | 68.33 | 0 | 0.91 |
| AG-333-F13 | GORG | 0 | 0 | NA | 1 | 0 | orphan | 0 | 0 | NA | 0 | 0 | NA | 68.29 | 0 | 0.71 |
| AG-916-J16 | GORG | 0 | 0 | NA | 1 | 0 | orphan | 0 | 0 | NA | 0 | 0 | NA | 68.29 | 0 | 1.16 |
| AG-457-D17 | GORG | 0 | 0 | NA | 2 | 0 | orphan | 0 | 0 | NA | 0 | 0 | NA | 68.27 | 0 | 1.28 |
| AG-912-I19 | GORG | 0 | 0 | NA | 2 | 0 | orphan | 0 | 0 | NA | 0 | 0 | NA | 68.25 | 0 | 1.02 |
| AG-422-C11 | GORG | 0 | 0 | NA | 0 | 0 | NA | 0 | 0 | NA | 0 | 1 | orphan | 68.21 | 0 | 1.12 |
| AG-422-O15 | GORG | 1 | 1 | complete | 0 | 0 | NA | 0 | 0 | NA | 0 | 0 | NA | 68.12 | 0 | 1.01 |
| AG-893-P08 | GORG | 0 | 0 | NA | 1 | 0 | orphan | 0 | 0 | NA | 0 | 0 | NA | 68.1 | 0 | 1.04 |
| AG-313-E13 | GORG | 0 | 0 | NA | 1 | 0 | orphan | 0 | 0 | NA | 0 | 0 | NA | 68.08 | 0 | 1.27 |
| AH-287-E16 | GORG | 0 | 0 | NA | 1 | 0 | orphan | 0 | 0 | NA | 0 | 2 | orphan | 68.07 | 0 | 1.1 |
| AG-426-L20 | GORG | 0 | 0 | NA | 1 | 0 | orphan | 0 | 0 | NA | 0 | 0 | NA | 68.07 | 0 | 0.81 |

| | | | | | | | | | | | | | | | | |
|---|---|---|---|---|---|---|---|---|---|---|---|---|---|---|---|---|
| AG-901-M08 | GORG | 0 | 0 | NA | 1 | 0 | orphan | 0 | 0 | NA | 0 | 0 | NA | 68.07 | 0 | 0.92 |
| AG-917-F15 | GORG | 2 | 1 | complete | 0 | 0 | NA | 0 | 0 | NA | 0 | 0 | NA | 68.07 | 0 | 0.78 |
| AG-920-P06 | GORG | 0 | 0 | NA | 1 | 0 | orphan | 0 | 0 | NA | 0 | 0 | NA | 68.07 | 0 | 0.95 |
| AG-894-L16 | GORG | 0 | 0 | NA | 1 | 0 | orphan | 0 | 0 | NA | 0 | 0 | NA | 68.06 | 0 | 1.22 |
| AH-287-C22 | GORG | 0 | 0 | NA | 1 | 0 | orphan | 0 | 0 | NA | 0 | 0 | NA | 68.03 | 0 | 1.2 |
| AG-404-J21 | GORG | 0 | 0 | NA | 1 | 0 | orphan | 0 | 0 | NA | 0 | 0 | NA | 68.02 | 0 | 0.91 |
| AG-910-I09 | GORG | 0 | 0 | NA | 1 | 0 | orphan | 0 | 0 | NA | 0 | 0 | NA | 68.02 | 0 | 0.97 |
| AG-917-O22 | GORG | 0 | 0 | NA | 1 | 0 | orphan | 0 | 0 | NA | 0 | 0 | NA | 68.02 | 0 | 0.89 |
| AH-326-B07 | GORG | 0 | 0 | NA | 1 | 0 | orphan | 0 | 0 | NA | 0 | 0 | NA | 68.02 | 0 | 0.91 |
| AG-430-I08 | GORG | 0 | 0 | NA | 1 | 0 | orphan | 0 | 0 | NA | 0 | 0 | NA | 68.01 | 0 | 1.06 |
| AG-319-F09 | GORG | 0 | 0 | NA | 1 | 0 | orphan | 1 | 0 | orphan | 0 | 2 | orphan | 67.96 | 0 | 0.85 |
| AG-319-J13 | GORG | 1 | 1 | complete | 1 | 0 | orphan | 0 | 0 | NA | 0 | 0 | NA | 67.95 | 0 | 1.21 |
| AH-321-C15 | GORG | 0 | 0 | NA | 1 | 0 | orphan | 0 | 0 | NA | 0 | 2 | orphan | 67.93 | 0 | 1.08 |
| AG-360-A05 | GORG | 0 | 0 | NA | 1 | 0 | orphan | 0 | 0 | NA | 0 | 0 | NA | 67.92 | 0 | 0.73 |
| AG-447-O06 | GORG | 0 | 0 | NA | 1 | 0 | orphan | 0 | 0 | NA | 0 | 0 | NA | 67.92 | 0 | 1.08 |
| AG-464-B15 | GORG | 0 | 0 | NA | 1 | 0 | orphan | 0 | 0 | NA | 0 | 0 | NA | 67.92 | 0 | 0.97 |
| AG-901-P18 | GORG | 0 | 0 | NA | 2 | 0 | orphan | 0 | 0 | NA | 0 | 0 | NA | 67.92 | 0 | 0.9 |
| AG-905-M17 | GORG | 0 | 0 | NA | 1 | 0 | orphan | 0 | 0 | NA | 0 | 0 | NA | 67.92 | 0 | 0.99 |
| AG-909-D23 | GORG | 0 | 0 | NA | 1 | 0 | orphan | 0 | 0 | NA | 0 | 0 | NA | 67.92 | 0 | 0.85 |
| AG-911-A10 | GORG | 0 | 0 | NA | 1 | 0 | orphan | 0 | 0 | NA | 0 | 0 | NA | 67.92 | 0 | 0.96 |
| AH-324-D10 | GORG | 0 | 0 | NA | 1 | 0 | orphan | 0 | 0 | NA | 0 | 0 | NA | 67.92 | 0 | 1.08 |
| AG-899-M02 | GORG | 0 | 0 | NA | 1 | 0 | orphan | 0 | 0 | NA | 0 | 0 | NA | 67.91 | 0 | 1.03 |
| AG-905-G05 | GORG | 0 | 0 | NA | 2 | 0 | orphan | 0 | 0 | NA | 0 | 1 | orphan | 67.9 | 0 | 1.15 |
| AG-325-A06 | GORG | 0 | 0 | NA | 1 | 0 | orphan | 0 | 0 | NA | 0 | 0 | NA | 67.9 | 0 | 0.73 |
| AG-898-A11 | GORG | 0 | 1 | NA | 1 | 1 | complete | 0 | 0 | NA | 0 | 0 | NA | 67.87 | 0 | 1.1 |

| | | | | | | | | | | | | | | | | |
|---|---|---|---|---|---|---|---|---|---|---|---|---|---|---|---|---|
| AG-916-C05 | GORG | 0 | 0 | NA | 1 | 0 | orphan | 0 | 0 | NA | 0 | 0 | NA | 67.87 | 0 | 1.08 |
| AG-910-J04 | GORG | 0 | 0 | NA | 3 | 0 | orphan | 0 | 0 | NA | 0 | 0 | NA | 67.85 | 0 | 1.2 |
| AG-920-C23 | GORG | 0 | 0 | NA | 1 | 0 | orphan | 0 | 0 | NA | 0 | 1 | orphan | 67.82 | 0 | 1.03 |
| AG-914-B23 | GORG | 0 | 0 | NA | 1 | 0 | orphan | 0 | 0 | NA | 0 | 0 | NA | 67.81 | 0 | 0.74 |
| AG-893-N14 | GORG | 0 | 0 | NA | 1 | 0 | orphan | 0 | 0 | NA | 0 | 0 | NA | 67.8 | 0 | 0.83 |
| AG-333-P14 | GORG | 1 | 0 | orphan | 0 | 0 | NA | 0 | 0 | NA | 0 | 0 | NA | 67.77 | 0 | 0.79 |
| AG-917-M21 | GORG | 0 | 0 | NA | 1 | 0 | orphan | 0 | 0 | NA | 0 | 0 | NA | 67.76 | 0 | 1.06 |
| AG-349-F21 | GORG | 0 | 0 | NA | 1 | 0 | orphan | 0 | 0 | NA | 0 | 0 | NA | 67.74 | 0 | 0.88 |
| AG-898-O03 | GORG | 0 | 0 | NA | 1 | 0 | orphan | 0 | 0 | NA | 0 | 0 | NA | 67.68 | 0 | 1.01 |
| AH-321-K19 | GORG | 0 | 0 | NA | 1 | 0 | orphan | 0 | 0 | NA | 0 | 0 | NA | 67.63 | 0 | 0.85 |
| AG-349-G19 | GORG | 0 | 0 | NA | 2 | 0 | orphan | 0 | 0 | NA | 0 | 0 | NA | 67.61 | 0 | 0.93 |
| AG-899-N23 | GORG | 1 | 1 | complete | 0 | 0 | NA | 0 | 0 | NA | 0 | 0 | NA | 67.59 | 0 | 1.25 |
| AG-911-L13 | GORG | 0 | 0 | NA | 1 | 0 | orphan | 0 | 0 | NA | 0 | 0 | NA | 67.59 | 0 | 0.9 |
| AG-313-F11 | GORG | 0 | 0 | NA | 2 | 0 | orphan | 0 | 0 | NA | 0 | 0 | NA | 67.56 | 0 | 1.25 |
| AG-894-F21 | GORG | 0 | 0 | NA | 1 | 0 | orphan | 0 | 0 | NA | 0 | 0 | NA | 67.56 | 0 | 1.07 |
| AG-891-M11 | GORG | 0 | 0 | NA | 1 | 0 | orphan | 0 | 0 | NA | 0 | 0 | NA | 67.55 | 0 | 0.79 |
| AG-325-B13 | GORG | 0 | 0 | NA | 2 | 1 | complete | 0 | 0 | NA | 0 | 0 | NA | 67.47 | 0 | 1.27 |
| AG-404-C04 | GORG | 0 | 0 | NA | 3 | 1 | complete | 0 | 0 | NA | 0 | 0 | NA | 67.47 | 0 | 1.05 |
| AG-453-D17 | GORG | 0 | 0 | NA | 1 | 0 | orphan | 0 | 0 | NA | 0 | 0 | NA | 67.47 | 0 | 0.77 |
| AG-461-L03 | GORG | 0 | 0 | NA | 3 | 0 | orphan | 0 | 0 | NA | 0 | 0 | NA | 67.47 | 0 | 0.9 |
| AG-892-J13 | GORG | 0 | 0 | NA | 3 | 1 | orphan | 0 | 0 | NA | 0 | 0 | NA | 67.47 | 0 | 1.2 |
| AG-896-B08 | GORG | 0 | 0 | NA | 1 | 0 | orphan | 0 | 0 | NA | 0 | 0 | NA | 67.47 | 0 | 1.27 |
| AG-904-B07 | GORG | 0 | 0 | NA | 2 | 1 | orphan | 0 | 0 | NA | 0 | 0 | NA | 67.47 | 0 | 0.72 |
| AG-913-A06 | GORG | 1 | 1 | complete | 1 | 0 | orphan | 0 | 0 | NA | 0 | 0 | NA | 67.47 | 0 | 0.98 |
| AG-916-I08 | GORG | 0 | 0 | NA | 1 | 0 | orphan | 0 | 1 | NA | 0 | 0 | NA | 67.47 | 0 | 1.19 |

| | | | | | | | | | | | | | | | | |
|---|---|---|---|---|---|---|---|---|---|---|---|---|---|---|---|---|
| AH-287-A01 | GORG | 0 | 0 | NA | 2 | 0 | orphan | 0 | 0 | NA | 0 | 0 | NA | 67.47 | 0 | 1.21 |
| AG-893-M18 | GORG | 0 | 0 | NA | 1 | 0 | orphan | 0 | 0 | NA | 0 | 0 | NA | 67.42 | 0 | 1.76 |
| AH-287-J22 | GORG | 0 | 0 | NA | 1 | 0 | orphan | 0 | 0 | NA | 0 | 0 | NA | 67.37 | 0 | 1.7 |
| AG-325-P18 | GORG | 0 | 0 | NA | 1 | 0 | orphan | 0 | 0 | NA | 0 | 0 | NA | 67.36 | 0 | 0.87 |
| AG-453-C10 | GORG | 1 | 1 | complete | 1 | 0 | orphan | 0 | 0 | NA | 0 | 0 | NA | 67.36 | 0 | 1.2 |
| AG-913-P01 | GORG | 0 | 0 | NA | 1 | 0 | orphan | 3 | 1 | complete | 0 | 0 | NA | 67.31 | 0 | 0.97 |
| AG-453-K23 | GORG | 0 | 0 | NA | 1 | 0 | orphan | 0 | 0 | NA | 0 | 0 | NA | 67.3 | 0 | 0.91 |
| AG-313-A14 | GORG | 0 | 0 | NA | 1 | 0 | orphan | 0 | 0 | NA | 0 | 0 | NA | 67.29 | 0 | 1.01 |
| AG-912-N19 | GORG | 0 | 1 | NA | 0 | 0 | NA | 0 | 0 | NA | 0 | 0 | NA | 67.21 | 0 | 1.19 |
| AG-920-M04 | GORG | 0 | 0 | NA | 1 | 0 | orphan | 0 | 0 | NA | 0 | 0 | NA | 67.16 | 0 | 1.42 |
| AH-326-P21 | GORG | 0 | 0 | NA | 2 | 0 | orphan | 0 | 0 | NA | 0 | 0 | NA | 67.14 | 0 | 0.77 |
| AG-892-C19 | GORG | 0 | 0 | NA | 3 | 0 | orphan | 0 | 0 | NA | 0 | 0 | NA | 67.11 | 0 | 1.73 |
| AG-461-J14 | GORG | 0 | 0 | NA | 1 | 0 | orphan | 0 | 0 | NA | 0 | 0 | NA | 67.1 | 0 | 1.3 |
| AG-464-B21 | GORG | 0 | 0 | NA | 1 | 0 | orphan | 0 | 0 | NA | 0 | 0 | NA | 67.07 | 0 | 0.87 |
| AG-917-O15 | GORG | 0 | 0 | NA | 1 | 0 | orphan | 0 | 0 | NA | 0 | 0 | NA | 67.05 | 0 | 0.86 |
| AG-915-E04 | GORG | 1 | 1 | complete | 0 | 0 | NA | 0 | 0 | NA | 0 | 0 | NA | 67.03 | 0 | 0.72 |
| AG-911-M11 | GORG | 1 | 1 | complete | 1 | 0 | orphan | 0 | 0 | NA | 0 | 0 | NA | 66.99 | 0 | 0.93 |
| AG-903-J11 | GORG | 0 | 0 | NA | 0 | 0 | NA | 0 | 0 | NA | 0 | 1 | orphan | 66.98 | 0 | 1.1 |
| AG-900-N20 | GORG | 0 | 0 | NA | 1 | 0 | orphan | 0 | 0 | NA | 0 | 0 | NA | 66.98 | 0 | 0.77 |
| AG-917-A22 | GORG | 0 | 0 | NA | 2 | 0 | orphan | 0 | 0 | NA | 0 | 0 | NA | 66.98 | 0 | 1.03 |
| AG-913-C20 | GORG | 0 | 0 | NA | 1 | 0 | orphan | 0 | 0 | NA | 0 | 0 | NA | 66.96 | 0 | 0.96 |
| AG-894-J02 | GORG | 0 | 0 | NA | 3 | 0 | orphan | 0 | 0 | NA | 0 | 0 | NA | 66.94 | 0 | 1.17 |
| AG-319-E11 | GORG | 0 | 1 | NA | 0 | 0 | NA | 0 | 0 | NA | 0 | 0 | NA | 66.9 | 0 | 1.27 |
| AG-911-C02 | GORG | 0 | 0 | NA | 1 | 0 | orphan | 0 | 0 | NA | 0 | 0 | NA | 66.87 | 0 | 1.12 |
| AG-912-L14 | GORG | 0 | 0 | NA | 1 | 0 | orphan | 0 | 0 | NA | 0 | 0 | NA | 66.87 | 0 | 1.09 |

| | | | | | | | | | | | | | | | | |
|---|---|---|---|---|---|---|---|---|---|---|---|---|---|---|---|---|
| AH-326-F13 | GORG | 1 | 1 | complete | 2 | 1 | orphan | 0 | 0 | NA | 0 | 0 | NA | 66.87 | 0 | 1.25 |
| AG-914-N13 | GORG | 0 | 0 | NA | 0 | 0 | NA | 0 | 0 | NA | 0 | 1 | orphan | 66.85 | 0 | 1.28 |
| AG-917-F20 | GORG | 0 | 0 | NA | 1 | 0 | orphan | 0 | 0 | NA | 0 | 1 | orphan | 66.85 | 0 | 1.69 |
| AG-325-G22 | GORG | 0 | 0 | NA | 2 | 0 | orphan | 0 | 0 | NA | 0 | 0 | NA | 66.84 | 0 | 0.8 |
| AG-894-P21 | GORG | 0 | 0 | NA | 2 | 0 | orphan | 0 | 0 | NA | 0 | 0 | NA | 66.83 | 0 | 1.02 |
| AG-447-M21 | GORG | 0 | 0 | NA | 1 | 0 | orphan | 0 | 0 | NA | 0 | 0 | NA | 66.81 | 0 | 0.79 |
| AG-900-I14 | GORG | 0 | 0 | NA | 1 | 1 | complete | 0 | 0 | NA | 0 | 0 | NA | 66.72 | 0 | 1.12 |
| AG-917-F21 | GORG | 0 | 0 | NA | 1 | 0 | orphan | 0 | 1 | NA | 0 | 0 | NA | 66.7 | 0 | 0.86 |
| AG-464-K18 | GORG | 0 | 0 | NA | 2 | 0 | orphan | 0 | 0 | NA | 0 | 0 | NA | 66.67 | 0 | 0.83 |
| AG-464-P19 | GORG | 0 | 0 | NA | 3 | 0 | orphan | 0 | 0 | NA | 0 | 0 | NA | 66.67 | 0 | 1.12 |
| AG-898-C15 | GORG | 1 | 1 | complete | 0 | 0 | NA | 0 | 0 | NA | 0 | 0 | NA | 66.67 | 0 | 1.11 |
| AG-901-F09 | GORG | 0 | 0 | NA | 1 | 0 | orphan | 0 | 0 | NA | 0 | 0 | NA | 66.67 | 0 | 1.2 |
| AG-918-M13 | GORG | 0 | 0 | NA | 1 | 0 | orphan | 0 | 0 | NA | 0 | 0 | NA | 66.65 | 0 | 1.05 |
| AG-390-L06 | GORG | 0 | 0 | NA | 1 | 0 | orphan | 0 | 0 | NA | 0 | 0 | NA | 66.59 | 0 | 1.25 |
| AG-333-G21 | GORG | 0 | 0 | NA | 1 | 0 | orphan | 0 | 0 | NA | 0 | 0 | NA | 66.57 | 0 | 0.84 |
| AG-917-J22 | GORG | 0 | 0 | NA | 2 | 0 | orphan | 0 | 0 | NA | 0 | 0 | NA | 66.57 | 0 | 0.8 |
| AG-905-O09 | GORG | 0 | 0 | NA | 1 | 0 | orphan | 0 | 0 | NA | 0 | 0 | NA | 66.52 | 0 | 0.76 |
| AG-917-I10 | GORG | 0 | 0 | NA | 2 | 0 | orphan | 0 | 0 | NA | 0 | 0 | NA | 66.51 | 0 | 1.16 |
| AH-324-J21 | GORG | 0 | 0 | NA | 1 | 0 | orphan | 0 | 0 | NA | 0 | 0 | NA | 66.49 | 0 | 1.17 |
| AG-470-O11 | GORG | 0 | 0 | NA | 4 | 0 | orphan | 0 | 0 | NA | 0 | 1 | orphan | 66.48 | 0 | 1.03 |
| AG-439-B06 | GORG | 0 | 0 | NA | 2 | 1 | complete | 0 | 0 | NA | 0 | 0 | NA | 66.48 | 0 | 0.73 |
| AG-912-G03 | GORG | 1 | 1 | complete | 1 | 0 | orphan | 0 | 0 | NA | 0 | 0 | NA | 66.47 | 0 | 1 |
| AG-914-I04 | GORG | 0 | 0 | NA | 1 | 0 | orphan | 0 | 0 | NA | 0 | 0 | NA | 66.46 | 0 | 0.71 |
| AG-916-B05 | GORG | 0 | 0 | NA | 1 | 0 | orphan | 0 | 0 | NA | 0 | 0 | NA | 66.44 | 0 | 1.06 |

| | | | | | | | | | | | | | | | | |
|---|---|---|---|---|---|---|---|---|---|---|---|---|---|---|---|---|
| AG-360-B02 | GORG | 0 | 0 | NA | 1 | 0 | orphan | 0 | 0 | NA | 0 | 0 | NA | 66.43 | 0 | 0.96 |
| AG-453-E19 | GORG | 0 | 0 | NA | 2 | 0 | orphan | 0 | 0 | NA | 0 | 0 | NA | 66.43 | 0 | 0.89 |
| AG-913-L07 | GORG | 1 | 1 | complete | 1 | 0 | orphan | 0 | 0 | NA | 0 | 0 | NA | 66.43 | 0 | 0.88 |
| AG-410-K20 | GORG | 1 | 1 | complete | 2 | 0 | orphan | 0 | 0 | NA | 0 | 0 | NA | 66.42 | 0 | 1.17 |
| AG-892-K09 | GORG | 0 | 0 | NA | 0 | 0 | NA | 1 | 1 | orphan | 0 | 0 | NA | 66.42 | 0 | 1.05 |
| AG-905-J08 | GORG | 0 | 0 | NA | 1 | 0 | orphan | 0 | 0 | NA | 0 | 0 | NA | 66.4 | 0 | 1.23 |
| AG-918-B19 | GORG | 0 | 0 | NA | 0 | 0 | NA | 0 | 0 | NA | 0 | 1 | orphan | 66.37 | 0 | 1.13 |
| AG-410-J15 | GORG | 0 | 0 | NA | 1 | 0 | orphan | 0 | 0 | NA | 0 | 0 | NA | 66.37 | 0 | 0.87 |
| AG-414-N04 | GORG | 0 | 0 | NA | 3 | 1 | complete | 0 | 0 | NA | 0 | 0 | NA | 66.37 | 0 | 0.8 |
| AG-343-O05 | GORG | 0 | 0 | NA | 2 | 0 | orphan | 0 | 0 | NA | 0 | 0 | NA | 66.3 | 0 | 1.08 |
| AG-359-L07 | GORG | 0 | 0 | NA | 1 | 0 | orphan | 0 | 0 | NA | 0 | 0 | NA | 66.27 | 0 | 1.01 |
| AG-360-M08 | GORG | 0 | 0 | NA | 1 | 0 | orphan | 0 | 0 | NA | 0 | 0 | NA | 66.27 | 0 | 0.74 |
| AG-426-F23 | GORG | 1 | 0 | orphan | 1 | 0 | orphan | 0 | 0 | NA | 0 | 0 | NA | 66.27 | 0 | 0.98 |
| AG-453-L03 | GORG | 0 | 0 | NA | 3 | 0 | orphan | 0 | 0 | NA | 0 | 0 | NA | 66.27 | 0 | 0.72 |
| AG-905-L17 | GORG | 0 | 0 | NA | 1 | 0 | orphan | 0 | 0 | NA | 0 | 0 | NA | 66.27 | 0 | 0.98 |
| AG-910-K07 | GORG | 0 | 0 | NA | 1 | 0 | orphan | 0 | 0 | NA | 0 | 0 | NA | 66.27 | 0 | 0.99 |
| AG-912-J11 | GORG | 0 | 0 | NA | 2 | 0 | orphan | 0 | 0 | NA | 0 | 0 | NA | 66.27 | 0 | 0.82 |
| AG-913-N07 | GORG | 0 | 0 | NA | 2 | 0 | orphan | 0 | 0 | NA | 0 | 0 | NA | 66.27 | 0 | 1.02 |
| AG-915-C03 | GORG | 0 | 0 | NA | 1 | 0 | orphan | 0 | 0 | NA | 0 | 0 | NA | 66.27 | 0 | 0.81 |
| AH-287-M14 | GORG | 0 | 0 | NA | 1 | 0 | orphan | 0 | 0 | NA | 0 | 0 | NA | 66.25 | 0 | 1.22 |
| AG-910-C02 | GORG | 0 | 0 | NA | 1 | 0 | orphan | 0 | 0 | NA | 0 | 0 | NA | 66.2 | 0 | 0.94 |
| AG-365-M21 | GORG | 0 | 0 | NA | 1 | 0 | orphan | 0 | 0 | NA | 0 | 0 | NA | 66.19 | 0 | 0.75 |
| AH-326-E21 | GORG | 0 | 0 | NA | 1 | 0 | orphan | 0 | 0 | NA | 0 | 0 | NA | 66.16 | 0 | 0.83 |
| AG-319-P13 | GORG | 1 | 1 | complete | 0 | 0 | NA | 0 | 0 | NA | 0 | 0 | NA | 66.15 | 0 | 1.25 |
| AG-333-C19 | GORG | 0 | 0 | NA | 2 | 0 | orphan | 0 | 0 | NA | 0 | 0 | NA | 66.15 | 0 | 0.76 |

| | | | | | | | | | | | | | | | | |
|---|---|---|---|---|---|---|---|---|---|---|---|---|---|---|---|---|
| AG-911-K23 | GORG | 0 | 0 | NA | 1 | 0 | orphan | 0 | 0 | NA | 0 | 0 | NA | 66.14 | 0 | 0.9 |
| AG-319-N20 | GORG | 1 | 1 | complete | 2 | 0 | orphan | 0 | 0 | NA | 0 | 0 | NA | 66.13 | 0 | 0.87 |
| AG-439-I13 | GORG | 0 | 0 | NA | 2 | 1 | complete | 0 | 0 | NA | 0 | 0 | NA | 66.13 | 0 | 1.31 |
| AG-365-O23 | GORG | 0 | 0 | NA | 2 | 0 | orphan | 0 | 0 | NA | 0 | 0 | NA | 66.12 | 0 | 1.09 |
| AG-435-K15 | GORG | 0 | 0 | NA | 1 | 0 | orphan | 0 | 0 | NA | 0 | 0 | NA | 66.1 | 0 | 1.79 |
| AG-390-M08 | GORG | 0 | 0 | NA | 2 | 1 | complete | 0 | 0 | NA | 0 | 0 | NA | 66.09 | 0 | 1.65 |
| AG-343-N19 | GORG | 0 | 0 | NA | 1 | 0 | orphan | 0 | 0 | NA | 0 | 0 | NA | 66.08 | 0 | 1.85 |
| AG-447-I23 | GORG | 0 | 0 | NA | 1 | 0 | orphan | 0 | 0 | NA | 0 | 0 | NA | 66.08 | 0 | 0.91 |
| AG-430-A20 | GORG | 1 | 1 | complete | 1 | 0 | orphan | 0 | 0 | NA | 0 | 0 | NA | 66.05 | 0 | 1.09 |
| AG-910-A22 | GORG | 0 | 0 | NA | 1 | 0 | orphan | 0 | 0 | NA | 0 | 0 | NA | 66.05 | 0 | 0.78 |
| AG-325-P08 | GORG | 0 | 0 | NA | 1 | 0 | orphan | 0 | 0 | NA | 0 | 0 | NA | 66.04 | 0 | 0.92 |
| AG-359-K09 | GORG | 0 | 0 | NA | 1 | 0 | orphan | 0 | 0 | NA | 0 | 0 | NA | 66.04 | 0 | 0.99 |
| AG-430-I15 | GORG | 0 | 0 | NA | 1 | 0 | orphan | 0 | 0 | NA | 0 | 0 | NA | 66.04 | 0 | 1.05 |
| AG-447-M11 | GORG | 0 | 0 | NA | 1 | 0 | orphan | 0 | 0 | NA | 0 | 0 | NA | 66.04 | 0 | 0.93 |
| AG-891-C19 | GORG | 0 | 0 | NA | 1 | 0 | orphan | 0 | 0 | NA | 0 | 0 | NA | 66.04 | 0 | 0.92 |
| AG-891-L21 | GORG | 0 | 0 | NA | 1 | 0 | orphan | 0 | 0 | NA | 0 | 0 | NA | 66.04 | 0 | 1.01 |
| AG-894-O19 | GORG | 0 | 0 | NA | 1 | 0 | orphan | 0 | 0 | NA | 0 | 0 | NA | 66.04 | 0 | 1.08 |
| AG-898-A14 | GORG | 0 | 0 | NA | 3 | 0 | orphan | 0 | 0 | NA | 0 | 0 | NA | 66.04 | 0 | 1.07 |
| AG-899-P18 | GORG | 0 | 0 | NA | 1 | 0 | orphan | 0 | 0 | NA | 0 | 0 | NA | 66.04 | 0 | 0.75 |
| AG-909-O11 | GORG | 0 | 0 | NA | 1 | 0 | orphan | 0 | 0 | NA | 0 | 0 | NA | 66.04 | 0 | 1.32 |
| AG-911-D08 | GORG | 1 | 1 | complete | 1 | 0 | orphan | 0 | 0 | NA | 0 | 0 | NA | 66.04 | 0 | 0.92 |
| AG-914-L09 | GORG | 0 | 0 | NA | 1 | 0 | orphan | 0 | 0 | NA | 0 | 0 | NA | 66.04 | 0 | 1.06 |
| AG-920-G10 | GORG | 1 | 1 | complete | 1 | 0 | orphan | 0 | 0 | NA | 0 | 0 | NA | 66.04 | 0 | 0.87 |
| AH-287-K21 | GORG | 0 | 0 | NA | 1 | 0 | orphan | 0 | 0 | NA | 0 | 0 | NA | 66.04 | 0 | 1.1 |
| AG-422-M11 | GORG | 0 | 0 | NA | 2 | 0 | orphan | 0 | 0 | NA | 0 | 0 | NA | 66.03 | 0 | 0.8 |

| | | | | | | | | | | | | | | | | |
|---|---|---|---|---|---|---|---|---|---|---|---|---|---|---|---|---|
| AG-898-F03 | GORG | 0 | 0 | NA | 1 | 0 | orphan | 0 | 0 | NA | 0 | 0 | NA | 66.03 | 0 | 1.22 |
| AG-900-B11 | GORG | 0 | 0 | NA | 0 | 0 | NA | 0 | 0 | NA | 0 | 1 | orphan | 65.99 | 0 | 1.17 |
| AG-896-O06 | GORG | 1 | 0 | orphan | 0 | 0 | NA | 0 | 0 | NA | 0 | 0 | NA | 65.98 | 0 | 0.9 |
| AG-337-A16 | GORG | 0 | 0 | NA | 1 | 0 | orphan | 0 | 0 | NA | 0 | 0 | NA | 65.97 | 0 | 0.8 |
| AG-337-K19 | GORG | 0 | 0 | NA | 3 | 0 | orphan | 0 | 0 | NA | 0 | 0 | NA | 65.95 | 0 | 0.82 |
| AH-321-B21 | GORG | 0 | 0 | NA | 4 | 0 | orphan | 0 | 0 | NA | 0 | 0 | NA | 65.95 | 0 | 0.93 |
| AG-464-L03 | GORG | 0 | 0 | NA | 2 | 0 | orphan | 0 | 0 | NA | 0 | 0 | NA | 65.94 | 0 | 1.21 |
| AG-897-D18 | GORG | 0 | 0 | NA | 4 | 0 | orphan | 0 | 0 | NA | 0 | 0 | NA | 65.94 | 0 | 1.05 |
| AG-457-L05 | GORG | 0 | 0 | NA | 2 | 0 | orphan | 0 | 0 | NA | 0 | 0 | NA | 65.91 | 0 | 0.74 |
| AG-919-P10 | GORG | 0 | 1 | NA | 0 | 0 | NA | 0 | 0 | NA | 0 | 0 | NA | 65.91 | 0 | 1.2 |
| AG-920-D13 | GORG | 0 | 0 | NA | 2 | 0 | orphan | 0 | 1 | NA | 0 | 0 | NA | 65.91 | 0 | 0.99 |
| AG-426-A04 | GORG | 0 | 0 | NA | 1 | 0 | orphan | 0 | 0 | NA | 0 | 0 | NA | 65.88 | 0 | 0.75 |
| AG-430-P10 | GORG | 0 | 0 | NA | 2 | 0 | orphan | 0 | 0 | NA | 0 | 0 | NA | 65.88 | 0 | 0.89 |
| AH-287-B11 | GORG | 0 | 0 | NA | 2 | 0 | orphan | 0 | 0 | NA | 0 | 0 | NA | 65.81 | 0 | 0.97 |
| AG-337-M06 | GORG | 0 | 0 | NA | 1 | 0 | orphan | 0 | 0 | NA | 0 | 0 | NA | 65.78 | 0 | 0.83 |
| AG-903-O06 | GORG | 0 | 0 | NA | 1 | 0 | orphan | 0 | 0 | NA | 0 | 1 | orphan | 65.77 | 0 | 1.01 |
| AG-349-M17 | GORG | 0 | 0 | NA | 0 | 0 | NA | 0 | 0 | NA | 0 | 1 | orphan | 65.76 | 0 | 1.06 |
| AG-337-B16 | GORG | 0 | 0 | NA | 1 | 0 | orphan | 0 | 1 | NA | 0 | 0 | NA | 65.76 | 0 | 1.11 |
| AH-324-P04 | GORG | 0 | 0 | NA | 3 | 0 | orphan | 0 | 0 | NA | 0 | 0 | NA | 65.75 | 0 | 0.79 |
| AG-891-O14 | GORG | 0 | 0 | NA | 1 | 0 | orphan | 0 | 0 | NA | 0 | 0 | NA | 65.71 | 0 | 0.87 |
| AH-324-N17 | GORG | 0 | 0 | NA | 1 | 0 | orphan | 0 | 0 | NA | 0 | 0 | NA | 65.69 | 0 | 0.81 |
| AG-461-G03 | GORG | 0 | 0 | NA | 1 | 0 | orphan | 0 | 0 | NA | 0 | 0 | NA | 65.67 | 0 | 0.84 |
| AH-326-N13 | GORG | 0 | 0 | NA | 4 | 1 | complete | 0 | 0 | NA | 0 | 0 | NA | 65.67 | 0 | 1.22 |
| AG-337-I13 | GORG | 0 | 0 | NA | 1 | 0 | orphan | 0 | 0 | NA | 0 | 0 | NA | 65.66 | 0 | 0.8 |
| AG-907-L05 | GORG | 0 | 0 | NA | 1 | 0 | orphan | 0 | 0 | NA | 0 | 0 | NA | 65.66 | 0 | 0.95 |
| AG-909-A20 | GORG | 0 | 0 | NA | 2 | 0 | orphan | 0 | 0 | NA | 0 | 0 | NA | 65.66 | 0 | 0.83 |

| | | | | | | | | | | | | | | | | |
|---|---|---|---|---|---|---|---|---|---|---|---|---|---|---|---|---|
| AG-909-E19 | GORG | 0 | 0 | NA | 1 | 0 | orphan | 0 | 0 | NA | 0 | 0 | NA | 65.66 | 0 | 0.72 |
| AG-909-P17 | GORG | 0 | 0 | NA | 2 | 0 | orphan | 0 | 1 | NA | 0 | 0 | NA | 65.66 | 0 | 1 |
| AG-919-A13 | GORG | 0 | 0 | NA | 1 | 0 | orphan | 0 | 0 | NA | 0 | 0 | NA | 65.66 | 0 | 0.88 |
| AH-324-A19 | GORG | 0 | 0 | NA | 3 | 0 | orphan | 0 | 0 | NA | 0 | 0 | NA | 65.63 | 0 | 0.72 |
| AG-911-A01 | GORG | 0 | 0 | NA | 1 | 0 | orphan | 0 | 0 | NA | 0 | 0 | NA | 65.59 | 0 | 0.68 |
| AG-892-O11 | GORG | 0 | 0 | NA | 1 | 0 | orphan | 0 | 0 | NA | 0 | 0 | NA | 65.53 | 0 | 1.01 |
| AG-319-D05 | GORG | 0 | 0 | NA | 0 | 0 | NA | 0 | 0 | NA | 0 | 1 | orpha n | 65.46 | 0 | 1.15 |
| AG-343-J22 | GORG | 0 | 0 | NA | 1 | 0 | orphan | 0 | 0 | NA | 0 | 0 | NA | 65.44 | 0 | 0.86 |
| AG-404-K16 | GORG | 0 | 0 | NA | 2 | 0 | orphan | 0 | 0 | NA | 0 | 0 | NA | 65.41 | 0 | 1.38 |
| AG-904-N15 | GORG | 0 | 0 | NA | 1 | 0 | orphan | 0 | 0 | NA | 0 | 0 | NA | 65.41 | 0 | 0.85 |
| AG-905-G02 | GORG | 0 | 0 | NA | 1 | 1 | orphan | 0 | 0 | NA | 0 | 0 | NA | 65.41 | 0 | 0.99 |
| AG-910-C21 | GORG | 0 | 0 | NA | 1 | 0 | orphan | 0 | 0 | NA | 0 | 0 | NA | 65.38 | 0 | 0.76 |
| AG-918-L17 | GORG | 0 | 0 | NA | 1 | 0 | orphan | 0 | 0 | NA | 0 | 0 | NA | 65.35 | 0 | 0.73 |
| AH-287-F17 | GORG | 0 | 0 | NA | 2 | 0 | orphan | 0 | 0 | NA | 0 | 0 | NA | 65.29 | 0 | 0.72 |
| AG-414-P19 | GORG | 0 | 0 | NA | 1 | 0 | orphan | 0 | 0 | NA | 0 | 0 | NA | 65.28 | 0 | 0.98 |
| AG-915-K20 | GORG | 0 | 0 | NA | 1 | 0 | orphan | 0 | 0 | NA | 0 | 1 | orpha n | 65.24 | 0 | 0.79 |
| AG-404-J18 | GORG | 0 | 0 | NA | 1 | 0 | orphan | 0 | 0 | NA | 0 | 0 | NA | 65.2 | 0 | 1.84 |
| AG-919-D18 | GORG | 0 | 0 | NA | 1 | 0 | orphan | 0 | 0 | NA | 0 | 0 | NA | 65.18 | 0 | 1.11 |
| AG-325-E11 | GORG | 0 | 0 | NA | 1 | 0 | orphan | 0 | 0 | NA | 0 | 0 | NA | 65.17 | 0 | 0.84 |
| AG-919-F13 | GORG | 0 | 0 | NA | 1 | 0 | orphan | 0 | 0 | NA | 0 | 0 | NA | 65.17 | 0 | 0.82 |
| AG-343-J10 | GORG | 0 | 0 | NA | 1 | 0 | orphan | 0 | 0 | NA | 0 | 0 | NA | 65.15 | 0 | 1.05 |
| AG-319-I20 | GORG | 0 | 0 | NA | 2 | 0 | orphan | 0 | 0 | NA | 0 | 0 | NA | 65.11 | 0 | 1.05 |
| AG-899-N16 | GORG | 1 | 1 | complet e | 0 | 0 | NA | 0 | 0 | NA | 0 | 0 | NA | 65.1 | 0 | 0.82 |
| AG-892-G14 | GORG | 0 | 0 | NA | 1 | 0 | orphan | 0 | 0 | NA | 0 | 0 | NA | 65.09 | 0 | 1.2 |
| AG-914-M13 | GORG | 1 | 1 | complet e | 0 | 0 | NA | 0 | 0 | NA | 0 | 0 | NA | 65.09 | 0 | 0.97 |
| AG-901-A01 | GORG | 0 | 0 | NA | 0 | 0 | NA | 0 | 0 | NA | 0 | 1 | orpha | 65.06 | 0 | 0.82 |

| | | | | | | | | | | | | | | | | |
|---|---|---|---|---|---|---|---|---|---|---|---|---|---|---|---|---|
| | | | | | | | | | | | | | n | | | |
| AG-325-J07 | GORG | 0 | 0 | NA | 1 | 0 | orphan | 0 | 0 | NA | 0 | 0 | NA | 65.06 | 0 | 1.02 |
| AG-901-G21 | GORG | 0 | 0 | NA | 2 | 0 | orphan | 0 | 0 | NA | 0 | 0 | NA | 65.06 | 0 | 1.06 |
| AG-903-F06 | GORG | 0 | 0 | NA | 1 | 0 | orphan | 0 | 0 | NA | 0 | 0 | NA | 65.06 | 0 | 1.03 |
| AG-905-L16 | GORG | 0 | 0 | NA | 1 | 0 | orphan | 0 | 0 | NA | 0 | 0 | NA | 65.06 | 0 | 0.77 |
| AG-917-F17 | GORG | 0 | 0 | NA | 1 | 0 | orphan | 0 | 0 | NA | 0 | 0 | NA | 65.06 | 0 | 0.91 |
| AG-464-A02 | GORG | 0 | 0 | NA | 1 | 0 | orphan | 0 | 0 | NA | 0 | 0 | NA | 65.05 | 0 | 0.73 |
| AG-895-I11 | GORG | 0 | 0 | NA | 1 | 0 | orphan | 1 | 1 | complete | 0 | 1 | orphan | 65 | 0 | 1.04 |
| AG-319-K19 | GORG | 0 | 0 | NA | 1 | 0 | orphan | 0 | 0 | NA | 0 | 0 | NA | 64.99 | 0 | 1.22 |
| AG-892-J17 | GORG | 0 | 0 | NA | 1 | 0 | orphan | 0 | 0 | NA | 0 | 0 | NA | 64.95 | 0 | 1.04 |
| AG-920-O16 | GORG | 0 | 0 | NA | 2 | 0 | orphan | 0 | 0 | NA | 0 | 0 | NA | 64.95 | 0 | 0.87 |
| AG-343-K22 | GORG | 0 | 0 | NA | 2 | 0 | orphan | 0 | 0 | NA | 0 | 0 | NA | 64.92 | 0 | 0.69 |
| AG-410-M23 | GORG | 0 | 0 | NA | 3 | 0 | orphan | 0 | 0 | NA | 0 | 0 | NA | 64.91 | 0 | 0.84 |
| AG-895-N09 | GORG | 1 | 1 | orphan | 0 | 0 | NA | 0 | 0 | NA | 0 | 0 | NA | 64.87 | 0 | 0.73 |
| AG-903-M17 | GORG | 0 | 0 | NA | 2 | 0 | orphan | 0 | 0 | NA | 0 | 0 | NA | 64.86 | 0 | 0.83 |
| AG-461-E06 | GORG | 0 | 0 | NA | 1 | 0 | orphan | 0 | 0 | NA | 0 | 0 | NA | 64.84 | 0 | 0.86 |
| AH-287-G16 | GORG | 0 | 0 | NA | 2 | 0 | orphan | 0 | 0 | NA | 0 | 0 | NA | 64.84 | 0 | 0.67 |
| AG-470-M09 | GORG | 0 | 0 | NA | 2 | 0 | orphan | 0 | 0 | NA | 0 | 0 | NA | 64.83 | 0 | 0.83 |
| AG-900-A17 | GORG | 0 | 0 | NA | 0 | 0 | NA | 0 | 0 | NA | 0 | 1 | orphan | 64.81 | 0 | 1.02 |
| AH-287-E15 | GORG | 0 | 0 | NA | 0 | 0 | NA | 0 | 0 | NA | 0 | 1 | orphan | 64.81 | 0 | 1.06 |
| AG-904-M22 | GORG | 0 | 0 | NA | 2 | 0 | orphan | 0 | 0 | NA | 0 | 0 | NA | 64.77 | 0 | 0.79 |
| AG-333-I22 | GORG | 0 | 0 | NA | 3 | 1 | complete | 0 | 0 | NA | 0 | 1 | orphan | 64.76 | 0 | 1.42 |
| AH-287-N15 | GORG | 0 | 0 | NA | 3 | 0 | orphan | 0 | 0 | NA | 0 | 0 | NA | 64.71 | 0 | 0.73 |
| AH-326-O20 | GORG | 0 | 0 | NA | 2 | 0 | orphan | 0 | 0 | NA | 0 | 0 | NA | 64.71 | 0 | 0.68 |
| AG-430-C07 | GORG | 1 | 0 | orphan | 3 | 0 | orphan | 0 | 0 | NA | 0 | 1 | orphan | 64.68 | 0 | 2.36 |

| | | | | | | | | | | | | | | | | |
|---|---|---|---|---|---|---|---|---|---|---|---|---|---|---|---|---|
| AG-447-I14 | GORG | 0 | 0 | NA | 2 | 0 | orphan | 0 | 0 | NA | 0 | 0 | NA | 64.67 | 0 | 1.21 |
| AG-893-E04 | GORG | 0 | 0 | NA | 1 | 0 | orphan | 0 | 0 | NA | 0 | 0 | NA | 64.66 | 0 | 0.95 |
| AG-899-L07 | GORG | 0 | 0 | NA | 1 | 1 | orphan | 0 | 0 | NA | 0 | 0 | NA | 64.66 | 0 | 0.81 |
| AG-325-P21 | GORG | 0 | 0 | NA | 2 | 0 | orphan | 0 | 0 | NA | 0 | 0 | NA | 64.56 | 0 | 0.75 |
| AG-453-D22 | GORG | 0 | 0 | NA | 1 | 0 | orphan | 0 | 0 | NA | 0 | 0 | NA | 64.46 | 0 | 0.65 |
| AG-897-E16 | GORG | 0 | 0 | NA | 2 | 0 | orphan | 0 | 0 | NA | 0 | 0 | NA | 64.46 | 0 | 0.98 |
| AG-911-P08 | GORG | 0 | 0 | NA | 1 | 0 | orphan | 0 | 0 | NA | 0 | 0 | NA | 64.4 | 0 | 0.65 |
| AG-892-O08 | GORG | 0 | 0 | NA | 1 | 0 | orphan | 0 | 0 | NA | 0 | 0 | NA | 64.32 | 0 | 0.87 |
| AG-903-C13 | GORG | 0 | 0 | NA | 2 | 0 | orphan | 0 | 0 | NA | 0 | 0 | NA | 64.32 | 0 | 0.73 |
| AG-892-J14 | GORG | 0 | 0 | NA | 1 | 0 | orphan | 0 | 0 | NA | 0 | 1 | orphan | 64.31 | 0 | 1.06 |
| AG-915-G04 | GORG | 0 | 0 | NA | 2 | 1 | complete | 0 | 0 | NA | 0 | 1 | orphan | 64.31 | 0 | 1.14 |
| AH-326-K11 | GORG | 0 | 0 | NA | 1 | 0 | orphan | 0 | 0 | NA | 0 | 0 | NA | 64.31 | 0 | 0.68 |
| AG-464-G09 | GORG | 0 | 0 | NA | 1 | 0 | orphan | 0 | 0 | NA | 0 | 0 | NA | 64.18 | 0 | 0.85 |
| AG-414-G02 | GORG | 0 | 0 | NA | 1 | 0 | orphan | 0 | 0 | NA | 0 | 0 | NA | 64.15 | 0 | 1.1 |
| AG-422-A09 | GORG | 0 | 0 | NA | 1 | 0 | orphan | 0 | 0 | NA | 0 | 0 | NA | 64.15 | 0 | 0.64 |
| AG-892-A10 | GORG | 0 | 0 | NA | 1 | 0 | orphan | 0 | 0 | NA | 0 | 0 | NA | 64.15 | 0 | 1.35 |
| AG-898-B19 | GORG | 0 | 0 | NA | 2 | 0 | orphan | 0 | 0 | NA | 0 | 0 | NA | 64.15 | 0 | 0.8 |
| AG-901-G05 | GORG | 0 | 0 | NA | 1 | 0 | orphan | 0 | 0 | NA | 0 | 0 | NA | 64.15 | 0 | 0.91 |
| AG-904-O07 | GORG | 0 | 0 | NA | 1 | 0 | orphan | 0 | 0 | NA | 0 | 0 | NA | 64.15 | 0 | 0.76 |
| AG-910-E23 | GORG | 0 | 0 | NA | 1 | 0 | orphan | 0 | 0 | NA | 0 | 0 | NA | 64.15 | 0 | 0.96 |
| AG-912-B10 | GORG | 0 | 0 | NA | 3 | 0 | orphan | 0 | 0 | NA | 0 | 0 | NA | 64.15 | 0 | 0.9 |
| AG-912-F15 | GORG | 0 | 0 | NA | 1 | 0 | orphan | 0 | 0 | NA | 0 | 0 | NA | 64.15 | 0 | 0.9 |
| AG-912-M23 | GORG | 0 | 0 | NA | 1 | 0 | orphan | 0 | 0 | NA | 0 | 0 | NA | 64.15 | 0 | 0.74 |
| AH-324-G05 | GORG | 0 | 0 | NA | 2 | 0 | orphan | 0 | 0 | NA | 0 | 0 | NA | 64.15 | 0 | 0.98 |
| AH-326-O21 | GORG | 0 | 0 | NA | 4 | 0 | orphan | 0 | 0 | NA | 0 | 0 | NA | 64.15 | 0 | 0.76 |
| AG-905-N19 | GORG | 1 | 1 | complete | 0 | 0 | NA | 0 | 0 | NA | 0 | 0 | NA | 64.13 | 0 | 1.36 |

| | | | | | | | | | | | | | | | | |
|---|---|---|---|---|---|---|---|---|---|---|---|---|---|---|---|---|
| AG-422-M23 | GORG | 1 | 1 | orphan | 0 | 0 | NA | 0 | 0 | NA | 0 | 0 | NA | 64.09 | 0 | 1.16 |
| AG-915-I21 | GORG | 0 | 0 | NA | 2 | 0 | orphan | 0 | 0 | NA | 0 | 0 | NA | 64.07 | 0 | 1.06 |
| AG-365-I20 | GORG | 0 | 0 | NA | 1 | 0 | orphan | 0 | 0 | NA | 0 | 0 | NA | 64.05 | 0 | 1.04 |
| AG-426-A07 | GORG | 2 | 0 | orphan | 0 | 0 | NA | 0 | 0 | NA | 0 | 0 | NA | 64.05 | 0 | 1.01 |
| AG-457-P15 | GORG | 0 | 0 | NA | 2 | 0 | orphan | 0 | 0 | NA | 0 | 0 | NA | 64.02 | 0 | 0.78 |
| AG-891-I14 | GORG | 0 | 0 | NA | 1 | 0 | orphan | 0 | 0 | NA | 0 | 0 | NA | 64 | 0 | 1.17 |
| AG-892-A08 | GORG | 0 | 0 | NA | 1 | 0 | orphan | 0 | 0 | NA | 0 | 0 | NA | 64 | 0 | 0.99 |
| AG-414-M02 | GORG | 0 | 0 | NA | 2 | 1 | complete | 0 | 0 | NA | 0 | 0 | NA | 63.96 | 0 | 0.93 |
| AG-915-G10 | GORG | 0 | 0 | NA | 2 | 0 | orphan | 0 | 0 | NA | 0 | 0 | NA | 63.93 | 0 | 1 |
| AG-457-K17 | GORG | 1 | 2 | complete | 1 | 0 | orphan | 0 | 0 | NA | 0 | 1 | orphan | 63.91 | 0 | 1.22 |
| AG-430-F09 | GORG | 0 | 0 | NA | 1 | 0 | orphan | 0 | 0 | NA | 0 | 0 | NA | 63.9 | 0 | 1.65 |
| AG-337-I16 | GORG | 0 | 0 | NA | 1 | 0 | orphan | 0 | 0 | NA | 0 | 0 | NA | 63.88 | 0 | 1.24 |
| AG-894-A10 | GORG | 0 | 0 | NA | 1 | 0 | orphan | 0 | 0 | NA | 0 | 0 | NA | 63.87 | 0 | 1 |
| AG-337-C22 | GORG | 0 | 0 | NA | 2 | 0 | orphan | 0 | 0 | NA | 0 | 0 | NA | 63.86 | 0 | 0.9 |
| AG-359-M02 | GORG | 0 | 0 | NA | 1 | 0 | orphan | 0 | 0 | NA | 0 | 0 | NA | 63.86 | 0 | 1.02 |
| AG-426-O23 | GORG | 0 | 0 | NA | 2 | 1 | complete | 0 | 0 | NA | 0 | 0 | NA | 63.84 | 0 | 0.92 |
| AG-470-G09 | GORG | 0 | 0 | NA | 1 | 0 | orphan | 0 | 0 | NA | 0 | 0 | NA | 63.79 | 0 | 1 |
| AG-470-L07 | GORG | 0 | 0 | NA | 1 | 0 | orphan | 0 | 0 | NA | 0 | 0 | NA | 63.77 | 0 | 0.99 |
| AG-422-O14 | GORG | 0 | 0 | NA | 2 | 0 | orphan | 0 | 0 | NA | 0 | 0 | NA | 63.75 | 0 | 0.83 |
| AG-319-E22 | GORG | 1 | 1 | complete | 0 | 0 | NA | 0 | 0 | NA | 0 | 0 | NA | 63.71 | 0 | 0.95 |
| AG-891-C06 | GORG | 0 | 0 | NA | 2 | 0 | orphan | 0 | 0 | NA | 0 | 1 | orphan | 63.7 | 0 | 2.41 |
| AG-899-O08 | GORG | 1 | 1 | complete | 1 | 0 | orphan | 0 | 0 | NA | 0 | 0 | NA | 63.69 | 0 | 1.06 |
| AG-365-N15 | GORG | 0 | 0 | NA | 2 | 0 | orphan | 0 | 0 | NA | 0 | 0 | NA | 63.68 | 0 | 0.93 |
| AG-447-E16 | GORG | 0 | 0 | NA | 2 | 0 | orphan | 0 | 0 | NA | 0 | 0 | NA | 63.67 | 0 | 1.11 |
| AG-909-O10 | GORG | 0 | 0 | NA | 2 | 1 | complete | 0 | 0 | NA | 0 | 0 | NA | 63.66 | 0 | 0.79 |

| | | | | | | | | | | | | | | | | |
|---|---|---|---|---|---|---|---|---|---|---|---|---|---|---|---|---|
| AG-404-D10 | GORG | 0 | 0 | NA | 1 | 0 | orphan | 1 | 1 | complete | 0 | 0 | NA | 63.61 | 0 | 1.17 |
| AG-447-P08 | GORG | 0 | 0 | NA | 1 | 0 | orphan | 0 | 0 | NA | 0 | 0 | NA | 63.6 | 0 | 1.45 |
| AG-918-D14 | GORG | 0 | 0 | NA | 1 | 0 | orphan | 0 | 0 | NA | 0 | 0 | NA | 63.6 | 0 | 0.86 |
| AG-439-E07 | GORG | 0 | 0 | NA | 2 | 0 | orphan | 0 | 0 | NA | 0 | 0 | NA | 63.52 | 0 | 0.83 |
| AG-430-F14 | GORG | 1 | 1 | complete | 0 | 0 | NA | 0 | 0 | NA | 0 | 0 | NA | 63.48 | 0 | 0.73 |
| AG-457-A01 | GORG | 0 | 0 | NA | 1 | 1 | complete | 0 | 0 | NA | 0 | 0 | NA | 63.41 | 0 | 1.18 |
| AG-337-P10 | GORG | 0 | 0 | NA | 2 | 0 | orphan | 0 | 0 | NA | 0 | 0 | NA | 63.4 | 0 | 1.03 |
| AG-359-J14 | GORG | 0 | 0 | NA | 2 | 0 | orphan | 0 | 0 | NA | 0 | 0 | NA | 63.37 | 0 | 1.32 |
| AG-920-G14 | GORG | 0 | 0 | NA | 1 | 0 | orphan | 0 | 0 | NA | 0 | 0 | NA | 63.36 | 0 | 0.78 |
| AG-895-I15 | GORG | 1 | 1 | complete | 0 | 0 | NA | 0 | 0 | NA | 0 | 1 | orphan | 63.34 | 0 | 1.09 |
| AG-319-P10 | GORG | 0 | 0 | NA | 1 | 0 | orphan | 0 | 0 | NA | 0 | 0 | NA | 63.29 | 0 | 1.14 |
| AG-447-N14 | GORG | 0 | 0 | NA | 1 | 0 | orphan | 0 | 0 | NA | 0 | 0 | NA | 63.25 | 0 | 0.98 |
| AG-914-L14 | GORG | 0 | 0 | NA | 1 | 0 | orphan | 0 | 0 | NA | 0 | 0 | NA | 63.25 | 0 | 1.01 |
| AG-914-P19 | GORG | 0 | 0 | NA | 2 | 0 | orphan | 0 | 1 | NA | 0 | 0 | NA | 63.25 | 0 | 0.87 |
| AG-915-L11 | GORG | 0 | 0 | NA | 2 | 0 | orphan | 0 | 0 | NA | 0 | 0 | NA | 63.25 | 0 | 0.83 |
| AG-464-P04 | GORG | 0 | 0 | NA | 1 | 0 | orphan | 0 | 0 | NA | 0 | 0 | NA | 63.23 | 0 | 0.89 |
| AG-457-P16 | GORG | 0 | 0 | NA | 2 | 0 | orphan | 0 | 0 | NA | 0 | 0 | NA | 63.21 | 0 | 0.97 |
| AG-911-J20 | GORG | 0 | 0 | NA | 2 | 0 | orphan | 0 | 0 | NA | 0 | 0 | NA | 63.21 | 0 | 0.89 |
| AG-913-A16 | GORG | 0 | 0 | NA | 1 | 0 | orphan | 0 | 0 | NA | 0 | 0 | NA | 63.21 | 0 | 1.03 |
| AG-916-D05 | GORG | 0 | 0 | NA | 1 | 0 | orphan | 0 | 0 | NA | 0 | 0 | NA | 63.21 | 0 | 0.75 |
| AG-899-E17 | GORG | 0 | 0 | NA | 1 | 0 | orphan | 0 | 0 | NA | 0 | 0 | NA | 63.2 | 0 | 0.7 |
| AG-453-A03 | GORG | 0 | 0 | NA | 1 | 0 | orphan | 0 | 0 | NA | 0 | 0 | NA | 63.19 | 0 | 0.68 |
| AG-422-E14 | GORG | 0 | 0 | NA | 1 | 0 | orphan | 0 | 0 | NA | 0 | 0 | NA | 63.17 | 0 | 0.76 |
| AG-911-P06 | GORG | 0 | 0 | NA | 1 | 0 | orphan | 0 | 0 | NA | 0 | 0 | NA | 63.17 | 0 | 0.83 |
| AG-426-B21 | GORG | 1 | 1 | complete | 1 | 0 | orphan | 0 | 0 | NA | 0 | 0 | NA | 63.16 | 0 | 1.37 |

| | | | | | | | | | | | | | | | | |
|---|---|---|---|---|---|---|---|---|---|---|---|---|---|---|---|---|
| AH-326-D16 | GORG | 0 | 0 | NA | 1 | 0 | orphan | 0 | 0 | NA | 0 | 0 | NA | 63.14 | 0 | 1.17 |
| AG-453-C15 | GORG | 0 | 0 | NA | 2 | 0 | orphan | 0 | 0 | NA | 0 | 0 | NA | 63.09 | 0 | 0.73 |
| AG-913-A09 | GORG | 0 | 0 | NA | 1 | 0 | orphan | 0 | 0 | NA | 0 | 0 | NA | 63.09 | 0 | 0.83 |
| AG-359-G20 | GORG | 0 | 0 | NA | 2 | 0 | orphan | 0 | 0 | NA | 0 | 0 | NA | 63.08 | 0 | 0.98 |
| AG-422-G19 | GORG | 0 | 0 | NA | 1 | 0 | orphan | 0 | 0 | NA | 0 | 0 | NA | 63.03 | 0 | 0.6 |
| AG-892-J18 | GORG | 0 | 0 | NA | 0 | 0 | NA | 0 | 0 | NA | 0 | 1 | orphan | 63 | 0 | 1.08 |
| AG-430-A09 | GORG | 0 | 0 | NA | 1 | 0 | orphan | 0 | 0 | NA | 0 | 0 | NA | 62.95 | 0 | 0.96 |
| AG-896-I08 | GORG | 0 | 0 | NA | 4 | 0 | orphan | 0 | 0 | NA | 0 | 0 | NA | 62.95 | 0 | 0.89 |
| AG-410-O05 | GORG | 0 | 0 | NA | 1 | 0 | orphan | 0 | 0 | NA | 0 | 0 | NA | 62.94 | 0 | 0.79 |
| AG-390-L13 | GORG | 0 | 0 | NA | 0 | 0 | NA | 0 | 0 | NA | 0 | 1 | orphan | 62.93 | 0 | 1.01 |
| AG-892-D19 | GORG | 0 | 0 | NA | 2 | 0 | orphan | 0 | 0 | NA | 0 | 0 | NA | 62.93 | 0 | 2.18 |
| AG-453-N20 | GORG | 0 | 0 | NA | 2 | 0 | orphan | 0 | 0 | NA | 0 | 0 | NA | 62.92 | 0 | 1.86 |
| AG-447-M06 | GORG | 0 | 0 | NA | 1 | 0 | orphan | 0 | 0 | NA | 0 | 0 | NA | 62.91 | 0 | 0.7 |
| AG-894-L14 | GORG | 0 | 0 | NA | 0 | 0 | NA | 0 | 0 | NA | 0 | 1 | orphan | 62.86 | 0 | 1 |
| AG-457-I17 | GORG | 0 | 0 | NA | 1 | 0 | orphan | 0 | 0 | NA | 0 | 0 | NA | 62.86 | 0 | 0.81 |
| AG-911-J03 | GORG | 0 | 0 | NA | 2 | 0 | orphan | 0 | 0 | NA | 0 | 0 | NA | 62.85 | 0 | 0.67 |
| AG-913-F10 | GORG | 0 | 0 | NA | 1 | 0 | orphan | 0 | 0 | NA | 0 | 0 | NA | 62.82 | 0 | 0.86 |
| AG-447-P14 | GORG | 0 | 0 | NA | 1 | 0 | orphan | 0 | 0 | NA | 0 | 0 | NA | 62.81 | 0 | 1.18 |
| AG-898-L21 | GORG | 0 | 0 | NA | 1 | 0 | orphan | 0 | 1 | NA | 0 | 0 | NA | 62.72 | 0 | 0.98 |
| AG-901-C21 | GORG | 1 | 0 | orphan | 1 | 0 | orphan | 0 | 0 | NA | 0 | 0 | NA | 62.69 | 0 | 0.86 |
| AG-913-O08 | GORG | 0 | 0 | NA | 1 | 0 | orphan | 0 | 0 | NA | 0 | 0 | NA | 62.69 | 0 | 0.81 |
| AG-905-C06 | GORG | 0 | 0 | NA | 1 | 0 | orphan | 0 | 0 | NA | 0 | 0 | NA | 62.66 | 0 | 0.8 |
| AG-435-G10 | GORG | 0 | 0 | NA | 2 | 0 | orphan | 0 | 0 | NA | 0 | 0 | NA | 62.65 | 0 | 0.83 |
| AG-435-M22 | GORG | 0 | 0 | NA | 1 | 0 | orphan | 0 | 0 | NA | 0 | 0 | NA | 62.65 | 0 | 0.72 |
| AG-900-O17 | GORG | 0 | 0 | NA | 1 | 0 | orphan | 0 | 0 | NA | 0 | 0 | NA | 62.65 | 0 | 1.01 |
| AG-915-K23 | GORG | 0 | 0 | NA | 1 | 0 | orphan | 0 | 0 | NA | 0 | 0 | NA | 62.65 | 0 | 1.01 |

| | | | | | | | | | | | | | | | | |
|---|---|---|---|---|---|---|---|---|---|---|---|---|---|---|---|---|
| AG-920-C13 | GORG | 0 | 0 | NA | 1 | 0 | orphan | 0 | 0 | NA | 0 | 0 | NA | 62.65 | 0 | 0.77 |
| AG-894-N18 | GORG | 0 | 0 | NA | 1 | 0 | orphan | 0 | 0 | NA | 0 | 0 | NA | 62.63 | 0 | 1.04 |
| AG-337-N17 | GORG | 0 | 0 | NA | 1 | 0 | orphan | 0 | 0 | NA | 0 | 0 | NA | 62.58 | 0 | 0.78 |
| AG-414-G21 | GORG | 0 | 0 | NA | 1 | 0 | orphan | 0 | 0 | NA | 0 | 0 | NA | 62.54 | 0 | 0.84 |
| AG-909-I06 | GORG | 0 | 0 | NA | 2 | 0 | orphan | 0 | 1 | NA | 0 | 0 | NA | 62.54 | 0 | 0.72 |
| AG-905-K17 | GORG | 0 | 0 | NA | 1 | 0 | orphan | 0 | 0 | NA | 0 | 0 | NA | 62.5 | 0 | 1.04 |
| AG-337-O13 | GORG | 0 | 0 | NA | 1 | 0 | orphan | 0 | 0 | NA | 0 | 0 | NA | 62.49 | 0 | 0.98 |
| AG-899-K23 | GORG | 0 | 0 | NA | 2 | 1 | complete | 0 | 0 | NA | 0 | 0 | NA | 62.44 | 0 | 0.79 |
| AH-324-N20 | GORG | 0 | 0 | NA | 1 | 0 | orphan | 0 | 0 | NA | 0 | 0 | NA | 62.44 | 0 | 0.97 |
| AH-326-F07 | GORG | 0 | 0 | NA | 1 | 0 | orphan | 0 | 0 | NA | 0 | 0 | NA | 62.4 | 0 | 0.72 |
| AG-457-M04 | GORG | 0 | 0 | NA | 1 | 0 | orphan | 0 | 0 | NA | 0 | 0 | NA | 62.37 | 0 | 1.15 |
| AG-898-J20 | GORG | 0 | 0 | NA | 0 | 0 | NA | 0 | 0 | NA | 0 | 1 | orphan | 62.36 | 0 | 1.11 |
| AG-365-L13 | GORG | 1 | 1 | complete | 0 | 0 | NA | 0 | 0 | NA | 0 | 0 | NA | 62.33 | 0 | 1.05 |
| AG-908-C20 | GORG | 1 | 1 | complete | 0 | 0 | NA | 0 | 0 | NA | 0 | 0 | NA | 62.31 | 0 | 1.06 |
| AG-919-N06 | GORG | 0 | 0 | NA | 2 | 1 | complete | 0 | 0 | NA | 0 | 0 | NA | 62.31 | 0 | 0.99 |
| AG-313-M21 | GORG | 0 | 0 | NA | 1 | 0 | orphan | 0 | 0 | NA | 0 | 0 | NA | 62.26 | 0 | 0.91 |
| AG-325-K07 | GORG | 0 | 0 | NA | 2 | 0 | orphan | 0 | 0 | NA | 0 | 0 | NA | 62.26 | 0 | 0.88 |
| AG-390-L21 | GORG | 0 | 0 | NA | 1 | 0 | orphan | 0 | 0 | NA | 0 | 0 | NA | 62.26 | 0 | 0.86 |
| AG-414-L21 | GORG | 0 | 0 | NA | 3 | 0 | orphan | 0 | 0 | NA | 0 | 0 | NA | 62.26 | 0 | 0.56 |
| AG-895-P10 | GORG | 0 | 0 | NA | 3 | 0 | orphan | 0 | 0 | NA | 0 | 0 | NA | 62.26 | 0 | 1.06 |
| AG-898-B05 | GORG | 0 | 0 | NA | 1 | 0 | orphan | 0 | 0 | NA | 0 | 0 | NA | 62.26 | 0 | 0.75 |
| AG-910-K19 | GORG | 0 | 0 | NA | 1 | 0 | orphan | 0 | 0 | NA | 0 | 0 | NA | 62.26 | 0 | 0.69 |
| AG-911-K02 | GORG | 0 | 0 | NA | 1 | 0 | orphan | 0 | 0 | NA | 0 | 0 | NA | 62.26 | 0 | 0.8 |
| AH-324-N05 | GORG | 0 | 0 | NA | 2 | 0 | orphan | 0 | 1 | NA | 0 | 0 | NA | 62.26 | 0 | 0.91 |
| AG-439-L09 | GORG | 0 | 0 | NA | 1 | 0 | orphan | 0 | 0 | NA | 0 | 1 | orphan | 62.25 | 0 | 0.7 |

| | | | | | | | | | | | | | | | | |
|---|---|---|---|---|---|---|---|---|---|---|---|---|---|---|---|---|
| AG-891-B08 | GORG | 1 | 1 | complete | 0 | 0 | NA | 0 | 0 | NA | 0 | 0 | NA | 62.23 | 0 | 1.06 |
| AG-313-B04 | GORG | 0 | 0 | NA | 1 | 1 | complete | 0 | 0 | NA | 0 | 0 | NA | 62.22 | 0 | 1.68 |
| AG-325-G13 | GORG | 0 | 0 | NA | 1 | 0 | orphan | 0 | 0 | NA | 0 | 0 | NA | 62.16 | 0 | 0.96 |
| AG-907-A01 | GORG | 0 | 0 | NA | 1 | 0 | orphan | 0 | 0 | NA | 0 | 0 | NA | 62.16 | 0 | 0.69 |
| AG-313-I15 | GORG | 0 | 0 | NA | 3 | 0 | orphan | 0 | 0 | NA | 0 | 0 | NA | 62.15 | 0 | 0.91 |
| AG-904-E07 | GORG | 1 | 1 | complete | 0 | 0 | NA | 0 | 0 | NA | 0 | 0 | NA | 62.15 | 0 | 0.85 |
| AG-917-L19 | GORG | 0 | 0 | NA | 1 | 0 | orphan | 0 | 0 | NA | 0 | 0 | NA | 62.15 | 0 | 0.75 |
| AH-287-O10 | GORG | 0 | 0 | NA | 2 | 0 | orphan | 0 | 0 | NA | 0 | 1 | orphan | 62.14 | 0 | 1.06 |
| AG-343-K16 | GORG | 0 | 0 | NA | 3 | 0 | orphan | 0 | 0 | NA | 0 | 0 | NA | 62.14 | 0 | 1.94 |
| AG-912-J14 | GORG | 0 | 0 | NA | 0 | 1 | NA | 0 | 0 | NA | 0 | 0 | NA | 62.14 | 0 | 1.04 |
| AG-464-B13 | GORG | 0 | 0 | NA | 2 | 0 | orphan | 0 | 0 | NA | 0 | 0 | NA | 62.13 | 0 | 0.84 |
| AG-908-E19 | GORG | 1 | 1 | complete | 0 | 0 | NA | 0 | 0 | NA | 0 | 0 | NA | 62.12 | 0 | 1.1 |
| AG-461-J11 | GORG | 0 | 0 | NA | 2 | 0 | orphan | 0 | 0 | NA | 0 | 0 | NA | 62.11 | 0 | 0.82 |
| AG-904-N10 | GORG | 0 | 0 | NA | 1 | 0 | orphan | 0 | 0 | NA | 0 | 0 | NA | 62.09 | 0 | 0.84 |
| AG-457-P06 | GORG | 1 | 1 | complete | 2 | 0 | orphan | 0 | 0 | NA | 0 | 0 | NA | 62.07 | 0 | 0.97 |
| AG-470-D15 | GORG | 0 | 0 | NA | 4 | 0 | orphan | 0 | 0 | NA | 0 | 0 | NA | 62.07 | 0 | 0.91 |
| AG-349-C03 | GORG | 0 | 0 | NA | 1 | 0 | orphan | 0 | 0 | NA | 0 | 0 | NA | 62.05 | 0 | 1.11 |
| AG-422-A01 | GORG | 0 | 0 | NA | 3 | 0 | orphan | 0 | 0 | NA | 0 | 0 | NA | 62.05 | 0 | 0.72 |
| AG-430-I04 | GORG | 0 | 0 | NA | 2 | 0 | orphan | 0 | 0 | NA | 0 | 0 | NA | 62.05 | 0 | 1.13 |
| AG-453-E17 | GORG | 0 | 0 | NA | 2 | 0 | orphan | 0 | 0 | NA | 0 | 0 | NA | 62.05 | 0 | 0.82 |
| AG-895-F19 | GORG | 0 | 0 | NA | 1 | 0 | orphan | 0 | 0 | NA | 0 | 0 | NA | 62.05 | 0 | 0.94 |
| AG-898-F04 | GORG | 0 | 0 | NA | 1 | 0 | orphan | 0 | 0 | NA | 0 | 0 | NA | 62.05 | 0 | 0.88 |
| AG-908-K05 | GORG | 0 | 0 | NA | 1 | 0 | orphan | 0 | 0 | NA | 0 | 0 | NA | 62.05 | 0 | 0.96 |
| AG-911-M22 | GORG | 0 | 0 | NA | 1 | 0 | orphan | 0 | 0 | NA | 0 | 0 | NA | 62.05 | 0 | 0.65 |
| AG-439-P04 | GORG | 0 | 0 | NA | 1 | 0 | orphan | 0 | 0 | NA | 0 | 0 | NA | 61.99 | 0 | 0.61 |

| | | | | | | | | | | | | | | | | |
|---|---|---|---|---|---|---|---|---|---|---|---|---|---|---|---|---|
| AG-895-P22 | GORG | 0 | 0 | NA | 2 | 0 | orphan | 0 | 0 | NA | 0 | 0 | NA | 61.94 | 0 | 1.06 |
| AG-439-M09 | GORG | 0 | 0 | NA | 1 | 0 | orphan | 0 | 0 | NA | 0 | 0 | NA | 61.91 | 0 | 0.65 |
| AG-325-D02 | GORG | 0 | 0 | NA | 1 | 0 | orphan | 0 | 0 | NA | 0 | 0 | NA | 61.9 | 0 | 0.75 |
| AG-917-K13 | GORG | 0 | 0 | NA | 0 | 0 | NA | 0 | 0 | NA | 0 | 2 | orphan | 61.89 | 0 | 1.26 |
| AG-319-K03 | GORG | 1 | 1 | complete | 1 | 0 | orphan | 0 | 0 | NA | 0 | 0 | NA | 61.88 | 0 | 1.24 |
| AG-909-D04 | GORG | 0 | 0 | NA | 0 | 0 | NA | 0 | 0 | NA | 0 | 1 | orphan | 61.87 | 0 | 0.97 |
| AG-464-J13 | GORG | 0 | 0 | NA | 1 | 0 | orphan | 0 | 0 | NA | 0 | 0 | NA | 61.87 | 0 | 0.8 |
| AG-325-B19 | GORG | 1 | 1 | complete | 2 | 0 | orphan | 0 | 0 | NA | 0 | 0 | NA | 61.83 | 0 | 1 |
| AG-457-B20 | GORG | 0 | 0 | NA | 1 | 0 | orphan | 0 | 0 | NA | 0 | 0 | NA | 61.83 | 0 | 0.74 |
| AG-919-O21 | GORG | 0 | 0 | NA | 1 | 0 | orphan | 0 | 0 | NA | 0 | 0 | NA | 61.83 | 0 | 1.59 |
| AG-892-I21 | GORG | 0 | 0 | NA | 3 | 0 | orphan | 0 | 0 | NA | 0 | 2 | orphan | 61.82 | 0 | 2.03 |
| AG-919-A15 | GORG | 1 | 1 | complete | 0 | 0 | NA | 0 | 0 | NA | 0 | 1 | orphan | 61.82 | 0 | 1.1 |
| AG-904-E19 | GORG | 0 | 0 | NA | 2 | 0 | orphan | 0 | 0 | NA | 0 | 0 | NA | 61.81 | 0 | 0.95 |
| AG-325-F16 | GORG | 0 | 0 | NA | 1 | 0 | orphan | 0 | 0 | NA | 0 | 0 | NA | 61.72 | 0 | 0.83 |
| AG-892-B08 | GORG | 1 | 0 | orphan | 0 | 0 | NA | 0 | 0 | NA | 0 | 0 | NA | 61.67 | 0 | 0.93 |
| AG-422-L16 | GORG | 0 | 0 | NA | 2 | 0 | orphan | 0 | 0 | NA | 0 | 0 | NA | 61.66 | 0 | 0.72 |
| AG-894-E17 | GORG | 0 | 0 | NA | 0 | 0 | NA | 0 | 0 | NA | 0 | 1 | orphan | 61.63 | 0 | 1.01 |
| AG-461-A07 | GORG | 0 | 0 | NA | 1 | 0 | orphan | 0 | 0 | NA | 0 | 0 | NA | 61.63 | 0 | 0.82 |
| AG-349-N22 | GORG | 0 | 0 | NA | 0 | 0 | NA | 0 | 0 | NA | 0 | 1 | orphan | 61.55 | 0 | 1 |
| AG-918-K05 | GORG | 0 | 0 | NA | 1 | 1 | orphan | 0 | 0 | NA | 0 | 0 | NA | 61.55 | 0 | 1.07 |
| AG-461-G18 | GORG | 0 | 0 | NA | 2 | 0 | orphan | 0 | 0 | NA | 0 | 0 | NA | 61.54 | 0 | 0.96 |
| AG-453-I07 | GORG | 0 | 0 | NA | 2 | 1 | complete | 0 | 0 | NA | 0 | 0 | NA | 61.52 | 0 | 1.19 |
| AG-892-D07 | GORG | 0 | 0 | NA | 1 | 1 | orphan | 0 | 0 | NA | 0 | 0 | NA | 61.52 | 0 | 1.09 |
| AG-896-P19 | GORG | 0 | 0 | NA | 1 | 0 | orphan | 0 | 0 | NA | 0 | 0 | NA | 61.52 | 0 | 0.93 |

| | | | | | | | | | | | | | | | | |
|---|---|---|---|---|---|---|---|---|---|---|---|---|---|---|---|---|
| AG-390-P05 | GORG | 0 | 0 | NA | 0 | 1 | NA | 0 | 0 | NA | 0 | 0 | NA | 61.49 | 0 | 1.09 |
| AG-903-G15 | GORG | 0 | 0 | NA | 1 | 0 | orphan | 0 | 0 | NA | 0 | 0 | NA | 61.49 | 0 | 0.66 |
| AG-349-P02 | GORG | 0 | 0 | NA | 0 | 0 | NA | 0 | 0 | NA | 0 | 1 | orphan | 61.47 | 0 | 1.02 |
| AG-918-C11 | GORG | 1 | 0 | orphan | 1 | 0 | orphan | 0 | 0 | NA | 0 | 0 | NA | 61.46 | 0 | 3.27 |
| AG-333-K22 | GORG | 0 | 0 | NA | 1 | 0 | orphan | 0 | 0 | NA | 0 | 0 | NA | 61.45 | 0 | 1.1 |
| AG-464-J02 | GORG | 0 | 0 | NA | 1 | 0 | orphan | 0 | 0 | NA | 0 | 0 | NA | 61.45 | 0 | 0.95 |
| AG-894-B05 | GORG | 0 | 0 | NA | 1 | 0 | orphan | 0 | 0 | NA | 0 | 0 | NA | 61.45 | 0 | 0.68 |
| AG-894-M06 | GORG | 0 | 0 | NA | 1 | 0 | orphan | 0 | 0 | NA | 0 | 0 | NA | 61.45 | 0 | 1.21 |
| AG-898-I20 | GORG | 1 | 0 | orphan | 1 | 0 | orphan | 0 | 0 | NA | 0 | 0 | NA | 61.45 | 0 | 0.88 |
| AG-905-P16 | GORG | 0 | 0 | NA | 1 | 0 | orphan | 0 | 0 | NA | 0 | 0 | NA | 61.45 | 0 | 0.82 |
| AG-907-J17 | GORG | 0 | 0 | NA | 1 | 0 | orphan | 0 | 0 | NA | 0 | 0 | NA | 61.45 | 0 | 0.95 |
| AG-911-N13 | GORG | 0 | 0 | NA | 1 | 0 | orphan | 0 | 0 | NA | 0 | 0 | NA | 61.45 | 0 | 0.62 |
| AG-912-L16 | GORG | 0 | 0 | NA | 1 | 0 | orphan | 0 | 0 | NA | 0 | 0 | NA | 61.45 | 0 | 1.03 |
| AG-914-M21 | GORG | 0 | 0 | NA | 1 | 0 | orphan | 0 | 0 | NA | 0 | 0 | NA | 61.45 | 0 | 0.87 |
| AH-321-J05 | GORG | 0 | 0 | NA | 2 | 0 | orphan | 0 | 0 | NA | 0 | 0 | NA | 61.45 | 0 | 1.19 |
| AH-324-P23 | GORG | 0 | 0 | NA | 1 | 0 | orphan | 0 | 0 | NA | 0 | 0 | NA | 61.45 | 0 | 0.88 |
| AG-891-I22 | GORG | 0 | 0 | NA | 1 | 0 | orphan | 0 | 0 | NA | 0 | 0 | NA | 61.4 | 0 | 0.83 |
| AG-910-K09 | GORG | 0 | 0 | NA | 2 | 2 | complete | 0 | 0 | NA | 0 | 1 | orphan | 61.38 | 0 | 1.56 |
| AG-917-B10 | GORG | 0 | 0 | NA | 1 | 0 | orphan | 0 | 0 | NA | 0 | 0 | NA | 61.37 | 0 | 1.15 |
| AG-918-L19 | GORG | 0 | 0 | NA | 2 | 0 | orphan | 0 | 0 | NA | 0 | 0 | NA | 61.32 | 0 | 0.92 |
| AG-435-I23 | GORG | 0 | 0 | NA | 1 | 0 | orphan | 0 | 0 | NA | 0 | 0 | NA | 61.28 | 0 | 0.83 |
| AG-910-B07 | GORG | 0 | 0 | NA | 1 | 0 | orphan | 0 | 0 | NA | 0 | 0 | NA | 61.28 | 0 | 0.75 |
| AG-333-E23 | GORG | 0 | 0 | NA | 1 | 0 | orphan | 0 | 0 | NA | 0 | 0 | NA | 61.24 | 0 | 0.73 |
| AG-343-D23 | GORG | 0 | 0 | NA | 1 | 0 | orphan | 0 | 0 | NA | 0 | 0 | NA | 61.24 | 0 | 0.62 |
| AG-461-J20 | GORG | 0 | 0 | NA | 1 | 0 | orphan | 0 | 0 | NA | 0 | 0 | NA | 61.24 | 0 | 0.87 |
| AG-908-D21 | GORG | 0 | 0 | NA | 1 | 0 | orphan | 0 | 0 | NA | 0 | 0 | NA | 61.24 | 0 | 0.9 |

| | | | | | | | | | | | | | | | | |
|---|---|---|---|---|---|---|---|---|---|---|---|---|---|---|---|---|
| AG-913-M20 | GORG | 0 | 0 | NA | 2 | 0 | orphan | 0 | 0 | NA | 0 | 0 | NA | 61.23 | 0 | 0.76 |
| AH-326-L14 | GORG | 1 | 1 | complete | 0 | 0 | NA | 0 | 0 | NA | 0 | 0 | NA | 61.21 | 0 | 0.8 |
| AG-422-P15 | GORG | 0 | 0 | NA | 0 | 0 | NA | 0 | 0 | NA | 0 | 1 | orphan | 61.19 | 0 | 0.99 |
| AG-919-I06 | GORG | 1 | 1 | complete | 2 | 0 | orphan | 0 | 0 | NA | 0 | 0 | NA | 61.17 | 0 | 0.99 |
| AG-457-E16 | GORG | 0 | 0 | NA | 1 | 0 | orphan | 0 | 0 | NA | 0 | 0 | NA | 61.15 | 0 | 0.72 |
| AG-453-O04 | GORG | 1 | 1 | complete | 1 | 0 | orphan | 0 | 0 | NA | 0 | 0 | NA | 61.14 | 0 | 0.71 |
| AG-422-C03 | GORG | 0 | 0 | NA | 0 | 0 | NA | 0 | 0 | NA | 0 | 1 | orphan | 61.13 | 0 | 1.02 |
| AG-457-A14 | GORG | 0 | 0 | NA | 2 | 0 | orphan | 0 | 0 | NA | 0 | 0 | NA | 61.06 | 0 | 0.62 |
| AG-919-N14 | GORG | 0 | 0 | NA | 1 | 0 | orphan | 0 | 0 | NA | 0 | 0 | NA | 61.06 | 0 | 0.88 |
| AG-337-M07 | GORG | 0 | 0 | NA | 2 | 0 | orphan | 0 | 0 | NA | 0 | 0 | NA | 61.04 | 0 | 0.72 |
| AG-913-G06 | GORG | 0 | 0 | NA | 1 | 0 | orphan | 0 | 0 | NA | 0 | 0 | NA | 61.04 | 0 | 0.69 |
| AG-439-L07 | GORG | 0 | 0 | NA | 2 | 0 | orphan | 0 | 0 | NA | 0 | 0 | NA | 61.02 | 0 | 0.9 |
| AG-899-E20 | GORG | 1 | 1 | complete | 0 | 0 | NA | 0 | 0 | NA | 0 | 0 | NA | 61.02 | 0 | 1.07 |
| AG-918-C23 | GORG | 0 | 0 | NA | 1 | 0 | orphan | 0 | 0 | NA | 0 | 0 | NA | 61 | 0 | 1.3 |
| AG-919-F17 | GORG | 0 | 0 | NA | 1 | 0 | orphan | 0 | 0 | NA | 0 | 0 | NA | 61 | 0 | 0.86 |
| AG-915-I02 | GORG | 0 | 0 | NA | 1 | 0 | orphan | 0 | 0 | NA | 0 | 0 | NA | 60.98 | 0 | 0.54 |
| AG-898-J09 | GORG | 0 | 1 | NA | 1 | 0 | orphan | 0 | 0 | NA | 0 | 0 | NA | 60.97 | 0 | 1.01 |
| AG-365-G17 | GORG | 0 | 0 | NA | 1 | 0 | orphan | 0 | 0 | NA | 0 | 0 | NA | 60.95 | 0 | 0.8 |
| AG-457-B16 | GORG | 1 | 1 | complete | 0 | 0 | NA | 0 | 0 | NA | 0 | 0 | NA | 60.95 | 0 | 1.38 |
| AG-898-O15 | GORG | 0 | 0 | NA | 1 | 0 | orphan | 0 | 0 | NA | 0 | 0 | NA | 60.9 | 0 | 0.92 |
| AG-913-B19 | GORG | 1 | 1 | complete | 1 | 0 | orphan | 0 | 0 | NA | 0 | 0 | NA | 60.88 | 0 | 0.87 |
| AG-422-A04 | GORG | 0 | 0 | NA | 0 | 0 | NA | 0 | 0 | NA | 0 | 2 | orphan | 60.87 | 0 | 0.97 |
| AG-899-E07 | GORG | 0 | 0 | NA | 1 | 0 | orphan | 0 | 0 | NA | 0 | 0 | NA | 60.87 | 0 | 0.76 |
| AH-324-C03 | GORG | 0 | 0 | NA | 1 | 0 | orphan | 0 | 0 | NA | 0 | 0 | NA | 60.84 | 0 | 0.98 |

| | | | | | | | | | | | | | | | | |
|---|---|---|---|---|---|---|---|---|---|---|---|---|---|---|---|---|
| AG-900-M19 | GORG | 0 | 0 | NA | 1 | 0 | orphan | 0 | 0 | NA | 0 | 0 | NA | 60.82 | 0 | 0.89 |
| AG-905-A11 | GORG | 0 | 0 | NA | 1 | 0 | orphan | 0 | 0 | NA | 0 | 0 | NA | 60.81 | 0 | 0.87 |
| AG-319-I09 | GORG | 0 | 0 | NA | 1 | 0 | orphan | 0 | 0 | NA | 0 | 0 | NA | 60.78 | 0 | 2.29 |
| AG-895-L09 | GORG | 0 | 0 | NA | 2 | 0 | orphan | 0 | 0 | NA | 0 | 0 | NA | 60.73 | 0 | 1.71 |
| AG-903-A05 | GORG | 0 | 0 | NA | 1 | 0 | orphan | 0 | 0 | NA | 0 | 0 | NA | 60.71 | 0 | 0.81 |
| AG-910-C06 | GORG | 0 | 0 | NA | 4 | 0 | orphan | 0 | 0 | NA | 0 | 0 | NA | 60.69 | 0 | 0.85 |
| AG-907-O16 | GORG | 0 | 0 | NA | 1 | 0 | orphan | 0 | 0 | NA | 0 | 0 | NA | 60.64 | 0 | 1.02 |
| AG-426-K20 | GORG | 0 | 0 | NA | 1 | 1 | complete | 0 | 0 | NA | 0 | 0 | NA | 60.63 | 0 | 0.9 |
| AG-904-F15 | GORG | 0 | 0 | NA | 1 | 0 | orphan | 0 | 0 | NA | 0 | 0 | NA | 60.63 | 0 | 0.83 |
| AG-360-O21 | GORG | 0 | 0 | NA | 1 | 0 | orphan | 0 | 0 | NA | 0 | 0 | NA | 60.62 | 0 | 0.67 |
| AG-360-A21 | GORG | 0 | 0 | NA | 1 | 0 | orphan | 0 | 0 | NA | 0 | 0 | NA | 60.6 | 0 | 0.72 |
| AG-898-L16 | GORG | 0 | 0 | NA | 1 | 0 | orphan | 0 | 0 | NA | 0 | 0 | NA | 60.59 | 0 | 1 |
| AH-287-G21 | GORG | 0 | 0 | NA | 2 | 0 | orphan | 0 | 0 | NA | 0 | 0 | NA | 60.56 | 0 | 0.71 |
| AG-390-G09 | GORG | 0 | 0 | NA | 1 | 0 | orphan | 0 | 0 | NA | 0 | 0 | NA | 60.55 | 0 | 0.65 |
| AG-390-O16 | GORG | 0 | 0 | NA | 1 | 0 | orphan | 0 | 0 | NA | 0 | 0 | NA | 60.55 | 0 | 0.93 |
| AG-343-E19 | GORG | 0 | 0 | NA | 1 | 0 | orphan | 0 | 0 | NA | 0 | 0 | NA | 60.49 | 0 | 0.65 |
| AG-896-P11 | GORG | 1 | 1 | complete | 1 | 0 | orphan | 0 | 0 | NA | 0 | 0 | NA | 60.49 | 0 | 1.15 |
| AH-326-E11 | GORG | 0 | 0 | NA | 1 | 0 | orphan | 0 | 0 | NA | 0 | 0 | NA | 60.48 | 0 | 0.65 |
| AG-470-A10 | GORG | 0 | 0 | NA | 1 | 0 | orphan | 0 | 0 | NA | 0 | 0 | NA | 60.47 | 0 | 0.91 |
| AG-422-D02 | GORG | 0 | 0 | NA | 1 | 0 | orphan | 0 | 0 | NA | 0 | 0 | NA | 60.46 | 0 | 0.89 |
| AG-905-P17 | GORG | 0 | 0 | NA | 1 | 0 | orphan | 0 | 0 | NA | 0 | 0 | NA | 60.46 | 0 | 0.83 |
| AG-349-J23 | GORG | 0 | 0 | NA | 0 | 0 | NA | 0 | 0 | NA | 0 | 1 | orphan | 60.44 | 0 | 0.97 |
| AG-349-C08 | GORG | 0 | 0 | NA | 0 | 0 | NA | 0 | 0 | NA | 0 | 2 | orphan | 60.42 | 0 | 1.02 |
| AG-410-E14 | GORG | 0 | 0 | NA | 1 | 0 | orphan | 0 | 0 | NA | 0 | 0 | NA | 60.41 | 0 | 0.79 |
| AG-343-J07 | GORG | 0 | 0 | NA | 1 | 0 | orphan | 0 | 0 | NA | 0 | 1 | orphan | 60.4 | 0 | 0.97 |

| | | | | | | | | | | | | | | | | |
|---|---|---|---|---|---|---|---|---|---|---|---|---|---|---|---|---|
| AG-916-B03 | GORG | 0 | 0 | NA | 1 | 0 | orphan | 0 | 0 | NA | 0 | 0 | NA | 60.4 | 0 | 0.75 |
| AG-333-E18 | GORG | 0 | 0 | NA | 1 | 0 | orphan | 0 | 0 | NA | 0 | 0 | NA | 60.38 | 0 | 0.67 |
| AG-337-G09 | GORG | 1 | 1 | complete | 1 | 0 | orphan | 0 | 0 | NA | 0 | 0 | NA | 60.38 | 0 | 0.94 |
| AG-390-A03 | GORG | 0 | 0 | NA | 2 | 0 | orphan | 0 | 0 | NA | 0 | 0 | NA | 60.38 | 0 | 1.02 |
| AG-430-J17 | GORG | 0 | 0 | NA | 1 | 0 | orphan | 0 | 0 | NA | 0 | 0 | NA | 60.38 | 0 | 0.99 |
| AG-447-P18 | GORG | 0 | 0 | NA | 2 | 0 | orphan | 0 | 0 | NA | 0 | 0 | NA | 60.38 | 0 | 0.9 |
| AG-457-M07 | GORG | 0 | 0 | NA | 1 | 0 | orphan | 0 | 0 | NA | 0 | 0 | NA | 60.38 | 0 | 0.86 |
| AG-891-G20 | GORG | 0 | 0 | NA | 1 | 0 | orphan | 0 | 0 | NA | 0 | 0 | NA | 60.38 | 0 | 1.09 |
| AG-896-P03 | GORG | 1 | 0 | orphan | 1 | 0 | orphan | 0 | 0 | NA | 0 | 0 | NA | 60.38 | 0 | 0.99 |
| AG-898-K21 | GORG | 0 | 0 | NA | 1 | 0 | orphan | 0 | 0 | NA | 0 | 0 | NA | 60.38 | 0 | 0.98 |
| AG-904-F16 | GORG | 0 | 0 | NA | 1 | 0 | orphan | 0 | 0 | NA | 0 | 0 | NA | 60.38 | 0 | 0.7 |
| AG-913-G15 | GORG | 0 | 0 | NA | 1 | 0 | orphan | 0 | 0 | NA | 0 | 0 | NA | 60.38 | 0 | 0.73 |
| AG-914-D13 | GORG | 0 | 0 | NA | 1 | 0 | orphan | 0 | 0 | NA | 0 | 0 | NA | 60.38 | 0 | 0.92 |
| AG-914-D21 | GORG | 0 | 0 | NA | 1 | 0 | orphan | 0 | 0 | NA | 0 | 0 | NA | 60.38 | 0 | 0.73 |
| AG-915-L04 | GORG | 1 | 1 | complete | 1 | 0 | orphan | 0 | 0 | NA | 0 | 0 | NA | 60.38 | 0 | 0.86 |
| AG-915-M10 | GORG | 0 | 0 | NA | 2 | 0 | orphan | 0 | 0 | NA | 0 | 0 | NA | 60.38 | 0 | 0.87 |
| AG-919-M03 | GORG | 0 | 0 | NA | 1 | 0 | orphan | 0 | 0 | NA | 0 | 0 | NA | 60.38 | 0 | 0.99 |
| AH-321-G07 | GORG | 0 | 0 | NA | 0 | 0 | NA | 0 | 0 | NA | 0 | 1 | orphan | 60.37 | 0 | 1.01 |
| AG-457-N18 | GORG | 0 | 0 | NA | 1 | 0 | orphan | 0 | 0 | NA | 0 | 0 | NA | 60.35 | 0 | 0.8 |
| AG-895-P19 | GORG | 0 | 0 | NA | 0 | 0 | NA | 0 | 0 | NA | 0 | 1 | orphan | 60.34 | 0 | 1.15 |
| AG-349-L15 | GORG | 0 | 0 | NA | 2 | 0 | orphan | 0 | 0 | NA | 0 | 0 | NA | 60.34 | 0 | 0.87 |
| AG-897-M14 | GORG | 1 | 0 | orphan | 1 | 0 | orphan | 0 | 0 | NA | 0 | 0 | NA | 60.34 | 0 | 0.91 |
| AG-915-N18 | GORG | 0 | 0 | NA | 1 | 0 | orphan | 0 | 0 | NA | 0 | 0 | NA | 60.34 | 0 | 1.42 |
| AG-895-E17 | GORG | 0 | 0 | NA | 1 | 0 | orphan | 0 | 0 | NA | 0 | 0 | NA | 60.31 | 0 | 0.89 |
| AH-321-K11 | GORG | 0 | 0 | NA | 3 | 2 | complete | 0 | 0 | NA | 0 | 0 | NA | 60.3 | 0 | 1.14 |

| | | | | | | | | | | | | | | | | |
|---|---|---|---|---|---|---|---|---|---|---|---|---|---|---|---|---|
| AG-333-C21 | GORG | 0 | 0 | NA | 5 | 0 | orphan | 0 | 0 | NA | 0 | 0 | NA | 60.28 | 0 | 1.08 |
| AG-903-D15 | GORG | 0 | 0 | NA | 2 | 0 | orphan | 0 | 0 | NA | 0 | 0 | NA | 60.28 | 0 | 1.23 |
| AG-457-P23 | GORG | 0 | 0 | NA | 1 | 0 | orphan | 0 | 0 | NA | 0 | 0 | NA | 60.27 | 0 | 1.23 |
| AG-901-B22 | GORG | 0 | 0 | NA | 1 | 0 | orphan | 0 | 0 | NA | 0 | 0 | NA | 60.26 | 0 | 0.82 |
| AG-447-P21 | GORG | 0 | 0 | NA | 1 | 0 | orphan | 0 | 0 | NA | 0 | 0 | NA | 60.24 | 0 | 0.88 |
| AG-901-E14 | GORG | 0 | 0 | NA | 1 | 0 | orphan | 0 | 0 | NA | 0 | 0 | NA | 60.24 | 0 | 0.75 |
| AG-903-O20 | GORG | 0 | 0 | NA | 1 | 0 | orphan | 0 | 0 | NA | 0 | 0 | NA | 60.24 | 0 | 1.03 |
| AG-470-O07 | GORG | 2 | 0 | orphan | 1 | 0 | orphan | 0 | 0 | NA | 0 | 0 | NA | 60.22 | 0 | 1.36 |
| AG-343-M05 | GORG | 0 | 0 | NA | 0 | 0 | NA | 0 | 0 | NA | 0 | 2 | orpha n | 60.2 | 0 | 1.02 |
| AG-447-K14 | GORG | 0 | 0 | NA | 1 | 0 | orphan | 0 | 0 | NA | 0 | 0 | NA | 60.03 | 0 | 0.76 |
| AG-895-A01 | GORG | 0 | 0 | NA | 1 | 0 | orphan | 0 | 0 | NA | 0 | 0 | NA | 60.03 | 0 | 0.91 |
| AG-333-F17 | GORG | 0 | 0 | NA | 1 | 0 | orphan | 0 | 0 | NA | 0 | 0 | NA | 60.02 | 0 | 0.7 |
| AG-410-J18 | GORG | 0 | 0 | NA | 1 | 0 | orphan | 0 | 0 | NA | 0 | 0 | NA | 60.01 | 0 | 0.62 |
| AG-907-D22 | GORG | 0 | 0 | NA | 1 | 0 | orphan | 0 | 0 | NA | 0 | 0 | NA | 59.96 | 0 | 0.97 |
| AG-910-L06 | GORG | 0 | 0 | NA | 2 | 0 | orphan | 0 | 0 | NA | 0 | 0 | NA | 59.96 | 0 | 0.84 |
| AG-349-P05 | GORG | 0 | 0 | NA | 1 | 0 | orphan | 0 | 0 | NA | 0 | 0 | NA | 59.95 | 0 | 0.88 |
| AG-898-D03 | GORG | 0 | 0 | NA | 1 | 0 | orphan | 0 | 0 | NA | 0 | 0 | NA | 59.95 | 0 | 0.79 |
| AG-414-L13 | GORG | 0 | 0 | NA | 1 | 0 | orphan | 0 | 0 | NA | 0 | 0 | NA | 59.93 | 0 | 0.74 |
| AG-898-I19 | GORG | 0 | 0 | NA | 1 | 0 | orphan | 0 | 0 | NA | 0 | 0 | NA | 59.93 | 0 | 1.07 |
| AG-325-M17 | GORG | 0 | 0 | NA | 1 | 0 | orphan | 0 | 0 | NA | 0 | 0 | NA | 59.87 | 0 | 0.78 |
| AH-326-D05 | GORG | 0 | 0 | NA | 1 | 0 | orphan | 0 | 0 | NA | 0 | 0 | NA | 59.87 | 0 | 1.06 |
| AG-426-C15 | GORG | 0 | 0 | NA | 1 | 0 | orphan | 0 | 1 | NA | 0 | 0 | NA | 59.85 | 0 | 1.19 |
| AG-464-J03 | GORG | 0 | 0 | NA | 1 | 0 | orphan | 0 | 0 | NA | 0 | 0 | NA | 59.84 | 0 | 0.62 |
| AG-897-K15 | GORG | 0 | 0 | NA | 1 | 0 | orphan | 0 | 0 | NA | 0 | 0 | NA | 59.84 | 0 | 0.69 |
| AG-898-P19 | GORG | 0 | 0 | NA | 1 | 0 | orphan | 0 | 0 | NA | 0 | 0 | NA | 59.83 | 0 | 0.79 |
| AG-918-F18 | GORG | 0 | 0 | NA | 1 | 0 | orphan | 1 | 0 | orphan | 0 | 0 | NA | 59.82 | 0 | 0.87 |
| AG-898-N11 | GORG | 0 | 0 | NA | 1 | 0 | orphan | 0 | 0 | NA | 0 | 0 | NA | 59.8 | 0 | 0.82 |

| | | | | | | | | | | | | | | | | |
|---|---|---|---|---|---|---|---|---|---|---|---|---|---|---|---|---|
| AG-359-K02 | GORG | 1 | 0 | orphan | 0 | 0 | NA | 0 | 0 | NA | 0 | 0 | NA | 59.76 | 0 | 1.05 |
| AH-326-N16 | GORG | 0 | 0 | NA | 1 | 0 | orphan | 0 | 0 | NA | 0 | 0 | NA | 59.66 | 0 | 1.68 |
| AG-359-I05 | GORG | 1 | 1 | complete | 0 | 0 | NA | 0 | 0 | NA | 0 | 0 | NA | 59.65 | 0 | 0.96 |
| AG-892-B13 | GORG | 0 | 0 | NA | 1 | 0 | orphan | 0 | 0 | NA | 0 | 0 | NA | 59.65 | 0 | 0.99 |
| AG-898-P11 | GORG | 0 | 0 | NA | 1 | 0 | orphan | 0 | 0 | NA | 0 | 0 | NA | 59.65 | 0 | 1 |
| AG-349-F08 | GORG | 0 | 0 | NA | 1 | 0 | orphan | 0 | 0 | NA | 0 | 0 | NA | 59.64 | 0 | 0.76 |
| AG-457-I08 | GORG | 0 | 0 | NA | 2 | 0 | orphan | 0 | 0 | NA | 0 | 0 | NA | 59.64 | 0 | 1.09 |
| AG-907-A10 | GORG | 0 | 0 | NA | 1 | 0 | orphan | 0 | 0 | NA | 0 | 0 | NA | 59.64 | 0 | 0.9 |
| AG-422-P05 | GORG | 0 | 0 | NA | 2 | 0 | orphan | 0 | 0 | NA | 0 | 0 | NA | 59.62 | 0 | 0.94 |
| AG-919-I03 | GORG | 0 | 0 | NA | 1 | 0 | orphan | 0 | 0 | NA | 0 | 0 | NA | 59.6 | 0 | 0.75 |
| AG-895-A22 | GORG | 0 | 0 | NA | 1 | 0 | orphan | 0 | 0 | NA | 0 | 0 | NA | 59.59 | 0 | 1.86 |
| AG-365-G13 | GORG | 0 | 0 | NA | 1 | 0 | orphan | 0 | 0 | NA | 0 | 0 | NA | 59.55 | 0 | 0.84 |
| AH-324-J20 | GORG | 0 | 0 | NA | 1 | 0 | orphan | 0 | 0 | NA | 0 | 0 | NA | 59.51 | 0 | 0.94 |
| AG-439-F07 | GORG | 0 | 0 | NA | 2 | 0 | orphan | 0 | 0 | NA | 0 | 0 | NA | 59.48 | 0 | 1.47 |
| AG-893-J13 | GORG | 0 | 0 | NA | 1 | 0 | orphan | 0 | 0 | NA | 0 | 0 | NA | 59.48 | 0 | 1.72 |
| AG-319-G23 | GORG | 1 | 2 | complete | 0 | 0 | NA | 0 | 0 | NA | 0 | 0 | NA | 59.47 | 0 | 1.1 |
| AG-435-C13 | GORG | 0 | 0 | NA | 1 | 0 | orphan | 0 | 0 | NA | 0 | 0 | NA | 59.46 | 0 | 0.71 |
| AG-359-E07 | GORG | 0 | 0 | NA | 1 | 0 | orphan | 0 | 0 | NA | 0 | 0 | NA | 59.43 | 0 | 0.91 |
| AG-414-J08 | GORG | 0 | 0 | NA | 1 | 0 | orphan | 0 | 0 | NA | 0 | 0 | NA | 59.43 | 0 | 0.85 |
| AG-901-B02 | GORG | 0 | 0 | NA | 1 | 0 | orphan | 0 | 0 | NA | 0 | 0 | NA | 59.43 | 0 | 1.11 |
| AG-414-M20 | GORG | 0 | 0 | NA | 2 | 0 | orphan | 0 | 0 | NA | 0 | 0 | NA | 59.41 | 0 | 1.03 |
| AG-333-C20 | GORG | 0 | 0 | NA | 2 | 0 | orphan | 0 | 0 | NA | 0 | 0 | NA | 59.4 | 0 | 0.78 |
| AG-908-B04 | GORG | 0 | 0 | NA | 2 | 0 | orphan | 0 | 0 | NA | 0 | 0 | NA | 59.4 | 0 | 2.2 |
| AG-918-M22 | GORG | 0 | 0 | NA | 2 | 0 | orphan | 0 | 0 | NA | 0 | 0 | NA | 59.39 | 0 | 1.05 |
| AG-919-M15 | GORG | 0 | 1 | NA | 0 | 0 | NA | 0 | 0 | NA | 0 | 0 | NA | 59.36 | 0 | 1.05 |
| AG-461-G07 | GORG | 0 | 0 | NA | 1 | 0 | orphan | 0 | 0 | NA | 0 | 0 | NA | 59.32 | 0 | 0.76 |

| | | | | | | | | | | | | | | | | |
|---|---|---|---|---|---|---|---|---|---|---|---|---|---|---|---|---|
| AG-916-P10 | GORG | 0 | 0 | NA | 2 | 0 | orphan | 0 | 0 | NA | 0 | 0 | NA | 59.3 | 0 | 0.99 |
| AG-898-A07 | GORG | 0 | 0 | NA | 1 | 0 | orphan | 0 | 0 | NA | 0 | 0 | NA | 59.26 | 0 | 0.78 |
| AG-895-F06 | GORG | 0 | 0 | NA | 1 | 0 | orphan | 0 | 0 | NA | 0 | 0 | NA | 59.25 | 0 | 1.5 |
| AG-917-M14 | GORG | 1 | 2 | complete | 0 | 0 | NA | 0 | 0 | NA | 0 | 0 | NA | 59.24 | 0 | 0.87 |
| AG-439-L13 | GORG | 0 | 0 | NA | 1 | 0 | orphan | 0 | 0 | NA | 0 | 0 | NA | 59.22 | 0 | 0.77 |
| AG-404-A19 | GORG | 0 | 0 | NA | 0 | 1 | NA | 0 | 0 | NA | 0 | 0 | NA | 59.2 | 0 | 1.1 |
| AG-898-C03 | GORG | 0 | 0 | NA | 1 | 0 | orphan | 0 | 0 | NA | 0 | 0 | NA | 59.19 | 0 | 0.93 |
| AG-909-N03 | GORG | 0 | 1 | NA | 0 | 0 | NA | 0 | 0 | NA | 0 | 0 | NA | 59.19 | 0 | 0.95 |
| AG-337-K14 | GORG | 1 | 0 | orphan | 1 | 0 | orphan | 0 | 0 | NA | 0 | 0 | NA | 59.15 | 0 | 0.88 |
| AG-900-D02 | GORG | 0 | 0 | NA | 2 | 0 | orphan | 0 | 0 | NA | 0 | 0 | NA | 59.12 | 0 | 1.89 |
| AG-899-K05 | GORG | 0 | 0 | NA | 0 | 0 | NA | 0 | 0 | NA | 0 | 1 | orphan | 59.11 | 0 | 0.98 |
| AG-422-K15 | GORG | 0 | 0 | NA | 0 | 0 | NA | 0 | 0 | NA | 0 | 1 | orphan | 59.08 | 0 | 0.76 |
| AG-390-L02 | GORG | 0 | 0 | NA | 1 | 0 | orphan | 0 | 0 | NA | 0 | 0 | NA | 59.07 | 0 | 1.05 |
| AG-325-D07 | GORG | 0 | 0 | NA | 1 | 0 | orphan | 0 | 0 | NA | 0 | 0 | NA | 59.05 | 0 | 0.71 |
| AG-325-I05 | GORG | 1 | 1 | complete | 1 | 0 | orphan | 0 | 0 | NA | 0 | 0 | NA | 59.05 | 0 | 0.68 |
| AG-903-D03 | GORG | 0 | 0 | NA | 0 | 0 | NA | 0 | 0 | NA | 0 | 1 | orphan | 59.04 | 0 | 0.85 |
| AG-360-D04 | GORG | 0 | 0 | NA | 2 | 0 | orphan | 0 | 0 | NA | 0 | 0 | NA | 59.04 | 0 | 0.74 |
| AG-899-G20 | GORG | 0 | 0 | NA | 1 | 0 | orphan | 0 | 0 | NA | 0 | 0 | NA | 59.04 | 0 | 0.83 |
| AG-904-B19 | GORG | 0 | 0 | NA | 1 | 0 | orphan | 0 | 0 | NA | 0 | 0 | NA | 59.04 | 0 | 0.62 |
| AH-321-I13 | GORG | 0 | 0 | NA | 0 | 0 | NA | 0 | 0 | NA | 0 | 1 | orphan | 59.01 | 0 | 1.03 |
| AG-365-E14 | GORG | 0 | 0 | NA | 1 | 0 | orphan | 0 | 0 | NA | 0 | 0 | NA | 59.01 | 0 | 0.73 |
| AG-903-F04 | GORG | 0 | 0 | NA | 1 | 0 | orphan | 0 | 0 | NA | 0 | 0 | NA | 59 | 0 | 0.68 |
| AG-911-P03 | GORG | 1 | 0 | orphan | 0 | 0 | NA | 0 | 0 | NA | 0 | 0 | NA | 58.83 | 0 | 0.73 |
| AG-919-B19 | GORG | 0 | 0 | NA | 1 | 0 | orphan | 0 | 0 | NA | 0 | 0 | NA | 58.79 | 0 | 0.78 |
| AG-390-P19 | GORG | 0 | 0 | NA | 1 | 0 | orphan | 0 | 0 | NA | 0 | 0 | NA | 58.77 | 0 | 0.92 |

| | | | | | | | | | | | | | | | | |
|---|---|---|---|---|---|---|---|---|---|---|---|---|---|---|---|---|
| AG-430-O03 | GORG | 1 | 1 | complete | 3 | 0 | orphan | 0 | 0 | NA | 0 | 0 | NA | 58.76 | 0 | 1.14 |
| AG-457-O11 | GORG | 0 | 0 | NA | 1 | 0 | orphan | 0 | 0 | NA | 0 | 0 | NA | 58.76 | 0 | 0.79 |
| AG-914-I22 | GORG | 0 | 0 | NA | 1 | 0 | orphan | 0 | 0 | NA | 0 | 0 | NA | 58.75 | 0 | 1.08 |
| AG-359-C21 | GORG | 0 | 0 | NA | 2 | 0 | orphan | 0 | 0 | NA | 0 | 0 | NA | 58.73 | 0 | 0.59 |
| AG-447-A01 | GORG | 1 | 1 | complete | 0 | 0 | NA | 0 | 0 | NA | 0 | 0 | NA | 58.72 | 0 | 1.06 |
| AG-917-N07 | GORG | 0 | 0 | NA | 1 | 1 | orphan | 0 | 0 | NA | 0 | 0 | NA | 58.67 | 0 | 0.69 |
| AG-422-G23 | GORG | 0 | 0 | NA | 1 | 0 | orphan | 0 | 0 | NA | 0 | 0 | NA | 58.66 | 0 | 0.84 |
| AG-426-F14 | GORG | 0 | 0 | NA | 1 | 0 | orphan | 0 | 0 | NA | 0 | 0 | NA | 58.66 | 0 | 0.73 |
| AG-920-K13 | GORG | 0 | 0 | NA | 2 | 0 | orphan | 0 | 0 | NA | 0 | 0 | NA | 58.63 | 0 | 0.77 |
| AG-900-F03 | GORG | 0 | 0 | NA | 1 | 0 | orphan | 0 | 1 | NA | 0 | 2 | orphan | 58.62 | 0 | 1.1 |
| AG-919-N16 | GORG | 1 | 1 | complete | 1 | 0 | orphan | 0 | 0 | NA | 0 | 1 | orphan | 58.62 | 0 | 1.54 |
| AG-319-G16 | GORG | 0 | 0 | NA | 1 | 0 | orphan | 0 | 0 | NA | 0 | 0 | NA | 58.6 | 0 | 1.06 |
| AG-426-M23 | GORG | 0 | 0 | NA | 1 | 1 | complete | 0 | 0 | NA | 0 | 0 | NA | 58.6 | 0 | 1.09 |
| AG-918-L22 | GORG | 2 | 2 | complete | 0 | 0 | NA | 0 | 0 | NA | 0 | 0 | NA | 58.6 | 0 | 1.96 |
| AG-916-N10 | GORG | 0 | 0 | NA | 2 | 0 | orphan | 0 | 0 | NA | 0 | 0 | NA | 58.56 | 0 | 0.69 |
| AG-404-P11 | GORG | 0 | 0 | NA | 1 | 0 | orphan | 0 | 0 | NA | 0 | 1 | orphan | 58.54 | 0 | 0.97 |
| AG-365-A04 | GORG | 0 | 0 | NA | 1 | 0 | orphan | 0 | 0 | NA | 0 | 0 | NA | 58.49 | 0 | 0.88 |
| AG-905-C04 | GORG | 0 | 0 | NA | 1 | 0 | orphan | 0 | 0 | NA | 0 | 0 | NA | 58.49 | 0 | 0.75 |
| AG-909-E22 | GORG | 0 | 0 | NA | 2 | 0 | orphan | 0 | 0 | NA | 0 | 0 | NA | 58.49 | 0 | 0.59 |
| AG-909-O15 | GORG | 0 | 0 | NA | 1 | 0 | orphan | 0 | 0 | NA | 0 | 0 | NA | 58.49 | 0 | 1 |
| AG-913-C02 | GORG | 0 | 0 | NA | 1 | 0 | orphan | 0 | 0 | NA | 0 | 0 | NA | 58.49 | 0 | 1.1 |
| AG-914-I03 | GORG | 0 | 0 | NA | 1 | 0 | orphan | 0 | 0 | NA | 0 | 0 | NA | 58.49 | 0 | 0.97 |
| AG-915-F17 | GORG | 0 | 0 | NA | 1 | 0 | orphan | 0 | 0 | NA | 0 | 0 | NA | 58.49 | 0 | 1.06 |
| AH-287-K14 | GORG | 1 | 0 | orphan | 1 | 0 | orphan | 0 | 0 | NA | 0 | 0 | NA | 58.49 | 0 | 0.82 |
| AH-321-E06 | GORG | 0 | 0 | NA | 1 | 0 | orphan | 0 | 0 | NA | 0 | 0 | NA | 58.49 | 0 | 0.82 |

| | | | | | | | | | | | | | | | | |
|---|---|---|---|---|---|---|---|---|---|---|---|---|---|---|---|---|
| AG-337-J06 | GORG | 0 | 0 | NA | 1 | 0 | orphan | 0 | 0 | NA | 0 | 0 | NA | 58.48 | 0 | 0.71 |
| AG-333-B23 | GORG | 1 | 1 | orphan | 1 | 0 | orphan | 0 | 0 | NA | 0 | 0 | NA | 58.47 | 0 | 0.9 |
| AG-903-I13 | GORG | 0 | 0 | NA | 1 | 0 | orphan | 0 | 1 | NA | 0 | 0 | NA | 58.44 | 0 | 0.75 |
| AG-343-N17 | GORG | 0 | 0 | NA | 3 | 0 | orphan | 0 | 0 | NA | 0 | 0 | NA | 58.43 | 0 | 0.93 |
| AG-365-O06 | GORG | 0 | 0 | NA | 1 | 0 | orphan | 0 | 0 | NA | 0 | 0 | NA | 58.43 | 0 | 1.15 |
| AG-439-J05 | GORG | 0 | 0 | NA | 1 | 0 | orphan | 0 | 0 | NA | 0 | 0 | NA | 58.42 | 0 | 0.7 |
| AG-349-N20 | GORG | 0 | 1 | NA | 0 | 0 | NA | 0 | 0 | NA | 0 | 1 | orphan | 58.41 | 0 | 1.04 |
| AG-390-E15 | GORG | 0 | 0 | NA | 2 | 0 | orphan | 0 | 0 | NA | 0 | 0 | NA | 58.41 | 0 | 1.14 |
| AG-319-E18 | GORG | 1 | 2 | complete | 0 | 0 | NA | 0 | 0 | NA | 0 | 0 | NA | 58.4 | 0 | 1.12 |
| AG-333-J13 | GORG | 0 | 0 | NA | 1 | 0 | orphan | 0 | 0 | NA | 0 | 0 | NA | 58.4 | 0 | 1 |
| AH-287-P14 | GORG | 0 | 0 | NA | 2 | 0 | orphan | 0 | 0 | NA | 0 | 0 | NA | 58.4 | 0 | 0.69 |
| AG-915-J21 | GORG | 0 | 0 | NA | 1 | 0 | orphan | 0 | 0 | NA | 0 | 0 | NA | 58.39 | 0 | 0.69 |
| AG-390-F05 | GORG | 0 | 0 | NA | 3 | 0 | orphan | 0 | 0 | NA | 0 | 0 | NA | 58.38 | 0 | 0.67 |
| AG-453-O21 | GORG | 0 | 0 | NA | 2 | 0 | orphan | 0 | 0 | NA | 0 | 0 | NA | 58.37 | 0 | 0.83 |
| AG-891-C05 | GORG | 0 | 0 | NA | 0 | 0 | NA | 0 | 0 | NA | 0 | 1 | orphan | 58.33 | 0 | 0.87 |
| AG-337-E22 | GORG | 0 | 1 | NA | 1 | 0 | orphan | 0 | 0 | NA | 0 | 0 | NA | 58.33 | 0 | 0.68 |
| AG-916-J14 | GORG | 0 | 0 | NA | 1 | 0 | orphan | 0 | 0 | NA | 0 | 0 | NA | 58.32 | 0 | 1.05 |
| AG-325-A13 | GORG | 0 | 0 | NA | 1 | 0 | orphan | 0 | 0 | NA | 0 | 0 | NA | 58.29 | 0 | 0.83 |
| AH-321-G14 | GORG | 0 | 0 | NA | 1 | 0 | orphan | 0 | 0 | NA | 0 | 0 | NA | 58.29 | 0 | 0.87 |
| AG-313-G08 | GORG | 0 | 0 | NA | 1 | 0 | orphan | 0 | 0 | NA | 0 | 0 | NA | 58.28 | 0 | 0.98 |
| AG-464-O02 | GORG | 0 | 0 | NA | 1 | 0 | orphan | 0 | 0 | NA | 0 | 0 | NA | 58.28 | 0 | 0.64 |
| AG-343-J20 | GORG | 0 | 0 | NA | 1 | 0 | orphan | 0 | 0 | NA | 0 | 0 | NA | 58.27 | 0 | 0.83 |
| AG-435-P09 | GORG | 1 | 1 | complete | 0 | 0 | NA | 0 | 0 | NA | 0 | 0 | NA | 58.26 | 0 | 0.98 |
| AG-907-C13 | GORG | 0 | 0 | NA | 0 | 0 | NA | 0 | 0 | NA | 0 | 1 | orphan | 58.24 | 0 | 0.88 |
| AG-907-G04 | GORG | 0 | 0 | NA | 3 | 0 | orphan | 0 | 0 | NA | 0 | 0 | NA | 58.2 | 0 | 0.72 |

| | | | | | | | | | | | | | | | | |
|---|---|---|---|---|---|---|---|---|---|---|---|---|---|---|---|---|
| AG-899-L08 | GORG | 0 | 0 | NA | 2 | 0 | orphan | 0 | 0 | NA | 0 | 0 | NA | 58.19 | 0 | 1.45 |
| AG-913-O16 | GORG | 0 | 0 | NA | 2 | 0 | orphan | 0 | 0 | NA | 0 | 0 | NA | 58.12 | 0 | 0.71 |
| AG-447-L02 | GORG | 0 | 0 | NA | 2 | 0 | orphan | 0 | 0 | NA | 0 | 0 | NA | 58.11 | 0 | 1 |
| AG-313-N13 | GORG | 0 | 0 | NA | 1 | 0 | orphan | 0 | 0 | NA | 0 | 0 | NA | 58.08 | 0 | 1.12 |
| AG-349-P08 | GORG | 0 | 0 | NA | 3 | 1 | orphan | 0 | 0 | NA | 0 | 0 | NA | 58.06 | 0 | 0.81 |
| AG-461-G23 | GORG | 0 | 0 | NA | 4 | 0 | orphan | 0 | 0 | NA | 0 | 0 | NA | 57.98 | 0 | 1.45 |
| AG-895-G04 | GORG | 0 | 0 | NA | 1 | 0 | orphan | 0 | 0 | NA | 0 | 0 | NA | 57.95 | 0 | 0.72 |
| AG-333-O07 | GORG | 0 | 0 | NA | 1 | 0 | orphan | 0 | 0 | NA | 0 | 0 | NA | 57.92 | 0 | 0.88 |
| AG-913-N04 | GORG | 0 | 0 | NA | 2 | 0 | orphan | 0 | 0 | NA | 0 | 0 | NA | 57.87 | 0 | 0.98 |
| AG-457-O16 | GORG | 0 | 0 | NA | 1 | 0 | orphan | 0 | 0 | NA | 0 | 0 | NA | 57.83 | 0 | 0.8 |
| AG-891-F19 | GORG | 1 | 1 | complete | 1 | 0 | orphan | 0 | 0 | NA | 0 | 0 | NA | 57.83 | 0 | 0.93 |
| AG-898-J15 | GORG | 0 | 0 | NA | 1 | 0 | orphan | 0 | 0 | NA | 0 | 0 | NA | 57.83 | 0 | 0.87 |
| AG-913-M03 | GORG | 0 | 0 | NA | 1 | 0 | orphan | 0 | 0 | NA | 0 | 0 | NA | 57.77 | 0 | 1.27 |
| AG-919-K21 | GORG | 0 | 0 | NA | 1 | 0 | orphan | 0 | 0 | NA | 0 | 1 | orphan | 57.76 | 0 | 0.79 |
| AG-435-E14 | GORG | 0 | 1 | NA | 1 | 0 | orphan | 0 | 0 | NA | 0 | 0 | NA | 57.72 | 0 | 0.75 |
| AG-919-N18 | GORG | 0 | 0 | NA | 1 | 0 | orphan | 0 | 0 | NA | 0 | 0 | NA | 57.71 | 0 | 0.97 |
| AH-324-F05 | GORG | 0 | 0 | NA | 1 | 0 | orphan | 0 | 0 | NA | 0 | 0 | NA | 57.71 | 0 | 1.05 |
| AG-461-M06 | GORG | 0 | 1 | NA | 1 | 0 | orphan | 0 | 0 | NA | 0 | 0 | NA | 57.66 | 0 | 0.74 |
| AG-359-N23 | GORG | 0 | 0 | NA | 1 | 0 | orphan | 0 | 0 | NA | 0 | 0 | NA | 57.63 | 0 | 2.02 |
| AG-447-B03 | GORG | 0 | 0 | NA | 1 | 0 | orphan | 0 | 0 | NA | 0 | 0 | NA | 57.63 | 0 | 0.83 |
| AG-453-O03 | GORG | 0 | 0 | NA | 1 | 0 | orphan | 0 | 0 | NA | 0 | 0 | NA | 57.63 | 0 | 0.82 |
| AG-325-B15 | GORG | 0 | 0 | NA | 0 | 0 | NA | 0 | 0 | NA | 0 | 1 | orphan | 57.6 | 0 | 0.94 |
| AH-287-A11 | GORG | 0 | 0 | NA | 1 | 0 | orphan | 0 | 0 | NA | 0 | 0 | NA | 57.59 | 0 | 1.98 |
| AG-447-C11 | GORG | 0 | 0 | NA | 1 | 0 | orphan | 0 | 0 | NA | 0 | 0 | NA | 57.58 | 0 | 0.58 |
| AG-891-E22 | GORG | 0 | 0 | NA | 2 | 0 | orphan | 0 | 0 | NA | 0 | 0 | NA | 57.55 | 0 | 0.96 |
| AG-904-P08 | GORG | 0 | 0 | NA | 1 | 0 | orphan | 0 | 0 | NA | 0 | 0 | NA | 57.55 | 0 | 0.76 |

| | | | | | | | | | | | | | | | | |
|---|---|---|---|---|---|---|---|---|---|---|---|---|---|---|---|---|
| AG-916-P19 | GORG | 0 | 0 | NA | 1 | 0 | orphan | 0 | 0 | NA | 0 | 2 | orphan | 57.54 | 0 | 0.99 |
| AG-898-B14 | GORG | 0 | 0 | NA | 1 | 0 | orphan | 0 | 0 | NA | 0 | 1 | orphan | 57.52 | 0 | 1.71 |
| AG-893-L09 | GORG | 0 | 0 | NA | 1 | 0 | orphan | 0 | 0 | NA | 0 | 1 | orphan | 57.5 | 0 | 0.65 |
| AG-422-M10 | GORG | 0 | 0 | NA | 1 | 0 | orphan | 0 | 0 | NA | 0 | 0 | NA | 57.49 | 0 | 1.08 |
| AG-359-D02 | GORG | 0 | 0 | NA | 3 | 0 | orphan | 0 | 0 | NA | 0 | 0 | NA | 57.48 | 0 | 0.73 |
| AG-414-I19 | GORG | 0 | 0 | NA | 1 | 0 | orphan | 0 | 0 | NA | 0 | 0 | NA | 57.48 | 0 | 0.86 |
| AG-359-N02 | GORG | 0 | 0 | NA | 1 | 0 | orphan | 0 | 0 | NA | 0 | 0 | NA | 57.41 | 0 | 0.92 |
| AG-464-J21 | GORG | 0 | 0 | NA | 1 | 0 | orphan | 0 | 0 | NA | 0 | 0 | NA | 57.41 | 0 | 0.6 |
| AG-422-A17 | GORG | 0 | 0 | NA | 1 | 0 | orphan | 0 | 0 | NA | 0 | 0 | NA | 57.39 | 0 | 1 |
| AG-337-A22 | GORG | 0 | 0 | NA | 3 | 0 | orphan | 0 | 0 | NA | 0 | 0 | NA | 57.36 | 0 | 0.82 |
| AG-914-P10 | GORG | 0 | 0 | NA | 1 | 0 | orphan | 0 | 0 | NA | 0 | 0 | NA | 57.36 | 0 | 0.82 |
| AG-349-I21 | GORG | 0 | 0 | NA | 1 | 0 | orphan | 0 | 0 | NA | 0 | 0 | NA | 57.34 | 0 | 0.84 |
| AG-915-I06 | GORG | 0 | 0 | NA | 1 | 0 | orphan | 0 | 0 | NA | 0 | 0 | NA | 57.34 | 0 | 1.72 |
| AG-319-A01 | GORG | 0 | 0 | NA | 2 | 0 | orphan | 0 | 0 | NA | 0 | 0 | NA | 57.28 | 0 | 0.76 |
| AG-461-F16 | GORG | 0 | 0 | NA | 1 | 0 | orphan | 0 | 0 | NA | 0 | 0 | NA | 57.28 | 0 | 0.87 |
| AG-337-P20 | GORG | 0 | 0 | NA | 1 | 0 | orphan | 0 | 0 | NA | 0 | 0 | NA | 57.23 | 0 | 0.91 |
| AG-390-F16 | GORG | 0 | 0 | NA | 1 | 0 | orphan | 0 | 0 | NA | 0 | 0 | NA | 57.23 | 0 | 0.76 |
| AG-414-G10 | GORG | 0 | 0 | NA | 1 | 0 | orphan | 0 | 0 | NA | 0 | 0 | NA | 57.23 | 0 | 0.76 |
| AG-439-I17 | GORG | 0 | 0 | NA | 3 | 0 | orphan | 0 | 0 | NA | 0 | 0 | NA | 57.23 | 0 | 0.67 |
| AG-903-N10 | GORG | 0 | 0 | NA | 2 | 0 | orphan | 0 | 0 | NA | 0 | 0 | NA | 57.23 | 0 | 0.97 |
| AG-325-O07 | GORG | 0 | 0 | NA | 1 | 0 | orphan | 0 | 0 | NA | 0 | 0 | NA | 57.22 | 0 | 0.96 |
| AG-313-K16 | GORG | 0 | 0 | NA | 1 | 0 | orphan | 0 | 0 | NA | 0 | 0 | NA | 57.21 | 0 | 2.2 |
| AG-337-I03 | GORG | 0 | 1 | NA | 0 | 0 | NA | 0 | 0 | NA | 0 | 0 | NA | 57.21 | 0 | 1.08 |
| AG-447-F14 | GORG | 0 | 0 | NA | 2 | 0 | orphan | 0 | 0 | NA | 0 | 0 | NA | 57.21 | 0 | 1.05 |
| AG-919-N20 | GORG | 0 | 0 | NA | 2 | 0 | orphan | 0 | 0 | NA | 0 | 0 | NA | 57.17 | 0 | 0.95 |
| AG-404-C08 | GORG | 0 | 0 | NA | 1 | 0 | orphan | 0 | 0 | NA | 0 | 0 | NA | 57.15 | 0 | 1.46 |

| | | | | | | | | | | | | | | | | |
|---|---|---|---|---|---|---|---|---|---|---|---|---|---|---|---|---|
| AG-910-O11 | GORG | 0 | 1 | NA | 1 | 0 | orphan | 0 | 0 | NA | 0 | 0 | NA | 57.12 | 0 | 0.72 |
| AG-359-L04 | GORG | 0 | 0 | NA | 2 | 0 | orphan | 0 | 0 | NA | 0 | 0 | NA | 57.08 | 0 | 1.26 |
| AG-897-D05 | GORG | 1 | 1 | complete | 1 | 0 | orphan | 0 | 0 | NA | 0 | 0 | NA | 57.05 | 0 | 1.37 |
| AG-435-P21 | GORG | 0 | 0 | NA | 1 | 0 | orphan | 0 | 0 | NA | 0 | 0 | NA | 57.02 | 0 | 0.52 |
| AH-324-B08 | GORG | 0 | 0 | NA | 3 | 0 | orphan | 0 | 0 | NA | 0 | 0 | NA | 57.02 | 0 | 0.86 |
| AG-447-L04 | GORG | 1 | 1 | complete | 1 | 0 | orphan | 0 | 0 | NA | 0 | 0 | NA | 56.98 | 0 | 0.9 |
| AG-422-N07 | GORG | 0 | 0 | NA | 1 | 0 | orphan | 0 | 0 | NA | 0 | 0 | NA | 56.96 | 0 | 0.82 |
| AG-913-C18 | GORG | 0 | 0 | NA | 1 | 0 | orphan | 0 | 0 | NA | 0 | 0 | NA | 56.95 | 0 | 0.77 |
| AG-899-M04 | GORG | 0 | 0 | NA | 0 | 0 | NA | 0 | 0 | NA | 0 | 1 | orphan | 56.9 | 0 | 1.06 |
| AG-359-A03 | GORG | 0 | 0 | NA | 1 | 0 | orphan | 0 | 0 | NA | 0 | 0 | NA | 56.9 | 0 | 2.68 |
| AG-435-C06 | GORG | 0 | 0 | NA | 1 | 0 | orphan | 0 | 0 | NA | 0 | 0 | NA | 56.9 | 0 | 0.96 |
| AG-899-P17 | GORG | 0 | 0 | NA | 3 | 0 | orphan | 0 | 0 | NA | 0 | 0 | NA | 56.9 | 0 | 0.95 |
| AG-900-G07 | GORG | 0 | 0 | NA | 2 | 0 | orphan | 0 | 0 | NA | 0 | 0 | NA | 56.9 | 0 | 1.27 |
| AG-912-E06 | GORG | 0 | 0 | NA | 2 | 0 | orphan | 0 | 0 | NA | 0 | 0 | NA | 56.9 | 0 | 1.61 |
| AG-913-K10 | GORG | 0 | 0 | NA | 4 | 0 | orphan | 0 | 0 | NA | 0 | 0 | NA | 56.9 | 0 | 0.83 |
| AH-287-M23 | GORG | 0 | 0 | NA | 2 | 0 | orphan | 0 | 0 | NA | 0 | 0 | NA | 56.9 | 0 | 1.52 |
| AG-912-E21 | GORG | 0 | 0 | NA | 1 | 0 | orphan | 0 | 0 | NA | 0 | 0 | NA | 56.87 | 0 | 0.68 |
| AG-917-D10 | GORG | 0 | 0 | NA | 1 | 0 | orphan | 0 | 0 | NA | 0 | 0 | NA | 56.87 | 0 | 0.77 |
| AG-912-C20 | GORG | 0 | 0 | NA | 2 | 1 | complete | 0 | 0 | NA | 0 | 1 | orphan | 56.83 | 0 | 0.98 |
| AH-326-P20 | GORG | 0 | 0 | NA | 1 | 0 | orphan | 0 | 0 | NA | 0 | 0 | NA | 56.83 | 0 | 0.54 |
| AG-325-C05 | GORG | 1 | 0 | orphan | 2 | 0 | orphan | 0 | 0 | NA | 0 | 0 | NA | 56.8 | 0 | 1.34 |
| AG-457-M17 | GORG | 0 | 0 | NA | 1 | 0 | orphan | 0 | 0 | NA | 0 | 0 | NA | 56.8 | 0 | 0.73 |
| AH-287-C03 | GORG | 0 | 0 | NA | 1 | 0 | orphan | 0 | 0 | NA | 0 | 0 | NA | 56.78 | 0 | 0.8 |
| AG-461-P20 | GORG | 0 | 0 | NA | 1 | 0 | orphan | 0 | 0 | NA | 0 | 0 | NA | 56.77 | 0 | 0.66 |
| AG-891-I10 | GORG | 2 | 1 | orphan | 0 | 0 | NA | 0 | 0 | NA | 0 | 0 | NA | 56.74 | 0 | 1.47 |
| AG-439-B07 | GORG | 0 | 0 | NA | 2 | 0 | orphan | 0 | 0 | NA | 0 | 0 | NA | 56.72 | 0 | 1.03 |

| | | | | | | | | | | | | | | | | |
|---|---|---|---|---|---|---|---|---|---|---|---|---|---|---|---|---|
| AG-900-C22 | GORG | 0 | 0 | NA | 1 | 0 | orphan | 0 | 0 | NA | 0 | 0 | NA | 56.71 | 0 | 0.9 |
| AG-901-E10 | GORG | 0 | 0 | NA | 1 | 0 | orphan | 0 | 0 | NA | 0 | 0 | NA | 56.64 | 0 | 0.64 |
| AG-337-D23 | GORG | 0 | 0 | NA | 1 | 0 | orphan | 0 | 0 | NA | 0 | 0 | NA | 56.63 | 0 | 0.67 |
| AG-430-J18 | GORG | 0 | 0 | NA | 1 | 0 | orphan | 0 | 0 | NA | 0 | 0 | NA | 56.63 | 0 | 0.96 |
| AG-430-K22 | GORG | 0 | 0 | NA | 1 | 0 | orphan | 0 | 0 | NA | 0 | 0 | NA | 56.63 | 0 | 0.79 |
| AG-900-K20 | GORG | 0 | 0 | NA | 1 | 0 | orphan | 0 | 0 | NA | 0 | 0 | NA | 56.63 | 0 | 0.78 |
| AG-910-A17 | GORG | 0 | 0 | NA | 1 | 0 | orphan | 0 | 0 | NA | 0 | 0 | NA | 56.63 | 0 | 0.93 |
| AG-912-D13 | GORG | 0 | 0 | NA | 1 | 0 | orphan | 0 | 0 | NA | 0 | 0 | NA | 56.63 | 0 | 0.94 |
| AG-917-L20 | GORG | 0 | 0 | NA | 1 | 0 | orphan | 0 | 0 | NA | 0 | 0 | NA | 56.63 | 0 | 0.56 |
| AG-430-C22 | GORG | 0 | 0 | NA | 1 | 0 | orphan | 0 | 0 | NA | 0 | 1 | orphan | 56.6 | 0 | 0.78 |
| AG-325-P17 | GORG | 0 | 0 | NA | 1 | 0 | orphan | 0 | 0 | NA | 0 | 0 | NA | 56.6 | 0 | 0.75 |
| AG-349-O21 | GORG | 0 | 0 | NA | 1 | 0 | orphan | 0 | 0 | NA | 0 | 0 | NA | 56.6 | 0 | 0.74 |
| AG-435-N03 | GORG | 0 | 0 | NA | 1 | 0 | orphan | 0 | 0 | NA | 0 | 0 | NA | 56.6 | 0 | 0.82 |
| AG-453-O19 | GORG | 0 | 0 | NA | 2 | 0 | orphan | 0 | 0 | NA | 0 | 0 | NA | 56.6 | 0 | 0.7 |
| AG-470-O18 | GORG | 0 | 0 | NA | 1 | 0 | orphan | 0 | 0 | NA | 0 | 0 | NA | 56.6 | 0 | 0.76 |
| AG-892-N17 | GORG | 0 | 0 | NA | 1 | 0 | orphan | 0 | 0 | NA | 0 | 0 | NA | 56.6 | 0 | 0.81 |
| AG-896-N02 | GORG | 0 | 0 | NA | 1 | 0 | orphan | 0 | 0 | NA | 0 | 0 | NA | 56.6 | 0 | 0.83 |
| AG-903-N09 | GORG | 0 | 0 | NA | 1 | 0 | orphan | 0 | 0 | NA | 0 | 0 | NA | 56.6 | 0 | 0.71 |
| AG-905-B07 | GORG | 0 | 0 | NA | 2 | 0 | orphan | 0 | 0 | NA | 0 | 0 | NA | 56.6 | 0 | 0.87 |
| AG-908-B21 | GORG | 0 | 0 | NA | 1 | 0 | orphan | 0 | 0 | NA | 0 | 0 | NA | 56.6 | 0 | 0.81 |
| AG-910-J06 | GORG | 0 | 0 | NA | 2 | 0 | orphan | 0 | 0 | NA | 0 | 0 | NA | 56.6 | 0 | 0.73 |
| AG-912-P17 | GORG | 1 | 1 | complete | 0 | 0 | NA | 0 | 0 | NA | 0 | 0 | NA | 56.6 | 0 | 0.7 |
| AG-913-E03 | GORG | 0 | 0 | NA | 1 | 0 | orphan | 0 | 0 | NA | 0 | 0 | NA | 56.6 | 0 | 0.86 |
| AG-914-J18 | GORG | 0 | 0 | NA | 1 | 0 | orphan | 0 | 0 | NA | 0 | 0 | NA | 56.6 | 0 | 0.86 |
| AG-918-I22 | GORG | 0 | 0 | NA | 2 | 0 | orphan | 0 | 0 | NA | 0 | 0 | NA | 56.6 | 0 | 0.65 |
| AH-321-G11 | GORG | 0 | 0 | NA | 1 | 0 | orphan | 0 | 0 | NA | 0 | 0 | NA | 56.6 | 0 | 0.7 |

| | | | | | | | | | | | | | | | | |
|---|---|---|---|---|---|---|---|---|---|---|---|---|---|---|---|---|
| AG-892-C06 | GORG | 0 | 0 | NA | 1 | 0 | orphan | 0 | 0 | NA | 0 | 0 | NA | 56.59 | 0 | 1.05 |
| AG-343-F06 | GORG | 0 | 0 | NA | 2 | 0 | orphan | 0 | 0 | NA | 0 | 0 | NA | 56.52 | 0 | 0.67 |
| AG-430-O13 | GORG | 0 | 0 | NA | 1 | 0 | orphan | 0 | 0 | NA | 0 | 0 | NA | 56.52 | 0 | 0.75 |
| AG-892-D03 | GORG | 1 | 1 | complete | 0 | 0 | NA | 0 | 0 | NA | 0 | 0 | NA | 56.52 | 0 | 0.83 |
| AG-894-O22 | GORG | 0 | 0 | NA | 2 | 0 | orphan | 0 | 0 | NA | 0 | 0 | NA | 56.51 | 0 | 0.96 |
| AG-430-O16 | GORG | 0 | 0 | NA | 1 | 0 | orphan | 0 | 0 | NA | 0 | 0 | NA | 56.49 | 0 | 1.27 |
| AG-919-K06 | GORG | 0 | 0 | NA | 1 | 0 | orphan | 0 | 0 | NA | 0 | 0 | NA | 56.48 | 0 | 0.81 |
| AG-919-K15 | GORG | 0 | 2 | NA | 0 | 0 | NA | 0 | 0 | NA | 0 | 0 | NA | 56.48 | 0 | 1 |
| AG-461-M08 | GORG | 0 | 0 | NA | 1 | 0 | orphan | 0 | 0 | NA | 0 | 0 | NA | 56.41 | 0 | 0.82 |
| AG-892-I04 | GORG | 0 | 0 | NA | 1 | 1 | orphan | 0 | 0 | NA | 0 | 0 | NA | 56.39 | 0 | 1.01 |
| AG-325-B04 | GORG | 0 | 0 | NA | 1 | 0 | orphan | 0 | 0 | NA | 0 | 0 | NA | 56.35 | 0 | 1.08 |
| AG-910-F17 | GORG | 0 | 0 | NA | 1 | 0 | orphan | 0 | 0 | NA | 0 | 0 | NA | 56.35 | 0 | 0.89 |
| AG-911-J17 | GORG | 0 | 0 | NA | 2 | 0 | orphan | 0 | 0 | NA | 0 | 1 | orphan | 56.29 | 0 | 2.07 |
| AH-324-E09 | GORG | 0 | 0 | NA | 1 | 0 | orphan | 0 | 0 | NA | 0 | 0 | NA | 56.24 | 0 | 1.02 |
| AG-343-D22 | GORG | 0 | 0 | NA | 2 | 0 | orphan | 0 | 0 | NA | 0 | 0 | NA | 56.23 | 0 | 0.72 |
| AG-915-N19 | GORG | 0 | 0 | NA | 0 | 0 | NA | 0 | 0 | NA | 0 | 1 | orphan | 56.2 | 0 | 0.92 |
| AG-898-P20 | GORG | 0 | 0 | NA | 2 | 0 | orphan | 0 | 0 | NA | 0 | 0 | NA | 56.17 | 0 | 0.85 |
| AG-901-J06 | GORG | 0 | 0 | NA | 1 | 0 | orphan | 0 | 0 | NA | 0 | 1 | orphan | 56.16 | 0 | 0.68 |
| AH-287-F23 | GORG | 0 | 0 | NA | 2 | 0 | orphan | 0 | 0 | NA | 0 | 0 | NA | 56.16 | 0 | 0.62 |
| AG-453-I03 | GORG | 0 | 0 | NA | 1 | 0 | orphan | 0 | 0 | NA | 0 | 0 | NA | 56.15 | 0 | 0.97 |
| AG-891-I07 | GORG | 1 | 1 | complete | 0 | 0 | NA | 0 | 0 | NA | 0 | 0 | NA | 56.14 | 0 | 0.81 |
| AG-914-D09 | GORG | 0 | 0 | NA | 0 | 0 | NA | 0 | 0 | NA | 0 | 1 | orphan | 56.07 | 0 | 0.95 |
| AG-913-L08 | GORG | 0 | 0 | NA | 2 | 1 | complete | 0 | 0 | NA | 0 | 0 | NA | 56.07 | 0 | 0.94 |
| AG-918-G23 | GORG | 0 | 0 | NA | 2 | 0 | orphan | 0 | 0 | NA | 0 | 0 | NA | 56.07 | 0 | 0.93 |
| AG-422-D22 | GORG | 0 | 0 | NA | 2 | 0 | orphan | 0 | 0 | NA | 0 | 0 | NA | 56.04 | 0 | 0.86 |

| | | | | | | | | | | | | | | | | |
|---|---|---|---|---|---|---|---|---|---|---|---|---|---|---|---|---|
| AG-896-A20 | GORG | 0 | 0 | NA | 1 | 0 | orphan | 0 | 0 | NA | 0 | 0 | NA | 56.03 | 0 | 1.14 |
| AG-439-K19 | GORG | 1 | 1 | complet e | 1 | 0 | orphan | 0 | 0 | NA | 0 | 0 | NA | 56.02 | 0 | 0.71 |
| AG-457-F14 | GORG | 0 | 0 | NA | 1 | 0 | orphan | 0 | 0 | NA | 0 | 0 | NA | 56.02 | 0 | 1.03 |
| AG-904-J19 | GORG | 0 | 0 | NA | 2 | 0 | orphan | 0 | 0 | NA | 0 | 0 | NA | 56.02 | 0 | 0.9 |
| AG-915-N02 | GORG | 0 | 0 | NA | 1 | 0 | orphan | 0 | 0 | NA | 0 | 0 | NA | 56 | 0 | 0.58 |
| AG-919-I17 | GORG | 2 | 0 | orphan | 0 | 0 | NA | 0 | 0 | NA | 0 | 0 | NA | 56 | 0 | 1.59 |
| AG-900-B07 | GORG | 2 | 1 | complet e | 1 | 0 | orphan | 0 | 0 | NA | 0 | 0 | NA | 55.98 | 0 | 1.32 |
| AG-894-G05 | GORG | 0 | 0 | NA | 1 | 0 | orphan | 0 | 0 | NA | 0 | 0 | NA | 55.97 | 0 | 1.09 |
| AG-914-B03 | GORG | 0 | 0 | NA | 1 | 0 | orphan | 0 | 0 | NA | 0 | 0 | NA | 55.92 | 0 | 0.85 |
| AG-439-A03 | GORG | 0 | 0 | NA | 0 | 1 | NA | 0 | 0 | NA | 0 | 0 | NA | 55.81 | 0 | 1 |
| AG-470-G22 | GORG | 0 | 0 | NA | 1 | 0 | orphan | 0 | 0 | NA | 0 | 0 | NA | 55.74 | 0 | 0.69 |
| AG-337-A17 | GORG | 0 | 0 | NA | 1 | 0 | orphan | 0 | 0 | NA | 0 | 0 | NA | 55.71 | 0 | 0.89 |
| AG-426-O18 | GORG | 0 | 0 | NA | 1 | 0 | orphan | 0 | 0 | NA | 0 | 0 | NA | 55.67 | 0 | 0.69 |
| AG-343-N15 | GORG | 0 | 0 | NA | 1 | 0 | orphan | 0 | 0 | NA | 0 | 0 | NA | 55.66 | 0 | 0.84 |
| AG-365-N23 | GORG | 0 | 0 | NA | 1 | 0 | orphan | 0 | 0 | NA | 0 | 0 | NA | 55.66 | 0 | 0.63 |
| AG-430-B08 | GORG | 0 | 0 | NA | 1 | 0 | orphan | 0 | 0 | NA | 0 | 0 | NA | 55.66 | 0 | 0.84 |
| AG-457-L02 | GORG | 0 | 0 | NA | 1 | 0 | orphan | 0 | 0 | NA | 0 | 0 | NA | 55.66 | 0 | 0.79 |
| AG-365-E17 | GORG | 0 | 0 | NA | 1 | 0 | orphan | 0 | 0 | NA | 0 | 0 | NA | 55.63 | 0 | 0.59 |
| AG-325-C09 | GORG | 0 | 0 | NA | 0 | 1 | NA | 0 | 0 | NA | 0 | 0 | NA | 55.61 | 0 | 1.01 |
| AG-920-O20 | GORG | 1 | 1 | complet e | 1 | 0 | orphan | 0 | 0 | NA | 0 | 0 | NA | 55.61 | 0 | 0.57 |
| AG-453-A18 | GORG | 0 | 0 | NA | 1 | 0 | orphan | 0 | 1 | NA | 0 | 0 | NA | 55.59 | 0 | 1.06 |
| AG-453-G03 | GORG | 0 | 0 | NA | 1 | 0 | orphan | 0 | 0 | NA | 0 | 0 | NA | 55.53 | 0 | 0.77 |
| AG-894-G19 | GORG | 0 | 0 | NA | 1 | 0 | orphan | 0 | 0 | NA | 0 | 0 | NA | 55.5 | 0 | 0.98 |
| AG-895-I21 | GORG | 0 | 0 | NA | 1 | 0 | orphan | 0 | 0 | NA | 0 | 0 | NA | 55.42 | 0 | 0.62 |
| AG-907-D05 | GORG | 1 | 0 | orphan | 2 | 0 | orphan | 0 | 0 | NA | 0 | 0 | NA | 55.42 | 0 | 0.99 |
| AG-910-N14 | GORG | 0 | 0 | NA | 1 | 0 | orphan | 0 | 0 | NA | 0 | 0 | NA | 55.42 | 0 | 1.03 |

| | | | | | | | | | | | | | | | | |
|---|---|---|---|---|---|---|---|---|---|---|---|---|---|---|---|---|
| AG-917-G14 | GORG | 0 | 0 | NA | 3 | 0 | orphan | 0 | 1 | NA | 0 | 0 | NA | 55.42 | 0 | 0.91 |
| AH-324-O04 | GORG | 0 | 0 | NA | 1 | 0 | orphan | 0 | 0 | NA | 0 | 0 | NA | 55.42 | 0 | 0.96 |
| AG-911-B19 | GORG | 0 | 0 | NA | 1 | 0 | orphan | 0 | 0 | NA | 0 | 0 | NA | 55.36 | 0 | 0.59 |
| AG-891-J10 | GORG | 0 | 0 | NA | 2 | 0 | orphan | 0 | 0 | NA | 0 | 0 | NA | 55.33 | 0 | 1.55 |
| AG-390-N06 | GORG | 0 | 0 | NA | 1 | 0 | orphan | 0 | 0 | NA | 0 | 0 | NA | 55.32 | 0 | 0.62 |
| AG-913-P05 | GORG | 0 | 0 | NA | 0 | 0 | NA | 0 | 0 | NA | 0 | 1 | orphan | 55.3 | 0 | 0.96 |
| AG-333-A07 | GORG | 0 | 0 | NA | 3 | 0 | orphan | 0 | 0 | NA | 0 | 0 | NA | 55.28 | 0 | 0.64 |
| AG-457-J16 | GORG | 0 | 0 | NA | 1 | 0 | orphan | 0 | 0 | NA | 0 | 0 | NA | 55.27 | 0 | 0.7 |
| AG-422-F22 | GORG | 0 | 0 | NA | 1 | 0 | orphan | 0 | 0 | NA | 0 | 0 | NA | 55.24 | 0 | 0.84 |
| AG-916-A17 | GORG | 0 | 0 | NA | 1 | 0 | orphan | 0 | 0 | NA | 0 | 0 | NA | 55.24 | 0 | 1.2 |
| AG-919-I04 | GORG | 1 | 1 | complete | 0 | 0 | NA | 0 | 0 | NA | 0 | 0 | NA | 55.19 | 0 | 1.27 |
| AG-337-J16 | GORG | 1 | 1 | complete | 0 | 0 | NA | 0 | 0 | NA | 0 | 1 | orphan | 55.18 | 0 | 1.27 |
| AG-896-I04 | GORG | 0 | 0 | NA | 1 | 0 | orphan | 0 | 0 | NA | 0 | 0 | NA | 55.18 | 0 | 0.84 |
| AG-337-A21 | GORG | 0 | 1 | NA | 0 | 0 | NA | 0 | 0 | NA | 0 | 1 | orphan | 55.17 | 0 | 1.35 |
| AG-349-N09 | GORG | 0 | 0 | NA | 1 | 0 | orphan | 0 | 0 | NA | 0 | 0 | NA | 55.17 | 0 | 0.74 |
| AG-426-M17 | GORG | 1 | 1 | complete | 0 | 0 | NA | 0 | 0 | NA | 0 | 0 | NA | 55.17 | 0 | 1.33 |
| AG-891-E05 | GORG | 0 | 0 | NA | 1 | 1 | orphan | 0 | 0 | NA | 0 | 0 | NA | 55.17 | 0 | 0.92 |
| AH-287-J03 | GORG | 0 | 0 | NA | 1 | 0 | orphan | 1 | 1 | complete | 0 | 0 | NA | 55.17 | 0 | 1.12 |
| AG-910-I07 | GORG | 0 | 0 | NA | 1 | 0 | orphan | 0 | 0 | NA | 0 | 0 | NA | 55.15 | 0 | 0.96 |
| AH-326-N10 | GORG | 0 | 0 | NA | 2 | 0 | orphan | 0 | 0 | NA | 0 | 0 | NA | 55.11 | 0 | 0.63 |
| AG-453-B10 | GORG | 0 | 0 | NA | 1 | 0 | orphan | 0 | 0 | NA | 0 | 0 | NA | 55.08 | 0 | 0.77 |
| AG-390-F23 | GORG | 1 | 1 | complete | 0 | 0 | NA | 0 | 0 | NA | 0 | 1 | orphan | 55.07 | 0 | 0.99 |
| AG-457-N16 | GORG | 0 | 0 | NA | 3 | 0 | orphan | 0 | 0 | NA | 0 | 0 | NA | 55.07 | 0 | 1.08 |
| AG-422-I10 | GORG | 0 | 0 | NA | 1 | 0 | orphan | 0 | 0 | NA | 0 | 0 | NA | 55.04 | 0 | 0.68 |
| AG-349-A01 | GORG | 0 | 0 | NA | 1 | 0 | orphan | 0 | 0 | NA | 0 | 0 | NA | 55.03 | 0 | 0.91 |

| | | | | | | | | | | | | | | | | |
|---|---|---|---|---|---|---|---|---|---|---|---|---|---|---|---|---|
| AG-319-O22 | GORG | 0 | 0 | NA | 1 | 0 | orphan | 0 | 0 | NA | 0 | 1 | orphan | 55.02 | 0 | 0.88 |
| AG-447-J13 | GORG | 0 | 0 | NA | 1 | 0 | orphan | 0 | 0 | NA | 0 | 0 | NA | 55.02 | 0 | 0.74 |
| AG-447-M04 | GORG | 0 | 0 | NA | 1 | 0 | orphan | 0 | 0 | NA | 0 | 0 | NA | 55.02 | 0 | 0.69 |
| AG-457-G18 | GORG | 0 | 0 | NA | 1 | 0 | orphan | 0 | 0 | NA | 0 | 0 | NA | 54.95 | 0 | 0.64 |
| AH-287-F03 | GORG | 0 | 0 | NA | 0 | 0 | NA | 0 | 0 | NA | 0 | 1 | orphan | 54.94 | 0 | 0.99 |
| AG-895-J18 | GORG | 0 | 0 | NA | 1 | 0 | orphan | 0 | 0 | NA | 0 | 0 | NA | 54.93 | 0 | 1.24 |
| AG-461-O03 | GORG | 0 | 0 | NA | 0 | 0 | NA | 0 | 0 | NA | 0 | 1 | orphan | 54.89 | 0 | 0.89 |
| AG-325-D03 | GORG | 0 | 0 | NA | 1 | 0 | orphan | 0 | 0 | NA | 0 | 0 | NA | 54.89 | 0 | 0.6 |
| AG-414-L18 | GORG | 0 | 0 | NA | 1 | 0 | orphan | 0 | 0 | NA | 0 | 0 | NA | 54.89 | 0 | 1.66 |
| AG-910-M08 | GORG | 0 | 0 | NA | 1 | 0 | orphan | 0 | 0 | NA | 0 | 0 | NA | 54.89 | 0 | 0.76 |
| AG-909-E18 | GORG | 0 | 0 | NA | 1 | 0 | orphan | 0 | 0 | NA | 0 | 0 | NA | 54.84 | 0 | 0.73 |
| AG-390-F09 | GORG | 0 | 0 | NA | 1 | 0 | orphan | 0 | 0 | NA | 0 | 0 | NA | 54.82 | 0 | 1.07 |
| AG-457-K13 | GORG | 0 | 0 | NA | 1 | 0 | orphan | 0 | 0 | NA | 0 | 0 | NA | 54.82 | 0 | 0.82 |
| AG-915-N04 | GORG | 0 | 0 | NA | 1 | 0 | orphan | 0 | 0 | NA | 0 | 0 | NA | 54.82 | 0 | 0.73 |
| AG-898-K05 | GORG | 0 | 0 | NA | 1 | 0 | orphan | 1 | 1 | orphan | 0 | 0 | NA | 54.79 | 0 | 0.82 |
| AG-325-K17 | GORG | 1 | 1 | orphan | 2 | 1 | complete | 0 | 0 | NA | 0 | 0 | NA | 54.72 | 0 | 0.8 |
| AG-349-L08 | GORG | 0 | 0 | NA | 1 | 0 | orphan | 0 | 0 | NA | 0 | 0 | NA | 54.72 | 0 | 0.64 |
| AG-430-A10 | GORG | 0 | 0 | NA | 2 | 1 | complete | 0 | 0 | NA | 0 | 0 | NA | 54.72 | 0 | 0.82 |
| AG-461-A11 | GORG | 0 | 0 | NA | 2 | 0 | orphan | 0 | 0 | NA | 0 | 0 | NA | 54.72 | 0 | 0.9 |
| AG-893-O09 | GORG | 0 | 0 | NA | 2 | 0 | orphan | 0 | 1 | NA | 0 | 0 | NA | 54.72 | 0 | 1.11 |
| AG-896-G14 | GORG | 0 | 0 | NA | 1 | 0 | orphan | 0 | 0 | NA | 0 | 0 | NA | 54.72 | 0 | 0.69 |
| AG-901-J10 | GORG | 0 | 0 | NA | 1 | 0 | orphan | 0 | 0 | NA | 0 | 0 | NA | 54.72 | 0 | 0.83 |
| AG-904-C13 | GORG | 0 | 0 | NA | 1 | 0 | orphan | 0 | 0 | NA | 0 | 0 | NA | 54.72 | 0 | 0.92 |
| AG-904-O11 | GORG | 0 | 0 | NA | 1 | 0 | orphan | 0 | 0 | NA | 0 | 0 | NA | 54.72 | 0 | 0.93 |
| AG-911-J21 | GORG | 0 | 0 | NA | 2 | 0 | orphan | 0 | 0 | NA | 0 | 0 | NA | 54.72 | 0 | 0.93 |

| | | | | | | | | | | | | | | | | |
|---|---|---|---|---|---|---|---|---|---|---|---|---|---|---|---|---|
| AG-911-O11 | GORG | 0 | 0 | NA | 1 | 0 | orphan | 0 | 0 | NA | 0 | 0 | NA | 54.72 | 0 | 0.78 |
| AG-913-L05 | GORG | 0 | 0 | NA | 1 | 0 | orphan | 0 | 0 | NA | 0 | 0 | NA | 54.72 | 0 | 0.68 |
| AG-915-P10 | GORG | 0 | 0 | NA | 1 | 0 | orphan | 0 | 0 | NA | 0 | 0 | NA | 54.72 | 0 | 0.69 |
| AG-919-I10 | GORG | 0 | 0 | NA | 1 | 0 | orphan | 0 | 0 | NA | 0 | 0 | NA | 54.72 | 0 | 0.84 |
| AG-422-P02 | GORG | 0 | 0 | NA | 2 | 0 | orphan | 0 | 1 | NA | 0 | 0 | NA | 54.7 | 0 | 0.89 |
| AG-900-K10 | GORG | 0 | 0 | NA | 1 | 0 | orphan | 0 | 0 | NA | 0 | 0 | NA | 54.69 | 0 | 0.86 |
| AH-287-G14 | GORG | 0 | 0 | NA | 2 | 0 | orphan | 0 | 0 | NA | 0 | 0 | NA | 54.68 | 0 | 0.53 |
| AG-903-O09 | GORG | 0 | 0 | NA | 1 | 0 | orphan | 0 | 0 | NA | 0 | 1 | orphan | 54.66 | 0 | 0.93 |
| AG-901-A23 | GORG | 1 | 1 | complete | 1 | 0 | orphan | 0 | 0 | NA | 0 | 0 | NA | 54.62 | 0 | 0.85 |
| AG-916-E18 | GORG | 1 | 1 | complete | 0 | 0 | NA | 0 | 0 | NA | 0 | 0 | NA | 54.62 | 0 | 0.84 |
| AG-900-F02 | GORG | 0 | 0 | NA | 3 | 0 | orphan | 0 | 0 | NA | 0 | 0 | NA | 54.59 | 0 | 1.09 |
| AG-359-F07 | GORG | 0 | 0 | NA | 1 | 1 | complete | 0 | 0 | NA | 0 | 0 | NA | 54.56 | 0 | 1.05 |
| AG-903-G22 | GORG | 0 | 0 | NA | 2 | 0 | orphan | 0 | 0 | NA | 0 | 0 | NA | 54.56 | 0 | 0.63 |
| AG-404-G02 | GORG | 0 | 0 | NA | 0 | 0 | NA | 0 | 0 | NA | 0 | 1 | orphan | 54.55 | 0 | 1.65 |
| AG-453-D18 | GORG | 1 | 1 | complete | 0 | 0 | NA | 0 | 0 | NA | 0 | 0 | NA | 54.55 | 0 | 1.56 |
| AG-911-D15 | GORG | 0 | 0 | NA | 0 | 0 | NA | 0 | 0 | NA | 0 | 1 | orphan | 54.49 | 0 | 1.34 |
| AH-321-O11 | GORG | 0 | 0 | NA | 2 | 0 | orphan | 0 | 0 | NA | 0 | 0 | NA | 54.49 | 0 | 0.62 |
| AG-319-N19 | GORG | 0 | 0 | NA | 0 | 0 | NA | 0 | 0 | NA | 0 | 1 | orphan | 54.47 | 0 | 0.96 |
| AG-453-O22 | GORG | 0 | 0 | NA | 1 | 0 | orphan | 0 | 0 | NA | 0 | 0 | NA | 54.45 | 0 | 1.59 |
| AG-903-C07 | GORG | 0 | 0 | NA | 1 | 0 | orphan | 0 | 0 | NA | 0 | 0 | NA | 54.4 | 0 | 0.76 |
| AH-324-C07 | GORG | 0 | 0 | NA | 3 | 0 | orphan | 0 | 0 | NA | 0 | 0 | NA | 54.39 | 0 | 0.93 |
| AG-337-N20 | GORG | 0 | 0 | NA | 2 | 0 | orphan | 0 | 0 | NA | 0 | 0 | NA | 54.38 | 0 | 0.84 |
| AG-919-D16 | GORG | 0 | 0 | NA | 1 | 0 | orphan | 0 | 0 | NA | 0 | 0 | NA | 54.35 | 0 | 0.68 |
| AG-360-N11 | GORG | 0 | 0 | NA | 1 | 0 | orphan | 0 | 0 | NA | 0 | 0 | NA | 54.34 | 0 | 0.65 |

| | | | | | | | | | | | | | | | | |
|---|---|---|---|---|---|---|---|---|---|---|---|---|---|---|---|---|
| AG-899-F03 | GORG | 0 | 0 | NA | 1 | 0 | orphan | 0 | 0 | NA | 0 | 0 | NA | 54.33 | 0 | 1.07 |
| AG-390-L23 | GORG | 0 | 0 | NA | 1 | 0 | orphan | 0 | 0 | NA | 0 | 0 | NA | 54.31 | 0 | 0.98 |
| AG-435-D20 | GORG | 0 | 0 | NA | 1 | 0 | orphan | 0 | 0 | NA | 0 | 0 | NA | 54.31 | 0 | 1.34 |
| AG-916-L16 | GORG | 0 | 0 | NA | 2 | 0 | orphan | 0 | 0 | NA | 0 | 0 | NA | 54.31 | 0 | 0.87 |
| AG-470-F02 | GORG | 0 | 0 | NA | 1 | 0 | orphan | 0 | 0 | NA | 0 | 0 | NA | 54.27 | 0 | 0.85 |
| AG-337-O02 | GORG | 0 | 0 | NA | 2 | 0 | orphan | 0 | 0 | NA | 0 | 0 | NA | 54.22 | 0 | 0.76 |
| AG-365-K15 | GORG | 0 | 0 | NA | 1 | 0 | orphan | 0 | 0 | NA | 0 | 0 | NA | 54.22 | 0 | 0.77 |
| AG-912-O16 | GORG | 0 | 0 | NA | 2 | 0 | orphan | 0 | 1 | NA | 0 | 0 | NA | 54.22 | 0 | 0.69 |
| AG-915-O13 | GORG | 0 | 0 | NA | 1 | 0 | orphan | 0 | 0 | NA | 0 | 0 | NA | 54.22 | 0 | 0.7 |
| AG-915-A11 | GORG | 0 | 0 | NA | 0 | 0 | NA | 0 | 0 | NA | 0 | 1 | orphan | 54.21 | 0 | 0.88 |
| AG-914-A11 | GORG | 0 | 0 | NA | 1 | 1 | complete | 0 | 0 | NA | 0 | 0 | NA | 54.17 | 0 | 1.04 |
| AG-325-O18 | GORG | 0 | 0 | NA | 1 | 0 | orphan | 0 | 0 | NA | 0 | 0 | NA | 54.08 | 0 | 1.04 |
| AG-917-G18 | GORG | 0 | 0 | NA | 1 | 0 | orphan | 0 | 0 | NA | 0 | 0 | NA | 54.08 | 0 | 2.13 |
| AG-343-N02 | GORG | 0 | 0 | NA | 4 | 0 | orphan | 0 | 0 | NA | 0 | 2 | orphan | 54.07 | 0 | 0.86 |
| AG-414-K06 | GORG | 0 | 0 | NA | 1 | 0 | orphan | 0 | 0 | NA | 0 | 0 | NA | 54 | 0 | 0.98 |
| AG-910-F15 | GORG | 0 | 0 | NA | 1 | 0 | orphan | 0 | 0 | NA | 0 | 0 | NA | 53.99 | 0 | 0.79 |
| AG-343-B07 | GORG | 0 | 0 | NA | 1 | 0 | orphan | 0 | 0 | NA | 0 | 0 | NA | 53.95 | 0 | 0.77 |
| AG-911-D20 | GORG | 0 | 0 | NA | 1 | 0 | orphan | 0 | 0 | NA | 0 | 0 | NA | 53.95 | 0 | 0.73 |
| AG-909-L15 | GORG | 0 | 0 | NA | 1 | 0 | orphan | 0 | 0 | NA | 0 | 0 | NA | 53.94 | 0 | 0.73 |
| AG-422-I22 | GORG | 0 | 0 | NA | 2 | 0 | orphan | 0 | 0 | NA | 0 | 0 | NA | 53.92 | 0 | 0.98 |
| AG-325-I03 | GORG | 0 | 0 | NA | 1 | 0 | orphan | 0 | 0 | NA | 0 | 0 | NA | 53.89 | 0 | 0.89 |
| AG-422-F19 | GORG | 2 | 1 | complete | 0 | 0 | NA | 0 | 0 | NA | 0 | 0 | NA | 53.88 | 0 | 0.64 |
| AG-447-D23 | GORG | 0 | 0 | NA | 2 | 0 | orphan | 0 | 0 | NA | 0 | 0 | NA | 53.86 | 0 | 0.83 |
| AG-426-O11 | GORG | 0 | 0 | NA | 1 | 0 | orphan | 0 | 0 | NA | 0 | 0 | NA | 53.85 | 0 | 1.27 |
| AG-337-N19 | GORG | 0 | 0 | NA | 1 | 0 | orphan | 0 | 0 | NA | 0 | 0 | NA | 53.84 | 0 | 0.92 |
| AG-900-B21 | GORG | 1 | 1 | complet | 0 | 0 | NA | 0 | 0 | NA | 0 | 0 | NA | 53.83 | 0 | 1.26 |

| | | | | | | | | | | | | | | | | |
|---|---|---|---|---|---|---|---|---|---|---|---|---|---|---|---|---|
| | | | | e | | | | | | | | | | | | |
| AG-360-P19 | GORG | 0 | 0 | NA | 1 | 0 | orphan | 0 | 0 | NA | 0 | 0 | NA | 53.78 | 0 | 0.66 |
| AG-404-O23 | GORG | 0 | 0 | NA | 1 | 0 | orphan | 0 | 0 | NA | 0 | 0 | NA | 53.77 | 0 | 0.63 |
| AG-904-B15 | GORG | 0 | 0 | NA | 1 | 0 | orphan | 0 | 0 | NA | 0 | 0 | NA | 53.74 | 0 | 0.74 |
| AG-913-N19 | GORG | 0 | 0 | NA | 1 | 0 | orphan | 0 | 0 | NA | 0 | 0 | NA | 53.71 | 0 | 0.77 |
| AG-905-M08 | GORG | 0 | 0 | NA | 2 | 0 | orphan | 0 | 0 | NA | 0 | 0 | NA | 53.7 | 0 | 0.94 |
| AG-898-G10 | GORG | 0 | 0 | NA | 1 | 0 | orphan | 0 | 0 | NA | 0 | 0 | NA | 53.67 | 0 | 0.84 |
| AG-910-D07 | GORG | 0 | 0 | NA | 1 | 0 | orphan | 0 | 0 | NA | 0 | 1 | orphan | 53.63 | 0 | 0.84 |
| AH-321-M03 | GORG | 0 | 0 | NA | 1 | 0 | orphan | 0 | 0 | NA | 0 | 0 | NA | 53.63 | 0 | 0.68 |
| AG-893-O23 | GORG | 0 | 0 | NA | 0 | 0 | NA | 0 | 0 | NA | 0 | 1 | orphan | 53.62 | 0 | 0.8 |
| AG-892-D02 | GORG | 0 | 0 | NA | 5 | 1 | complete | 0 | 0 | NA | 0 | 1 | orphan | 53.61 | 0 | 1.45 |
| AG-903-E05 | GORG | 1 | 1 | complete | 1 | 0 | orphan | 0 | 0 | NA | 0 | 0 | NA | 53.61 | 0 | 0.98 |
| AG-909-J09 | GORG | 1 | 1 | complete | 1 | 0 | orphan | 0 | 0 | NA | 0 | 0 | NA | 53.57 | 0 | 0.55 |
| AG-430-K11 | GORG | 0 | 0 | NA | 3 | 0 | orphan | 0 | 1 | NA | 0 | 0 | NA | 53.56 | 0 | 0.69 |
| AG-426-B02 | GORG | 1 | 1 | complete | 0 | 0 | NA | 0 | 0 | NA | 0 | 2 | orphan | 53.55 | 0 | 1.26 |
| AG-917-L16 | GORG | 0 | 0 | NA | 1 | 0 | orphan | 0 | 0 | NA | 0 | 0 | NA | 53.48 | 0 | 0.82 |
| AG-337-J18 | GORG | 1 | 0 | orphan | 0 | 0 | NA | 0 | 0 | NA | 0 | 0 | NA | 53.45 | 0 | 1.1 |
| AG-426-L02 | GORG | 0 | 0 | NA | 1 | 0 | orphan | 0 | 0 | NA | 0 | 0 | NA | 53.45 | 0 | 0.94 |
| AG-900-F13 | GORG | 0 | 0 | NA | 1 | 0 | orphan | 0 | 0 | NA | 0 | 0 | NA | 53.45 | 0 | 1.07 |
| AG-905-D11 | GORG | 0 | 0 | NA | 1 | 0 | orphan | 1 | 3 | complete | 0 | 0 | NA | 53.45 | 0 | 0.83 |
| AG-435-C18 | GORG | 0 | 0 | NA | 1 | 0 | orphan | 0 | 0 | NA | 0 | 0 | NA | 53.44 | 0 | 0.64 |
| AG-333-L23 | GORG | 0 | 0 | NA | 1 | 0 | orphan | 0 | 0 | NA | 0 | 0 | NA | 53.41 | 0 | 1.39 |
| AG-891-D07 | GORG | 0 | 0 | NA | 1 | 0 | orphan | 0 | 0 | NA | 0 | 0 | NA | 53.4 | 0 | 0.96 |
| AG-912-J22 | GORG | 0 | 0 | NA | 1 | 0 | orphan | 0 | 0 | NA | 0 | 0 | NA | 53.4 | 0 | 0.82 |
| AG-439-J13 | GORG | 0 | 0 | NA | 1 | 0 | orphan | 0 | 0 | NA | 0 | 0 | NA | 53.37 | 0 | 0.84 |

| | | | | | | | | | | | | | | | | |
|---|---|---|---|---|---|---|---|---|---|---|---|---|---|---|---|---|
| AG-464-F23 | GORG | 1 | 1 | complete | 1 | 0 | orphan | 0 | 0 | NA | 0 | 0 | NA | 53.37 | 0 | 0.54 |
| AG-899-L15 | GORG | 0 | 0 | NA | 1 | 1 | orphan | 0 | 0 | NA | 0 | 0 | NA | 53.34 | 0 | 0.66 |
| AG-903-B06 | GORG | 0 | 0 | NA | 1 | 0 | orphan | 0 | 0 | NA | 0 | 0 | NA | 53.3 | 0 | 0.6 |
| AG-426-I23 | GORG | 0 | 0 | NA | 0 | 0 | NA | 0 | 0 | NA | 0 | 1 | orphan | 53.29 | 0 | 0.92 |
| AG-896-L15 | GORG | 0 | 0 | NA | 1 | 0 | orphan | 0 | 0 | NA | 0 | 0 | NA | 53.29 | 0 | 0.71 |
| AG-910-J20 | GORG | 0 | 0 | NA | 1 | 0 | orphan | 0 | 0 | NA | 0 | 0 | NA | 53.29 | 0 | 0.61 |
| AG-422-M20 | GORG | 0 | 0 | NA | 2 | 0 | orphan | 0 | 0 | NA | 0 | 1 | orphan | 53.28 | 0 | 0.98 |
| AG-907-J20 | GORG | 0 | 0 | NA | 1 | 0 | orphan | 0 | 0 | NA | 0 | 0 | NA | 53.28 | 0 | 0.85 |
| AG-313-I14 | GORG | 0 | 0 | NA | 1 | 0 | orphan | 0 | 0 | NA | 0 | 0 | NA | 53.14 | 0 | 1.12 |
| AG-915-P07 | GORG | 0 | 0 | NA | 1 | 0 | orphan | 0 | 0 | NA | 0 | 0 | NA | 53.11 | 0 | 0.93 |
| AG-892-O21 | GORG | 0 | 0 | NA | 0 | 0 | NA | 0 | 0 | NA | 0 | 1 | orphan | 53.08 | 0 | 0.95 |
| AG-430-L07 | GORG | 0 | 0 | NA | 2 | 0 | orphan | 0 | 0 | NA | 0 | 0 | NA | 53.08 | 0 | 0.61 |
| AG-333-E02 | GORG | 1 | 1 | complete | 1 | 0 | orphan | 0 | 0 | NA | 0 | 0 | NA | 53.01 | 0 | 0.61 |
| AG-414-I06 | GORG | 0 | 0 | NA | 2 | 0 | orphan | 0 | 0 | NA | 0 | 0 | NA | 53.01 | 0 | 0.49 |
| AG-903-K13 | GORG | 0 | 0 | NA | 1 | 0 | orphan | 0 | 1 | NA | 0 | 0 | NA | 53.01 | 0 | 0.68 |
| AG-904-O02 | GORG | 0 | 0 | NA | 1 | 0 | orphan | 0 | 0 | NA | 0 | 0 | NA | 53.01 | 0 | 0.54 |
| AG-914-K13 | GORG | 0 | 0 | NA | 3 | 0 | orphan | 0 | 0 | NA | 0 | 0 | NA | 53.01 | 0 | 0.99 |
| AH-324-E10 | GORG | 0 | 0 | NA | 1 | 0 | orphan | 0 | 0 | NA | 0 | 0 | NA | 53.01 | 0 | 0.7 |
| AH-324-J11 | GORG | 1 | 1 | complete | 1 | 0 | orphan | 0 | 0 | NA | 0 | 0 | NA | 53.01 | 0 | 0.6 |
| AH-326-D08 | GORG | 0 | 0 | NA | 1 | 1 | complete | 0 | 0 | NA | 0 | 0 | NA | 53.01 | 0 | 0.89 |
| AG-365-G10 | GORG | 0 | 0 | NA | 1 | 0 | orphan | 0 | 0 | NA | 0 | 0 | NA | 53 | 0 | 0.77 |
| AG-896-L16 | GORG | 1 | 1 | complete | 1 | 0 | orphan | 0 | 0 | NA | 0 | 0 | NA | 53 | 0 | 0.92 |
| AG-414-N05 | GORG | 2 | 1 | complete | 0 | 0 | NA | 0 | 0 | NA | 0 | 0 | NA | 52.99 | 0 | 0.84 |
| AG-453-E11 | GORG | 0 | 0 | NA | 1 | 0 | orphan | 0 | 0 | NA | 0 | 0 | NA | 52.99 | 0 | 0.73 |

| | | | | | | | | | | | | | | | | |
|---|---|---|---|---|---|---|---|---|---|---|---|---|---|---|---|---|
| AG-461-C13 | GORG | 1 | 1 | complete | 0 | 0 | NA | 0 | 0 | NA | 0 | 0 | NA | 52.99 | 0 | 1.18 |
| AG-337-I20 | GORG | 0 | 0 | NA | 1 | 0 | orphan | 0 | 0 | NA | 0 | 0 | NA | 52.95 | 0 | 0.91 |
| AG-892-A04 | GORG | 0 | 0 | NA | 1 | 1 | orphan | 0 | 0 | NA | 0 | 0 | NA | 52.94 | 0 | 1.43 |
| AG-319-F19 | GORG | 1 | 1 | complete | 2 | 0 | orphan | 0 | 0 | NA | 0 | 0 | NA | 52.91 | 0 | 0.61 |
| AG-325-I13 | GORG | 0 | 0 | NA | 1 | 0 | orphan | 0 | 0 | NA | 0 | 0 | NA | 52.9 | 0 | 0.74 |
| AG-464-G19 | GORG | 1 | 0 | orphan | 1 | 0 | orphan | 0 | 0 | NA | 0 | 0 | NA | 52.9 | 0 | 0.77 |
| AG-891-K07 | GORG | 1 | 1 | complete | 0 | 0 | NA | 0 | 0 | NA | 0 | 0 | NA | 52.9 | 0 | 0.89 |
| AG-892-M17 | GORG | 0 | 0 | NA | 3 | 0 | orphan | 0 | 0 | NA | 0 | 0 | NA | 52.88 | 0 | 1.43 |
| AG-910-E03 | GORG | 0 | 0 | NA | 2 | 0 | orphan | 0 | 0 | NA | 0 | 2 | orphan | 52.83 | 0 | 0.77 |
| AG-337-E13 | GORG | 0 | 0 | NA | 2 | 0 | orphan | 0 | 0 | NA | 0 | 0 | NA | 52.83 | 0 | 0.78 |
| AG-349-D16 | GORG | 0 | 0 | NA | 2 | 0 | orphan | 0 | 0 | NA | 0 | 0 | NA | 52.83 | 0 | 0.96 |
| AG-359-P04 | GORG | 0 | 0 | NA | 1 | 0 | orphan | 0 | 0 | NA | 0 | 0 | NA | 52.83 | 0 | 0.8 |
| AG-365-P13 | GORG | 0 | 0 | NA | 1 | 0 | orphan | 0 | 0 | NA | 0 | 0 | NA | 52.83 | 0 | 0.96 |
| AG-390-P04 | GORG | 0 | 0 | NA | 2 | 0 | orphan | 0 | 0 | NA | 0 | 0 | NA | 52.83 | 0 | 0.82 |
| AG-414-L03 | GORG | 0 | 0 | NA | 1 | 0 | orphan | 0 | 0 | NA | 0 | 0 | NA | 52.83 | 0 | 0.84 |
| AG-422-J14 | GORG | 0 | 0 | NA | 2 | 0 | orphan | 0 | 0 | NA | 0 | 0 | NA | 52.83 | 0 | 0.94 |
| AG-435-P07 | GORG | 0 | 0 | NA | 1 | 0 | orphan | 0 | 0 | NA | 0 | 0 | NA | 52.83 | 0 | 0.79 |
| AG-457-F22 | GORG | 0 | 0 | NA | 1 | 0 | orphan | 0 | 0 | NA | 0 | 0 | NA | 52.83 | 0 | 0.56 |
| AG-461-O08 | GORG | 1 | 1 | complete | 1 | 0 | orphan | 0 | 0 | NA | 0 | 0 | NA | 52.83 | 0 | 0.72 |
| AG-470-P13 | GORG | 0 | 0 | NA | 1 | 0 | orphan | 0 | 0 | NA | 0 | 0 | NA | 52.83 | 0 | 0.64 |
| AG-893-G06 | GORG | 0 | 0 | NA | 2 | 0 | orphan | 0 | 0 | NA | 0 | 0 | NA | 52.83 | 0 | 0.63 |
| AG-901-B19 | GORG | 0 | 0 | NA | 1 | 0 | orphan | 0 | 0 | NA | 0 | 0 | NA | 52.83 | 0 | 0.79 |
| AG-901-C16 | GORG | 0 | 0 | NA | 2 | 1 | complete | 0 | 0 | NA | 0 | 0 | NA | 52.83 | 0 | 0.72 |
| AG-903-C03 | GORG | 0 | 0 | NA | 1 | 0 | orphan | 0 | 0 | NA | 0 | 0 | NA | 52.83 | 0 | 0.79 |
| AG-903-D18 | GORG | 0 | 0 | NA | 1 | 0 | orphan | 0 | 0 | NA | 0 | 0 | NA | 52.83 | 0 | 0.65 |

| | | | | | | | | | | | | | | | | |
|---|---|---|---|---|---|---|---|---|---|---|---|---|---|---|---|---|
| AG-905-M20 | GORG | 0 | 0 | NA | 1 | 0 | orphan | 0 | 0 | NA | 0 | 0 | NA | 52.83 | 0 | 0.92 |
| AG-912-L09 | GORG | 0 | 0 | NA | 1 | 0 | orphan | 0 | 0 | NA | 0 | 0 | NA | 52.83 | 0 | 0.6 |
| AG-913-D16 | GORG | 0 | 0 | NA | 1 | 0 | orphan | 0 | 0 | NA | 0 | 0 | NA | 52.83 | 0 | 0.81 |
| AH-326-N23 | GORG | 0 | 0 | NA | 1 | 0 | orphan | 0 | 0 | NA | 0 | 0 | NA | 52.83 | 0 | 0.72 |
| AG-904-J16 | GORG | 1 | 1 | complet e | 1 | 0 | orphan | 0 | 0 | NA | 0 | 0 | NA | 52.82 | 0 | 0.99 |
| AG-892-M18 | GORG | 0 | 0 | NA | 0 | 0 | NA | 0 | 0 | NA | 0 | 1 | orpha n | 52.81 | 0 | 0.88 |
| AG-912-P11 | GORG | 0 | 0 | NA | 1 | 0 | orphan | 0 | 0 | NA | 0 | 0 | NA | 52.8 | 0 | 0.96 |
| AG-914-C20 | GORG | 0 | 0 | NA | 1 | 0 | orphan | 0 | 0 | NA | 0 | 0 | NA | 52.74 | 0 | 0.83 |
| AG-333-P23 | GORG | 0 | 0 | NA | 1 | 0 | orphan | 0 | 0 | NA | 0 | 0 | NA | 52.72 | 0 | 0.92 |
| AG-905-E08 | GORG | 0 | 0 | NA | 1 | 0 | orphan | 0 | 0 | NA | 0 | 0 | NA | 52.72 | 0 | 0.59 |
| AG-896-L05 | GORG | 0 | 0 | NA | 1 | 0 | orphan | 0 | 0 | NA | 0 | 0 | NA | 52.68 | 0 | 0.65 |
| AG-390-K06 | GORG | 0 | 0 | NA | 1 | 0 | orphan | 0 | 0 | NA | 0 | 0 | NA | 52.67 | 0 | 0.66 |
| AG-915-B20 | GORG | 0 | 0 | NA | 1 | 0 | orphan | 0 | 0 | NA | 0 | 0 | NA | 52.67 | 0 | 0.69 |
| AG-912-E20 | GORG | 0 | 0 | NA | 2 | 0 | orphan | 0 | 0 | NA | 0 | 0 | NA | 52.66 | 0 | 0.72 |
| AG-896-N09 | GORG | 0 | 0 | NA | 0 | 0 | NA | 0 | 0 | NA | 0 | 1 | orpha n | 52.65 | 0 | 0.91 |
| AG-349-L06 | GORG | 0 | 0 | NA | 1 | 0 | orphan | 0 | 0 | NA | 0 | 0 | NA | 52.63 | 0 | 1 |
| AG-898-P17 | GORG | 0 | 0 | NA | 1 | 0 | orphan | 0 | 0 | NA | 0 | 0 | NA | 52.62 | 0 | 0.8 |
| AG-905-D21 | GORG | 0 | 0 | NA | 2 | 0 | orphan | 0 | 0 | NA | 0 | 0 | NA | 52.61 | 0 | 0.81 |
| AG-390-F10 | GORG | 1 | 1 | complet e | 0 | 0 | NA | 0 | 0 | NA | 0 | 0 | NA | 52.59 | 0 | 1.47 |
| AG-912-P13 | GORG | 0 | 0 | NA | 1 | 0 | orphan | 0 | 0 | NA | 0 | 0 | NA | 52.59 | 0 | 0.68 |
| AG-904-I16 | GORG | 0 | 0 | NA | 2 | 0 | orphan | 0 | 0 | NA | 0 | 0 | NA | 52.58 | 0 | 0.72 |
| AG-892-N14 | GORG | 0 | 1 | NA | 0 | 0 | NA | 0 | 0 | NA | 0 | 0 | NA | 52.57 | 0 | 0.69 |
| AG-343-A13 | GORG | 0 | 0 | NA | 1 | 0 | orphan | 0 | 0 | NA | 0 | 1 | orpha n | 52.54 | 0 | 0.7 |
| AG-893-D02 | GORG | 0 | 0 | NA | 0 | 0 | NA | 0 | 0 | NA | 0 | 1 | orpha n | 52.53 | 0 | 0.9 |
| AG-337-M23 | GORG | 1 | 1 | complet e | 0 | 0 | NA | 0 | 0 | NA | 0 | 1 | orpha n | 52.52 | 0 | 0.99 |

| | | | | | | | | | | | | | | | | |
|---|---|---|---|---|---|---|---|---|---|---|---|---|---|---|---|---|
| AG-349-B19 | GORG | 0 | 0 | NA | 1 | 0 | orphan | 0 | 0 | NA | 0 | 0 | NA | 52.52 | 0 | 0.54 |
| AG-313-O08 | GORG | 0 | 0 | NA | 1 | 0 | orphan | 0 | 0 | NA | 0 | 0 | NA | 52.51 | 0 | 1.14 |
| AG-914-K03 | GORG | 0 | 0 | NA | 0 | 1 | NA | 0 | 0 | NA | 0 | 0 | NA | 52.48 | 0 | 0.79 |
| AG-313-J15 | GORG | 0 | 0 | NA | 1 | 0 | orphan | 0 | 0 | NA | 0 | 0 | NA | 52.46 | 0 | 1.22 |
| AG-894-M16 | GORG | 0 | 0 | NA | 1 | 0 | orphan | 0 | 0 | NA | 0 | 0 | NA | 52.44 | 0 | 0.91 |
| AG-461-E22 | GORG | 1 | 1 | complete | 1 | 0 | orphan | 0 | 0 | NA | 0 | 0 | NA | 52.41 | 0 | 1.16 |
| AG-892-M13 | GORG | 0 | 0 | NA | 1 | 0 | orphan | 0 | 0 | NA | 0 | 0 | NA | 52.41 | 0 | 0.88 |
| AG-898-L13 | GORG | 0 | 0 | NA | 1 | 0 | orphan | 0 | 0 | NA | 0 | 0 | NA | 52.41 | 0 | 0.95 |
| AG-905-L21 | GORG | 0 | 0 | NA | 2 | 0 | orphan | 0 | 0 | NA | 0 | 0 | NA | 52.41 | 0 | 0.99 |
| AG-365-N05 | GORG | 1 | 1 | complete | 0 | 0 | NA | 0 | 0 | NA | 0 | 0 | NA | 52.35 | 0 | 1.55 |
| AG-899-I20 | GORG | 0 | 0 | NA | 2 | 0 | orphan | 0 | 0 | NA | 0 | 0 | NA | 52.35 | 0 | 0.71 |
| AG-461-F13 | GORG | 0 | 0 | NA | 2 | 0 | orphan | 0 | 0 | NA | 0 | 0 | NA | 52.34 | 0 | 1.24 |
| AG-919-M17 | GORG | 0 | 0 | NA | 1 | 0 | orphan | 0 | 0 | NA | 0 | 0 | NA | 52.34 | 0 | 0.84 |
| AG-908-I06 | GORG | 0 | 0 | NA | 1 | 0 | orphan | 0 | 0 | NA | 0 | 0 | NA | 52.32 | 0 | 0.72 |
| AG-907-J09 | GORG | 0 | 0 | NA | 1 | 0 | orphan | 0 | 0 | NA | 0 | 0 | NA | 52.31 | 0 | 0.95 |
| AH-287-A06 | GORG | 0 | 0 | NA | 0 | 0 | NA | 0 | 0 | NA | 0 | 3 | orphan | 52.29 | 0 | 0.85 |
| AG-919-J06 | GORG | 0 | 0 | NA | 1 | 0 | orphan | 1 | 1 | complete | 0 | 0 | NA | 52.25 | 0 | 0.81 |
| AH-324-J15 | GORG | 0 | 0 | NA | 2 | 0 | orphan | 0 | 0 | NA | 0 | 0 | NA | 52.25 | 0 | 2.09 |
| AH-321-B08 | GORG | 0 | 0 | NA | 2 | 0 | orphan | 0 | 0 | NA | 0 | 0 | NA | 52.23 | 0 | 0.83 |
| AG-891-O09 | GORG | 0 | 0 | NA | 1 | 0 | orphan | 0 | 0 | NA | 0 | 2 | orphan | 52.22 | 0 | 1.6 |
| AG-390-N21 | GORG | 0 | 0 | NA | 2 | 0 | orphan | 0 | 0 | NA | 0 | 1 | orphan | 52.2 | 0 | 1.21 |
| AG-916-C04 | GORG | 0 | 0 | NA | 1 | 0 | orphan | 0 | 0 | NA | 0 | 0 | NA | 52.2 | 0 | 0.73 |
| AG-435-I03 | GORG | 0 | 0 | NA | 2 | 0 | orphan | 0 | 0 | NA | 0 | 0 | NA | 52.19 | 0 | 0.59 |
| AG-893-K02 | GORG | 0 | 0 | NA | 1 | 0 | orphan | 0 | 0 | NA | 0 | 0 | NA | 52.19 | 0 | 0.95 |
| AG-422-C09 | GORG | 0 | 0 | NA | 4 | 0 | orphan | 0 | 0 | NA | 0 | 0 | NA | 52.15 | 0 | 1.03 |

| | | | | | | | | | | | | | | | | |
|---|---|---|---|---|---|---|---|---|---|---|---|---|---|---|---|---|
| AG-319-P11 | GORG | 0 | 0 | NA | 1 | 0 | orphan | 0 | 0 | NA | 0 | 0 | NA | 52.1 | 0 | 0.91 |
| AG-914-O04 | GORG | 0 | 0 | NA | 0 | 1 | NA | 0 | 0 | NA | 0 | 0 | NA | 52.06 | 0 | 1.05 |
| AG-337-J10 | GORG | 0 | 0 | NA | 1 | 0 | orphan | 0 | 0 | NA | 0 | 0 | NA | 52.05 | 0 | 0.58 |
| AG-359-B02 | GORG | 0 | 0 | NA | 0 | 0 | NA | 0 | 0 | NA | 0 | 1 | orphan | 52.04 | 0 | 0.89 |
| AG-426-O15 | GORG | 0 | 0 | NA | 1 | 0 | orphan | 0 | 0 | NA | 0 | 0 | NA | 52.04 | 0 | 0.99 |
| AG-390-J19 | GORG | 0 | 0 | NA | 1 | 0 | orphan | 0 | 0 | NA | 0 | 0 | NA | 52 | 0 | 1.25 |
| AG-457-C19 | GORG | 1 | 0 | orphan | 0 | 0 | NA | 0 | 0 | NA | 0 | 0 | NA | 51.95 | 0 | 0.96 |
| AG-343-L18 | GORG | 0 | 0 | NA | 1 | 0 | orphan | 0 | 0 | NA | 0 | 0 | NA | 51.89 | 0 | 0.78 |
| AG-430-P05 | GORG | 0 | 0 | NA | 1 | 0 | orphan | 0 | 0 | NA | 0 | 0 | NA | 51.89 | 0 | 0.88 |
| AG-896-P17 | GORG | 0 | 0 | NA | 1 | 0 | orphan | 0 | 0 | NA | 0 | 0 | NA | 51.89 | 0 | 0.93 |
| AG-904-C18 | GORG | 0 | 0 | NA | 1 | 0 | orphan | 0 | 0 | NA | 0 | 0 | NA | 51.89 | 0 | 0.71 |
| AG-918-M05 | GORG | 0 | 0 | NA | 1 | 0 | orphan | 0 | 1 | NA | 0 | 0 | NA | 51.89 | 0 | 0.63 |
| AG-914-E11 | GORG | 0 | 0 | NA | 1 | 0 | orphan | 0 | 0 | NA | 0 | 0 | NA | 51.88 | 0 | 0.54 |
| AG-912-O17 | GORG | 0 | 0 | NA | 0 | 0 | NA | 0 | 0 | NA | 0 | 1 | orphan | 51.86 | 0 | 0.91 |
| AG-313-C20 | GORG | 0 | 0 | NA | 1 | 0 | orphan | 0 | 0 | NA | 0 | 0 | NA | 51.83 | 0 | 1.04 |
| AG-390-B07 | GORG | 0 | 0 | NA | 2 | 0 | orphan | 0 | 0 | NA | 0 | 0 | NA | 51.81 | 0 | 1.46 |
| AG-891-G08 | GORG | 0 | 0 | NA | 1 | 0 | orphan | 0 | 0 | NA | 0 | 0 | NA | 51.81 | 0 | 0.89 |
| AG-909-O09 | GORG | 0 | 0 | NA | 1 | 0 | orphan | 0 | 0 | NA | 0 | 0 | NA | 51.81 | 0 | 0.85 |
| AG-907-O13 | GORG | 0 | 0 | NA | 1 | 0 | orphan | 0 | 0 | NA | 0 | 0 | NA | 51.79 | 0 | 1.61 |
| AG-333-O18 | GORG | 0 | 0 | NA | 1 | 1 | complete | 0 | 0 | NA | 0 | 0 | NA | 51.78 | 0 | 0.73 |
| AG-916-I02 | GORG | 0 | 0 | NA | 1 | 0 | orphan | 0 | 0 | NA | 0 | 0 | NA | 51.76 | 0 | 0.73 |
| AG-325-O22 | GORG | 0 | 0 | NA | 1 | 0 | orphan | 0 | 0 | NA | 0 | 0 | NA | 51.74 | 0 | 0.71 |
| AG-343-P21 | GORG | 0 | 0 | NA | 1 | 0 | orphan | 0 | 0 | NA | 0 | 0 | NA | 51.73 | 0 | 0.9 |
| AG-903-E23 | GORG | 0 | 0 | NA | 1 | 0 | orphan | 0 | 0 | NA | 0 | 1 | orphan | 51.72 | 0 | 0.85 |
| AG-404-I19 | GORG | 0 | 0 | NA | 1 | 0 | orphan | 0 | 0 | NA | 0 | 0 | NA | 51.72 | 0 | 1.43 |
| AG-430-N15 | GORG | 0 | 1 | NA | 1 | 0 | orphan | 0 | 0 | NA | 0 | 0 | NA | 51.72 | 0 | 1.08 |

| | | | | | | | | | | | | | | | | |
|---|---|---|---|---|---|---|---|---|---|---|---|---|---|---|---|---|
| AG-894-M05 | GORG | 0 | 0 | NA | 1 | 0 | orphan | 0 | 0 | NA | 0 | 0 | NA | 51.72 | 0 | 1.36 |
| AG-900-G21 | GORG | 1 | 1 | complete | 1 | 0 | orphan | 0 | 0 | NA | 0 | 0 | NA | 51.72 | 0 | 0.68 |
| AG-910-K20 | GORG | 0 | 0 | NA | 1 | 0 | orphan | 0 | 0 | NA | 0 | 0 | NA | 51.72 | 0 | 0.77 |
| AH-324-L07 | GORG | 0 | 0 | NA | 1 | 0 | orphan | 0 | 0 | NA | 0 | 0 | NA | 51.72 | 0 | 0.69 |
| AG-896-C22 | GORG | 0 | 0 | NA | 1 | 0 | orphan | 0 | 0 | NA | 0 | 0 | NA | 51.7 | 0 | 0.65 |
| AG-915-K04 | GORG | 2 | 1 | complete | 0 | 0 | NA | 0 | 0 | NA | 0 | 0 | NA | 51.64 | 0 | 1.29 |
| AG-453-L23 | GORG | 0 | 0 | NA | 1 | 0 | orphan | 0 | 0 | NA | 0 | 0 | NA | 51.61 | 0 | 0.6 |
| AG-896-B07 | GORG | 0 | 0 | NA | 1 | 0 | orphan | 0 | 0 | NA | 0 | 0 | NA | 51.56 | 0 | 0.79 |
| AG-914-C17 | GORG | 0 | 0 | NA | 1 | 0 | orphan | 0 | 0 | NA | 0 | 0 | NA | 51.56 | 0 | 0.78 |
| AG-426-A17 | GORG | 0 | 0 | NA | 1 | 0 | orphan | 0 | 0 | NA | 0 | 0 | NA | 51.54 | 0 | 1.03 |
| AG-464-M02 | GORG | 0 | 0 | NA | 4 | 0 | orphan | 0 | 0 | NA | 0 | 0 | NA | 51.54 | 0 | 1.21 |
| AG-915-D07 | GORG | 0 | 0 | NA | 1 | 0 | orphan | 0 | 0 | NA | 0 | 0 | NA | 51.54 | 0 | 0.82 |
| AG-404-E02 | GORG | 0 | 0 | NA | 2 | 0 | orphan | 0 | 0 | NA | 0 | 0 | NA | 51.52 | 0 | 0.77 |
| AG-899-D15 | GORG | 0 | 0 | NA | 3 | 0 | orphan | 0 | 0 | NA | 0 | 0 | NA | 51.51 | 0 | 0.67 |
| AG-319-K08 | GORG | 0 | 0 | NA | 3 | 0 | orphan | 0 | 0 | NA | 0 | 0 | NA | 51.48 | 0 | 1.09 |
| AH-321-E05 | GORG | 0 | 0 | NA | 0 | 0 | NA | 0 | 0 | NA | 0 | 1 | orphan | 51.46 | 0 | 0.66 |
| AG-908-O07 | GORG | 0 | 0 | NA | 1 | 0 | orphan | 0 | 0 | NA | 0 | 0 | NA | 51.45 | 0 | 0.98 |
| AG-916-I05 | GORG | 0 | 0 | NA | 1 | 0 | orphan | 0 | 0 | NA | 0 | 1 | orphan | 51.4 | 0 | 0.66 |
| AG-390-J18 | GORG | 0 | 0 | NA | 2 | 0 | orphan | 0 | 0 | NA | 0 | 0 | NA | 51.4 | 0 | 1.05 |
| AG-901-J04 | GORG | 0 | 0 | NA | 1 | 0 | orphan | 0 | 0 | NA | 0 | 1 | orphan | 51.36 | 0 | 0.85 |
| AG-422-P11 | GORG | 1 | 1 | complete | 0 | 0 | NA | 0 | 0 | NA | 0 | 0 | NA | 51.36 | 0 | 0.78 |
| AG-915-D09 | GORG | 0 | 0 | NA | 2 | 0 | orphan | 0 | 0 | NA | 0 | 0 | NA | 51.35 | 0 | 0.84 |
| AG-898-A23 | GORG | 0 | 0 | NA | 2 | 0 | orphan | 0 | 0 | NA | 0 | 0 | NA | 51.31 | 0 | 1.13 |
| AG-333-N13 | GORG | 0 | 0 | NA | 2 | 0 | orphan | 0 | 0 | NA | 0 | 0 | NA | 51.29 | 0 | 0.53 |
| AG-892-L02 | GORG | 0 | 0 | NA | 1 | 0 | orphan | 0 | 0 | NA | 0 | 1 | orphan | 51.27 | 0 | 0.87 |

| | | | | | | | | | | | | | | | | |
|---|---|---|---|---|---|---|---|---|---|---|---|---|---|---|---|---|
| AG-896-A21 | GORG | 0 | 0 | NA | 1 | 0 | orphan | 0 | 0 | NA | 0 | 0 | NA | 51.26 | 0 | 0.76 |
| AG-894-L03 | GORG | 0 | 0 | NA | 1 | 0 | orphan | 0 | 0 | NA | 0 | 0 | NA | 51.24 | 0 | 1.08 |
| AG-908-O13 | GORG | 0 | 0 | NA | 2 | 0 | orphan | 0 | 0 | NA | 0 | 0 | NA | 51.24 | 0 | 0.67 |
| AG-447-G17 | GORG | 0 | 0 | NA | 0 | 0 | NA | 0 | 0 | NA | 0 | 1 | orpha n | 51.23 | 0 | 0.99 |
| AG-893-P19 | GORG | 0 | 0 | NA | 1 | 0 | orphan | 0 | 0 | NA | 0 | 0 | NA | 51.14 | 0 | 1 |
| AG-359-J03 | GORG | 0 | 0 | NA | 3 | 0 | orphan | 0 | 0 | NA | 0 | 0 | NA | 51.11 | 0 | 0.62 |
| AG-905-A08 | GORG | 0 | 0 | NA | 1 | 0 | orphan | 0 | 0 | NA | 0 | 0 | NA | 51.11 | 0 | 0.76 |
| AG-343-O03 | GORG | 0 | 0 | NA | 2 | 0 | orphan | 0 | 0 | NA | 0 | 2 | orpha n | 51.08 | 0 | 1.15 |
| AG-447-J19 | GORG | 0 | 0 | NA | 1 | 0 | orphan | 0 | 0 | NA | 0 | 0 | NA | 51.06 | 0 | 0.65 |
| AG-904-B18 | GORG | 0 | 0 | NA | 1 | 0 | orphan | 0 | 0 | NA | 0 | 0 | NA | 51.04 | 0 | 0.53 |
| AG-333-F08 | GORG | 0 | 0 | NA | 1 | 0 | orphan | 0 | 0 | NA | 0 | 0 | NA | 50.97 | 0 | 0.56 |
| AG-896-E04 | GORG | 0 | 0 | NA | 1 | 0 | orphan | 0 | 0 | NA | 0 | 0 | NA | 50.96 | 0 | 0.77 |
| AG-349-A23 | GORG | 0 | 0 | NA | 0 | 0 | NA | 0 | 0 | NA | 0 | 1 | orpha n | 50.95 | 0 | 0.92 |
| AG-439-M22 | GORG | 0 | 0 | NA | 1 | 0 | orphan | 0 | 0 | NA | 0 | 0 | NA | 50.95 | 0 | 0.76 |
| AG-333-C17 | GORG | 0 | 0 | NA | 2 | 0 | orphan | 0 | 0 | NA | 0 | 0 | NA | 50.94 | 0 | 0.79 |
| AG-333-O20 | GORG | 0 | 0 | NA | 1 | 0 | orphan | 0 | 0 | NA | 0 | 0 | NA | 50.94 | 0 | 0.53 |
| AG-359-L11 | GORG | 0 | 0 | NA | 1 | 0 | orphan | 0 | 0 | NA | 0 | 0 | NA | 50.94 | 0 | 0.48 |
| AG-457-P20 | GORG | 0 | 0 | NA | 1 | 0 | orphan | 0 | 0 | NA | 0 | 0 | NA | 50.94 | 0 | 0.76 |
| AG-464-D20 | GORG | 0 | 0 | NA | 1 | 0 | orphan | 0 | 0 | NA | 0 | 0 | NA | 50.94 | 0 | 0.57 |
| AG-470-M17 | GORG | 0 | 0 | NA | 1 | 0 | orphan | 0 | 0 | NA | 0 | 0 | NA | 50.94 | 0 | 1.08 |
| AG-898-F08 | GORG | 0 | 0 | NA | 1 | 0 | orphan | 0 | 0 | NA | 0 | 0 | NA | 50.94 | 0 | 0.55 |
| AG-901-D21 | GORG | 1 | 0 | orphan | 1 | 0 | orphan | 0 | 0 | NA | 0 | 0 | NA | 50.94 | 0 | 0.88 |
| AG-903-G04 | GORG | 0 | 0 | NA | 1 | 0 | orphan | 0 | 0 | NA | 0 | 0 | NA | 50.94 | 0 | 0.68 |
| AG-908-J02 | GORG | 0 | 0 | NA | 1 | 0 | orphan | 0 | 0 | NA | 0 | 0 | NA | 50.94 | 0 | 0.68 |
| AG-909-C13 | GORG | 0 | 0 | NA | 1 | 0 | orphan | 0 | 0 | NA | 0 | 0 | NA | 50.94 | 0 | 0.62 |
| AG-915-D22 | GORG | 0 | 0 | NA | 1 | 0 | orphan | 0 | 0 | NA | 0 | 0 | NA | 50.94 | 0 | 0.55 |

| | | | | | | | | | | | | | | | | |
|---|---|---|---|---|---|---|---|---|---|---|---|---|---|---|---|---|
| AH-287-P13 | GORG | 0 | 0 | NA | 1 | 0 | orphan | 0 | 0 | NA | 0 | 0 | NA | 50.94 | 0 | 0.79 |
| AG-414-A11 | GORG | 0 | 0 | NA | 2 | 1 | complet e | 0 | 0 | NA | 0 | 0 | NA | 50.88 | 0 | 0.74 |
| AH-326-A17 | GORG | 0 | 0 | NA | 2 | 0 | orphan | 0 | 0 | NA | 0 | 0 | NA | 50.88 | 0 | 0.98 |
| AG-901-D06 | GORG | 0 | 1 | NA | 0 | 0 | NA | 0 | 0 | NA | 0 | 0 | NA | 50.83 | 0 | 0.93 |
| AG-919-C08 | GORG | 0 | 0 | NA | 0 | 0 | NA | 0 | 0 | NA | 0 | 1 | orpha n | 50.77 | 0 | 0.87 |
| AG-907-B22 | GORG | 0 | 0 | NA | 1 | 0 | orphan | 0 | 0 | NA | 0 | 0 | NA | 50.77 | 0 | 0.51 |
| AG-920-B04 | GORG | 0 | 0 | NA | 2 | 0 | orphan | 0 | 0 | NA | 0 | 0 | NA | 50.74 | 0 | 1.24 |
| AG-919-G08 | GORG | 0 | 0 | NA | 1 | 0 | orphan | 0 | 0 | NA | 0 | 1 | orpha n | 50.68 | 0 | 1.82 |
| AG-910-P17 | GORG | 0 | 0 | NA | 1 | 0 | orphan | 0 | 0 | NA | 0 | 0 | NA | 50.66 | 0 | 0.72 |
| AG-426-G14 | GORG | 0 | 0 | NA | 1 | 1 | complet e | 0 | 0 | NA | 0 | 0 | NA | 50.65 | 0 | 0.74 |
| AG-894-B10 | GORG | 0 | 1 | NA | 0 | 0 | NA | 0 | 0 | NA | 0 | 0 | NA | 50.65 | 0 | 1.57 |
| AG-349-F15 | GORG | 0 | 0 | NA | 1 | 0 | orphan | 0 | 0 | NA | 0 | 0 | NA | 50.6 | 0 | 0.59 |
| AG-892-P08 | GORG | 0 | 0 | NA | 1 | 0 | orphan | 0 | 0 | NA | 0 | 0 | NA | 50.6 | 0 | 0.56 |
| AG-909-N17 | GORG | 0 | 0 | NA | 1 | 0 | orphan | 0 | 0 | NA | 0 | 0 | NA | 50.6 | 0 | 0.71 |
| AG-915-C20 | GORG | 0 | 0 | NA | 1 | 0 | orphan | 0 | 0 | NA | 0 | 0 | NA | 50.6 | 0 | 0.81 |
| AG-917-I20 | GORG | 0 | 0 | NA | 1 | 0 | orphan | 0 | 0 | NA | 0 | 0 | NA | 50.6 | 0 | 0.61 |
| AG-920-J11 | GORG | 0 | 0 | NA | 1 | 1 | complet e | 0 | 0 | NA | 0 | 0 | NA | 50.6 | 0 | 0.72 |
| AG-404-K22 | GORG | 1 | 1 | complet e | 0 | 0 | NA | 0 | 0 | NA | 0 | 0 | NA | 50.54 | 0 | 1.36 |
| AG-390-B21 | GORG | 1 | 1 | complet e | 1 | 0 | orphan | 0 | 0 | NA | 0 | 0 | NA | 50.47 | 0 | 1.18 |
| AG-435-A07 | GORG | 0 | 0 | NA | 1 | 0 | orphan | 0 | 0 | NA | 0 | 0 | NA | 50.41 | 0 | 0.99 |
| AG-333-N14 | GORG | 0 | 0 | NA | 1 | 0 | orphan | 0 | 0 | NA | 0 | 0 | NA | 50.4 | 0 | 1.17 |
| AG-337-C21 | GORG | 0 | 0 | NA | 1 | 0 | orphan | 0 | 0 | NA | 0 | 0 | NA | 50.39 | 0 | 0.55 |
| AG-918-K22 | GORG | 0 | 0 | NA | 1 | 0 | orphan | 0 | 0 | NA | 0 | 0 | NA | 50.39 | 0 | 0.54 |
| AG-898-I14 | GORG | 0 | 0 | NA | 1 | 0 | orphan | 0 | 0 | NA | 0 | 0 | NA | 50.36 | 0 | 0.71 |
| AG-895-K06 | GORG | 1 | 1 | complet | 0 | 0 | NA | 0 | 0 | NA | 0 | 0 | NA | 50.32 | 0 | 0.77 |

| | | | | | | | | | | | | | | | | |
|---|---|---|---|---|---|---|---|---|---|---|---|---|---|---|---|---|
| | | | | e | | | | | | | | | | | | |
| AG-319-F16 | GORG | 0 | 0 | NA | 0 | 0 | NA | 0 | 0 | NA | 0 | 1 | orphan | 50.31 | 0 | 1.22 |
| AG-907-A04 | GORG | 0 | 0 | NA | 1 | 0 | orphan | 0 | 1 | NA | 0 | 0 | NA | 50.31 | 0 | 0.72 |
| AG-435-C17 | GORG | 1 | 1 | complete | 2 | 0 | orphan | 0 | 0 | NA | 0 | 0 | NA | 50.3 | 0 | 0.75 |
| AG-325-J04 | GORG | 0 | 0 | NA | 1 | 0 | orphan | 0 | 0 | NA | 0 | 0 | NA | 50.29 | 0 | 0.69 |
| AG-914-K17 | GORG | 1 | 1 | complete | 0 | 0 | NA | 0 | 0 | NA | 0 | 0 | NA | 50.29 | 0 | 0.87 |
| AG-917-P01 | GORG | 0 | 0 | NA | 1 | 0 | orphan | 0 | 0 | NA | 0 | 1 | orphan | 50.27 | 0 | 0.88 |
| AG-457-J09 | GORG | 0 | 0 | NA | 1 | 0 | orphan | 0 | 0 | NA | 0 | 0 | NA | 50.26 | 0 | 0.65 |
| AG-912-G19 | GORG | 0 | 0 | NA | 2 | 0 | orphan | 0 | 0 | NA | 0 | 0 | NA | 50.24 | 0 | 0.57 |
| AH-287-I17 | GORG | 0 | 1 | NA | 0 | 0 | NA | 0 | 0 | NA | 0 | 0 | NA | 50.23 | 0 | 0.97 |
| AG-900-P04 | GORG | 1 | 1 | complete | 1 | 0 | orphan | 0 | 0 | NA | 0 | 0 | NA | 50.2 | 0 | 0.77 |
| AG-325-D19 | GORG | 0 | 1 | NA | 1 | 0 | orphan | 0 | 0 | NA | 0 | 0 | NA | 50.19 | 0 | 0.58 |
| AG-920-B10 | GORG | 0 | 0 | NA | 1 | 0 | orphan | 0 | 0 | NA | 0 | 0 | NA | 50.15 | 0 | 0.58 |
| AG-900-L21 | GORG | 0 | 0 | NA | 0 | 0 | NA | 0 | 0 | NA | 0 | 1 | orphan | 50.14 | 0 | 0.63 |
| AG-333-C03 | GORG | 0 | 0 | NA | 2 | 0 | orphan | 0 | 0 | NA | 0 | 0 | NA | 50.13 | 0 | 0.61 |
| AG-325-A20 | GORG | 1 | 1 | complete | 0 | 0 | NA | 0 | 0 | NA | 0 | 0 | NA | 50.04 | 0 | 0.94 |
| AG-464-G04 | GORG | 0 | 0 | NA | 1 | 0 | orphan | 0 | 0 | NA | 0 | 0 | NA | 50.02 | 0 | 0.62 |
| AG-897-D08 | GORG | 0 | 0 | NA | 1 | 0 | orphan | 0 | 0 | NA | 0 | 0 | NA | 65.04 | 0.01 | 1.19 |
| AG-899-E03 | GORG | 0 | 0 | NA | 1 | 0 | orphan | 0 | 0 | NA | 0 | 0 | NA | 63.81 | 0.01 | 1.21 |
| AG-337-G03 | GORG | 0 | 0 | NA | 3 | 1 | complete | 0 | 0 | NA | 0 | 0 | NA | 60.86 | 0.01 | 1.04 |
| AG-390-O20 | GORG | 0 | 0 | NA | 1 | 0 | orphan | 0 | 0 | NA | 0 | 0 | NA | 66.52 | 0.02 | 0.91 |
| AG-899-M13 | GORG | 1 | 1 | complete | 0 | 0 | NA | 0 | 0 | NA | 0 | 1 | orphan | 89.79 | 0.03 | 1.47 |
| AG-917-F02 | GORG | 0 | 0 | NA | 1 | 0 | orphan | 0 | 0 | NA | 0 | 0 | NA | 81.81 | 0.03 | 1.22 |
| AG-904-M20 | GORG | 0 | 0 | NA | 1 | 0 | orphan | 0 | 0 | NA | 0 | 0 | NA | 77.77 | 0.03 | 2.19 |

| | | | | | | | | | | | | | | | | |
|---|---|---|---|---|---|---|---|---|---|---|---|---|---|---|---|---|
| AG-337-P05 | GORG | 1 | 1 | complete | 0 | 0 | NA | 0 | 0 | NA | 0 | 0 | NA | 69.43 | 0.03 | 0.91 |
| AG-319-E21 | GORG | 0 | 0 | NA | 1 | 0 | orphan | 0 | 0 | NA | 0 | 0 | NA | 60.39 | 0.03 | 1.11 |
| AG-914-B14 | GORG | 0 | 0 | NA | 1 | 0 | orphan | 0 | 0 | NA | 0 | 0 | NA | 90.03 | 0.04 | 1.2 |
| AG-891-L02 | GORG | 0 | 0 | NA | 1 | 0 | orphan | 0 | 0 | NA | 0 | 0 | NA | 89.39 | 0.04 | 0.97 |
| AG-910-C18 | GORG | 0 | 0 | NA | 2 | 0 | orphan | 0 | 0 | NA | 0 | 0 | NA | 79.52 | 0.04 | 0.98 |
| AG-901-G20 | GORG | 0 | 0 | NA | 2 | 0 | orphan | 0 | 0 | NA | 0 | 0 | NA | 76.94 | 0.04 | 0.95 |
| AG-325-C07 | GORG | 0 | 0 | NA | 1 | 0 | orphan | 0 | 0 | NA | 0 | 0 | NA | 76.57 | 0.04 | 0.87 |
| AG-896-G05 | GORG | 0 | 0 | NA | 1 | 0 | orphan | 0 | 0 | NA | 0 | 0 | NA | 73.68 | 0.04 | 0.92 |
| AG-909-J19 | GORG | 1 | 1 | complete | 1 | 0 | orphan | 0 | 0 | NA | 0 | 0 | NA | 73.28 | 0.04 | 0.8 |
| AH-326-L19 | GORG | 0 | 0 | NA | 2 | 0 | orphan | 0 | 0 | NA | 0 | 0 | NA | 69.15 | 0.04 | 1.9 |
| AG-916-K23 | GORG | 0 | 0 | NA | 2 | 0 | orphan | 0 | 1 | NA | 0 | 0 | NA | 66.13 | 0.04 | 0.81 |
| AG-917-J07 | GORG | 0 | 0 | NA | 2 | 0 | orphan | 0 | 0 | NA | 0 | 0 | NA | 65.79 | 0.04 | 0.77 |
| AG-461-P07 | GORG | 0 | 0 | NA | 1 | 0 | orphan | 0 | 0 | NA | 0 | 0 | NA | 63.68 | 0.04 | 1.81 |
| AG-907-G02 | GORG | 0 | 0 | NA | 1 | 0 | orphan | 0 | 0 | NA | 0 | 0 | NA | 63.68 | 0.04 | 0.74 |
| AG-464-K08 | GORG | 0 | 0 | NA | 0 | 0 | NA | 0 | 0 | NA | 0 | 1 | orphan | 58.84 | 0.04 | 1.02 |
| AH-324-A20 | GORG | 0 | 0 | NA | 1 | 0 | orphan | 0 | 0 | NA | 0 | 0 | NA | 55.03 | 0.04 | 0.67 |
| AG-899-K02 | GORG | 0 | 0 | NA | 1 | 0 | orphan | 0 | 0 | NA | 0 | 0 | NA | 84.96 | 0.05 | 2.08 |
| AG-390-B14 | GORG | 1 | 1 | complete | 1 | 0 | orphan | 0 | 0 | NA | 0 | 0 | NA | 68.62 | 0.05 | 1.76 |
| AG-390-A18 | GORG | 0 | 0 | NA | 2 | 0 | orphan | 0 | 0 | NA | 0 | 0 | NA | 67.67 | 0.05 | 1.85 |
| AH-324-A10 | GORG | 0 | 0 | NA | 1 | 0 | orphan | 0 | 0 | NA | 0 | 1 | orphan | 55.43 | 0.05 | 1.13 |
| AG-422-J09 | GORG | 0 | 0 | NA | 2 | 0 | orphan | 0 | 0 | NA | 0 | 0 | NA | 53.87 | 0.05 | 1.53 |
| AG-439-P13 | GORG | 0 | 0 | NA | 1 | 0 | orphan | 0 | 0 | NA | 0 | 0 | NA | 66.79 | 0.06 | 2.11 |
| AG-915-J13 | GORG | 0 | 0 | NA | 3 | 0 | orphan | 0 | 0 | NA | 0 | 0 | NA | 61.9 | 0.06 | 1.17 |
| AG-337-L06 | GORG | 1 | 1 | complete | 0 | 0 | NA | 0 | 0 | NA | 0 | 0 | NA | 56.67 | 0.06 | 1.24 |
| AG-457-F09 | GORG | 0 | 0 | NA | 1 | 0 | orphan | 0 | 0 | NA | 0 | 0 | NA | 53.69 | 0.06 | 1.38 |

| | | | | | | | | | | | | | | | | |
|---|---|---|---|---|---|---|---|---|---|---|---|---|---|---|---|---|
| AG-916-E19 | GORG | 0 | 0 | NA | 1 | 0 | orphan | 0 | 0 | NA | 0 | 0 | NA | 90.15 | 0.07 | 1.25 |
| AG-333-L11 | GORG | 0 | 0 | NA | 2 | 1 | complete | 0 | 0 | NA | 0 | 0 | NA | 87.35 | 0.07 | 1.13 |
| AG-447-O04 | GORG | 0 | 1 | NA | 3 | 0 | orphan | 0 | 0 | NA | 0 | 0 | NA | 64.4 | 0.07 | 1.21 |
| AG-426-M08 | GORG | 0 | 0 | NA | 2 | 0 | orphan | 0 | 0 | NA | 0 | 0 | NA | 62.73 | 0.07 | 0.69 |
| AH-321-O14 | GORG | 0 | 0 | NA | 1 | 0 | orphan | 0 | 0 | NA | 0 | 0 | NA | 62.69 | 0.07 | 1.22 |
| AG-313-P16 | GORG | 0 | 0 | NA | 1 | 0 | orphan | 0 | 0 | NA | 0 | 0 | NA | 62.43 | 0.07 | 1.39 |
| AH-287-G19 | GORG | 0 | 0 | NA | 1 | 0 | orphan | 0 | 0 | NA | 0 | 0 | NA | 61.23 | 0.07 | 0.99 |
| AG-430-E15 | GORG | 0 | 0 | NA | 1 | 0 | orphan | 0 | 0 | NA | 0 | 0 | NA | 59.53 | 0.07 | 1.04 |
| AG-920-P09 | GORG | 0 | 0 | NA | 1 | 0 | orphan | 0 | 0 | NA | 0 | 0 | NA | 56.88 | 0.07 | 1.14 |
| AG-325-P10 | GORG | 1 | 1 | complete | 0 | 0 | NA | 0 | 0 | NA | 0 | 0 | NA | 55.52 | 0.07 | 0.98 |
| AG-903-A17 | GORG | 0 | 0 | NA | 1 | 0 | orphan | 0 | 0 | NA | 0 | 0 | NA | 54.94 | 0.07 | 0.7 |
| AG-325-E16 | GORG | 0 | 0 | NA | 2 | 0 | orphan | 0 | 0 | NA | 0 | 0 | NA | 54.5 | 0.07 | 1.15 |
| AG-365-D17 | GORG | 1 | 1 | complete | 1 | 0 | orphan | 0 | 0 | NA | 0 | 0 | NA | 54 | 0.07 | 0.92 |
| AG-404-N17 | GORG | 0 | 0 | NA | 3 | 0 | orphan | 0 | 0 | NA | 0 | 0 | NA | 52.75 | 0.07 | 0.99 |
| AG-349-I20 | GORG | 0 | 0 | NA | 1 | 0 | orphan | 0 | 0 | NA | 0 | 0 | NA | 75.09 | 0.08 | 1.56 |
| AG-313-B13 | GORG | 0 | 0 | NA | 1 | 0 | orphan | 0 | 0 | NA | 0 | 0 | NA | 56.71 | 0.08 | 1.15 |
| AG-892-M16 | GORG | 1 | 1 | complete | 0 | 0 | NA | 0 | 0 | NA | 0 | 0 | NA | 56.13 | 0.08 | 1.19 |
| AG-914-L10 | GORG | 0 | 0 | NA | 2 | 0 | orphan | 0 | 0 | NA | 0 | 0 | NA | 85.15 | 0.09 | 0.94 |
| AG-895-K23 | GORG | 0 | 0 | NA | 3 | 2 | complete | 0 | 0 | NA | 0 | 0 | NA | 64.81 | 0.09 | 1.77 |
| AG-470-P20 | GORG | 0 | 0 | NA | 2 | 0 | orphan | 0 | 0 | NA | 0 | 0 | NA | 51.58 | 0.09 | 2.36 |
| AG-898-J17 | GORG | 0 | 0 | NA | 1 | 0 | orphan | 0 | 0 | NA | 0 | 0 | NA | 81.49 | 0.1 | 2.34 |
| AG-426-E20 | GORG | 0 | 0 | NA | 1 | 0 | orphan | 0 | 0 | NA | 0 | 0 | NA | 72.23 | 0.1 | 0.8 |
| AG-319-A04 | GORG | 0 | 1 | NA | 0 | 0 | NA | 0 | 0 | NA | 0 | 0 | NA | 55.57 | 0.1 | 0.61 |
| AH-287-F15 | GORG | 0 | 0 | NA | 2 | 0 | orphan | 0 | 0 | NA | 0 | 0 | NA | 98.36 | 0.11 | 1.32 |
| AG-911-I16 | GORG | 0 | 0 | NA | 1 | 0 | orphan | 0 | 0 | NA | 0 | 0 | NA | 87.84 | 0.11 | 1.07 |

| | | | | | | | | | | | | | | | | |
|---|---|---|---|---|---|---|---|---|---|---|---|---|---|---|---|---|
| AG-916-A21 | GORG | 0 | 0 | NA | 1 | 0 | orphan | 0 | 0 | NA | 0 | 0 | NA | 84.94 | 0.11 | 1.04 |
| AG-404-P13 | GORG | 0 | 0 | NA | 2 | 0 | orphan | 0 | 0 | NA | 0 | 0 | NA | 83.46 | 0.11 | 1.17 |
| AG-359-J18 | GORG | 0 | 0 | NA | 1 | 0 | orphan | 0 | 0 | NA | 0 | 0 | NA | 82.2 | 0.11 | 1.15 |
| AG-907-C09 | GORG | 0 | 0 | NA | 1 | 0 | orphan | 0 | 0 | NA | 0 | 0 | NA | 81.93 | 0.11 | 1.21 |
| AG-891-E06 | GORG | 0 | 0 | NA | 2 | 0 | orphan | 0 | 0 | NA | 0 | 0 | NA | 81.33 | 0.11 | 1.01 |
| AG-893-G20 | GORG | 0 | 0 | NA | 1 | 0 | orphan | 0 | 0 | NA | 0 | 0 | NA | 80.42 | 0.11 | 1.38 |
| AG-333-P08 | GORG | 1 | 0 | orphan | 1 | 0 | orphan | 0 | 0 | NA | 0 | 0 | NA | 80.12 | 0.11 | 0.92 |
| AG-426-D13 | GORG | 0 | 0 | NA | 2 | 0 | orphan | 0 | 0 | NA | 0 | 0 | NA | 79.52 | 0.11 | 1.11 |
| AG-457-I22 | GORG | 0 | 0 | NA | 1 | 0 | orphan | 0 | 0 | NA | 0 | 0 | NA | 79.38 | 0.11 | 1.04 |
| AG-903-G09 | GORG | 0 | 0 | NA | 1 | 0 | orphan | 0 | 0 | NA | 0 | 0 | NA | 79.38 | 0.11 | 0.72 |
| AG-901-M15 | GORG | 1 | 1 | complete | 2 | 0 | orphan | 0 | 0 | NA | 0 | 0 | NA | 78.31 | 0.11 | 1.31 |
| AG-337-K21 | GORG | 0 | 0 | NA | 1 | 0 | orphan | 0 | 0 | NA | 0 | 0 | NA | 77.71 | 0.11 | 1.09 |
| AG-912-J04 | GORG | 0 | 0 | NA | 1 | 0 | orphan | 0 | 0 | NA | 0 | 0 | NA | 77.38 | 0.11 | 1.2 |
| AG-891-P17 | GORG | 1 | 1 | complete | 1 | 0 | orphan | 0 | 0 | NA | 0 | 0 | NA | 77.11 | 0.11 | 1.23 |
| AG-912-O15 | GORG | 0 | 0 | NA | 1 | 0 | orphan | 0 | 0 | NA | 0 | 0 | NA | 75.3 | 0.11 | 1.17 |
| AG-464-I13 | GORG | 0 | 0 | NA | 1 | 1 | complete | 0 | 0 | NA | 0 | 0 | NA | 74.7 | 0.11 | 0.93 |
| AG-337-O03 | GORG | 0 | 0 | NA | 1 | 0 | orphan | 0 | 0 | NA | 0 | 0 | NA | 72.54 | 0.11 | 0.79 |
| AG-430-J08 | GORG | 0 | 0 | NA | 2 | 0 | orphan | 0 | 0 | NA | 0 | 0 | NA | 72.03 | 0.11 | 0.81 |
| AG-422-J08 | GORG | 1 | 1 | orphan | 2 | 0 | orphan | 0 | 0 | NA | 0 | 0 | NA | 71.03 | 0.11 | 0.98 |
| AG-908-G17 | GORG | 0 | 0 | NA | 1 | 0 | orphan | 0 | 0 | NA | 0 | 0 | NA | 69.7 | 0.11 | 0.97 |
| AG-894-D03 | GORG | 0 | 0 | NA | 1 | 0 | orphan | 0 | 0 | NA | 0 | 0 | NA | 69.46 | 0.11 | 1.16 |
| AG-894-K13 | GORG | 0 | 0 | NA | 1 | 0 | orphan | 0 | 0 | NA | 0 | 0 | NA | 69.46 | 0.11 | 1.01 |
| AG-337-J23 | GORG | 0 | 0 | NA | 1 | 0 | orphan | 0 | 0 | NA | 0 | 0 | NA | 69.28 | 0.11 | 1.04 |
| AG-390-B06 | GORG | 0 | 0 | NA | 1 | 0 | orphan | 0 | 0 | NA | 0 | 0 | NA | 68.67 | 0.11 | 0.68 |
| AG-899-E09 | GORG | 0 | 0 | NA | 1 | 0 | orphan | 0 | 0 | NA | 0 | 0 | NA | 68.62 | 0.11 | 0.84 |
| AG-365-K21 | GORG | 0 | 0 | NA | 1 | 0 | orphan | 0 | 0 | NA | 0 | 0 | NA | 68.46 | 0.11 | 0.82 |

| | | | | | | | | | | | | | | | | |
|---|---|---|---|---|---|---|---|---|---|---|---|---|---|---|---|---|
| AG-904-C06 | GORG | 0 | 0 | NA | 2 | 0 | orphan | 0 | 0 | NA | 0 | 0 | NA | 66.98 | 0.11 | 0.95 |
| AG-917-N04 | GORG | 0 | 0 | NA | 1 | 0 | orphan | 0 | 1 | NA | 0 | 0 | NA | 65.71 | 0.11 | 0.94 |
| AG-896-M21 | GORG | 0 | 0 | NA | 1 | 0 | orphan | 0 | 0 | NA | 0 | 0 | NA | 65.06 | 0.11 | 0.97 |
| AG-901-P17 | GORG | 0 | 0 | NA | 1 | 0 | orphan | 0 | 0 | NA | 0 | 0 | NA | 63.86 | 0.11 | 0.9 |
| AG-359-M18 | GORG | 0 | 0 | NA | 1 | 0 | orphan | 0 | 0 | NA | 0 | 0 | NA | 62.65 | 0.11 | 0.57 |
| AG-439-L02 | GORG | 0 | 0 | NA | 2 | 0 | orphan | 0 | 0 | NA | 0 | 0 | NA | 61.42 | 0.11 | 0.93 |
| AG-905-M07 | GORG | 0 | 0 | NA | 2 | 0 | orphan | 0 | 0 | NA | 0 | 0 | NA | 57.61 | 0.11 | 0.69 |
| AG-910-K03 | GORG | 0 | 0 | NA | 1 | 0 | orphan | 0 | 0 | NA | 0 | 0 | NA | 56.34 | 0.11 | 1.07 |
| AG-891-F02 | GORG | 0 | 1 | NA | 1 | 1 | complet e | 0 | 0 | NA | 0 | 0 | NA | 56.32 | 0.11 | 1.01 |
| AG-910-O15 | GORG | 0 | 0 | NA | 1 | 0 | orphan | 0 | 0 | NA | 0 | 0 | NA | 51.26 | 0.11 | 0.93 |
| AG-464-M18 | GORG | 0 | 0 | NA | 2 | 0 | orphan | 0 | 0 | NA | 0 | 0 | NA | 57.82 | 0.12 | 1.25 |
| AG-426-A10 | GORG | 1 | 0 | orphan | 0 | 0 | NA | 0 | 0 | NA | 0 | 1 | orpha n | 53.45 | 0.12 | 2.39 |
| AG-916-G15 | GORG | 0 | 0 | NA | 1 | 0 | orphan | 0 | 0 | NA | 0 | 0 | NA | 51.6 | 0.12 | 0.88 |
| AG-414-O15 | GORG | 0 | 0 | NA | 0 | 1 | NA | 0 | 0 | NA | 0 | 0 | NA | 76.61 | 0.13 | 2.16 |
| AG-359-L21 | GORG | 0 | 0 | NA | 1 | 0 | orphan | 0 | 0 | NA | 0 | 0 | NA | 75.23 | 0.13 | 1.73 |
| AG-916-E11 | GORG | 0 | 0 | NA | 2 | 0 | orphan | 0 | 0 | NA | 0 | 0 | NA | 72.13 | 0.13 | 1.83 |
| AG-915-E13 | GORG | 0 | 0 | NA | 1 | 0 | orphan | 0 | 0 | NA | 0 | 0 | NA | 70.74 | 0.13 | 1.97 |
| AG-894-B17 | GORG | 0 | 0 | NA | 2 | 1 | complet e | 0 | 0 | NA | 0 | 0 | NA | 69.04 | 0.13 | 1.82 |
| AG-319-E10 | GORG | 0 | 0 | NA | 0 | 0 | NA | 0 | 0 | NA | 0 | 1 | orpha n | 56.47 | 0.13 | 1 |
| AG-894-L05 | GORG | 0 | 0 | NA | 1 | 0 | orphan | 0 | 0 | NA | 0 | 0 | NA | 55.39 | 0.13 | 1.62 |
| AG-319-O07 | GORG | 1 | 1 | orphan | 0 | 0 | NA | 0 | 0 | NA | 0 | 0 | NA | 54.62 | 0.13 | 1.14 |
| AG-891-P07 | GORG | 0 | 0 | NA | 3 | 0 | orphan | 0 | 0 | NA | 0 | 0 | NA | 52.64 | 0.13 | 1.09 |
| AG-893-P05 | GORG | 0 | 0 | NA | 3 | 0 | orphan | 0 | 0 | NA | 0 | 0 | NA | 87.84 | 0.14 | 2.08 |
| AG-893-G05 | GORG | 0 | 0 | NA | 1 | 0 | orphan | 0 | 0 | NA | 0 | 0 | NA | 85.64 | 0.14 | 2.11 |
| AG-892-O16 | GORG | 0 | 0 | NA | 3 | 0 | orphan | 0 | 0 | NA | 0 | 0 | NA | 84.3 | 0.14 | 1.98 |
| AG-915-E09 | GORG | 1 | 1 | complet | 1 | 0 | orphan | 0 | 1 | NA | 0 | 0 | NA | 79.15 | 0.14 | 1.81 |

| | | | | | | | | | | | | | | | | |
|---|---|---|---|---|---|---|---|---|---|---|---|---|---|---|---|---|
| | | | | e | | | | | | | | | | | | |
| AG-920-I22 | GORG | 0 | 0 | NA | 2 | 0 | orphan | 0 | 1 | NA | 0 | 0 | NA | 78.89 | 0.14 | 1.52 |
| AG-915-B22 | GORG | 0 | 0 | NA | 2 | 0 | orphan | 0 | 1 | NA | 0 | 0 | NA | 78.5 | 0.14 | 1.72 |
| AG-918-I11 | GORG | 0 | 0 | NA | 3 | 0 | orphan | 0 | 0 | NA | 0 | 0 | NA | 74.31 | 0.14 | 1.76 |
| AG-911-B18 | GORG | 0 | 0 | NA | 1 | 0 | orphan | 0 | 0 | NA | 0 | 0 | NA | 73.1 | 0.14 | 1.58 |
| AG-919-I05 | GORG | 1 | 1 | complete | 2 | 0 | orphan | 0 | 0 | NA | 0 | 0 | NA | 72.61 | 0.14 | 1.66 |
| AG-426-K10 | GORG | 0 | 0 | NA | 0 | 0 | NA | 0 | 0 | NA | 0 | 1 | orphan | 71.06 | 0.14 | 1.16 |
| AG-359-O10 | GORG | 0 | 0 | NA | 2 | 0 | orphan | 0 | 0 | NA | 0 | 0 | NA | 70.36 | 0.14 | 1.9 |
| AH-326-I13 | GORG | 0 | 0 | NA | 1 | 0 | orphan | 0 | 0 | NA | 0 | 0 | NA | 70.33 | 0.14 | 1.57 |
| AG-903-P05 | GORG | 0 | 0 | NA | 2 | 0 | orphan | 0 | 0 | NA | 0 | 0 | NA | 69.39 | 0.14 | 1.88 |
| AG-908-P09 | GORG | 0 | 0 | NA | 3 | 0 | orphan | 0 | 0 | NA | 0 | 0 | NA | 68.58 | 0.14 | 2.38 |
| AG-337-O15 | GORG | 3 | 2 | complete | 0 | 0 | NA | 0 | 0 | NA | 0 | 0 | NA | 66.81 | 0.14 | 1.72 |
| AG-349-K03 | GORG | 0 | 0 | NA | 0 | 0 | NA | 0 | 0 | NA | 0 | 1 | orphan | 66.58 | 0.14 | 1.07 |
| AG-913-G13 | GORG | 0 | 0 | NA | 2 | 1 | orphan | 0 | 0 | NA | 0 | 0 | NA | 66.09 | 0.14 | 1.18 |
| AG-895-P14 | GORG | 0 | 0 | NA | 1 | 0 | orphan | 0 | 0 | NA | 0 | 0 | NA | 66.04 | 0.14 | 1.53 |
| AG-911-O05 | GORG | 0 | 0 | NA | 2 | 0 | orphan | 0 | 0 | NA | 0 | 0 | NA | 64.95 | 0.14 | 1.35 |
| AG-903-E20 | GORG | 0 | 0 | NA | 2 | 0 | orphan | 0 | 0 | NA | 0 | 0 | NA | 64.94 | 0.14 | 1.6 |
| AH-321-J15 | GORG | 2 | 1 | complete | 0 | 0 | NA | 0 | 0 | NA | 0 | 1 | orphan | 63.53 | 0.14 | 1.08 |
| AG-313-B09 | GORG | 0 | 0 | NA | 1 | 0 | orphan | 0 | 0 | NA | 0 | 0 | NA | 63.39 | 0.14 | 1.88 |
| AG-404-N23 | GORG | 0 | 0 | NA | 1 | 0 | orphan | 0 | 0 | NA | 0 | 0 | NA | 61.25 | 0.14 | 1.25 |
| AG-907-E16 | GORG | 0 | 0 | NA | 1 | 0 | orphan | 0 | 0 | NA | 0 | 0 | NA | 59.52 | 0.14 | 1.34 |
| AG-904-C08 | GORG | 0 | 0 | NA | 0 | 0 | NA | 0 | 0 | NA | 0 | 1 | orphan | 56.32 | 0.14 | 0.83 |
| AG-349-B03 | GORG | 0 | 0 | NA | 1 | 0 | orphan | 0 | 0 | NA | 0 | 0 | NA | 53.99 | 0.14 | 1.48 |
| AG-912-E19 | GORG | 0 | 0 | NA | 0 | 0 | NA | 0 | 0 | NA | 0 | 1 | orphan | 51.63 | 0.14 | 0.86 |
| AG-422-I13 | GORG | 0 | 0 | NA | 0 | 0 | NA | 0 | 0 | NA | 0 | 1 | orpha | 50.71 | 0.14 | 0.75 |

| | | | | | | | | | | | | | | | | |
|---|---|---|---|---|---|---|---|---|---|---|---|---|---|---|---|---|
| | | | | | | | | | | | | | n | | | |
| AG-422-D16 | GORG | 0 | 0 | NA | 0 | 0 | NA | 0 | 0 | NA | 0 | 1 | orphan | 77.18 | 0.15 | 1.28 |
| AG-913-B07 | GORG | 0 | 0 | NA | 1 | 0 | orphan | 0 | 0 | NA | 0 | 0 | NA | 72.05 | 0.15 | 1.87 |
| AG-920-D05 | GORG | 0 | 0 | NA | 1 | 0 | orphan | 0 | 0 | NA | 0 | 0 | NA | 96.67 | 0.16 | 1.23 |
| AG-919-O02 | GORG | 0 | 0 | NA | 1 | 0 | orphan | 0 | 0 | NA | 0 | 0 | NA | 90.46 | 0.16 | 1.03 |
| AG-920-L03 | GORG | 0 | 0 | NA | 1 | 0 | orphan | 0 | 0 | NA | 0 | 0 | NA | 85.06 | 0.16 | 0.97 |
| AG-325-L05 | GORG | 0 | 0 | NA | 1 | 0 | orphan | 0 | 0 | NA | 0 | 0 | NA | 81.17 | 0.16 | 1.04 |
| AG-916-K04 | GORG | 0 | 0 | NA | 2 | 0 | orphan | 0 | 0 | NA | 0 | 0 | NA | 80.49 | 0.16 | 0.96 |
| AG-337-C08 | GORG | 0 | 0 | NA | 1 | 0 | orphan | 0 | 0 | NA | 0 | 0 | NA | 79.03 | 0.16 | 0.92 |
| AG-891-A17 | GORG | 0 | 0 | NA | 1 | 0 | orphan | 0 | 0 | NA | 0 | 0 | NA | 76.23 | 0.16 | 0.88 |
| AH-287-E18 | GORG | 0 | 0 | NA | 2 | 0 | orphan | 0 | 0 | NA | 0 | 0 | NA | 74.76 | 0.16 | 0.92 |
| AG-920-M22 | GORG | 0 | 0 | NA | 1 | 0 | orphan | 0 | 0 | NA | 0 | 0 | NA | 68.81 | 0.16 | 0.81 |
| AH-287-E14 | GORG | 0 | 0 | NA | 2 | 0 | orphan | 0 | 0 | NA | 0 | 0 | NA | 67.25 | 0.16 | 1.55 |
| AG-908-D18 | GORG | 0 | 0 | NA | 1 | 0 | orphan | 0 | 0 | NA | 0 | 0 | NA | 67.22 | 0.16 | 0.88 |
| AG-915-L05 | GORG | 0 | 0 | NA | 2 | 0 | orphan | 0 | 0 | NA | 0 | 0 | NA | 67.06 | 0.16 | 0.84 |
| AG-909-O06 | GORG | 0 | 0 | NA | 1 | 0 | orphan | 0 | 0 | NA | 0 | 0 | NA | 66.71 | 0.16 | 0.83 |
| AG-426-D22 | GORG | 0 | 0 | NA | 0 | 0 | NA | 0 | 0 | NA | 0 | 1 | orphan | 61.18 | 0.16 | 0.81 |
| AG-360-A16 | GORG | 1 | 1 | complete | 1 | 0 | orphan | 0 | 0 | NA | 0 | 0 | NA | 58.42 | 0.16 | 0.67 |
| AG-360-N15 | GORG | 0 | 0 | NA | 2 | 0 | orphan | 0 | 0 | NA | 0 | 0 | NA | 57.88 | 0.16 | 0.67 |
| AH-321-D14 | GORG | 0 | 0 | NA | 1 | 0 | orphan | 0 | 0 | NA | 0 | 0 | NA | 76.67 | 0.17 | 1.49 |
| AG-920-I17 | GORG | 0 | 0 | NA | 1 | 0 | orphan | 0 | 0 | NA | 0 | 0 | NA | 74.52 | 0.17 | 1.53 |
| AG-359-I22 | GORG | 0 | 0 | NA | 1 | 0 | orphan | 0 | 0 | NA | 0 | 0 | NA | 67.37 | 0.17 | 1.27 |
| AG-907-I16 | GORG | 1 | 1 | complete | 0 | 0 | NA | 0 | 0 | NA | 0 | 0 | NA | 66.54 | 0.17 | 1.29 |
| AG-899-P13 | GORG | 1 | 1 | complete | 0 | 0 | NA | 0 | 0 | NA | 0 | 0 | NA | 63.65 | 0.17 | 1.15 |
| AH-287-K10 | GORG | 0 | 0 | NA | 2 | 0 | orphan | 0 | 0 | NA | 0 | 0 | NA | 60.88 | 0.17 | 1.12 |

| | | | | | | | | | | | | | | | | |
|---|---|---|---|---|---|---|---|---|---|---|---|---|---|---|---|---|
| AG-891-P18 | GORG | 1 | 1 | complet e | 0 | 0 | NA | 0 | 0 | NA | 0 | 0 | NA | 59.97 | 0.17 | 1.12 |
| AG-390-M20 | GORG | 1 | 1 | complet e | 0 | 0 | NA | 0 | 0 | NA | 0 | 0 | NA | 59.8 | 0.17 | 1.17 |
| AG-313-A15 | GORG | 0 | 0 | NA | 1 | 0 | orphan | 0 | 0 | NA | 0 | 0 | NA | 56.95 | 0.17 | 1.08 |
| AG-430-N11 | GORG | 0 | 1 | NA | 2 | 0 | orphan | 0 | 0 | NA | 0 | 0 | NA | 70.33 | 0.18 | 0.8 |
| AG-911-M18 | GORG | 0 | 0 | NA | 3 | 0 | orphan | 0 | 0 | NA | 0 | 0 | NA | 66.9 | 0.18 | 1.65 |
| AG-914-N16 | GORG | 0 | 1 | NA | 0 | 0 | NA | 0 | 0 | NA | 0 | 0 | NA | 78.35 | 0.19 | 1.58 |
| AG-343-D11 | GORG | 0 | 0 | NA | 4 | 0 | orphan | 0 | 0 | NA | 0 | 0 | NA | 67.53 | 0.19 | 0.87 |
| AG-905-L09 | GORG | 0 | 0 | NA | 2 | 0 | orphan | 0 | 0 | NA | 0 | 0 | NA | 63.06 | 0.19 | 0.83 |
| AG-325-I23 | GORG | 0 | 0 | NA | 2 | 1 | orphan | 0 | 0 | NA | 0 | 2 | orpha n | 59.67 | 0.19 | 1.15 |
| AG-898-G08 | GORG | 0 | 0 | NA | 1 | 0 | orphan | 0 | 0 | NA | 0 | 0 | NA | 59.2 | 0.19 | 0.79 |
| AG-343-G10 | GORG | 0 | 0 | NA | 3 | 0 | orphan | 0 | 1 | NA | 0 | 0 | NA | 56.01 | 0.19 | 1.11 |
| AG-899-J15 | GORG | 0 | 0 | NA | 1 | 0 | orphan | 0 | 0 | NA | 0 | 0 | NA | 74.23 | 0.2 | 2.16 |
| AG-912-I02 | GORG | 0 | 0 | NA | 2 | 0 | orphan | 0 | 0 | NA | 0 | 0 | NA | 92.77 | 0.22 | 1.33 |
| AH-287-K18 | GORG | 0 | 0 | NA | 1 | 0 | orphan | 0 | 0 | NA | 0 | 0 | NA | 76.51 | 0.22 | 0.75 |
| AG-918-A03 | GORG | 0 | 0 | NA | 1 | 0 | orphan | 0 | 0 | NA | 0 | 0 | NA | 75.39 | 0.22 | 1.16 |
| AG-898-O17 | GORG | 0 | 0 | NA | 1 | 0 | orphan | 0 | 0 | NA | 0 | 0 | NA | 71.55 | 0.22 | 1.02 |
| AH-287-A08 | GORG | 0 | 0 | NA | 1 | 0 | orphan | 0 | 0 | NA | 0 | 0 | NA | 67.65 | 0.22 | 1.39 |
| AH-326-N07 | GORG | 1 | 1 | complet e | 0 | 0 | NA | 0 | 0 | NA | 0 | 0 | NA | 50.41 | 0.22 | 0.92 |
| AG-908-B16 | GORG | 1 | 1 | complet e | 0 | 0 | NA | 0 | 0 | NA | 0 | 1 | orpha n | 63 | 0.23 | 2.02 |
| AG-896-N20 | GORG | 0 | 0 | NA | 1 | 0 | orphan | 0 | 0 | NA | 0 | 0 | NA | 55.17 | 0.23 | 1.28 |
| AG-390-G23 | GORG | 0 | 0 | NA | 1 | 0 | orphan | 0 | 0 | NA | 0 | 0 | NA | 92.89 | 0.24 | 1.27 |
| AG-891-A05 | GORG | 0 | 0 | NA | 1 | 0 | orphan | 0 | 0 | NA | 0 | 1 | orpha n | 89.73 | 0.24 | 1.22 |
| AH-287-E09 | GORG | 1 | 1 | complet e | 1 | 0 | orphan | 0 | 0 | NA | 0 | 0 | NA | 86.33 | 0.24 | 1.15 |
| AG-390-L16 | GORG | 0 | 0 | NA | 1 | 0 | orphan | 0 | 0 | NA | 0 | 0 | NA | 85.19 | 0.24 | 1.09 |
| AG-916-J21 | GORG | 0 | 0 | NA | 1 | 0 | orphan | 0 | 0 | NA | 0 | 1 | orpha | 83.65 | 0.24 | 1.04 |

| | | | | | | | | | | | | | | | | |
|---|---|---|---|---|---|---|---|---|---|---|---|---|---|---|---|---|
| | | | | | | | | | | | | | n | | | |
| AH-287-E08 | GORG | 1 | 1 | complete | 0 | 0 | NA | 0 | 0 | NA | 0 | 0 | NA | 81.75 | 0.24 | 1.02 |
| AG-899-E08 | GORG | 0 | 0 | NA | 2 | 1 | complete | 0 | 0 | NA | 0 | 0 | NA | 81.12 | 0.24 | 1.06 |
| AG-892-C08 | GORG | 0 | 0 | NA | 1 | 0 | orphan | 0 | 0 | NA | 0 | 0 | NA | 79.76 | 0.24 | 0.95 |
| AG-899-P11 | GORG | 0 | 0 | NA | 1 | 0 | orphan | 0 | 1 | NA | 0 | 0 | NA | 78.44 | 0.24 | 1 |
| AG-390-N22 | GORG | 0 | 0 | NA | 2 | 0 | orphan | 0 | 0 | NA | 0 | 0 | NA | 77.48 | 0.24 | 1.18 |
| AG-426-A16 | GORG | 0 | 0 | NA | 2 | 0 | orphan | 0 | 0 | NA | 0 | 0 | NA | 76.98 | 0.24 | 0.88 |
| AG-390-A20 | GORG | 0 | 0 | NA | 1 | 0 | orphan | 0 | 0 | NA | 0 | 0 | NA | 76.75 | 0.24 | 0.88 |
| AG-910-B10 | GORG | 0 | 0 | NA | 2 | 0 | orphan | 0 | 0 | NA | 0 | 0 | NA | 75.87 | 0.24 | 0.91 |
| AG-911-B05 | GORG | 0 | 0 | NA | 1 | 0 | orphan | 0 | 0 | NA | 0 | 0 | NA | 75.43 | 0.24 | 0.93 |
| AG-447-P13 | GORG | 0 | 0 | NA | 2 | 0 | orphan | 0 | 0 | NA | 0 | 0 | NA | 75.12 | 0.24 | 1.09 |
| AG-414-K18 | GORG | 0 | 0 | NA | 2 | 0 | orphan | 0 | 0 | NA | 0 | 0 | NA | 74.68 | 0.24 | 0.9 |
| AG-426-M15 | GORG | 0 | 0 | NA | 2 | 0 | orphan | 0 | 0 | NA | 0 | 0 | NA | 73.85 | 0.24 | 0.96 |
| AH-324-K04 | GORG | 0 | 0 | NA | 1 | 0 | orphan | 0 | 0 | NA | 0 | 1 | orphan | 68.73 | 0.24 | 0.83 |
| AG-404-L19 | GORG | 0 | 0 | NA | 2 | 0 | orphan | 0 | 0 | NA | 0 | 0 | NA | 67.98 | 0.24 | 0.88 |
| AG-422-K22 | GORG | 0 | 0 | NA | 1 | 0 | orphan | 0 | 0 | NA | 0 | 0 | NA | 67.04 | 0.24 | 0.95 |
| AG-919-P18 | GORG | 0 | 0 | NA | 1 | 0 | orphan | 0 | 0 | NA | 0 | 0 | NA | 66.9 | 0.24 | 0.8 |
| AG-461-E23 | GORG | 0 | 0 | NA | 1 | 0 | orphan | 0 | 0 | NA | 0 | 0 | NA | 66.28 | 0.24 | 0.71 |
| AG-900-I09 | GORG | 0 | 0 | NA | 1 | 0 | orphan | 0 | 0 | NA | 0 | 0 | NA | 65.99 | 0.24 | 0.84 |
| AG-447-E02 | GORG | 0 | 0 | NA | 2 | 0 | orphan | 0 | 0 | NA | 0 | 0 | NA | 65 | 0.24 | 0.94 |
| AG-447-A22 | GORG | 0 | 0 | NA | 1 | 0 | orphan | 0 | 0 | NA | 0 | 0 | NA | 61.93 | 0.24 | 0.78 |
| AG-333-I10 | GORG | 1 | 1 | complete | 1 | 0 | orphan | 0 | 0 | NA | 0 | 0 | NA | 61.89 | 0.24 | 0.8 |
| AH-321-I22 | GORG | 0 | 0 | NA | 1 | 0 | orphan | 0 | 0 | NA | 0 | 1 | orphan | 56 | 0.24 | 0.67 |
| AH-287-O20 | GORG | 0 | 0 | NA | 1 | 0 | orphan | 0 | 0 | NA | 0 | 0 | NA | 55.4 | 0.24 | 0.73 |
| AG-390-C20 | GORG | 0 | 0 | NA | 1 | 0 | orphan | 0 | 0 | NA | 0 | 0 | NA | 54.93 | 0.24 | 0.84 |

| | | | | | | | | | | | | | | | | |
|---|---|---|---|---|---|---|---|---|---|---|---|---|---|---|---|---|
| AG-900-C18 | GORG | 0 | 0 | NA | 1 | 0 | orphan | 0 | 0 | NA | 0 | 0 | NA | 53.21 | 0.24 | 0.72 |
| AG-359-L15 | GORG | 0 | 0 | NA | 1 | 0 | orphan | 0 | 0 | NA | 0 | 0 | NA | 51.27 | 0.25 | 2.17 |
| AG-915-O02 | GORG | 1 | 1 | complete | 0 | 0 | NA | 0 | 0 | NA | 0 | 1 | orphan | 93.39 | 0.27 | 1.58 |
| AG-894-I06 | GORG | 0 | 0 | NA | 0 | 0 | NA | 0 | 0 | NA | 0 | 1 | orphan | 92.92 | 0.27 | 1.49 |
| AG-404-D14 | GORG | 0 | 0 | NA | 0 | 0 | NA | 0 | 0 | NA | 0 | 1 | orphan | 90.14 | 0.27 | 1.57 |
| AG-913-N17 | GORG | 0 | 0 | NA | 0 | 0 | NA | 0 | 0 | NA | 0 | 2 | orphan | 86.32 | 0.27 | 1.39 |
| AG-896-A03 | GORG | 0 | 0 | NA | 0 | 0 | NA | 0 | 0 | NA | 0 | 1 | orphan | 85.37 | 0.27 | 1.41 |
| AG-894-G22 | GORG | 1 | 0 | orphan | 0 | 0 | NA | 0 | 0 | NA | 0 | 2 | orphan | 83.65 | 0.27 | 1.36 |
| AG-911-F16 | GORG | 0 | 0 | NA | 1 | 0 | orphan | 0 | 0 | NA | 0 | 0 | NA | 81.78 | 0.27 | 1.96 |
| AG-917-E11 | GORG | 0 | 0 | NA | 1 | 0 | orphan | 0 | 0 | NA | 0 | 0 | NA | 81.75 | 0.27 | 1.27 |
| AG-894-M14 | GORG | 0 | 0 | NA | 1 | 0 | orphan | 0 | 0 | NA | 0 | 1 | orphan | 78.67 | 0.27 | 1.27 |
| AG-898-A19 | GORG | 0 | 0 | NA | 0 | 0 | NA | 0 | 0 | NA | 0 | 1 | orphan | 77.43 | 0.27 | 1.23 |
| AG-435-M14 | GORG | 0 | 0 | NA | 1 | 0 | orphan | 0 | 0 | NA | 0 | 0 | NA | 77.13 | 0.27 | 1.13 |
| AG-907-P07 | GORG | 0 | 0 | NA | 1 | 0 | orphan | 0 | 0 | NA | 0 | 0 | NA | 77.11 | 0.27 | 1.43 |
| AG-920-C05 | GORG | 0 | 0 | NA | 0 | 1 | NA | 0 | 0 | NA | 0 | 0 | NA | 74.77 | 0.27 | 1.16 |
| AG-917-D20 | GORG | 0 | 0 | NA | 1 | 0 | orphan | 0 | 0 | NA | 0 | 1 | orphan | 74.59 | 0.27 | 1.18 |
| AH-287-M08 | GORG | 0 | 0 | NA | 2 | 0 | orphan | 1 | 2 | complete | 0 | 1 | orphan | 74.46 | 0.27 | 1.19 |
| AG-414-C16 | GORG | 0 | 0 | NA | 4 | 1 | orphan | 0 | 0 | NA | 0 | 1 | orphan | 73.84 | 0.27 | 1.92 |
| AG-895-O22 | GORG | 1 | 1 | complete | 0 | 0 | NA | 0 | 0 | NA | 0 | 0 | NA | 71.47 | 0.27 | 1.3 |
| AG-909-M08 | GORG | 0 | 0 | NA | 0 | 0 | NA | 0 | 0 | NA | 0 | 1 | orphan | 71.2 | 0.27 | 1.13 |
| AG-919-B02 | GORG | 0 | 0 | NA | 1 | 0 | orphan | 0 | 0 | NA | 0 | 1 | orphan | 70.2 | 0.27 | 1.17 |
| AG-349-B16 | GORG | 0 | 0 | NA | 1 | 0 | orphan | 0 | 0 | NA | 0 | 0 | NA | 66.53 | 0.27 | 0.98 |

| | | | | | | | | | | | | | | | | |
|---|---|---|---|---|---|---|---|---|---|---|---|---|---|---|---|---|
| AG-916-B09 | GORG | 0 | 0 | NA | 1 | 0 | orphan | 0 | 0 | NA | 0 | 2 | orphan | 66.12 | 0.27 | 1.17 |
| AG-900-L19 | GORG | 0 | 0 | NA | 0 | 0 | NA | 0 | 0 | NA | 0 | 1 | orphan | 65.67 | 0.27 | 1.06 |
| AG-422-I06 | GORG | 1 | 1 | complete | 0 | 0 | NA | 0 | 0 | NA | 0 | 1 | orphan | 63.86 | 0.27 | 0.98 |
| AG-899-F05 | GORG | 0 | 0 | NA | 0 | 0 | NA | 0 | 0 | NA | 0 | 1 | orphan | 63.25 | 0.27 | 1.05 |
| AG-919-L15 | GORG | 0 | 0 | NA | 3 | 0 | orphan | 0 | 0 | NA | 0 | 1 | orphan | 63.18 | 0.27 | 0.92 |
| AG-899-B22 | GORG | 0 | 0 | NA | 1 | 0 | orphan | 0 | 0 | NA | 0 | 1 | orphan | 63.05 | 0.27 | 0.96 |
| AG-918-A18 | GORG | 0 | 0 | NA | 2 | 0 | orphan | 0 | 0 | NA | 0 | 0 | NA | 62.64 | 0.27 | 1.07 |
| AG-908-A22 | GORG | 0 | 0 | NA | 1 | 0 | orphan | 0 | 0 | NA | 0 | 1 | orphan | 62.18 | 0.27 | 0.93 |
| AG-900-I06 | GORG | 0 | 0 | NA | 1 | 0 | orphan | 0 | 0 | NA | 0 | 1 | orphan | 60.73 | 0.27 | 1.04 |
| AG-453-F21 | GORG | 0 | 0 | NA | 1 | 0 | orphan | 0 | 0 | NA | 0 | 0 | NA | 59.94 | 0.27 | 1.02 |
| AG-911-F20 | GORG | 0 | 0 | NA | 1 | 0 | orphan | 0 | 0 | NA | 0 | 0 | NA | 59.83 | 0.27 | 0.92 |
| AG-916-O09 | GORG | 0 | 0 | NA | 2 | 0 | orphan | 0 | 0 | NA | 0 | 1 | orphan | 59.69 | 0.27 | 1.02 |
| AG-909-M16 | GORG | 0 | 0 | NA | 1 | 0 | orphan | 0 | 0 | NA | 0 | 0 | NA | 59.28 | 0.27 | 0.92 |
| AG-891-L11 | GORG | 0 | 0 | NA | 0 | 0 | NA | 0 | 0 | NA | 0 | 2 | orphan | 58.56 | 0.27 | 1.03 |
| AG-359-F16 | GORG | 0 | 0 | NA | 0 | 0 | NA | 0 | 0 | NA | 0 | 1 | orphan | 55.48 | 0.27 | 0.95 |
| AH-326-D22 | GORG | 0 | 0 | NA | 1 | 0 | orphan | 0 | 0 | NA | 0 | 1 | orphan | 54.95 | 0.27 | 0.93 |
| AG-422-K18 | GORG | 0 | 0 | NA | 0 | 0 | NA | 0 | 0 | NA | 0 | 1 | orphan | 51.95 | 0.27 | 0.84 |
| AG-914-N20 | GORG | 0 | 0 | NA | 1 | 1 | complete | 0 | 0 | NA | 0 | 1 | orphan | 51.31 | 0.27 | 0.83 |
| AG-359-P11 | GORG | 0 | 0 | NA | 1 | 0 | orphan | 0 | 0 | NA | 0 | 1 | orphan | 59.72 | 0.28 | 0.75 |
| AH-321-C05 | GORG | 0 | 0 | NA | 2 | 0 | orphan | 0 | 0 | NA | 0 | 0 | NA | 72.46 | 0.29 | 1.09 |
| AG-907-K13 | GORG | 0 | 0 | NA | 3 | 0 | orphan | 0 | 1 | NA | 0 | 0 | NA | 63.79 | 0.29 | 0.95 |
| AG-349-D03 | GORG | 0 | 0 | NA | 1 | 0 | orphan | 0 | 0 | NA | 0 | 0 | NA | 53.48 | 0.29 | 0.68 |

| | | | | | | | | | | | | | | | | |
|---|---|---|---|---|---|---|---|---|---|---|---|---|---|---|---|---|
| AG-899-C06 | GORG | 0 | 0 | NA | 1 | 0 | orphan | 0 | 0 | NA | 0 | 0 | NA | 75.27 | 0.3 | 1.99 |
| AG-457-F15 | GORG | 1 | 1 | complete | 1 | 0 | orphan | 0 | 0 | NA | 0 | 0 | NA | 71.64 | 0.3 | 1.85 |
| AG-447-G11 | GORG | 0 | 0 | NA | 1 | 0 | orphan | 0 | 0 | NA | 0 | 1 | orphan | 67.26 | 0.3 | 1.94 |
| AG-447-P15 | GORG | 0 | 0 | NA | 2 | 0 | orphan | 0 | 0 | NA | 0 | 0 | NA | 65.9 | 0.3 | 1.6 |
| AG-895-P05 | GORG | 0 | 0 | NA | 2 | 0 | orphan | 0 | 0 | NA | 0 | 0 | NA | 64.92 | 0.3 | 1.64 |
| AH-321-I09 | GORG | 1 | 0 | orphan | 1 | 0 | orphan | 0 | 0 | NA | 0 | 0 | NA | 64.66 | 0.3 | 1.73 |
| AG-390-A10 | GORG | 0 | 0 | NA | 2 | 0 | orphan | 0 | 0 | NA | 0 | 0 | NA | 62.77 | 0.3 | 1.79 |
| AG-447-E04 | GORG | 0 | 0 | NA | 1 | 0 | orphan | 0 | 0 | NA | 0 | 0 | NA | 59 | 0.3 | 1.61 |
| AG-313-J17 | GORG | 0 | 0 | NA | 1 | 0 | orphan | 0 | 0 | NA | 0 | 0 | NA | 58.85 | 0.3 | 1.64 |
| AG-919-D14 | GORG | 0 | 0 | NA | 1 | 0 | orphan | 0 | 0 | NA | 0 | 0 | NA | 54.57 | 0.3 | 1.39 |
| AH-324-N03 | GORG | 0 | 0 | NA | 1 | 0 | orphan | 0 | 0 | NA | 0 | 0 | NA | 93.43 | 0.32 | 1.2 |
| AG-899-M07 | GORG | 0 | 0 | NA | 1 | 0 | orphan | 0 | 0 | NA | 0 | 0 | NA | 93.1 | 0.32 | 1.16 |
| AG-894-I19 | GORG | 0 | 0 | NA | 3 | 0 | orphan | 0 | 0 | NA | 0 | 0 | NA | 87.46 | 0.32 | 1.01 |
| AG-343-M11 | GORG | 0 | 0 | NA | 1 | 0 | orphan | 0 | 0 | NA | 0 | 0 | NA | 76.7 | 0.32 | 0.97 |
| AG-422-L23 | GORG | 0 | 0 | NA | 6 | 0 | orphan | 0 | 0 | NA | 0 | 2 | orphan | 55.65 | 0.32 | 2.02 |
| AG-909-P15 | GORG | 0 | 0 | NA | 2 | 0 | orphan | 0 | 0 | NA | 0 | 0 | NA | 75.63 | 0.33 | 1.93 |
| AG-313-J10 | GORG | 0 | 0 | NA | 1 | 0 | orphan | 0 | 0 | NA | 0 | 0 | NA | 73.66 | 0.33 | 1.5 |
| AG-913-F17 | GORG | 0 | 0 | NA | 0 | 0 | NA | 1 | 0 | orphan | 0 | 0 | NA | 69.87 | 0.33 | 1.35 |
| AG-447-P01 | GORG | 0 | 0 | NA | 1 | 0 | orphan | 0 | 0 | NA | 0 | 0 | NA | 69.26 | 0.33 | 1.06 |
| AG-333-D18 | GORG | 0 | 1 | NA | 1 | 0 | orphan | 0 | 0 | NA | 0 | 0 | NA | 63.35 | 0.33 | 1.24 |
| AG-904-E23 | GORG | 1 | 1 | complete | 0 | 0 | NA | 0 | 0 | NA | 0 | 0 | NA | 61.24 | 0.33 | 1.1 |
| AG-313-I17 | GORG | 0 | 0 | NA | 1 | 1 | complete | 0 | 0 | NA | 0 | 0 | NA | 59.06 | 0.33 | 1.16 |
| AG-894-P10 | GORG | 0 | 0 | NA | 1 | 0 | orphan | 0 | 0 | NA | 0 | 0 | NA | 73.63 | 0.35 | 0.97 |
| AG-917-D08 | GORG | 0 | 0 | NA | 2 | 0 | orphan | 0 | 0 | NA | 0 | 0 | NA | 73.19 | 0.35 | 1.87 |
| AG-390-G08 | GORG | 0 | 0 | NA | 2 | 0 | orphan | 0 | 0 | NA | 0 | 0 | NA | 73 | 0.35 | 2.13 |

| | | | | | | | | | | | | | | | | |
|---|---|---|---|---|---|---|---|---|---|---|---|---|---|---|---|---|
| AG-914-M11 | GORG | 0 | 0 | NA | 1 | 0 | orphan | 0 | 0 | NA | 0 | 0 | NA | 68.34 | 0.35 | 0.89 |
| AH-324-A07 | GORG | 0 | 0 | NA | 1 | 0 | orphan | 0 | 0 | NA | 0 | 0 | NA | 66.41 | 0.35 | 0.91 |
| AG-912-K02 | GORG | 0 | 1 | NA | 0 | 0 | NA | 0 | 0 | NA | 0 | 0 | NA | 65.46 | 0.35 | 1.24 |
| AG-319-F21 | GORG | 0 | 0 | NA | 1 | 1 | complete | 0 | 0 | NA | 0 | 0 | NA | 61.05 | 0.35 | 1.21 |
| AG-349-K14 | GORG | 1 | 1 | complete | 0 | 0 | NA | 0 | 0 | NA | 0 | 0 | NA | 74.97 | 0.37 | 1.66 |
| AG-898-J13 | GORG | 0 | 0 | NA | 1 | 0 | orphan | 0 | 0 | NA | 0 | 0 | NA | 72.82 | 0.37 | 1.77 |
| AG-893-C16 | GORG | 0 | 0 | NA | 1 | 1 | complete | 0 | 0 | NA | 0 | 0 | NA | 72.17 | 0.37 | 1.51 |
| AG-903-K11 | GORG | 1 | 2 | complete | 1 | 0 | orphan | 0 | 0 | NA | 0 | 0 | NA | 70.44 | 0.37 | 1.47 |
| AG-359-J04 | GORG | 0 | 0 | NA | 1 | 1 | complete | 0 | 0 | NA | 0 | 0 | NA | 66.26 | 0.37 | 1.23 |
| AG-903-L08 | GORG | 0 | 0 | NA | 2 | 0 | orphan | 0 | 0 | NA | 0 | 0 | NA | 66.23 | 0.37 | 1.11 |
| AG-893-A01 | GORG | 0 | 0 | NA | 1 | 0 | orphan | 0 | 1 | NA | 0 | 0 | NA | 66.21 | 0.37 | 1.21 |
| AG-897-E21 | GORG | 0 | 0 | NA | 1 | 0 | orphan | 0 | 0 | NA | 0 | 0 | NA | 65.84 | 0.37 | 1.2 |
| AG-390-C16 | GORG | 0 | 0 | NA | 1 | 0 | orphan | 0 | 0 | NA | 0 | 0 | NA | 65.39 | 0.37 | 1.2 |
| AG-893-E22 | GORG | 1 | 1 | complete | 0 | 0 | NA | 0 | 0 | NA | 0 | 0 | NA | 65.16 | 0.37 | 1.21 |
| AG-325-I14 | GORG | 0 | 0 | NA | 1 | 0 | orphan | 0 | 0 | NA | 0 | 0 | NA | 64.32 | 0.37 | 1.35 |
| AG-905-B21 | GORG | 0 | 0 | NA | 0 | 1 | NA | 0 | 0 | NA | 0 | 0 | NA | 63.86 | 0.37 | 1.95 |
| AG-899-J22 | GORG | 0 | 0 | NA | 1 | 0 | orphan | 0 | 0 | NA | 0 | 0 | NA | 63.84 | 0.37 | 1.08 |
| AG-914-M02 | GORG | 0 | 0 | NA | 1 | 0 | orphan | 0 | 0 | NA | 0 | 1 | orphan | 63.76 | 0.37 | 1.11 |
| AG-337-G20 | GORG | 0 | 0 | NA | 1 | 0 | orphan | 0 | 0 | NA | 0 | 0 | NA | 63.64 | 0.37 | 1.18 |
| AG-915-D04 | GORG | 0 | 0 | NA | 1 | 1 | complete | 0 | 0 | NA | 0 | 0 | NA | 63.64 | 0.37 | 1.14 |
| AG-359-K21 | GORG | 0 | 0 | NA | 2 | 2 | complete | 0 | 0 | NA | 0 | 0 | NA | 63.63 | 0.37 | 1.31 |
| AG-390-J09 | GORG | 0 | 0 | NA | 2 | 0 | orphan | 0 | 0 | NA | 0 | 0 | NA | 63.2 | 0.37 | 1.16 |
| AG-891-K16 | GORG | 0 | 0 | NA | 1 | 0 | orphan | 0 | 0 | NA | 0 | 0 | NA | 62.96 | 0.37 | 1.09 |
| AG-895-G19 | GORG | 0 | 0 | NA | 5 | 0 | orphan | 0 | 0 | NA | 0 | 0 | NA | 62.77 | 0.37 | 1.1 |

| | | | | | | | | | | | | | | | | |
|---|---|---|---|---|---|---|---|---|---|---|---|---|---|---|---|---|
| AG-894-B09 | GORG | 0 | 0 | NA | 1 | 0 | orphan | 0 | 0 | NA | 0 | 0 | NA | 62.09 | 0.37 | 1.17 |
| AG-913-B17 | GORG | 0 | 0 | NA | 1 | 0 | orphan | 0 | 0 | NA | 0 | 0 | NA | 61.91 | 0.37 | 1.09 |
| AG-461-D04 | GORG | 0 | 0 | NA | 1 | 0 | orphan | 0 | 0 | NA | 0 | 0 | NA | 61.2 | 0.37 | 1.41 |
| AG-904-C17 | GORG | 0 | 0 | NA | 1 | 0 | orphan | 0 | 0 | NA | 0 | 0 | NA | 60.79 | 0.37 | 1.09 |
| AG-899-B14 | GORG | 0 | 0 | NA | 1 | 0 | orphan | 0 | 0 | NA | 0 | 0 | NA | 60 | 0.37 | 1.34 |
| AG-390-B18 | GORG | 0 | 0 | NA | 1 | 0 | orphan | 0 | 0 | NA | 0 | 0 | NA | 59.95 | 0.37 | 1.36 |
| AG-893-J18 | GORG | 1 | 1 | complete | 4 | 0 | orphan | 0 | 0 | NA | 0 | 1 | orphan | 59.56 | 0.37 | 1.11 |
| AG-893-O10 | GORG | 0 | 0 | NA | 1 | 0 | orphan | 0 | 0 | NA | 0 | 0 | NA | 58.63 | 0.37 | 1.31 |
| AG-920-P13 | GORG | 0 | 0 | NA | 1 | 0 | orphan | 0 | 0 | NA | 0 | 0 | NA | 58.34 | 0.37 | 1 |
| AG-907-E06 | GORG | 0 | 0 | NA | 2 | 1 | complete | 0 | 0 | NA | 0 | 0 | NA | 58.33 | 0.37 | 1.04 |
| AG-893-C13 | GORG | 1 | 0 | orphan | 0 | 0 | NA | 0 | 0 | NA | 0 | 0 | NA | 57.77 | 0.37 | 0.99 |
| AG-414-O08 | GORG | 0 | 0 | NA | 2 | 0 | orphan | 0 | 0 | NA | 0 | 0 | NA | 57.69 | 0.37 | 1.29 |
| AG-891-G15 | GORG | 0 | 0 | NA | 1 | 0 | orphan | 0 | 0 | NA | 0 | 0 | NA | 57.64 | 0.37 | 1.12 |
| AG-915-G15 | GORG | 0 | 0 | NA | 0 | 0 | NA | 1 | 1 | complete | 0 | 0 | NA | 57.57 | 0.37 | 1.01 |
| AG-359-J23 | GORG | 0 | 0 | NA | 1 | 0 | orphan | 0 | 0 | NA | 0 | 0 | NA | 56.89 | 0.37 | 0.96 |
| AG-359-K10 | GORG | 0 | 0 | NA | 2 | 0 | orphan | 0 | 0 | NA | 0 | 0 | NA | 56.55 | 0.37 | 0.97 |
| AG-893-M06 | GORG | 0 | 0 | NA | 1 | 0 | orphan | 0 | 0 | NA | 0 | 0 | NA | 55.77 | 0.37 | 1 |
| AG-903-I05 | GORG | 0 | 0 | NA | 1 | 0 | orphan | 0 | 0 | NA | 0 | 0 | NA | 55.64 | 0.37 | 1.01 |
| AG-894-B20 | GORG | 0 | 0 | NA | 1 | 0 | orphan | 0 | 0 | NA | 0 | 0 | NA | 55.49 | 0.37 | 0.97 |
| AG-920-O09 | GORG | 1 | 1 | complete | 4 | 0 | orphan | 0 | 0 | NA | 0 | 1 | orphan | 55.27 | 0.37 | 1.05 |
| AG-360-N22 | GORG | 0 | 0 | NA | 1 | 0 | orphan | 0 | 0 | NA | 0 | 0 | NA | 55.25 | 0.37 | 1.05 |
| AG-900-C19 | GORG | 0 | 0 | NA | 3 | 0 | orphan | 0 | 0 | NA | 0 | 1 | orphan | 55.15 | 0.37 | 1 |
| AG-349-N21 | GORG | 0 | 0 | NA | 1 | 0 | orphan | 0 | 0 | NA | 0 | 0 | NA | 54.9 | 0.37 | 1 |
| AG-447-F09 | GORG | 0 | 0 | NA | 1 | 0 | orphan | 0 | 0 | NA | 0 | 0 | NA | 54.71 | 0.37 | 0.93 |
| AG-909-O03 | GORG | 1 | 1 | complete | 0 | 0 | NA | 0 | 0 | NA | 0 | 0 | NA | 54.65 | 0.37 | 1.53 |

| | | | | | | | | | | | | | | | |
|---|---|---|---|---|---|---|---|---|---|---|---|---|---|---|---|
| AG-912-D20 | GORG | 0 | 0 | NA | 1 | 0 | orphan | 0 | 0 | NA | 0 | 0 | NA | 54.04 | 0.37 | 0.94 |
| AG-913-J18 | GORG | 0 | 0 | NA | 3 | 0 | orphan | 1 | 1 | complete | 0 | 0 | NA | 53.97 | 0.37 | 0.94 |
| AG-337-E18 | GORG | 0 | 0 | NA | 1 | 0 | orphan | 0 | 0 | NA | 0 | 0 | NA | 53.78 | 0.37 | 0.93 |
| AG-916-P15 | GORG | 0 | 0 | NA | 1 | 0 | orphan | 0 | 0 | NA | 0 | 0 | NA | 52.02 | 0.37 | 0.97 |
| AG-913-P07 | GORG | 0 | 0 | NA | 3 | 0 | orphan | 0 | 0 | NA | 0 | 0 | NA | 51.85 | 0.37 | 0.96 |
| AG-337-J07 | GORG | 0 | 0 | NA | 1 | 0 | orphan | 0 | 0 | NA | 0 | 0 | NA | 51.31 | 0.37 | 0.95 |
| AG-359-O07 | GORG | 0 | 0 | NA | 1 | 0 | orphan | 0 | 0 | NA | 0 | 0 | NA | 50.68 | 0.37 | 1.11 |
| AG-337-D20 | GORG | 0 | 0 | NA | 1 | 0 | orphan | 0 | 0 | NA | 0 | 0 | NA | 50.62 | 0.37 | 0.84 |
| AG-915-O09 | GORG | 0 | 0 | NA | 1 | 0 | orphan | 0 | 0 | NA | 0 | 0 | NA | 60.64 | 0.38 | 1.1 |
| AG-319-A02 | GORG | 0 | 0 | NA | 2 | 0 | orphan | 0 | 0 | NA | 0 | 1 | orphan | 53.55 | 0.39 | 1.16 |
| AG-892-M03 | GORG | 0 | 0 | NA | 1 | 0 | orphan | 0 | 0 | NA | 0 | 0 | NA | 84.7 | 0.4 | 2.28 |
| AG-918-I18 | GORG | 0 | 0 | NA | 1 | 0 | orphan | 0 | 0 | NA | 0 | 0 | NA | 82.49 | 0.4 | 2.04 |
| AG-896-O22 | GORG | 0 | 0 | NA | 1 | 0 | orphan | 0 | 0 | NA | 0 | 0 | NA | 74.57 | 0.4 | 1.77 |
| AG-343-F17 | GORG | 0 | 0 | NA | 4 | 2 | complete | 0 | 0 | NA | 0 | 0 | NA | 72.53 | 0.4 | 2.06 |
| AG-920-E03 | GORG | 1 | 1 | complete | 2 | 1 | complete | 0 | 0 | NA | 0 | 0 | NA | 68.24 | 0.4 | 1.77 |
| AG-904-O14 | GORG | 0 | 0 | NA | 1 | 1 | orphan | 0 | 0 | NA | 0 | 1 | orphan | 63.27 | 0.4 | 1.6 |
| AG-893-N10 | GORG | 0 | 0 | NA | 1 | 0 | orphan | 0 | 0 | NA | 0 | 0 | NA | 62.59 | 0.4 | 1.49 |
| AG-892-F07 | GORG | 0 | 0 | NA | 2 | 0 | orphan | 0 | 0 | NA | 0 | 0 | NA | 58.4 | 0.4 | 1.49 |
| AG-414-C21 | GORG | 0 | 0 | NA | 2 | 0 | orphan | 0 | 0 | NA | 0 | 0 | NA | 55.77 | 0.4 | 1.51 |
| AG-333-I03 | GORG | 0 | 0 | NA | 1 | 0 | orphan | 0 | 0 | NA | 0 | 0 | NA | 55.56 | 0.4 | 1.25 |
| AG-430-D23 | GORG | 0 | 0 | NA | 1 | 0 | orphan | 0 | 0 | NA | 0 | 0 | NA | 55.31 | 0.4 | 1.47 |
| AG-893-O07 | GORG | 0 | 0 | NA | 0 | 0 | NA | 0 | 0 | NA | 0 | 1 | orphan | 60.45 | 0.41 | 1.04 |
| AG-893-I10 | GORG | 0 | 0 | NA | 1 | 0 | orphan | 0 | 0 | NA | 0 | 0 | NA | 55.7 | 0.41 | 0.99 |
| AG-349-N23 | GORG | 0 | 0 | NA | 0 | 0 | NA | 0 | 0 | NA | 0 | 1 | orphan | 54.37 | 0.41 | 0.79 |
| AG-894-A16 | GORG | 0 | 0 | NA | 1 | 0 | orphan | 0 | 0 | NA | 0 | 1 | orpha | 84.73 | 0.43 | 2.04 |

| | | | | | | | | | | | | | | | |
|---|---|---|---|---|---|---|---|---|---|---|---|---|---|---|---|
| | | | | | | | | | | | | | n | | | |
| AG-453-B23 | GORG | 0 | 0 | NA | 2 | 0 | orphan | 0 | 0 | NA | 0 | 0 | NA | 80.82 | 0.43 | 1.28 |
| AG-891-K18 | GORG | 1 | 1 | complete | 1 | 0 | orphan | 0 | 0 | NA | 0 | 0 | NA | 80.08 | 0.43 | 1.2 |
| AG-343-O04 | GORG | 0 | 0 | NA | 1 | 0 | orphan | 0 | 0 | NA | 0 | 0 | NA | 76.92 | 0.43 | 0.98 |
| AG-333-G05 | GORG | 1 | 1 | complete | 1 | 0 | orphan | 0 | 0 | NA | 0 | 0 | NA | 76.07 | 0.43 | 1.09 |
| AG-333-G15 | GORG | 0 | 0 | NA | 1 | 0 | orphan | 0 | 0 | NA | 0 | 0 | NA | 76.07 | 0.43 | 0.96 |
| AG-915-F11 | GORG | 1 | 1 | complete | 1 | 0 | orphan | 0 | 0 | NA | 0 | 0 | NA | 76.07 | 0.43 | 1.1 |
| AG-919-E16 | GORG | 0 | 0 | NA | 2 | 0 | orphan | 0 | 0 | NA | 0 | 0 | NA | 75.31 | 0.43 | 1.78 |
| AG-891-I09 | GORG | 0 | 0 | NA | 3 | 0 | orphan | 0 | 0 | NA | 0 | 0 | NA | 75.21 | 0.43 | 1.07 |
| AG-390-E09 | GORG | 0 | 0 | NA | 1 | 0 | orphan | 0 | 0 | NA | 0 | 0 | NA | 74.79 | 0.43 | 1.03 |
| AG-439-B19 | GORG | 0 | 0 | NA | 1 | 0 | orphan | 0 | 0 | NA | 0 | 0 | NA | 74.79 | 0.43 | 1.28 |
| AG-895-L13 | GORG | 0 | 0 | NA | 3 | 0 | orphan | 0 | 0 | NA | 0 | 0 | NA | 74.71 | 0.43 | 1.04 |
| AG-422-C10 | GORG | 0 | 0 | NA | 1 | 0 | orphan | 0 | 0 | NA | 0 | 0 | NA | 74.55 | 0.43 | 1.03 |
| AG-919-G14 | GORG | 0 | 0 | NA | 1 | 0 | orphan | 0 | 0 | NA | 0 | 0 | NA | 74.02 | 0.43 | 1.07 |
| AG-903-G18 | GORG | 0 | 0 | NA | 1 | 0 | orphan | 0 | 0 | NA | 0 | 0 | NA | 72.65 | 0.43 | 1.06 |
| AG-919-L04 | GORG | 0 | 1 | NA | 1 | 0 | orphan | 0 | 0 | NA | 0 | 0 | NA | 72.65 | 0.43 | 1 |
| AH-321-A09 | GORG | 0 | 0 | NA | 1 | 0 | orphan | 0 | 0 | NA | 0 | 0 | NA | 72.65 | 0.43 | 1.02 |
| AG-911-D11 | GORG | 0 | 0 | NA | 1 | 0 | orphan | 0 | 0 | NA | 0 | 0 | NA | 72 | 0.43 | 1 |
| AG-920-J05 | GORG | 0 | 0 | NA | 3 | 0 | orphan | 0 | 0 | NA | 0 | 0 | NA | 71.79 | 0.43 | 1.08 |
| AG-359-K22 | GORG | 0 | 0 | NA | 2 | 0 | orphan | 0 | 0 | NA | 0 | 0 | NA | 71.37 | 0.43 | 1.06 |
| AG-349-C02 | GORG | 0 | 0 | NA | 1 | 0 | orphan | 0 | 0 | NA | 0 | 0 | NA | 70.94 | 0.43 | 0.99 |
| AG-404-I14 | GORG | 0 | 0 | NA | 1 | 0 | orphan | 0 | 0 | NA | 0 | 0 | NA | 70.94 | 0.43 | 1.02 |
| AG-897-D06 | GORG | 0 | 0 | NA | 1 | 0 | orphan | 0 | 0 | NA | 0 | 0 | NA | 70.94 | 0.43 | 0.98 |
| AG-899-O20 | GORG | 0 | 0 | NA | 2 | 0 | orphan | 0 | 0 | NA | 0 | 0 | NA | 70.59 | 0.43 | 1.68 |
| AG-447-O20 | GORG | 0 | 0 | NA | 1 | 0 | orphan | 0 | 0 | NA | 0 | 0 | NA | 70.44 | 0.43 | 0.94 |
| AG-426-J16 | GORG | 0 | 0 | NA | 4 | 0 | orphan | 0 | 0 | NA | 0 | 0 | NA | 70.09 | 0.43 | 1.02 |

| | | | | | | | | | | | | | | | | |
|---|---|---|---|---|---|---|---|---|---|---|---|---|---|---|---|---|
| AG-913-G17 | GORG | 1 | 1 | complete | 1 | 0 | orphan | 0 | 0 | NA | 0 | 0 | NA | 69.23 | 0.43 | 0.97 |
| AG-916-I18 | GORG | 0 | 0 | NA | 1 | 0 | orphan | 0 | 0 | NA | 0 | 0 | NA | 69.23 | 0.43 | 1.05 |
| AG-333-M17 | GORG | 0 | 0 | NA | 1 | 0 | orphan | 0 | 0 | NA | 0 | 0 | NA | 68.38 | 0.43 | 0.94 |
| AG-426-M18 | GORG | 0 | 0 | NA | 1 | 0 | orphan | 0 | 0 | NA | 0 | 0 | NA | 68.38 | 0.43 | 1.03 |
| AG-359-N14 | GORG | 0 | 0 | NA | 1 | 0 | orphan | 0 | 0 | NA | 0 | 0 | NA | 67.81 | 0.43 | 0.96 |
| AH-324-A14 | GORG | 0 | 0 | NA | 2 | 0 | orphan | 0 | 0 | NA | 0 | 0 | NA | 67.52 | 0.43 | 1.43 |
| AG-918-I09 | GORG | 0 | 0 | NA | 3 | 0 | orphan | 0 | 0 | NA | 0 | 0 | NA | 67.09 | 0.43 | 0.99 |
| AG-911-K21 | GORG | 0 | 0 | NA | 1 | 0 | orphan | 0 | 0 | NA | 0 | 0 | NA | 66.67 | 0.43 | 1.01 |
| AG-917-A11 | GORG | 0 | 0 | NA | 1 | 0 | orphan | 0 | 0 | NA | 0 | 0 | NA | 66.67 | 0.43 | 1.03 |
| AG-461-K19 | GORG | 0 | 0 | NA | 1 | 0 | orphan | 0 | 0 | NA | 0 | 0 | NA | 66.24 | 0.43 | 0.87 |
| AG-908-E09 | GORG | 0 | 0 | NA | 2 | 0 | orphan | 0 | 0 | NA | 0 | 0 | NA | 65.81 | 0.43 | 0.98 |
| AG-447-L16 | GORG | 0 | 0 | NA | 4 | 0 | orphan | 0 | 0 | NA | 0 | 0 | NA | 64.96 | 0.43 | 0.93 |
| AG-461-C10 | GORG | 0 | 0 | NA | 1 | 0 | orphan | 0 | 0 | NA | 0 | 0 | NA | 64.96 | 0.43 | 0.94 |
| AG-430-M11 | GORG | 0 | 0 | NA | 4 | 0 | orphan | 0 | 0 | NA | 0 | 0 | NA | 64.2 | 0.43 | 1.02 |
| AG-916-D09 | GORG | 0 | 0 | NA | 1 | 0 | orphan | 0 | 0 | NA | 0 | 0 | NA | 63.59 | 0.43 | 1 |
| AG-426-J07 | GORG | 0 | 0 | NA | 1 | 0 | orphan | 0 | 0 | NA | 0 | 0 | NA | 63.25 | 0.43 | 0.89 |
| AG-919-O23 | GORG | 0 | 0 | NA | 1 | 0 | orphan | 0 | 0 | NA | 0 | 0 | NA | 63.25 | 0.43 | 0.91 |
| AH-326-D23 | GORG | 0 | 0 | NA | 1 | 0 | orphan | 0 | 0 | NA | 0 | 0 | NA | 63.25 | 0.43 | 1 |
| AG-901-G13 | GORG | 0 | 0 | NA | 2 | 0 | orphan | 0 | 0 | NA | 0 | 0 | NA | 63.03 | 0.43 | 0.84 |
| AG-325-J06 | GORG | 0 | 0 | NA | 1 | 0 | orphan | 0 | 0 | NA | 0 | 0 | NA | 62.39 | 0.43 | 0.95 |
| AG-917-G04 | GORG | 0 | 0 | NA | 2 | 0 | orphan | 0 | 0 | NA | 0 | 0 | NA | 62.35 | 0.43 | 1.56 |
| AH-287-C18 | GORG | 0 | 0 | NA | 1 | 0 | orphan | 0 | 0 | NA | 0 | 0 | NA | 61.96 | 0.43 | 2.41 |
| AG-915-A08 | GORG | 0 | 0 | NA | 1 | 0 | orphan | 0 | 0 | NA | 0 | 0 | NA | 61.88 | 0.43 | 0.91 |
| AG-912-N21 | GORG | 0 | 0 | NA | 1 | 0 | orphan | 0 | 0 | NA | 0 | 0 | NA | 61.78 | 0.43 | 0.85 |
| AG-912-I13 | GORG | 0 | 0 | NA | 1 | 0 | orphan | 0 | 0 | NA | 0 | 0 | NA | 61.54 | 0.43 | 0.96 |
| AG-903-D11 | GORG | 0 | 0 | NA | 4 | 0 | orphan | 0 | 0 | NA | 0 | 2 | orphan | 60.68 | 0.43 | 0.91 |

| | | | | | | | | | | | | | | | | |
|---|---|---|---|---|---|---|---|---|---|---|---|---|---|---|---|---|
| AG-447-G15 | GORG | 0 | 0 | NA | 1 | 0 | orphan | 0 | 0 | NA | 0 | 0 | NA | 60.68 | 0.43 | 0.84 |
| AG-313-L19 | GORG | 0 | 0 | NA | 1 | 0 | orphan | 0 | 0 | NA | 0 | 0 | NA | 60.57 | 0.43 | 0.81 |
| AG-908-K19 | GORG | 0 | 0 | NA | 1 | 0 | orphan | 0 | 0 | NA | 0 | 0 | NA | 60.26 | 0.43 | 0.95 |
| AG-912-P15 | GORG | 0 | 0 | NA | 1 | 0 | orphan | 0 | 0 | NA | 0 | 0 | NA | 60.26 | 0.43 | 0.96 |
| AG-453-E15 | GORG | 0 | 0 | NA | 1 | 0 | orphan | 0 | 0 | NA | 0 | 0 | NA | 59.4 | 0.43 | 1.43 |
| AH-324-I15 | GORG | 0 | 0 | NA | 2 | 0 | orphan | 0 | 0 | NA | 0 | 0 | NA | 59.27 | 0.43 | 1.39 |
| AG-918-M14 | GORG | 0 | 0 | NA | 2 | 0 | orphan | 0 | 0 | NA | 0 | 0 | NA | 59.21 | 0.43 | 0.94 |
| AH-287-K22 | GORG | 0 | 0 | NA | 1 | 0 | orphan | 0 | 0 | NA | 0 | 0 | NA | 58.12 | 0.43 | 0.7 |
| AH-321-L10 | GORG | 0 | 0 | NA | 3 | 0 | orphan | 0 | 0 | NA | 0 | 0 | NA | 58.12 | 0.43 | 0.81 |
| AG-908-O17 | GORG | 0 | 0 | NA | 1 | 0 | orphan | 0 | 0 | NA | 0 | 0 | NA | 57.09 | 0.43 | 0.83 |
| AG-404-P04 | GORG | 0 | 0 | NA | 2 | 0 | orphan | 0 | 0 | NA | 0 | 0 | NA | 56.8 | 0.43 | 0.79 |
| AG-913-L11 | GORG | 0 | 0 | NA | 2 | 0 | orphan | 0 | 0 | NA | 0 | 0 | NA | 56.7 | 0.43 | 0.93 |
| AG-447-L17 | GORG | 0 | 0 | NA | 2 | 1 | complete | 0 | 0 | NA | 0 | 0 | NA | 56.62 | 0.43 | 0.92 |
| AG-453-I20 | GORG | 0 | 0 | NA | 0 | 1 | NA | 0 | 0 | NA | 0 | 0 | NA | 56.53 | 0.43 | 1.36 |
| AG-457-F08 | GORG | 0 | 0 | NA | 1 | 0 | orphan | 0 | 0 | NA | 0 | 0 | NA | 54.93 | 0.43 | 1.64 |
| AG-333-K09 | GORG | 0 | 0 | NA | 1 | 0 | orphan | 0 | 0 | NA | 0 | 0 | NA | 54.82 | 0.43 | 0.8 |
| AG-414-K08 | GORG | 0 | 0 | NA | 1 | 0 | orphan | 0 | 0 | NA | 0 | 0 | NA | 54.7 | 0.43 | 0.76 |
| AG-910-M10 | GORG | 0 | 0 | NA | 2 | 0 | orphan | 0 | 0 | NA | 0 | 0 | NA | 54.19 | 0.43 | 0.76 |
| AG-430-I21 | GORG | 0 | 0 | NA | 1 | 0 | orphan | 0 | 0 | NA | 0 | 0 | NA | 53.85 | 0.43 | 0.89 |
| AG-903-I02 | GORG | 0 | 0 | NA | 1 | 0 | orphan | 0 | 0 | NA | 0 | 0 | NA | 53.85 | 0.43 | 0.79 |
| AG-915-E18 | GORG | 0 | 0 | NA | 1 | 0 | orphan | 0 | 0 | NA | 0 | 0 | NA | 53.73 | 0.43 | 0.68 |
| AG-915-C17 | GORG | 0 | 0 | NA | 1 | 0 | orphan | 0 | 0 | NA | 0 | 0 | NA | 53.42 | 0.43 | 0.85 |
| AH-287-P09 | GORG | 0 | 0 | NA | 1 | 0 | orphan | 0 | 0 | NA | 0 | 0 | NA | 53.13 | 0.43 | 0.81 |
| AG-918-B15 | GORG | 0 | 0 | NA | 1 | 0 | orphan | 0 | 0 | NA | 0 | 0 | NA | 53.05 | 0.43 | 0.71 |
| AG-325-E09 | GORG | 0 | 0 | NA | 1 | 0 | orphan | 0 | 0 | NA | 0 | 0 | NA | 52.99 | 0.43 | 0.83 |
| AG-360-L09 | GORG | 1 | 1 | complete | 1 | 0 | orphan | 0 | 0 | NA | 0 | 0 | NA | 52.53 | 0.43 | 0.86 |

| | | | | | | | | | | | | | | | | |
|---|---|---|---|---|---|---|---|---|---|---|---|---|---|---|---|---|
| AG-892-O14 | GORG | 0 | 0 | NA | 3 | 0 | orphan | 0 | 0 | NA | 0 | 0 | NA | 52.23 | 0.43 | 0.97 |
| AG-439-N06 | GORG | 0 | 0 | NA | 3 | 0 | orphan | 0 | 0 | NA | 0 | 0 | NA | 52.14 | 0.43 | 0.78 |
| AG-917-A10 | GORG | 0 | 0 | NA | 2 | 0 | orphan | 0 | 0 | NA | 0 | 0 | NA | 50.47 | 0.43 | 0.74 |
| AH-287-B15 | GORG | 0 | 0 | NA | 0 | 0 | NA | 0 | 0 | NA | 0 | 2 | orphan | 82.26 | 0.44 | 2.39 |
| AG-894-E05 | GORG | 0 | 0 | NA | 3 | 0 | orphan | 0 | 0 | NA | 0 | 0 | NA | 75.25 | 0.44 | 1.94 |
| AG-325-M05 | GORG | 0 | 1 | NA | 2 | 0 | orphan | 0 | 0 | NA | 0 | 0 | NA | 66.3 | 0.44 | 1.47 |
| AG-891-G07 | GORG | 0 | 0 | NA | 0 | 0 | NA | 0 | 0 | NA | 0 | 1 | orphan | 61.2 | 0.44 | 1.9 |
| AG-912-K11 | GORG | 0 | 0 | NA | 2 | 0 | orphan | 0 | 1 | NA | 0 | 0 | NA | 59.61 | 0.44 | 1.5 |
| AG-919-I23 | GORG | 0 | 1 | NA | 1 | 0 | orphan | 0 | 0 | NA | 0 | 0 | NA | 59.61 | 0.44 | 1.85 |
| AG-896-C14 | GORG | 0 | 0 | NA | 3 | 0 | orphan | 0 | 0 | NA | 0 | 0 | NA | 57.29 | 0.44 | 1.81 |
| AG-907-J10 | GORG | 0 | 0 | NA | 2 | 0 | orphan | 0 | 0 | NA | 0 | 0 | NA | 55.78 | 0.44 | 1.62 |
| AG-914-O23 | GORG | 0 | 0 | NA | 2 | 0 | orphan | 0 | 0 | NA | 0 | 0 | NA | 52.96 | 0.44 | 1.98 |
| AG-896-K16 | GORG | 1 | 1 | complete | 0 | 0 | NA | 0 | 0 | NA | 0 | 0 | NA | 52.71 | 0.44 | 1.71 |
| AG-422-D21 | GORG | 0 | 0 | NA | 0 | 0 | NA | 0 | 0 | NA | 0 | 1 | orphan | 73.18 | 0.45 | 1.22 |
| AG-898-M15 | GORG | 0 | 0 | NA | 1 | 0 | orphan | 0 | 0 | NA | 0 | 0 | NA | 74.69 | 0.46 | 1.93 |
| AG-919-N15 | GORG | 0 | 0 | NA | 1 | 0 | orphan | 0 | 0 | NA | 0 | 0 | NA | 52.95 | 0.46 | 0.94 |
| AG-893-F11 | GORG | 0 | 0 | NA | 1 | 0 | orphan | 0 | 0 | NA | 0 | 1 | orphan | 100 | 0.47 | 1.19 |
| AG-894-C14 | GORG | 0 | 0 | NA | 1 | 0 | orphan | 0 | 0 | NA | 0 | 0 | NA | 99.37 | 0.47 | 1.22 |
| AG-447-C08 | GORG | 0 | 0 | NA | 1 | 0 | orphan | 0 | 0 | NA | 0 | 0 | NA | 95.5 | 0.47 | 1.25 |
| AG-904-F17 | GORG | 0 | 0 | NA | 1 | 0 | orphan | 0 | 0 | NA | 0 | 0 | NA | 95.38 | 0.47 | 1.2 |
| AG-913-C03 | GORG | 0 | 0 | NA | 2 | 0 | orphan | 0 | 1 | NA | 0 | 0 | NA | 95.26 | 0.47 | 1.21 |
| AG-894-B07 | GORG | 0 | 0 | NA | 3 | 0 | orphan | 0 | 0 | NA | 0 | 0 | NA | 94.51 | 0.47 | 1.14 |
| AG-913-F03 | GORG | 0 | 0 | NA | 1 | 0 | orphan | 0 | 0 | NA | 0 | 1 | orphan | 93.68 | 0.47 | 1.17 |
| AG-899-J23 | GORG | 0 | 0 | NA | 1 | 0 | orphan | 0 | 0 | NA | 0 | 0 | NA | 93.52 | 0.47 | 1.1 |
| AG-916-N14 | GORG | 0 | 0 | NA | 2 | 0 | orphan | 0 | 0 | NA | 0 | 0 | NA | 93.05 | 0.47 | 1.27 |

| | | | | | | | | | | | | | | | | |
|---|---|---|---|---|---|---|---|---|---|---|---|---|---|---|---|---|
| AG-914-L04 | GORG | 1 | 1 | complet<br>e | 1 | 0 | orphan | 0 | 0 | NA | 0 | 0 | NA | 92.65 | 0.47 | 1.09 |
| AG-909-G08 | GORG | 0 | 0 | NA | 3 | 0 | orphan | 0 | 0 | NA | 0 | 0 | NA | 91.94 | 0.47 | 1.12 |
| AG-914-G22 | GORG | 0 | 0 | NA | 2 | 0 | orphan | 0 | 0 | NA | 0 | 0 | NA | 91.91 | 0.47 | 1.04 |
| AG-918-L21 | GORG | 0 | 0 | NA | 3 | 0 | orphan | 0 | 0 | NA | 0 | 0 | NA | 91.36 | 0.47 | 1.18 |
| AG-899-J02 | GORG | 0 | 0 | NA | 1 | 0 | orphan | 0 | 0 | NA | 0 | 0 | NA | 91.04 | 0.47 | 1.21 |
| AG-905-G04 | GORG | 0 | 0 | NA | 2 | 0 | orphan | 0 | 0 | NA | 0 | 0 | NA | 90.05 | 0.47 | 1.03 |
| AG-916-M15 | GORG | 0 | 0 | NA | 3 | 0 | orphan | 0 | 0 | NA | 0 | 0 | NA | 90.03 | 0.47 | 1.13 |
| AG-903-D02 | GORG | 1 | 0 | orphan | 1 | 0 | orphan | 0 | 0 | NA | 0 | 0 | NA | 89.4 | 0.47 | 1.15 |
| AG-920-J04 | GORG | 0 | 0 | NA | 1 | 0 | orphan | 0 | 0 | NA | 0 | 0 | NA | 89.26 | 0.47 | 1.13 |
| AG-892-E08 | GORG | 0 | 0 | NA | 2 | 0 | orphan | 0 | 0 | NA | 0 | 0 | NA | 89.1 | 0.47 | 1.25 |
| AG-333-D08 | GORG | 0 | 0 | NA | 2 | 0 | orphan | 0 | 0 | NA | 0 | 0 | NA | 88.7 | 0.47 | 1.01 |
| AG-422-B22 | GORG | 0 | 0 | NA | 2 | 0 | orphan | 0 | 0 | NA | 0 | 0 | NA | 88.63 | 0.47 | 1.16 |
| AG-919-L06 | GORG | 0 | 0 | NA | 1 | 0 | orphan | 0 | 0 | NA | 0 | 0 | NA | 88.39 | 0.47 | 1.13 |
| AH-287-B07 | GORG | 0 | 0 | NA | 2 | 0 | orphan | 0 | 0 | NA | 0 | 0 | NA | 87.44 | 0.47 | 1.21 |
| AG-913-B21 | GORG | 0 | 0 | NA | 1 | 0 | orphan | 0 | 0 | NA | 0 | 0 | NA | 87.32 | 0.47 | 0.98 |
| AG-337-D17 | GORG | 0 | 0 | NA | 1 | 0 | orphan | 0 | 0 | NA | 0 | 0 | NA | 86.65 | 0.47 | 1.06 |
| AG-390-I15 | GORG | 0 | 0 | NA | 2 | 0 | orphan | 0 | 0 | NA | 0 | 0 | NA | 86.35 | 0.47 | 1.14 |
| AG-897-N19 | GORG | 0 | 0 | NA | 1 | 0 | orphan | 0 | 0 | NA | 0 | 0 | NA | 86.01 | 0.47 | 1.08 |
| AG-920-O11 | GORG | 1 | 1 | complet<br>e | 0 | 0 | NA | 0 | 0 | NA | 0 | 0 | NA | 85.74 | 0.47 | 1.02 |
| AG-414-D10 | GORG | 0 | 0 | NA | 1 | 0 | orphan | 0 | 0 | NA | 0 | 0 | NA | 84.04 | 0.47 | 1.08 |
| AG-894-M23 | GORG | 0 | 0 | NA | 1 | 0 | orphan | 0 | 0 | NA | 0 | 0 | NA | 83.77 | 0.47 | 1.04 |
| AG-319-M14 | GORG | 0 | 0 | NA | 2 | 0 | orphan | 0 | 0 | NA | 0 | 0 | NA | 83.65 | 0.47 | 1.2 |
| AG-349-O13 | GORG | 1 | 0 | orphan | 2 | 0 | orphan | 0 | 0 | NA | 0 | 0 | NA | 82.63 | 0.47 | 0.95 |
| AG-457-B14 | GORG | 0 | 0 | NA | 2 | 0 | orphan | 0 | 0 | NA | 0 | 0 | NA | 82.37 | 0.47 | 1.04 |
| AG-447-G05 | GORG | 0 | 0 | NA | 1 | 0 | orphan | 0 | 0 | NA | 0 | 0 | NA | 81.2 | 0.47 | 1.06 |
| AH-321-P10 | GORG | 0 | 0 | NA | 2 | 0 | orphan | 0 | 0 | NA | 0 | 0 | NA | 80.96 | 0.47 | 1.14 |

| | | | | | | | | | | | | | | | | |
|---|---|---|---|---|---|---|---|---|---|---|---|---|---|---|---|---|
| AG-916-P21 | GORG | 0 | 0 | NA | 2 | 0 | orphan | 0 | 0 | NA | 0 | 0 | NA | 80.82 | 0.47 | 0.95 |
| AG-430-K05 | GORG | 0 | 0 | NA | 1 | 0 | orphan | 0 | 0 | NA | 0 | 0 | NA | 80.78 | 0.47 | 0.92 |
| AG-461-M14 | GORG | 0 | 0 | NA | 1 | 0 | orphan | 0 | 0 | NA | 0 | 0 | NA | 80.07 | 0.47 | 1.03 |
| AG-911-B23 | GORG | 1 | 1 | complete | 1 | 0 | orphan | 0 | 0 | NA | 0 | 0 | NA | 79.9 | 0.47 | 1.03 |
| AG-919-A21 | GORG | 1 | 1 | complete | 1 | 0 | orphan | 0 | 0 | NA | 0 | 0 | NA | 79.79 | 0.47 | 1.01 |
| AG-894-J16 | GORG | 0 | 0 | NA | 4 | 0 | orphan | 0 | 0 | NA | 0 | 0 | NA | 79.58 | 0.47 | 0.9 |
| AG-447-M19 | GORG | 0 | 0 | NA | 1 | 0 | orphan | 0 | 0 | NA | 0 | 0 | NA | 79.3 | 0.47 | 0.93 |
| AG-447-F02 | GORG | 0 | 0 | NA | 1 | 0 | orphan | 0 | 0 | NA | 0 | 0 | NA | 78.85 | 0.47 | 1.1 |
| AG-349-M03 | GORG | 0 | 0 | NA | 1 | 0 | orphan | 0 | 0 | NA | 0 | 1 | orphan | 77.88 | 0.47 | 1 |
| AG-920-N08 | GORG | 0 | 0 | NA | 1 | 0 | orphan | 0 | 0 | NA | 0 | 0 | NA | 77.77 | 0.47 | 0.96 |
| AG-325-K09 | GORG | 0 | 0 | NA | 1 | 0 | orphan | 0 | 0 | NA | 0 | 0 | NA | 77.22 | 0.47 | 1.01 |
| AG-895-O16 | GORG | 0 | 0 | NA | 1 | 0 | orphan | 0 | 0 | NA | 0 | 0 | NA | 77.12 | 0.47 | 1.03 |
| AG-893-J20 | GORG | 0 | 0 | NA | 1 | 0 | orphan | 0 | 0 | NA | 0 | 0 | NA | 76.86 | 0.47 | 1.08 |
| AG-914-P09 | GORG | 0 | 0 | NA | 1 | 0 | orphan | 0 | 0 | NA | 0 | 0 | NA | 76.78 | 0.47 | 0.94 |
| AG-913-O03 | GORG | 0 | 0 | NA | 3 | 0 | orphan | 0 | 0 | NA | 0 | 0 | NA | 75.75 | 0.47 | 0.84 |
| AG-909-D16 | GORG | 0 | 0 | NA | 1 | 0 | orphan | 0 | 0 | NA | 0 | 0 | NA | 75.05 | 0.47 | 0.95 |
| AG-447-P03 | GORG | 0 | 0 | NA | 1 | 0 | orphan | 0 | 0 | NA | 0 | 0 | NA | 74.72 | 0.47 | 1.16 |
| AG-461-A22 | GORG | 0 | 0 | NA | 1 | 0 | orphan | 0 | 0 | NA | 0 | 0 | NA | 74.33 | 0.47 | 0.92 |
| AG-333-E22 | GORG | 0 | 0 | NA | 2 | 0 | orphan | 0 | 0 | NA | 0 | 0 | NA | 73.57 | 0.47 | 0.87 |
| AG-911-M05 | GORG | 1 | 0 | orphan | 2 | 0 | orphan | 0 | 0 | NA | 0 | 0 | NA | 72.02 | 0.47 | 0.79 |
| AG-896-K07 | GORG | 0 | 0 | NA | 2 | 0 | orphan | 0 | 0 | NA | 0 | 0 | NA | 70.94 | 0.47 | 0.98 |
| AH-324-A02 | GORG | 0 | 0 | NA | 1 | 0 | orphan | 0 | 0 | NA | 0 | 0 | NA | 70.77 | 0.47 | 0.91 |
| AG-914-O09 | GORG | 0 | 0 | NA | 1 | 0 | orphan | 0 | 0 | NA | 0 | 0 | NA | 70.67 | 0.47 | 0.91 |
| AG-904-B09 | GORG | 0 | 0 | NA | 1 | 0 | orphan | 0 | 0 | NA | 0 | 0 | NA | 69.96 | 0.47 | 0.92 |
| AH-287-B14 | GORG | 0 | 1 | NA | 1 | 0 | orphan | 0 | 0 | NA | 0 | 0 | NA | 68.72 | 0.47 | 0.85 |
| AG-457-O13 | GORG | 0 | 0 | NA | 1 | 0 | orphan | 0 | 0 | NA | 0 | 0 | NA | 67.85 | 0.47 | 0.83 |

| | | | | | | | | | | | | | | | | |
|---|---|---|---|---|---|---|---|---|---|---|---|---|---|---|---|---|
| AG-319-F05 | GORG | 0 | 0 | NA | 2 | 0 | orphan | 0 | 0 | NA | 0 | 0 | NA | 67.12 | 0.47 | 0.9 |
| AG-916-F20 | GORG | 0 | 0 | NA | 1 | 0 | orphan | 0 | 0 | NA | 0 | 0 | NA | 67.06 | 0.47 | 0.89 |
| AH-287-B05 | GORG | 0 | 0 | NA | 1 | 0 | orphan | 0 | 0 | NA | 0 | 0 | NA | 66.7 | 0.47 | 0.86 |
| AH-321-B23 | GORG | 1 | 0 | orphan | 1 | 0 | orphan | 0 | 0 | NA | 0 | 0 | NA | 65.02 | 0.47 | 0.92 |
| AG-343-L02 | GORG | 0 | 0 | NA | 1 | 0 | orphan | 0 | 0 | NA | 0 | 0 | NA | 64.98 | 0.47 | 0.87 |
| AG-422-L21 | GORG | 0 | 0 | NA | 1 | 0 | orphan | 0 | 0 | NA | 0 | 0 | NA | 64.77 | 0.47 | 1.01 |
| AG-914-D20 | GORG | 0 | 0 | NA | 2 | 0 | orphan | 0 | 0 | NA | 0 | 0 | NA | 64.22 | 0.47 | 0.92 |
| AG-908-A06 | GORG | 0 | 0 | NA | 3 | 0 | orphan | 0 | 0 | NA | 0 | 0 | NA | 63.92 | 0.47 | 0.79 |
| AG-916-C11 | GORG | 0 | 0 | NA | 1 | 0 | orphan | 0 | 0 | NA | 0 | 0 | NA | 63.82 | 0.47 | 0.75 |
| AG-390-M06 | GORG | 0 | 0 | NA | 1 | 0 | orphan | 0 | 0 | NA | 0 | 0 | NA | 62.6 | 0.47 | 0.77 |
| AG-892-O13 | GORG | 0 | 0 | NA | 1 | 0 | orphan | 0 | 0 | NA | 0 | 1 | orphan | 62.55 | 0.47 | 0.63 |
| AG-912-O07 | GORG | 1 | 1 | complete | 0 | 0 | NA | 0 | 0 | NA | 0 | 0 | NA | 62.37 | 0.47 | 0.74 |
| AG-916-L04 | GORG | 1 | 1 | complete | 0 | 0 | NA | 0 | 0 | NA | 0 | 0 | NA | 62.15 | 0.47 | 0.76 |
| AG-908-E04 | GORG | 0 | 0 | NA | 1 | 0 | orphan | 0 | 0 | NA | 0 | 0 | NA | 60.91 | 0.47 | 0.76 |
| AG-912-B06 | GORG | 0 | 0 | NA | 1 | 0 | orphan | 0 | 0 | NA | 0 | 0 | NA | 60.19 | 0.47 | 0.76 |
| AG-360-K11 | GORG | 0 | 0 | NA | 1 | 0 | orphan | 0 | 0 | NA | 0 | 0 | NA | 60.02 | 0.47 | 0.9 |
| AG-390-A21 | GORG | 1 | 0 | orphan | 0 | 0 | NA | 0 | 0 | NA | 0 | 0 | NA | 59.38 | 0.47 | 0.8 |
| AH-326-A08 | GORG | 0 | 0 | NA | 1 | 0 | orphan | 0 | 0 | NA | 0 | 0 | NA | 58.45 | 0.47 | 1.02 |
| AG-910-O20 | GORG | 1 | 1 | complete | 1 | 0 | orphan | 0 | 0 | NA | 0 | 0 | NA | 56.7 | 0.47 | 0.8 |
| AG-909-G06 | GORG | 0 | 0 | NA | 1 | 0 | orphan | 0 | 0 | NA | 0 | 0 | NA | 56.58 | 0.47 | 0.79 |
| AG-390-J11 | GORG | 0 | 0 | NA | 1 | 0 | orphan | 0 | 0 | NA | 0 | 0 | NA | 53.96 | 0.47 | 0.65 |
| AG-919-P03 | GORG | 0 | 0 | NA | 1 | 0 | orphan | 1 | 1 | complete | 0 | 0 | NA | 53.72 | 0.47 | 0.76 |
| AH-321-M18 | GORG | 1 | 1 | complete | 1 | 0 | orphan | 0 | 0 | NA | 0 | 0 | NA | 53.17 | 0.47 | 0.63 |
| AG-447-L14 | GORG | 0 | 0 | NA | 1 | 0 | orphan | 0 | 0 | NA | 0 | 0 | NA | 90.89 | 0.48 | 1.05 |
| AG-359-F04 | GORG | 0 | 0 | NA | 2 | 0 | orphan | 0 | 1 | NA | 0 | 0 | NA | 89.6 | 0.48 | 1.13 |

| | | | | | | | | | | | | | | | | |
|---|---|---|---|---|---|---|---|---|---|---|---|---|---|---|---|---|
| AG-430-J07 | GORG | 0 | 0 | NA | 2 | 0 | orphan | 0 | 0 | NA | 0 | 0 | NA | 87.21 | 0.48 | 0.97 |
| AG-333-M03 | GORG | 0 | 0 | NA | 1 | 0 | orphan | 0 | 0 | NA | 0 | 0 | NA | 84.6 | 0.48 | 1.02 |
| AG-905-E10 | GORG | 0 | 0 | NA | 1 | 0 | orphan | 0 | 0 | NA | 0 | 0 | NA | 83.57 | 0.48 | 1.07 |
| AG-359-G06 | GORG | 0 | 0 | NA | 1 | 0 | orphan | 0 | 0 | NA | 0 | 1 | orphan | 82.14 | 0.48 | 0.92 |
| AG-915-K17 | GORG | 0 | 0 | NA | 2 | 0 | orphan | 0 | 0 | NA | 0 | 0 | NA | 81.19 | 0.48 | 0.99 |
| AG-918-P15 | GORG | 1 | 1 | complete | 1 | 0 | orphan | 0 | 0 | NA | 0 | 0 | NA | 80.76 | 0.48 | 0.96 |
| AG-914-M07 | GORG | 0 | 0 | NA | 2 | 0 | orphan | 0 | 0 | NA | 0 | 0 | NA | 74.46 | 0.48 | 0.81 |
| AG-892-B21 | GORG | 0 | 0 | NA | 1 | 0 | orphan | 0 | 0 | NA | 0 | 0 | NA | 74.11 | 0.48 | 0.93 |
| AG-904-C10 | GORG | 0 | 0 | NA | 1 | 0 | orphan | 0 | 1 | NA | 0 | 0 | NA | 72.54 | 0.48 | 0.87 |
| AH-287-P03 | GORG | 0 | 0 | NA | 1 | 0 | orphan | 0 | 0 | NA | 0 | 0 | NA | 71.93 | 0.48 | 0.87 |
| AH-326-I06 | GORG | 0 | 0 | NA | 2 | 0 | orphan | 0 | 0 | NA | 0 | 0 | NA | 71.19 | 0.48 | 0.8 |
| AG-470-P06 | GORG | 0 | 0 | NA | 1 | 0 | orphan | 0 | 0 | NA | 0 | 0 | NA | 69.93 | 0.48 | 0.91 |
| AG-916-C06 | GORG | 0 | 1 | NA | 3 | 0 | orphan | 0 | 1 | NA | 0 | 0 | NA | 68.59 | 0.48 | 2.24 |
| AG-911-D19 | GORG | 0 | 0 | NA | 1 | 0 | orphan | 0 | 0 | NA | 0 | 0 | NA | 65.71 | 0.48 | 0.77 |
| AG-900-C09 | GORG | 0 | 0 | NA | 2 | 0 | orphan | 0 | 0 | NA | 0 | 0 | NA | 65.3 | 0.48 | 0.74 |
| AG-910-D18 | GORG | 0 | 0 | NA | 2 | 0 | orphan | 0 | 0 | NA | 0 | 0 | NA | 63.25 | 0.48 | 0.72 |
| AG-905-B10 | GORG | 0 | 0 | NA | 1 | 0 | orphan | 0 | 0 | NA | 0 | 1 | orphan | 61.83 | 0.48 | 1.1 |
| AG-909-P05 | GORG | 1 | 1 | complete | 1 | 0 | orphan | 0 | 0 | NA | 0 | 0 | NA | 59.05 | 0.48 | 0.67 |
| AG-359-I14 | GORG | 0 | 0 | NA | 2 | 0 | orphan | 0 | 0 | NA | 0 | 0 | NA | 55.52 | 0.48 | 0.65 |
| AG-453-F22 | GORG | 0 | 0 | NA | 1 | 0 | orphan | 0 | 0 | NA | 0 | 0 | NA | 63.81 | 0.49 | 1.72 |
| AG-457-N23 | GORG | 1 | 1 | orphan | 0 | 0 | NA | 0 | 0 | NA | 0 | 0 | NA | 64.81 | 0.5 | 1.21 |
| AG-426-D09 | GORG | 0 | 0 | NA | 3 | 0 | orphan | 0 | 0 | NA | 0 | 0 | NA | 91.6 | 0.52 | 1.17 |
| AG-337-J09 | GORG | 0 | 0 | NA | 1 | 0 | orphan | 0 | 0 | NA | 0 | 0 | NA | 73.47 | 0.52 | 0.98 |
| AG-337-C07 | GORG | 0 | 0 | NA | 1 | 0 | orphan | 0 | 0 | NA | 0 | 0 | NA | 68.72 | 0.52 | 0.72 |
| AG-903-P01 | GORG | 0 | 0 | NA | 3 | 0 | orphan | 0 | 0 | NA | 0 | 0 | NA | 52.45 | 0.52 | 0.59 |
| AG-916-C10 | GORG | 1 | 1 | complet | 1 | 0 | orphan | 0 | 0 | NA | 0 | 0 | NA | 83.58 | 0.53 | 1.97 |

| | | | | | | | | | | | | | | | | |
|---|---|---|---|---|---|---|---|---|---|---|---|---|---|---|---|---|
| | | | | e | | | | | | | | | | | | |
| AG-422-M03 | GORG | 1 | 1 | complete | 1 | 0 | orphan | 0 | 0 | NA | 0 | 0 | NA | 79.53 | 0.53 | 2.09 |
| AG-899-D10 | GORG | 0 | 0 | NA | 1 | 0 | orphan | 0 | 0 | NA | 0 | 0 | NA | 75.12 | 0.53 | 1.66 |
| AG-891-A21 | GORG | 1 | 1 | complete | 1 | 0 | orphan | 0 | 0 | NA | 0 | 1 | orphan | 69.74 | 0.53 | 1.62 |
| AG-899-J06 | GORG | 0 | 0 | NA | 2 | 0 | orphan | 0 | 0 | NA | 0 | 0 | NA | 68.68 | 0.53 | 1.64 |
| AG-435-J13 | GORG | 0 | 0 | NA | 2 | 0 | orphan | 0 | 0 | NA | 0 | 0 | NA | 66.67 | 0.53 | 1.84 |
| AG-899-C03 | GORG | 0 | 0 | NA | 1 | 0 | orphan | 0 | 0 | NA | 0 | 0 | NA | 62.73 | 0.53 | 1.61 |
| AG-904-C11 | GORG | 0 | 0 | NA | 1 | 0 | orphan | 0 | 0 | NA | 0 | 1 | orphan | 59.5 | 0.53 | 1.69 |
| AG-917-C06 | GORG | 0 | 0 | NA | 1 | 0 | orphan | 0 | 0 | NA | 0 | 0 | NA | 50.82 | 0.53 | 1.44 |
| AG-894-O08 | GORG | 0 | 0 | NA | 0 | 0 | NA | 0 | 0 | NA | 0 | 1 | orphan | 88.01 | 0.54 | 1.45 |
| AG-917-B09 | GORG | 0 | 0 | NA | 0 | 0 | NA | 0 | 0 | NA | 0 | 1 | orphan | 83.33 | 0.54 | 1.36 |
| AG-895-M21 | GORG | 0 | 0 | NA | 0 | 0 | NA | 0 | 0 | NA | 0 | 1 | orphan | 82.2 | 0.54 | 1.4 |
| AG-404-O15 | GORG | 0 | 0 | NA | 0 | 0 | NA | 0 | 0 | NA | 0 | 1 | orphan | 80.39 | 0.54 | 1.36 |
| AG-422-G21 | GORG | 0 | 0 | NA | 1 | 0 | orphan | 0 | 0 | NA | 0 | 0 | NA | 76.42 | 0.54 | 1.33 |
| AH-287-B22 | GORG | 0 | 1 | NA | 0 | 0 | NA | 0 | 0 | NA | 0 | 0 | NA | 74.95 | 0.54 | 1.21 |
| AG-917-O08 | GORG | 0 | 0 | NA | 1 | 0 | orphan | 0 | 0 | NA | 0 | 0 | NA | 73.23 | 0.54 | 1.21 |
| AG-359-P22 | GORG | 0 | 0 | NA | 0 | 0 | NA | 0 | 0 | NA | 0 | 1 | orphan | 65.94 | 0.54 | 1.04 |
| AG-333-G13 | GORG | 1 | 2 | complete | 0 | 0 | NA | 0 | 0 | NA | 0 | 0 | NA | 60.9 | 0.54 | 1.05 |
| AG-898-L05 | GORG | 0 | 0 | NA | 0 | 0 | NA | 0 | 0 | NA | 0 | 1 | orphan | 58.83 | 0.54 | 0.94 |
| AG-917-I07 | GORG | 0 | 0 | NA | 1 | 0 | orphan | 0 | 0 | NA | 0 | 0 | NA | 57.34 | 0.54 | 0.98 |
| AG-349-P06 | GORG | 0 | 1 | NA | 0 | 0 | NA | 0 | 0 | NA | 0 | 1 | orphan | 55.32 | 0.54 | 0.91 |
| AG-913-P14 | GORG | 0 | 0 | NA | 1 | 0 | orphan | 0 | 0 | NA | 0 | 0 | NA | 69.18 | 0.56 | 1.46 |
| AG-319-N23 | GORG | 0 | 0 | NA | 1 | 0 | orphan | 0 | 0 | NA | 0 | 0 | NA | 68.87 | 0.56 | 0.91 |

| | | | | | | | | | | | | | | | | |
|---|---|---|---|---|---|---|---|---|---|---|---|---|---|---|---|---|
| AG-404-F02 | GORG | 0 | 0 | NA | 1 | 0 | orphan | 0 | 0 | NA | 0 | 0 | NA | 50.28 | 0.56 | 1.77 |
| AG-912-C22 | GORG | 0 | 0 | NA | 3 | 0 | orphan | 0 | 0 | NA | 0 | 0 | NA | 83.05 | 0.57 | 1.5 |
| AG-908-B08 | GORG | 0 | 0 | NA | 1 | 0 | orphan | 0 | 0 | NA | 0 | 0 | NA | 81.82 | 0.57 | 1.82 |
| AG-917-G06 | GORG | 0 | 0 | NA | 1 | 0 | orphan | 0 | 0 | NA | 0 | 0 | NA | 77.69 | 0.57 | 0.86 |
| AG-911-M04 | GORG | 1 | 1 | complete | 0 | 0 | NA | 0 | 0 | NA | 0 | 0 | NA | 73.8 | 0.57 | 1 |
| AG-337-G10 | GORG | 0 | 0 | NA | 0 | 1 | NA | 0 | 0 | NA | 0 | 0 | NA | 71.12 | 0.57 | 1.07 |
| AG-917-K07 | GORG | 0 | 0 | NA | 1 | 0 | orphan | 0 | 0 | NA | 0 | 0 | NA | 69 | 0.57 | 1.01 |
| AG-457-O06 | GORG | 1 | 1 | complete | 0 | 0 | NA | 0 | 0 | NA | 0 | 0 | NA | 68.34 | 0.57 | 1.22 |
| AG-903-N07 | GORG | 0 | 0 | NA | 2 | 0 | orphan | 0 | 0 | NA | 0 | 0 | NA | 66.48 | 0.57 | 0.96 |
| AG-912-F17 | GORG | 0 | 0 | NA | 1 | 0 | orphan | 0 | 0 | NA | 0 | 0 | NA | 61.9 | 0.57 | 0.93 |
| AG-911-K13 | GORG | 0 | 0 | NA | 2 | 0 | orphan | 0 | 0 | NA | 0 | 0 | NA | 61.54 | 0.57 | 0.87 |
| AG-337-J15 | GORG | 0 | 0 | NA | 1 | 0 | orphan | 0 | 0 | NA | 0 | 0 | NA | 59.42 | 0.57 | 0.77 |
| AG-913-L06 | GORG | 0 | 0 | NA | 1 | 0 | orphan | 0 | 0 | NA | 0 | 0 | NA | 56.61 | 0.57 | 0.77 |
| AG-893-J17 | GORG | 0 | 0 | NA | 1 | 0 | orphan | 0 | 0 | NA | 0 | 0 | NA | 96.94 | 0.58 | 2.61 |
| AG-892-O07 | GORG | 0 | 0 | NA | 2 | 0 | orphan | 0 | 0 | NA | 0 | 0 | NA | 91.04 | 0.58 | 2.16 |
| AG-390-P06 | GORG | 0 | 0 | NA | 1 | 0 | orphan | 0 | 0 | NA | 0 | 0 | NA | 70.57 | 0.58 | 1.71 |
| AG-914-E17 | GORG | 0 | 0 | NA | 3 | 0 | orphan | 0 | 0 | NA | 0 | 0 | NA | 65.75 | 0.58 | 1.56 |
| AG-905-F13 | GORG | 0 | 0 | NA | 1 | 0 | orphan | 0 | 0 | NA | 0 | 0 | NA | 53.86 | 0.58 | 1.44 |
| AG-917-O05 | GORG | 0 | 0 | NA | 1 | 0 | orphan | 0 | 0 | NA | 0 | 0 | NA | 52.76 | 0.58 | 1.19 |
| AG-894-O04 | GORG | 0 | 0 | NA | 3 | 1 | orphan | 0 | 0 | NA | 0 | 0 | NA | 92.77 | 0.6 | 1.34 |
| AG-897-E09 | GORG | 0 | 0 | NA | 2 | 0 | orphan | 0 | 0 | NA | 0 | 0 | NA | 91.97 | 0.6 | 1.28 |
| AG-894-E10 | GORG | 0 | 0 | NA | 1 | 0 | orphan | 0 | 0 | NA | 0 | 0 | NA | 90.36 | 0.6 | 1.2 |
| AG-895-F07 | GORG | 0 | 0 | NA | 1 | 0 | orphan | 0 | 0 | NA | 0 | 0 | NA | 90.36 | 0.6 | 1.27 |
| AG-903-I23 | GORG | 0 | 0 | NA | 3 | 0 | orphan | 0 | 0 | NA | 0 | 0 | NA | 84.74 | 0.6 | 1.1 |
| AG-430-B21 | GORG | 0 | 0 | NA | 1 | 0 | orphan | 0 | 0 | NA | 0 | 0 | NA | 84.61 | 0.6 | 1.16 |
| AG-313-D21 | GORG | 0 | 0 | NA | 1 | 0 | orphan | 0 | 0 | NA | 0 | 0 | NA | 81.93 | 0.6 | 1.27 |

| | | | | | | | | | | | | | | | | |
|---|---|---|---|---|---|---|---|---|---|---|---|---|---|---|---|---|
| AG-910-G09 | GORG | 0 | 0 | NA | 3 | 0 | orphan | 0 | 0 | NA | 0 | 0 | NA | 81.33 | 0.6 | 1.25 |
| AG-911-L10 | GORG | 0 | 0 | NA | 2 | 0 | orphan | 0 | 0 | NA | 0 | 0 | NA | 80.72 | 0.6 | 1.24 |
| AG-912-N03 | GORG | 0 | 0 | NA | 1 | 0 | orphan | 0 | 0 | NA | 0 | 0 | NA | 79.52 | 0.6 | 1.01 |
| AG-918-C13 | GORG | 0 | 0 | NA | 2 | 0 | orphan | 0 | 1 | NA | 0 | 0 | NA | 77.71 | 0.6 | 1.24 |
| AG-920-J13 | GORG | 0 | 0 | NA | 1 | 0 | orphan | 0 | 0 | NA | 0 | 2 | orphan | 75.3 | 0.6 | 1.23 |
| AH-326-D14 | GORG | 0 | 0 | NA | 1 | 0 | orphan | 0 | 0 | NA | 0 | 1 | orphan | 73.59 | 0.6 | 0.82 |
| AG-891-D20 | GORG | 0 | 0 | NA | 1 | 0 | orphan | 0 | 0 | NA | 0 | 0 | NA | 73.49 | 0.6 | 0.99 |
| AG-911-A07 | GORG | 0 | 0 | NA | 3 | 0 | orphan | 0 | 0 | NA | 0 | 0 | NA | 66.16 | 0.6 | 1.08 |
| AG-918-C10 | GORG | 0 | 0 | NA | 2 | 0 | orphan | 0 | 0 | NA | 0 | 0 | NA | 64.62 | 0.6 | 1.07 |
| AG-414-M07 | GORG | 0 | 0 | NA | 1 | 0 | orphan | 0 | 0 | NA | 0 | 0 | NA | 63.65 | 0.6 | 0.58 |
| AG-325-A16 | GORG | 0 | 0 | NA | 1 | 0 | orphan | 0 | 0 | NA | 0 | 0 | NA | 61.24 | 0.6 | 0.98 |
| AG-365-D19 | GORG | 0 | 0 | NA | 2 | 0 | orphan | 0 | 0 | NA | 0 | 0 | NA | 58.63 | 0.6 | 0.81 |
| AG-895-N06 | GORG | 0 | 0 | NA | 1 | 0 | orphan | 0 | 0 | NA | 0 | 0 | NA | 57.72 | 0.6 | 0.62 |
| AH-326-K23 | GORG | 0 | 0 | NA | 1 | 0 | orphan | 0 | 0 | NA | 0 | 0 | NA | 56.22 | 0.6 | 0.56 |
| AG-896-A01 | GORG | 0 | 0 | NA | 1 | 0 | orphan | 0 | 0 | NA | 0 | 0 | NA | 76.16 | 0.61 | 2.09 |
| AG-426-E05 | GORG | 0 | 0 | NA | 2 | 1 | orphan | 0 | 0 | NA | 0 | 0 | NA | 70.58 | 0.61 | 1.29 |
| AG-430-P06 | GORG | 0 | 0 | NA | 1 | 0 | orphan | 0 | 0 | NA | 0 | 0 | NA | 67.55 | 0.61 | 1.18 |
| AG-447-D03 | GORG | 0 | 0 | NA | 1 | 0 | orphan | 0 | 0 | NA | 0 | 1 | orphan | 66.07 | 0.61 | 1.2 |
| AG-435-A22 | GORG | 0 | 0 | NA | 1 | 0 | orphan | 0 | 0 | NA | 0 | 0 | NA | 65.24 | 0.61 | 1.12 |
| AG-913-B20 | GORG | 0 | 0 | NA | 2 | 0 | orphan | 0 | 0 | NA | 0 | 1 | orphan | 59.4 | 0.61 | 1.13 |
| AG-447-C17 | GORG | 0 | 0 | NA | 1 | 0 | orphan | 0 | 0 | NA | 0 | 1 | orphan | 57.91 | 0.62 | 1.23 |
| AH-321-K16 | GORG | 0 | 0 | NA | 2 | 0 | orphan | 0 | 0 | NA | 0 | 0 | NA | 77.22 | 0.63 | 0.94 |
| AG-911-G05 | GORG | 0 | 0 | NA | 2 | 0 | orphan | 1 | 1 | complete | 0 | 0 | NA | 72.85 | 0.63 | 0.93 |
| AG-905-P20 | GORG | 0 | 0 | NA | 1 | 0 | orphan | 0 | 0 | NA | 0 | 0 | NA | 67.93 | 0.63 | 0.82 |
| AG-435-F10 | GORG | 0 | 0 | NA | 2 | 0 | orphan | 0 | 0 | NA | 0 | 0 | NA | 65.01 | 0.63 | 0.79 |

| | | | | | | | | | | | | | | | | |
|---|---|---|---|---|---|---|---|---|---|---|---|---|---|---|---|---|
| AG-319-D09 | GORG | 0 | 0 | NA | 1 | 1 | complete | 0 | 0 | NA | 0 | 0 | NA | 91.75 | 0.66 | 1.48 |
| AG-897-G20 | GORG | 1 | 1 | complete | 1 | 0 | orphan | 0 | 0 | NA | 0 | 0 | NA | 85.07 | 0.66 | 2.35 |
| AG-913-E23 | GORG | 0 | 1 | NA | 0 | 0 | NA | 0 | 0 | NA | 0 | 0 | NA | 74.35 | 0.66 | 1.36 |
| AG-904-O05 | GORG | 0 | 0 | NA | 1 | 0 | orphan | 0 | 0 | NA | 0 | 0 | NA | 67.59 | 0.66 | 1.99 |
| AG-910-B21 | GORG | 0 | 0 | NA | 1 | 0 | orphan | 0 | 0 | NA | 0 | 0 | NA | 75.14 | 0.67 | 1.93 |
| AG-439-G03 | GORG | 0 | 0 | NA | 1 | 0 | orphan | 0 | 0 | NA | 0 | 0 | NA | 71.07 | 0.68 | 3.16 |
| AG-439-O23 | GORG | 0 | 1 | NA | 0 | 0 | NA | 0 | 0 | NA | 0 | 0 | NA | 56.04 | 0.68 | 2.18 |
| AG-422-E10 | GORG | 0 | 0 | NA | 2 | 0 | orphan | 0 | 0 | NA | 0 | 0 | NA | 51.09 | 0.68 | 1.07 |
| AG-390-I23 | GORG | 0 | 0 | NA | 1 | 0 | orphan | 0 | 0 | NA | 0 | 0 | NA | 66.76 | 0.7 | 1.5 |
| AG-343-P19 | GORG | 0 | 0 | NA | 0 | 1 | NA | 0 | 0 | NA | 0 | 0 | NA | 50.46 | 0.7 | 0.82 |
| AG-447-M05 | GORG | 1 | 1 | complete | 2 | 0 | orphan | 0 | 0 | NA | 0 | 0 | NA | 96.54 | 0.71 | 1.4 |
| AG-918-L07 | GORG | 1 | 1 | complete | 1 | 0 | orphan | 0 | 0 | NA | 0 | 0 | NA | 91.32 | 0.71 | 1.18 |
| AG-390-D02 | GORG | 0 | 0 | NA | 2 | 0 | orphan | 0 | 0 | NA | 0 | 0 | NA | 88.41 | 0.71 | 1.1 |
| AG-917-O20 | GORG | 0 | 0 | NA | 1 | 0 | orphan | 0 | 0 | NA | 0 | 0 | NA | 86.73 | 0.71 | 1.11 |
| AG-453-I04 | GORG | 0 | 0 | NA | 1 | 0 | orphan | 0 | 0 | NA | 0 | 0 | NA | 79.86 | 0.71 | 1.12 |
| AG-447-N15 | GORG | 0 | 0 | NA | 2 | 0 | orphan | 0 | 0 | NA | 0 | 0 | NA | 79.29 | 0.71 | 0.94 |
| AG-447-J03 | GORG | 0 | 0 | NA | 0 | 0 | NA | 0 | 0 | NA | 0 | 1 | orphan | 79.15 | 0.71 | 0.97 |
| AG-910-F03 | GORG | 0 | 0 | NA | 1 | 0 | orphan | 0 | 0 | NA | 0 | 1 | orphan | 78.64 | 0.71 | 0.96 |
| AG-325-A21 | GORG | 1 | 1 | complete | 4 | 0 | orphan | 0 | 0 | NA | 0 | 0 | NA | 76.78 | 0.71 | 0.97 |
| AG-337-P04 | GORG | 0 | 0 | NA | 1 | 0 | orphan | 0 | 0 | NA | 0 | 0 | NA | 75.86 | 0.71 | 0.81 |
| AG-337-I11 | GORG | 1 | 2 | complete | 6 | 0 | orphan | 0 | 0 | NA | 0 | 0 | NA | 67.63 | 0.71 | 1.88 |
| AG-908-K15 | GORG | 0 | 0 | NA | 1 | 0 | orphan | 0 | 0 | NA | 0 | 0 | NA | 64.01 | 0.71 | 0.94 |
| AG-430-P14 | GORG | 0 | 0 | NA | 0 | 0 | NA | 0 | 0 | NA | 0 | 1 | orphan | 70.15 | 0.73 | 2.01 |
| AG-916-A19 | GORG | 1 | 1 | complete | 0 | 0 | NA | 0 | 0 | NA | 0 | 0 | NA | 90.81 | 0.74 | 1.77 |

| | | | | | | | | | | | | | | | | |
|---|---|---|---|---|---|---|---|---|---|---|---|---|---|---|---|---|
| AG-891-F08 | GORG | 1 | 1 | complete | 0 | 0 | NA | 0 | 0 | NA | 0 | 0 | NA | 88.97 | 0.74 | 1.43 |
| AG-892-K21 | GORG | 1 | 1 | complete | 0 | 0 | NA | 0 | 0 | NA | 0 | 0 | NA | 87.13 | 0.74 | 1.79 |
| AG-896-F13 | GORG | 0 | 0 | NA | 1 | 0 | orphan | 0 | 1 | NA | 0 | 0 | NA | 86.05 | 0.74 | 1.91 |
| AG-916-D20 | GORG | 0 | 0 | NA | 1 | 0 | orphan | 0 | 0 | NA | 0 | 0 | NA | 74.18 | 0.74 | 1.68 |
| AG-913-J07 | GORG | 1 | 1 | complete | 0 | 0 | NA | 0 | 0 | NA | 0 | 0 | NA | 71.98 | 0.74 | 1.52 |
| AG-447-D22 | GORG | 0 | 0 | NA | 1 | 0 | orphan | 0 | 0 | NA | 0 | 0 | NA | 71.69 | 0.74 | 1.39 |
| AG-898-G05 | GORG | 0 | 0 | NA | 1 | 0 | orphan | 0 | 0 | NA | 0 | 0 | NA | 70.96 | 0.74 | 1.01 |
| AG-453-N14 | GORG | 0 | 0 | NA | 1 | 0 | orphan | 0 | 0 | NA | 0 | 0 | NA | 70.8 | 0.74 | 1.66 |
| AG-896-K13 | GORG | 1 | 1 | complete | 0 | 0 | NA | 0 | 0 | NA | 0 | 0 | NA | 69.68 | 0.74 | 2.13 |
| AG-918-B02 | GORG | 0 | 0 | NA | 1 | 0 | orphan | 0 | 0 | NA | 0 | 0 | NA | 68.99 | 0.74 | 1.52 |
| AG-390-P11 | GORG | 0 | 0 | NA | 1 | 0 | orphan | 0 | 0 | NA | 0 | 0 | NA | 67.93 | 0.74 | 1.31 |
| AG-343-K03 | GORG | 0 | 0 | NA | 4 | 0 | orphan | 0 | 0 | NA | 0 | 0 | NA | 65.21 | 0.74 | 1.17 |
| AG-920-D21 | GORG | 0 | 0 | NA | 1 | 0 | orphan | 0 | 0 | NA | 0 | 0 | NA | 65.01 | 0.74 | 1.37 |
| AG-900-J21 | GORG | 0 | 0 | NA | 2 | 0 | orphan | 0 | 0 | NA | 0 | 0 | NA | 64.58 | 0.74 | 1.14 |
| AG-892-F18 | GORG | 1 | 1 | complete | 0 | 0 | NA | 0 | 0 | NA | 0 | 0 | NA | 62.39 | 0.74 | 1.39 |
| AG-894-D21 | GORG | 0 | 0 | NA | 1 | 0 | orphan | 0 | 0 | NA | 0 | 0 | NA | 61.93 | 0.74 | 1.07 |
| AG-891-G11 | GORG | 0 | 0 | NA | 2 | 0 | orphan | 0 | 0 | NA | 0 | 0 | NA | 60.62 | 0.74 | 1.08 |
| AG-892-G04 | GORG | 0 | 0 | NA | 0 | 0 | NA | 0 | 0 | NA | 0 | 1 | orphan | 59.06 | 0.74 | 1.07 |
| AG-914-N21 | GORG | 1 | 1 | complete | 0 | 0 | NA | 0 | 0 | NA | 0 | 0 | NA | 57.72 | 0.74 | 0.93 |
| AG-893-N17 | GORG | 1 | 1 | complete | 0 | 0 | NA | 0 | 0 | NA | 0 | 0 | NA | 55.17 | 0.74 | 1.16 |
| AG-908-B09 | GORG | 0 | 0 | NA | 1 | 0 | orphan | 0 | 0 | NA | 0 | 0 | NA | 54.68 | 0.74 | 1.32 |
| AG-899-P07 | GORG | 0 | 0 | NA | 1 | 0 | orphan | 0 | 0 | NA | 0 | 0 | NA | 63.16 | 0.75 | 1.34 |
| AG-918-G13 | GORG | 0 | 0 | NA | 1 | 1 | complete | 0 | 0 | NA | 0 | 0 | NA | 67.5 | 0.76 | 1.61 |
| AG-390-F02 | GORG | 2 | 2 | complete | 0 | 0 | NA | 0 | 0 | NA | 0 | 0 | NA | 72.96 | 0.79 | 2.34 |

| | | | | | | | | | | | | | | | | |
|---|---|---|---|---|---|---|---|---|---|---|---|---|---|---|---|---|
| AG-439-D13 | GORG | 0 | 0 | NA | 1 | 0 | orphan | 0 | 0 | NA | 0 | 0 | NA | 62.72 | 0.79 | 0.81 |
| AG-422-O22 | GORG | 0 | 0 | NA | 1 | 0 | orphan | 0 | 0 | NA | 0 | 0 | NA | 78.13 | 0.8 | 1.34 |
| AG-895-B04 | GORG | 0 | 0 | NA | 1 | 0 | orphan | 0 | 0 | NA | 0 | 0 | NA | 64.94 | 0.8 | 1.83 |
| AG-447-B06 | GORG | 0 | 0 | NA | 1 | 0 | orphan | 0 | 0 | NA | 0 | 0 | NA | 61.73 | 0.8 | 1.46 |
| AG-898-K23 | GORG | 1 | 0 | orphan | 1 | 0 | orphan | 0 | 0 | NA | 0 | 0 | NA | 59.67 | 0.8 | 1.45 |
| AG-453-K15 | GORG | 0 | 0 | NA | 2 | 1 | complete | 0 | 0 | NA | 0 | 0 | NA | 58.98 | 0.8 | 1.7 |
| AG-905-N21 | GORG | 0 | 0 | NA | 1 | 0 | orphan | 0 | 0 | NA | 0 | 0 | NA | 57.93 | 0.8 | 1.28 |
| AG-916-P14 | GORG | 0 | 0 | NA | 1 | 0 | orphan | 0 | 0 | NA | 0 | 0 | NA | 76.06 | 0.82 | 1.23 |
| AG-470-J15 | GORG | 0 | 0 | NA | 0 | 0 | NA | 0 | 0 | NA | 0 | 1 | orphan | 65.62 | 0.82 | 1 |
| AG-905-I06 | GORG | 1 | 0 | orphan | 1 | 0 | orphan | 0 | 0 | NA | 0 | 0 | NA | 55.8 | 0.82 | 0.87 |
| AH-326-E05 | GORG | 1 | 0 | orphan | 0 | 0 | NA | 0 | 0 | NA | 0 | 0 | NA | 63.22 | 0.83 | 1.39 |
| AG-447-C23 | GORG | 0 | 0 | NA | 1 | 0 | orphan | 0 | 0 | NA | 0 | 0 | NA | 59.12 | 0.83 | 1.09 |
| AG-430-N19 | GORG | 0 | 0 | NA | 1 | 0 | orphan | 0 | 0 | NA | 0 | 0 | NA | 57.31 | 0.83 | 1.13 |
| AG-913-K07 | GORG | 0 | 0 | NA | 2 | 0 | orphan | 0 | 1 | NA | 0 | 2 | orphan | 78.63 | 0.85 | 1.12 |
| AG-414-E04 | GORG | 0 | 0 | NA | 2 | 0 | orphan | 0 | 0 | NA | 0 | 0 | NA | 72.84 | 0.85 | 0.98 |
| AG-365-B02 | GORG | 1 | 0 | orphan | 0 | 0 | NA | 0 | 0 | NA | 0 | 0 | NA | 70.65 | 0.85 | 1.67 |
| AG-439-A17 | GORG | 0 | 0 | NA | 4 | 0 | orphan | 0 | 0 | NA | 0 | 0 | NA | 69.77 | 0.85 | 1.02 |
| AH-324-P01 | GORG | 0 | 0 | NA | 2 | 0 | orphan | 0 | 0 | NA | 0 | 0 | NA | 66.76 | 0.85 | 0.96 |
| AG-343-C08 | GORG | 0 | 0 | NA | 2 | 0 | orphan | 0 | 0 | NA | 0 | 0 | NA | 64.76 | 0.85 | 0.86 |
| AG-920-F19 | GORG | 1 | 1 | complete | 1 | 0 | orphan | 0 | 0 | NA | 0 | 2 | orphan | 64.1 | 0.85 | 1.05 |
| AG-343-I08 | GORG | 0 | 0 | NA | 2 | 0 | orphan | 0 | 0 | NA | 0 | 0 | NA | 62.25 | 0.85 | 1.15 |
| AG-410-G14 | GORG | 0 | 1 | NA | 0 | 0 | NA | 0 | 0 | NA | 0 | 0 | NA | 61.97 | 0.85 | 1.46 |
| AH-321-N21 | GORG | 0 | 0 | NA | 2 | 0 | orphan | 0 | 0 | NA | 0 | 0 | NA | 57.93 | 0.85 | 0.88 |
| AG-313-C18 | GORG | 0 | 0 | NA | 1 | 0 | orphan | 0 | 0 | NA | 0 | 0 | NA | 57.09 | 0.85 | 0.89 |
| AG-905-B08 | GORG | 0 | 0 | NA | 1 | 0 | orphan | 0 | 0 | NA | 0 | 0 | NA | 55.34 | 0.85 | 0.8 |
| AG-447-N10 | GORG | 0 | 0 | NA | 2 | 0 | orphan | 0 | 0 | NA | 0 | 0 | NA | 54.91 | 0.85 | 0.9 |

| | | | | | | | | | | | | | | | | |
|---|---|---|---|---|---|---|---|---|---|---|---|---|---|---|---|---|
| AG-360-C06 | GORG | 0 | 0 | NA | 1 | 0 | orphan | 0 | 0 | NA | 0 | 0 | NA | 54.81 | 0.85 | 0.83 |
| AG-464-B09 | GORG | 0 | 0 | NA | 2 | 0 | orphan | 0 | 0 | NA | 0 | 0 | NA | 53.87 | 0.85 | 0.71 |
| AG-422-O17 | GORG | 0 | 0 | NA | 2 | 0 | orphan | 0 | 0 | NA | 0 | 0 | NA | 53.77 | 0.85 | 0.87 |
| AG-905-D04 | GORG | 0 | 0 | NA | 2 | 0 | orphan | 0 | 0 | NA | 0 | 2 | orphan | 76.37 | 0.87 | 1.74 |
| AG-919-E18 | GORG | 0 | 1 | NA | 3 | 0 | orphan | 0 | 0 | NA | 0 | 0 | NA | 64.78 | 0.87 | 1.53 |
| AG-901-J20 | GORG | 0 | 0 | NA | 1 | 0 | orphan | 0 | 0 | NA | 0 | 1 | orphan | 58.52 | 0.87 | 1.67 |
| AG-894-J17 | GORG | 1 | 1 | complete | 1 | 0 | orphan | 0 | 0 | NA | 0 | 0 | NA | 65.14 | 0.88 | 2.14 |
| AG-899-B02 | GORG | 0 | 0 | NA | 1 | 0 | orphan | 0 | 0 | NA | 0 | 0 | NA | 83.83 | 0.93 | 2.11 |
| AG-911-I18 | GORG | 1 | 1 | complete | 1 | 0 | orphan | 0 | 0 | NA | 0 | 0 | NA | 63.7 | 0.93 | 1.53 |
| AG-910-G06 | GORG | 0 | 0 | NA | 1 | 0 | orphan | 0 | 0 | NA | 0 | 0 | NA | 69.81 | 0.94 | 1.16 |
| AG-899-G14 | GORG | 0 | 0 | NA | 1 | 0 | orphan | 0 | 0 | NA | 0 | 0 | NA | 66.04 | 0.94 | 0.91 |
| AG-897-I16 | GORG | 0 | 0 | NA | 2 | 0 | orphan | 0 | 0 | NA | 0 | 0 | NA | 58.49 | 0.94 | 0.95 |
| AG-430-B09 | GORG | 0 | 0 | NA | 1 | 0 | orphan | 0 | 0 | NA | 0 | 0 | NA | 96.21 | 0.95 | 1.16 |
| AG-896-C18 | GORG | 0 | 0 | NA | 2 | 0 | orphan | 0 | 0 | NA | 0 | 0 | NA | 92.7 | 0.95 | 1.08 |
| AG-390-N03 | GORG | 0 | 0 | NA | 1 | 0 | orphan | 0 | 0 | NA | 0 | 0 | NA | 78.75 | 0.95 | 0.92 |
| AG-447-D04 | GORG | 0 | 0 | NA | 2 | 0 | orphan | 0 | 0 | NA | 0 | 0 | NA | 78.27 | 0.95 | 0.97 |
| AG-422-C07 | GORG | 0 | 0 | NA | 1 | 0 | orphan | 0 | 0 | NA | 0 | 0 | NA | 76.21 | 0.95 | 0.9 |
| AG-916-O23 | GORG | 0 | 0 | NA | 1 | 0 | orphan | 1 | 1 | complete | 0 | 0 | NA | 75.99 | 0.95 | 0.97 |
| AG-390-N17 | GORG | 0 | 0 | NA | 1 | 0 | orphan | 0 | 0 | NA | 0 | 0 | NA | 73.04 | 0.95 | 0.91 |
| AG-893-N09 | GORG | 0 | 0 | NA | 2 | 0 | orphan | 0 | 0 | NA | 0 | 0 | NA | 72.79 | 0.95 | 0.89 |
| AG-333-C06 | GORG | 0 | 0 | NA | 3 | 0 | orphan | 0 | 0 | NA | 0 | 0 | NA | 66.59 | 0.95 | 0.97 |
| AG-904-I09 | GORG | 0 | 0 | NA | 1 | 0 | orphan | 0 | 0 | NA | 0 | 0 | NA | 65.76 | 0.95 | 0.79 |
| AH-326-O05 | GORG | 1 | 1 | complete | 1 | 0 | orphan | 0 | 0 | NA | 0 | 0 | NA | 59.45 | 0.95 | 0.7 |
| AG-904-G11 | GORG | 0 | 0 | NA | 1 | 0 | orphan | 0 | 0 | NA | 0 | 0 | NA | 58.93 | 0.95 | 0.72 |
| AG-325-C04 | GORG | 0 | 0 | NA | 1 | 0 | orphan | 0 | 0 | NA | 0 | 0 | NA | 58.14 | 0.95 | 0.74 |

| | | | | | | | | | | | | | | | | |
|---|---|---|---|---|---|---|---|---|---|---|---|---|---|---|---|---|
| AG-313-J19 | GORG | 0 | 0 | NA | 1 | 0 | orphan | 0 | 0 | NA | 0 | 0 | NA | 56.01 | 0.95 | 0.72 |
| AG-349-N17 | GORG | 1 | 1 | complete | 2 | 1 | orphan | 0 | 0 | NA | 0 | 0 | NA | 53.54 | 0.95 | 1.28 |
| AG-360-C15 | GORG | 0 | 0 | NA | 0 | 0 | NA | 2 | 1 | complete | 0 | 0 | NA | 51.66 | 0.95 | 0.66 |
| AG-898-J11 | GORG | 0 | 0 | NA | 1 | 0 | orphan | 0 | 0 | NA | 0 | 0 | NA | 50.42 | 0.95 | 0.51 |
| AG-464-F05 | GORG | 0 | 0 | NA | 2 | 0 | orphan | 0 | 0 | NA | 0 | 0 | NA | 53.8 | 0.97 | 0.57 |
| AG-426-G08 | GORG | 0 | 0 | NA | 1 | 0 | orphan | 0 | 0 | NA | 0 | 0 | NA | 61.48 | 1.01 | 1.54 |
| AG-404-B10 | GORG | 0 | 0 | NA | 1 | 0 | orphan | 0 | 0 | NA | 0 | 0 | NA | 58.73 | 1.01 | 2.75 |
| AG-461-L13 | GORG | 1 | 1 | complete | 0 | 0 | NA | 0 | 0 | NA | 0 | 0 | NA | 55.76 | 1.01 | 1.28 |
| AG-390-E03 | GORG | 0 | 0 | NA | 3 | 0 | orphan | 0 | 0 | NA | 0 | 0 | NA | 66.29 | 1.02 | 1.71 |
| AG-470-E04 | GORG | 0 | 0 | NA | 1 | 0 | orphan | 0 | 0 | NA | 0 | 0 | NA | 64.88 | 1.02 | 2.76 |
| AH-321-C20 | GORG | 1 | 1 | complete | 0 | 0 | NA | 0 | 0 | NA | 0 | 0 | NA | 78.17 | 1.09 | 1.55 |
| AG-901-P11 | GORG | 0 | 0 | NA | 1 | 0 | orphan | 0 | 0 | NA | 0 | 0 | NA | 70.87 | 1.09 | 1.43 |
| AG-911-C05 | GORG | 0 | 0 | NA | 0 | 0 | NA | 0 | 0 | NA | 0 | 1 | orphan | 56.88 | 1.09 | 0.95 |
| AG-430-E04 | GORG | 0 | 0 | NA | 0 | 0 | NA | 0 | 0 | NA | 0 | 1 | orphan | 51.63 | 1.09 | 0.83 |
| AG-349-G20 | GORG | 0 | 0 | NA | 1 | 0 | orphan | 0 | 1 | NA | 0 | 0 | NA | 92.31 | 1.1 | 1.42 |
| AG-892-J04 | GORG | 0 | 0 | NA | 1 | 1 | complete | 0 | 0 | NA | 0 | 0 | NA | 90.11 | 1.1 | 1.26 |
| AG-892-F10 | GORG | 0 | 0 | NA | 0 | 0 | NA | 0 | 0 | NA | 0 | 2 | orphan | 89.01 | 1.1 | 1.43 |
| AG-895-B20 | GORG | 0 | 0 | NA | 2 | 0 | orphan | 0 | 0 | NA | 0 | 0 | NA | 88.62 | 1.1 | 1.33 |
| AG-404-M02 | GORG | 0 | 0 | NA | 2 | 1 | complete | 0 | 0 | NA | 0 | 0 | NA | 85.71 | 1.1 | 1.27 |
| AG-896-A16 | GORG | 0 | 0 | NA | 2 | 0 | orphan | 0 | 0 | NA | 0 | 0 | NA | 83.88 | 1.1 | 1.33 |
| AG-899-D02 | GORG | 0 | 0 | NA | 1 | 0 | orphan | 0 | 0 | NA | 0 | 0 | NA | 83.75 | 1.1 | 1.23 |
| AG-359-F09 | GORG | 0 | 0 | NA | 2 | 0 | orphan | 0 | 0 | NA | 0 | 0 | NA | 80.96 | 1.1 | 1.21 |
| AG-337-K15 | GORG | 1 | 1 | complete | 0 | 0 | NA | 0 | 0 | NA | 0 | 0 | NA | 80.95 | 1.1 | 1.26 |
| AG-390-K05 | GORG | 0 | 0 | NA | 2 | 1 | orphan | 0 | 0 | NA | 0 | 0 | NA | 73.08 | 1.1 | 1.31 |

| | | | | | | | | | | | | | | | | |
|---|---|---|---|---|---|---|---|---|---|---|---|---|---|---|---|---|
| AG-914-I17 | GORG | 0 | 0 | NA | 0 | 0 | NA | 0 | 0 | NA | 0 | 1 | orphan | 72.43 | 1.1 | 1.07 |
| AG-919-E11 | GORG | 0 | 0 | NA | 1 | 0 | orphan | 0 | 0 | NA | 0 | 0 | NA | 72.43 | 1.1 | 1.31 |
| AG-453-C09 | GORG | 0 | 0 | NA | 1 | 0 | orphan | 0 | 0 | NA | 0 | 0 | NA | 68.74 | 1.1 | 1.03 |
| AG-426-B06 | GORG | 1 | 1 | complete | 0 | 0 | NA | 0 | 0 | NA | 0 | 0 | NA | 65.38 | 1.1 | 1.08 |
| AG-365-O11 | GORG | 0 | 0 | NA | 1 | 0 | orphan | 0 | 0 | NA | 0 | 0 | NA | 60.99 | 1.1 | 1.21 |
| AH-287-P08 | GORG | 0 | 0 | NA | 2 | 0 | orphan | 0 | 0 | NA | 0 | 0 | NA | 59.92 | 1.1 | 0.97 |
| AG-916-G23 | GORG | 0 | 0 | NA | 1 | 1 | complete | 0 | 0 | NA | 0 | 0 | NA | 59.34 | 1.1 | 1.1 |
| AG-919-A04 | GORG | 0 | 0 | NA | 1 | 0 | orphan | 0 | 0 | NA | 0 | 0 | NA | 56.79 | 1.1 | 1.07 |
| AG-916-N07 | GORG | 0 | 0 | NA | 2 | 0 | orphan | 0 | 0 | NA | 0 | 0 | NA | 54.62 | 1.1 | 0.9 |
| AG-359-L02 | GORG | 0 | 0 | NA | 3 | 1 | orphan | 0 | 0 | NA | 0 | 0 | NA | 51.45 | 1.1 | 1.07 |
| AG-895-J23 | GORG | 1 | 1 | complete | 1 | 0 | orphan | 0 | 0 | NA | 0 | 0 | NA | 75.72 | 1.11 | 1.01 |
| AG-349-A14 | GORG | 1 | 0 | orphan | 0 | 0 | NA | 0 | 0 | NA | 0 | 0 | NA | 55.1 | 1.11 | 2.06 |
| AG-895-A04 | GORG | 1 | 1 | complete | 0 | 0 | NA | 0 | 0 | NA | 0 | 0 | NA | 70.82 | 1.13 | 1.09 |
| AG-360-D18 | GORG | 0 | 0 | NA | 1 | 0 | orphan | 0 | 0 | NA | 0 | 0 | NA | 57.82 | 1.15 | 0.86 |
| AG-325-C15 | GORG | 0 | 0 | NA | 1 | 0 | orphan | 0 | 0 | NA | 0 | 0 | NA | 55.37 | 1.15 | 2.12 |
| AH-324-O11 | GORG | 0 | 0 | NA | 2 | 0 | orphan | 0 | 0 | NA | 0 | 0 | NA | 62.74 | 1.16 | 1.32 |
| AH-287-A14 | GORG | 0 | 0 | NA | 1 | 0 | orphan | 0 | 0 | NA | 0 | 1 | orphan | 63.51 | 1.18 | 0.96 |
| AG-893-L13 | GORG | 1 | 1 | complete | 2 | 0 | orphan | 0 | 0 | NA | 0 | 0 | NA | 92.88 | 1.2 | 1.08 |
| AG-447-O15 | GORG | 0 | 0 | NA | 1 | 0 | orphan | 0 | 0 | NA | 0 | 0 | NA | 90.61 | 1.2 | 1.47 |
| AG-414-B08 | GORG | 0 | 0 | NA | 1 | 0 | orphan | 0 | 0 | NA | 0 | 0 | NA | 89.16 | 1.2 | 1.13 |
| AG-343-K02 | GORG | 0 | 0 | NA | 2 | 0 | orphan | 0 | 0 | NA | 0 | 0 | NA | 86.86 | 1.2 | 0.97 |
| AG-909-G09 | GORG | 1 | 0 | orphan | 1 | 0 | orphan | 0 | 0 | NA | 0 | 0 | NA | 86.75 | 1.2 | 1.26 |
| AG-439-C22 | GORG | 0 | 0 | NA | 1 | 0 | orphan | 0 | 0 | NA | 0 | 0 | NA | 85.98 | 1.2 | 1.04 |
| AG-892-O03 | GORG | 0 | 0 | NA | 1 | 0 | orphan | 0 | 0 | NA | 0 | 0 | NA | 82.28 | 1.2 | 1 |
| AG-414-F13 | GORG | 0 | 0 | NA | 2 | 0 | orphan | 0 | 0 | NA | 0 | 0 | NA | 81.05 | 1.2 | 0.92 |

| | | | | | | | | | | | | | | | | |
|---|---|---|---|---|---|---|---|---|---|---|---|---|---|---|---|---|
| AG-917-C10 | GORG | 0 | 0 | NA | 1 | 0 | orphan | 0 | 0 | NA | 0 | 0 | NA | 74.04 | 1.2 | 0.95 |
| AG-919-I20 | GORG | 0 | 0 | NA | 1 | 0 | orphan | 1 | 1 | complete | 0 | 0 | NA | 72.89 | 1.2 | 0.91 |
| AG-414-B07 | GORG | 0 | 0 | NA | 1 | 0 | orphan | 0 | 0 | NA | 0 | 0 | NA | 71.08 | 1.2 | 0.81 |
| AG-447-C07 | GORG | 0 | 0 | NA | 2 | 0 | orphan | 0 | 0 | NA | 0 | 0 | NA | 68.4 | 1.2 | 1.33 |
| AG-430-P03 | GORG | 0 | 0 | NA | 1 | 0 | orphan | 0 | 0 | NA | 0 | 0 | NA | 66.27 | 1.2 | 1.1 |
| AG-365-F10 | GORG | 0 | 0 | NA | 2 | 0 | orphan | 0 | 0 | NA | 0 | 0 | NA | 62.65 | 1.2 | 0.86 |
| AG-439-C02 | GORG | 0 | 0 | NA | 1 | 0 | orphan | 0 | 0 | NA | 0 | 0 | NA | 60.24 | 1.2 | 0.84 |
| AG-414-B10 | GORG | 0 | 0 | NA | 1 | 0 | orphan | 0 | 0 | NA | 0 | 0 | NA | 59.46 | 1.2 | 0.96 |
| AG-360-G06 | GORG | 0 | 0 | NA | 1 | 0 | orphan | 0 | 0 | NA | 0 | 0 | NA | 57.6 | 1.2 | 0.74 |
| AG-319-K04 | GORG | 0 | 0 | NA | 1 | 0 | orphan | 0 | 0 | NA | 0 | 0 | NA | 57.05 | 1.2 | 0.48 |
| AG-414-N11 | GORG | 0 | 0 | NA | 1 | 0 | orphan | 0 | 0 | NA | 0 | 0 | NA | 56.63 | 1.2 | 0.71 |
| AG-453-E06 | GORG | 0 | 0 | NA | 1 | 0 | orphan | 0 | 0 | NA | 0 | 0 | NA | 55.17 | 1.2 | 0.67 |
| AG-435-M09 | GORG | 0 | 0 | NA | 1 | 0 | orphan | 0 | 0 | NA | 0 | 0 | NA | 54.39 | 1.2 | 0.8 |
| AG-414-L14 | GORG | 0 | 0 | NA | 1 | 0 | orphan | 0 | 0 | NA | 0 | 0 | NA | 54.22 | 1.2 | 0.87 |
| AG-439-P17 | GORG | 0 | 0 | NA | 1 | 0 | orphan | 0 | 0 | NA | 0 | 0 | NA | 51.2 | 1.2 | 0.61 |
| AG-349-J18 | GORG | 0 | 0 | NA | 2 | 1 | complete | 0 | 0 | NA | 0 | 0 | NA | 73.62 | 1.21 | 1.92 |
| AG-426-I05 | GORG | 0 | 0 | NA | 0 | 1 | NA | 0 | 0 | NA | 0 | 0 | NA | 79.98 | 1.22 | 1.41 |
| AG-410-K11 | GORG | 0 | 0 | NA | 0 | 0 | NA | 0 | 0 | NA | 0 | 1 | orphan | 83.2 | 1.25 | 3.14 |
| AG-917-O09 | GORG | 0 | 0 | NA | 1 | 0 | orphan | 0 | 0 | NA | 0 | 0 | NA | 77.78 | 1.28 | 1.1 |
| AG-426-G06 | GORG | 1 | 1 | complete | 0 | 0 | NA | 0 | 0 | NA | 0 | 0 | NA | 75.21 | 1.28 | 1.7 |
| AG-447-P23 | GORG | 0 | 0 | NA | 2 | 0 | orphan | 0 | 0 | NA | 0 | 0 | NA | 75.21 | 1.28 | 1.1 |
| AG-911-M23 | GORG | 0 | 0 | NA | 2 | 0 | orphan | 0 | 0 | NA | 0 | 0 | NA | 74.36 | 1.28 | 1 |
| AG-447-I21 | GORG | 0 | 0 | NA | 2 | 0 | orphan | 0 | 0 | NA | 0 | 0 | NA | 73.5 | 1.28 | 1.07 |
| AH-287-M22 | GORG | 0 | 0 | NA | 1 | 0 | orphan | 0 | 0 | NA | 0 | 0 | NA | 73.5 | 1.28 | 1.07 |
| AG-426-F13 | GORG | 0 | 0 | NA | 3 | 0 | orphan | 1 | 1 | complete | 0 | 0 | NA | 73.08 | 1.28 | 0.99 |

| | | | | | | | | | | | | | | | | |
|---|---|---|---|---|---|---|---|---|---|---|---|---|---|---|---|---|
| AG-343-C03 | GORG | 0 | 0 | NA | 2 | 0 | orphan | 0 | 0 | NA | 0 | 0 | NA | 72.22 | 1.28 | 0.96 |
| AG-426-L21 | GORG | 0 | 0 | NA | 1 | 0 | orphan | 0 | 0 | NA | 0 | 0 | NA | 71.42 | 1.28 | 0.99 |
| AG-404-B23 | GORG | 0 | 0 | NA | 1 | 0 | orphan | 0 | 0 | NA | 0 | 0 | NA | 70.3 | 1.28 | 1.01 |
| AG-453-L02 | GORG | 0 | 0 | NA | 2 | 0 | orphan | 0 | 0 | NA | 0 | 0 | NA | 70.09 | 1.28 | 1.02 |
| AG-916-D18 | GORG | 0 | 0 | NA | 2 | 0 | orphan | 0 | 0 | NA | 0 | 0 | NA | 70.09 | 1.28 | 1.03 |
| AH-324-G14 | GORG | 0 | 0 | NA | 1 | 0 | orphan | 0 | 0 | NA | 0 | 0 | NA | 69.23 | 1.28 | 0.93 |
| AG-919-L19 | GORG | 0 | 0 | NA | 1 | 0 | orphan | 0 | 0 | NA | 0 | 0 | NA | 68.9 | 1.28 | 1.04 |
| AG-917-L07 | GORG | 0 | 0 | NA | 1 | 0 | orphan | 0 | 0 | NA | 0 | 0 | NA | 67.52 | 1.28 | 1.01 |
| AG-447-N19 | GORG | 1 | 1 | complete | 1 | 0 | orphan | 0 | 0 | NA | 0 | 0 | NA | 67.37 | 1.28 | 1.05 |
| AG-911-M08 | GORG | 0 | 0 | NA | 1 | 0 | orphan | 0 | 0 | NA | 0 | 0 | NA | 67.18 | 1.28 | 1.61 |
| AG-435-L19 | GORG | 2 | 1 | complete | 2 | 0 | orphan | 0 | 0 | NA | 0 | 0 | NA | 67.09 | 1.28 | 1.62 |
| AG-313-L14 | GORG | 0 | 0 | NA | 3 | 0 | orphan | 0 | 0 | NA | 0 | 0 | NA | 66.91 | 1.28 | 0.98 |
| AG-439-P07 | GORG | 1 | 1 | complete | 1 | 0 | orphan | 0 | 0 | NA | 0 | 0 | NA | 66.67 | 1.28 | 1.29 |
| AG-390-B19 | GORG | 0 | 0 | NA | 1 | 0 | orphan | 0 | 0 | NA | 0 | 0 | NA | 65.81 | 1.28 | 1.01 |
| AH-326-P07 | GORG | 1 | 1 | complete | 1 | 0 | orphan | 0 | 0 | NA | 0 | 0 | NA | 64.96 | 1.28 | 0.98 |
| AG-343-L11 | GORG | 0 | 0 | NA | 2 | 0 | orphan | 0 | 0 | NA | 0 | 0 | NA | 61.69 | 1.28 | 1.48 |
| AH-287-A16 | GORG | 0 | 0 | NA | 3 | 0 | orphan | 0 | 0 | NA | 0 | 0 | NA | 60.26 | 1.28 | 0.99 |
| AG-410-B14 | GORG | 2 | 1 | complete | 1 | 0 | orphan | 0 | 0 | NA | 0 | 0 | NA | 59.83 | 1.28 | 1.35 |
| AG-447-D11 | GORG | 0 | 0 | NA | 4 | 0 | orphan | 0 | 0 | NA | 0 | 0 | NA | 59.6 | 1.28 | 0.91 |
| AG-343-B13 | GORG | 2 | 1 | complete | 2 | 0 | orphan | 0 | 0 | NA | 0 | 0 | NA | 57.66 | 1.28 | 1.36 |
| AG-349-M08 | GORG | 1 | 1 | complete | 1 | 0 | orphan | 0 | 0 | NA | 0 | 0 | NA | 57.26 | 1.28 | 0.89 |
| AG-457-E08 | GORG | 0 | 0 | NA | 1 | 0 | orphan | 0 | 0 | NA | 0 | 0 | NA | 56.92 | 1.28 | 0.86 |
| AG-470-E13 | GORG | 0 | 0 | NA | 3 | 0 | orphan | 0 | 0 | NA | 0 | 0 | NA | 56.74 | 1.28 | 0.85 |
| AG-313-C08 | GORG | 0 | 0 | NA | 1 | 0 | orphan | 0 | 0 | NA | 0 | 0 | NA | 56.41 | 1.28 | 0.9 |
| AH-321-N15 | GORG | 0 | 0 | NA | 1 | 0 | orphan | 0 | 0 | NA | 0 | 0 | NA | 55.13 | 1.28 | 0.91 |

| | | | | | | | | | | | | | | | | |
|---|---|---|---|---|---|---|---|---|---|---|---|---|---|---|---|---|
| AG-313-I09 | GORG | 0 | 0 | NA | 2 | 0 | orphan | 0 | 0 | NA | 0 | 0 | NA | 54.7 | 1.28 | 0.83 |
| AG-447-C10 | GORG | 0 | 0 | NA | 2 | 0 | orphan | 0 | 0 | NA | 0 | 0 | NA | 54.04 | 1.28 | 0.92 |
| AG-365-N16 | GORG | 0 | 0 | NA | 1 | 0 | orphan | 0 | 0 | NA | 0 | 0 | NA | 53.77 | 1.28 | 0.85 |
| AG-447-M08 | GORG | 0 | 0 | NA | 2 | 0 | orphan | 0 | 0 | NA | 0 | 0 | NA | 53.7 | 1.28 | 0.91 |
| AH-287-I15 | GORG | 0 | 0 | NA | 1 | 0 | orphan | 0 | 0 | NA | 0 | 0 | NA | 53.48 | 1.28 | 0.91 |
| AG-439-N21 | GORG | 0 | 0 | NA | 3 | 0 | orphan | 0 | 0 | NA | 0 | 0 | NA | 52.99 | 1.28 | 0.85 |
| AG-414-M13 | GORG | 0 | 0 | NA | 2 | 0 | orphan | 0 | 0 | NA | 0 | 0 | NA | 50.85 | 1.28 | 0.94 |
| AH-324-A03 | GORG | 0 | 0 | NA | 2 | 0 | orphan | 0 | 0 | NA | 0 | 0 | NA | 50.43 | 1.28 | 0.88 |
| AG-410-B17 | GORG | 1 | 1 | complet e | 2 | 0 | orphan | 1 | 1 | complet e | 0 | 0 | NA | 80.25 | 1.3 | 1.76 |
| AG-349-E15 | GORG | 0 | 0 | NA | 1 | 0 | orphan | 0 | 0 | NA | 0 | 0 | NA | 60.68 | 1.3 | 1.92 |
| AG-917-P14 | GORG | 1 | 0 | orphan | 1 | 0 | orphan | 0 | 0 | NA | 0 | 0 | NA | 80.37 | 1.31 | 1.03 |
| AG-439-N08 | GORG | 0 | 1 | NA | 1 | 0 | orphan | 0 | 0 | NA | 0 | 0 | NA | 53.01 | 1.31 | 0.59 |
| AG-390-K07 | GORG | 0 | 0 | NA | 1 | 1 | complet e | 0 | 0 | NA | 0 | 0 | NA | 57.21 | 1.32 | 1.14 |
| AG-907-D18 | GORG | 0 | 0 | NA | 1 | 0 | orphan | 0 | 0 | NA | 0 | 0 | NA | 81.08 | 1.35 | 1.23 |
| AG-447-E05 | GORG | 0 | 0 | NA | 1 | 0 | orphan | 0 | 0 | NA | 0 | 0 | NA | 58.24 | 1.35 | 2.36 |
| AG-337-J14 | GORG | 1 | 0 | orphan | 4 | 0 | orphan | 0 | 0 | NA | 0 | 1 | orpha n | 51.53 | 1.35 | 2.22 |
| AG-918-G06 | GORG | 0 | 0 | NA | 2 | 0 | orphan | 0 | 0 | NA | 0 | 0 | NA | 96.51 | 1.42 | 1.21 |
| AG-917-C11 | GORG | 0 | 0 | NA | 2 | 0 | orphan | 0 | 0 | NA | 0 | 0 | NA | 83.79 | 1.42 | 1.13 |
| AG-365-K10 | GORG | 0 | 0 | NA | 1 | 0 | orphan | 0 | 0 | NA | 0 | 0 | NA | 78.66 | 1.42 | 0.91 |
| AG-917-B08 | GORG | 0 | 0 | NA | 1 | 0 | orphan | 0 | 0 | NA | 0 | 0 | NA | 67.54 | 1.42 | 0.87 |
| AH-324-I08 | GORG | 1 | 1 | complet e | 1 | 0 | orphan | 0 | 0 | NA | 0 | 0 | NA | 66.36 | 1.42 | 0.93 |
| AG-912-K07 | GORG | 0 | 0 | NA | 0 | 0 | NA | 0 | 0 | NA | 0 | 1 | orpha n | 55.28 | 1.42 | 0.62 |
| AG-914-M08 | GORG | 0 | 0 | NA | 1 | 0 | orphan | 0 | 0 | NA | 0 | 0 | NA | 53.58 | 1.42 | 0.74 |
| AG-337-P03 | GORG | 0 | 0 | NA | 1 | 0 | orphan | 0 | 0 | NA | 0 | 0 | NA | 62.38 | 1.46 | 0.78 |
| AG-900-G19 | GORG | 0 | 0 | NA | 0 | 0 | NA | 0 | 0 | NA | 0 | 2 | orpha n | 89.25 | 1.47 | 1.65 |

| | | | | | | | | | | | | | | | | |
|---|---|---|---|---|---|---|---|---|---|---|---|---|---|---|---|---|
| AG-447-P07 | GORG | 0 | 0 | NA | 1 | 0 | orphan | 0 | 0 | NA | 0 | 0 | NA | 54.1 | 1.58 | 1.16 |
| AG-426-D04 | GORG | 1 | 1 | complet e | 1 | 0 | orphan | 0 | 0 | NA | 0 | 1 | orpha n | 64.09 | 1.6 | 0.95 |
| AG-360-E10 | GORG | 0 | 0 | NA | 1 | 0 | orphan | 0 | 0 | NA | 0 | 0 | NA | 63.4 | 1.64 | 0.92 |
| AG-453-J21 | GORG | 0 | 0 | NA | 0 | 0 | NA | 0 | 0 | NA | 0 | 2 | orpha n | 79.88 | 1.71 | 1.75 |
| AG-319-N07 | GORG | 0 | 1 | NA | 1 | 0 | orphan | 0 | 0 | NA | 0 | 1 | orpha n | 57.34 | 1.71 | 1.25 |
| AG-410-M04 | GORG | 0 | 0 | NA | 2 | 0 | orphan | 0 | 0 | NA | 0 | 0 | NA | 65.52 | 1.72 | 1.93 |
| AG-892-I13 | GORG | 0 | 0 | NA | 1 | 0 | orphan | 0 | 0 | NA | 0 | 0 | NA | 63.79 | 1.72 | 1.55 |
| AG-359-I23 | GORG | 0 | 0 | NA | 1 | 0 | orphan | 0 | 0 | NA | 0 | 0 | NA | 58.93 | 1.72 | 1.55 |
| AG-896-K09 | GORG | 0 | 0 | NA | 2 | 0 | orphan | 0 | 0 | NA | 0 | 0 | NA | 57.76 | 1.72 | 1.08 |
| AG-891-J15 | GORG | 0 | 0 | NA | 1 | 0 | orphan | 0 | 0 | NA | 0 | 0 | NA | 56.9 | 1.72 | 1.37 |
| AG-904-E10 | GORG | 0 | 0 | NA | 2 | 0 | orphan | 0 | 0 | NA | 0 | 1 | orpha n | 55.17 | 1.72 | 1.13 |
| AG-333-F19 | GORG | 0 | 0 | NA | 1 | 1 | complet e | 0 | 0 | NA | 0 | 0 | NA | 51.61 | 1.72 | 0.7 |
| AG-914-K05 | GORG | 0 | 0 | NA | 2 | 0 | orphan | 0 | 0 | NA | 0 | 0 | NA | 63.16 | 1.75 | 1.08 |
| AG-894-B03 | GORG | 0 | 0 | NA | 2 | 0 | orphan | 0 | 0 | NA | 0 | 0 | NA | 60.53 | 1.75 | 1.03 |
| AG-912-G23 | GORG | 0 | 1 | NA | 0 | 0 | NA | 0 | 0 | NA | 0 | 0 | NA | 60.53 | 1.75 | 0.94 |
| AG-892-N22 | GORG | 0 | 0 | NA | 1 | 0 | orphan | 0 | 0 | NA | 0 | 0 | NA | 59.65 | 1.75 | 0.93 |
| AG-895-J07 | GORG | 1 | 1 | complet e | 1 | 0 | orphan | 0 | 0 | NA | 0 | 0 | NA | 50.88 | 1.75 | 1.33 |
| AG-896-P20 | GORG | 0 | 0 | NA | 1 | 0 | orphan | 0 | 0 | NA | 0 | 0 | NA | 50.88 | 1.75 | 0.76 |
| AG-319-C10 | GORG | 0 | 0 | NA | 1 | 0 | orphan | 0 | 0 | NA | 0 | 0 | NA | 61.64 | 1.76 | 1.47 |
| AG-337-E14 | GORG | 0 | 0 | NA | 1 | 0 | orphan | 0 | 0 | NA | 0 | 0 | NA | 73.56 | 1.8 | 1.95 |
| AG-422-B10 | GORG | 0 | 0 | NA | 2 | 0 | orphan | 0 | 0 | NA | 0 | 0 | NA | 73.3 | 1.81 | 0.89 |
| AG-900-E05 | GORG | 0 | 0 | NA | 1 | 0 | orphan | 0 | 0 | NA | 0 | 0 | NA | 57.78 | 1.81 | 0.72 |
| AG-919-G03 | GORG | 0 | 0 | NA | 1 | 0 | orphan | 0 | 0 | NA | 0 | 0 | NA | 84.19 | 1.83 | 1.18 |
| AG-898-K13 | GORG | 0 | 0 | NA | 0 | 0 | NA | 0 | 0 | NA | 0 | 1 | orpha n | 60.38 | 1.89 | 0.78 |
| AG-319-F14 | GORG | 0 | 0 | NA | 1 | 0 | orphan | 0 | 0 | NA | 0 | 0 | NA | 58.49 | 1.89 | 0.86 |

| | | | | | | | | | | | | | | | | |
|---|---|---|---|---|---|---|---|---|---|---|---|---|---|---|---|---|
| AG-901-J13 | GORG | 0 | 0 | NA | 1 | 0 | orphan | 0 | 0 | NA | 0 | 0 | NA | 58.49 | 1.89 | 0.89 |
| AG-913-E02 | GORG | 1 | 0 | orphan | 1 | 0 | orphan | 0 | 0 | NA | 0 | 0 | NA | 57.29 | 1.89 | 1.05 |
| AH-287-E22 | GORG | 0 | 0 | NA | 2 | 0 | orphan | 0 | 0 | NA | 0 | 0 | NA | 54.4 | 1.89 | 0.76 |
| AG-919-A16 | GORG | 0 | 0 | NA | 1 | 0 | orphan | 0 | 0 | NA | 0 | 0 | NA | 52.83 | 1.89 | 0.59 |
| AG-905-P04 | GORG | 0 | 0 | NA | 2 | 0 | orphan | 0 | 0 | NA | 0 | 0 | NA | 52.39 | 1.89 | 0.81 |
| AG-908-J11 | GORG | 0 | 0 | NA | 1 | 0 | orphan | 0 | 0 | NA | 0 | 0 | NA | 50.94 | 1.89 | 0.59 |
| AG-897-O11 | GORG | 1 | 1 | complete | 1 | 0 | orphan | 0 | 0 | NA | 0 | 1 | orphan | 81.28 | 1.9 | 1.09 |
| AG-917-A09 | GORG | 0 | 0 | NA | 2 | 0 | orphan | 0 | 0 | NA | 0 | 0 | NA | 80.82 | 1.9 | 1.07 |
| AG-470-M06 | GORG | 0 | 0 | NA | 1 | 0 | orphan | 0 | 0 | NA | 0 | 0 | NA | 65.88 | 1.9 | 0.76 |
| AG-333-A19 | GORG | 0 | 0 | NA | 2 | 0 | orphan | 0 | 0 | NA | 0 | 0 | NA | 64.84 | 1.9 | 0.68 |
| AG-414-C19 | GORG | 0 | 0 | NA | 2 | 0 | orphan | 0 | 0 | NA | 0 | 0 | NA | 58.54 | 1.9 | 0.88 |
| AG-903-G03 | GORG | 0 | 0 | NA | 1 | 0 | orphan | 0 | 0 | NA | 0 | 0 | NA | 55.46 | 1.9 | 0.64 |
| AG-918-O17 | GORG | 0 | 0 | NA | 0 | 0 | NA | 0 | 0 | NA | 0 | 1 | orphan | 70.34 | 1.97 | 1.22 |
| AG-453-E14 | GORG | 0 | 0 | NA | 1 | 0 | orphan | 0 | 0 | NA | 0 | 0 | NA | 71.31 | 2.11 | 1.86 |
| AG-470-B18 | GORG | 0 | 0 | NA | 1 | 0 | orphan | 0 | 0 | NA | 0 | 0 | NA | 51 | 2.2 | 0.82 |
| AG-319-F15 | GORG | 0 | 0 | NA | 2 | 0 | orphan | 0 | 0 | NA | 0 | 0 | NA | 65.42 | 2.26 | 2.39 |
| AG-414-D07 | GORG | 0 | 0 | NA | 1 | 0 | orphan | 0 | 0 | NA | 0 | 0 | NA | 88.59 | 2.41 | 0.99 |
| AG-337-I10 | GORG | 0 | 0 | NA | 2 | 0 | orphan | 0 | 0 | NA | 0 | 0 | NA | 87.14 | 2.41 | 1.06 |
| AG-349-K05 | GORG | 0 | 0 | NA | 1 | 0 | orphan | 0 | 0 | NA | 0 | 1 | orphan | 77.11 | 2.41 | 1.04 |
| AG-414-A22 | GORG | 1 | 1 | complete | 1 | 0 | orphan | 0 | 0 | NA | 0 | 0 | NA | 75.13 | 2.41 | 1.19 |
| AG-333-C07 | GORG | 0 | 0 | NA | 1 | 0 | orphan | 0 | 0 | NA | 0 | 0 | NA | 69.16 | 2.41 | 1 |
| AG-343-N07 | GORG | 0 | 0 | NA | 1 | 0 | orphan | 0 | 0 | NA | 0 | 0 | NA | 59.04 | 2.52 | 0.75 |
| AG-414-N07 | GORG | 0 | 0 | NA | 1 | 0 | orphan | 0 | 0 | NA | 0 | 0 | NA | 67.67 | 2.61 | 0.83 |
| AG-900-I20 | GORG | 0 | 0 | NA | 1 | 0 | orphan | 0 | 0 | NA | 0 | 0 | NA | 58.62 | 2.68 | 0.84 |
| AG-410-G07 | GORG | 0 | 0 | NA | 2 | 0 | orphan | 0 | 0 | NA | 0 | 0 | NA | 57.03 | 2.99 | 0.92 |
| AG-439-N22 | GORG | 0 | 0 | NA | 2 | 0 | orphan | 0 | 0 | NA | 0 | 0 | NA | 59.12 | 3.2 | 0.97 |

| | | | | | | | | | | | | | | | | |
|---|---|---|---|---|---|---|---|---|---|---|---|---|---|---|---|---|
| AG-414-F21 | GORG | 0 | 0 | NA | 2 | 0 | orphan | 0 | 0 | NA | 0 | 0 | NA | 82.01 | 3.61 | 1.12 |
| AG-410-O08 | GORG | 0 | 0 | NA | 2 | 0 | orphan | 0 | 0 | NA | 0 | 0 | NA | 67.47 | 3.61 | 0.92 |
| AG-464-D14 | GORG | 0 | 0 | NA | 1 | 0 | orphan | 0 | 0 | NA | 0 | 0 | NA | 51.89 | 3.77 | 0.57 |
| AG-410-P15 | GORG | 0 | 0 | NA | 3 | 0 | orphan | 0 | 0 | NA | 0 | 0 | NA | 78.08 | 4.82 | 1.2 |
| AG-410-K17 | GORG | 0 | 0 | NA | 1 | 0 | orphan | 0 | 0 | NA | 0 | 0 | NA | 65.05 | 6.25 | 0.9 |
| AG-897-N06 | GORG | 0 | 0 | NA | 1 | 0 | orphan | 0 | 0 | NA | 0 | 2 | orphan | 65.14 | 6.72 | 1.18 |
| AG-410-M07 | GORG | 0 | 0 | NA | 1 | 0 | orphan | 0 | 0 | NA | 0 | 0 | NA | 56.96 | 7.23 | 0.67 |
| AG-337-D16 | GORG | 0 | 0 | NA | 1 | 0 | orphan | 0 | 0 | NA | 0 | 0 | NA | 86.21 | 9.43 | 1.2 |

RM system: Restriction-modification system; M: modification; R: restriction; TARA: Tara Oceans, SHIPPO: Shipborne Pole-to-Pole Observations; GORG: Global Ocean Reference Genomes

**table S3. RM system of SHIPPO MAGs**

| RM system | Family | Type I | | | Type II | | | Type III | | | Type IV | | |
|---|---|---|---|---|---|---|---|---|---|---|---|---|---|
| | | M | R | system | M | R | system | M | R | system | M | R | system |
| SHIPPO_EUK_1 | Bathycoccaceae | 0 | 0 | NA | 4 | 0 | orphan | 0 | 0 | NA | 0 | 0 | NA |
| SHIPPO_EUK_2 | Mamiellaceae | 0 | 0 | NA | 5 | 0 | orphan | 0 | 0 | NA | 0 | 0 | NA |
| SHIPPO_EUK_3 | Bathycoccaceae | 0 | 0 | NA | 3 | 0 | orphan | 0 | 0 | NA | 0 | 0 | NA |
| SHIPPO_EUK_4 | Mamiellaceae | 0 | 0 | NA | 8 | 0 | orphan | 0 | 0 | NA | 0 | 0 | NA |
| SHIPPO_EUK_5 | Mamiellaceae | 0 | 0 | NA | 5 | 0 | orphan | 0 | 0 | NA | 0 | 0 | NA |
| SHIPPO_EUK_6 | Bathycoccaceae | 0 | 0 | NA | 5 | 0 | orphan | 0 | 0 | NA | 0 | 0 | NA |
| SHIPPO_PRO_101 | MGIIA | 0 | 0 | NA | 1 | 0 | orphan | 0 | 0 | NA | 0 | 0 | NA |
| SHIPPO_PRO_102 | MGIIA | 0 | 0 | NA | 1 | 0 | orphan | 0 | 0 | NA | 0 | 0 | NA |
| SHIPPO_PRO_103 | MGIIA | 0 | 0 | NA | 1 | 0 | orphan | 0 | 0 | NA | 0 | 0 | NA |
| SHIPPO_PRO_104 | MGIIA | 0 | 0 | NA | 1 | 0 | orphan | 0 | 0 | NA | 0 | 0 | NA |
| SHIPPO_PRO_105 | MGIIA | 0 | 0 | NA | 1 | 0 | orphan | 0 | 0 | NA | 0 | 0 | NA |
| SHIPPO_PRO_127 | UBA1268 | 0 | 0 | NA | 1 | 0 | orphan | 0 | 0 | NA | 0 | 0 | NA |
| SHIPPO_PRO_128 | UBA1268 | 0 | 0 | NA | 1 | 0 | orphan | 0 | 0 | NA | 0 | 0 | NA |
| SHIPPO_PRO_13 | Emcibacteraceae | 0 | 0 | NA | 1 | 0 | orphan | 0 | 0 | NA | 0 | 0 | NA |
| SHIPPO_PRO_130 | Nisaeaceae | 0 | 0 | NA | 2 | 0 | orphan | 0 | 0 | NA | 0 | 0 | NA |
| SHIPPO_PRO_132 | Microbacteriaceae | 0 | 0 | NA | 1 | 0 | orphan | 0 | 0 | NA | 0 | 0 | NA |
| SHIPPO_PRO_135 | UBA955 | 1 | 0 | orphan | 0 | 0 | NA | 0 | 0 | NA | 0 | 0 | NA |
| SHIPPO_PRO_136 | Flavobacteriaceae | 0 | 1 | NA | 0 | 0 | NA | 0 | 0 | NA | 0 | 0 | NA |
| SHIPPO_PRO_140 | Flavobacteriaceae | 0 | 1 | NA | 0 | 0 | NA | 0 | 0 | NA | 0 | 0 | NA |
| SHIPPO_PRO_141 | Flavobacteriaceae | 0 | 0 | NA | 1 | 0 | orphan | 0 | 0 | NA | 0 | 0 | NA |
| SHIPPO_PRO_143 | Flavobacteriaceae | 0 | 0 | NA | 2 | 0 | orphan | 0 | 0 | NA | 0 | 0 | NA |
| SHIPPO_PRO_147 | BACL11 | 1 | 1 | orphan | 0 | 0 | NA | 0 | 0 | NA | 0 | 0 | NA |
| SHIPPO_PRO_152 | Flavobacteriaceae | 0 | 0 | NA | 4 | 1 | complete | 0 | 0 | NA | 0 | 0 | NA |

| | | | | | | | | | | | | | |
|---|---|---|---|---|---|---|---|---|---|---|---|---|---|
| SHIPPO_PRO_16 | Cyanobiaceae | 0 | 0 | NA | 1 | 0 | orphan | 0 | 0 | NA | 0 | 0 | NA |
| SHIPPO_PRO_161 | unclassified_Pelagibacterales | 0 | 0 | NA | 1 | 0 | orphan | 0 | 0 | NA | 0 | 0 | NA |
| SHIPPO_PRO_164 | Flavobacteriaceae | 0 | 0 | NA | 0 | 1 | NA | 0 | 0 | NA | 0 | 0 | NA |
| SHIPPO_PRO_17 | Rhodobacteraceae | 0 | 0 | NA | 1 | 0 | orphan | 0 | 0 | NA | 0 | 0 | NA |
| SHIPPO_PRO_171 | UBA8229 | 0 | 0 | NA | 5 | 0 | orphan | 0 | 0 | NA | 0 | 2 | orphan |
| SHIPPO_PRO_173 | UBA9320 | 0 | 0 | NA | 2 | 0 | orphan | 0 | 0 | NA | 0 | 0 | NA |
| SHIPPO_PRO_174 | UBA9320 | 0 | 0 | NA | 1 | 0 | orphan | 0 | 0 | NA | 0 | 0 | NA |
| SHIPPO_PRO_176 | Rhodobacteraceae | 0 | 0 | NA | 2 | 0 | orphan | 0 | 0 | NA | 0 | 0 | NA |
| SHIPPO_PRO_177 | koll-22 | 0 | 0 | NA | 0 | 0 | NA | 1 | 0 | orphan | 0 | 0 | NA |
| SHIPPO_PRO_179 | UBA7430 | 0 | 2 | NA | 0 | 0 | NA | 0 | 0 | NA | 0 | 0 | NA |
| SHIPPO_PRO_180 | 1G12 | 0 | 0 | NA | 1 | 0 | orphan | 0 | 0 | NA | 0 | 0 | NA |
| SHIPPO_PRO_181 | unclassified_Fibrobacterales | 0 | 1 | NA | 0 | 0 | NA | 0 | 0 | NA | 0 | 1 | orphan |
| SHIPPO_PRO_184 | MGIIA | 0 | 0 | NA | 2 | 0 | orphan | 0 | 0 | NA | 0 | 0 | NA |
| SHIPPO_PRO_185 | MGIIA | 0 | 0 | NA | 3 | 0 | orphan | 0 | 0 | NA | 0 | 0 | NA |
| SHIPPO_PRO_186 | MGIIA | 0 | 0 | NA | 1 | 0 | orphan | 0 | 0 | NA | 0 | 0 | NA |
| SHIPPO_PRO_187 | MGIIA | 0 | 0 | NA | 1 | 0 | orphan | 0 | 0 | NA | 0 | 0 | NA |
| SHIPPO_PRO_188 | MGIIB | 0 | 0 | NA | 1 | 0 | orphan | 0 | 0 | NA | 0 | 0 | NA |
| SHIPPO_PRO_194 | unclassified_Nanopelagicales | 0 | 3 | NA | 1 | 0 | orphan | 0 | 0 | NA | 0 | 0 | NA |
| SHIPPO_PRO_195 | HIMB59 | 0 | 0 | NA | 1 | 0 | orphan | 0 | 0 | NA | 0 | 0 | NA |
| SHIPPO_PRO_198 | Emcibacteraceae | 0 | 0 | NA | 1 | 0 | orphan | 0 | 0 | NA | 0 | 0 | NA |
| SHIPPO_PRO_2 | Burkholderiaceae | 0 | 0 | NA | 1 | 0 | orphan | 0 | 0 | NA | 0 | 0 | NA |
| SHIPPO_PRO_200 | Saprospiraceae | 0 | 0 | NA | 1 | 0 | orphan | 0 | 0 | NA | 0 | 0 | NA |
| SHIPPO_PRO_202 | TMED189 | 2 | 1 | complete | 3 | 0 | orphan | 0 | 1 | NA | 0 | 2 | orphan |
| SHIPPO_PRO_203 | UBA1268 | 0 | 0 | NA | 1 | 0 | orphan | 0 | 0 | NA | 0 | 0 | NA |
| SHIPPO_PRO_204 | Flavobacteriaceae | 0 | 0 | NA | 1 | 0 | orphan | 0 | 0 | NA | 0 | 0 | NA |

| | | | | | | | | | | | | | |
|---|---|---|---|---|---|---|---|---|---|---|---|---|---|
| SHIPPO_PRO_21 | Rhodobacteraceae | 0 | 0 | NA | 1 | 0 | orphan | 0 | 0 | NA | 0 | 0 | NA |
| SHIPPO_PRO_210 | Flavobacteriaceae | 0 | 1 | NA | 5 | 0 | orphan | 0 | 1 | NA | 0 | 2 | orpha n |
| SHIPPO_PRO_22 | Rhodobacteraceae | 0 | 0 | NA | 1 | 0 | orphan | 0 | 0 | NA | 0 | 0 | NA |
| SHIPPO_PRO_224 | Flavobacteriaceae | 1 | 0 | orphan | 0 | 0 | NA | 0 | 0 | NA | 0 | 0 | NA |
| SHIPPO_PRO_225 | Flavobacteriaceae | 0 | 0 | NA | 1 | 0 | orphan | 0 | 0 | NA | 0 | 0 | NA |
| SHIPPO_PRO_227 | Flavobacteriaceae | 1 | 0 | orphan | 0 | 0 | NA | 0 | 0 | NA | 0 | 0 | NA |
| SHIPPO_PRO_23 | Rhodobacteraceae | 0 | 0 | NA | 2 | 0 | orphan | 0 | 0 | NA | 0 | 0 | NA |
| SHIPPO_PRO_233 | GCA-002716945 | 1 | 2 | orphan | 0 | 0 | NA | 0 | 0 | NA | 0 | 0 | NA |
| SHIPPO_PRO_238 | Rhodobacteraceae | 0 | 0 | NA | 1 | 0 | orphan | 0 | 0 | NA | 0 | 0 | NA |
| SHIPPO_PRO_240 | AAA536-G10 | 0 | 0 | NA | 2 | 0 | orphan | 0 | 0 | NA | 0 | 0 | NA |
| SHIPPO_PRO_241 | AAA536-G10 | 0 | 0 | NA | 1 | 0 | orphan | 0 | 0 | NA | 0 | 0 | NA |
| SHIPPO_PRO_242 | MB11C04 | 0 | 0 | NA | 5 | 0 | orphan | 0 | 0 | NA | 0 | 0 | NA |
| SHIPPO_PRO_244 | Puniceicoccaceae | 0 | 0 | NA | 1 | 0 | orphan | 0 | 0 | NA | 0 | 0 | NA |
| SHIPPO_PRO_245 | Pelagibacteraceae | 0 | 0 | NA | 2 | 0 | orphan | 0 | 0 | NA | 0 | 0 | NA |
| SHIPPO_PRO_246 | Pelagibacteraceae | 0 | 0 | NA | 1 | 0 | orphan | 0 | 0 | NA | 0 | 0 | NA |
| SHIPPO_PRO_248 | Pelagibacteraceae | 0 | 0 | NA | 1 | 0 | orphan | 0 | 0 | NA | 0 | 0 | NA |
| SHIPPO_PRO_249 | Puniceispirillaceae | 0 | 0 | NA | 1 | 0 | orphan | 0 | 0 | NA | 0 | 0 | NA |
| SHIPPO_PRO_253 | Rhodobacteraceae | 0 | 0 | NA | 6 | 0 | orphan | 0 | 0 | NA | 0 | 0 | NA |
| SHIPPO_PRO_254 | MB11C04 | 0 | 0 | NA | 7 | 0 | orphan | 0 | 0 | NA | 0 | 0 | NA |
| SHIPPO_PRO_262 | D2472 | 0 | 0 | NA | 0 | 1 | NA | 0 | 0 | NA | 0 | 0 | NA |
| SHIPPO_PRO_263 | D2472 | 0 | 0 | NA | 0 | 1 | NA | 0 | 0 | NA | 0 | 0 | NA |
| SHIPPO_PRO_265 | D2472 | 0 | 0 | NA | 0 | 1 | NA | 0 | 0 | NA | 0 | 0 | NA |
| SHIPPO_PRO_27 | MB11C04 | 0 | 1 | NA | 2 | 0 | orphan | 0 | 0 | NA | 0 | 0 | NA |
| SHIPPO_PRO_272 | Thioglobaceae | 0 | 0 | NA | 2 | 0 | orphan | 0 | 0 | NA | 0 | 0 | NA |
| SHIPPO_PRO_277 | Puniceispirillaceae | 0 | 1 | NA | 1 | 0 | orphan | 0 | 0 | NA | 0 | 0 | NA |
| SHIPPO_PRO_278 | Woeseiaceae | 0 | 0 | NA | 2 | 0 | orphan | 0 | 0 | NA | 0 | 0 | NA |
| SHIPPO_PRO_279 | Rhodobacteraceae | 0 | 0 | NA | 1 | 0 | orphan | 0 | 0 | NA | 0 | 0 | NA |

| | | | | | | | | | | | | | |
|---|---|---|---|---|---|---|---|---|---|---|---|---|---|
| SHIPPO_PRO_280 | Rhodobacteraceae | 0 | 0 | NA | 1 | 0 | orphan | 0 | 0 | NA | 0 | 0 | NA |
| SHIPPO_PRO_281 | Rhodobacteraceae | 0 | 0 | NA | 1 | 0 | orphan | 0 | 0 | NA | 0 | 0 | NA |
| SHIPPO_PRO_282 | Rhodobacteraceae | 0 | 0 | NA | 2 | 0 | orphan | 0 | 0 | NA | 0 | 1 | orphan |
| SHIPPO_PRO_283 | Rhodobacteraceae | 0 | 0 | NA | 1 | 0 | orphan | 0 | 0 | NA | 0 | 0 | NA |
| SHIPPO_PRO_284 | Rhodobacteraceae | 0 | 0 | NA | 1 | 0 | orphan | 0 | 0 | NA | 0 | 0 | NA |
| SHIPPO_PRO_285 | HTCC2089 | 1 | 0 | orphan | 0 | 0 | NA | 0 | 0 | NA | 0 | 0 | NA |
| SHIPPO_PRO_287 | Casp-alpha2 | 0 | 0 | NA | 2 | 0 | orphan | 0 | 0 | NA | 0 | 0 | NA |
| SHIPPO_PRO_288 | Methylophagaceae | 0 | 0 | NA | 1 | 0 | orphan | 0 | 0 | NA | 0 | 0 | NA |
| SHIPPO_PRO_289 | Methylophagaceae | 0 | 0 | NA | 1 | 0 | orphan | 0 | 0 | NA | 0 | 0 | NA |
| SHIPPO_PRO_294 | Nitrincolaceae | 0 | 0 | NA | 2 | 0 | orphan | 0 | 0 | NA | 0 | 0 | NA |
| SHIPPO_PRO_30 | MB11C04 | 1 | 0 | orphan | 4 | 0 | orphan | 0 | 0 | NA | 0 | 0 | NA |
| SHIPPO_PRO_33 | Pelagibacteraceae | 0 | 0 | NA | 1 | 0 | orphan | 0 | 0 | NA | 0 | 0 | NA |
| SHIPPO_PRO_49 | HTCC2089 | 0 | 0 | NA | 1 | 0 | orphan | 0 | 0 | NA | 0 | 0 | NA |
| SHIPPO_PRO_5 | Microbacteriaceae | 0 | 0 | NA | 1 | 0 | orphan | 0 | 0 | NA | 0 | 0 | NA |
| SHIPPO_PRO_52 | HTCC2089 | 0 | 0 | NA | 1 | 0 | orphan | 0 | 0 | NA | 0 | 0 | NA |
| SHIPPO_PRO_6 | Rhodobacteraceae | 0 | 0 | NA | 1 | 0 | orphan | 0 | 0 | NA | 0 | 0 | NA |
| SHIPPO_PRO_62 | Ilumatobacteraceae | 0 | 0 | NA | 1 | 0 | orphan | 0 | 0 | NA | 0 | 0 | NA |
| SHIPPO_PRO_63 | Ilumatobacteraceae | 0 | 3 | NA | 1 | 0 | orphan | 0 | 0 | NA | 0 | 0 | NA |
| SHIPPO_PRO_64 | unclassified_Pelagibacterales | 0 | 0 | NA | 1 | 0 | orphan | 0 | 0 | NA | 0 | 0 | NA |
| SHIPPO_PRO_65 | TMED25 | 0 | 0 | NA | 1 | 1 | orphan | 0 | 0 | NA | 0 | 0 | NA |
| SHIPPO_PRO_66 | AAA536-G10 | 0 | 0 | NA | 1 | 0 | orphan | 0 | 0 | NA | 0 | 0 | NA |
| SHIPPO_PRO_67 | TMED109 | 0 | 0 | NA | 1 | 0 | orphan | 0 | 0 | NA | 0 | 0 | NA |
| SHIPPO_PRO_68 | Rhodobacteraceae | 0 | 0 | NA | 1 | 0 | orphan | 0 | 0 | NA | 0 | 0 | NA |
| SHIPPO_PRO_71 | Rhodobacteraceae | 0 | 0 | NA | 1 | 0 | orphan | 0 | 0 | NA | 0 | 0 | NA |
| SHIPPO_PRO_73 | Rhodobacteraceae | 0 | 0 | NA | 1 | 0 | orphan | 0 | 0 | NA | 0 | 0 | NA |
| SHIPPO_PRO_77 | Methylophilaceae | 0 | 0 | NA | 1 | 0 | orphan | 0 | 0 | NA | 0 | 0 | NA |
| SHIPPO_PRO_8 | Rhodobacteraceae | 0 | 0 | NA | 1 | 0 | orphan | 0 | 0 | NA | 0 | 0 | NA |

| | | | | | | | | | | | | | |
|---|---|---|---|---|---|---|---|---|---|---|---|---|---|
| SHIPPO_PRO_82 | SG8-40 | 0 | 0 | NA | 2 | 0 | orphan | 0 | 0 | NA | 0 | 0 | NA |
| SHIPPO_PRO_84 | Rhodobacteraceae | 0 | 0 | NA | 1 | 0 | orphan | 0 | 0 | NA | 0 | 0 | NA |
| SHIPPO_PRO_85 | Rhodobacteraceae | 0 | 0 | NA | 0 | 0 | NA | 0 | 1 | NA | 0 | 0 | NA |
| SHIPPO_PRO_94 | UBA8229 | 0 | 0 | NA | 4 | 0 | orphan | 0 | 0 | NA | 0 | 1 | orphan |
| SHIPPO_PRO_95 | UBA8229 | 0 | 0 | NA | 1 | 0 | orphan | 0 | 0 | NA | 0 | 0 | NA |
| SHIPPO_PRO_98 | MGIIA | 0 | 0 | NA | 1 | 0 | orphan | 0 | 0 | NA | 0 | 0 | NA |
| vOTU_102__Co_k141_2590266 | Myoviridae | 0 | 0 | NA | 1 | 0 | orphan | 0 | 0 | NA | 0 | 0 | NA |
| vOTU_1035__S6_k141_345104 | Podoviridae | 0 | 0 | NA | 1 | 0 | orphan | 0 | 0 | NA | 0 | 0 | NA |
| vOTU_1068__S2_k141_247496 | Myoviridae | 0 | 1 | NA | 2 | 0 | orphan | 0 | 0 | NA | 0 | 0 | NA |
| vOTU_1068__S7_k141_582884 | Myoviridae | 0 | 1 | NA | 2 | 0 | orphan | 0 | 0 | NA | 0 | 0 | NA |
| vOTU_1116__S3_k141_4013 | Unclassified_Caudovirales | 0 | 0 | NA | 1 | 0 | orphan | 0 | 0 | NA | 0 | 0 | NA |
| vOTU_1147__S3_k141_263968 | Myoviridae | 0 | 0 | NA | 0 | 0 | NA | 0 | 1 | NA | 0 | 0 | NA |
| vOTU_1179__S4_k141_575152 | Unclassified_Caudovirales | 0 | 0 | NA | 1 | 0 | orphan | 0 | 0 | NA | 0 | 0 | NA |
| vOTU_1179__S6_k141_111516 | Unclassified_Caudovirales | 0 | 0 | NA | 1 | 0 | orphan | 0 | 0 | NA | 0 | 0 | NA |
| vOTU_1179__S9_k141_552693 | Unclassified_Caudovirales | 0 | 0 | NA | 1 | 0 | orphan | 0 | 0 | NA | 0 | 0 | NA |
| vOTU_1204__S8_k141_182509 | Unknown_Virus | 0 | 0 | NA | 1 | 0 | orphan | 0 | 0 | NA | 0 | 0 | NA |
| vOTU_1274__Co_k141_777427 | Podoviridae | 0 | 0 | NA | 1 | 0 | orphan | 0 | 0 | NA | 0 | 0 | NA |
| vOTU_1274__S5_k141_677773 | Podoviridae | 0 | 0 | NA | 1 | 0 | orphan | 0 | 0 | NA | 0 | 0 | NA |
| vOTU_1277__S10_k141_375502 | Siphoviridae | 0 | 0 | NA | 1 | 0 | orphan | 0 | 0 | NA | 0 | 0 | NA |
| vOTU_1277__S11_k141_507932 | Siphoviridae | 0 | 0 | NA | 1 | 0 | orphan | 0 | 0 | NA | 0 | 0 | NA |
| vOTU_1295__S3_k141_24098 | Podoviridae | 0 | 0 | NA | 1 | 0 | orphan | 0 | 0 | NA | 0 | 0 | NA |
| vOTU_1295__S5_k141_631154 | Podoviridae | 0 | 0 | NA | 1 | 0 | orphan | 0 | 0 | NA | 0 | 0 | NA |
| vOTU_1450__Co_k141_50577 | Myoviridae | 0 | 1 | NA | 1 | 0 | orphan | 0 | 0 | NA | 0 | 0 | NA |
| vOTU_1459__Co_k141_2565134 | Unclassified_Caudovirales | 0 | 0 | NA | 1 | 0 | orphan | 0 | 0 | NA | 0 | 0 | NA |
| vOTU_1476__Co_k141_2212073 | Myoviridae | 0 | 1 | NA | 0 | 0 | NA | 0 | 0 | NA | 0 | 0 | NA |
| vOTU_1503__S2_k141_686267 | Podoviridae | 0 | 0 | NA | 1 | 0 | orphan | 0 | 0 | NA | 0 | 0 | NA |
| vOTU_1571__Co_k141_857091 | Unclassified_Caudovirales | 0 | 0 | NA | 1 | 0 | orphan | 0 | 0 | NA | 0 | 0 | NA |

| | | | | | | | | | | | | | |
|---|---|---|---|---|---|---|---|---|---|---|---|---|---|
| vOTU_1571__S11_k141_282489 | Unclassified_Caudovirales | 0 | 0 | NA | 1 | 0 | orphan | 0 | 0 | NA | 0 | 0 | NA |
| vOTU_1601__S9_k141_259051 | Podoviridae | 0 | 0 | NA | 1 | 0 | orphan | 0 | 0 | NA | 0 | 0 | NA |
| vOTU_1606__Co_k141_194888 | Unclassified_Caudovirales | 0 | 0 | NA | 1 | 0 | orphan | 0 | 0 | NA | 0 | 0 | NA |
| vOTU_1670__Co_k141_2480578 | Podoviridae | 0 | 0 | NA | 1 | 0 | orphan | 0 | 0 | NA | 0 | 0 | NA |
| vOTU_1670__S9_k141_305768 | Podoviridae | 0 | 0 | NA | 1 | 0 | orphan | 0 | 0 | NA | 0 | 0 | NA |
| vOTU_1681__S10_k141_80435 | Myoviridae | 0 | 0 | NA | 1 | 0 | orphan | 0 | 0 | NA | 0 | 0 | NA |
| vOTU_1691__S11_k141_749468 | Podoviridae | 0 | 0 | NA | 1 | 0 | orphan | 0 | 0 | NA | 0 | 0 | NA |
| vOTU_1710__Co_k141_2350775 | Phycodnaviridae | 0 | 0 | NA | 1 | 0 | orphan | 0 | 0 | NA | 0 | 0 | NA |
| vOTU_172__Co_contig_625_pilon_pilon | Myoviridae | 0 | 0 | NA | 1 | 0 | orphan | 0 | 0 | NA | 0 | 0 | NA |
| vOTU_1724__Co_k141_623856 | Unclassified_Caudovirales | 0 | 0 | NA | 1 | 0 | orphan | 0 | 0 | NA | 0 | 0 | NA |
| vOTU_1747__S10_k141_602633 | Myoviridae | 0 | 1 | NA | 0 | 0 | NA | 0 | 0 | NA | 0 | 0 | NA |
| vOTU_1795__S9_k141_96341 | Myoviridae | 0 | 0 | NA | 1 | 0 | orphan | 0 | 0 | NA | 0 | 0 | NA |
| vOTU_1806__S11_k141_279959 | Siphoviridae | 0 | 0 | NA | 1 | 0 | orphan | 0 | 0 | NA | 0 | 0 | NA |
| vOTU_1806__S5_k141_420705 | Siphoviridae | 0 | 0 | NA | 1 | 0 | orphan | 0 | 0 | NA | 0 | 0 | NA |
| vOTU_184__Co_contig_4228_pilon_pilon | Phycodnaviridae | 0 | 0 | NA | 1 | 0 | orphan | 0 | 0 | NA | 0 | 0 | NA |
| vOTU_1888__S2_k141_797290 | Unclassified_Caudovirales | 0 | 0 | NA | 1 | 0 | orphan | 0 | 0 | NA | 0 | 0 | NA |
| vOTU_1888__S5_k141_236639 | Unclassified_Caudovirales | 0 | 0 | NA | 1 | 0 | orphan | 0 | 0 | NA | 0 | 0 | NA |
| vOTU_1894__S10_k141_234113 | Unknown_Virus | 0 | 0 | NA | 1 | 0 | orphan | 0 | 0 | NA | 0 | 0 | NA |
| vOTU_191__S11_k141_573013 | Phycodnaviridae | 0 | 0 | NA | 1 | 0 | orphan | 0 | 0 | NA | 0 | 0 | NA |
| vOTU_1924__S2_k141_339393 | Siphoviridae | 0 | 0 | NA | 1 | 0 | orphan | 0 | 0 | NA | 0 | 0 | NA |
| vOTU_1924__S4_k141_612117 | Siphoviridae | 0 | 0 | NA | 1 | 0 | orphan | 0 | 0 | NA | 0 | 0 | NA |
| vOTU_1931__Co_k141_2192817 | Unknown_Virus | 0 | 0 | NA | 1 | 0 | orphan | 0 | 0 | NA | 0 | 0 | NA |
| vOTU_1945__Co_k141_2547596 | Unknown_Virus | 0 | 0 | NA | 2 | 0 | orphan | 0 | 0 | NA | 0 | 0 | NA |
| vOTU_1959__Co_contig_1736_pilon_pilon | Myoviridae | 0 | 0 | NA | 1 | 0 | orphan | 0 | 0 | NA | 0 | 0 | NA |
| vOTU_1959__Co_contig_6230_pilon_pilon | Myoviridae | 0 | 0 | NA | 1 | 0 | orphan | 0 | 0 | NA | 0 | 0 | NA |
| vOTU_1959__S11_contig_253_pilon_pilon | Myoviridae | 0 | 0 | NA | 1 | 0 | orphan | 0 | 0 | NA | 0 | 0 | NA |

| | | | | | | | | | | | | | |
|---|---|---|---|---|---|---|---|---|---|---|---|---|---|
| vOTU_1981__S2_k141_569071 | Myoviridae | 0 | 0 | NA | 3 | 0 | orphan | 0 | 0 | NA | 0 | 0 | NA |
| vOTU_200__Co_k141_471493 | Myoviridae | 0 | 0 | NA | 1 | 0 | orphan | 0 | 0 | NA | 0 | 0 | NA |
| vOTU_201__Co_contig_1300_pilon_pilon | Siphoviridae | 0 | 0 | NA | 1 | 0 | orphan | 0 | 0 | NA | 0 | 0 | NA |
| vOTU_201__Co_k141_235388 | Siphoviridae | 0 | 0 | NA | 1 | 0 | orphan | 0 | 0 | NA | 0 | 0 | NA |
| vOTU_201__S11_contig_181_pilon_pilon | Siphoviridae | 0 | 0 | NA | 1 | 0 | orphan | 0 | 0 | NA | 0 | 0 | NA |
| vOTU_201__S11_contig_80_pilon_pilon | Siphoviridae | 0 | 0 | NA | 1 | 0 | orphan | 0 | 0 | NA | 0 | 0 | NA |
| vOTU_2011__S2_k141_30249 | Unclassified_Caudovirales | 0 | 0 | NA | 1 | 0 | orphan | 0 | 0 | NA | 0 | 0 | NA |
| vOTU_2012__Co_contig_6405_pilon_pilon | Siphoviridae | 0 | 0 | NA | 1 | 0 | orphan | 0 | 0 | NA | 0 | 0 | NA |
| vOTU_2012__S10_contig_246_pilon_pilon | Siphoviridae | 0 | 0 | NA | 1 | 0 | orphan | 0 | 0 | NA | 0 | 0 | NA |
| vOTU_2094__S2_k141_303417 | Unclassified_Caudovirales | 0 | 0 | NA | 1 | 0 | orphan | 0 | 0 | NA | 0 | 0 | NA |
| vOTU_2103__S6_k141_774823 | Podoviridae | 0 | 1 | NA | 0 | 0 | NA | 0 | 0 | NA | 0 | 0 | NA |
| vOTU_2103__S7_k141_363705 | Podoviridae | 0 | 1 | NA | 0 | 0 | NA | 0 | 0 | NA | 0 | 0 | NA |
| vOTU_216__S8_k141_252089 | Phycodnaviridae | 0 | 0 | NA | 1 | 0 | orphan | 0 | 0 | NA | 0 | 0 | NA |
| vOTU_2175__S4_k141_203219 | Myoviridae | 0 | 0 | NA | 1 | 0 | orphan | 0 | 0 | NA | 0 | 0 | NA |
| vOTU_2175__S5_k141_179830 | Myoviridae | 0 | 0 | NA | 1 | 0 | orphan | 0 | 0 | NA | 0 | 0 | NA |
| vOTU_2234__S6_k141_703493 | Myoviridae | 0 | 0 | NA | 2 | 0 | orphan | 0 | 0 | NA | 0 | 0 | NA |
| vOTU_2234__S8_k141_161249 | Myoviridae | 0 | 0 | NA | 2 | 0 | orphan | 0 | 0 | NA | 0 | 0 | NA |
| vOTU_2255__S10_k141_7922 | Siphoviridae | 0 | 0 | NA | 1 | 0 | orphan | 0 | 0 | NA | 0 | 0 | NA |
| vOTU_2277__Co_contig_4536_pilon_pilon | Podoviridae | 0 | 0 | NA | 1 | 0 | orphan | 0 | 0 | NA | 0 | 0 | NA |
| vOTU_2278__S5_k141_647684 | Unknown_Virus | 0 | 0 | NA | 1 | 0 | orphan | 0 | 0 | NA | 0 | 0 | NA |
| vOTU_2279__S10_k141_536998 | Myoviridae | 0 | 0 | NA | 1 | 0 | orphan | 0 | 0 | NA | 0 | 0 | NA |
| vOTU_2279__S11_k141_218838-circular | Myoviridae | 0 | 0 | NA | 1 | 0 | orphan | 0 | 0 | NA | 0 | 0 | NA |
| vOTU_2326__S3_k141_87451 | Unclassified_Caudovirales | 0 | 1 | NA | 0 | 0 | NA | 0 | 0 | NA | 0 | 0 | NA |
| vOTU_2326__S4_contig_89_pilon_pilon | Unclassified_Caudovirales | 0 | 1 | NA | 0 | 0 | NA | 0 | 0 | NA | 0 | 0 | NA |
| vOTU_2326__S8_contig_227_pilon_pilon | Unclassified_Caudovirales | 0 | 1 | NA | 0 | 0 | NA | 0 | 0 | NA | 0 | 0 | NA |
| vOTU_2371__S10_k141_35579 | Myoviridae | 0 | 0 | NA | 1 | 0 | orphan | 0 | 0 | NA | 0 | 0 | NA |

| | | | | | | | | | | | | | |
|---|---|---|---|---|---|---|---|---|---|---|---|---|---|
| vOTU_2421__S2_k141_254465 | Myoviridae | 0 | 0 | NA | 0 | 1 | NA | 0 | 0 | NA | 0 | 0 | NA |
| vOTU_2457__S11_k141_785167 | Unclassified_Caudovirales | 0 | 0 | NA | 1 | 0 | orphan | 0 | 0 | NA | 0 | 0 | NA |
| vOTU_2516__S6_k141_435094 | Myoviridae | 0 | 0 | NA | 1 | 0 | orphan | 0 | 0 | NA | 0 | 0 | NA |
| vOTU_2518__S6_k141_76422 | Myoviridae | 0 | 1 | NA | 0 | 0 | NA | 0 | 0 | NA | 0 | 0 | NA |
| vOTU_2626__Co_k141_2253228 | Siphoviridae | 0 | 0 | NA | 1 | 0 | orphan | 0 | 0 | NA | 0 | 0 | NA |
| vOTU_2626__S10_k141_164009 | Siphoviridae | 0 | 0 | NA | 1 | 0 | orphan | 0 | 0 | NA | 0 | 0 | NA |
| vOTU_2639__S6_k141_552678 | Phycodnaviridae | 0 | 0 | NA | 1 | 0 | orphan | 0 | 0 | NA | 0 | 0 | NA |
| vOTU_2639__S9_k141_259090 | Phycodnaviridae | 0 | 0 | NA | 1 | 0 | orphan | 0 | 0 | NA | 0 | 0 | NA |
| vOTU_2651__S6_k141_160800 | Podoviridae | 0 | 0 | NA | 1 | 0 | orphan | 0 | 0 | NA | 0 | 0 | NA |
| vOTU_2687__S8_k141_655084 | Myoviridae | 0 | 0 | NA | 1 | 0 | orphan | 0 | 0 | NA | 0 | 0 | NA |
| vOTU_2692__Co_k141_1918461 | Podoviridae | 0 | 0 | NA | 1 | 0 | orphan | 0 | 0 | NA | 0 | 0 | NA |
| vOTU_2692__S10_k141_383214 | Podoviridae | 0 | 0 | NA | 1 | 0 | orphan | 0 | 0 | NA | 0 | 0 | NA |
| vOTU_2702__S11_k141_252820 | Podoviridae | 0 | 0 | NA | 1 | 0 | orphan | 0 | 0 | NA | 0 | 0 | NA |
| vOTU_2731__S10_k141_105741 | Siphoviridae | 0 | 0 | NA | 1 | 0 | orphan | 0 | 0 | NA | 0 | 0 | NA |
| vOTU_2737__S9_k141_171713 | Myoviridae | 0 | 1 | NA | 0 | 0 | NA | 0 | 0 | NA | 0 | 0 | NA |
| vOTU_2757__S9_k141_428776 | Siphoviridae | 0 | 0 | NA | 1 | 0 | orphan | 0 | 0 | NA | 0 | 0 | NA |
| vOTU_2772__Co_k141_1893663 | Myoviridae | 0 | 0 | NA | 1 | 0 | orphan | 0 | 0 | NA | 0 | 0 | NA |
| vOTU_2847__S11_k141_181973 | Unclassified_Caudovirales | 0 | 0 | NA | 1 | 0 | orphan | 0 | 0 | NA | 0 | 0 | NA |
| vOTU_2872__S8_contig_253_pilon_pilon | Phycodnaviridae | 0 | 0 | NA | 1 | 0 | orphan | 0 | 0 | NA | 0 | 0 | NA |
| vOTU_2893__S5_k141_478807 | Myoviridae | 0 | 0 | NA | 1 | 0 | orphan | 0 | 0 | NA | 0 | 0 | NA |
| vOTU_3026__S6_k141_612303 | Myoviridae | 0 | 0 | NA | 1 | 0 | orphan | 0 | 0 | NA | 0 | 0 | NA |
| vOTU_3026__S7_k141_639711 | Myoviridae | 0 | 0 | NA | 1 | 0 | orphan | 0 | 0 | NA | 0 | 0 | NA |
| vOTU_3033__S8_k141_323664 | Myoviridae | 0 | 0 | NA | 1 | 0 | orphan | 0 | 0 | NA | 0 | 0 | NA |
| vOTU_3044__Co_contig_5281_pilon_pilon | Phycodnaviridae | 0 | 0 | NA | 1 | 0 | orphan | 0 | 0 | NA | 0 | 0 | NA |
| vOTU_3044__S9_contig_290_pilon_pilon | Phycodnaviridae | 0 | 0 | NA | 1 | 0 | orphan | 0 | 0 | NA | 0 | 0 | NA |
| vOTU_3065__Co_contig_272_pilon_pilon | Siphoviridae | 0 | 0 | NA | 2 | 0 | orphan | 0 | 0 | NA | 0 | 0 | NA |
| vOTU_3065__S11_contig_363_pilon_pilo | Siphoviridae | 0 | 0 | NA | 1 | 0 | orphan | 0 | 0 | NA | 0 | 0 | NA |

| | | | | | | | | | | | | | |
|---|---|---|---|---|---|---|---|---|---|---|---|---|---|
| n | | | | | | | | | | | | | |
| vOTU_3065__S4_contig_299_pilon_pilon | Siphoviridae | 0 | 0 | NA | 1 | 0 | orphan | 0 | 0 | NA | 0 | 0 | NA |
| vOTU_3065__S6_k141_635399 | Siphoviridae | 0 | 0 | NA | 1 | 0 | orphan | 0 | 0 | NA | 0 | 0 | NA |
| vOTU_3065__S7_k141_78528 | Siphoviridae | 0 | 0 | NA | 1 | 0 | orphan | 0 | 0 | NA | 0 | 0 | NA |
| vOTU_3065__S8_contig_23_pilon_pilon | Siphoviridae | 0 | 0 | NA | 1 | 0 | orphan | 0 | 0 | NA | 0 | 0 | NA |
| vOTU_3065__S9_contig_387_pilon_pilon | Siphoviridae | 0 | 0 | NA | 1 | 0 | orphan | 0 | 0 | NA | 0 | 0 | NA |
| vOTU_3074__Co_k141_1666744 | Myoviridae | 0 | 0 | NA | 1 | 0 | orphan | 0 | 0 | NA | 0 | 0 | NA |
| vOTU_3074__S6_k141_778652 | Myoviridae | 0 | 0 | NA | 1 | 0 | orphan | 0 | 0 | NA | 0 | 0 | NA |
| vOTU_3140__Co_k141_271333-circular | Unknown_Virus | 0 | 0 | NA | 2 | 0 | orphan | 0 | 0 | NA | 0 | 0 | NA |
| vOTU_317__Co_contig_589_pilon_pilon | Myoviridae | 0 | 1 | NA | 0 | 0 | NA | 0 | 0 | NA | 0 | 0 | NA |
| vOTU_3188__Co_k141_83801 | Myoviridae | 0 | 0 | NA | 1 | 0 | orphan | 0 | 0 | NA | 0 | 0 | NA |
| vOTU_3188__S6_k141_253086 | Myoviridae | 0 | 0 | NA | 1 | 0 | orphan | 0 | 0 | NA | 0 | 0 | NA |
| vOTU_3260__S5_k141_619516 | Podoviridae | 0 | 0 | NA | 1 | 0 | orphan | 0 | 0 | NA | 0 | 0 | NA |
| vOTU_3278__S10_k141_125015 | Podoviridae | 0 | 0 | NA | 1 | 0 | orphan | 0 | 0 | NA | 0 | 0 | NA |
| vOTU_3282__Co_contig_4897_pilon_pilon | Siphoviridae | 0 | 0 | NA | 1 | 0 | orphan | 0 | 0 | NA | 0 | 0 | NA |
| vOTU_3307__S2_k141_16326 | Siphoviridae | 0 | 0 | NA | 1 | 0 | orphan | 0 | 0 | NA | 0 | 0 | NA |
| vOTU_3360__Co_k141_1826557-circular | Unknown_Virus | 0 | 0 | NA | 1 | 0 | orphan | 0 | 0 | NA | 0 | 0 | NA |
| vOTU_3360__S11_k141_761796-circular | Unknown_Virus | 0 | 0 | NA | 1 | 0 | orphan | 0 | 0 | NA | 0 | 0 | NA |
| vOTU_3414__Co_contig_7378_pilon_pilon | Myoviridae | 0 | 1 | NA | 1 | 0 | orphan | 0 | 0 | NA | 0 | 0 | NA |
| vOTU_3414__S11_contig_44_pilon_pilon | Myoviridae | 0 | 1 | NA | 1 | 0 | orphan | 0 | 0 | NA | 0 | 0 | NA |
| vOTU_3414__S11_contig_744_pilon_pilon | Myoviridae | 0 | 1 | NA | 1 | 0 | orphan | 0 | 0 | NA | 0 | 0 | NA |
| vOTU_3416__S6_k141_742799 | Podoviridae | 0 | 0 | NA | 1 | 0 | orphan | 0 | 0 | NA | 0 | 0 | NA |
| vOTU_3442__Co_contig_1855_pilon_pilon | Myoviridae | 0 | 0 | NA | 4 | 0 | orphan | 0 | 0 | NA | 0 | 0 | NA |
| vOTU_3442__S5_contig_250_pilon_pilon | Myoviridae | 0 | 0 | NA | 1 | 0 | orphan | 0 | 0 | NA | 0 | 0 | NA |
| vOTU_3442__S7_contig_607_pilon_pilon | Myoviridae | 0 | 0 | NA | 3 | 0 | orphan | 0 | 0 | NA | 0 | 0 | NA |

| | | | | | | | | | | | | | |
|---|---|---|---|---|---|---|---|---|---|---|---|---|---|
| vOTU_3442__S8_contig_36_pilon_pilon | Myoviridae | 0 | 0 | NA | 2 | 0 | orphan | 0 | 0 | NA | 0 | 0 | NA |
| vOTU_3442__S9_contig_184_pilon_pilon | Myoviridae | 0 | 0 | NA | 1 | 0 | orphan | 0 | 0 | NA | 0 | 0 | NA |
| vOTU_3463__S8_k141_183042 | Phycodnaviridae | 0 | 0 | NA | 1 | 0 | orphan | 0 | 0 | NA | 0 | 0 | NA |
| vOTU_3482__Co_contig_6833_pilon_pilon | Siphoviridae | 0 | 0 | NA | 1 | 0 | orphan | 0 | 0 | NA | 0 | 0 | NA |
| vOTU_3519__S4_k141_249690 | Myoviridae | 0 | 0 | NA | 1 | 0 | orphan | 0 | 0 | NA | 0 | 0 | NA |
| vOTU_3519__S7_k141_478914 | Myoviridae | 0 | 0 | NA | 2 | 0 | orphan | 0 | 0 | NA | 0 | 0 | NA |
| vOTU_3521__S9_k141_183084 | Podoviridae | 0 | 0 | NA | 1 | 0 | orphan | 0 | 0 | NA | 0 | 0 | NA |
| vOTU_3580__S7_contig_422_pilon_pilon | Myoviridae | 0 | 0 | NA | 1 | 0 | orphan | 0 | 0 | NA | 0 | 0 | NA |
| vOTU_3580__S8_contig_271_pilon_pilon | Myoviridae | 0 | 0 | NA | 1 | 0 | orphan | 0 | 0 | NA | 0 | 0 | NA |
| vOTU_360__S10_k141_165370 | Myoviridae | 0 | 1 | NA | 0 | 0 | NA | 0 | 0 | NA | 0 | 0 | NA |
| vOTU_3608__Co_contig_8123_pilon_pilon | Podoviridae | 0 | 0 | NA | 2 | 0 | orphan | 0 | 0 | NA | 0 | 0 | NA |
| vOTU_3643__Co_k141_2554370 | Unclassified_Caudovirales | 0 | 0 | NA | 1 | 0 | orphan | 0 | 0 | NA | 0 | 0 | NA |
| vOTU_3681__S6_k141_242785 | Myoviridae | 0 | 0 | NA | 1 | 0 | orphan | 0 | 0 | NA | 0 | 0 | NA |
| vOTU_3681__S7_k141_46140 | Myoviridae | 0 | 0 | NA | 1 | 0 | orphan | 0 | 0 | NA | 0 | 0 | NA |
| vOTU_3681__S8_k141_146571 | Myoviridae | 0 | 0 | NA | 1 | 0 | orphan | 0 | 0 | NA | 0 | 0 | NA |
| vOTU_3737__Co_contig_6041_pilon_pilon | Siphoviridae | 0 | 0 | NA | 1 | 0 | orphan | 0 | 0 | NA | 0 | 0 | NA |
| vOTU_3775__S9_k141_140367 | Siphoviridae | 0 | 0 | NA | 1 | 0 | orphan | 0 | 0 | NA | 0 | 0 | NA |
| vOTU_378__S3_k141_333273 | Phycodnaviridae | 0 | 0 | NA | 1 | 0 | orphan | 0 | 0 | NA | 0 | 0 | NA |
| vOTU_3780__S10_k141_271602 | Siphoviridae | 0 | 0 | NA | 1 | 0 | orphan | 0 | 0 | NA | 0 | 0 | NA |
| vOTU_3792__S11_k141_373006 | Siphoviridae | 0 | 0 | NA | 1 | 0 | orphan | 0 | 0 | NA | 0 | 0 | NA |
| vOTU_38__S5_k141_367900 | Phycodnaviridae | 0 | 0 | NA | 2 | 0 | orphan | 0 | 0 | NA | 0 | 0 | NA |
| vOTU_3847__S2_k141_237061 | Siphoviridae | 0 | 0 | NA | 1 | 0 | orphan | 0 | 0 | NA | 0 | 0 | NA |
| vOTU_3847__S4_k141_14506 | Siphoviridae | 0 | 0 | NA | 1 | 0 | orphan | 0 | 0 | NA | 0 | 0 | NA |
| vOTU_3847__S8_k141_32443 | Siphoviridae | 0 | 0 | NA | 1 | 0 | orphan | 0 | 0 | NA | 0 | 0 | NA |
| vOTU_3847__S9_k141_14437 | Siphoviridae | 0 | 0 | NA | 1 | 0 | orphan | 0 | 0 | NA | 0 | 0 | NA |
| vOTU_3852__S11_k141_214378 | Podoviridae | 0 | 0 | NA | 1 | 0 | orphan | 0 | 0 | NA | 0 | 0 | NA |

| | | | | | | | | | | | | | |
|---|---|---|---|---|---|---|---|---|---|---|---|---|---|
| vOTU_3861__Co_k141_12717 | Podoviridae | 0 | 0 | NA | 1 | 0 | orphan | 0 | 0 | NA | 0 | 0 | NA |
| vOTU_3861__S9_k141_473176 | Podoviridae | 0 | 0 | NA | 1 | 0 | orphan | 0 | 0 | NA | 0 | 0 | NA |
| vOTU_3866__S3_k141_618434 | Siphoviridae | 0 | 0 | NA | 1 | 0 | orphan | 0 | 0 | NA | 0 | 0 | NA |
| vOTU_3907__S7_k141_391709 | Unknown_Virus | 0 | 0 | NA | 1 | 0 | orphan | 0 | 0 | NA | 0 | 0 | NA |
| vOTU_3936__S10_k141_367452 | Siphoviridae | 0 | 0 | NA | 1 | 0 | orphan | 0 | 0 | NA | 0 | 0 | NA |
| vOTU_3941__Co_k141_2881661 | Myoviridae | 0 | 1 | NA | 0 | 0 | NA | 0 | 0 | NA | 0 | 0 | NA |
| vOTU_3952__S2_k141_727365 | Unclassified_Caudovirales | 0 | 0 | NA | 1 | 0 | orphan | 0 | 0 | NA | 0 | 0 | NA |
| vOTU_4006__Co_k141_380694 | Siphoviridae | 0 | 0 | NA | 1 | 0 | orphan | 0 | 0 | NA | 0 | 0 | NA |
| vOTU_4023__S4_k141_373990 | Podoviridae | 0 | 0 | NA | 1 | 0 | orphan | 0 | 0 | NA | 0 | 0 | NA |
| vOTU_4023__S6_k141_540444 | Podoviridae | 0 | 0 | NA | 1 | 0 | orphan | 0 | 0 | NA | 0 | 0 | NA |
| vOTU_4027__Co_contig_4624_pilon_pilon | Myoviridae | 0 | 1 | NA | 2 | 0 | orphan | 0 | 0 | NA | 0 | 0 | NA |
| vOTU_4027__S10_k141_79946 | Myoviridae | 0 | 0 | NA | 1 | 0 | orphan | 0 | 0 | NA | 0 | 0 | NA |
| vOTU_4027__S8_contig_385_pilon_pilon | Myoviridae | 0 | 0 | NA | 2 | 0 | orphan | 0 | 0 | NA | 0 | 0 | NA |
| vOTU_4058__S11_k141_691567 | Podoviridae | 0 | 1 | NA | 0 | 0 | NA | 0 | 0 | NA | 0 | 0 | NA |
| vOTU_4085__S2_k141_465986-circular | Podoviridae | 0 | 0 | NA | 2 | 0 | orphan | 0 | 0 | NA | 0 | 0 | NA |
| vOTU_4085__S3_k141_397594 | Podoviridae | 0 | 0 | NA | 2 | 0 | orphan | 0 | 0 | NA | 0 | 0 | NA |
| vOTU_4088__Co_k141_2348156-circular | Podoviridae | 0 | 0 | NA | 1 | 0 | orphan | 0 | 0 | NA | 0 | 0 | NA |
| vOTU_4088__S2_k141_745199-circular | Podoviridae | 0 | 0 | NA | 1 | 0 | orphan | 0 | 0 | NA | 0 | 0 | NA |
| vOTU_4090__S2_k141_130170 | Siphoviridae | 0 | 0 | NA | 1 | 0 | orphan | 0 | 0 | NA | 0 | 0 | NA |
| vOTU_4125__Co_k141_1313068 | Siphoviridae | 0 | 0 | NA | 1 | 0 | orphan | 0 | 0 | NA | 0 | 0 | NA |
| vOTU_4129__Co_k141_1357260 | Myoviridae | 0 | 0 | NA | 1 | 0 | orphan | 0 | 0 | NA | 0 | 0 | NA |
| vOTU_4204__S10_k141_87131 | Siphoviridae | 0 | 0 | NA | 1 | 0 | orphan | 0 | 0 | NA | 0 | 0 | NA |
| vOTU_4247__Co_contig_4360_pilon_pilon | Myoviridae | 0 | 1 | NA | 0 | 0 | NA | 0 | 0 | NA | 0 | 0 | NA |
| vOTU_4247__S2_k141_403888 | Myoviridae | 0 | 1 | NA | 0 | 0 | NA | 0 | 0 | NA | 0 | 0 | NA |
| vOTU_4252__Co_k141_1896564 | Unclassified_Caudovirales | 0 | 0 | NA | 1 | 0 | orphan | 0 | 0 | NA | 0 | 0 | NA |
| vOTU_4267__S2_k141_6045 | Myoviridae | 0 | 0 | NA | 1 | 0 | orphan | 0 | 0 | NA | 0 | 0 | NA |

| | | | | | | | | | | | | | |
|---|---|---|---|---|---|---|---|---|---|---|---|---|---|
| vOTU_4267__S4_k141_434896 | Myoviridae | 0 | 0 | NA | 1 | 0 | orphan | 0 | 0 | NA | 0 | 0 | NA |
| vOTU_4267__S5_k141_460020 | Myoviridae | 0 | 0 | NA | 1 | 0 | orphan | 0 | 0 | NA | 0 | 0 | NA |
| vOTU_4271__S4_k141_306690 | Siphoviridae | 0 | 0 | NA | 1 | 0 | orphan | 0 | 0 | NA | 0 | 0 | NA |
| vOTU_4285__Co_k141_417837 | Myoviridae | 0 | 0 | NA | 1 | 0 | orphan | 0 | 0 | NA | 0 | 0 | NA |
| vOTU_4350__S3_k141_86843 | Podoviridae | 0 | 0 | NA | 1 | 0 | orphan | 0 | 0 | NA | 0 | 0 | NA |
| vOTU_4350__S7_k141_280872 | Podoviridae | 0 | 0 | NA | 1 | 0 | orphan | 0 | 0 | NA | 0 | 0 | NA |
| vOTU_4404__Co_contig_5593_pilon_pilon | Siphoviridae | 0 | 0 | NA | 1 | 0 | orphan | 0 | 0 | NA | 0 | 0 | NA |
| vOTU_4423__S3_k141_55240 | Unclassified_Caudovirales | 0 | 0 | NA | 1 | 0 | orphan | 0 | 0 | NA | 0 | 0 | NA |
| vOTU_4423__S4_k141_558661 | Unclassified_Caudovirales | 0 | 0 | NA | 1 | 0 | orphan | 0 | 0 | NA | 0 | 0 | NA |
| vOTU_4423__S7_k141_518970 | Unclassified_Caudovirales | 0 | 0 | NA | 1 | 0 | orphan | 0 | 0 | NA | 0 | 0 | NA |
| vOTU_4452__S2_k141_769993 | Phycodnaviridae | 0 | 0 | NA | 2 | 0 | orphan | 0 | 0 | NA | 0 | 0 | NA |
| vOTU_4452__S3_k141_287470 | Phycodnaviridae | 0 | 0 | NA | 2 | 0 | orphan | 0 | 0 | NA | 0 | 0 | NA |
| vOTU_4497__Co_k141_1815981 | Unknown_Virus | 0 | 0 | NA | 1 | 0 | orphan | 0 | 0 | NA | 0 | 0 | NA |
| vOTU_4513__S5_k141_121934 | Phycodnaviridae | 0 | 0 | NA | 1 | 0 | orphan | 0 | 0 | NA | 0 | 0 | NA |
| vOTU_4563__Co_contig_408_pilon_pilon | Myoviridae | 0 | 0 | NA | 2 | 0 | orphan | 0 | 0 | NA | 0 | 0 | NA |
| vOTU_4594__Co_k141_2201052 | Siphoviridae | 0 | 0 | NA | 1 | 0 | orphan | 0 | 0 | NA | 0 | 0 | NA |
| vOTU_4665__S7_k141_92008 | Myoviridae | 0 | 0 | NA | 1 | 0 | orphan | 0 | 0 | NA | 0 | 0 | NA |
| vOTU_4688__Co_k141_168950 | Unclassified_Caudovirales | 0 | 0 | NA | 1 | 0 | orphan | 0 | 0 | NA | 0 | 0 | NA |
| vOTU_4688__S11_k141_514187 | Unclassified_Caudovirales | 0 | 0 | NA | 1 | 0 | orphan | 0 | 0 | NA | 0 | 0 | NA |
| vOTU_4715__S11_k141_218301 | Myoviridae | 0 | 0 | NA | 1 | 0 | orphan | 0 | 0 | NA | 0 | 0 | NA |
| vOTU_4732__S11_k141_751626 | Siphoviridae | 0 | 0 | NA | 1 | 0 | orphan | 0 | 0 | NA | 0 | 0 | NA |
| vOTU_475__S8_k141_271435 | Phycodnaviridae | 0 | 0 | NA | 1 | 0 | orphan | 0 | 0 | NA | 0 | 0 | NA |
| vOTU_4773__S3_k141_129556-circular | Siphoviridae | 0 | 0 | NA | 1 | 0 | orphan | 0 | 0 | NA | 0 | 0 | NA |
| vOTU_4781__S11_k141_683637 | Unclassified_Caudovirales | 0 | 0 | NA | 1 | 0 | orphan | 0 | 0 | NA | 0 | 0 | NA |
| vOTU_4803__S2_k141_192162 | Podoviridae | 0 | 0 | NA | 1 | 0 | orphan | 0 | 0 | NA | 0 | 0 | NA |
| vOTU_4803__S8_k141_222890 | Podoviridae | 0 | 0 | NA | 1 | 0 | orphan | 0 | 0 | NA | 0 | 0 | NA |
| vOTU_4803__S9_k141_194410 | Podoviridae | 0 | 0 | NA | 1 | 0 | orphan | 0 | 0 | NA | 0 | 0 | NA |

| | | | | | | | | | | | | | |
|---|---|---|---|---|---|---|---|---|---|---|---|---|---|
| vOTU_4821__Co_contig_4761_pilon_pilon | Siphoviridae | 0 | 0 | NA | 1 | 0 | orphan | 0 | 0 | NA | 0 | 0 | NA |
| vOTU_4873__S11_k141_777520 | Siphoviridae | 0 | 0 | NA | 1 | 0 | orphan | 0 | 0 | NA | 0 | 0 | NA |
| vOTU_4899__Co_contig_4998_pilon_pilon | Siphoviridae | 0 | 0 | NA | 1 | 0 | orphan | 0 | 0 | NA | 0 | 0 | NA |
| vOTU_4905__Co_k141_2654467 | Unknown_Virus | 0 | 0 | NA | 1 | 0 | orphan | 0 | 0 | NA | 0 | 0 | NA |
| vOTU_4913__Co_k141_948615 | Myoviridae | 0 | 0 | NA | 2 | 0 | orphan | 0 | 0 | NA | 0 | 0 | NA |
| vOTU_4918__Co_k141_486984 | Unknown_Virus | 0 | 0 | NA | 1 | 0 | orphan | 0 | 0 | NA | 0 | 0 | NA |
| vOTU_492__S4_k141_576793 | Siphoviridae | 0 | 0 | NA | 1 | 0 | orphan | 0 | 0 | NA | 0 | 0 | NA |
| vOTU_4926__Co_k141_630277 | Siphoviridae | 0 | 0 | NA | 1 | 0 | orphan | 0 | 0 | NA | 0 | 0 | NA |
| vOTU_4926__S11_k141_469712 | Siphoviridae | 0 | 0 | NA | 1 | 0 | orphan | 0 | 0 | NA | 0 | 0 | NA |
| vOTU_4949__S6_k141_684615 | Myoviridae | 0 | 0 | NA | 1 | 0 | orphan | 0 | 0 | NA | 0 | 0 | NA |
| vOTU_4949__S9_k141_380183 | Myoviridae | 0 | 0 | NA | 1 | 0 | orphan | 0 | 0 | NA | 0 | 0 | NA |
| vOTU_4965__Co_contig_193_pilon_pilon | Siphoviridae | 0 | 0 | NA | 1 | 0 | orphan | 0 | 0 | NA | 0 | 0 | NA |
| vOTU_4965__Co_contig_8503_pilon_pilon | Siphoviridae | 0 | 0 | NA | 1 | 0 | orphan | 0 | 0 | NA | 0 | 0 | NA |
| vOTU_4965__S10_contig_684_pilon_pilon | Siphoviridae | 0 | 0 | NA | 1 | 0 | orphan | 0 | 0 | NA | 0 | 0 | NA |
| vOTU_4965__S11_contig_799_pilon_pilon | Siphoviridae | 0 | 0 | NA | 1 | 0 | orphan | 0 | 0 | NA | 0 | 0 | NA |
| vOTU_4969__S10_k141_316010 | Podoviridae | 0 | 0 | NA | 1 | 0 | orphan | 0 | 0 | NA | 0 | 0 | NA |
| vOTU_4981__S2_k141_588784 | Siphoviridae | 0 | 0 | NA | 1 | 0 | orphan | 0 | 0 | NA | 0 | 0 | NA |
| vOTU_4996__Co_contig_7898_pilon_pilon | Myoviridae | 0 | 1 | NA | 0 | 0 | NA | 0 | 0 | NA | 0 | 0 | NA |
| vOTU_5006__Co_k141_2019658 | Myoviridae | 0 | 0 | NA | 1 | 0 | orphan | 0 | 0 | NA | 0 | 0 | NA |
| vOTU_504__Co_k141_2690441 | Unclassified_Caudovirales | 0 | 0 | NA | 1 | 0 | orphan | 0 | 0 | NA | 0 | 0 | NA |
| vOTU_5059__S6_k141_157201 | Unclassified_Caudovirales | 0 | 0 | NA | 1 | 0 | orphan | 0 | 0 | NA | 0 | 0 | NA |
| vOTU_5106__Co_k141_132044 | Podoviridae | 0 | 0 | NA | 1 | 0 | orphan | 0 | 0 | NA | 0 | 0 | NA |
| vOTU_5106__S2_k141_44729 | Podoviridae | 0 | 0 | NA | 1 | 0 | orphan | 0 | 0 | NA | 0 | 0 | NA |
| vOTU_5114__S2_k141_71671 | Podoviridae | 0 | 0 | NA | 1 | 0 | orphan | 0 | 0 | NA | 0 | 0 | NA |

| | | | | | | | | | | | | | |
|---|---|---|---|---|---|---|---|---|---|---|---|---|---|
| vOTU_5114__S9_k141_54024 | Podoviridae | 0 | 0 | NA | 1 | 0 | orphan | 0 | 0 | NA | 0 | 0 | NA |
| vOTU_513__S8_k141_76105 | Podoviridae | 0 | 0 | NA | 1 | 0 | orphan | 0 | 0 | NA | 0 | 0 | NA |
| vOTU_5177__Co_k141_1010547 | Myoviridae | 0 | 1 | NA | 0 | 0 | NA | 0 | 0 | NA | 0 | 0 | NA |
| vOTU_5223__S3_k141_226384 | Myoviridae | 0 | 0 | NA | 1 | 0 | orphan | 0 | 0 | NA | 0 | 0 | NA |
| vOTU_5229__S6_k141_589038 | Podoviridae | 0 | 0 | NA | 1 | 0 | orphan | 0 | 0 | NA | 0 | 0 | NA |
| vOTU_658__S8_k141_211165 | Unclassified_Caudovirales | 0 | 0 | NA | 1 | 0 | orphan | 0 | 0 | NA | 0 | 0 | NA |
| vOTU_71__S2_k141_75580 | Podoviridae | 0 | 0 | NA | 1 | 0 | orphan | 0 | 0 | NA | 0 | 0 | NA |
| vOTU_743__S4_k141_295985 | Myoviridae | 0 | 0 | NA | 1 | 0 | orphan | 0 | 0 | NA | 0 | 0 | NA |
| vOTU_743__S5_k141_285040 | Myoviridae | 0 | 0 | NA | 1 | 0 | orphan | 0 | 0 | NA | 0 | 0 | NA |
| vOTU_774__Co_k141_1064006 | Podoviridae | 0 | 0 | NA | 1 | 0 | orphan | 0 | 0 | NA | 0 | 0 | NA |
| vOTU_789__Co_contig_8103_pilon_pilon | Phycodnaviridae | 0 | 0 | NA | 1 | 0 | orphan | 0 | 0 | NA | 0 | 0 | NA |
| vOTU_792__Co_k141_1730617 | Podoviridae | 0 | 0 | NA | 1 | 0 | orphan | 0 | 0 | NA | 0 | 0 | NA |
| vOTU_792__S10_k141_108380-circular | Podoviridae | 0 | 0 | NA | 1 | 0 | orphan | 0 | 0 | NA | 0 | 0 | NA |
| vOTU_861__Co_contig_4361_pilon_pilon | Myoviridae | 0 | 1 | NA | 0 | 0 | NA | 0 | 0 | NA | 0 | 0 | NA |
| vOTU_886__Co_k141_1178114 | Siphoviridae | 0 | 0 | NA | 1 | 0 | orphan | 0 | 0 | NA | 0 | 0 | NA |
| vOTU_902__S6_k141_480930 | Podoviridae | 0 | 0 | NA | 1 | 0 | orphan | 0 | 0 | NA | 0 | 0 | NA |
| vOTU_904__S11_k141_41059 | Siphoviridae | 0 | 0 | NA | 1 | 0 | orphan | 0 | 0 | NA | 0 | 0 | NA |
| vOTU_91__S11_k141_72600 | Unclassified_Caudovirales | 0 | 0 | NA | 1 | 0 | orphan | 0 | 0 | NA | 0 | 0 | NA |
| vOTU_91__S3_k141_520546 | Unclassified_Caudovirales | 0 | 0 | NA | 1 | 0 | orphan | 0 | 0 | NA | 0 | 0 | NA |
| vOTU_925__S8_contig_409_pilon_pilon | Unclassified_Caudovirales | 0 | 0 | NA | 1 | 0 | orphan | 0 | 0 | NA | 0 | 0 | NA |
| vOTU_928__S3_k141_232795 | Phycodnaviridae | 0 | 0 | NA | 2 | 0 | orphan | 0 | 0 | NA | 0 | 0 | NA |
| vOTU_941__S10_k141_459331 | Unclassified_Caudovirales | 0 | 0 | NA | 1 | 0 | orphan | 0 | 0 | NA | 0 | 0 | NA |
| vOTU_948__Co_k141_774231 | Siphoviridae | 0 | 0 | NA | 1 | 0 | orphan | 0 | 0 | NA | 0 | 0 | NA |
| vOTU_948__S10_k141_472697 | Siphoviridae | 0 | 0 | NA | 1 | 0 | orphan | 0 | 0 | NA | 0 | 0 | NA |
| vOTU_965__Co_k141_744553 | Podoviridae | 0 | 0 | NA | 1 | 0 | orphan | 0 | 0 | NA | 0 | 0 | NA |
| vOTU_965__S10_k141_472680 | Podoviridae | 0 | 0 | NA | 1 | 0 | orphan | 0 | 0 | NA | 0 | 0 | NA |
| vOTU_975__Co_contig_1509_pilon_pilon | Myoviridae | 0 | 0 | NA | 1 | 0 | orphan | 0 | 0 | NA | 0 | 0 | NA |

| | | | | | | | | | | | | | |
|---|---|---|---|---|---|---|---|---|---|---|---|---|---|
| vOTU_975__S7_contig_509_pilon_pilon | Myoviridae | 0 | 0 | NA | 1 | 0 | orphan | 0 | 0 | NA | 0 | 0 | NA |
| vOTU_992__S11_k141_282486 | Unclassified_Caudovirales | 0 | 0 | NA | 1 | 0 | orphan | 0 | 0 | NA | 0 | 0 | NA |
| vOTU_GV_127 | Phycodnaviridae | 0 | 0 | NA | 2 | 0 | orphan | 0 | 0 | NA | 0 | 2 | orphan |
| vOTU_GV_14 | Phycodnaviridae | 0 | 0 | NA | 2 | 0 | orphan | 0 | 0 | NA | 0 | 0 | NA |
| vOTU_GV_16 | Phycodnaviridae | 3 | 0 | orphan | 0 | 0 | NA | 0 | 0 | NA | 0 | 0 | NA |
| vOTU_GV_210 | Phycodnaviridae | 0 | 0 | NA | 1 | 0 | orphan | 0 | 0 | NA | 0 | 0 | NA |
| vOTU_GV_214 | Phycodnaviridae | 0 | 0 | NA | 1 | 0 | orphan | 0 | 0 | NA | 0 | 0 | NA |
| vOTU_GV_22 | Phycodnaviridae | 0 | 0 | NA | 7 | 0 | orphan | 0 | 0 | NA | 0 | 0 | NA |
| vOTU_GV_222 | Phycodnaviridae | 3 | 0 | orphan | 1 | 0 | orphan | 0 | 0 | NA | 0 | 0 | NA |
| vOTU_GV_234 | Phycodnaviridae | 0 | 0 | NA | 2 | 0 | orphan | 0 | 0 | NA | 0 | 0 | NA |
| vOTU_GV_246 | Phycodnaviridae | 0 | 0 | NA | 1 | 0 | orphan | 0 | 0 | NA | 0 | 0 | NA |
| vOTU_GV_247 | Phycodnaviridae | 0 | 0 | NA | 1 | 0 | orphan | 0 | 0 | NA | 0 | 0 | NA |
| vOTU_GV_25 | Phycodnaviridae | 0 | 0 | NA | 4 | 0 | orphan | 0 | 0 | NA | 0 | 0 | NA |
| vOTU_GV_270 | Phycodnaviridae | 0 | 0 | NA | 3 | 0 | orphan | 0 | 0 | NA | 0 | 0 | NA |
| vOTU_GV_276 | Phycodnaviridae | 0 | 0 | NA | 3 | 0 | orphan | 0 | 0 | NA | 0 | 0 | NA |
| vOTU_GV_302 | Phycodnaviridae | 0 | 0 | NA | 1 | 0 | orphan | 0 | 0 | NA | 0 | 0 | NA |
| vOTU_GV_305 | Phycodnaviridae | 0 | 0 | NA | 1 | 0 | orphan | 0 | 0 | NA | 0 | 0 | NA |
| vOTU_GV_330 | Phycodnaviridae | 0 | 0 | NA | 3 | 0 | orphan | 0 | 0 | NA | 0 | 0 | NA |
| vOTU_GV_346 | Phycodnaviridae | 0 | 0 | NA | 3 | 0 | orphan | 0 | 0 | NA | 0 | 0 | NA |
| vOTU_GV_360 | Phycodnaviridae | 0 | 0 | NA | 1 | 0 | orphan | 0 | 0 | NA | 0 | 0 | NA |
| vOTU_GV_364 | Phycodnaviridae | 0 | 0 | NA | 1 | 0 | orphan | 0 | 0 | NA | 0 | 0 | NA |
| vOTU_GV_376 | Phycodnaviridae | 0 | 0 | NA | 2 | 0 | orphan | 0 | 0 | NA | 0 | 0 | NA |
| vOTU_GV_38 | Phycodnaviridae | 0 | 0 | NA | 1 | 0 | orphan | 0 | 0 | NA | 0 | 0 | NA |
| vOTU_GV_389 | Phycodnaviridae | 0 | 0 | NA | 2 | 0 | orphan | 0 | 0 | NA | 0 | 0 | NA |
| vOTU_GV_407 | Phycodnaviridae | 0 | 0 | NA | 1 | 0 | orphan | 0 | 0 | NA | 0 | 0 | NA |
| vOTU_GV_410 | Phycodnaviridae | 0 | 0 | NA | 2 | 0 | orphan | 0 | 0 | NA | 0 | 0 | NA |
| vOTU_GV_414 | Phycodnaviridae | 0 | 0 | NA | 2 | 0 | orphan | 0 | 0 | NA | 0 | 0 | NA |

| | | | | | | | | | | | | | |
|---|---|---|---|---|---|---|---|---|---|---|---|---|---|
| vOTU_GV_449 | Phycodnaviridae | 1 | 0 | orphan | 1 | 0 | orphan | 0 | 0 | NA | 0 | 0 | NA |
| vOTU_GV_462 | Phycodnaviridae | 0 | 0 | NA | 1 | 0 | orphan | 0 | 0 | NA | 0 | 0 | NA |
| vOTU_GV_464 | Phycodnaviridae | 0 | 0 | NA | 2 | 0 | orphan | 0 | 0 | NA | 0 | 0 | NA |
| vOTU_GV_468 | Phycodnaviridae | 0 | 0 | NA | 4 | 0 | orphan | 0 | 0 | NA | 0 | 0 | NA |
| vOTU_GV_473 | Phycodnaviridae | 0 | 0 | NA | 3 | 0 | orphan | 0 | 0 | NA | 0 | 0 | NA |
| vOTU_GV_489 | Phycodnaviridae | 0 | 0 | NA | 2 | 0 | orphan | 0 | 0 | NA | 0 | 0 | NA |
| vOTU_GV_71 | Phycodnaviridae | 0 | 0 | NA | 4 | 0 | orphan | 0 | 0 | NA | 0 | 0 | NA |
| vOTU_GV_87 | Phycodnaviridae | 0 | 0 | NA | 1 | 0 | orphan | 0 | 0 | NA | 0 | 0 | NA |
| vOTU_GV_90 | Phycodnaviridae | 0 | 0 | NA | 3 | 0 | orphan | 0 | 0 | NA | 0 | 0 | NA |
| vOTU_GV_91 | Phycodnaviridae | 0 | 0 | NA | 1 | 0 | orphan | 0 | 0 | NA | 0 | 0 | NA |
| vOTU_N_0__S2_k141_378702 | Unknown_Virus | 0 | 0 | NA | 1 | 0 | orphan | 0 | 0 | NA | 0 | 0 | NA |
| vOTU_N_0__S3_k141_352826 | Unknown_Virus | 0 | 0 | NA | 1 | 0 | orphan | 0 | 0 | NA | 0 | 0 | NA |
| vOTU_N_1030__Co_contig_8015_pilon_pilon | Unknown_Virus | 0 | 0 | NA | 1 | 0 | orphan | 0 | 0 | NA | 0 | 0 | NA |
| vOTU_N_1030__S4_contig_410_pilon_pilon | Unknown_Virus | 0 | 0 | NA | 1 | 0 | orphan | 0 | 0 | NA | 0 | 0 | NA |
| vOTU_N_1030__S5_contig_60_pilon_pilon | Unknown_Virus | 0 | 0 | NA | 1 | 0 | orphan | 0 | 0 | NA | 0 | 0 | NA |
| vOTU_N_1030__S7_contig_129_pilon_pilon | Unknown_Virus | 0 | 0 | NA | 1 | 0 | orphan | 0 | 0 | NA | 0 | 0 | NA |
| vOTU_N_1030__S8_contig_204_pilon_pilon | Unknown_Virus | 0 | 0 | NA | 1 | 0 | orphan | 0 | 0 | NA | 0 | 0 | NA |
| vOTU_N_1048__S3_k141_350164 | Unknown_Virus | 0 | 0 | NA | 1 | 0 | orphan | 0 | 0 | NA | 0 | 0 | NA |
| vOTU_N_1048__S5_k141_484074 | Unknown_Virus | 0 | 0 | NA | 1 | 0 | orphan | 0 | 0 | NA | 0 | 0 | NA |
| vOTU_N_1067__S2_k141_361929 | Unknown_Virus | 0 | 0 | NA | 1 | 0 | orphan | 0 | 0 | NA | 0 | 0 | NA |
| vOTU_N_1067__S3_k141_291107 | Unknown_Virus | 0 | 0 | NA | 1 | 0 | orphan | 0 | 0 | NA | 0 | 0 | NA |
| vOTU_N_1097__S5_k141_267678 | Unknown_Virus | 0 | 0 | NA | 1 | 0 | orphan | 0 | 0 | NA | 0 | 0 | NA |
| vOTU_N_1133__S8_k141_433072 | Unknown_Virus | 0 | 0 | NA | 1 | 0 | orphan | 0 | 0 | NA | 0 | 0 | NA |
| vOTU_N_1133__S9_k141_379843 | Unknown_Virus | 0 | 0 | NA | 1 | 0 | orphan | 0 | 0 | NA | 0 | 0 | NA |
| vOTU_N_114__S8_k141_211900 | Unknown_Virus | 0 | 0 | NA | 1 | 0 | orphan | 0 | 0 | NA | 0 | 0 | NA |

| | | | | | | | | | | | | | |
|---|---|---|---|---|---|---|---|---|---|---|---|---|---|
| vOTU_N_1140__Co_contig_4794_pilon_pilon | Unknown_Virus | 0 | 0 | NA | 1 | 0 | orphan | 0 | 0 | NA | 0 | 0 | NA |
| vOTU_N_1169__Co_contig_3127_pilon_pilon | Unknown_Virus | 0 | 0 | NA | 2 | 0 | orphan | 0 | 0 | NA | 0 | 0 | NA |
| vOTU_N_1184__S4_k141_231533 | Unknown_Virus | 0 | 0 | NA | 1 | 0 | orphan | 0 | 0 | NA | 0 | 0 | NA |
| vOTU_N_1187__Co_k141_1664390 | Unknown_Virus | 0 | 0 | NA | 1 | 0 | orphan | 0 | 0 | NA | 0 | 0 | NA |
| vOTU_N_1187__S11_k141_396228 | Unknown_Virus | 0 | 0 | NA | 1 | 0 | orphan | 0 | 0 | NA | 0 | 0 | NA |
| vOTU_N_1242__S3_k141_209936 | Unknown_Virus | 0 | 0 | NA | 1 | 0 | orphan | 0 | 0 | NA | 0 | 0 | NA |
| vOTU_N_1242__S4_k141_173570 | Unknown_Virus | 0 | 0 | NA | 1 | 0 | orphan | 0 | 0 | NA | 0 | 0 | NA |
| vOTU_N_1283__Co_k141_1497779 | Unknown_Virus | 0 | 0 | NA | 1 | 0 | orphan | 0 | 0 | NA | 0 | 0 | NA |
| vOTU_N_129__Co_k141_2491092 | Unknown_Virus | 0 | 0 | NA | 1 | 0 | orphan | 0 | 0 | NA | 0 | 0 | NA |
| vOTU_N_129__S11_k141_551656 | Unknown_Virus | 0 | 0 | NA | 1 | 0 | orphan | 0 | 0 | NA | 0 | 0 | NA |
| vOTU_N_1340__S2_k141_443050 | Unknown_Virus | 0 | 0 | NA | 1 | 0 | orphan | 0 | 0 | NA | 0 | 0 | NA |
| vOTU_N_1340__S3_k141_244341 | Unknown_Virus | 0 | 0 | NA | 1 | 0 | orphan | 0 | 0 | NA | 0 | 0 | NA |
| vOTU_N_1354__S7_k141_444936 | Unknown_Virus | 0 | 0 | NA | 1 | 0 | orphan | 0 | 0 | NA | 0 | 0 | NA |
| vOTU_N_136__Co_contig_7241_pilon_pilon | Unknown_Virus | 0 | 0 | NA | 1 | 0 | orphan | 0 | 0 | NA | 0 | 0 | NA |
| vOTU_N_136__S10_k141_159340 | Unknown_Virus | 0 | 0 | NA | 1 | 0 | orphan | 0 | 0 | NA | 0 | 0 | NA |
| vOTU_N_136__S8_k141_348190 | Unknown_Virus | 0 | 0 | NA | 1 | 0 | orphan | 0 | 0 | NA | 0 | 0 | NA |
| vOTU_N_1383__S5_k141_466865 | Unknown_Virus | 0 | 0 | NA | 1 | 0 | orphan | 0 | 0 | NA | 0 | 0 | NA |
| vOTU_N_1383__S7_k141_511313 | Unknown_Virus | 0 | 0 | NA | 1 | 0 | orphan | 0 | 0 | NA | 0 | 0 | NA |
| vOTU_N_1414__S11_k141_132877 | Unknown_Virus | 0 | 0 | NA | 1 | 0 | orphan | 0 | 0 | NA | 0 | 0 | NA |
| vOTU_N_1444__Co_contig_134_pilon_pilon | Unknown_Virus | 0 | 0 | NA | 1 | 0 | orphan | 0 | 0 | NA | 0 | 0 | NA |
| vOTU_N_1444__S7_k141_2642 | Unknown_Virus | 0 | 0 | NA | 1 | 0 | orphan | 0 | 0 | NA | 0 | 0 | NA |
| vOTU_N_1444__S9_contig_162_pilon_pilon | Unknown_Virus | 0 | 0 | NA | 1 | 0 | orphan | 0 | 0 | NA | 0 | 0 | NA |
| vOTU_N_1457__Co_k141_141373 | Unknown_Virus | 0 | 0 | NA | 1 | 0 | orphan | 0 | 0 | NA | 0 | 0 | NA |
| vOTU_N_1493__S7_k141_716468 | Unknown_Virus | 0 | 0 | NA | 1 | 0 | orphan | 0 | 0 | NA | 0 | 0 | NA |
| vOTU_N_1508__Co_k141_71469 | Unknown_Virus | 0 | 1 | NA | 1 | 0 | orphan | 0 | 0 | NA | 0 | 0 | NA |

| | | | | | | | | | | | | | |
|---|---|---|---|---|---|---|---|---|---|---|---|---|---|
| vOTU_N_1508__S3_k141_565821 | Unknown_Virus | 0 | 1 | NA | 1 | 0 | orphan | 0 | 0 | NA | 0 | 0 | NA |
| vOTU_N_151__S11_k141_498879 | Unknown_Virus | 0 | 0 | NA | 1 | 0 | orphan | 0 | 0 | NA | 0 | 0 | NA |
| vOTU_N_1532__S4_k141_595601 | Unknown_Virus | 0 | 0 | NA | 1 | 0 | orphan | 0 | 0 | NA | 0 | 0 | NA |
| vOTU_N_1532__S6_k141_80229 | Unknown_Virus | 0 | 0 | NA | 1 | 0 | orphan | 0 | 0 | NA | 0 | 0 | NA |
| vOTU_N_1536__S11_k141_25373 | Unknown_Virus | 0 | 0 | NA | 1 | 0 | orphan | 0 | 0 | NA | 0 | 0 | NA |
| vOTU_N_1580__S4_k141_478784 | Unknown_Virus | 0 | 0 | NA | 1 | 0 | orphan | 0 | 0 | NA | 0 | 0 | NA |
| vOTU_N_1580__S6_k141_41334 | Unknown_Virus | 0 | 0 | NA | 1 | 0 | orphan | 0 | 0 | NA | 0 | 0 | NA |
| vOTU_N_159__S4_k141_195893 | Unknown_Virus | 0 | 1 | NA | 0 | 0 | NA | 0 | 0 | NA | 0 | 0 | NA |
| vOTU_N_159__S7_k141_163137 | Unknown_Virus | 0 | 1 | NA | 0 | 0 | NA | 0 | 0 | NA | 0 | 0 | NA |
| vOTU_N_1590__S3_k141_152657 | Unknown_Virus | 0 | 0 | NA | 1 | 0 | orphan | 0 | 0 | NA | 0 | 0 | NA |
| vOTU_N_1590__S5_k141_428724 | Unknown_Virus | 0 | 0 | NA | 1 | 0 | orphan | 0 | 0 | NA | 0 | 0 | NA |
| vOTU_N_1604__S7_k141_274273 | Unknown_Virus | 0 | 0 | NA | 1 | 0 | orphan | 0 | 0 | NA | 0 | 0 | NA |
| vOTU_N_1669__S4_contig_140_pilon_pilon | Unknown_Virus | 0 | 0 | NA | 1 | 0 | orphan | 0 | 0 | NA | 0 | 0 | NA |
| vOTU_N_1669__S5_contig_154_pilon_pilon | Unknown_Virus | 0 | 0 | NA | 1 | 0 | orphan | 0 | 0 | NA | 0 | 0 | NA |
| vOTU_N_1701__Co_contig_6510_pilon_pilon | Unknown_Virus | 0 | 0 | NA | 1 | 0 | orphan | 0 | 0 | NA | 0 | 0 | NA |
| vOTU_N_1701__S5_contig_349_pilon_pilon | Unknown_Virus | 0 | 0 | NA | 1 | 0 | orphan | 0 | 0 | NA | 0 | 0 | NA |
| vOTU_N_1722__Co_k141_1397407 | Unknown_Virus | 0 | 0 | NA | 1 | 0 | orphan | 0 | 0 | NA | 0 | 0 | NA |
| vOTU_N_1726__Co_contig_7040_pilon_pilon | Unknown_Virus | 0 | 0 | NA | 2 | 0 | orphan | 0 | 0 | NA | 0 | 0 | NA |
| vOTU_N_1726__Co_k141_289199 | Unknown_Virus | 0 | 0 | NA | 2 | 0 | orphan | 0 | 0 | NA | 0 | 0 | NA |
| vOTU_N_1726__S6_k141_448157 | Unknown_Virus | 0 | 0 | NA | 2 | 0 | orphan | 0 | 0 | NA | 0 | 0 | NA |
| vOTU_N_1726__S7_k141_640913 | Unknown_Virus | 0 | 0 | NA | 2 | 0 | orphan | 0 | 0 | NA | 0 | 0 | NA |
| vOTU_N_1726__S8_contig_6_pilon_pilon | Unknown_Virus | 0 | 0 | NA | 2 | 0 | orphan | 0 | 0 | NA | 0 | 0 | NA |
| vOTU_N_1726__S9_k141_35340 | Unknown_Virus | 0 | 0 | NA | 1 | 0 | orphan | 0 | 0 | NA | 0 | 0 | NA |
| vOTU_N_1749__S5_k141_668676 | Unknown_Virus | 0 | 0 | NA | 1 | 0 | orphan | 0 | 0 | NA | 0 | 0 | NA |

| | | | | | | | | | | | | | |
|---|---|---|---|---|---|---|---|---|---|---|---|---|---|
| vOTU_N_1754__Co_contig_412_pilon_pilon | Unknown_Virus | 0 | 0 | NA | 1 | 0 | orphan | 0 | 0 | NA | 0 | 0 | NA |
| vOTU_N_1754__S3_k141_103668 | Unknown_Virus | 0 | 0 | NA | 1 | 0 | orphan | 0 | 0 | NA | 0 | 0 | NA |
| vOTU_N_1754__S5_k141_584542 | Unknown_Virus | 0 | 0 | NA | 1 | 0 | orphan | 0 | 0 | NA | 0 | 0 | NA |
| vOTU_N_1780__S4_k141_691396 | Unknown_Virus | 0 | 0 | NA | 1 | 0 | orphan | 0 | 0 | NA | 0 | 0 | NA |
| vOTU_N_1787__S2_k141_153435 | Unknown_Virus | 0 | 0 | NA | 1 | 0 | orphan | 0 | 0 | NA | 0 | 0 | NA |
| vOTU_N_1787__S4_k141_357148 | Unknown_Virus | 0 | 0 | NA | 1 | 0 | orphan | 0 | 0 | NA | 0 | 0 | NA |
| vOTU_N_2002__S2_k141_561506 | Unknown_Virus | 0 | 0 | NA | 1 | 0 | orphan | 0 | 0 | NA | 0 | 0 | NA |
| vOTU_N_2030__Co_contig_6284_pilon_pilon | Unknown_Virus | 0 | 0 | NA | 1 | 0 | orphan | 0 | 0 | NA | 0 | 0 | NA |
| vOTU_N_2030__Co_k141_485061 | Unknown_Virus | 0 | 0 | NA | 1 | 0 | orphan | 0 | 0 | NA | 0 | 0 | NA |
| vOTU_N_2030__S4_k141_253484 | Unknown_Virus | 0 | 0 | NA | 1 | 0 | orphan | 0 | 0 | NA | 0 | 0 | NA |
| vOTU_N_2030__S7_k141_139328 | Unknown_Virus | 0 | 0 | NA | 1 | 0 | orphan | 0 | 0 | NA | 0 | 0 | NA |
| vOTU_N_2030__S8_k141_356023 | Unknown_Virus | 0 | 0 | NA | 1 | 0 | orphan | 0 | 0 | NA | 0 | 0 | NA |
| vOTU_N_2049__S4_k141_726211 | Unknown_Virus | 0 | 0 | NA | 1 | 0 | orphan | 0 | 0 | NA | 0 | 0 | NA |
| vOTU_N_2063__Co_k141_2397596 | Unknown_Virus | 0 | 0 | NA | 1 | 0 | orphan | 0 | 0 | NA | 0 | 0 | NA |
| vOTU_N_2065__S11_contig_790_pilon_pilon | Unknown_Virus | 0 | 0 | NA | 1 | 0 | orphan | 0 | 0 | NA | 0 | 0 | NA |
| vOTU_N_2073__Co_k141_1245851 | Unknown_Virus | 0 | 0 | NA | 1 | 0 | orphan | 0 | 0 | NA | 0 | 0 | NA |
| vOTU_N_2103__Co_k141_585579 | Unknown_Virus | 0 | 0 | NA | 1 | 0 | orphan | 0 | 0 | NA | 0 | 0 | NA |
| vOTU_N_2103__S3_k141_385580 | Unknown_Virus | 0 | 0 | NA | 1 | 0 | orphan | 0 | 0 | NA | 0 | 0 | NA |
| vOTU_N_214__Co_k141_2701152 | Unknown_Virus | 0 | 0 | NA | 1 | 0 | orphan | 0 | 0 | NA | 0 | 0 | NA |
| vOTU_N_2147__S6_k141_403099 | Unknown_Virus | 0 | 0 | NA | 1 | 0 | orphan | 0 | 0 | NA | 0 | 0 | NA |
| vOTU_N_2147__S8_k141_508504 | Unknown_Virus | 0 | 0 | NA | 1 | 0 | orphan | 0 | 0 | NA | 0 | 0 | NA |
| vOTU_N_2147__S9_k141_269636 | Unknown_Virus | 0 | 0 | NA | 1 | 0 | orphan | 0 | 0 | NA | 0 | 0 | NA |
| vOTU_N_248__S2_k141_225817 | Unknown_Virus | 0 | 0 | NA | 1 | 0 | orphan | 0 | 0 | NA | 0 | 0 | NA |
| vOTU_N_269__Co_k141_267590 | Unknown_Virus | 0 | 0 | NA | 1 | 0 | orphan | 0 | 0 | NA | 0 | 0 | NA |
| vOTU_N_286__S4_k141_97536 | Unknown_Virus | 0 | 0 | NA | 1 | 0 | orphan | 0 | 0 | NA | 0 | 0 | NA |
| vOTU_N_371__Co_k141_2007472- | Unknown_Virus | 0 | 0 | NA | 1 | 0 | orphan | 0 | 0 | NA | 0 | 0 | NA |

| | | | | | | | | | | | | | |
|---|---|---|---|---|---|---|---|---|---|---|---|---|---|
| circular | | | | | | | | | | | | | |
| vOTU_N_371__S2_k141_436713-circular | Unknown_Virus | 0 | 0 | NA | 1 | 0 | orphan | 0 | 0 | NA | 0 | 0 | NA |
| vOTU_N_371__S4_k141_736992 | Unknown_Virus | 0 | 0 | NA | 1 | 0 | orphan | 0 | 0 | NA | 0 | 0 | NA |
| vOTU_N_371__S7_k141_539961-circular | Unknown_Virus | 0 | 0 | NA | 1 | 0 | orphan | 0 | 0 | NA | 0 | 0 | NA |
| vOTU_N_380__S2_k141_199966 | Unknown_Virus | 0 | 0 | NA | 1 | 0 | orphan | 0 | 0 | NA | 0 | 0 | NA |
| vOTU_N_380__S3_k141_110231 | Unknown_Virus | 0 | 0 | NA | 1 | 0 | orphan | 0 | 0 | NA | 0 | 0 | NA |
| vOTU_N_382__S9_k141_63941-circular | Unknown_Virus | 0 | 0 | NA | 1 | 0 | orphan | 0 | 0 | NA | 0 | 0 | NA |
| vOTU_N_428__S7_k141_564877 | Unknown_Virus | 0 | 0 | NA | 1 | 0 | orphan | 0 | 0 | NA | 0 | 0 | NA |
| vOTU_N_440__Co_k141_999945 | Unknown_Virus | 0 | 0 | NA | 1 | 0 | orphan | 0 | 0 | NA | 0 | 0 | NA |
| vOTU_N_440__S10_k141_362159 | Unknown_Virus | 0 | 0 | NA | 1 | 0 | orphan | 0 | 0 | NA | 0 | 0 | NA |
| vOTU_N_444__Co_contig_3450_pilon_pilon | Unknown_Virus | 0 | 0 | NA | 2 | 0 | orphan | 0 | 0 | NA | 0 | 0 | NA |
| vOTU_N_444__S5_contig_413_pilon_pilon | Unknown_Virus | 0 | 0 | NA | 2 | 0 | orphan | 0 | 0 | NA | 0 | 0 | NA |
| vOTU_N_459__S4_k141_261583 | Unknown_Virus | 0 | 0 | NA | 1 | 0 | orphan | 0 | 0 | NA | 0 | 0 | NA |
| vOTU_N_462__Co_k141_757791 | Unknown_Virus | 0 | 0 | NA | 1 | 0 | orphan | 0 | 0 | NA | 0 | 0 | NA |
| vOTU_N_462__S8_k141_108177-circular | Unknown_Virus | 0 | 0 | NA | 1 | 0 | orphan | 0 | 0 | NA | 0 | 0 | NA |
| vOTU_N_462__S9_k141_102166 | Unknown_Virus | 0 | 0 | NA | 1 | 0 | orphan | 0 | 0 | NA | 0 | 0 | NA |
| vOTU_N_475__Co_contig_3479_pilon_pilon | Unknown_Virus | 0 | 0 | NA | 1 | 0 | orphan | 0 | 0 | NA | 0 | 0 | NA |
| vOTU_N_475__S3_k141_209930 | Unknown_Virus | 0 | 0 | NA | 1 | 0 | orphan | 0 | 0 | NA | 0 | 0 | NA |
| vOTU_N_475__S4_k141_133015 | Unknown_Virus | 0 | 0 | NA | 1 | 0 | orphan | 0 | 0 | NA | 0 | 0 | NA |
| vOTU_N_475__S5_k141_489400 | Unknown_Virus | 0 | 0 | NA | 1 | 0 | orphan | 0 | 0 | NA | 0 | 0 | NA |
| vOTU_N_489__S7_k141_174182 | Unknown_Virus | 0 | 0 | NA | 1 | 0 | orphan | 0 | 0 | NA | 0 | 0 | NA |
| vOTU_N_5__Co_k141_78126 | Unknown_Virus | 0 | 1 | NA | 0 | 0 | NA | 0 | 0 | NA | 0 | 0 | NA |
| vOTU_N_512__Co_k141_314978 | Unknown_Virus | 0 | 0 | NA | 1 | 0 | orphan | 0 | 0 | NA | 0 | 0 | NA |
| vOTU_N_512__S4_k141_364609 | Unknown_Virus | 0 | 0 | NA | 1 | 0 | orphan | 0 | 0 | NA | 0 | 0 | NA |
| vOTU_N_530__S9_k141_517060 | Unknown_Virus | 0 | 0 | NA | 1 | 0 | orphan | 0 | 0 | NA | 0 | 0 | NA |
| vOTU_N_540__Co_contig_6035_pilon_pi | Unknown_Virus | 0 | 0 | NA | 1 | 0 | orphan | 0 | 0 | NA | 0 | 0 | NA |

| | | | | | | | | | | | | | |
|---|---|---|---|---|---|---|---|---|---|---|---|---|---|
| lon | | | | | | | | | | | | | |
| vOTU_N_540__S8_contig_551_pilon_pilon | Unknown_Virus | 0 | 0 | NA | 1 | 0 | orphan | 0 | 0 | NA | 0 | 0 | NA |
| vOTU_N_592__Co_k141_1374739 | Unknown_Virus | 0 | 0 | NA | 1 | 0 | orphan | 0 | 0 | NA | 0 | 0 | NA |
| vOTU_N_592__S11_k141_387930 | Unknown_Virus | 0 | 0 | NA | 1 | 0 | orphan | 0 | 0 | NA | 0 | 0 | NA |
| vOTU_N_599__S11_k141_262500 | Unknown_Virus | 0 | 0 | NA | 1 | 0 | orphan | 0 | 0 | NA | 0 | 0 | NA |
| vOTU_N_6__Co_k141_811564 | Unknown_Virus | 0 | 0 | NA | 1 | 0 | orphan | 0 | 0 | NA | 0 | 0 | NA |
| vOTU_N_6__S10_k141_151574 | Unknown_Virus | 0 | 0 | NA | 1 | 0 | orphan | 0 | 0 | NA | 0 | 0 | NA |
| vOTU_N_609__Co_k141_2636756 | Unknown_Virus | 0 | 0 | NA | 1 | 0 | orphan | 0 | 0 | NA | 0 | 0 | NA |
| vOTU_N_648__Co_contig_4186_pilon_pilon | Unknown_Virus | 0 | 0 | NA | 1 | 0 | orphan | 0 | 0 | NA | 0 | 0 | NA |
| vOTU_N_648__S5_k141_429925 | Unknown_Virus | 0 | 0 | NA | 1 | 0 | orphan | 0 | 0 | NA | 0 | 0 | NA |
| vOTU_N_665__S2_contig_186_pilon_pilon | Unknown_Virus | 0 | 0 | NA | 1 | 0 | orphan | 0 | 0 | NA | 0 | 0 | NA |
| vOTU_N_687__Co_k141_1329792 | Unknown_Virus | 0 | 0 | NA | 1 | 0 | orphan | 0 | 0 | NA | 0 | 0 | NA |
| vOTU_N_690__Co_k141_2973538-circular | Unknown_Virus | 0 | 0 | NA | 2 | 0 | orphan | 0 | 0 | NA | 0 | 0 | NA |
| vOTU_N_690__S5_k141_632482 | Unknown_Virus | 0 | 0 | NA | 1 | 0 | orphan | 0 | 0 | NA | 0 | 0 | NA |
| vOTU_N_752__Co_k141_1009193 | Unknown_Virus | 0 | 0 | NA | 1 | 0 | orphan | 0 | 0 | NA | 0 | 0 | NA |
| vOTU_N_77__S2_k141_118260 | Unknown_Virus | 0 | 0 | NA | 0 | 0 | NA | 0 | 1 | NA | 0 | 0 | NA |
| vOTU_N_77__S3_k141_123426 | Unknown_Virus | 0 | 0 | NA | 0 | 0 | NA | 0 | 1 | NA | 0 | 0 | NA |
| vOTU_N_77__S4_k141_186037 | Unknown_Virus | 0 | 0 | NA | 0 | 0 | NA | 0 | 1 | NA | 0 | 0 | NA |
| vOTU_N_77__S5_k141_370673 | Unknown_Virus | 0 | 0 | NA | 0 | 0 | NA | 0 | 1 | NA | 0 | 0 | NA |
| vOTU_N_799__S7_contig_396_pilon_pilon | Unknown_Virus | 0 | 0 | NA | 1 | 0 | orphan | 0 | 0 | NA | 0 | 0 | NA |
| vOTU_N_816__Co_contig_5825_pilon_pilon | Unknown_Virus | 0 | 0 | NA | 1 | 0 | orphan | 0 | 0 | NA | 0 | 0 | NA |
| vOTU_N_822__S2_k141_653076 | Unknown_Virus | 0 | 0 | NA | 1 | 0 | orphan | 0 | 0 | NA | 0 | 0 | NA |
| vOTU_N_822__S6_k141_80578 | Unknown_Virus | 0 | 0 | NA | 1 | 0 | orphan | 0 | 0 | NA | 0 | 0 | NA |
| vOTU_N_832__Co_k141_1808514-circular | Unknown_Virus | 0 | 0 | NA | 1 | 0 | orphan | 0 | 0 | NA | 0 | 0 | NA |

| | | | | | | | | | | | | | |
|---|---|---|---|---|---|---|---|---|---|---|---|---|---|
| vOTU_N_832__S11_k141_308155-circular | Unknown_Virus | 0 | 0 | NA | 1 | 0 | orphan | 0 | 0 | NA | 0 | 0 | NA |
| vOTU_N_833__S6_k141_722603 | Unknown_Virus | 0 | 1 | NA | 0 | 0 | NA | 0 | 0 | NA | 0 | 0 | NA |
| vOTU_N_833__S8_k141_436188 | Unknown_Virus | 0 | 1 | NA | 0 | 0 | NA | 0 | 0 | NA | 0 | 0 | NA |
| vOTU_N_874__Co_contig_4095_pilon_pilon | Unknown_Virus | 0 | 0 | NA | 1 | 0 | orphan | 0 | 0 | NA | 0 | 0 | NA |
| vOTU_N_874__S4_k141_155941 | Unknown_Virus | 0 | 0 | NA | 1 | 0 | orphan | 0 | 0 | NA | 0 | 0 | NA |
| vOTU_N_892__Co_k141_380732 | Unknown_Virus | 0 | 0 | NA | 1 | 0 | orphan | 0 | 0 | NA | 0 | 0 | NA |
| vOTU_N_895__Co_contig_6186_pilon_pilon | Unknown_Virus | 0 | 0 | NA | 1 | 0 | orphan | 0 | 0 | NA | 0 | 0 | NA |
| vOTU_N_923__Co_k141_861972 | Unknown_Virus | 0 | 1 | NA | 0 | 0 | NA | 0 | 0 | NA | 0 | 0 | NA |
| vOTU_N_926__S8_k141_185931 | Unknown_Virus | 0 | 0 | NA | 2 | 0 | orphan | 0 | 0 | NA | 0 | 0 | NA |
| vOTU_N_973__Co_k141_830347-circular | Unknown_Virus | 0 | 0 | NA | 1 | 0 | orphan | 0 | 0 | NA | 0 | 0 | NA |
| vOTU_N_973__S3_k141_609624-circular | Unknown_Virus | 0 | 0 | NA | 1 | 0 | orphan | 0 | 0 | NA | 0 | 0 | NA |
| vOTU_N_987__Co_k141_158605 | Unknown_Virus | 0 | 0 | NA | 1 | 0 | orphan | 0 | 0 | NA | 0 | 0 | NA |
| vOTU_N_99__S3_k141_305617 | Unknown_Virus | 0 | 0 | NA | 1 | 0 | orphan | 0 | 0 | NA | 0 | 0 | NA |
| vOTU_sg_10193__S8_k141_392980 | Unknown_Virus | 0 | 0 | NA | 1 | 0 | orphan | 0 | 0 | NA | 0 | 0 | NA |
| vOTU_sg_10215__Co_k141_1884905 | Unclassified_Caudovirales | 0 | 0 | NA | 1 | 0 | orphan | 0 | 0 | NA | 0 | 0 | NA |
| vOTU_sg_10258__S11_k141_100897 | Podoviridae | 0 | 0 | NA | 1 | 0 | orphan | 0 | 0 | NA | 0 | 0 | NA |
| vOTU_sg_10518__Co_k141_1907519 | Phycodnaviridae | 0 | 0 | NA | 1 | 0 | orphan | 0 | 0 | NA | 0 | 0 | NA |
| vOTU_sg_10538__S10_k141_75557 | Siphoviridae | 0 | 0 | NA | 1 | 0 | orphan | 0 | 0 | NA | 0 | 0 | NA |
| vOTU_sg_10592__Co_k141_147594 | Siphoviridae | 0 | 0 | NA | 1 | 0 | orphan | 0 | 0 | NA | 0 | 1 | orphan |
| vOTU_sg_10713__S9_contig_359_pilon_pilon | Siphoviridae | 0 | 0 | NA | 2 | 0 | orphan | 0 | 0 | NA | 0 | 0 | NA |
| vOTU_sg_10724__Co_contig_6120_pilon_pilon | Siphoviridae | 0 | 0 | NA | 1 | 0 | orphan | 0 | 0 | NA | 0 | 0 | NA |
| vOTU_sg_1087__S10_contig_500_pilon_pilon | Siphoviridae | 0 | 0 | NA | 1 | 0 | orphan | 0 | 0 | NA | 0 | 0 | NA |
| vOTU_sg_11026__S3_k141_458088-circular | Myoviridae | 0 | 1 | NA | 0 | 0 | NA | 0 | 0 | NA | 0 | 0 | NA |

| | | | | | | | | | | | | | |
|---|---|---|---|---|---|---|---|---|---|---|---|---|---|
| vOTU_sg_11122__S8_contig_438_pilon_pilon | Siphoviridae | 0 | 0 | NA | 1 | 0 | orphan | 0 | 0 | NA | 0 | 0 | NA |
| vOTU_sg_11165__S5_contig_92_pilon_pilon | Siphoviridae | 0 | 0 | NA | 1 | 0 | orphan | 0 | 0 | NA | 0 | 0 | NA |
| vOTU_sg_11316__Co_k141_3054345 | Unknown_Virus | 0 | 0 | NA | 1 | 0 | orphan | 0 | 0 | NA | 0 | 0 | NA |
| vOTU_sg_11422__Co_k141_1165020 | Unclassified_Caudovirales | 0 | 0 | NA | 1 | 0 | orphan | 0 | 0 | NA | 0 | 0 | NA |
| vOTU_sg_11559__Co_k141_533399 | Unknown_Virus | 0 | 0 | NA | 0 | 0 | NA | 0 | 0 | NA | 0 | 1 | orphan |
| vOTU_sg_11656__S11_k141_627832 | Siphoviridae | 0 | 0 | NA | 1 | 0 | orphan | 0 | 0 | NA | 0 | 0 | NA |
| vOTU_sg_11778__S9_contig_334_pilon_pilon | Unknown_Virus | 0 | 0 | NA | 2 | 0 | orphan | 0 | 0 | NA | 0 | 0 | NA |
| vOTU_sg_11886__Co_k141_1410875 | Unclassified_no_order_Iridoviridae | 0 | 0 | NA | 1 | 0 | orphan | 0 | 0 | NA | 0 | 0 | NA |
| vOTU_sg_11940__Co_k141_32073 | Unknown_Virus | 0 | 0 | NA | 1 | 0 | orphan | 0 | 0 | NA | 0 | 0 | NA |
| vOTU_sg_12087__S11_k141_573128 | Unclassified_Caudovirales | 0 | 0 | NA | 1 | 0 | orphan | 0 | 0 | NA | 0 | 0 | NA |
| vOTU_sg_12100__Co_k141_1666574-circular | Unclassified_Caudovirales | 0 | 0 | NA | 1 | 0 | orphan | 0 | 0 | NA | 0 | 0 | NA |
| vOTU_sg_1613__Co_k141_434189 | Myoviridae | 0 | 0 | NA | 1 | 0 | orphan | 0 | 0 | NA | 0 | 0 | NA |
| vOTU_sg_163__Co_contig_4181_pilon_pilon | Unclassified_Caudovirales | 0 | 0 | NA | 1 | 0 | orphan | 0 | 0 | NA | 0 | 0 | NA |
| vOTU_sg_1701__S7_k141_277492 | Unclassified_Caudovirales | 0 | 0 | NA | 1 | 0 | orphan | 0 | 0 | NA | 0 | 0 | NA |
| vOTU_sg_1929__S7_k141_215008 | Unknown_Virus | 0 | 0 | NA | 1 | 0 | orphan | 0 | 0 | NA | 0 | 0 | NA |
| vOTU_sg_2181__Co_k141_1782479-circular | Podoviridae | 0 | 0 | NA | 1 | 0 | orphan | 0 | 0 | NA | 0 | 0 | NA |
| vOTU_sg_2402__Co_contig_4813_pilon_pilon | Unclassified_Caudovirales | 0 | 0 | NA | 1 | 0 | orphan | 0 | 0 | NA | 0 | 0 | NA |
| vOTU_sg_2403__Co_k141_948632 | Myoviridae | 0 | 0 | NA | 1 | 0 | orphan | 0 | 0 | NA | 0 | 0 | NA |
| vOTU_sg_251__S5_k141_374473 | Podoviridae | 0 | 0 | NA | 1 | 0 | orphan | 0 | 0 | NA | 0 | 0 | NA |
| vOTU_sg_2556__Co_contig_7920_pilon_pilon | Myoviridae | 0 | 0 | NA | 1 | 0 | orphan | 0 | 0 | NA | 0 | 0 | NA |
| vOTU_sg_2621__Co_contig_2832_pilon_pilon | Myoviridae | 0 | 0 | NA | 1 | 0 | orphan | 0 | 0 | NA | 0 | 0 | NA |
| vOTU_sg_2743__S10_k141_610836 | Siphoviridae | 0 | 0 | NA | 1 | 0 | orphan | 0 | 0 | NA | 0 | 0 | NA |

| | | | | | | | | | | | | | |
|---|---|---|---|---|---|---|---|---|---|---|---|---|---|
| vOTU_sg_2773__S4_contig_51_pilon_pilon | Siphoviridae | 0 | 0 | NA | 1 | 0 | orphan | 0 | 0 | NA | 0 | 0 | NA |
| vOTU_sg_2786__Co_contig_5853_pilon_pilon | Myoviridae | 0 | 1 | NA | 0 | 0 | NA | 0 | 0 | NA | 0 | 0 | NA |
| vOTU_sg_2804__Co_contig_6086_pilon_pilon | Siphoviridae | 0 | 0 | NA | 1 | 0 | orphan | 0 | 0 | NA | 0 | 0 | NA |
| vOTU_sg_2848__S11_k141_141931 | Unknown_Virus | 0 | 0 | NA | 1 | 0 | orphan | 0 | 0 | NA | 0 | 0 | NA |
| vOTU_sg_2864__Co_k141_97649 | Unknown_Virus | 0 | 0 | NA | 1 | 0 | orphan | 0 | 0 | NA | 0 | 0 | NA |
| vOTU_sg_2868__S5_contig_200_pilon_pilon | Siphoviridae | 0 | 0 | NA | 1 | 0 | orphan | 0 | 0 | NA | 0 | 0 | NA |
| vOTU_sg_2885__Co_contig_8090_pilon_pilon | Myoviridae | 0 | 0 | NA | 1 | 0 | orphan | 0 | 0 | NA | 0 | 0 | NA |
| vOTU_sg_2899__S2_contig_403_pilon_pilon | Phycodnaviridae | 0 | 0 | NA | 1 | 0 | orphan | 0 | 0 | NA | 0 | 0 | NA |
| vOTU_sg_2911__S8_contig_375_pilon_pilon | Phycodnaviridae | 0 | 0 | NA | 1 | 0 | orphan | 0 | 0 | NA | 0 | 0 | NA |
| vOTU_sg_2916__S8_k141_221192 | Siphoviridae | 0 | 0 | NA | 1 | 0 | orphan | 0 | 0 | NA | 0 | 0 | NA |
| vOTU_sg_2948__Co_contig_969_pilon_pilon | Siphoviridae | 0 | 0 | NA | 1 | 0 | orphan | 0 | 0 | NA | 0 | 0 | NA |
| vOTU_sg_2950__Co_k141_1950424 | Unclassified_Caudovirales | 0 | 0 | NA | 2 | 0 | orphan | 0 | 0 | NA | 0 | 0 | NA |
| vOTU_sg_2951__S11_k141_409548 | Siphoviridae | 0 | 0 | NA | 1 | 0 | orphan | 0 | 0 | NA | 0 | 0 | NA |
| vOTU_sg_2975__Co_contig_5357_pilon_pilon | Unknown_Virus | 0 | 0 | NA | 1 | 0 | orphan | 0 | 0 | NA | 0 | 0 | NA |
| vOTU_sg_2976__S11_k141_187229 | Podoviridae | 0 | 0 | NA | 1 | 0 | orphan | 0 | 0 | NA | 0 | 0 | NA |
| vOTU_sg_2983__S6_k141_251364 | Unclassified_Caudovirales | 0 | 0 | NA | 1 | 0 | orphan | 0 | 0 | NA | 0 | 0 | NA |
| vOTU_sg_2984__Co_contig_3086_pilon_pilon | Siphoviridae | 0 | 0 | NA | 3 | 0 | orphan | 0 | 0 | NA | 0 | 0 | NA |
| vOTU_sg_2986__S10_k141_149399 | Myoviridae | 0 | 0 | NA | 1 | 0 | orphan | 0 | 0 | NA | 0 | 0 | NA |
| vOTU_sg_2996__Co_k141_389227 | Unknown_Virus | 0 | 0 | NA | 1 | 0 | orphan | 0 | 0 | NA | 0 | 0 | NA |
| vOTU_sg_3025__S2_k141_575334 | Phycodnaviridae | 0 | 0 | NA | 1 | 0 | orphan | 0 | 0 | NA | 0 | 0 | NA |
| vOTU_sg_3059__S11_k141_492267 | Unclassified_Caudovirales | 0 | 0 | NA | 2 | 0 | orphan | 0 | 0 | NA | 0 | 0 | NA |
| vOTU_sg_3064__Co_contig_4009_pilon_pilon | Siphoviridae | 0 | 0 | NA | 1 | 0 | orphan | 0 | 0 | NA | 0 | 0 | NA |

| | | | | | | | | | | | | | |
|---|---|---|---|---|---|---|---|---|---|---|---|---|---|
| vOTU_sg_3079__Co_contig_2097_pilon_pilon | Myoviridae | 0 | 1 | NA | 0 | 0 | NA | 0 | 0 | NA | 0 | 0 | NA |
| vOTU_sg_3112__S7_k141_160638 | Myoviridae | 0 | 0 | NA | 1 | 0 | orphan | 0 | 0 | NA | 0 | 0 | NA |
| vOTU_sg_3189__Co_k141_2765900 | Phycodnaviridae | 0 | 0 | NA | 1 | 0 | orphan | 0 | 0 | NA | 0 | 0 | NA |
| vOTU_sg_3198__S5_k141_76080 | Myoviridae | 0 | 0 | NA | 1 | 0 | orphan | 0 | 0 | NA | 0 | 0 | NA |
| vOTU_sg_3213__S3_k141_301342 | Unclassified_Caudovirales | 0 | 0 | NA | 1 | 0 | orphan | 0 | 0 | NA | 0 | 0 | NA |
| vOTU_sg_3217__S8_k141_136717-circular | Siphoviridae | 0 | 0 | NA | 1 | 0 | orphan | 0 | 0 | NA | 0 | 0 | NA |
| vOTU_sg_3272__S10_k141_358398 | Unclassified_Caudovirales | 0 | 1 | NA | 0 | 0 | NA | 0 | 0 | NA | 0 | 0 | NA |
| vOTU_sg_3291__S10_k141_562051 | Podoviridae | 0 | 0 | NA | 1 | 0 | orphan | 0 | 0 | NA | 0 | 0 | NA |
| vOTU_sg_3302__Co_k141_975787 | Myoviridae | 0 | 1 | NA | 0 | 0 | NA | 0 | 0 | NA | 0 | 0 | NA |
| vOTU_sg_3307__Co_contig_2498_pilon_pilon | Podoviridae | 0 | 0 | NA | 1 | 0 | orphan | 0 | 0 | NA | 0 | 0 | NA |
| vOTU_sg_3328__S8_k141_354379 | Unknown_Virus | 0 | 0 | NA | 1 | 0 | orphan | 0 | 0 | NA | 0 | 0 | NA |
| vOTU_sg_3334__S3_k141_66352 | Podoviridae | 0 | 0 | NA | 1 | 0 | orphan | 0 | 0 | NA | 0 | 0 | NA |
| vOTU_sg_3339__S8_contig_34_pilon_pilon | Myoviridae | 0 | 1 | NA | 0 | 0 | NA | 0 | 0 | NA | 0 | 0 | NA |
| vOTU_sg_3356__Co_contig_3462_pilon_pilon | Podoviridae | 0 | 0 | NA | 1 | 0 | orphan | 0 | 0 | NA | 0 | 0 | NA |
| vOTU_sg_3373__S5_k141_448354 | Unclassified_Caudovirales | 0 | 0 | NA | 1 | 0 | orphan | 0 | 0 | NA | 0 | 0 | NA |
| vOTU_sg_3380__S8_k141_94712 | Myoviridae | 0 | 0 | NA | 1 | 0 | orphan | 0 | 0 | NA | 0 | 0 | NA |
| vOTU_sg_3398__Co_k141_706091 | Unknown_Virus | 0 | 0 | NA | 1 | 0 | orphan | 0 | 0 | NA | 0 | 0 | NA |
| vOTU_sg_3399__Co_contig_4967_pilon_pilon | Phycodnaviridae | 0 | 0 | NA | 1 | 0 | orphan | 0 | 0 | NA | 0 | 0 | NA |
| vOTU_sg_3412__Co_contig_3514_pilon_pilon | Unknown_Virus | 0 | 0 | NA | 1 | 0 | orphan | 0 | 0 | NA | 0 | 0 | NA |
| vOTU_sg_3422__Co_k141_2315511 | Myoviridae | 0 | 0 | NA | 1 | 0 | orphan | 0 | 0 | NA | 0 | 0 | NA |
| vOTU_sg_3448__S5_k141_307234 | Myoviridae | 0 | 1 | NA | 1 | 0 | orphan | 0 | 0 | NA | 0 | 0 | NA |
| vOTU_sg_3476__S8_k141_147874 | Unclassified_Caudovirales | 0 | 0 | NA | 1 | 0 | orphan | 0 | 0 | NA | 0 | 0 | NA |
| vOTU_sg_349__S8_contig_213_pilon_pilon | Myoviridae | 0 | 0 | NA | 1 | 0 | orphan | 0 | 0 | NA | 0 | 0 | NA |

| | | | | | | | | | | | | | |
|---|---|---|---|---|---|---|---|---|---|---|---|---|---|
| vOTU_sg_3491__S11_k141_786398 | Unclassified_Caudovirales | 0 | 0 | NA | 1 | 0 | orphan | 0 | 0 | NA | 0 | 0 | NA |
| vOTU_sg_3518__Co_contig_1904_pilon_pilon | Siphoviridae | 0 | 0 | NA | 1 | 0 | orphan | 0 | 0 | NA | 0 | 0 | NA |
| vOTU_sg_3522__Co_k141_1840531-circular | Unknown_Virus | 0 | 0 | NA | 1 | 0 | orphan | 0 | 0 | NA | 0 | 0 | NA |
| vOTU_sg_3523__S8_k141_177172-circular | Unknown_Virus | 0 | 0 | NA | 2 | 0 | orphan | 0 | 0 | NA | 0 | 0 | NA |
| vOTU_sg_3559__S11_k141_545449 | Unclassified_Caudovirales | 0 | 0 | NA | 1 | 0 | orphan | 0 | 0 | NA | 0 | 0 | NA |
| vOTU_sg_3579__Co_k141_1976971 | Siphoviridae | 0 | 0 | NA | 1 | 0 | orphan | 0 | 0 | NA | 0 | 0 | NA |
| vOTU_sg_3593__Co_k141_44372 | Siphoviridae | 0 | 0 | NA | 1 | 0 | orphan | 0 | 0 | NA | 0 | 0 | NA |
| vOTU_sg_3606__S9_k141_546660 | Unclassified_Caudovirales | 0 | 0 | NA | 1 | 0 | orphan | 0 | 0 | NA | 0 | 0 | NA |
| vOTU_sg_3611__S11_k141_464486 | Unclassified_Caudovirales | 0 | 0 | NA | 1 | 0 | orphan | 0 | 0 | NA | 0 | 0 | NA |
| vOTU_sg_3621__Co_k141_46150 | Podoviridae | 0 | 0 | NA | 1 | 0 | orphan | 0 | 0 | NA | 0 | 0 | NA |
| vOTU_sg_3646__S8_contig_378_pilon_pilon | Siphoviridae | 0 | 0 | NA | 1 | 0 | orphan | 0 | 0 | NA | 0 | 0 | NA |
| vOTU_sg_3667__S11_k141_22830 | Siphoviridae | 0 | 0 | NA | 1 | 0 | orphan | 0 | 0 | NA | 0 | 0 | NA |
| vOTU_sg_3674__Co_k141_868189 | Myoviridae | 0 | 0 | NA | 1 | 0 | orphan | 0 | 0 | NA | 0 | 0 | NA |
| vOTU_sg_3690__Co_k141_1261843 | Unknown_Virus | 0 | 0 | NA | 1 | 0 | orphan | 0 | 0 | NA | 0 | 0 | NA |
| vOTU_sg_3714__S7_contig_239_pilon_pilon | Myoviridae | 0 | 1 | NA | 0 | 0 | NA | 0 | 0 | NA | 0 | 0 | NA |
| vOTU_sg_3715__Co_contig_3745_pilon_pilon | Unclassified_Caudovirales | 0 | 0 | NA | 1 | 0 | orphan | 0 | 0 | NA | 0 | 0 | NA |
| vOTU_sg_3748__S5_k141_79972 | Unknown_Virus | 0 | 0 | NA | 2 | 0 | orphan | 0 | 0 | NA | 0 | 0 | NA |
| vOTU_sg_3752__S6_k141_298427 | Unknown_Virus | 0 | 0 | NA | 1 | 0 | orphan | 0 | 0 | NA | 0 | 0 | NA |
| vOTU_sg_3755__Co_contig_1028_pilon_pilon | Siphoviridae | 0 | 0 | NA | 1 | 0 | orphan | 0 | 0 | NA | 0 | 0 | NA |
| vOTU_sg_3758__Co_k141_2826655 | Phycodnaviridae | 0 | 0 | NA | 1 | 0 | orphan | 0 | 0 | NA | 0 | 0 | NA |
| vOTU_sg_3791__S3_k141_287092 | Siphoviridae | 0 | 0 | NA | 1 | 0 | orphan | 0 | 0 | NA | 0 | 0 | NA |
| vOTU_sg_3814__S9_k141_98376 | Podoviridae | 0 | 0 | NA | 1 | 0 | orphan | 0 | 0 | NA | 0 | 0 | NA |
| vOTU_sg_3823__Co_k141_2290519 | Unclassified_Caudovirales | 0 | 0 | NA | 1 | 0 | orphan | 0 | 0 | NA | 0 | 0 | NA |
| vOTU_sg_3826__Co_k141_585763 | Siphoviridae | 0 | 0 | NA | 1 | 0 | orphan | 0 | 0 | NA | 0 | 0 | NA |

| | | | | | | | | | | | | | |
|---|---|---|---|---|---|---|---|---|---|---|---|---|---|
| vOTU_sg_3848__S11_contig_1_pilon_pilon | Siphoviridae | 0 | 0 | NA | 1 | 0 | orphan | 0 | 0 | NA | 0 | 0 | NA |
| vOTU_sg_3850__S7_k141_453366 | Unknown_Virus | 0 | 0 | NA | 1 | 0 | orphan | 0 | 0 | NA | 0 | 0 | NA |
| vOTU_sg_3853__S6_k141_722688 | Siphoviridae | 0 | 0 | NA | 0 | 0 | NA | 0 | 0 | NA | 0 | 1 | orphan |
| vOTU_sg_3857__Co_k141_2102330 | Siphoviridae | 0 | 0 | NA | 1 | 0 | orphan | 0 | 0 | NA | 0 | 0 | NA |
| vOTU_sg_3884__S2_contig_79_pilon_pilon | Siphoviridae | 0 | 0 | NA | 1 | 0 | orphan | 0 | 0 | NA | 0 | 0 | NA |
| vOTU_sg_3894__S10_k141_188451 | Myoviridae | 0 | 0 | NA | 1 | 0 | orphan | 0 | 0 | NA | 0 | 0 | NA |
| vOTU_sg_3905__S3_k141_557220 | Unclassified_Caudovirales | 0 | 0 | NA | 1 | 0 | orphan | 0 | 0 | NA | 0 | 0 | NA |
| vOTU_sg_3928__Co_contig_61_pilon_pilon | Unclassified_Caudovirales | 0 | 0 | NA | 1 | 0 | orphan | 0 | 0 | NA | 0 | 0 | NA |
| vOTU_sg_3969__S6_k141_541471 | Unclassified_Caudovirales | 0 | 0 | NA | 1 | 0 | orphan | 0 | 0 | NA | 0 | 0 | NA |
| vOTU_sg_3975__Co_contig_3736_pilon_pilon | Siphoviridae | 0 | 0 | NA | 1 | 0 | orphan | 0 | 0 | NA | 0 | 0 | NA |
| vOTU_sg_4011__Co_k141_451587 | Phycodnaviridae | 0 | 0 | NA | 2 | 0 | orphan | 0 | 0 | NA | 0 | 0 | NA |
| vOTU_sg_4012__S9_k141_247686 | Podoviridae | 0 | 0 | NA | 1 | 0 | orphan | 0 | 0 | NA | 0 | 0 | NA |
| vOTU_sg_403__S3_k141_67787 | Siphoviridae | 0 | 0 | NA | 1 | 0 | orphan | 0 | 0 | NA | 0 | 0 | NA |
| vOTU_sg_4054__S11_k141_767117-circular | Unknown_Virus | 0 | 0 | NA | 1 | 0 | orphan | 0 | 0 | NA | 0 | 0 | NA |
| vOTU_sg_4064__Co_k141_193576 | Phycodnaviridae | 0 | 0 | NA | 1 | 0 | orphan | 0 | 0 | NA | 0 | 0 | NA |
| vOTU_sg_4140__S11_contig_392_pilon_pilon | Podoviridae | 0 | 0 | NA | 1 | 0 | orphan | 0 | 0 | NA | 0 | 0 | NA |
| vOTU_sg_4151__Co_k141_894923 | Siphoviridae | 0 | 0 | NA | 1 | 0 | orphan | 0 | 0 | NA | 0 | 0 | NA |
| vOTU_sg_4214__S11_contig_538_pilon_pilon | Myoviridae | 0 | 0 | NA | 1 | 0 | orphan | 0 | 0 | NA | 0 | 0 | NA |
| vOTU_sg_4220__S6_k141_254751 | Siphoviridae | 0 | 0 | NA | 1 | 0 | orphan | 0 | 0 | NA | 0 | 0 | NA |
| vOTU_sg_4229__S3_k141_521239 | Phycodnaviridae | 0 | 0 | NA | 1 | 0 | orphan | 0 | 0 | NA | 0 | 0 | NA |
| vOTU_sg_4250__Co_contig_3914_pilon_pilon | Podoviridae | 0 | 0 | NA | 1 | 0 | orphan | 0 | 0 | NA | 0 | 0 | NA |
| vOTU_sg_4253__Co_k141_2225356 | Phycodnaviridae | 0 | 0 | NA | 1 | 0 | orphan | 0 | 0 | NA | 0 | 0 | NA |
| vOTU_sg_4267__S11_k141_535869 | Phycodnaviridae | 0 | 0 | NA | 1 | 0 | orphan | 0 | 0 | NA | 0 | 0 | NA |

| | | | | | | | | | | | | | |
|---|---|---|---|---|---|---|---|---|---|---|---|---|---|
| vOTU_sg_4327__Co_k141_1282565 | Unknown_Virus | 0 | 0 | NA | 1 | 0 | orphan | 0 | 0 | NA | 0 | 0 | NA |
| vOTU_sg_4337__Co_k141_2073906 | Myoviridae | 0 | 1 | NA | 0 | 0 | NA | 0 | 0 | NA | 0 | 0 | NA |
| vOTU_sg_4355__Co_k141_36109 | Phycodnaviridae | 0 | 0 | NA | 1 | 0 | orphan | 0 | 0 | NA | 0 | 0 | NA |
| vOTU_sg_4367__S6_k141_508290-circular | Unclassified_Caudovirales | 0 | 0 | NA | 1 | 0 | orphan | 0 | 0 | NA | 0 | 0 | NA |
| vOTU_sg_4384__Co_k141_1639153 | Unclassified_Caudovirales | 0 | 0 | NA | 1 | 0 | orphan | 0 | 0 | NA | 0 | 0 | NA |
| vOTU_sg_4392__S5_contig_325_pilon_pilon | Siphoviridae | 0 | 0 | NA | 1 | 0 | orphan | 0 | 0 | NA | 0 | 0 | NA |
| vOTU_sg_4400__S2_k141_24742 | Myoviridae | 0 | 1 | NA | 0 | 0 | NA | 0 | 0 | NA | 0 | 0 | NA |
| vOTU_sg_4408__S10_k141_591287 | Myoviridae | 0 | 0 | NA | 1 | 0 | orphan | 0 | 0 | NA | 0 | 0 | NA |
| vOTU_sg_4431__Co_k141_6700 | Phycodnaviridae | 0 | 0 | NA | 0 | 0 | NA | 0 | 0 | NA | 0 | 1 | orphan |
| vOTU_sg_4439__Co_contig_6178_pilon_pilon | Unclassified_Caudovirales | 0 | 0 | NA | 1 | 0 | orphan | 0 | 0 | NA | 0 | 0 | NA |
| vOTU_sg_4476__S2_k141_73226 | Unclassified_Caudovirales | 0 | 0 | NA | 1 | 0 | orphan | 0 | 0 | NA | 0 | 0 | NA |
| vOTU_sg_4551__Co_k141_621093 | Siphoviridae | 0 | 0 | NA | 1 | 0 | orphan | 0 | 0 | NA | 0 | 0 | NA |
| vOTU_sg_4572__S5_k141_608884 | Unknown_Virus | 0 | 0 | NA | 1 | 0 | orphan | 0 | 0 | NA | 0 | 0 | NA |
| vOTU_sg_4582__S9_k141_172122-circular | Siphoviridae | 0 | 0 | NA | 1 | 0 | orphan | 0 | 0 | NA | 0 | 0 | NA |
| vOTU_sg_4588__Co_k141_828620 | Myoviridae | 0 | 1 | NA | 0 | 0 | NA | 0 | 0 | NA | 0 | 0 | NA |
| vOTU_sg_4611__Co_contig_2801_pilon_pilon | Unclassified_Caudovirales | 0 | 0 | NA | 1 | 0 | orphan | 0 | 0 | NA | 0 | 0 | NA |
| vOTU_sg_4649__S4_contig_580_pilon_pilon | Siphoviridae | 0 | 0 | NA | 1 | 0 | orphan | 0 | 0 | NA | 0 | 0 | NA |
| vOTU_sg_4668__Co_k141_1788030 | Podoviridae | 0 | 0 | NA | 1 | 0 | orphan | 0 | 0 | NA | 0 | 0 | NA |
| vOTU_sg_4691__S11_k141_455759 | Unknown_Virus | 0 | 1 | NA | 0 | 0 | NA | 0 | 0 | NA | 0 | 0 | NA |
| vOTU_sg_4734__S8_k141_219803 | Siphoviridae | 0 | 0 | NA | 2 | 0 | orphan | 0 | 0 | NA | 0 | 0 | NA |
| vOTU_sg_4753__S5_k141_54077 | Myoviridae | 0 | 0 | NA | 1 | 0 | orphan | 0 | 0 | NA | 0 | 0 | NA |
| vOTU_sg_4764__S10_k141_208531 | Phycodnaviridae | 0 | 0 | NA | 1 | 0 | orphan | 0 | 0 | NA | 0 | 0 | NA |
| vOTU_sg_4780__Co_contig_6082_pilon_pilon | Siphoviridae | 0 | 0 | NA | 1 | 0 | orphan | 0 | 0 | NA | 0 | 0 | NA |

| | | | | | | | | | | | | | |
|---|---|---|---|---|---|---|---|---|---|---|---|---|---|
| vOTU_sg_4791__S2_k141_538836-circular | Siphoviridae | 0 | 0 | NA | 1 | 0 | orphan | 0 | 0 | NA | 0 | 0 | NA |
| vOTU_sg_4796__Co_k141_914907 | Unclassified_Caudovirales | 0 | 0 | NA | 1 | 0 | orphan | 0 | 0 | NA | 0 | 0 | NA |
| vOTU_sg_4867__S10_k141_28320 | Siphoviridae | 0 | 0 | NA | 1 | 0 | orphan | 0 | 0 | NA | 0 | 0 | NA |
| vOTU_sg_4886__S3_k141_65250 | Unclassified_Caudovirales | 0 | 0 | NA | 1 | 0 | orphan | 0 | 0 | NA | 0 | 0 | NA |
| vOTU_sg_491__S8_k141_412961 | Myoviridae | 0 | 0 | NA | 1 | 0 | orphan | 0 | 0 | NA | 0 | 0 | NA |
| vOTU_sg_4968__S6_k141_546788 | Unknown_Virus | 0 | 0 | NA | 1 | 0 | orphan | 0 | 0 | NA | 0 | 0 | NA |
| vOTU_sg_4976__Co_contig_3965_pilon_pilon | Siphoviridae | 0 | 0 | NA | 1 | 0 | orphan | 0 | 0 | NA | 0 | 0 | NA |
| vOTU_sg_4978__Co_contig_7607_pilon_pilon | Unclassified_Caudovirales | 0 | 0 | NA | 1 | 0 | orphan | 0 | 0 | NA | 0 | 0 | NA |
| vOTU_sg_5006__S11_contig_276_pilon_pilon | Myoviridae | 0 | 1 | NA | 2 | 0 | orphan | 0 | 0 | NA | 0 | 0 | NA |
| vOTU_sg_5021__Co_contig_7034_pilon_pilon | Unclassified_Caudovirales | 0 | 0 | NA | 1 | 0 | orphan | 0 | 0 | NA | 0 | 0 | NA |
| vOTU_sg_5027__S3_k141_519174 | Siphoviridae | 0 | 0 | NA | 1 | 0 | orphan | 0 | 0 | NA | 0 | 0 | NA |
| vOTU_sg_5031__Co_contig_7611_pilon_pilon | Unclassified_Caudovirales | 0 | 0 | NA | 1 | 0 | orphan | 0 | 0 | NA | 0 | 0 | NA |
| vOTU_sg_5043__Co_contig_6678_pilon_pilon | Siphoviridae | 0 | 0 | NA | 1 | 0 | orphan | 0 | 0 | NA | 0 | 0 | NA |
| vOTU_sg_5086__Co_k141_2880682 | Iridoviridae | 0 | 0 | NA | 1 | 0 | orphan | 0 | 0 | NA | 0 | 0 | NA |
| vOTU_sg_5129__Co_contig_4896_pilon_pilon | Myoviridae | 0 | 0 | NA | 2 | 0 | orphan | 0 | 0 | NA | 0 | 0 | NA |
| vOTU_sg_5152__S2_k141_639214 | Unclassified_Caudovirales | 0 | 0 | NA | 1 | 0 | orphan | 0 | 0 | NA | 0 | 0 | NA |
| vOTU_sg_5153__Co_contig_994_pilon_pilon | Unclassified_Caudovirales | 0 | 0 | NA | 1 | 0 | orphan | 0 | 0 | NA | 0 | 0 | NA |
| vOTU_sg_5163__Co_k141_832535 | Podoviridae | 0 | 0 | NA | 1 | 0 | orphan | 0 | 0 | NA | 0 | 0 | NA |
| vOTU_sg_5185__S4_k141_374897 | Podoviridae | 0 | 0 | NA | 1 | 0 | orphan | 0 | 0 | NA | 0 | 0 | NA |
| vOTU_sg_5207__Co_contig_6967_pilon_pilon | Podoviridae | 0 | 0 | NA | 1 | 0 | orphan | 0 | 0 | NA | 0 | 0 | NA |
| vOTU_sg_5271__S7_k141_143179 | Unclassified_Caudovirales | 0 | 0 | NA | 1 | 0 | orphan | 0 | 0 | NA | 0 | 0 | NA |
| vOTU_sg_5272__S10_k141_532010 | Podoviridae | 0 | 0 | NA | 1 | 0 | orphan | 0 | 0 | NA | 0 | 0 | NA |

| | | | | | | | | | | | | | |
|---|---|---|---|---|---|---|---|---|---|---|---|---|---|
| vOTU_sg_5283__S9_k141_499429 | Unknown_Virus | 0 | 0 | NA | 1 | 0 | orphan | 0 | 0 | NA | 0 | 0 | NA |
| vOTU_sg_5304__Co_contig_7685_pilon_pilon | Phycodnaviridae | 0 | 0 | NA | 1 | 0 | orphan | 0 | 0 | NA | 0 | 0 | NA |
| vOTU_sg_5305__S10_k141_49399 | Unknown_Virus | 0 | 0 | NA | 1 | 0 | orphan | 0 | 0 | NA | 0 | 0 | NA |
| vOTU_sg_5372__S5_k141_246655 | Myoviridae | 0 | 0 | NA | 1 | 0 | orphan | 0 | 0 | NA | 0 | 0 | NA |
| vOTU_sg_5424__S4_contig_296_pilon_pilon | Siphoviridae | 0 | 0 | NA | 1 | 0 | orphan | 0 | 0 | NA | 0 | 0 | NA |
| vOTU_sg_5436__Co_contig_2574_pilon_pilon | Podoviridae | 0 | 0 | NA | 1 | 0 | orphan | 0 | 0 | NA | 0 | 0 | NA |
| vOTU_sg_5473__S8_contig_338_pilon_pilon | Unknown_Virus | 0 | 0 | NA | 1 | 0 | orphan | 0 | 0 | NA | 0 | 0 | NA |
| vOTU_sg_5514__Co_contig_4170_pilon_pilon | Phycodnaviridae | 0 | 0 | NA | 1 | 0 | orphan | 0 | 0 | NA | 0 | 0 | NA |
| vOTU_sg_5576__Co_k141_568825 | Myoviridae | 0 | 0 | NA | 1 | 0 | orphan | 0 | 0 | NA | 0 | 0 | NA |
| vOTU_sg_5597__Co_k141_1670894 | Unknown_Virus | 0 | 0 | NA | 2 | 0 | orphan | 0 | 0 | NA | 0 | 0 | NA |
| vOTU_sg_5628__Co_contig_7013_pilon_pilon | Myoviridae | 0 | 1 | NA | 1 | 0 | orphan | 0 | 0 | NA | 0 | 0 | NA |
| vOTU_sg_5648__S4_k141_302979 | Unknown_Virus | 0 | 0 | NA | 1 | 0 | orphan | 0 | 0 | NA | 0 | 0 | NA |
| vOTU_sg_5691__S9_k141_384924 | Phycodnaviridae | 0 | 0 | NA | 1 | 0 | orphan | 0 | 0 | NA | 0 | 0 | NA |
| vOTU_sg_5705__S2_k141_277740 | Siphoviridae | 0 | 0 | NA | 2 | 0 | orphan | 0 | 0 | NA | 0 | 0 | NA |
| vOTU_sg_5710__S11_k141_754352 | Unclassified_Caudovirales | 0 | 0 | NA | 1 | 0 | orphan | 0 | 0 | NA | 0 | 0 | NA |
| vOTU_sg_5715__S6_k141_500391 | Unclassified_Caudovirales | 0 | 0 | NA | 1 | 0 | orphan | 0 | 0 | NA | 0 | 0 | NA |
| vOTU_sg_5728__Co_k141_2973164 | Siphoviridae | 0 | 0 | NA | 1 | 0 | orphan | 0 | 0 | NA | 0 | 0 | NA |
| vOTU_sg_5755__S11_contig_236_pilon_pilon | Unclassified_Caudovirales | 0 | 0 | NA | 1 | 0 | orphan | 0 | 0 | NA | 0 | 0 | NA |
| vOTU_sg_5758__S3_k141_306935 | Siphoviridae | 0 | 0 | NA | 1 | 0 | orphan | 0 | 0 | NA | 0 | 0 | NA |
| vOTU_sg_5784__Co_contig_1753_pilon_pilon | Myoviridae | 0 | 1 | NA | 0 | 0 | NA | 0 | 0 | NA | 0 | 0 | NA |
| vOTU_sg_5785__S10_k141_64151 | Podoviridae | 0 | 0 | NA | 1 | 0 | orphan | 0 | 0 | NA | 0 | 0 | NA |
| vOTU_sg_5828__Co_k141_159323 | Myoviridae | 0 | 0 | NA | 1 | 0 | orphan | 0 | 0 | NA | 0 | 0 | NA |
| vOTU_sg_5875__S6_k141_191319 | Siphoviridae | 0 | 0 | NA | 2 | 0 | orphan | 0 | 0 | NA | 0 | 0 | NA |

| | | | | | | | | | | | | | |
|---|---|---|---|---|---|---|---|---|---|---|---|---|---|
| vOTU_sg_5919__Co_k141_1619408 | Unclassified_Caudovirales | 0 | 0 | NA | 1 | 0 | orphan | 0 | 0 | NA | 0 | 0 | NA |
| vOTU_sg_5920__Co_contig_5838_pilon_pilon | Podoviridae | 0 | 0 | NA | 1 | 0 | orphan | 0 | 0 | NA | 0 | 0 | NA |
| vOTU_sg_5935__S8_k141_229660 | Siphoviridae | 0 | 0 | NA | 2 | 0 | orphan | 0 | 0 | NA | 0 | 0 | NA |
| vOTU_sg_5977__S4_k141_115930 | Myoviridae | 0 | 0 | NA | 1 | 0 | orphan | 0 | 0 | NA | 0 | 0 | NA |
| vOTU_sg_6011__S4_k141_70341 | Unclassified_Caudovirales | 0 | 0 | NA | 1 | 0 | orphan | 0 | 0 | NA | 0 | 0 | NA |
| vOTU_sg_6094__Co_k141_112688 | Myoviridae | 0 | 0 | NA | 1 | 0 | orphan | 0 | 0 | NA | 0 | 0 | NA |
| vOTU_sg_6143__S11_contig_40_pilon_pilon | Siphoviridae | 0 | 0 | NA | 1 | 0 | orphan | 0 | 0 | NA | 0 | 0 | NA |
| vOTU_sg_6163__S11_k141_140482 | Unclassified_Caudovirales | 0 | 0 | NA | 1 | 0 | orphan | 0 | 0 | NA | 0 | 0 | NA |
| vOTU_sg_6198__S7_k141_393549 | Siphoviridae | 0 | 0 | NA | 1 | 0 | orphan | 0 | 0 | NA | 0 | 0 | NA |
| vOTU_sg_6202__Co_k141_2240107 | Myoviridae | 0 | 0 | NA | 1 | 0 | orphan | 0 | 0 | NA | 0 | 0 | NA |
| vOTU_sg_6206__S11_contig_459_pilon_pilon | Myoviridae | 0 | 0 | NA | 1 | 0 | orphan | 0 | 0 | NA | 0 | 0 | NA |
| vOTU_sg_6246__S9_k141_87935 | Siphoviridae | 0 | 0 | NA | 1 | 0 | orphan | 0 | 0 | NA | 0 | 0 | NA |
| vOTU_sg_6256__S10_k141_414648 | Siphoviridae | 0 | 0 | NA | 2 | 0 | orphan | 0 | 0 | NA | 0 | 0 | NA |
| vOTU_sg_6261__Co_k141_1309302 | Unknown_Virus | 0 | 0 | NA | 1 | 0 | orphan | 0 | 0 | NA | 0 | 0 | NA |
| vOTU_sg_6282__S8_k141_329843-circular | Siphoviridae | 0 | 0 | NA | 1 | 0 | orphan | 0 | 0 | NA | 0 | 0 | NA |
| vOTU_sg_6298__S8_k141_161473 | Siphoviridae | 0 | 0 | NA | 1 | 0 | orphan | 0 | 0 | NA | 0 | 0 | NA |
| vOTU_sg_6304__Co_contig_6865_pilon_pilon | Unknown_Virus | 0 | 0 | NA | 1 | 0 | orphan | 0 | 0 | NA | 0 | 0 | NA |
| vOTU_sg_6378__S4_k141_87135 | Unknown_Virus | 0 | 0 | NA | 1 | 0 | orphan | 0 | 0 | NA | 0 | 0 | NA |
| vOTU_sg_6390__S8_k141_246924-circular | Unclassified_Caudovirales | 0 | 0 | NA | 1 | 0 | orphan | 0 | 0 | NA | 0 | 0 | NA |
| vOTU_sg_6465__Co_contig_2455_pilon_pilon | Siphoviridae | 0 | 0 | NA | 1 | 0 | orphan | 0 | 0 | NA | 0 | 0 | NA |
| vOTU_sg_6474__S3_k141_216147 | Unclassified_Caudovirales | 0 | 0 | NA | 1 | 0 | orphan | 0 | 0 | NA | 0 | 0 | NA |
| vOTU_sg_6494__S2_k141_31600 | Phycodnaviridae | 0 | 0 | NA | 1 | 0 | orphan | 0 | 0 | NA | 0 | 0 | NA |
| vOTU_sg_6495__Co_k141_2133485-circular | Podoviridae | 0 | 0 | NA | 1 | 0 | orphan | 0 | 0 | NA | 0 | 0 | NA |

| | | | | | | | | | | | | | |
|---|---|---|---|---|---|---|---|---|---|---|---|---|---|
| vOTU_sg_6507__S5_k141_329986 | Siphoviridae | 0 | 0 | NA | 1 | 0 | orphan | 0 | 0 | NA | 0 | 0 | NA |
| vOTU_sg_6544__S11_k141_610848 | Unclassified_Caudovirales | 0 | 0 | NA | 1 | 0 | orphan | 0 | 0 | NA | 0 | 0 | NA |
| vOTU_sg_6571__Co_k141_479190 | Phycodnaviridae | 0 | 0 | NA | 1 | 0 | orphan | 0 | 0 | NA | 0 | 0 | NA |
| vOTU_sg_6582__Co_contig_3490_pilon_pilon | Siphoviridae | 0 | 0 | NA | 1 | 0 | orphan | 0 | 0 | NA | 0 | 0 | NA |
| vOTU_sg_6611__Co_k141_2804683 | Myoviridae | 0 | 0 | NA | 1 | 0 | orphan | 0 | 0 | NA | 0 | 0 | NA |
| vOTU_sg_6615__Co_k141_2623286 | Phycodnaviridae | 0 | 0 | NA | 1 | 0 | orphan | 0 | 0 | NA | 0 | 0 | NA |
| vOTU_sg_6647__S4_k141_318000 | Siphoviridae | 0 | 0 | NA | 1 | 0 | orphan | 0 | 0 | NA | 0 | 0 | NA |
| vOTU_sg_6692__S10_contig_88_pilon_pilon | Siphoviridae | 0 | 0 | NA | 1 | 0 | orphan | 0 | 0 | NA | 0 | 0 | NA |
| vOTU_sg_6704__Co_k141_1788287-circular | Siphoviridae | 0 | 0 | NA | 1 | 0 | orphan | 0 | 0 | NA | 0 | 0 | NA |
| vOTU_sg_6726__Co_k141_1120636 | Podoviridae | 0 | 0 | NA | 1 | 0 | orphan | 0 | 0 | NA | 0 | 0 | NA |
| vOTU_sg_6737__S10_k141_542305 | Phycodnaviridae | 0 | 0 | NA | 1 | 0 | orphan | 0 | 0 | NA | 0 | 0 | NA |
| vOTU_sg_6758__Co_contig_2218_pilon_pilon | Podoviridae | 0 | 0 | NA | 1 | 0 | orphan | 0 | 0 | NA | 0 | 0 | NA |
| vOTU_sg_6766__Co_k141_1374476 | Siphoviridae | 0 | 0 | NA | 1 | 0 | orphan | 0 | 0 | NA | 0 | 0 | NA |
| vOTU_sg_6779__Co_contig_4591_pilon_pilon | Siphoviridae | 0 | 0 | NA | 1 | 0 | orphan | 0 | 0 | NA | 0 | 0 | NA |
| vOTU_sg_6780__Co_k141_1865030 | Myoviridae | 0 | 1 | NA | 0 | 0 | NA | 0 | 0 | NA | 0 | 0 | NA |
| vOTU_sg_6811__S2_k141_289786 | Siphoviridae | 0 | 0 | NA | 2 | 0 | orphan | 0 | 0 | NA | 0 | 0 | NA |
| vOTU_sg_6840__Co_contig_3235_pilon_pilon | Myoviridae | 0 | 0 | NA | 1 | 0 | orphan | 0 | 0 | NA | 0 | 0 | NA |
| vOTU_sg_6924__Co_k141_1113502 | Podoviridae | 0 | 0 | NA | 1 | 0 | orphan | 0 | 0 | NA | 0 | 0 | NA |
| vOTU_sg_6975__Co_contig_6563_pilon_pilon | Unclassified_Caudovirales | 0 | 0 | NA | 1 | 0 | orphan | 0 | 0 | NA | 0 | 0 | NA |
| vOTU_sg_7079__Co_k141_627482 | Siphoviridae | 0 | 0 | NA | 1 | 0 | orphan | 0 | 0 | NA | 0 | 0 | NA |
| vOTU_sg_7111__Co_contig_4725_pilon_pilon | Unclassified_Caudovirales | 0 | 0 | NA | 1 | 0 | orphan | 0 | 0 | NA | 0 | 0 | NA |
| vOTU_sg_7173__S10_k141_108971 | Podoviridae | 0 | 0 | NA | 1 | 0 | orphan | 0 | 0 | NA | 0 | 0 | NA |
| vOTU_sg_7266__Co_contig_7372_pilon_pilon | Siphoviridae | 0 | 0 | NA | 1 | 0 | orphan | 0 | 0 | NA | 0 | 0 | NA |

| | | | | | | | | | | | | | |
|---|---|---|---|---|---|---|---|---|---|---|---|---|---|
| vOTU_sg_7301__S2_contig_230_pilon_pilon | Siphoviridae | 0 | 0 | NA | 1 | 0 | orphan | 0 | 0 | NA | 0 | 0 | NA |
| vOTU_sg_7312__S11_k141_317962 | Siphoviridae | 0 | 0 | NA | 1 | 0 | orphan | 0 | 0 | NA | 0 | 0 | NA |
| vOTU_sg_7344__S3_k141_51914 | Unclassified_Caudovirales | 0 | 0 | NA | 1 | 0 | orphan | 0 | 0 | NA | 0 | 0 | NA |
| vOTU_sg_7351__S8_k141_216698 | Unclassified_Caudovirales | 0 | 0 | NA | 1 | 0 | orphan | 0 | 0 | NA | 0 | 0 | NA |
| vOTU_sg_7352__S10_k141_345081-circular | Unclassified_Caudovirales | 0 | 0 | NA | 0 | 0 | NA | 0 | 0 | NA | 0 | 1 | orphan |
| vOTU_sg_7428__S8_k141_186256 | Siphoviridae | 0 | 0 | NA | 1 | 0 | orphan | 0 | 0 | NA | 0 | 0 | NA |
| vOTU_sg_7440__S2_k141_694257 | Phycodnaviridae | 0 | 0 | NA | 1 | 0 | orphan | 0 | 0 | NA | 0 | 0 | NA |
| vOTU_sg_7447__S6_k141_424039 | Myoviridae | 0 | 1 | NA | 0 | 0 | NA | 0 | 0 | NA | 0 | 0 | NA |
| vOTU_sg_7448__Co_k141_1199672 | Siphoviridae | 0 | 0 | NA | 1 | 0 | orphan | 0 | 0 | NA | 0 | 0 | NA |
| vOTU_sg_7464__S8_k141_281944 | Siphoviridae | 0 | 0 | NA | 1 | 0 | orphan | 0 | 0 | NA | 0 | 0 | NA |
| vOTU_sg_7471__Co_k141_3063885 | Siphoviridae | 0 | 0 | NA | 1 | 0 | orphan | 0 | 0 | NA | 0 | 0 | NA |
| vOTU_sg_7507__Co_k141_2469593 | Unclassified_Caudovirales | 0 | 0 | NA | 1 | 0 | orphan | 0 | 0 | NA | 0 | 0 | NA |
| vOTU_sg_7520__Co_contig_3942_pilon_pilon | Podoviridae | 0 | 0 | NA | 2 | 0 | orphan | 0 | 0 | NA | 0 | 0 | NA |
| vOTU_sg_7522__S2_k141_46609 | Unknown_Virus | 0 | 0 | NA | 1 | 0 | orphan | 0 | 0 | NA | 0 | 0 | NA |
| vOTU_sg_7533__S8_k141_213641 | Unclassified_Caudovirales | 0 | 0 | NA | 1 | 0 | orphan | 0 | 0 | NA | 0 | 0 | NA |
| vOTU_sg_7593__S9_k141_360752 | Myoviridae | 0 | 1 | NA | 0 | 0 | NA | 0 | 0 | NA | 0 | 0 | NA |
| vOTU_sg_764__Co_contig_4255_pilon_pilon | Myoviridae | 0 | 1 | NA | 0 | 0 | NA | 0 | 0 | NA | 0 | 0 | NA |
| vOTU_sg_7644__S3_k141_81356 | Unclassified_Caudovirales | 0 | 0 | NA | 1 | 0 | orphan | 0 | 0 | NA | 0 | 0 | NA |
| vOTU_sg_7685__S4_k141_506200 | Myoviridae | 0 | 0 | NA | 1 | 0 | orphan | 0 | 0 | NA | 0 | 0 | NA |
| vOTU_sg_7711__Co_contig_8340_pilon_pilon | Siphoviridae | 0 | 0 | NA | 1 | 0 | orphan | 0 | 0 | NA | 0 | 0 | NA |
| vOTU_sg_7766__Co_contig_2210_pilon_pilon | Myoviridae | 0 | 0 | NA | 1 | 0 | orphan | 0 | 0 | NA | 0 | 0 | NA |
| vOTU_sg_7772__S4_k141_740329 | Myoviridae | 0 | 1 | NA | 0 | 0 | NA | 0 | 0 | NA | 0 | 0 | NA |
| vOTU_sg_7795__Co_contig_2788_pilon_pilon | Siphoviridae | 0 | 0 | NA | 1 | 0 | orphan | 0 | 0 | NA | 0 | 0 | NA |
| vOTU_sg_7802__S2_k141_765720 | Unknown_Virus | 0 | 0 | NA | 1 | 0 | orphan | 0 | 0 | NA | 0 | 0 | NA |

| | | | | | | | | | | | | | |
|---|---|---|---|---|---|---|---|---|---|---|---|---|---|
| vOTU_sg_7821__Co_contig_254_pilon_pilon | Siphoviridae | 0 | 0 | NA | 1 | 0 | orphan | 0 | 0 | NA | 0 | 0 | NA |
| vOTU_sg_7914__S9_contig_411_pilon_pilon | Unclassified_Caudovirales | 0 | 0 | NA | 1 | 0 | orphan | 0 | 0 | NA | 0 | 0 | NA |
| vOTU_sg_7966__S11_k141_22227 | Siphoviridae | 0 | 0 | NA | 1 | 0 | orphan | 0 | 0 | NA | 0 | 0 | NA |
| vOTU_sg_7979__S10_k141_529329 | Siphoviridae | 0 | 0 | NA | 1 | 0 | orphan | 0 | 0 | NA | 0 | 0 | NA |
| vOTU_sg_8064__S10_k141_515727 | Phycodnaviridae | 0 | 0 | NA | 1 | 0 | orphan | 0 | 0 | NA | 0 | 0 | NA |
| vOTU_sg_8084__S9_k141_65045 | Unclassified_Caudovirales | 0 | 0 | NA | 1 | 0 | orphan | 0 | 0 | NA | 0 | 0 | NA |
| vOTU_sg_8091__Co_k141_874615 | Siphoviridae | 0 | 0 | NA | 1 | 0 | orphan | 0 | 0 | NA | 0 | 0 | NA |
| vOTU_sg_8093__S11_k141_425234 | Siphoviridae | 0 | 1 | NA | 0 | 0 | NA | 0 | 0 | NA | 0 | 0 | NA |
| vOTU_sg_8097__S8_k141_152932 | Unknown_Virus | 0 | 1 | NA | 0 | 0 | NA | 0 | 0 | NA | 0 | 0 | NA |
| vOTU_sg_8099__S6_k141_284803 | Podoviridae | 0 | 0 | NA | 1 | 0 | orphan | 0 | 0 | NA | 0 | 0 | NA |
| vOTU_sg_8120__Co_contig_618_pilon_pilon | Siphoviridae | 0 | 0 | NA | 1 | 0 | orphan | 0 | 0 | NA | 0 | 0 | NA |
| vOTU_sg_8177__Co_k141_1873378 | Siphoviridae | 0 | 0 | NA | 1 | 0 | orphan | 0 | 0 | NA | 0 | 0 | NA |
| vOTU_sg_8182__S10_k141_315545 | Podoviridae | 0 | 0 | NA | 1 | 0 | orphan | 0 | 0 | NA | 0 | 0 | NA |
| vOTU_sg_8212__Co_k141_1567871-circular | Siphoviridae | 0 | 0 | NA | 2 | 0 | orphan | 0 | 0 | NA | 0 | 0 | NA |
| vOTU_sg_8214__Co_k141_1229844 | Siphoviridae | 0 | 0 | NA | 1 | 0 | orphan | 0 | 0 | NA | 0 | 0 | NA |
| vOTU_sg_8247__Co_contig_3391_pilon_pilon | Podoviridae | 0 | 0 | NA | 1 | 0 | orphan | 0 | 0 | NA | 0 | 0 | NA |
| vOTU_sg_8277__S11_k141_557365 | Phycodnaviridae | 0 | 0 | NA | 1 | 0 | orphan | 0 | 0 | NA | 0 | 0 | NA |
| vOTU_sg_8321__S3_k141_209609 | Phycodnaviridae | 0 | 0 | NA | 1 | 0 | orphan | 0 | 0 | NA | 0 | 0 | NA |
| vOTU_sg_8356__S2_k141_567846 | Myoviridae | 0 | 1 | NA | 0 | 0 | NA | 0 | 0 | NA | 0 | 0 | NA |
| vOTU_sg_8427__S6_k141_500634-circular | Myoviridae | 0 | 0 | NA | 3 | 0 | orphan | 0 | 0 | NA | 0 | 0 | NA |
| vOTU_sg_8643__S8_k141_111285-circular | Myoviridae | 0 | 0 | NA | 1 | 0 | orphan | 0 | 0 | NA | 0 | 0 | NA |
| vOTU_sg_8719__S6_k141_202104 | Myoviridae | 0 | 0 | NA | 1 | 0 | orphan | 0 | 0 | NA | 0 | 0 | NA |
| vOTU_sg_8886__S2_k141_55893 | Phycodnaviridae | 0 | 0 | NA | 1 | 0 | orphan | 0 | 0 | NA | 0 | 0 | NA |
| vOTU_sg_8941__S9_contig_125_pilon_pi | Siphoviridae | 0 | 0 | NA | 1 | 0 | orphan | 0 | 0 | NA | 0 | 0 | NA |

| | | | | | | | | | | | | | |
|---|---|---|---|---|---|---|---|---|---|---|---|---|---|
| lon | | | | | | | | | | | | | |
| vOTU_sg_9134__Co_contig_2010_pilon_pilon | Siphoviridae | 0 | 0 | NA | 1 | 0 | orphan | 0 | 0 | NA | 0 | 0 | NA |
| vOTU_sg_9304__Co_k141_1713855 | Podoviridae | 0 | 0 | NA | 1 | 0 | orphan | 0 | 0 | NA | 0 | 0 | NA |
| vOTU_sg_9402__Co_k141_1059106 | Myoviridae | 0 | 1 | NA | 0 | 0 | NA | 0 | 0 | NA | 0 | 0 | NA |
| vOTU_sg_9445__Co_contig_3299_pilon_pilon | Unclassified_Caudovirales | 0 | 0 | NA | 2 | 0 | orphan | 0 | 0 | NA | 0 | 0 | NA |
| vOTU_sg_9553__S8_contig_42_pilon_pilon | Myoviridae | 0 | 0 | NA | 1 | 0 | orphan | 0 | 0 | NA | 0 | 0 | NA |
| vOTU_sg_9712__S8_k141_245476 | Myoviridae | 0 | 0 | NA | 1 | 0 | orphan | 0 | 0 | NA | 0 | 0 | NA |
| vOTU_sg_9885__S10_k141_495592 | Unclassified_Caudovirales | 0 | 0 | NA | 1 | 0 | orphan | 0 | 0 | NA | 0 | 0 | NA |

RM system: Restriction-modification system; M: modification; R: restriction; SHIPPO: Shipborne Pole-to-Pole Observations

**table S4. Methylated motifs of SHIPPO MAGs.**

| MAGs | Family | Methylated motifs (forward/reverse) | REBASE motif | MTase Type |
|---|---|---|---|---|
| SHIPPO_PRO_101 | MGIIA | GGTAG/None, CGCG/CGCG, TCGCGA/TCGCGA, GGTAG/CTACC | CGCG | Type II MTase |
| SHIPPO_PRO_102 | MGIIA | CGCG/CGCG | CGCG | Type II MTase |
| SHIPPO_PRO_132 | *Microbacteriaceae* | TTAA/TTAA, ATTAAT/ATTAAT, GTTAAT/None | TTAA | Type II MTase |
| SHIPPO_PRO_22 | *Rhodobacteraceae* | GANTC/GANTC, GASTC/GASTC, GAWTC/GAWTC, GAGTC/None | GANTC | Type II MTase |
| SHIPPO_PRO_241 | AAA536-G10 | GANTC/GANTC, GASTC/GASTC, GAWTC/GAWTC, GAGTC/None | GANTC | Type II MTase |
| SHIPPO_PRO_245 | *Pelagibacteraceae* | GANTC/GANTC, GASTC/GASTC, GAWTC/GAWTC, GAGTC/None, CGAN$_6$AATC/GATTN$_6$TCG, CGAYN$_4$TTCA/TGAAN$_4$RTCG, GAAN$_6$TCGG/CCGAN$_6$TTC, CGAN$_6$GTCY/RGACN$_6$TCG, GAAN$_5$TCGA/TCGAN$_5$TTC | GANTC | Type II MTase |
| SHIPPO_PRO_246 | *Pelagibacteraceae* | GANTC/GANTC, GASTC/GASTC, GAWTC/GAWTC, GAGTC/None, CGAAN$_5$TCC/GGAN$_5$TTCG | GANTC | Type II MTase |
| SHIPPO_PRO_33 | *Pelagibacteraceae* | GANTC/GANTC, GASTC/GASTC, GAWTC/GAWTC, GAGTC/None, CGAN$_7$CTTG/CAAGN$_7$TCG, TGACN$_5$CTC/GAGN$_5$GTCA, GTGAN$_5$RTCG/CGAYN$_5$TCAC, CCGCAG/None, ATGN$_5$GTCG/CGACN$_5$CAT, CGAN$_6$GTCY/RGACN$_6$TCG | GANTC | Type II MTase |
| SHIPPO_PRO_64 | o_*Pelagibacteraceae* | GANTC/GANTC, GASTC/GASTC, GAWTC/GAWTC, GAGTC/None, GAGN$_6$TAYG/CRTAN$_6$CTC | GANTC | Type II MTase |
| SHIPPO_PRO_66 | AAA536-G10 | GANTC/GANTC, GASTC/GASTC, GAWTC/GAWTC, GAGTC/None | GANTC | Type II MTase |

| | | | | |
|---|---|---|---|---|
| vOTU_N_648 | Unknown_Virus | GATC/GATC, GGATC/None, None/GATCC, TGATCA/TGATCA, RGATCY/RGATCY, GGATC/GATCC | GATC | Type II MTase |
| vOTU_sg_10720 | *Siphoviridae* | GGAG/None, GRRGA/None, GGWGA/None, GAAGATCTTC | CTYCAG | Type II MTase |

SHIPPO: Shipborne Pole-to-Pole Observations; MAG: metagenome-assembled genome; MTase: methyltransferase.; underscore: DNA modification

**table S5. Unmethylated GANTC sites across samples.**

| Position (Contig:Position:Strand) | Region | KEGG |
|---|---|---|
| SHIPPO_PRO_33_11:21967:- | Genic | K00215 |
| SHIPPO_PRO_33_10:5600:- | Genic | K00285 |
| SHIPPO_PRO_33_1:33961:+ | Genic | K00666 |
| SHIPPO_PRO_33_1:34117:+ | Genic | K00666 |
| SHIPPO_PRO_33_4:47162:+ | Genic | K01879 |
| SHIPPO_PRO_33_10:61000:+ | Genic | K01956 |
| SHIPPO_PRO_33_10:61092:+ | Genic | K01956 |
| SHIPPO_PRO_33_5:70038:- | Genic | K03702 |
| SHIPPO_PRO_33_11:32121:+ | Genic | K03980 |
| SHIPPO_PRO_33_6:5815:+ | Genic | K07090 |
| SHIPPO_PRO_33_1:85989:+ | Genic | K14980 |
| SHIPPO_PRO_33_2:49451:+ | Genic | Unclassified |
| SHIPPO_PRO_33_10:26785:- | Genic | Unclassified |
| SHIPPO_PRO_33_11:155821:+ | Genic | Unclassified |
| SHIPPO_PRO_33_1:88695:- | Intergenic | Unclassified |
| SHIPPO_PRO_33_1:33963:- | Intergenic | Unclassified |
| SHIPPO_PRO_33_1:34119:- | Intergenic | Unclassified |
| SHIPPO_PRO_33_1:85991:- | Intergenic | Unclassified |
| SHIPPO_PRO_33_2:49453:- | Intergenic | Unclassified |
| SHIPPO_PRO_33_2:7767:- | Intergenic | Unclassified |
| SHIPPO_PRO_33_3:78935:+ | Intergenic | Unclassified |
| SHIPPO_PRO_33_3:57851:+ | Intergenic | Unclassified |
| SHIPPO_PRO_33_3:4705:- | Intergenic | Unclassified |
| SHIPPO_PRO_33_5:70036:+ | Intergenic | Unclassified |
| SHIPPO_PRO_33_6:5817:- | Intergenic | Unclassified |

| | | |
|---|---|---|
| SHIPPO_PRO_33_8:61449:+ | Intergenic | Unclassified |
| SHIPPO_PRO_33_10:26783:+ | Intergenic | Unclassified |
| SHIPPO_PRO_33_10:5598:+ | Intergenic | Unclassified |
| SHIPPO_PRO_33_10:61094:- | Intergenic | Unclassified |
| SHIPPO_PRO_33_11:171226:+ | Intergenic | Unclassified |
| SHIPPO_PRO_33_11:146653:+ | Intergenic | Unclassified |
| SHIPPO_PRO_33_11:21965:+ | Intergenic | Unclassified |
| SHIPPO_PRO_33_11:86684:- | Intergenic | Unclassified |
| SHIPPO_PRO_33_11:99001:- | Intergenic | Unclassified |
| SHIPPO_PRO_33_11:32123:- | Intergenic | Unclassified |
| SHIPPO_PRO_33_3:78937:- | Regulatory | K02864 |
| SHIPPO_PRO_33_5:1650:- | Regulatory | K04077 |
| SHIPPO_PRO_33_6:11812:- | Regulatory | K04488 |
| SHIPPO_PRO_33_8:6914:- | Regulatory | K07462 |
| SHIPPO_PRO_33_11:114352:- | Regulatory | K09883 |
| SHIPPO_PRO_33_8:61451:- | Regulatory | K09969 |
| SHIPPO_PRO_33_5:61241:- | Regulatory | Unclassified |

SHIPPO: Shipborne Pole-to-Pole Observations; KEGG: Kyoto Encyclopedia of Genes and Genomes.
